# AI-Discovered Cognitive Models Reveal Novel Insights into Human and Animal Learning

**DOI:** 10.64898/2026.05.18.725921

**Authors:** Daniel Kasenberg, Pablo Samuel Castro, Maria K. Eckstein, Nóemi Éltető, Will Dabney, Caroline Wang, Martin Engelcke, Rishika Mohanta, Aparna Dev, Matthew M. Botvinick, Nenad Tomasev, Glenn C. Turner, Vincent Costa, Nathaniel D. Daw, Kimberly L. Stachenfeld, Kevin J. Miller

**Author notes:** These authors contributed equally to this work (Daniel Kasenberg, Pablo Samuel Castro). These authors contributed equally to this work (Kimberly L. Stachenfeld, Kevin J. Miller).

## Abstract

Scientific models are widely used across the natural sciences as an interface between scientific theories and empirical data [1]. Such models play a key role, for example, in the study of human and animal learning, where they express algorithmic hypotheses and relate them to psychology and neuroscience data [2, 3]. These models are traditionally handcrafted by expert researchers based on existing theory or new insights. Such handcrafted models, however, are now known to fall short of capturing the full richness of behavior, even in their narrow domains [4–7]. An alternative data-driven approach has emerged, seeking to discover new insights by fitting and interpreting flexible models [8–11]. However, these tools require substantial human effort to derive insight from data, and it has been unclear how to discover new ideas from data efficiently. Here, we present DataDIVER, a general approach for automatically discovering computational models from data, and demonstrate that these models surface novel mechanistic insights into human and animal learning. Our approach delivers models that take the form of short computer programs, which are optimized both to fit data well and to be simple. These programs explicitly connect with existing theoretical frameworks and are readily understandable by human scientists. They can also be used to make novel predictions, some of which we show are borne out in re-analysis of existing data. General-purpose tools for surfacing new ideas from data, especially in combination with the large datasets that are increasingly available in many fields, stand to dramatically accelerate scientific discovery.

---

Mechanistic computational models are an important tool in many sciences, allowing abstract theoretical ideas to be instantiated and tested quantitatively against data. In many domains, this has traditionally been a “theory-first” process, with models constructed based on existing theory and used as tools for exploring the implications of that theory [12, 13]. Recently, however, a new generation of data-driven model discovery tools has begun to invert this, enabling researchers to discover useful scientific models directly from data [9, 11, 14, 15]. The advent of strong and general-purpose generative AI in principle holds enormous potential for this practice, as machines can for the first time generate exactly the types of objects that human researchers use to express scientific models, including human-readable, literature-aware computer code. Recently, AI-optimized code has shown an impressive ability to propose statistical models and data processing pipelines [16, 17], and even identify unknown solutions to problems in mathematics and engineering [18, 19]. However, discovery in these domains is fundamentally unlike that in basic science, because progress in basic science requires surfacing novel insights into the underlying structure of the real-world processes.

A key open problem of the latter kind is identifying algorithms that humans and other animals use to learn from reward. Despite being one of the oldest questions in psychology [20, 21], it remains unsolved. Current computational approaches are heavily theory-driven, taking inspiration from reinforcement learning [22, 23] as well as Bayesian optimality [24, 25]. Recent work has demonstrated that these theory-driven models are decisively outperformed by blackbox predictive models in terms of quality-of-fit [4, 5, 26, 27]. This suggests that better mechanistic models very likely exist, but does not on its own suggest a way to discover them. Reward learning, like other problems in cognitive science, is particularly challenging as it involves the update over time of internal (“cognitive”) variables that are only indirectly reflected in observable behavior like choices. Model discovery in these contexts therefore requires inferring from data how many internal variables exist, how they are updated based on external input, and how they result in behavior.

Here, we present DataDIVER (Data-driven Discovery of Interpretable models Via Evolutionary Refinement), a tool for discovering scientifically interpretable symbolic models from data. DataDIVER relies on AlphaEvolve, a recently developed tool that has shown impressive performance on program optimization problems in mathematics and computer science by using LLMs to edit programs within an evolutionary algorithm that optimizes for a user-specified objective [19]. DataDIVER extends AlphaEvolve, allowing it to optimize programs based on both quality-of-fit to a dataset and on simplicity.

A challenge for the problem of automating model discovery is deciding on the target(s) of optimization, as scientifically useful models must meet two important criteria. The first is that they should capture data accurately, which in our setting can be quantified as predicting the observed data with high likelihood (“quality-of-fit”). While this is straightforward to quantify, it is difficult to optimize, and earlier attempts with LLM-based search techniques simpler than AlphaEvolve have not been able to match the performance of deep learning models at predicting learning behavior [28, 29]. The second is that they should convey human-interpretable insights about the data. While human interpretability is ultimately subjective, a close correlate is program complexity, for which a number of quantitative metrics have been proposed [30]. Furthermore, quality-of-fit often trades off against complexity metrics in practice, and there is no single answer to how much relative value to place on each. Different applications demand different tradeoffs between complexity and quality-of-fit, from simple abstract models that are “wrong but useful” [31] to detailed models that do not “surrender the adequate representation of a single datum of experience” [32].

Our approach to this tension is to optimize a set of models that each strikes a different balance between quality-of-fit and simplicity. We apply this approach to five datasets of learning behavior from a range of species and behavioral settings. We find that the best-fitting programs match the quality-of-fit of “blackbox” neural network models, which to our knowledge has never been achieved with purely symbolic models on learning behavior datasets [28, 29]. Perhaps unsurprisingly, these extremely well fitting programs are lengthy, redundant, complex programs that do not readily afford interpretation. The remaining spectrum of programs surfaces meaningfully novel insights in an accessible way, with the simplest models shedding light on the basic organization of learning behavior, and more complex programs revealing more detailed structure (Fig. 2a). Some of these discovered learning mechanisms suggested the presence of previously unknown patterns which we verified by reexamining the behavioral data. Broadly, these results showcase the usefulness of AI tools not just to predict behavior but also to explain it.

**Fig. 1:**
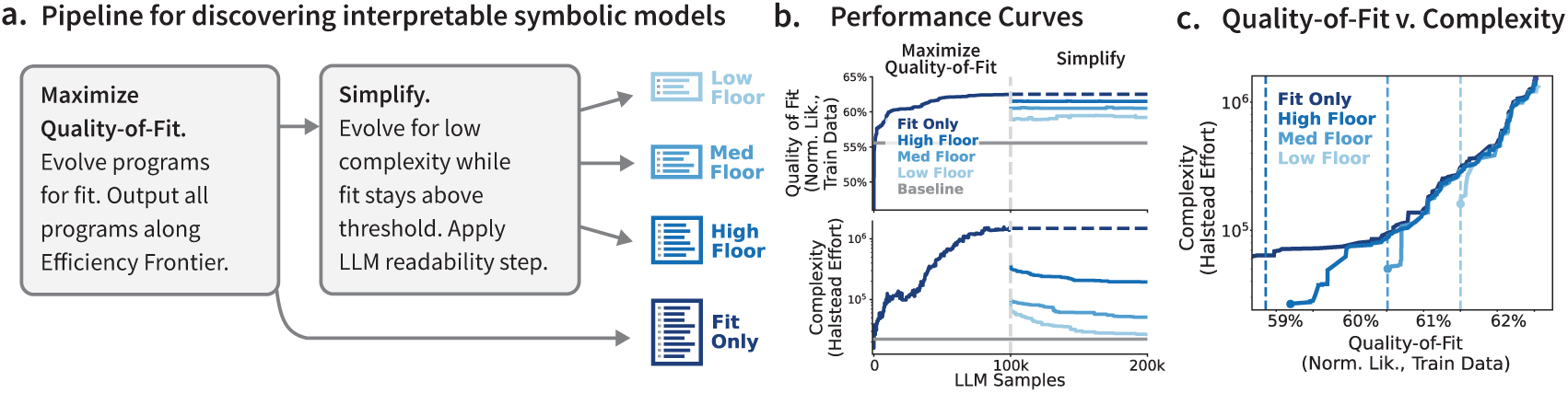
Overview of DataDIVER Pipeline. **(a)** Our pipeline consists of two stages: in the *Maximize Quality-of-Fit* stage, AlphaEvolve is used to optimize programs that maximize the normalized likelihood of the observed data. In the *Simplify* stage, a second AlphaEvolve run minimizes Halstead complexity metrics while keeping the quality-of-fit score above a floor, after which an LLM-powered code refactor improves human readability. These stages result in four programs: a very complex “Fit only” program emerging from stage one, and three simplified programs that strike different balances of complexity and quality-of-fit. **(b)** Performance curves illustrating likelihood and complexity scores over the course of evolution. In the first stage, both likelihood and complexity increase; in the second stage, likelihood is maintained above the specified floor while complexity is reduced. **(c)** Quality-of-fit versus complexity for programs that strike efficient tradeoffs of fit and complexity. Darkest line depicts programs generated during the *Maximize Quality-of-Fit* stage; lighter lines depict programs generated during the *Simplify* stages for different floors (vertical dashed lines). Each *Simplify* stage improves the frontier (i.e. identifying simpler programs that satisfy each floor).

**Fig. 2:**
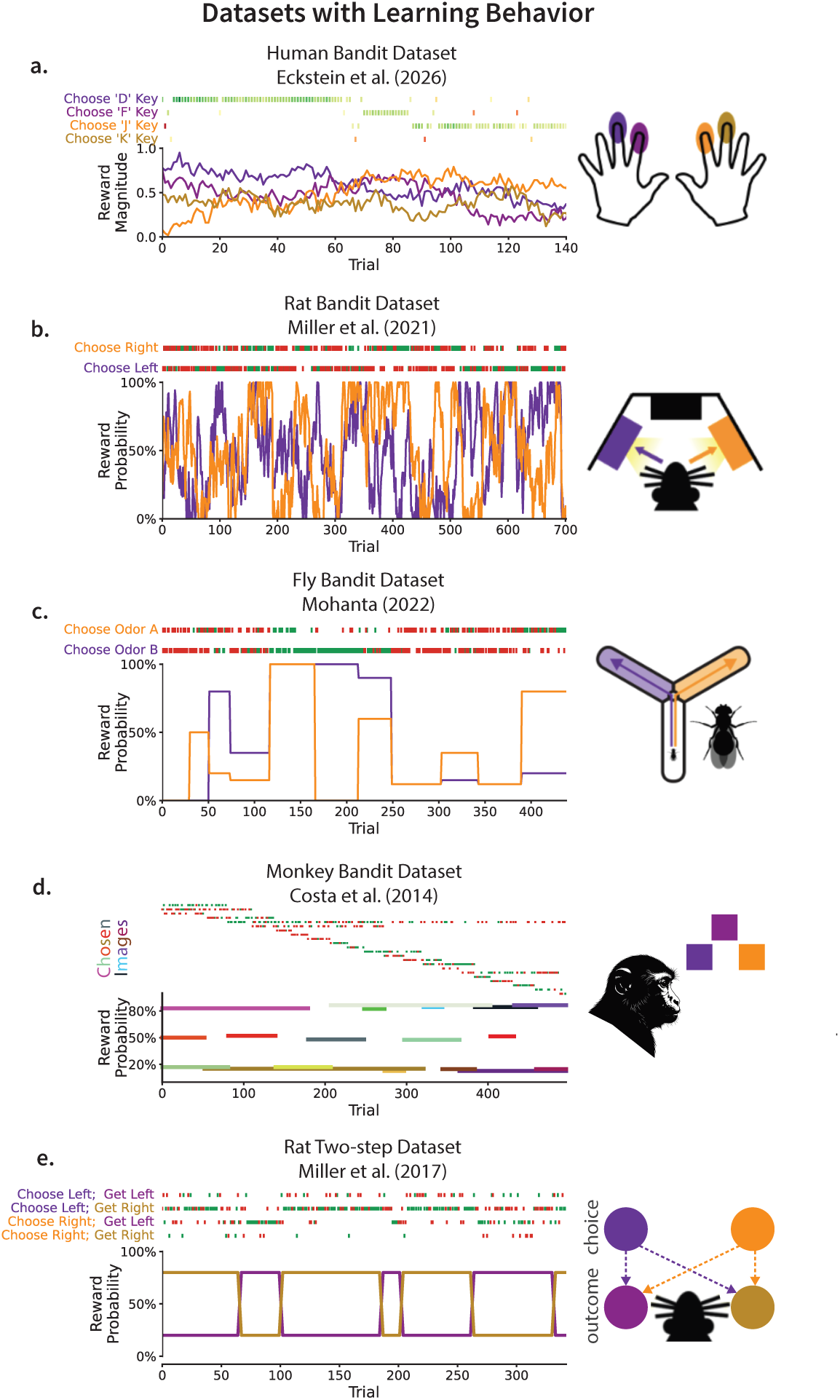
Illustration of datasets. Each dataset contains human or animal behavior in different reward-guided learning experiments. In each dataset, subjects chose among discrete choices to receive dynamically changing rewards, with each line above denoting the reward structure for each choice over time. The datasets vary in the subject species, the way the animal experiences stimuli and demonstrates choices, the number of choices available, and the latent reward structure.

## 1 Data-driven optimization of programs

The DataDIVER pipeline consists of two program optimization stages, both of which are powered by AlphaEvolve [19]. AlphaEvolve is an evolutionary algorithm that uses a large language model (LLM) to modify (or “mutate”) programs as instructed in user-specified prompts. These programs are evaluated according to a user-specified “fitness” objective, with higher scoring programs preferentially sampled for further modification. Following Castro et al. [28], our programs implement a learning rule that takes in relevant information about the subject’s past experience (previous choice, previous reward, other information about the previous and current trial if relevant, and an “agent state” maintaining memory) and output a probabilistic prediction about the animal’s next choice. The programs have individual parameters which are fit separately to data from each subject, and validated on a held-out portion of the subject’s data using two-fold cross-validation. These parameters might include learning rates, exploration parameters, or whatever individual parameters DataDIVER decides to implement. We run three separate instances of this evolutionary process for each dataset, considering half of the subjects in the dataset. The remaining subjects are held-out for use in evaluating the discovered models. Programs are implemented in Python using JAX [33] so that they are differentiable, allowing efficient parameter optimization. We run three independent runs of both DataDIVER stages.

In the first stage (*Maximize Quality-of-Fit*; Figure 1A), programs are optimized to maximize quality-of-fit on the behavioral data without regard to complexity(see Supplement C.1). This results in a large set of programs that vary widely both in quality-of-fit and in complexity. We quantify complexity using Halstead effort [30], a heuristic measure that uses the number of variables and operations in the program to estimate how much time and effort a human programmer would need to understand it (Figure 1C). We select for further consideration the single program with the greatest quality-of-fit.

The second stage (*Simplify*; Fig. 1A) aims to produce simpler programs which only modestly sacrifice quality-of-fit. We run three separate instances of this stage per dataset, each with a different quality-of-fit floor that programs cannot fall beneath. This enabled us to recover a set of programs that optimize for different tradeoffs between complexity and quality-of-fit. The stage is initialized with all programs from the *Maximize Quality-of-Fit* stage that efficiently trade off quality-of-fit and complexity: that is, programs on the “Pareto frontier” for which no discovered program improves one objective without sacrificing the other (Fig. 1c). Each *Simplify* stage succeeds if it succeeds in pushing this frontier, which they do by identifying simpler models that maintain quality-of-fit. In this stage, the LLM prompt encourages simplifying the programs (see Supplement C.2), and the fitness function rejects programs that fall beneath the quality-of-fit floor but otherwise selects programs based on their simplicity. We select for further consideration the single program with the greatest simplicity (smallest Halstead effort) from each *Simplify* run.

Minimizing Halstead complexity encourages the programs to implement simpler mechanisms, but ignores important human readability considerations like documentation, organization, and consistency. We use an LLM (Gemini 2.5 Pro) to rewrite each simplified program to improve readability without changing the program’s function (prompts in Supplement C.3). The final output of DataDIVER consists of the best-fitting programs from the *Maximize Quality-of-Fit* stage and the rewritten programs from the *Simplify* stage (see Figure 4 for an example program; additional examples for each dataset are found in Supplement A). In addition to these automatically-generated programs, we also manually constructed a single “synthesis” program for each dataset combining the key human-interpretable mechanisms discovered for that dataset.

## 2 Fit-only programs are strong predictive and generative models

We apply DataDIVER to five datasets containing humans and other animals performing diverse reward learning tasks: *Human Bandit* [10], *Rat Bandit* [8], *Fly Bandit* [34], *Monkey Bandit* [35, 36], *Rat Two-step* [37] (Figure 2). In each of these tasks, a subject repeatedly selects one of several available actions and receives rewards whose magnitude or probability depend on the chosen action and vary over time. Each dataset contains a large number of sessions, making them suitable for data-driven model discovery. Each has also been the focus of intensive previous computational modeling efforts which have yielded a strong handcrafted baseline model [8, 10, 34, 35, 38, 39], enabling us to robustly benchmark DataDIVER-discovered models.

First, we consider the best-fitting “fit-only” programs from the first *Maximize Quality-of-Fit* stage of DataDIVER by evaluating their quality-of-fit on the held-out subjects for each dataset using trial-normalized likelihood [2]. Each program significantly outperformed the baseline model for its dataset (average difference in normalized likelihood of 4.1 percentage points, all p *<*0.02 on fifteen separate t-tests over subjects; Figure B14). These differences were consistent over three independent runs of DataDIVER (average difference in normalized likelihood was 5.8 percentage points for the *Human Bandit* dataset, p=0.0004, t-test over three DataDIVER runs; *Rat Bandit* 0.50pp p=0.01; *Fly Bandit* 0.51pp, p=0.0002; *Monkey Bandit* 0.47pp, p= 0.002; *Rat Two-step* 1.4pp, p=0.0005; Figure 3A). We also compare our programs to generic recurrent neural networks (RNNs; see Section 7.9), which have been widely found to outperform symbolic models when applied to large behavioral datasets [5, 9, 10, 26, 28]. We find that the fit-only models discovered by DataDIVER overall perform similarly to RNNs (average performance difference 0.05pp, p=0.88, t-test across datasets). Considering individual datasets, the fit-only programs significantly outperformed the best RNNs on three datasets (*Fly Bandit*, average difference 0.14pp, p=0.003, t-test across DataDIVER seeds; *Rat Two-step*, average difference 0.17pp, p=0.03; and *Monkey Bandit*, the only dataset for which the RNN underperformed the baselines, likely due to smaller dataset size, average difference 1.0pp, p=0.0003) and significantly underperformed them on one (*Human Bandit*, average difference 0.9pp, p=0.02).

**Fig. 3:**
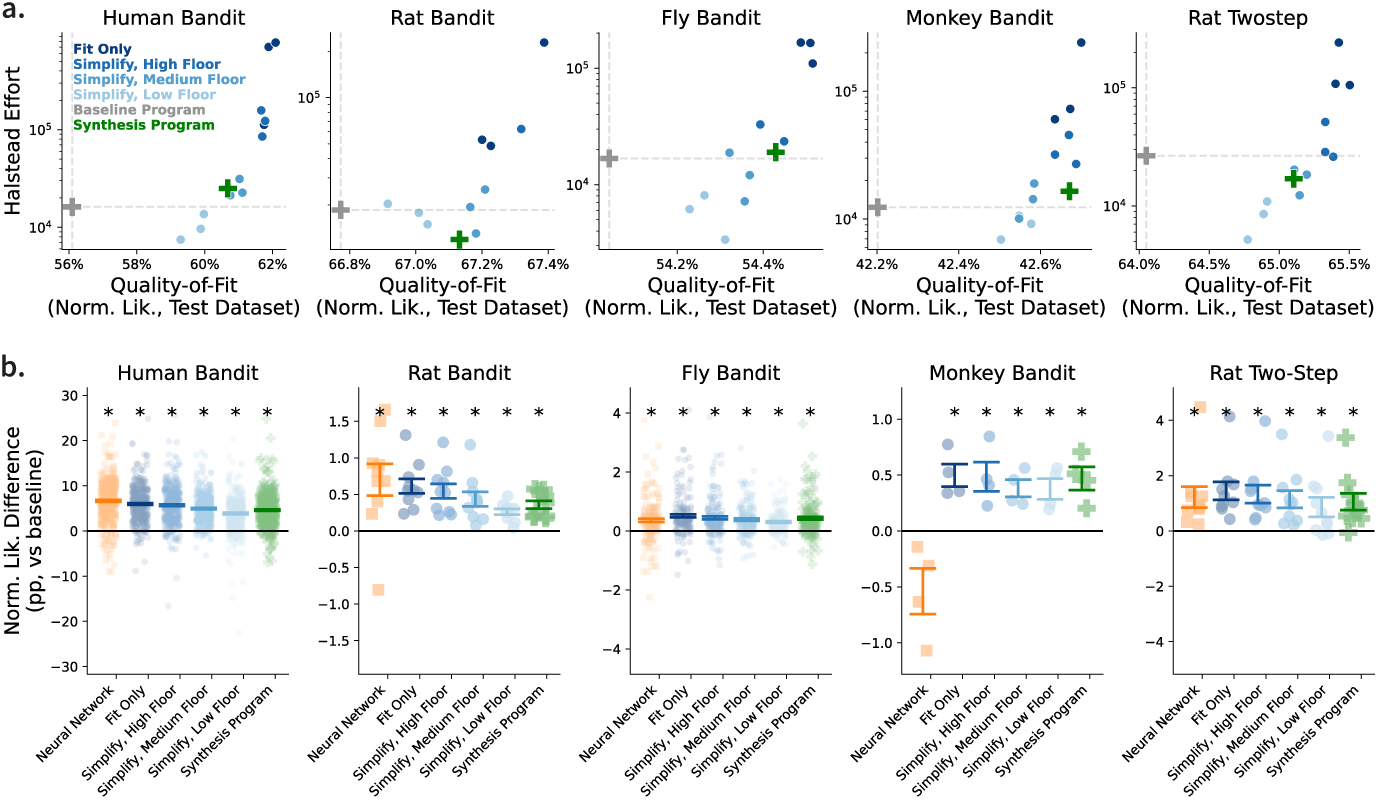
Evolved programs fit data well and trade off quality-of-fit and complexity. **(a)** Quality-of-fit (normalized likelihood on held-out data) against complexity (Halstead effort) for each DataDIVER program, the state-of-the-art handcrafted baseline program from the literature, and the synthesis program, plotted for each dataset. Discovered programs achieve higher quality-of-fit than the baseline program. Programs simplified to lower floors tend to have lower quality-of-fit. **(b)** Quality-of-fit difference compared to the baseline model, plotted separately for each participant, for the best-fitting program at each floor (see B14 for all programs). Points indicate the quality-of-fit difference between the indicated model and the baseline model for individual subjects. Error bars indicate standard errors over subjects. Asterisks indicate significant differences from the handcrafted baseline model (*p <* 0.05, t-test).

**Fig. 4:**
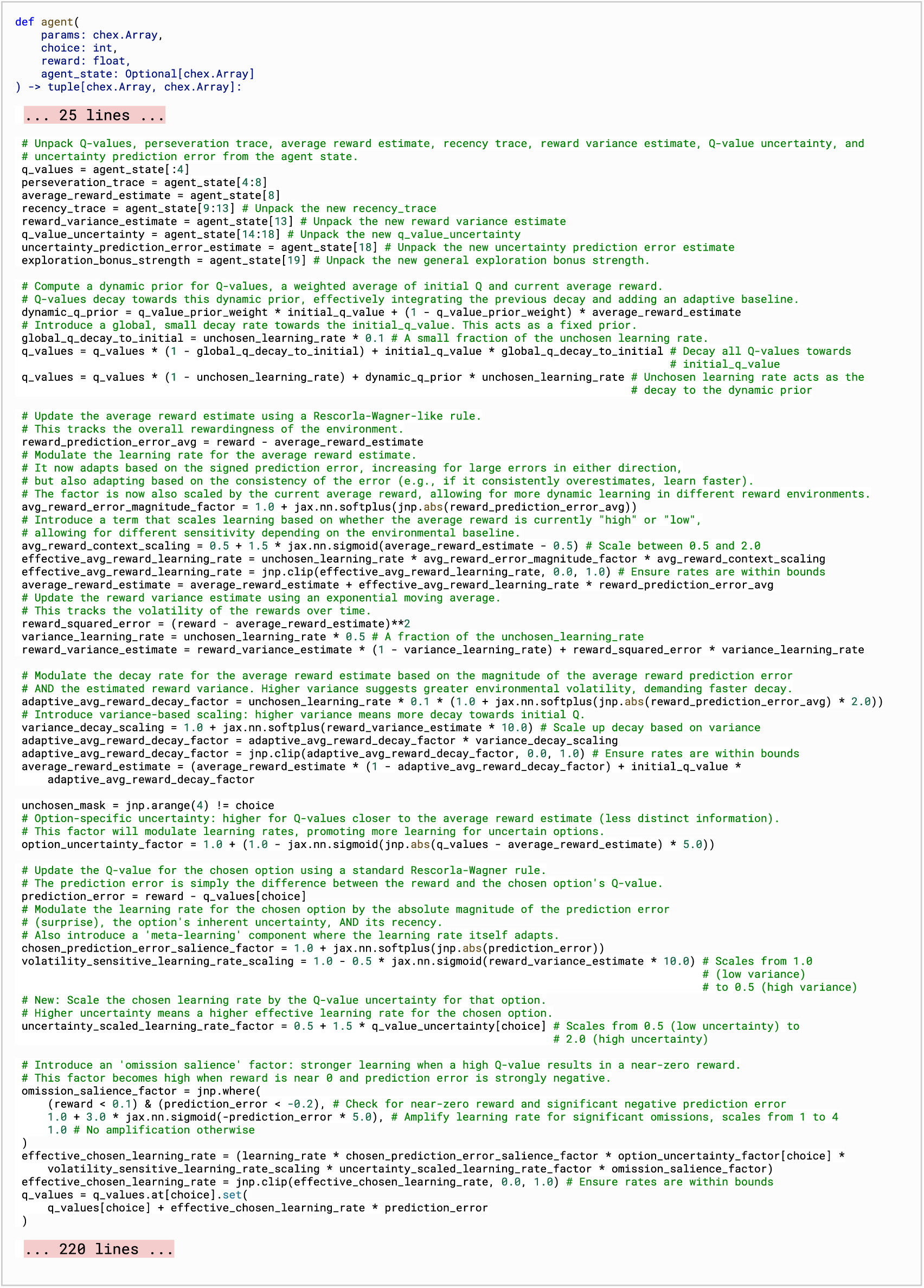
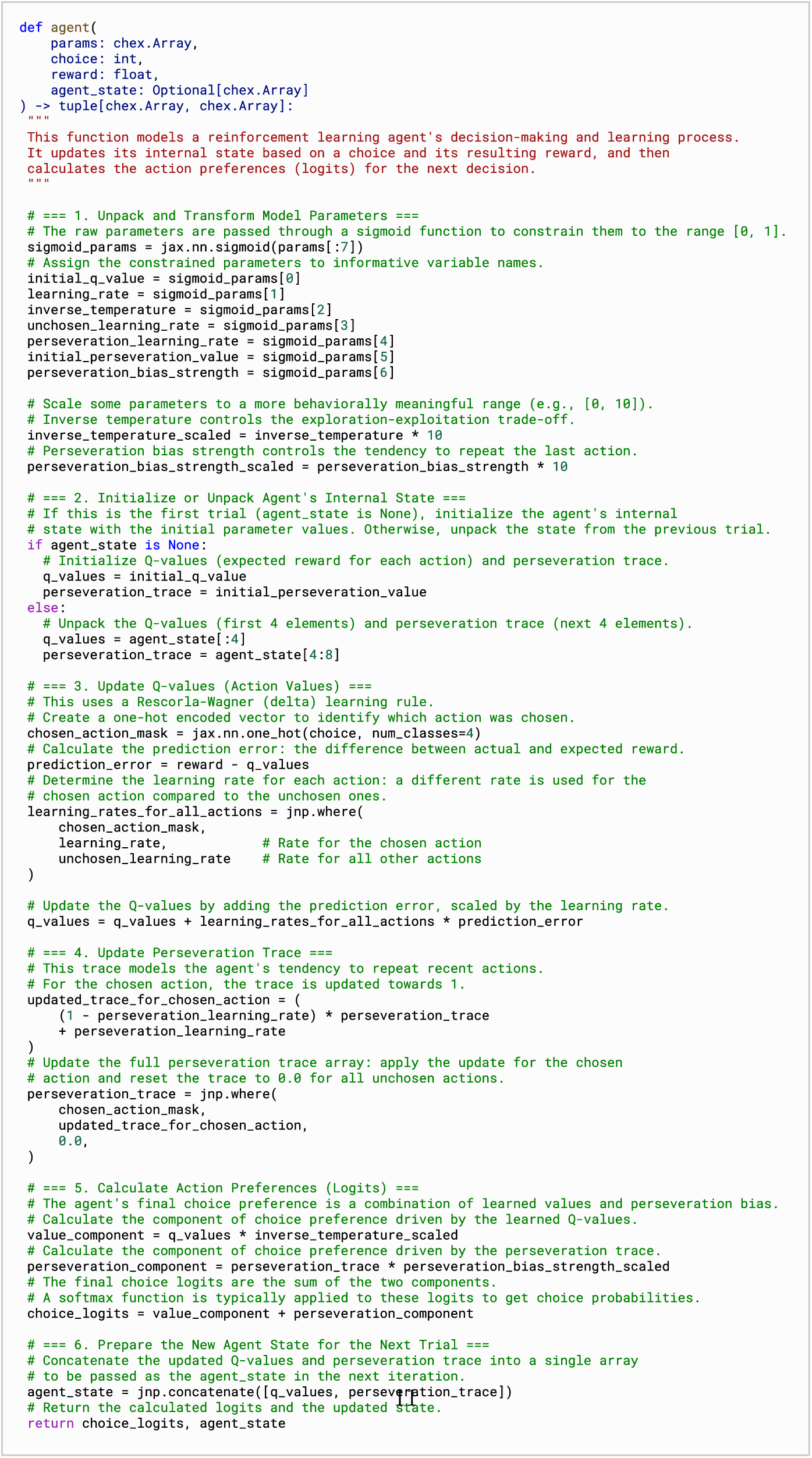
**(a) Example Discovered Fit-only Program.** This is a representative “fit-only” program outputted by the *Maximize Quality-of-fit* stage. It shows certain promising characteristics–informative variable names and recognizable computational motifs–but the program is quite complex (20 state variables are defined), the operations are complex and highly nonlinear, and the mechanisms are partially overlapping (e.g. multiple variations on reward prediction error-driven updates). Program is truncated and adjusted to fit on one page; it is otherwise unedited. We note that jnp = jax.numpy; programs were implemented in jax to permit gradient-based parameter optimization. **(b) Example Discovered Simplified, Low-floor Program.** This is an example program outputted by the *Simplify* stage (*Human Bandit*, low floor, run 1). It shows characteristics seen across other discovered programs: interpretable variable names, single computations per line, and comments describing hard-to-parse operations and organizing the computational structure. It is unedited except for adjustments to the spacing for compactness.

In order to test whether the discovered programs generate behavior that matches the characteristics of the real data [3, 40], we simulated choices in an environment matched to the experimental conditions to create an artificial dataset (see Section 7.10 for details). We then computed a trial-history regression model for the real and artificial datasets [41, 42] to quantify the effects of past trials’ outcomes on current choice (see Section 7.11). We find that behavior generated by our AI-discovered programs closely matches the patterns seen in the real datasets, while datasets generated by the handcrafted baseline models fail to capture many features (Figures A2, A6, A9, A11, A13). Taken together, these results indicate that the fit-only models discovered by DataDIVER are strong predictive and generative models, outperforming handcrafted baseline models and closing most or all of the gap with blackbox neural networks.

## 3 “Simplified” programs trade off complexity and quality-of-fit

The programs that emerge from the *Maximize Quality-of-Fit* stage are strong predictive and generative models, but they are very complex (Figure 4A, see full code for one program in A.1.6). We see that the program is thoroughly commented and includes recognizable variable names and mechanisms; however, it is highly repetitive and overwhelmingly long, and individual operations are very complex. These programs exceed the baseline programs in terms of Halstead effort (by a factor of 12.7 ± 3.7; mean ± standard error over all programs), number of lines of code (by a factor of 2.7 ± 0.6), and number of state variables (by a factor of 3.2 ± 0.4). This complexity substantially reduces their appeal to human scientists, as complex models are difficult to interpret and often considered to be less plausible as mechanistic models.

The *Simplify* stage of our pipeline (Figure 1A) succeeded in generating substantially simpler programs: the high, medium, and low floor programs respectively had 30 ± 6%, 14 ± 2.5%, and 9.8 ± 2.8% the Halstead effort of the fit-only program (Figure 3A). Strikingly, programs simplified to the low floor were simpler even than the handcrafted baseline programs, having on average about half the Halstead effort (59 ± 0.1 %). The simplification procedure did affect quality-of-fit, with programs simplified to a lower quality-of-fit floor having lower fit quality (*Human Bandit*: p=0.0001; *Rat Bandit* p=0.005; *Fly Bandit*: p=0.0001; *Monkey Bandit*, p=0.003; *Rat Two-step* p=0.0001; Page’s trend test [43] over DataDIVER seeds and quality-of-fit floors). Nevertheless, the vast majority of the AI-discovered models significantly outperformed our handcrafted benchmark models in terms of quality-of-fit (54/58 evolved programs significantly better than baseline at *p <* 0.05, paired t-tests over subjects; Figure B14). Qualitatively, these simplified programs were surprisingly readable, and far more accessible than the fit-only programs. Figure 4B shows an example low-floor program (additional programs are found in Supplement A). The discovered programs use literature-aware variable names like “learning rate”, “q-values”, and “perseveration” as well as mechanisms like error-driven learning and forgetting by decay. Unlike the fit-only program in Figure 4A, the entire program can be viewed on a single page, and does not contain obviously redundant code or unnecessarily complex operations. The simplified programs use comments to organize the code into numbered sections, which can be browsed as a “table of contents”, and additional comments describe complex steps. Some of these computations are unusual (e.g. the update on unchosen action values, the forgetting update on the perseveration trace), but the mechanisms can still be readily understood. Furthermore, a similar organization is applied across the different discovered programs for each dataset, allowing straightforward comparison and synthesis. In general, we found the low-floor programs to be readily understandable, the medium-floor programs could be fully understood with moderate effort, and the high floor programs often could not be fully understood, though it was nevertheless typically possible to glean testable insights from inspecting them.

## 4 Evolved models surface novel mechanistic insights

Several themes emerged across the discovered programs. First, although the programs are written in familiar, literature-aware vocabulary, there was often surprising structure to how internal variables were organized and updated and how they mapped onto decisions. One notable tendency across species and complexity levels was to introduce additional or different cognitive variables than those assumed by previous models. Such reorganizations challenge the prior interpretations of the baseline models’ putative subcomponents; for example, our discovered models break assumptions that novelty and reward preference are folded into a single common-currency reward expectancy (*Monkey Bandit*) or that learning rules update a scalar preference for one choice over the other rather than per-choice statistics (*Rat Bandit*, *Rat Two-step*). Second, discovered programs often included nonstationary, nonlinear modulation of various operations or parameters (e.g. softmax temperatures or decay targets). This potentially captures the subjects’ implicit adaptation to statistics of the task like the average reward level, a type of adaptive learning that has often been neglected when studying any individual dataset. Notably, a number of discovered motifs suggested novel insights about the data that were supported by reanalysis–a striking instance of AI leading to a novel data-driven discovery (e.g. Figure 7).

We briefly summarize insights from the DataDIVER discovered programs below (see Supplement A for more detail). To simplify visualization and verification of the many programs produced by DataDIVER, we also consolidated the different programs for each dataset into a single synthesis programs, which are depicted in Figures 5 and 6. Discovered motifs were excluded from the synthesis program if removing them had no effect on quality-of-fit, and prioritized if they evidently contributed to their program’s predictive or generative performance (see Section 7.14 for more detail). All insights discussed below are included in the synthesis program.

**Fig. 5:**
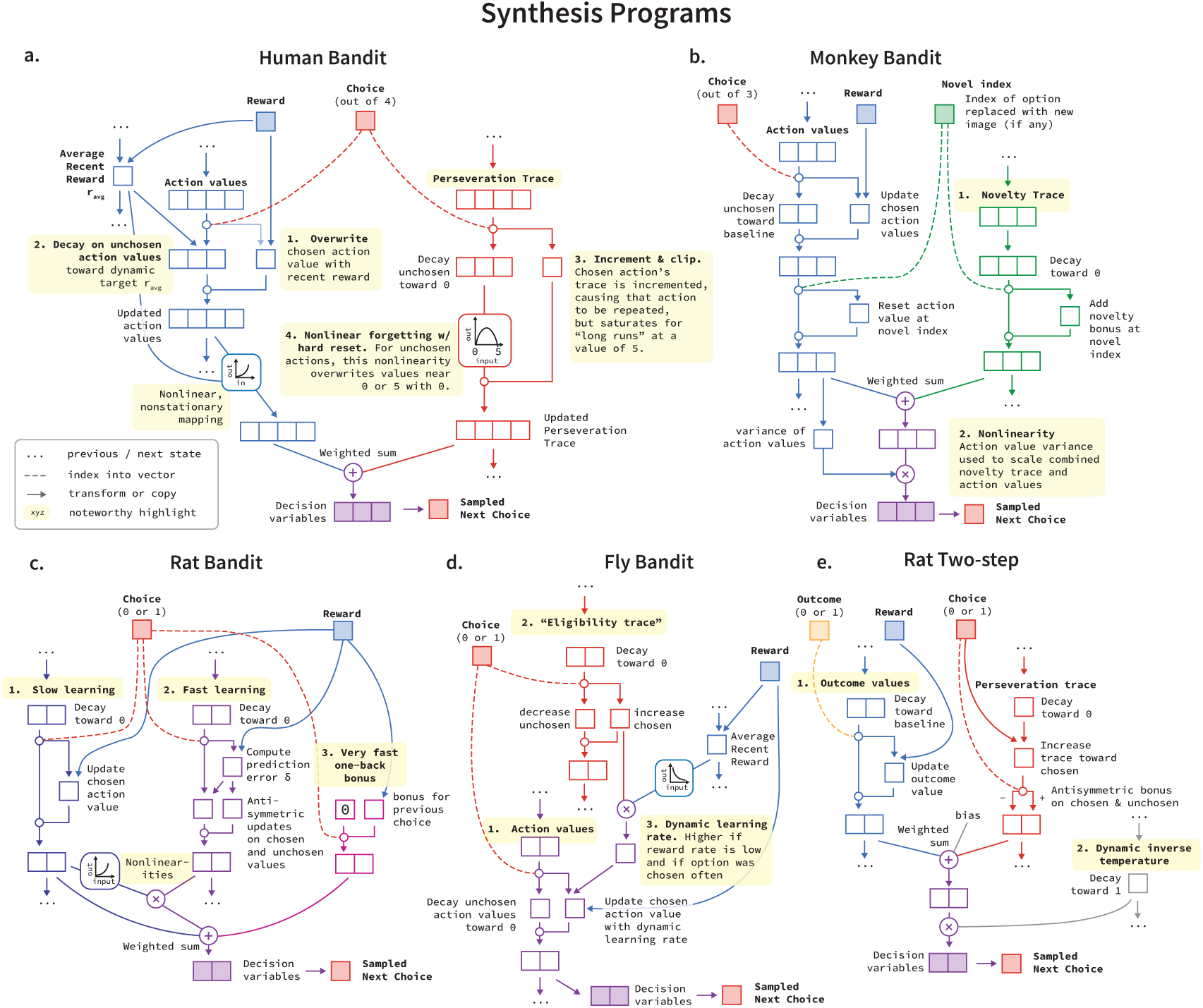
Human expert synthesis of mechanisms in evolved programs. Each diagram illustrates the logic of the synthesis programs. Distinct modules are denoted with different colors.

**Fig. 6:**
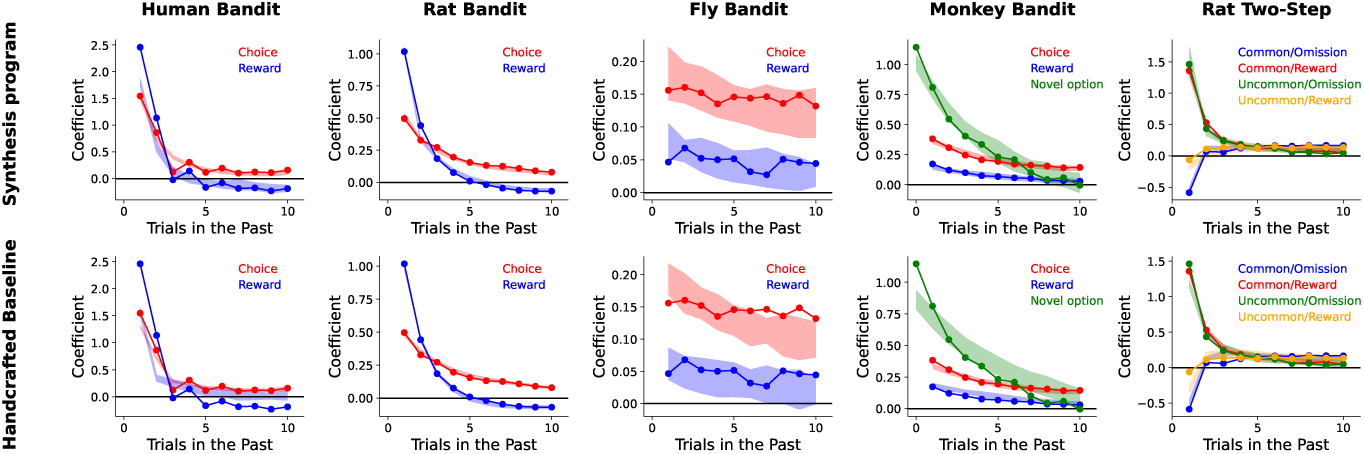
Synthesis programs are strong generative models. We generate artificial data using the synthesis programs, which aggregate information from across the discovered programs, and compute trial-history regression models, which measure the effects of trial variables (like choices and rewards) from earlier trials on the current choice. Coefficients estimated from the real experimental data are shown as dots and lines; and compared to the range given by the same analyses applied to simulations (where shaded patches show the 95% prediction intervals over artificial datasets). The datasets generated by our synthesis programs (top) closely match the patterns seen in the real datasets, often qualitatively better than the original handcrafted baselines (bottom). Lagged regression plots for all models can be found in Supplement A.

**Fig. 7:**
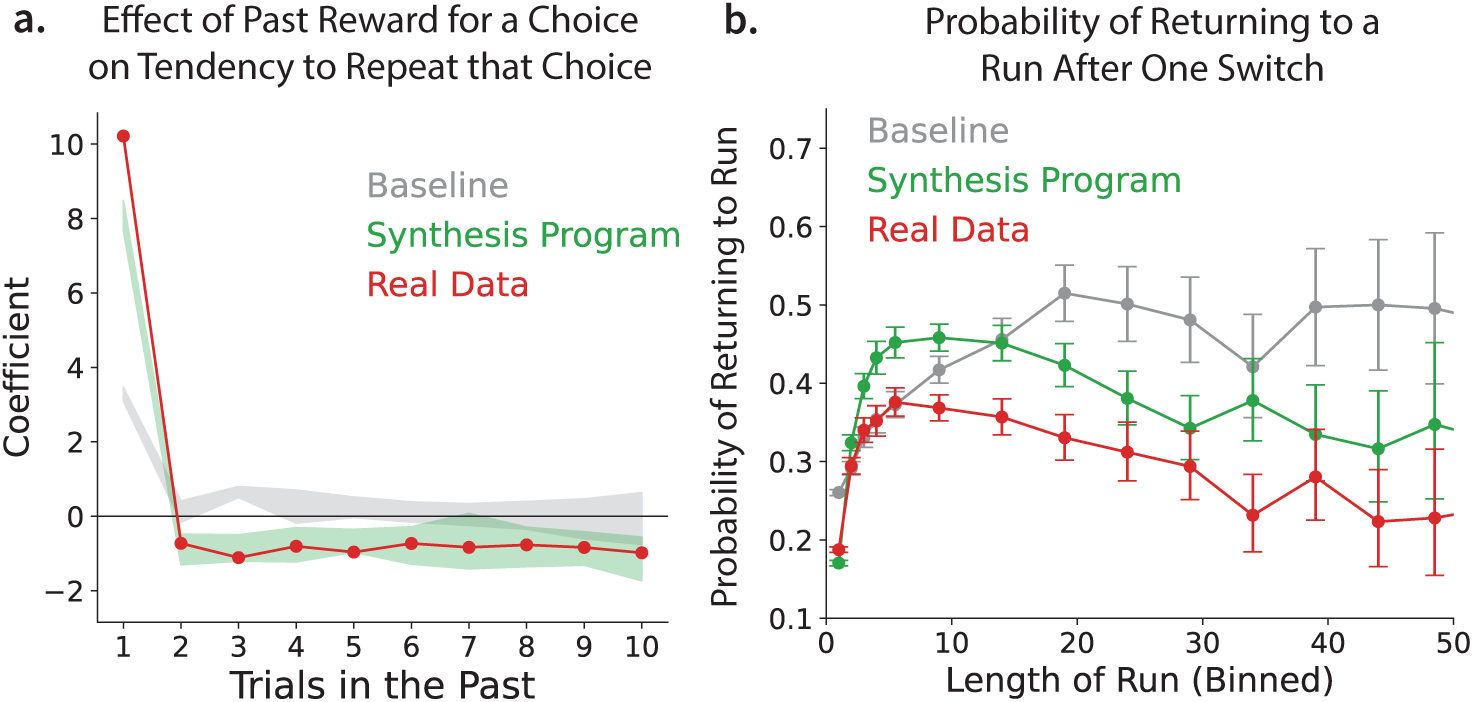
Inspecting discovered programs reveals unexpected patterns in *Human Bandit* data. **(a)** Trial-lagged regression analysis showing the effect (coefficient, positive promotes repeating) of being rewarded for a choice in a previous trial (1-10 trials ago) on tendency to repeat that choice. We note two features of interest predicted by our analysis of discovered programs. First, because previous reward overwrites the chosen action value, we predict that the immediately preceding trial should have an outsized effect, which we see for the real data and synthesis program’s generated data but not baseline program’s. Second, since the unchosen action values are slowly updated toward the recent average reward and chosen action values are overwritten, rewards that are more than one trial in the past actually have a longer-term effect on incrementing unchosen action values—regardless whether they were received at the same arm or a different one, thereby driving a tendency to switch. This is reflected in the negative coefficients on trial lags 2-10 seen for the real data and synthesis program’s generated data but not the baseline program’s. **(b)** Probability of returning to a “run” (a sequence of repeated choices) after a single switch as a function of how long that run was. The discovered update mechanism reflected in the synthesis program predicts that a single switch should “reset” the perseveration trace following long runs in particular. Accordingly, we see that the probability of returning to the run rises with the length of that run for runs of lengths 5 or less, then begins to decrease. The synthesis program’s generated data shows a similar pattern, while the baseline’s continues to rise mostly monotonically.

### 4.1 Human Bandit

In this experiment, human subjects choose repeatedly among four options by pressing one of four keys on their computer keyboard, and receive rewards indicated by a number of points between 1 and 100 (Figure 2A). A question in the literature regarding this behavior concerns the role of memory: recent analyses of this dataset and others suggest that humans may rely on rapid memorization of rewards received for chosen actions in working memory [10, 44] rather than the classically described incremental trial-and-error reward learning mechanism [39]. We noted upon first reading the discovered programs that the learning update on the chosen action value appeared to be an incremental error-driven rule. However, automatic ablations revealed that this could be simplified to a working memory rule where the previous value is overwritten, being replaced by the value of the reward received (Figure 5A.1). In contrast, we did see incremental updating on the values of the *unchosen* actions, which decayed gradually toward a recency-weighted average reward (Figure 5A.2). This means that recent rewards for one action positively update all other action values. Together, these update patterns make a surprising prediction about how past rewards affect the tendency to repeat a choice: the immediately previous reward should cause a tendency to repeat choice, but all other past rewards should promote switching, because they increment the alternative choices and are overwritten for the current choice. In contrast, a key signature of standard error-driven learning rules like our baseline model is the tendency to repeat choices that have led to higher rewards in the recent past [42]. We designed an additional lagged regression model to check for this pattern, and found that it is indeed present in the dataset (Figure 7A).

The discovered programs also exhibited novel patterns of forgetting on their perseveration variables. Models in the literature which incorporate perseveration [45] typically incrementally update choice statistics and use this to implement a tendency to repeat recently selected actions. In two of the evolved models (one low- and one medium-floor program), the perseveration trace had a mechanism that instead could reset to zero if that action went unchosen a single time (Figure 5A.3-4). This mechanism makes another surprising prediction validated in data: after a long run of choices of the same action, even a single choice of a different action entirely erases the perseverative tendency to repeat that choice. The model predicts that the longer the run (after an initial saturation point), the lower the chance of returning to it; this is in contrast to standard models, according to which longer runs predict strictly more likely returns. We verified this pattern in the dataset (Figure 7B).

#### 4.1.1 Rat Bandit

In this experiment, rats decide between two nose ports and receive binary rewards whose probabilities followed independent random walks for each port (Figure 2B). The baseline model developed for this task included three learning systems operating fully independently, each governing the update of a single decision variable, and the decision on each trial was based on the sum of these variables [8]. While the discovered programs did consistently contain multiple learning systems, these systems did not operate independently. Instead, they typically interacted, either serving as update targets for one another or nonlinearly modulating the impact of one another on choice (Figure 5C.1-3). The discovered programs also differed from the baseline model in that each learning system updated not a single decision variable representing a preference between the two actions, but instead a pair of variables each representing the value of one of the actions. While this was missing from the baseline model, it is a feature that is common in other models for datasets of these kinds [2, 39, 41]. They also differed in a number of details about the balance of learning and forgetting as well as in the relative influence of rewards and reward omissions on learning within each system.

#### 4.1.2 Fly Bandit

In this experiment, fruit flies decided repeatedly between two odors, indicating their choice by entering an arm in a Y-maze that was filled with this odor. They received binary rewards whose probabilities changed independently in blocks (Figure 2C). The evolved programs departed from the baseline models in that they exhibited a typical error-driven learning rule modulated by an atypical, nonstationary learning rate (Figure 5D). The evolved programs with highest quality-of-fit made use of “eligibility traces”, which dynamically modulated learning rate depending on how frequently an option had recently been chosen. This motif is novel with respect to models of learning behavior in flies (Figure 5D.2). Nearly all of our discovered programs maintained a form of reward history, which was also used to modulate learning and/or update rates (Figure 5D.3).

#### 4.1.3 Monkey Bandit

In this experiment, monkeys chose among three images on a screen and received binary rewards with a fixed probability for each image (Figure 2B). Periodically, a novel image with an unknown reward probabilities was exchanged for one of the existing images. This design allows studying how novelty preferences interact with reward learning to guide exploration. The discovered programs outperformed the handcrafted baseline model by restructuring how novelty preference interacts with reward-guided learning (8/9 programs). Previous models largely assumed a single action value tracking average reward which was initialized, for new options, with a fixed novelty bonus [35, 46]. This reflects a substantive theoretical idea about the neural mechanisms of exploration: that the value of exploring a novel option is accounted in common currency with other (e.g., primary) rewards, and processed equivalently as action value by the same brain systems such as dopamine [47]. Empirically, though, this unified approach consistently underestimates the monkeys’ initial novelty seeking (bottom panel of Figure 6), since in the combined model it is forced to decay at the same rate as other rewards. The discovered programs solve this problem, by replacing the unified values with two separate cognitive variables: “action values”; tracking average reward, and a decaying “novelty trace”; which independently tracks perceptual novelty (Figure 5B.1). Instead of the common-currency assumption, this architecture reinforces the theory that novelty-driven exploration instead relies on dissociable cognitive streams [46, 48], which in turn has testable implications for the neural correlates of these variables [36, 48–50]. Consistent with the theme of discovered nonlinear, nonstationary updates, all discovered programs (9/9) also exhibited a nonstationary, nonlinear operation in which the decision variables are scaled by the variance of the action values, which has the effect of making behavior more exploratory (i.e. more stochastic) when the action values are similar.

#### 4.1.4 Rat Two-step

In the *Rat Two-step* experiment (Figure 2E), the connection between a rat’s action and resulting reward is mediated by a stochastic intermediate state, so as to distinguish between *model-based* reinforcement learning (which forecasts action values indirectly via the environment’s dynamics) and *model-free* reinforcement learning (which does not). Accordingly, discovered programs typically use both model-based and model-free action values, though the model-free values can typically be ablated with little or no effect on quality-of-fit [37, 38]. Similar to the *Rat Bandit* dataset (above), all 9 discovered programs differed from the baseline [37, 38] in representing the action values as two-dimensional vectors (one for each option) rather than a single variable summarizing relative preference (Figure 2E.1). As with *Rat Bandit*, this distinction in the learned representation has behaviorally detectable consequences because it allows the model to implement asymmetries: in this case, to decay the reward history for unchosen actions asymmetrically from chosen ones. Discovered models also often agreed (3/6 of low and medium-floor programs) on a specific form for this decay (with shared forgetting for both options, applied before the learning update for the chosen one), which differed from previous models in the literature [39]. Finally, also as in other datasets, nonstationary modulation was sometimes noted; for instance, two discovered programs incorporate a dynamic inverse temperature parameter that increases the entropy of the first choices in each session, perhaps reflecting a difference in strategy early in the session [38].

## 5 Discussion

The fundamental goal of basic science is not prediction or control, but understanding. Applying AI tools to solve basic science problems therefore requires centering human understanding as a key output. An increasingly popular approach for AI modeling of scientific data is to build “foundation models”; that is, to train large-scale neural network models on diverse large-scale datasets [51–54] in order to simulate the system’s behavior in different settings. This approach centers predictive performance as the primary target of optimization and primary metric of success. Scientific understanding might come from probing these models post-hoc, for example using the tools of mechanistic interpretability, but this hope remains somewhat speculative. Our approach uses large-scale neural networks not as models in themselves but as tools for generating models, expressed in a human-readable form and optimized both to fit data well and to be simple.

Model discovery for cognitive science requires discovering stateful computational models from indirect observations. Recent work has developed interpretable neural network approaches to this problem [9–11, 55–57]. Neural models are appealing because they are easy to optimize, but lack the explainability, generalizability, formulaic precision, and identifiability of symbolic models. Symbolic models are more difficult to optimize as one cannot use gradient descent. Existing symbolic regression (SR) [58, 59] and program induction [60–63] use discrete optimization techniques to discover closed-form symbolic models. These methods require handcrafted libraries of basis functions and operations which simultaneously makes the search problem potentially tractable and limits their expressivity [62, 63]. Symbolic regression approaches also cannot directly be applied to problems like those we study here, where the timeseries being modeled are not directly observed (though see [64, 65]). Other recent work has used LLMs to discover symbolic models, as they can easily generate programs in general purpose programming languages [16–19, 66, 67]. Very recent work, including from our group, has applied LLM program synthesis to discover symbolic models from data [28, 29, 68]. These approaches have not aimed to identify programs that match the quality-of-fit of blackbox models. We demonstrate that DataDIVER is capable of discovering symbolic models of latent dynamics from data using general-purpose tools, and to produce novel predictions that can be verified in the data.

The discovered models are readily interpretable as mechanistic hypotheses. Interpreted in this way they bear both striking similarities and important differences to hypotheses that are currently popular in the field. For example, it is common in the literature to use quite similar models in tasks that are implemented using diverse species (humans, monkeys, rats, fruit flies), available actions (button presses, eye movements, whole body movements) and rewards (points, sugar water, optogenetic stimulation). In contrast, we find that the models discovered for the different datasets are quite different. For example, the *Human Bandit* model does not include incremental reward learning, and may resemble a working memory process [10, 44], and the *Fly* and *Rat Bandit* models contain multiple learning mechanisms operating at different timescales [8]. This suggests that these conceptually-similar reward learning tasks may recruit different cognitive algorithms, and therefore different neural mechanisms [69].

Despite this diversity, there are at least two common themes in how the discovered models depart from the literature. First, they frequently introduce novel cognitive variables, such as the novelty tracking in the monkey dataset and the eligibility traces in the fly dataset. Interpreted as mechanistic hypotheses, these make the prediction that there are distinct neural correlates of these novel variables, and that neural perturbation experiments to specific brain regions might specifically affect the aspects of behavior that they mediate in the model. Second, discovered models frequently introduce nonlinear transformations between cognitive variables that are propagated through time and the computation of choice. These make the prediction that learning processes and choice processes in the brain may be more separate than is typically thought.

An important note of caution is that, as with any data-driven discovery tool, there is no guarantee that models discovered by DataDIVER will correspond with the true cognitive mechanisms used by the brain. Instead, they are best viewed as candidate hypotheses, which must be evaluated by human scientists to determine which, if any, contain plausible new ideas. That they take the form of computer programs facilitates this evaluation process, and makes it easy to mix and match ideas between models. The ultimate test of a model is whether it can successfully make predictions for new experiments involving radically different types of data – for our models this might take the form of very different behavior experiments in the same species and experimental setups, of neural recording experiments seeking correlates of the novel cognitive variables our models identify, or neural perturbation experiments asking whether focal changes to the model result in simulated behavior that resembles that of animals with focal perturbations to their brains.

While we have applied DataDIVER to questions about reward learning, the tool is general. Many scientific fields, for example ecology, epidemiology, and economics, require inferring the structure of indirectly observable latent processes in situations where data is plentiful but theories explaining the data are limited. To date, AI for scientific discovery has largely focused on prediction, forecasting, or mathematical problems that can be formulated as maximizing a single score with a black-box model [52, 53, 70]. They have steered clear of scientific problems where a breakthrough consists instead of a novel explanation. Generative AI’s ability to automatically generate artifacts that humans can readily inspect and understand, be it computer code, natural language, or images and charts, provides a new opportunity to start taking on these new types of discovery problems. Our work represents a step toward a vision in which these tools are used to help scientists make sense of the natural world as captured by data.

## Supporting information

DataDIVER Discovered Programs

DataDIVER Ablation Plots

## Acknowledgments

We would like to thank Ferran Alet, Alhussein Fawzi, Bernardino Romera-Paredes, Kyle Levin, Siddhant Jain, Kuba Perlin, Esteban Real, Carter Wendelken, and Doina Precup for helpful conversations. We would also like to thank Gheorghe Comanici for thoughtful comments on the manuscript. We would like to thank the AlphaEvolve team [19], and especially Alexander Novikov, Ngân Vũ, Maria Cardoso, Matej Balog, and Po-sen Huang, for helpful conversations and technical support. We would finally like to thank all of Google DeepMind for the support.

The work at Janelia Research Campus was supported by funding from the Howard Hughes Medical Institute (G.C.T, R.M., A.D.). We thank Kaitlyn N. Boone (Janelia Project Technical Resources) for assistance with fly behavior assays. This research was also supported in part by the National Institutes of Health awards R01 MH125824 and P51 OD011132 (V.D.C.).

Finally, we would also like to thank the Python community [71, 72] for developing tools that enabled this work, including NumPy [73], Matplotlib [74], Jupyter [75], Pandas [76] and JAX [33].

## 6 Methods

### 6.1 Dataset structure

Each dataset consists of data from multiple *subjects*, with each subject having completed multiple *sessions* comprised of a sequence of *trials*^1^. Across all datasets, a trial entails a subject making a *choice* and receiving a *reward* based on that choice. Two of the datasets include additional information specific to the experimental design. For the *Monkey Bandit* dataset, in which images representing choices are occasionally exchanged for new images with unknown reward, trial information includes about which (if any) options have been replaced by a novel option. For the *Rat Two-step* dataset, in which the rat’s reward is mediated by an observed outcome which is stochastically sampled based on the rat’s choice, trial information includes this outcome. For each dataset, we follow Castro et al. [28] in assigning half of the subjects (those with even indices) to the training split and using them for program generation, and holding out the remaining subjects for post-hoc program evaluation only.

Individual datasets are described in more detail in Appendix A.

### 6.2 Program structure

Each program accepts as input parameters, information about the current trial, and the previous agent state, and outputs a probabilistic prediction about the next choice as well as an updated agent state. The parameters allow the program to modify its behavior to match individual participants, and do not change over time or across sessions. The names and roles assigned to each parameter are not set ahead of time, and are instead established by DataDIVER. The trial information contained recent choice and recent reward, as well as dataset-specific information for *Monkey Bandit* (novel option, if any) and *Rat Two-step* (recent outcome). The previous agent state was required to be an array that did not change in size across or within sessions; however, the dimensions of the array and the names and roles assigned to agent state variables could be determined by DataDIVER. The agent state had a null value if none was provided, meaning that the program had to define the agent state during the first trial.

All programs were written in JAX [33] so that parameters could be optimized with gradient descent.

### 6.3 Quality-of-fit

We follow the bi-level cross-validation process of Castro et al. [28] in which an outer loop optimizes programs across subjects and an inner loop optimizes per-subject parameters across a subject’s sessions. Parameters were optimized using two-fold cross-validation, with folds comprised of even and odd sessions, respectively. This involved fitting parameters to each fold by maximizing the summed log likelihood of the all choices under the program’s predictions, and validating the parameters on the opposite fold, producing two log likelihood validation scores. To produce a single score for the subject, the two log likelihood scores are summed, divided by the total number of trials across both folds, and exponentiated to produce a “normalized likelihood” score. This can be interpreted as the geometric average probability that the model would have made the choices that the participants made [2]. Our final “quality-of-fit” score for each program is the average of these normalized likelihoods across participants. Unless specifically indicated (e.g. in Figure 1B,C), quality-of-fit scores are always reported on held-out test subjects that were not used for program evolution.

Because in the *Fly Bandit* experiment each fly only participated in one session. Thus, both programs and parameters were optimized across subjects, and could be thought of as capturing fly behavior in aggregate. This is described in Appendix A.3.2. Maximum likelihood parameters were estimated using gradient-based optimization, with initial parameters sampled uniformly from *U*(−2, 2). Gradient descent terminated if the convergence criterion was met, the maximum number of steps *M*_gd_ was reached, or the score became undefined (e.g., due to exploding state variables).

To address the issue of nonconvexity, we conducted multiple fitting attempts per dataset using different initializations. After the minimum number of fitting attempts *m*_fit_ occurred, the process terminated once *n*_fit_ runs converged to values near the best obtained score *s*^∗^, defined as scores *s* satisfying 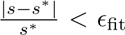, or once *M*_fit_ total attempts had elapsed.

During training, we used a set of optimization hyperparameters tuned for speed. For the final evaluation and reported results, we employed a more stringent set of hyperparameters and a different gradient-based optimizer (L-BFGS instead of AdaBelief) to ensure accuracy. The convergence criteria also differed: during training, we checked the score every *k*_gd_ steps and declared convergence if 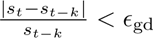, where *s_t_* is the current score and *s_t_*_−*k*_ is the score from *k* steps prior. During evaluation, we used the gradient infinity norm, declaring convergence if 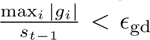, where *g_i_* is the gradient of the *i*-th parameter.

Additionally, we used distinct random seeds for evaluation to ensure results did not reflect overfitting to a specific initialization order. Because of the time-consuming nature of evaluating automatic ablations, these were computed using the training hyperparameters and seeds. See Table 1 for the complete list of optimization parameters.

**Table 1:**
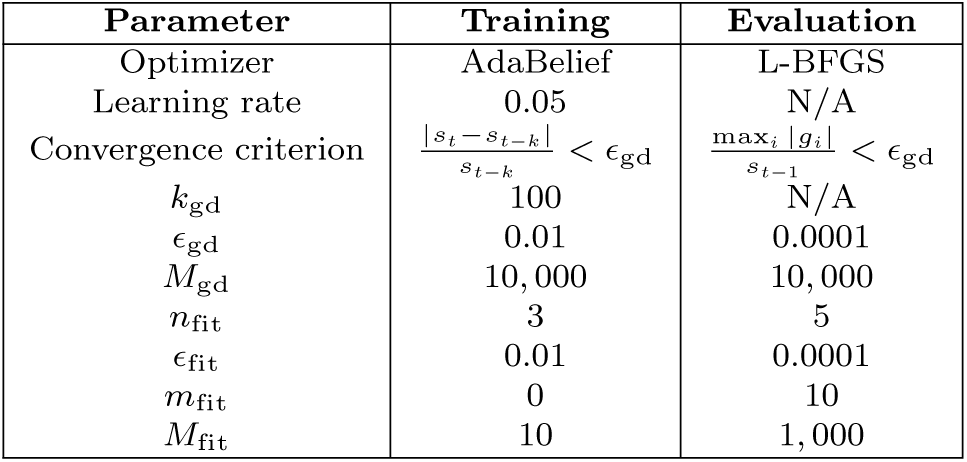
Optimization hyperparameters used to compute quality-of-fit.

Parameter optimization was done using custom code implemented in Python and JAX.

### 6.4 Halstead complexity metrics

We use Halstead complexity metrics to capture program complexity [30]. These are syntax-only analysis measures developed to quantify software complexity, maintainability, and size based on the number of operators and operands in source code. Our pipeline involves three of these metrics–volume *V*, difficulty *D*, and effort *E*– which can be defined in terms of the number of distinct operators *η*_1_, distinct operands *η*_2_, total operators *N*_1_, and total operands *N*_2_:

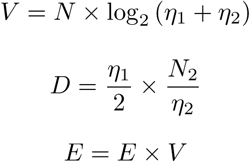

Volume represents the program’s total information content, difficulty represents the code density/obfuscation, and effort represents the mental work of implementing the code.

### 6.5 Optimizing programs with AlphaEvolve

Program optimization was performed using AlphaEvolve [19], which implements an evolutionary algorithm that optimizing programs using large language models as the mutation operator. The language model used for all AlphaEvolve runs was Gemini 2.5 Flash [77]. A simple dead code elimination algorithm was applied to each generated program to remove unused or overwritten variable assignments. All AlphaEvolve runs were halted after 100, 000 steps.

### 6.6 Two-stage program optimization

DataDIVER consists of two stages: a “Maximize Quality-of-Fit” first stage, and a “Simplify” second stage. Each stage involved a separate AlphaEvolve run with different objectives. Three independent runs of the entire two stage process were conducted, starting from different seeds.

Performing the multiobjective optimization in two stages permits discovery of models in the high quality-of-fit, high complexity region of the model space before simplicity constraints are introduced. This is similar to the strategy of KL-annealing for training variational autoencoders, wherein models are trained such that fit-to-data is prioritized early in training and complex latent activity is penalized later to prevent posterior collapse [78].

#### 6.6.1 Stage 1: “Maximize Quality-of-Fit”

In the first stage of DataDIVER, “Maximize Quality-of-Fit”, the target of optimization is the average normalized likelihood (described in Section 7.3). The LLM is prompted with text that contains context on the computational modeling task, examples of prior programs, a “parent program” that it will edit, and instructions for how to format edits (see the complete prompt in section C.1. Instructions suggesting different strategies for editing the program (e.g. “combine the strengths of the programs above”, “implement an idea not present not present but commonly used in the literature.”) are sampled stochastically (see Table C8 for full specifications).

The product of this stage is a vast set of programs generated by the LLM. The program with the highest quality-of-fit is referred to as the “fit only” program because it was optimized with only quality-of-fit in mind. The set of programs on the “Pareto frontier” of quality-of-fit versus Halstead effort are also extracted and used to initialize stage 2. The Pareto frontier refers to programs that are optimal for some trade-off of the two metrics; more specifically, any program for which no other program exists that is better at one metric without being worse at the other.

#### 6.6.2 Stage 2: “Simplify”

In a second “Simplify” stage, the Pareto frontier programs from the first stage are further evolved in order to better trade off complexity and quality-of-fit. Here, the LLM is prompted to simplify the program rather while maintaining similar quantitative performance (see the complete prompt in Section C.2, with stochastically sampled prompts suggesting different strategies for simplifying the code (see Table C9). AlphaEvolve was run in multiobjective mode with both Halstead difficulty and volume as optimization metrics at this stage. For each “Simplify” stage we specified a quality-of-fit floor, and any generated programs with quality-of-fit scores below this floor were discarded. Floors were selected to lie at fixed fractions *α* of the gap between the fixed hand-crafted baseline model’s quality-of-fit score *s_B_* and that of the best program discovered across all three runs *s*^∗^, such that each floor can be written as *s_B_* + *α*(*s*^∗^ − *s_B_*). For each of the three DataDIVER runs, we run this “Simplify” AlphaEvolve run three independent times with different values for *α*: a “low floor” *α* of 50%, a “medium floor” *α* of 75%, and a “high floor” *α* of 90%. From each “Simplify” run, we extract the program with the best Halstead effort, leaving us with nine programs in most cases (3 DataDIVER runs × 3 “Simplify” runs each). Occasionally, due to variance in performance of the best programs in the best first-stage runs, some stage 1 runs did not exceed the *α* = 90% performance threshold; these runs thus fail to produce any valid programs, resulting in some datasets having fewer than 9 total program simplification experiments.

For each of the (at most) 9 program simplification runs, we select the generated program with the minimum Halstead effort (the product of the difficulty and volume metrics optimized above). We then apply a final readability refactoring step, which involves prompting Gemini 2.5 Pro to rewrite the program to be more readable while ensuring the program’s functionality is unaffected (the rewriting prompt is specified in Section C.3). A hash function is used to ensure that the program’s functionality is unaffected. For each input program, a hash was computed by computing the output logits and state values across 100 rollouts of length 10 with randomly sampled parameters and inputs. Discrete inputs (e.g. choices) are sampled from a uniform multinomial distribution, continuous inputs (reward in *Human Bandit*) from a uniform distribution on the [0, 1] interval, and parameters from a normal distribution with mean 0 and standard deviation 1. The flattened and concatenated vector of outputted logits and state values constitutes an approximately unique signature of program behavior. The refactor is deemed successful if a program is generated whose hash matches the hash of the un-rewritten program, given the same random inputs. Rewriting is attempted a maximum of ten times, and with a deadline of 600 seconds total. Refactor succeeded for 41 out of 43 programs; the other two (*Human Bandit*, high floor, run 2; *Rat Two-step*, low floor, run 2) were kept in their unrefactored form.

### 6.7 Program evaluation

For each dataset, up to twelve total programs are evaluated: the three “fit-only” programs, which are optimized for quality-of-fit only and are outputted by the “Maximize Quality-of-Fit” stage (3 programs per dataset); and the simplified programs that emerge from the “Simplify” stages (up to 9 programs per dataset). For each of these programs, we calculate the quality-of-fit for the held out evaluation subjects, again using cross-validation procedure across each subject’s even/odd sessions as described in Section 7.3. Note that this entails two levels of validation: programs are validated on never-before-seen subjects, while parameters are validated across sessions for each subject.

### 6.8 Automatic ablations

In order to test which of a program’s many operations contributed to the program’s performance, we implemented an automated ablation procedure. This consists of generating all programs that modify the input program in one of the following ways:

- deleting a line of code; (e.g. removing a line x = x + y)
- setting the right-hand side of one variable assignment to zero or a size-matched array of zeros (e.g. with x = jnp.zeros like(x + y));
- setting the right-hand side of one variable assignment to one or a size-matched array of ones (e.g. with x = jnp.ones like(x + y));
- replacing a binary operator (e.g., + or *) with either its left or its right argument (e.g. x = x + y → x = x).

We then recompute the quality-of-fit across evaluation subjects for each of these programs, and measure the drop in score. This is useful for identifying whether any of the computations that the “Simplify” stage failed to prune are in fact unnecessary, and understand how much each operation is contributing to performance in terms of magnitude of explained likelihood and significance across subjects.

### 6.9 Recurrent neural network (RNN) baselines

We compare the programs generated using DataDIVER to recurrent neural network (RNN) baselines. The RNNs each consist of one gated recurrent unit (GRU) layer followed by a linear readout layer [79].

For the *Rat Bandit*, *Monkey Bandit*, and *Rat Two-step* datasets, in which each subject participated in a large number of sessions, separate RNN models were trained for each subject. This involved performing two-fold cross-validation by training one network on that subject’s even-indexed sessions and evaluating on that subject’s odd sessions (and vice versa). We performed a sweep over the learning rate (0.00001, 0.0001, 0.001), the number of hidden units (1, 2, 4, 8, 16, 32, 64, 128), and three random initialization seeds in order to find the optimal hyperparameters for each dataset. In all cases, we train for 100, 000 learning steps, and log parameters and quality-of-fit every 100 steps. The quality-of-fit, cross-validated over sessions, is computed for the even-indexed training subjects (the same set of subjects used for generating programs in DataDIVER) and used for hyperparameter selection: we select the learning rate, the number of hidden units, and an early stopping point, that maximize quality-of-fit across subjects and random seeds. The reported RNN performance is on the odd-indexed evaluation subjects (the same subjects on which we DataDIVER programs), using the hyperparameters and early stopping point selected from the even-indexed subjects. We report the results of the best-performing random seed.

For the *Fly Bandit* and *Human Bandit* datasets, in which each subject participated in few sessions (*Human Bandit*) or one session (*Fly Bandit*), there were not enough sessions to train and validate a separate RNN for each subjects [28]. Sessions were therefore combined across subjects, and RNNs were trained on this aggregate data. Even-indexed subjects’ sessions were combined into one training split, and odd-indexed subjects’ sessions were combined into an evaluation split. Note that this preserved the train and evaluation subject splits used by DataDIVER. For each split, these combined sessions were further subdivided in half for two-fold cross-validation. The cross-validated normalized likelihood computed for the training subjects was used for hyperparameter selection (learning rate, number of hidden units, and early stopping point). Reported performance scores are computed on the odd-indexed evaluation subjects. We note that rather than report the average performance across all combined sessions, we re-normalized across subjects for apples-to-apples comparison with DataDIVER programs. As with DataDIVER, we use the results for the best-performing random seed. The best-performing hyperparameters for each dataset can be found in Table 2.

**Table 2:** Hyperparameters which maximize RNN performance on even-indexed subjects or even split, per dataset.

| Dataset | # hidden units | Learning rate | Early stopping point (# steps) |
| --- | --- | --- | --- |
| *Human Bandit* | 16 | 0.0001 | 99800 |
| *Rat Bandit* | 128 | 0.001 | 200 |
| *Fly Bandit* | 128 | 0.001 | 200 |
| *Monkey Bandit* | 2 | 0.001 | 1900 |
| *Rat Two-step* | 128 | 0.001 | 100 |

### 6.10 Artificial Data Generation

In order to understand the behavior of the different models, artificial datasets were generated from each of DataDIVER-discovered programs, the synthesis programs, the handcrafted baselines, and the RNNs. This entailed running artificial experiments with the models and their optimized parameters using an *experimental environment* configured to match the outputs or possible outcomes of the experiment used to collect the real data.

To generate a simulated session of data, the model was initialized with the optimized parameters, a null agent state, the first action and reward observed in the real data, and any other relevant trial information, if applicable (outcome for *Rat Two-step*, novel option for *Monkey Bandit*). Optimized parameters used for artificial data generation were those obtained using cross-validation in order to score the models. This means that for each session of artificial data in one fold, the parameters used to generate that data were those obtained by optimizing parameters over the data in the opposite fold. On each timestep, the model would output an updated state and a probability distribution from which the next choice would be sampled. Whenever the choice matched the choice made by the animal, the same reward (and outcome, for *Rat Two-step*)) was observed; otherwise, rewards (and other outcomes) were sampled from the experimental environment given the experimental configuration. This was applied iteratively for as many trials as the animal completed in the corresponding session to collect artificial data. In all, five artificial datasets (with different random seeds governing the choice sampling) were generated per model.

The experimental environment was instantiated such that it would return rewards (and other outcomes, for *Rat Two-step*) that were observed in the real data whenever the model’s choices matched those of the real subject at the same trial. When choices differed, the environment would return rewards (and outcomes) sampled from the distribution specified by the experimental configuration (i.e. observations the subject might have seen had it made that choice). For *Monkey Bandit*, the experimental environment also contained when existing options were swapped out for novel options so that these were matched to the real data.

The rewards and other inputs are sampled in the following way, depending on dataset:

- For the *Human Bandit* dataset, rewards are determined by a payout matrix that indicates the exact reward that each participant would receive for each available choice in each trial of the experiment.
- The *Rat Bandit*, *Fly Bandit*, and *Monkey Bandit* datasets include the *probability of (binary) reward* for each (chosen or unchosen) choice in each trial of the experiment. For each trial of the synthetic experiment, if the choice does not match the choice made by the real subject, a binary reward is sampled according to the probability specified for the agent function’s choice in the equivalent natural trial.
- The *Rat Two-step* dataset also includes the probabilities of reward, but these depend only indirectly on the agent’s choice: instead, choices stochastically determine the binary *outcome*, and outcomes determine reward probabilities. For each trial, a choice could be “congruent” (choice and outcome located on the same side) or incongruent (choice and outcome located on opposite sides). When generating the artificial dataset, we first determine whether each trial’s choice and outcome were congruent or incongruent in the real data. This congruency relationship was preserved in the experimental environment regardless of the choice. Rewards are then sampled conditional on the outcome, using the reward probabilities specified for that trial in the natural dataset. When the sampled outcome matches the outcome observed in the real data, the rewards will also be matched.

### 6.11 Trial-Lagged Regression Plots

To determine the extent to which each model captured relevant features of the corresponding real data, we used trial-history regression analyses common in the literature [8, 37, 41, 42].

For the *Rat Bandit* and *Fly Bandit* datasets, the following logistic regression model was fit to the data (as in Castro et al. [28]):

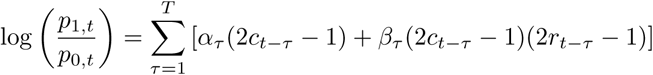

with the choices *c_t_* ∈ {0, 1}, rewards *r_t_* ∈ {0, 1}, and where we characterize 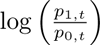, the log-odds of choosing *c_t_* = 1, in terms of the past choices and the past products of choice and reward (after normalizing both choice and reward be in {−1, 1}). The *α_τ_* coefficient characterizes the extent to which the subject tends to repeat the choice made *τ* trials ago; the *β* coefficients characterize the extent to which the subject tends to repeat rewarding choices and avoid non-rewarding choices from *τ* trials ago.

Because the *Human Bandit* and *Monkey Bandit* datasets are four- and three-armed bandit tasks respectively, we use a conditional logit regression model [80], where a linear model outputs a utility *U_i_*for each choice *i*, and choices are sampled according to the softmax of the utilities. For the *Monkey Bandit* dataset, the utility for choice *i* ∈ {0, 1, 2} is

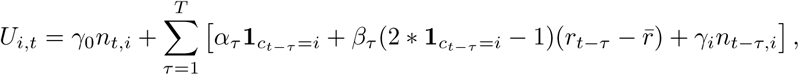

where **1***_ct_*_−_*_τ_* =*i* is 1 if the *τ*-back choice *c_t_*_−*τ*_ is equal to choice *i* and 0 otherwise, *r̅* is the mean reward over the entire dataset and *r_t_*_−*τ*_ – *r̅* is the mean-centered reward received *τ* trials back, and *n_t,i_* is 1 if choice *i* is the novel option at time *t* and 0 otherwise. Here *α_τ_*again represents the tendency to repeat a choice made *τ* trials back, *β_τ_* represents the tendency to repeat choices that yielded above-average reward *τ* trials ago and avoid those that did not, and *γ_i_*represents the tendency to select novel options added to the choice set *τ* trials ago.

The *Human Bandit* regression model is identical except it excludes the novel option terms and their coefficients.

For the *Rat Two-step* experiment, we repeat the trial-history lagged regression model introduced by Miller et al. [37]. In that experiment, the transitions between rats’ choices and the (observable) outcomes were stochastic, with one transition being *common* (happening 80% of the time) and one being *uncommon* (happening 20% of the time) for each choice (note: this is different from congruency, see Appendix A.5). The outcome that was common for one choice was the uncommon outcome for the other. Given choice at time *t c_t_* ∈ {0, 1}, reward *r_t_* ∈ {0, 1}, and *b_t_* ∈ {0, 1} indicating whether the transition at time *t* was a common transition, we can define regressors capturing relevant information for this task: CR(*t*) = (2*c_t_* − 1)*b_t_r_t_* (Common Reward, common outcome occurred and was followed by reward), CO(*t*) = (2*c_t_* − 1)*b_t_*(1 − *r_t_*) (Common Omission), UR(*t*) = (2*c_t_* − 1)(1 − *b_t_*)*r_t_* (Uncommon Reward), and UO(*t*) = (2*c_t_*−1)(1−*b_t_*)(1−*r_t_*) (Uncommon Omissions). The logistic regression model is defined as follows:

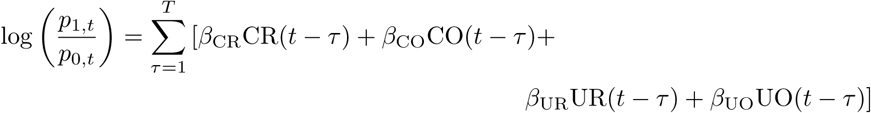

By breaking down the dependency of the current trial on past trials’ choices and rewards into terms depending on the commonness of the choice → outcome transition, we can distinguish between agents which update the values of choices in a model-free manner (and thus are more likely to repeat an action that resulted in reward, regardless if that reward resulted from the common transition) and those which engage in some planning (which are *less* likely to repeat an action that resulted in reward if that action was the result of an uncommon transition, since the outcome which is uncommon for one choice is common for the other).

The error bars for all trial lagged regression plots reflect the 95% prediction intervals. This indicates the range that is expected, with 95% probability, to contain the value of a single future observation.

### 6.12 Trial-lagged regression: repeating rewarding choices

To assess whether past reward promotes a tendency to stay with the same choice (*c_t_* = *c_t_*_+1_) or switch to a different choice (*c_t_ ≠ c_t_*_+1_), we performed a lagged regression model that was broken down into whether the *τ*-back reward had been received for a choice of the current arm (“same choice”) vs an alternative one (“different choice”). For this, the linear model could be written as:

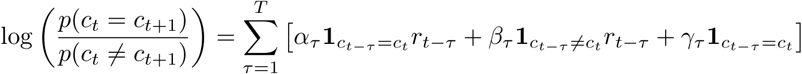

We choose *T* = 10. Figure 7a plots the values of *α_τ_* for *τ* ∈ {1, · · ·, 10}; the values of *β_τ_* and *γ_τ_* can be found in Figure A3 in the Supplement.

### 6.13 Run return probabilities

In order to test whether the discovered forgetting rule was present in the *Human Bandit* dataset, the probability of returning to a “run” (a sequence of repeated choices) after a single switch was observed. A run of length *n* is defined as a sequence in which the same choice is consecutively made for exactly *n* steps. For each session, all runs were identified and counted. For each run, we computed the number of times that the second choice made after ending a run was the same as the choice made during the run (the choice immediately following each run is, by definition, different). The fraction of runs in which the subject returned to the choice made during the run was computed for each run length and shown in Figure 7B. The error bars show 95% confidence intervals computed across all sessions and subjects combined.

### 6.14 Synthesis Programs

In order to unify the different discovered programs into a single programs, synthesis programs were manually constructed. We note that this step was not required to produce interpretable programs; rather, this allows us to verify that different motifs discovered across different programs do not interfere when combined and therefore can each be safely interpreted. Broadly, the procedure for constructing these programs was to first start with a discovered program, then remove elements that did not contribute to the overall likelihood (as assessed with automated ablations; see Section 7.8). Motifs from other programs were substituted in if they contributed to performance in other programs, or if they contributed to uniquely good performance for some diagnostic (as in *Human Bandit*, where the selected forgetting rule led to high performance on the Run Return statistics). Crucially, no extra motifs based on background knowledge of the literature were added.

## Appendix A Further Analysis of Individual Datasets

### A.1 Human Bandit

#### A.1.1 Details of the dataset

Eckstein et al. [10] consider human participants performing a four-alternative task with graded rewards. Participants performed the task online, and indicated their choice on each trial by pressing either ‘D’, ‘F’, ‘J’, or ‘K’ on their keyboard. Reward was indicated by displaying an integer number of ‘points’ between 0 and 100, which subjects were asked to maximize. Available rewards followed independent bounded random walks with additional trial-unique noise. Each participant performed up to five back-to-back sessions of up to 150 trials each. The dataset contains choices from 862 participants performing 4,134 total sessions and 617,871 total trials.

We obtained this dataset from the following URL, where it is freely available under a permissive open-source license: https://osf.io/8xz3w/

#### A.1.2 Handcrafted Baseline Program

Eckstein et al. [10] performed an extensive comparison of a wide variety of computational cognitive models on this dataset. Following that, we adopt a model we refer to as “Perseverative Forgetting Q-Learning” as the human-discovered benchmark model. Note that we normalize reward to be between 0.0 and 1.0 as input to all baseline and DataDIVER models.

#### A.1.3 Discussion of evolved programs

Subjectively, we found that the low- and medium-floor programs could be understood with low or moderate effort. The high-floor programs were far more complex, owing to their greater length and diminished modularity, and yielded fewer discernible insights. Example low, medium, and high-floor programs are provided in Supplement A.1.7.

##### Cognitive Variables

Two cognitive variables that consistently arose across discovered programs were action values (or “Q-values”), which tracked expected rewards for each choice, and a “perseveration trace”, which tracked choice history independent of reward (and went by a variety of different terms, e.g. “choice trace”, “recency trace”). In many discovered programs, these were the only cognitive variables defined (all low-floor programs, one medium-floor program (run 3), and one high-floor (run 3) program). Another cognitive variable that arose in two out of three medium-floor programs tracked average reward (independent of choice), which interacted with the Q-value updates and the decision variable (described in more detail below). An additional discovered cognitive variable was found in one medium-floor program (run 2) and used average prediction errors per choice to update Q-values. However, this program had a low quality-of-fit and performed poorly on diagnostics. Two of the three high-floor programs did define additional cognitive variables, although their role and contribution to quality of fit was not straightforward due to the complexity of these programs.

For all low and medium-floor programs, action values and Perseveration Trace were updated modularly, with no interactions between the reward-dependent action value update and reward-independent Perseveration Trace updates. This decomposition into modules parallels the modular hybrid neural network architecture identified by Eckstein et al. [10], who showed that neural network modules organized accordingly predicted behavior on this task better than other architectures. Here, we have the added benefit of being able to inspect these modules. This modularity broke down in high-floor programs, where reward-dependent terms are used to modulate perseveration updates.

##### Reward learning

Across all discovered programs, learning on the chosen action value *Q*(*c*) presented in the form of a standard reward prediction error driven update given reward *r*: *Q*(*c*) = *Q*(*c*) + *ϕ_i_*(*r* − *Q*(*c*)). However, this proved misleading: ablations revealed that for most programs, the equation can be reduced further into a less conventional but much more simple update, *Q*(*c*) = *r* with no loss in quality-of-fit. This is what we use for our synthesis program and discuss more in Section 4.1. Again, this parallels the observation from Eckstein et al. [10] that action values are not updated incrementally.

All discovered programs showed an update on all unchosen action values in which they slowly decayed toward some target that captured recent reward. While the particular statistic varied, this update generally has the effect of decaying *Q*(*c*^′^) toward the average over the recent reward history. For a given target *r̅* and an unchosen option *c*^′^, the updates had the form *Q*(*c*^′^) = *Q*(*c*^′^) + *ϕ*(*r̅* − *Q*(*c*^′^)). Across programs, discovered targets included the previous reward *r* (run 1 and 2, low-floor), in others toward *Q*(*c*) (run 3, all floors), and in others, toward a separate cognitive variable which tracked the average reward *r*_avg_ (run 1 and 2, medium- and high-floors).

##### Perseveration Updates

In many of our programs, as in many models in the literature, choice is not driven by past rewards (summarized by action values) alone, but also by a separate set of reward-independent mechanisms that capture a tendency to repeat actions that have been taken recently regardless of their outcome (perseveration).

The particular form of the discovered perseveration traces that implement this across our discovered programs showed interesting variations. Like the action values, the perseveration traces are 4-dimensional vectors in which each element corresponds to a different choice. All programs included some update that drives up the perseveration trace for the choice selected on the previous trial by incrementing it or setting it to 1. All programs implemented some kind of forgetting for the unchosen options, usually by decaying their perseveration traces towards 0 (7/9 programs). Collectively, this implements a tendency to repeat actions that were taken recently.

However, the remaining two discovered programs included an unusual forgetting pattern on the perseveration trace *P* for each unchosen option *c*^′^ in which *P* (*c*^′^) was *reset* to 0 rather than gradually *decayed* under certain conditions. This interrupts the perseverative bout if even a single different action is taken.

In low-floor (run 1), this amounted to simply resetting *P* (*c*^′^) to 0, erasing the value that had been accumulated for this variable when it had been chosen. This suggests that there is no extra perseverative momentum on an action once even a single different action is taken, even if that action has been taken many times prior to the deviation. There still might be some above chance likelihood of returning to that action, but it would have to be driven by the action values.

In medium-floor (run 3), the perseverative update had a more complex form of reset. The perseveration update for chosen options involved incrementing *P* (*c*), clipping it at 5.0, and scaling by a positive parameter *ϕ_c_*: *P* (*c*) ← *ϕ_c_* min{(*P* (*c*) + 1), 5.0}. The update on unchosen options is *P* (*c*^′^) ← *ϕ_u_P* (*c*^′^)(5.0 − *P* (*c*^′^)) for positive parameter *ϕ_u_*. This means that *P* (*c*^′^) will be reset to 0 when *P* (*c*^′^) is close to 5, and remain positive if it is between 0 and 5. Thus, this reset only occurs for very long runs, where *P* (*c*) has accumulated to the clipped value of 5. Intuitively, this means that after the subject has been making the same choice for a long time, their making a different choice means that they have stopped perseverating on that run: there is no extra perseverative momentum on that choice. However, for short runs (*<*5), some perseverative momentum remains.

##### Nonlinear, Nonstationary Exploration

In the medium- and high-floor programs, action values undergo a nonlinear, nonstationary transformation before merging with the perseveration component. This has the effect of making behavior more deterministic when action values are further from 0 or from the nonstationary average recent reward. This means that the nonlinearity mapping action values to decision variables is not merely a softmax.

#### A.1.4 Discussion of synthesis program

The synthesis program included versions of each of the motifs described above. It defined three cognitive variables: action values, a perseveration trace, and a term which tracked the recent average reward. The chosen action value was updated by being overwritten by the received reward; as such, there was no learning rate parameter needed to update the chosen action’s value. The unchosen action values were updated by decaying toward the recent average reward. The nonlinearity applied to the action values before their combination with the perseveration trace was

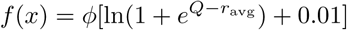

which was the particular nonlinearity from the medium-floor, run 1 program.

The perseveration trace was updated using the nonlinear forgetting rule from medium-floor, run 3, as this program exhibited the best performance at recovering the run return diagnostic.

The synthesis program is shown in Supplement A.1.5.

#### A.1.5 Code: synthesis program

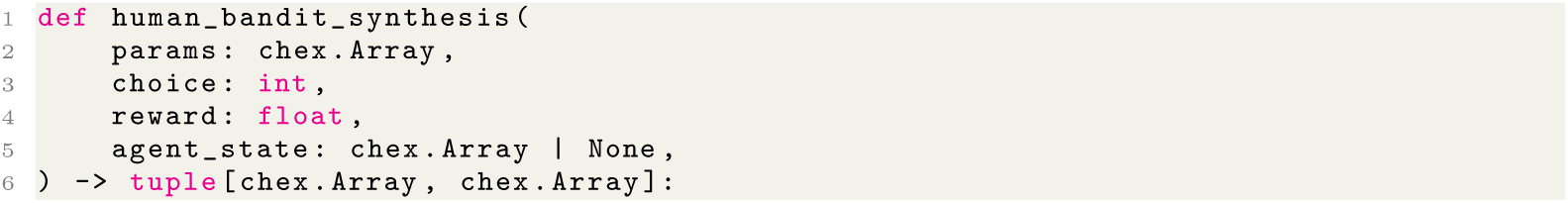
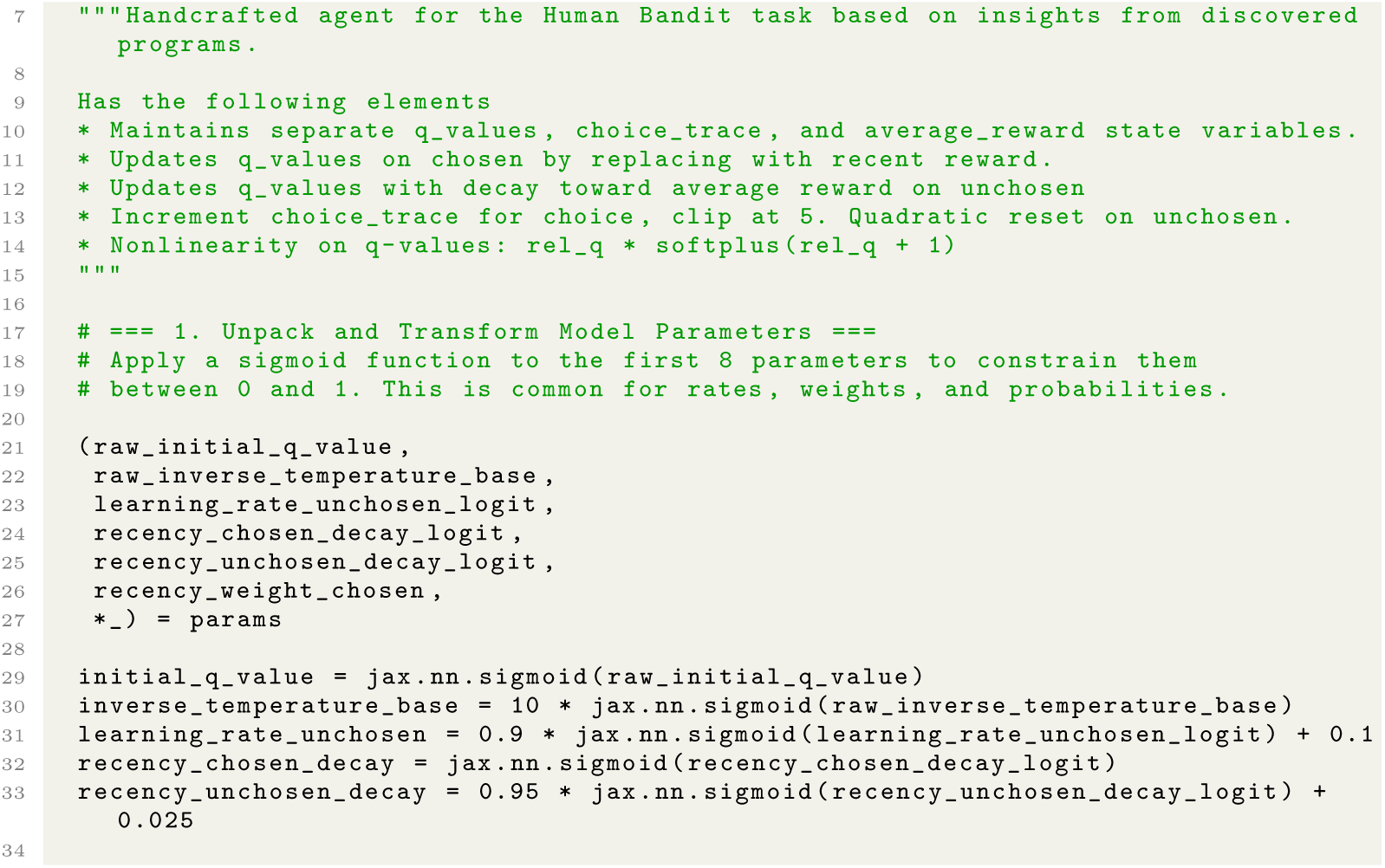
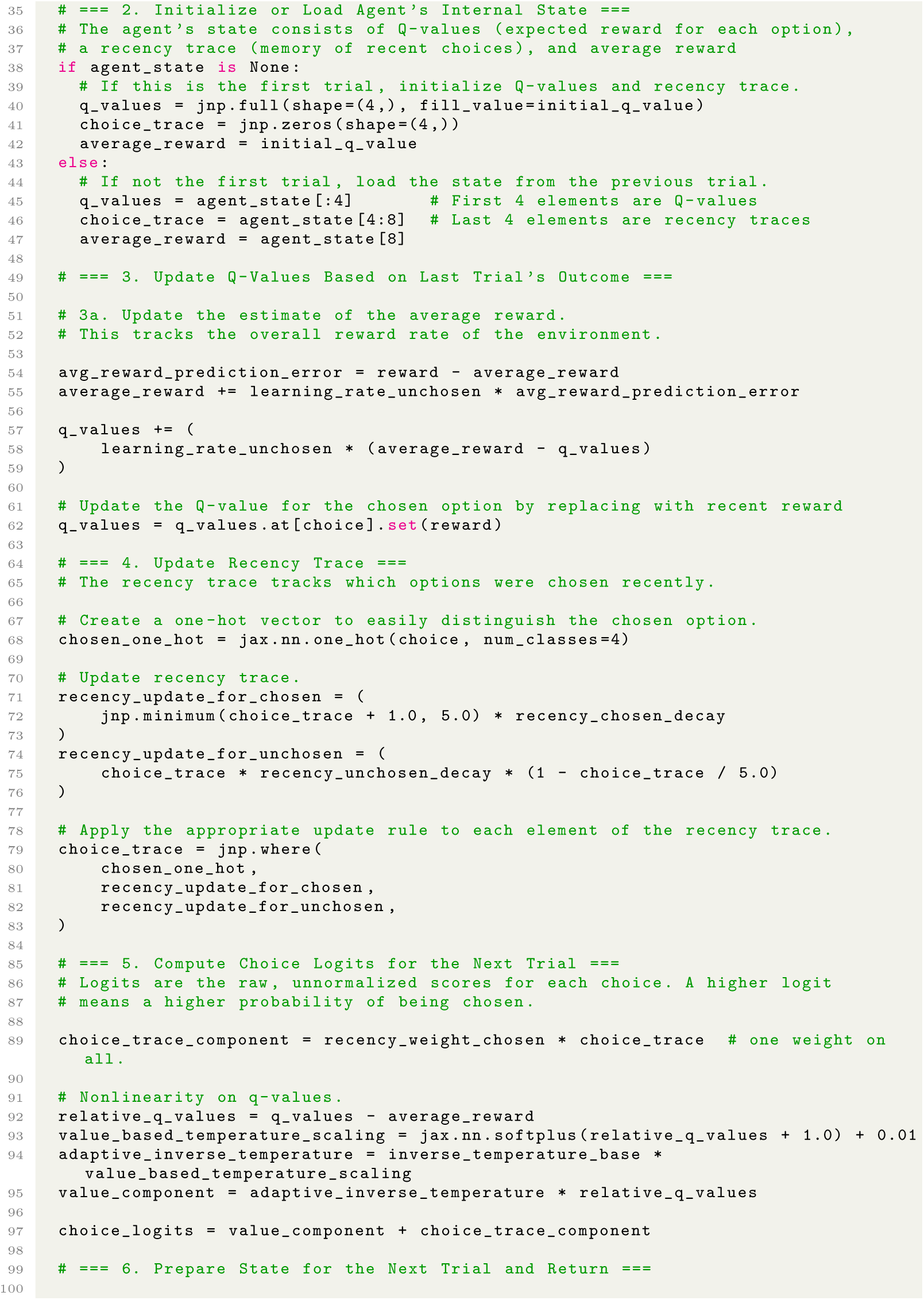
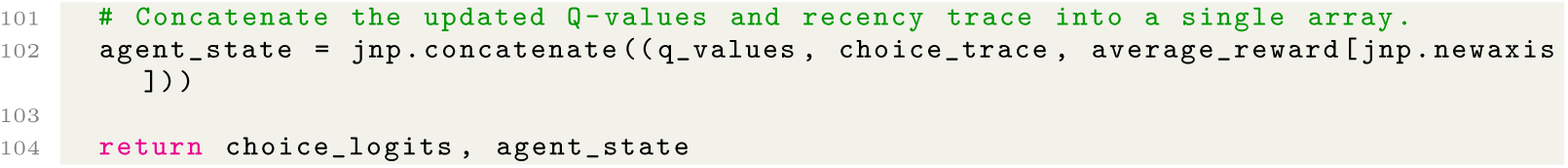

**Table A1:** Evaluation performance and program complexity for models in the *Human Bandit* dataset. For programs generated by the “Simplify” stage, Floor represents the quality-of-fit threshold below which programs are discarded (see Section 7.6.2). Score indicates the average normalized likelihood across evaluation subjects (see Section 7.3); Effort is Halstead effort. State, Params, and Lines indicate the number of state variables, per-subject parameters, and lines of code respectively.

| Model type | Floor | Run | Score | Effort | State | Params | Lines |
| --- | --- | --- | --- | --- | --- | --- | --- |
| Handcrafted Baseline | – | – | 0.5610 | 16,197 | 4 | 4 | 59 |
| RNN Baseline | – | – | 0.6274 | – | – | – | – |
| Stage 1: “Maximize Quality-of-Fit” | – | 1 | 0.6188 | 703,382 | 20 | 10 | 350 |
|  |  | 2 | 0.6209 | 780,146 | 30 | 10 | 555 |
|  |  | 3 | 0.6175 | 112,167 | 8 | 10 | 169 |
| Stage 2: “Simplify” | 50% | 1 | 0.5988 | 9,645 | 8 | 7 | 108 |
|  |  | 2 | 0.5998 | 13,660 | 8 | 7 | 114 |
|  |  | 3 | 0.5929 | 7,484 | 8 | 6 | 99 |
|  | 75% | 1 | 0.6075 | 21,296 | 9 | 10 | 141 |
|  |  | 2 | 0.6103 | 31,285 | 13 | 10 | 154 |
|  |  | 3 | 0.6111 | 22,647 | 8 | 9 | 138 |
|  | 90% | 1 | 0.6178 | 122,938 | 13 | 9 | 210 |
|  |  | 2 | 0.6166 | 157,612 | 22 | 10 | 307 |
|  |  | 3 | 0.6170 | 85,125 | 8 | 9 | 184 |
| Synthesis Program | – | – | 0.6068 | 25,047 | 9 | 7 | 106 |

Code 1: Synthesis program for the human bandit dataset.

#### A.1.6 Code: stage 1 (“Maximize Quality-of-Fit”) programs

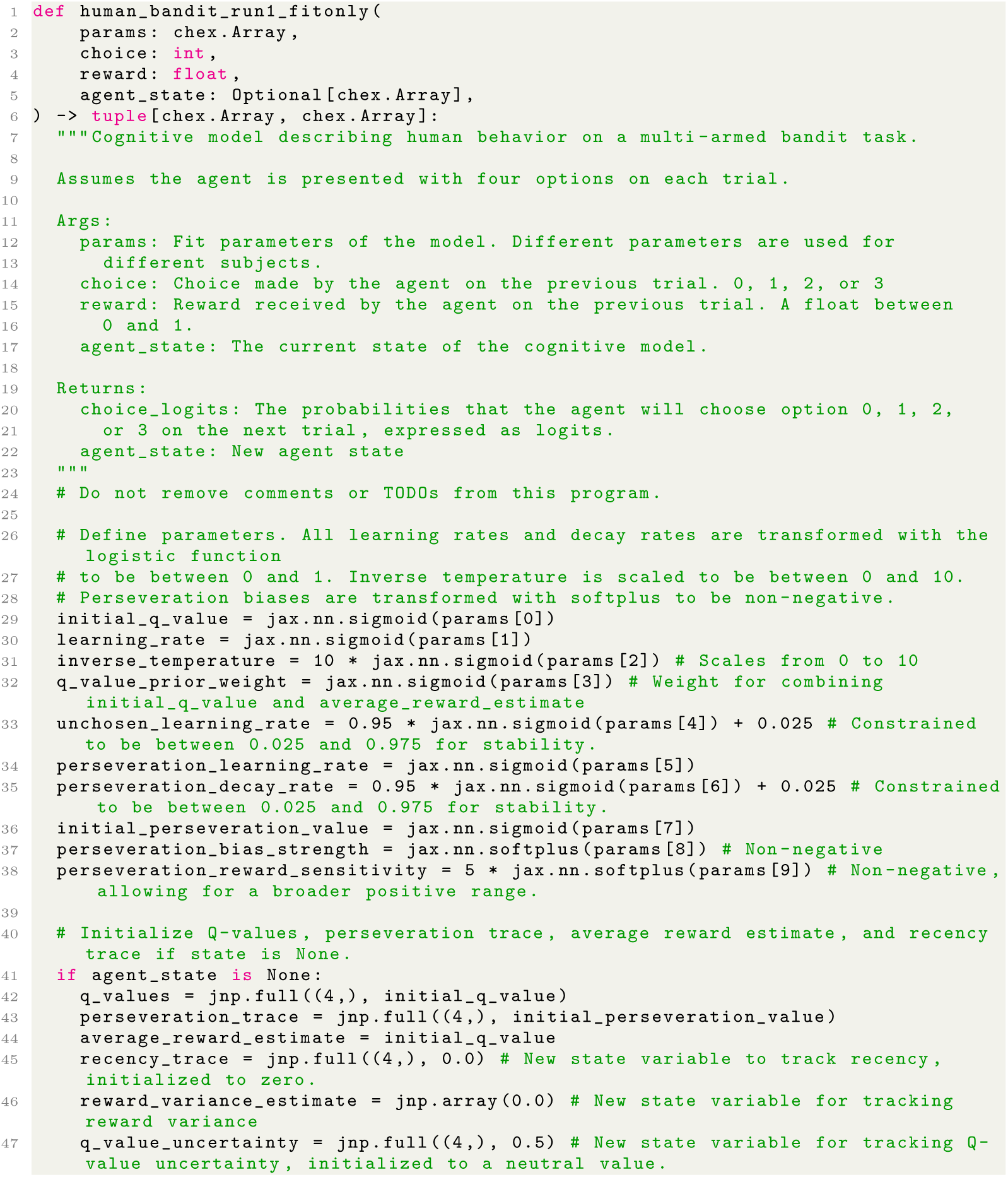
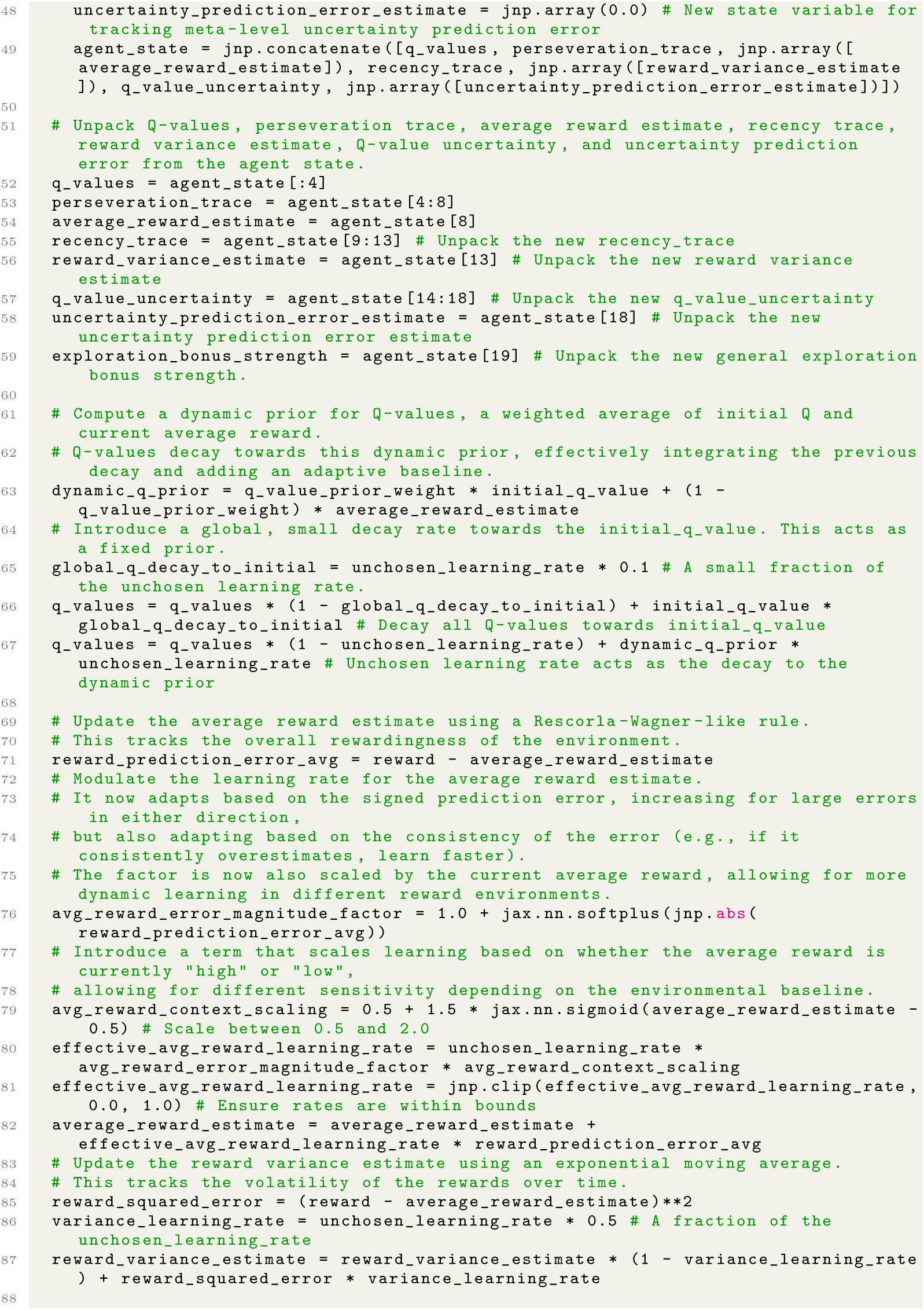
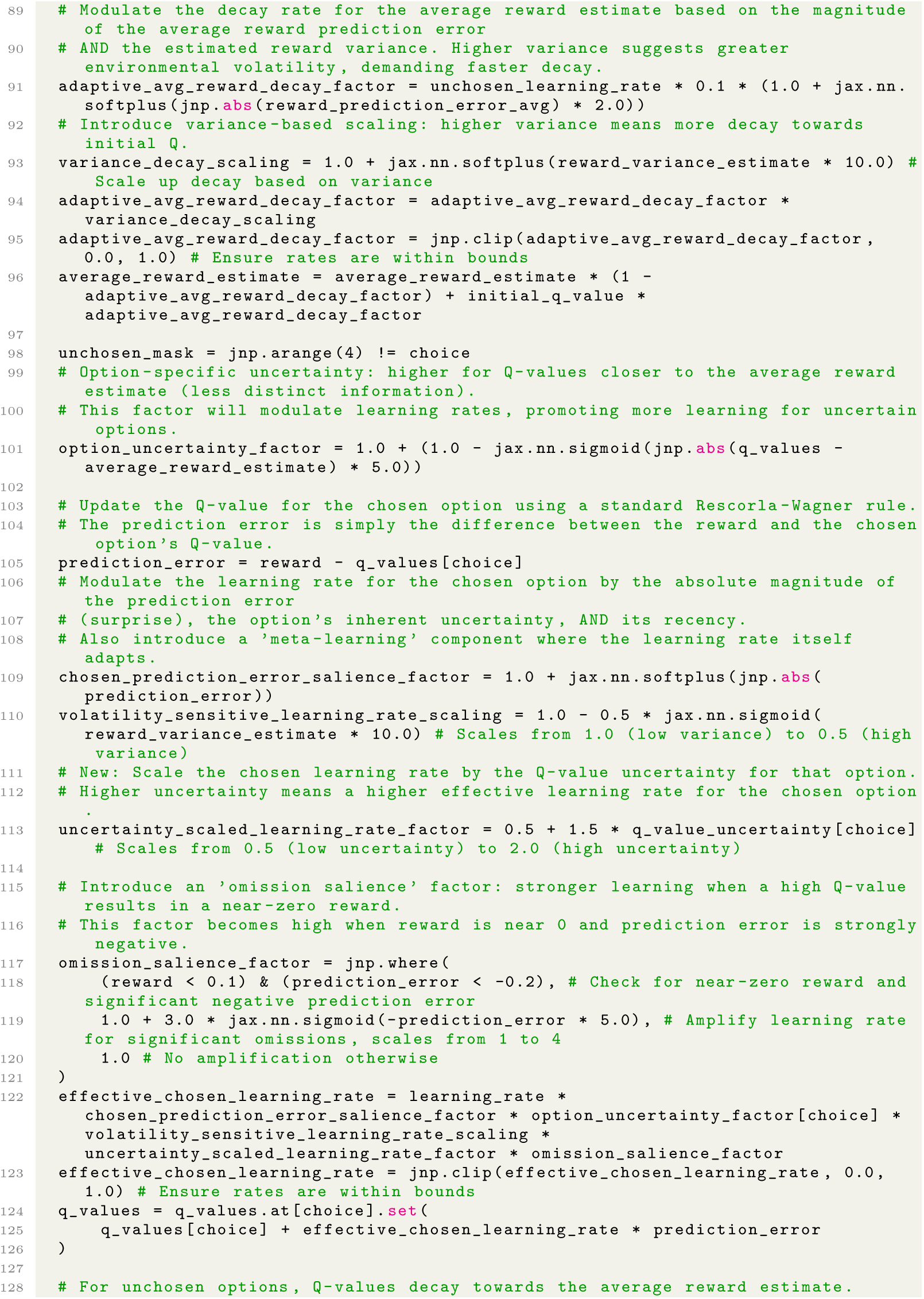
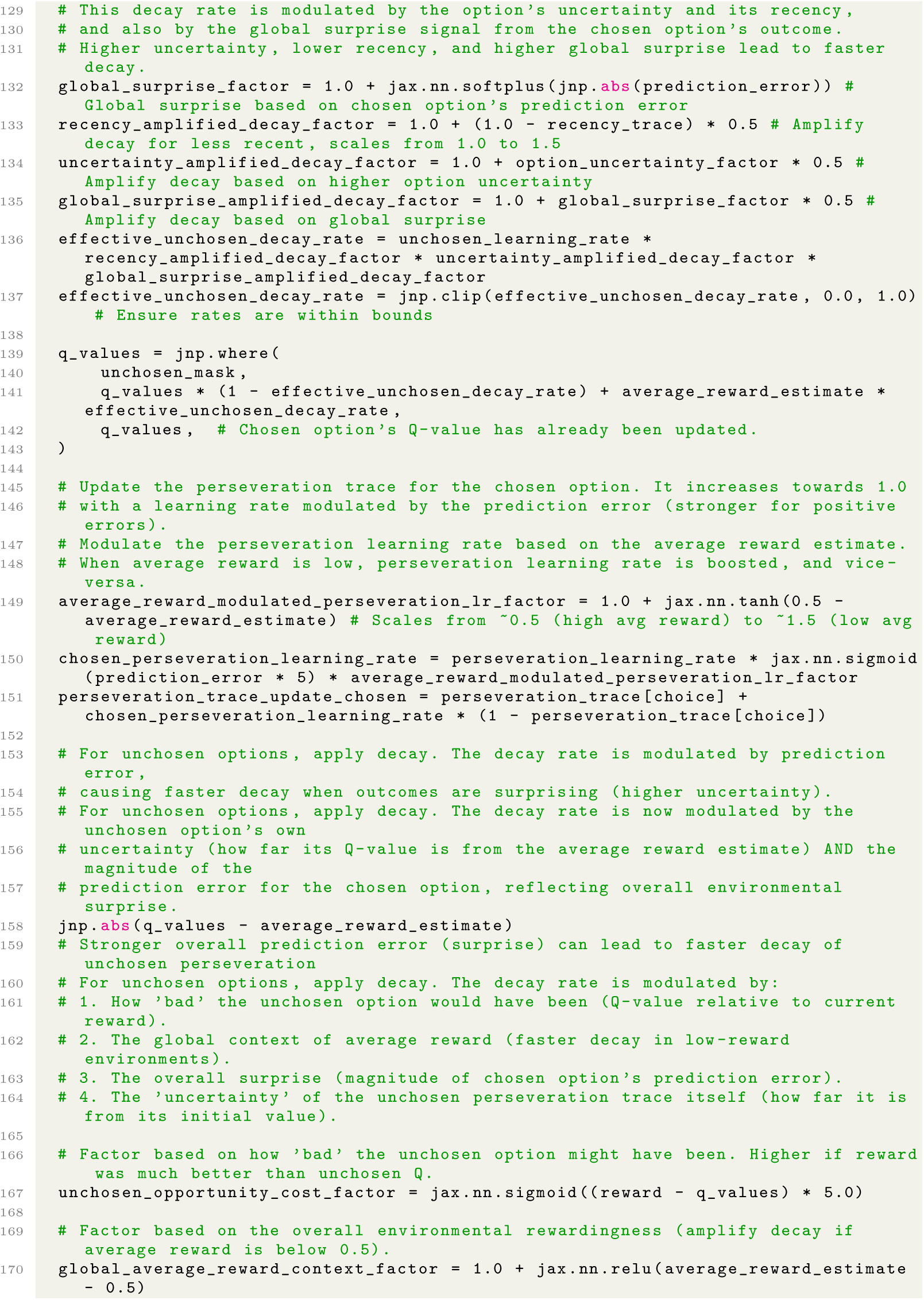
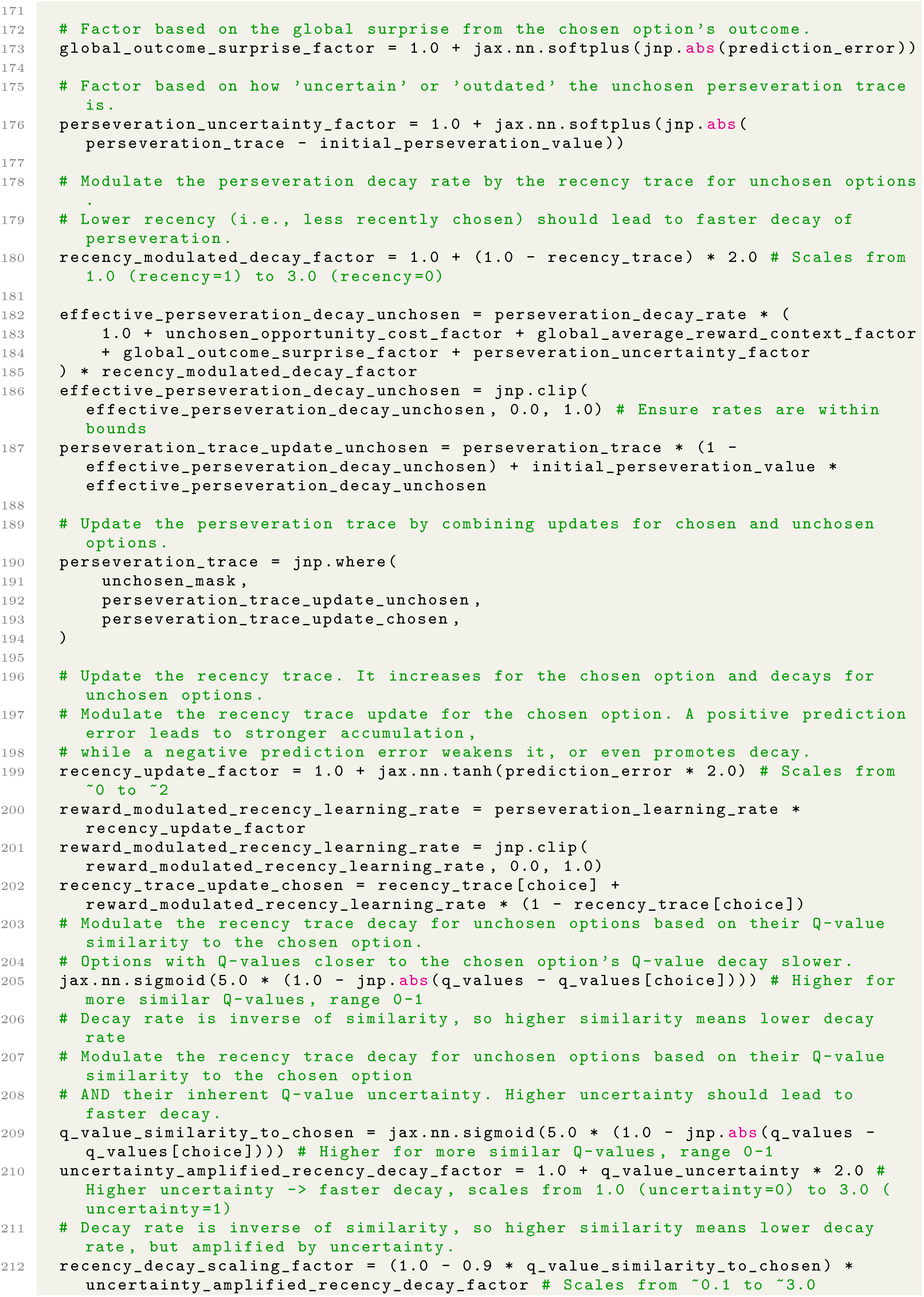
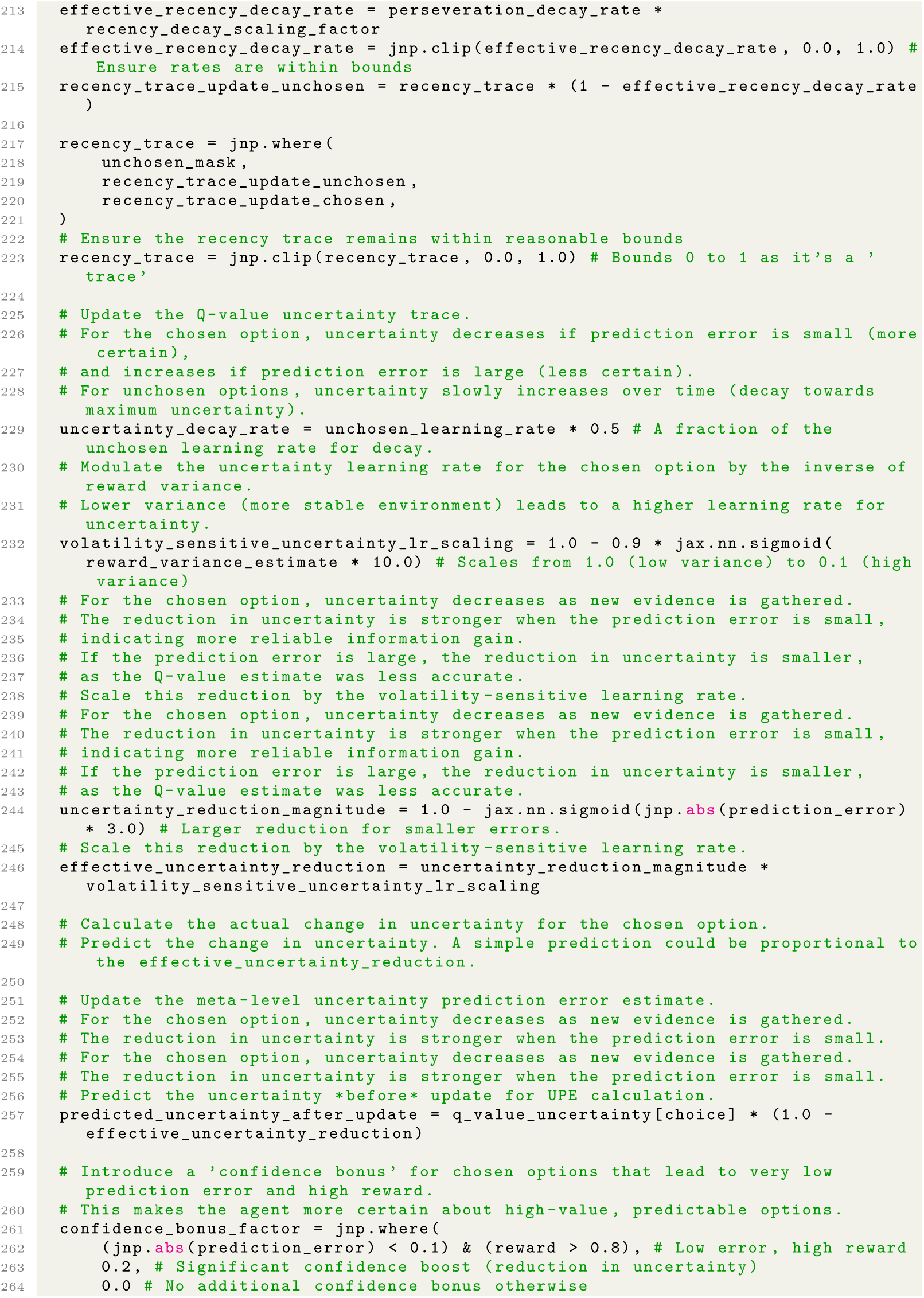
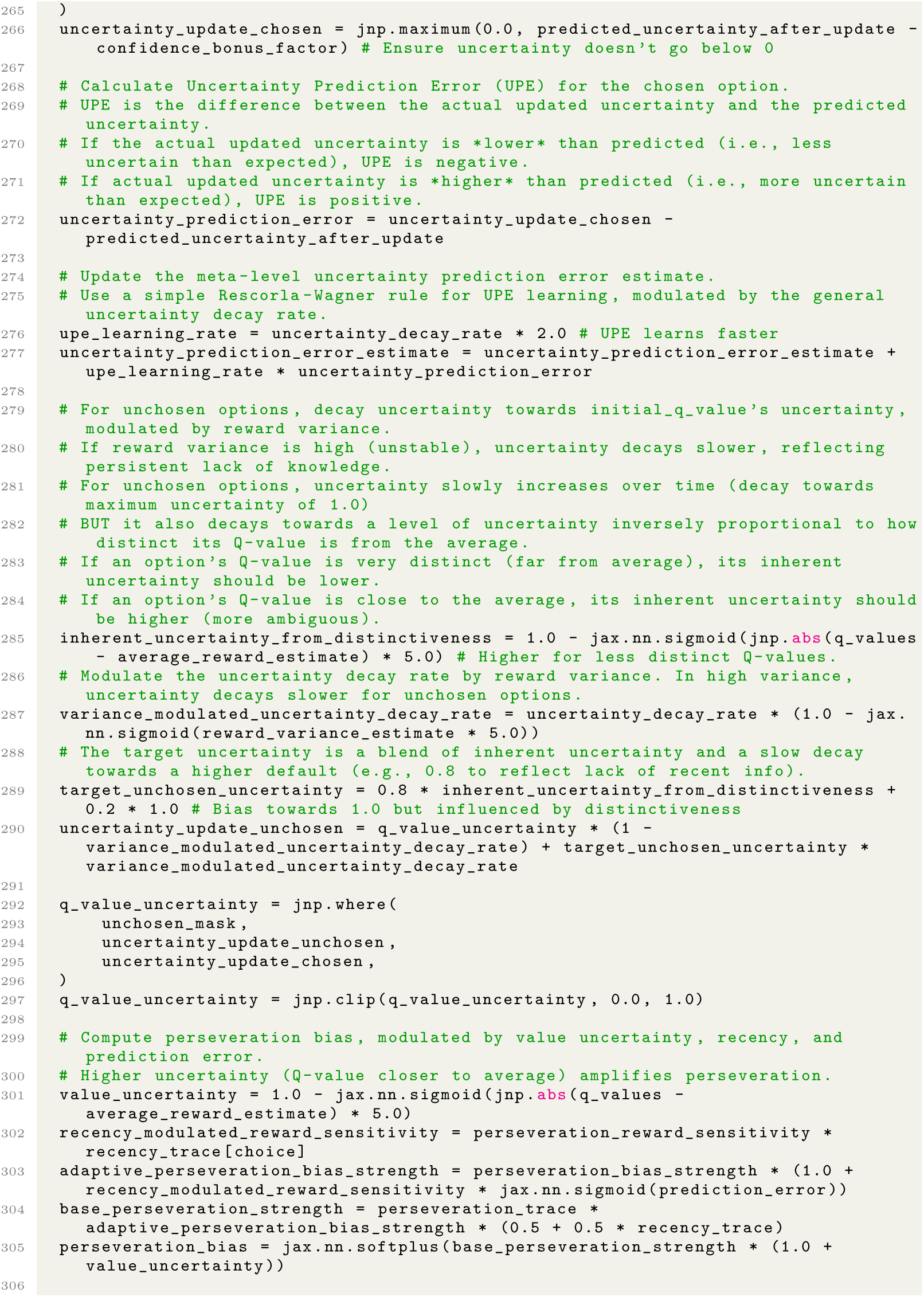
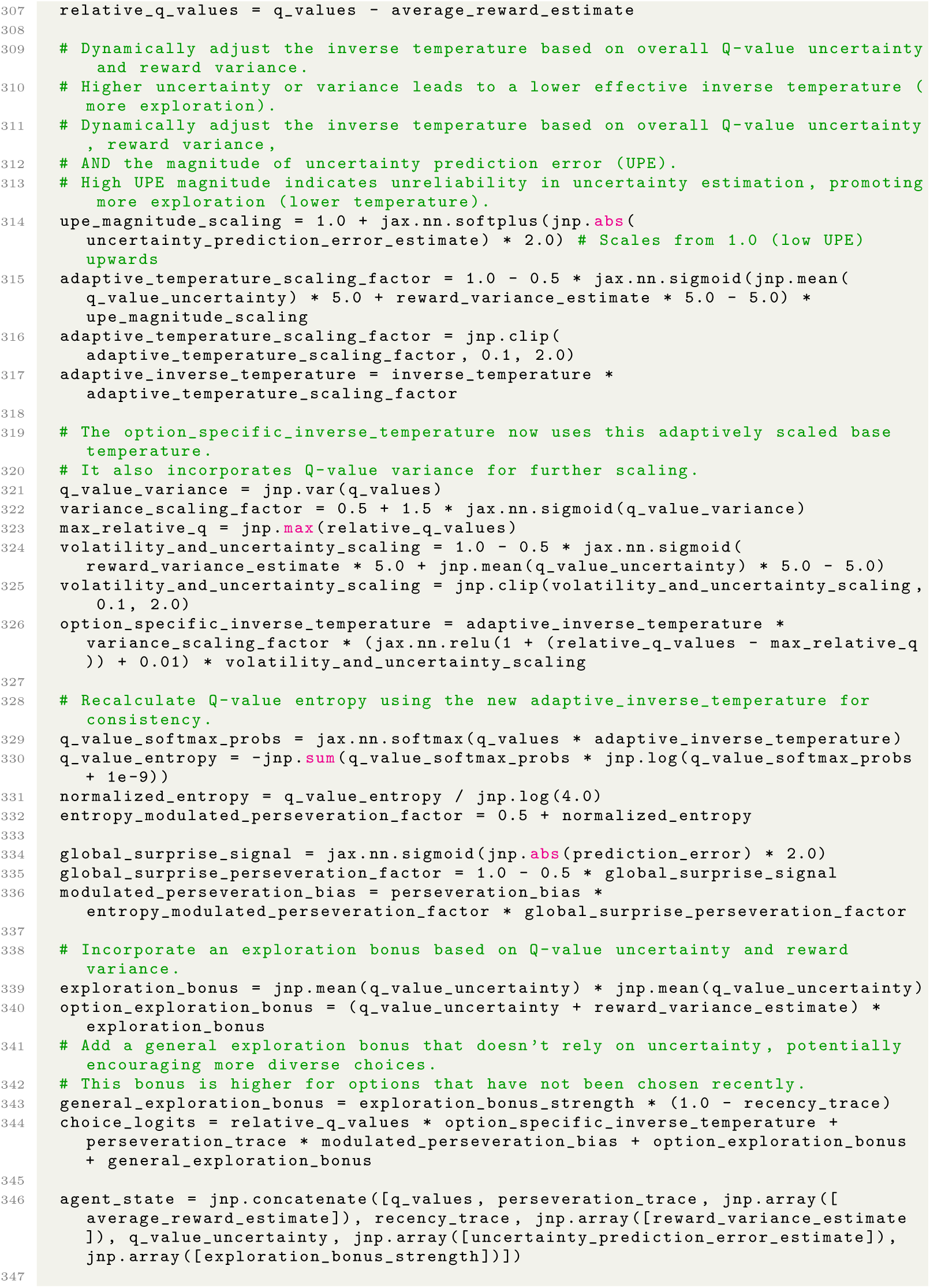
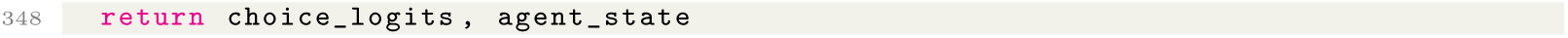

Code 2: Highest quality-of-fit program from the first independent Stage 1 AlphaEvolve run for the human bandit dataset.

#### A.1.7 Code: stage 2 (“Simplify”) programs

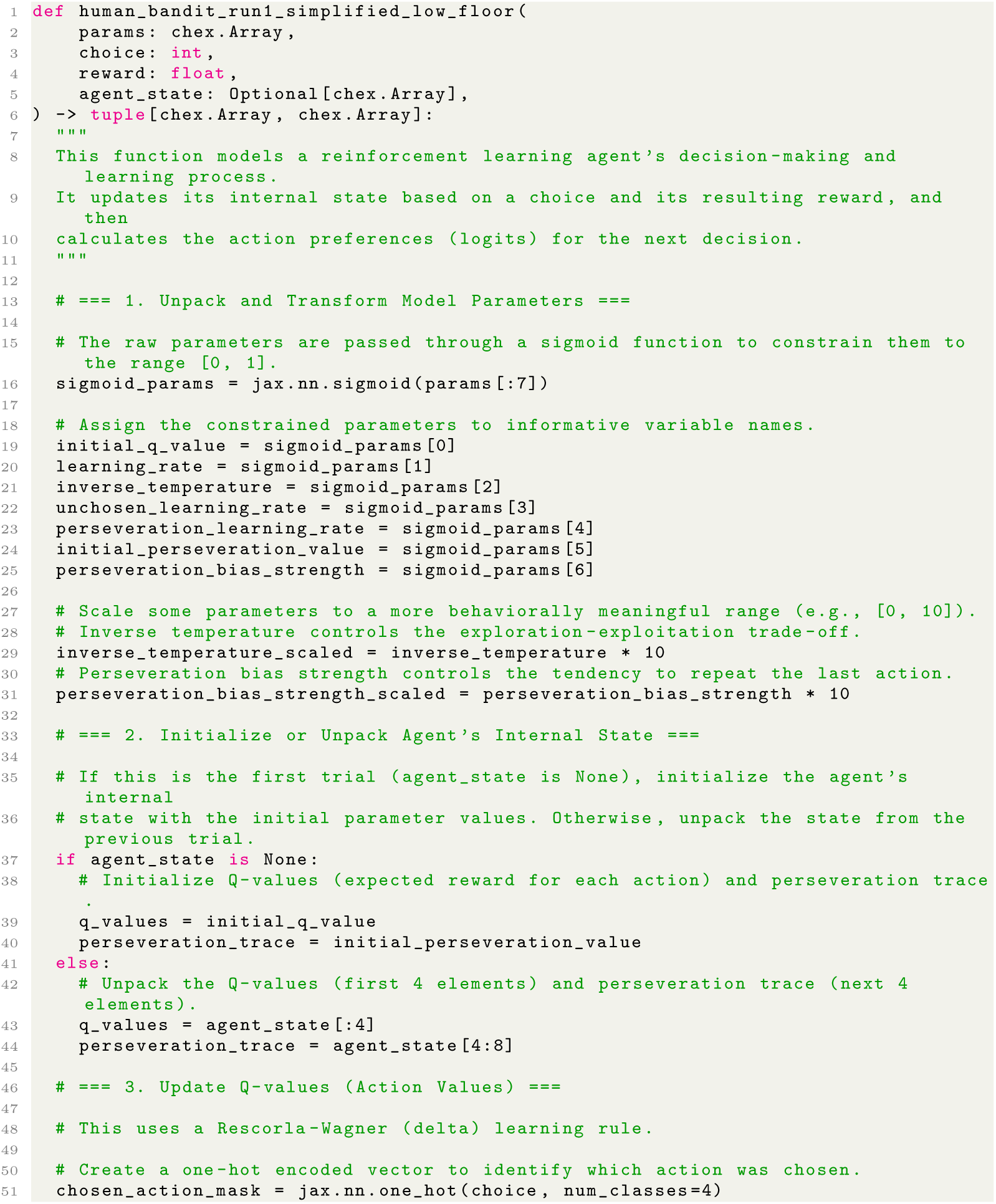
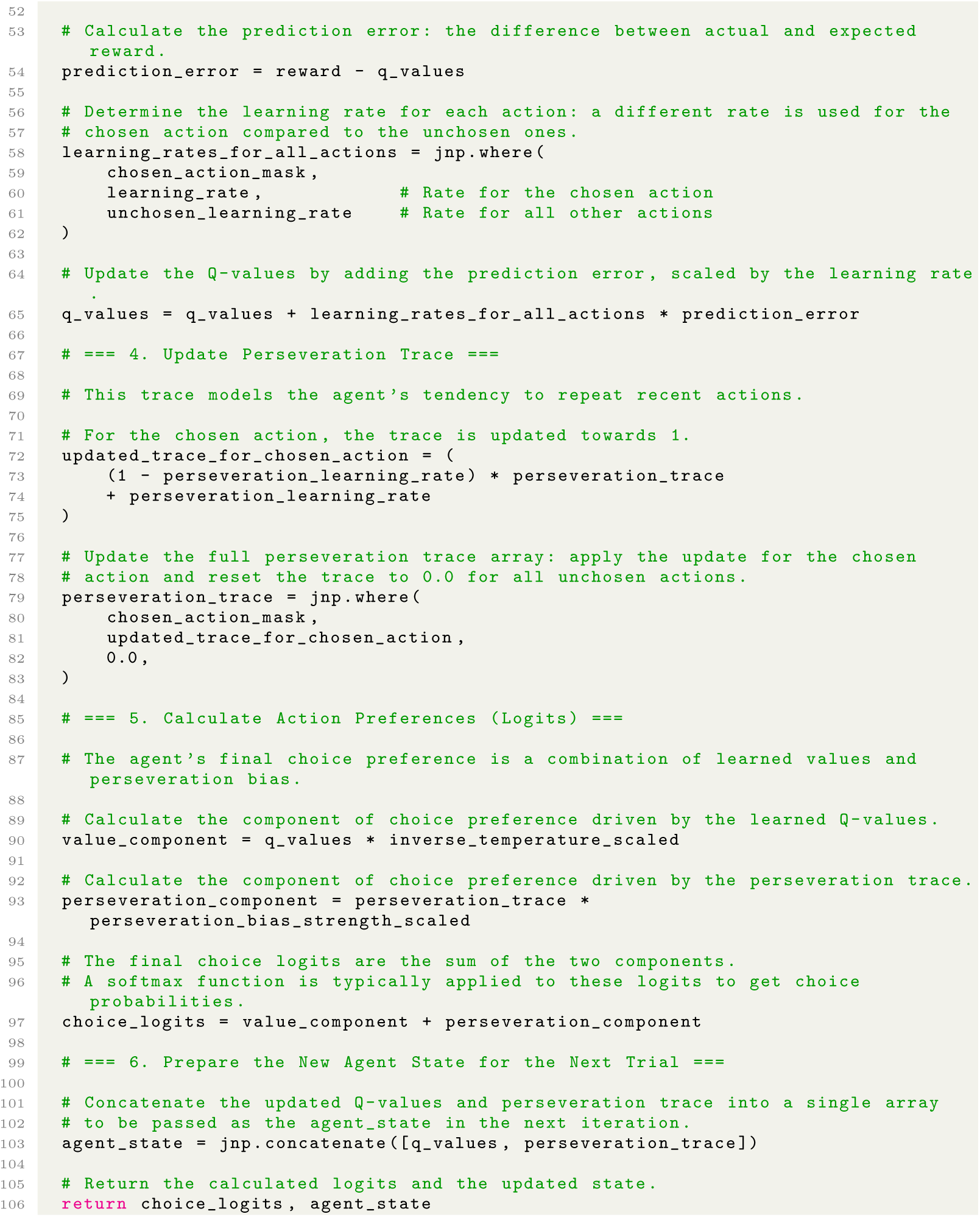

Code 3: Lowest-complexity program from Stage 2 AlphaEvolve run with 50% threshold for the human bandit dataset, evolved from programs in the first independent Stage 1 AlphaEvolve run and rewritten for readability (Stage 3).

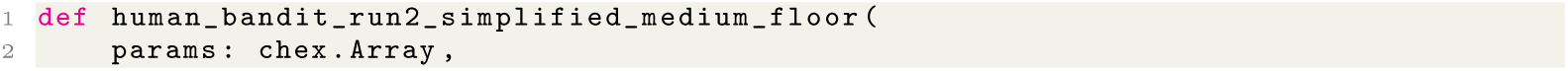
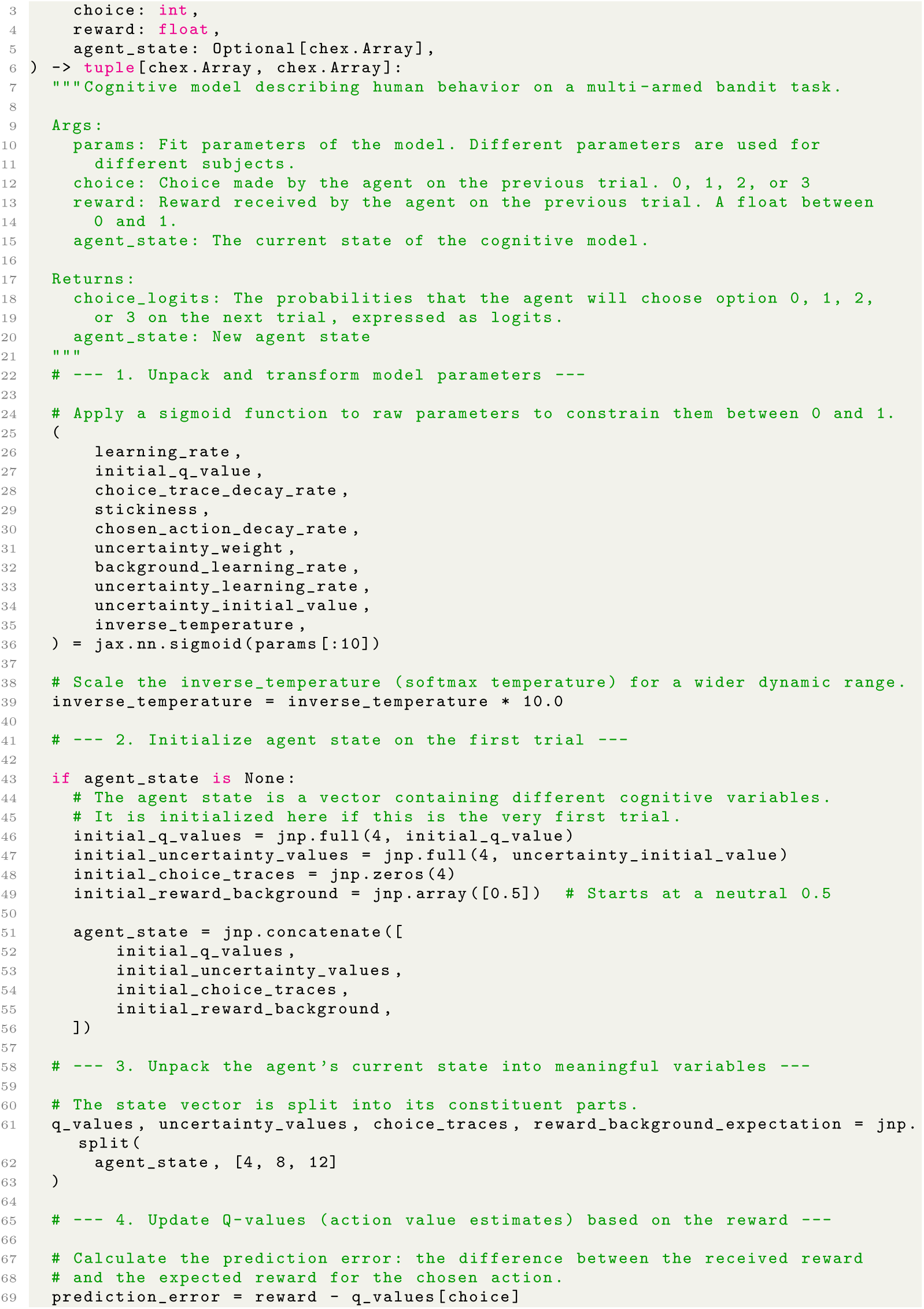
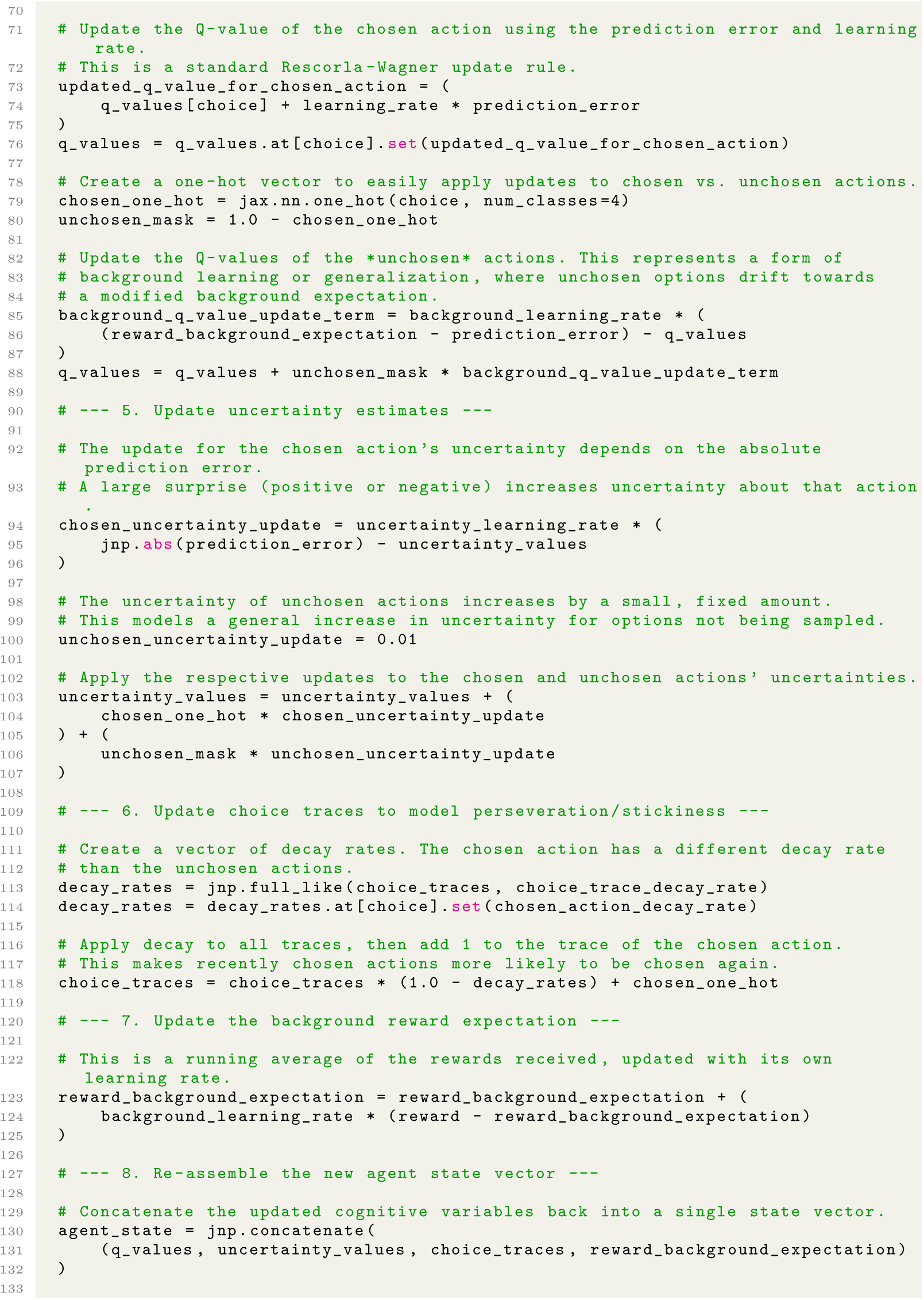
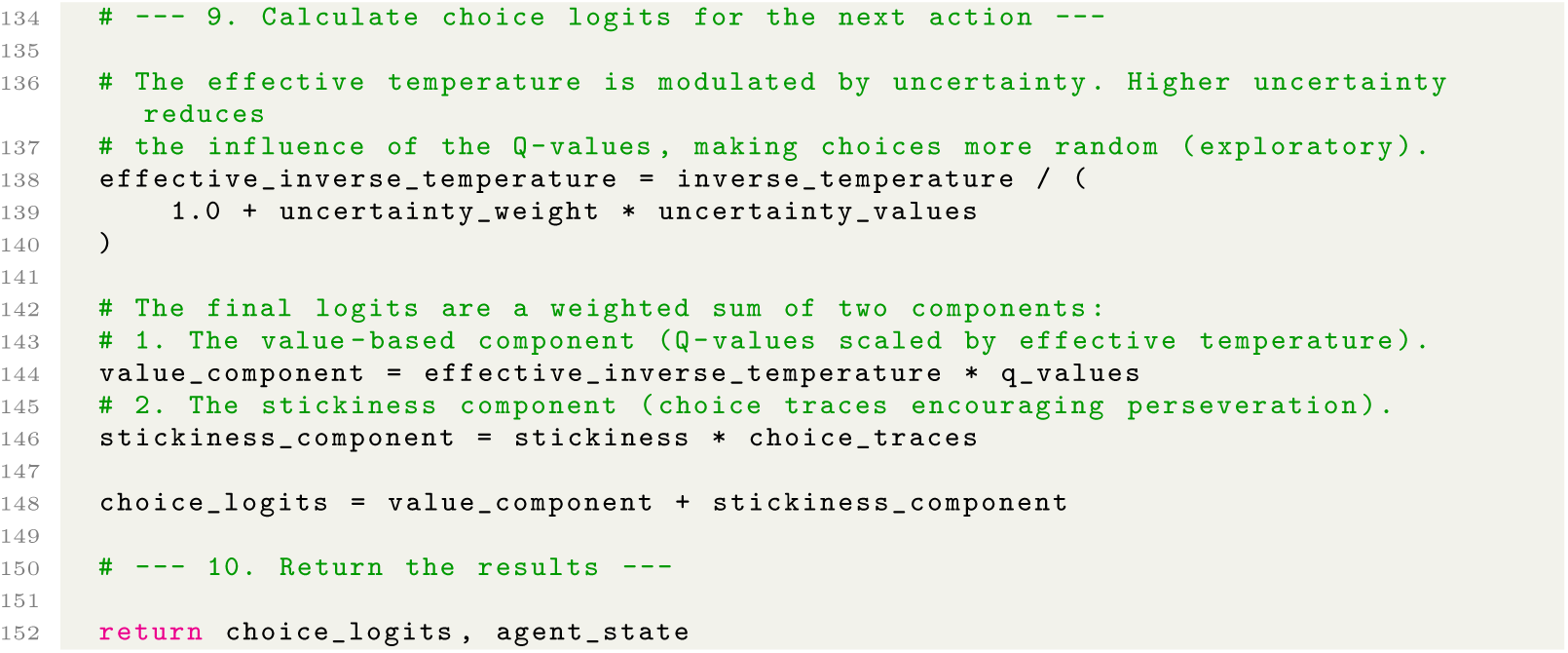

Code 4: Lowest-complexity program from Stage 2 AlphaEvolve run with 75% threshold for the human bandit dataset, evolved from programs in the second independent Stage 1 AlphaEvolve run and rewritten for readability (Stage 3).

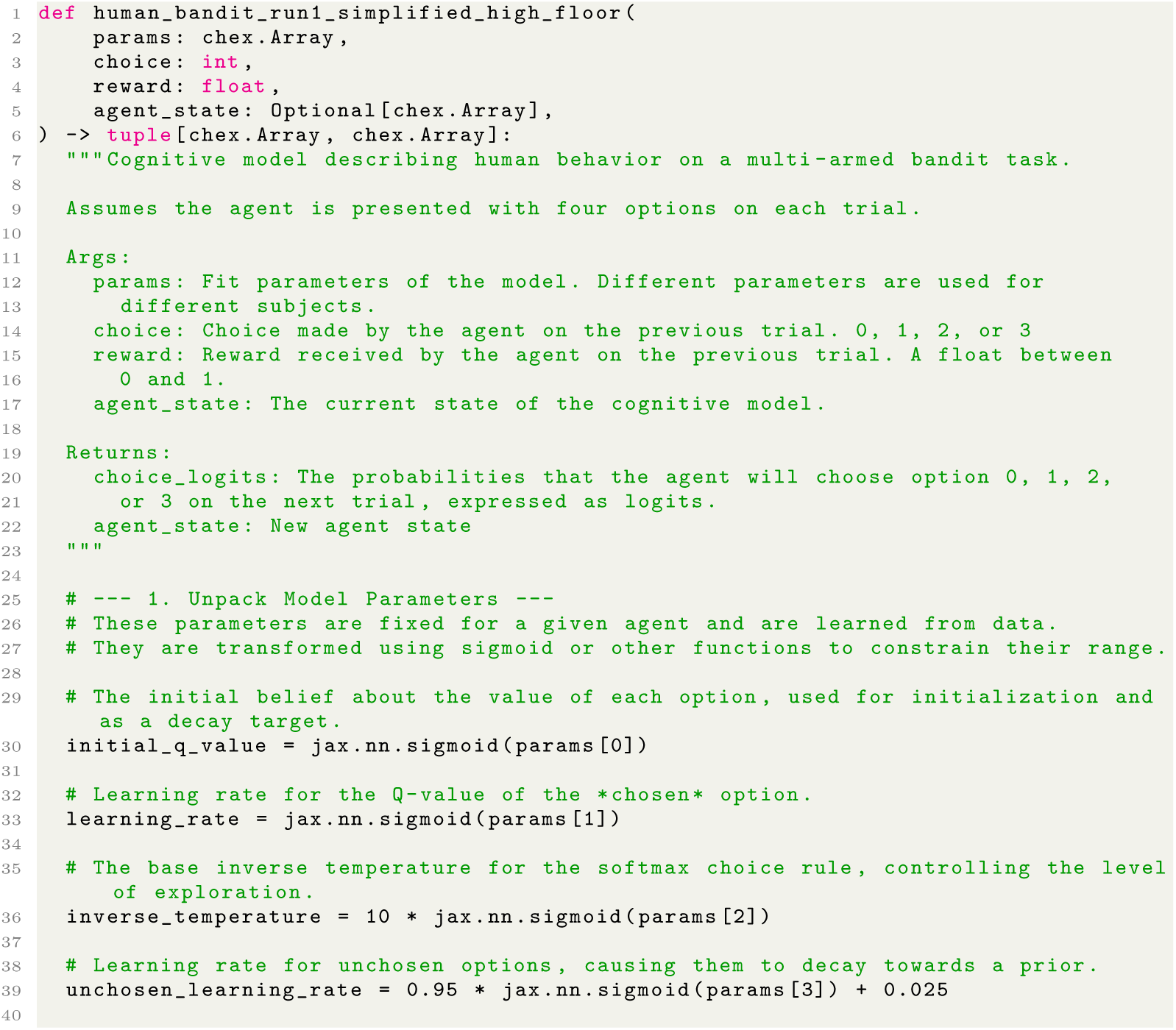
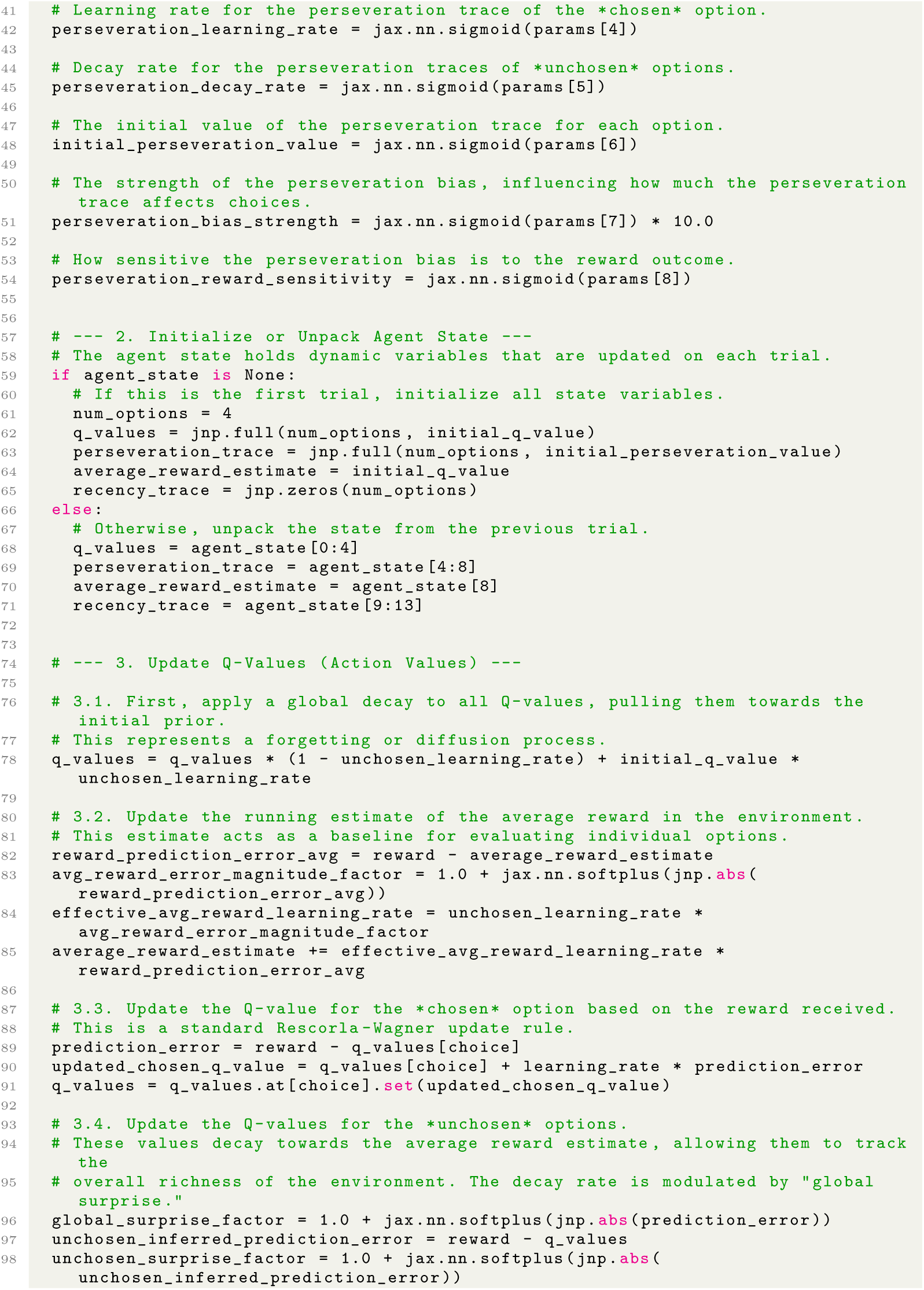
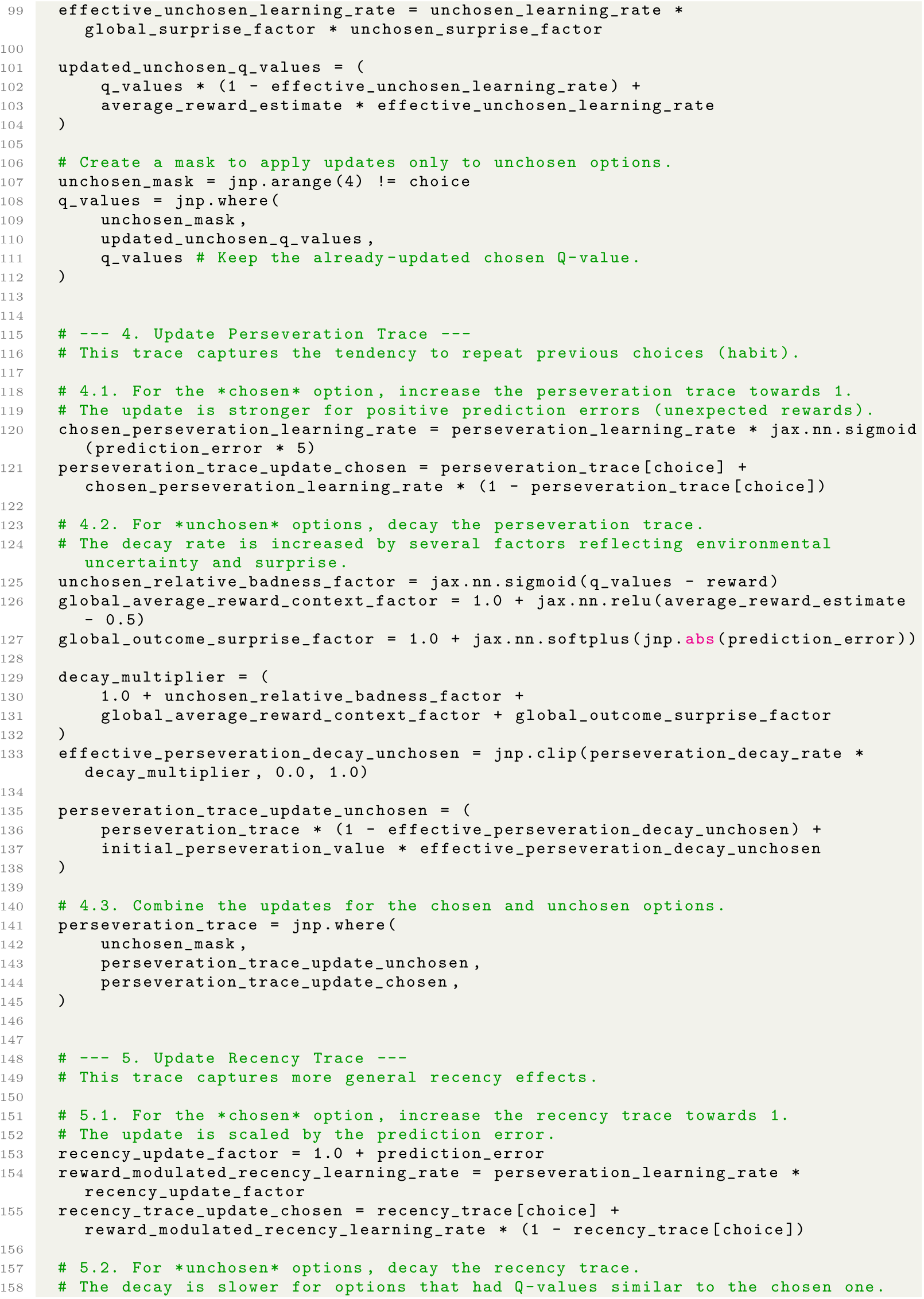
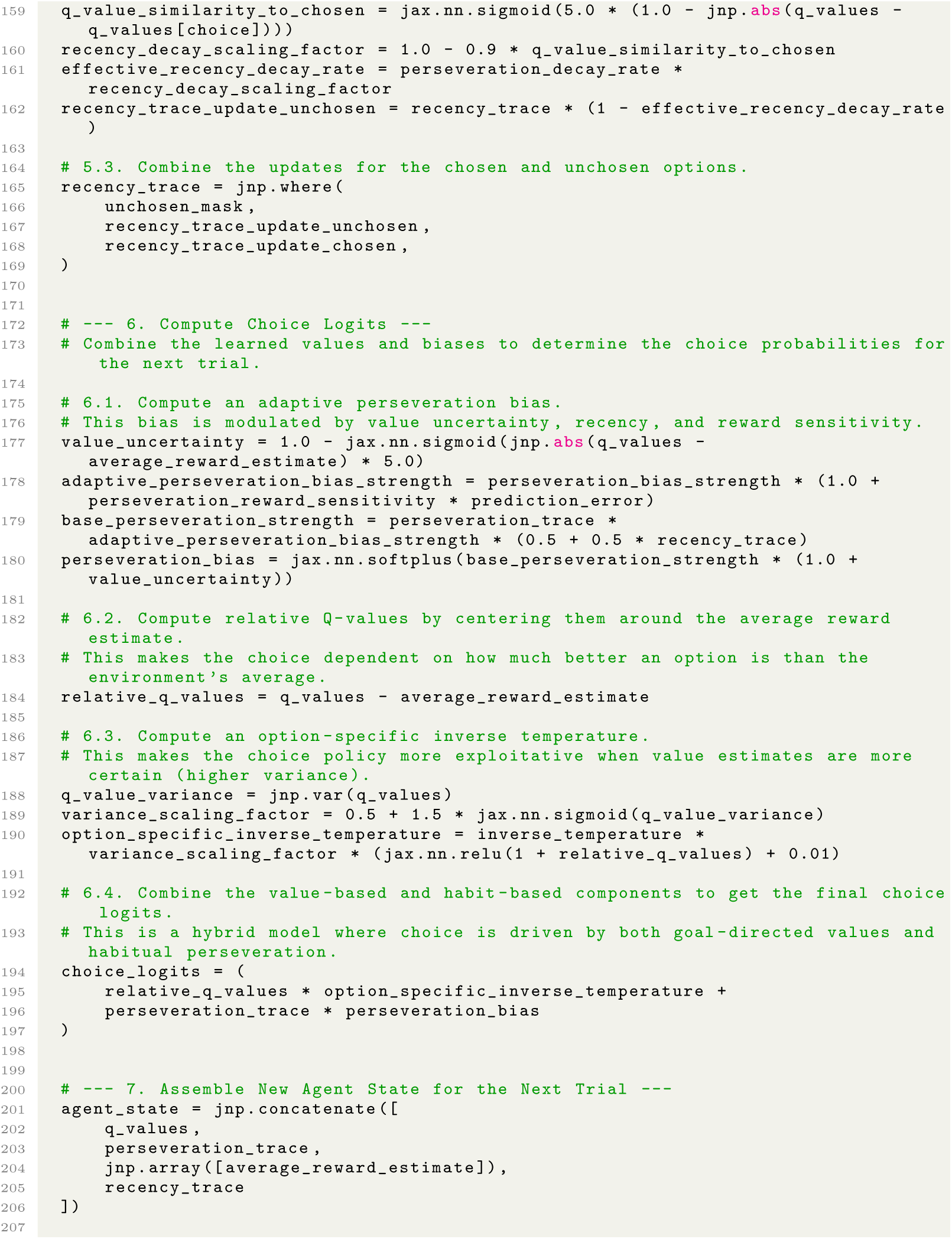
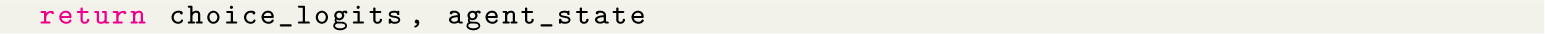

Code 5: Lowest-complexity program from Stage 2 AlphaEvolve run with 90% threshold for the human bandit dataset, evolved from programs in the first independent Stage 1 AlphaEvolve run and rewritten for readability (Stage 3).

#### A.1.8 Additional figures

**Fig. A1:**
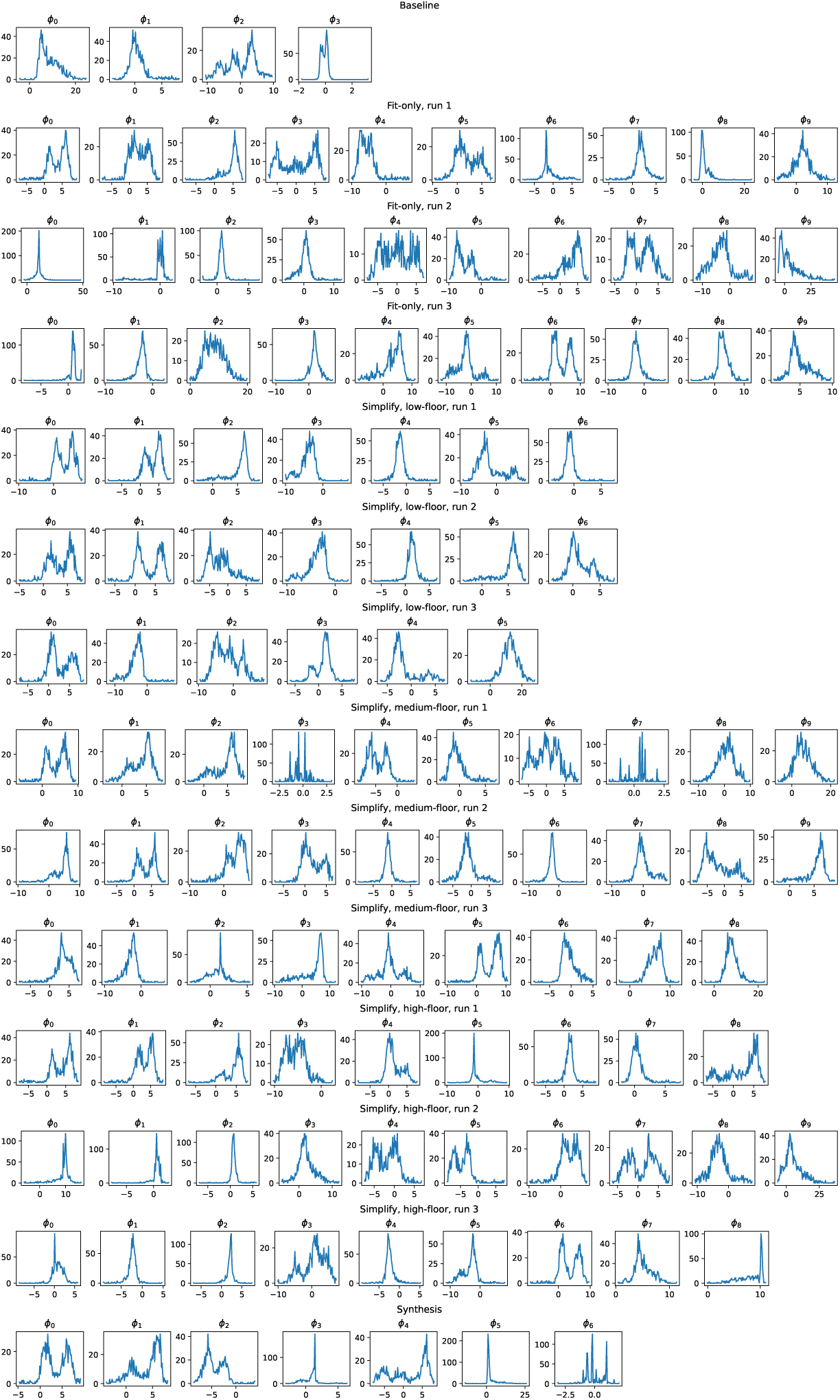
*Human Bandit* Dataset: Fit parameters for each program. The distribution of fit parameters for each fold of all discovered programs (fit-only and simplified), as well as the handcrafted baseline and synthesis program.

**Fig. A2:**
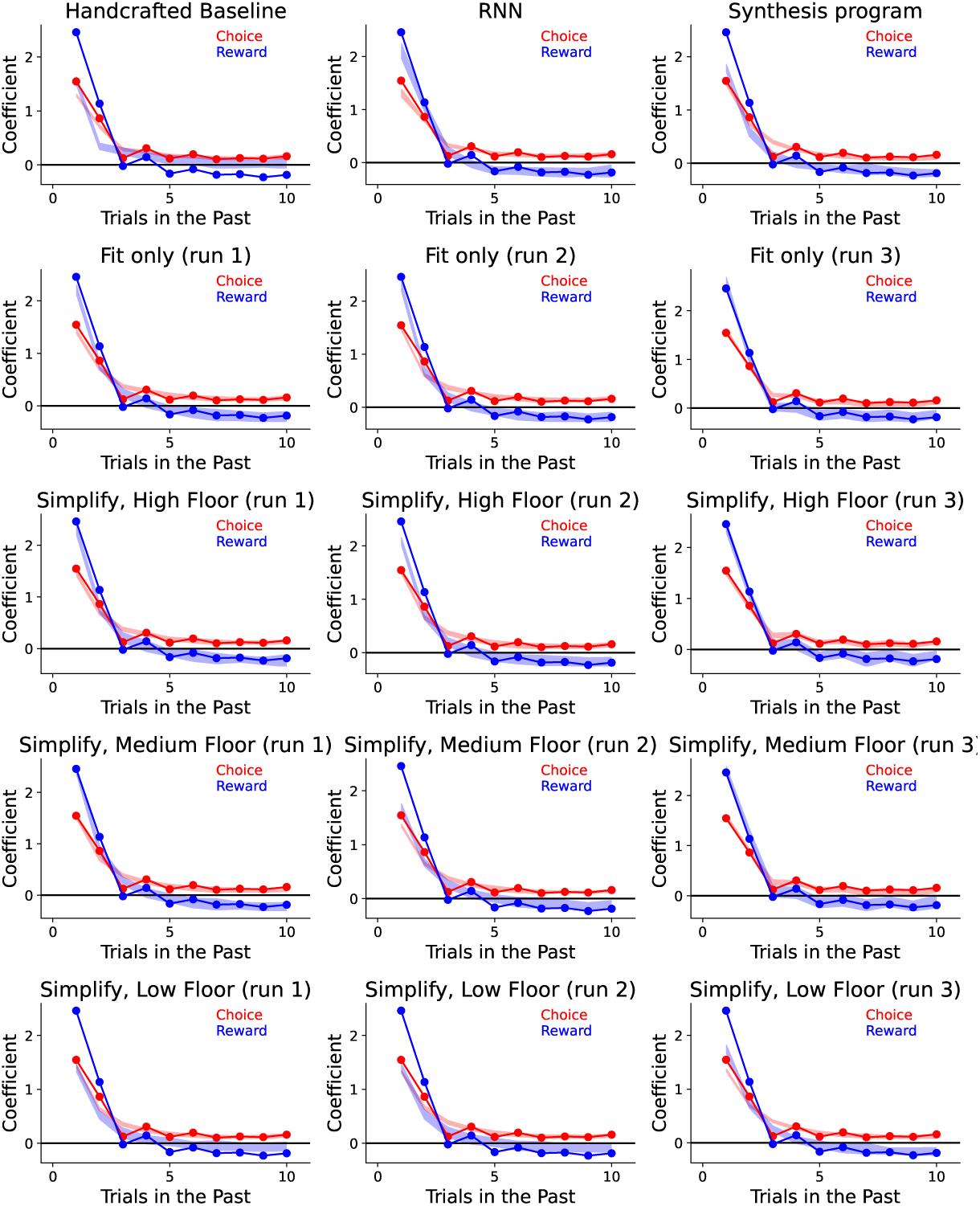
*Human Bandit* Dataset: Trial-lagged regression analyses. Here we see the trial-lagged regression analysis shown in Figure 6 for all discovered programs for this dataset. The coefficients for the real data are shown in solid lines, while the transparent patch shows the 95% prediction interval for the artificial data. We see that most discovered models capture the trial-lagged regression statistics better than the handcrafted model, and that programs obtained with higher floors tend to provide a tighter match. This demonstrates that the evolved programs are strong generative models.

**Fig. A3:**
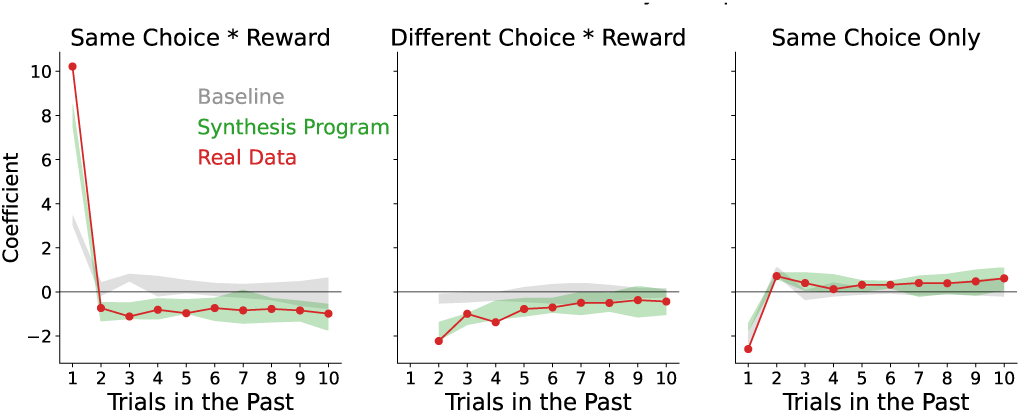
*Human Bandit* Dataset: Trial-lagged regression coefficients for tendency to repeat previous choice. Trial-lagged regression analysis showing the influence (coefficient, positive promotes repeating) of previous trials (1-10 trials ago) on tendency to repeat the agent’s current choice. The The leftmost plot (“Same Choice * Reward”) shows the influence of being rewarded in the past for choosing the same action the subject is considering repeating; the middle plot (“Different Choice * Reward”) shows the influence of being rewarded in the past for choosing a *different* action from the one the subject is considering repeating; the rightmost plot shows the influence of having performed the same action in the past, regardless of reward.

**Fig. A4:**
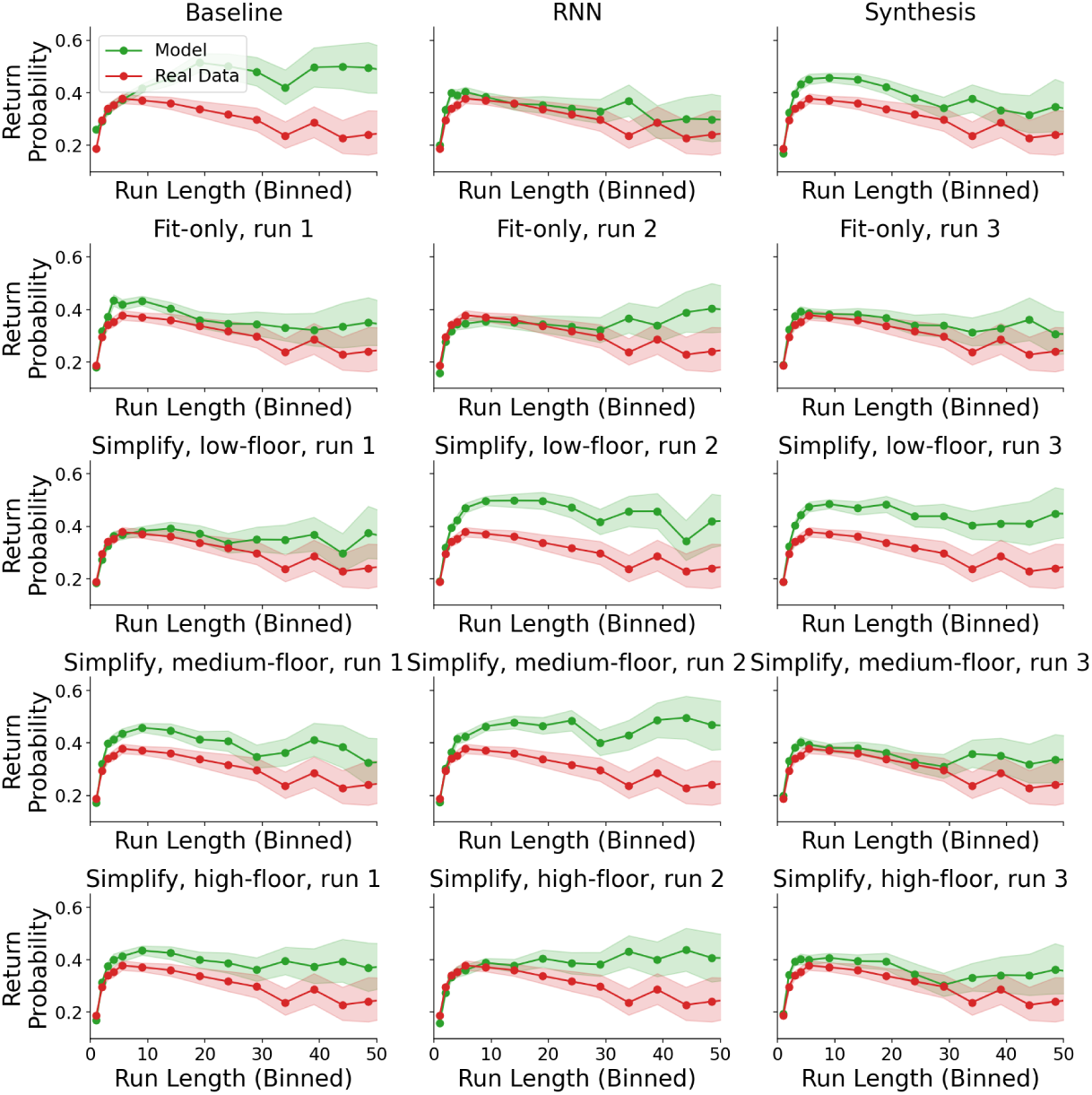
*Human Bandit* Dataset: Run Return Probability analysis for all models. Here we see the probability of returning to a run of repeated choices, for runs of different length, after a single different choice. Shaded intervals depict 95% confidence intervals.

### A.2 Rat Bandit

#### A.2.1 Dataset

We consider the behavioral dataset from Miller et al. [8], in which rats perform a two-alternative reward-learning task with binary rewards. Rats indicated their choice on each trial by entering one of two available nose ports, which were equipped to deliver small liquid rewards. Reward probabilities followed independent bounded random walks. Rats performed daily sessions of approximately one hour. The dataset contains choices from 20 rats performing 1,946 total sessions and 1,087,140 total trials.

We obtained this dataset from the following URL, where it is freely available under a permissive open-source license: https://figshare.com/articles/dataset/From predictive models to cognitive models Separable behavioral processes underlying reward learning in the rat/ 20449356

#### A.2.2 Baseline Model

Miller et al. [8] performed an intensive human process of data-driven model discovery on this dataset. This resulted in a model that we refer to as “Reward-Seeking/Habit/Gambler-Fallacy” (RHG), which we adopt as the handcrafted baseline model for this dataset. For compatibility with our pipeline, we re-implemented this model in jax.

#### A.2.3 Discussion of evolved programs

One fit-only seed achieved a much higher performance on the training set than the others. Because the floors for simplifying were based on the difference between this best fit-only program and the baseline program, this meant that the highest floor was actually higher than the fit-only performance on the training set for the other two seeds. As a result, there was only one high-floor simplify program for this dataset.

We therefore focused our analysis on the 6 medium- and low-floor programs, of which we had three each. Medium-floor programs had similar scores to one another on the held-out subjects, and similar scores to the fit-only programs from the two lower-scoring seeds. This indicates that the simplifications done did not substantially impact held-out performance. Low-floor programs had substantially worse scores, indicating that the simplifications done to get to this floor did substantially impact held-out performance.

##### Cognitive variables

All simplified programs defined two sets of variables for fast and slow reward-dependent learning processes respectively. Some programs (3/6 low- and medium-floor programs) also defined action perservation terms that encouraged actions to be repeated independent of whether they had been rewarding. This particular decomposition of cognitive variables was present in the handcrafted baseline model as well, although the particular discovered updates on the terms had slightly different forms and added nonlinearities.

##### Fast reward-guided learning

The fast learning modules often implemented a “linked” Q-learning variant in which the prediction error *δ* = *r*−*Q*(*c*) given the chosen action value was used to update both the chosen and the unchosen Q-values (3/6 programs). The chosen and unchosen action values were updated with different learning rates but otherwise antisymmetrically, with updates *Q*(*c*) ← *Q*(*c*) + *ϕ_i_δ* and *Q*(*c*^′^) ← *Q*(*c*^′^) − *ϕ_j_δ*. We call this linked because the update on the chosen and unchosen action values are linked. Other discovered programs found more complex reward-guided learning.

##### Slow learning of non-rewarding options

The slow learning modules often exhibited a pattern wherein it accumulated recent non-rewarding choices and ignored rewarding choices. This had the effect of driving choices *toward* choices from which rewards had been absent. This pattern was evident in the baseline RHG model as well, and was referred to as the “Gambler’s Fallacy” term. Gambler’s Fallacy refers to an often mistaken belief that options that have not returned rewards recently are due to payoff soon.

##### Mapping cognitive variables onto behavior

One discovered model exhibited the nonlinear pattern of behavior present in the synthesis program. However, this nonlinearity contributed substantially to behavior performance: across the medium-floor models, it is the simplest while maintaining the median quality-of-fit. For each slow learning value *x* and fast learning values *y*, the particular nonlinearity is *ye^x^*.

Programs also exhibited a “win-stay/lose-shift” contribution to behavior. This does not reflect “learning” *per se*, as it does not necessitate updating cognitive variables; rather, a bonus is applied to the previous action that depends on the reward.

#### A.2.4 Discussion of synthesis program

The synthesis program kept the linked Q-learning rule that appeared in 3/6 programs, as this was the most common learning rule to emerge and fit as well as the other more complex rules. The term tracking the slow learning of non-rewarding options was also included, as was the nonlinear rule for combining the fast and slow learning terms and the one-back win-stay/lose-switch bias, as these all contributed positively to behavior. Its slightly improved simplicity over the other medium-floor programs comes from the removal of terms that were revealed to not contribute to score.

#### A.2.5 Code: stage 2 (“Simplify”) programs

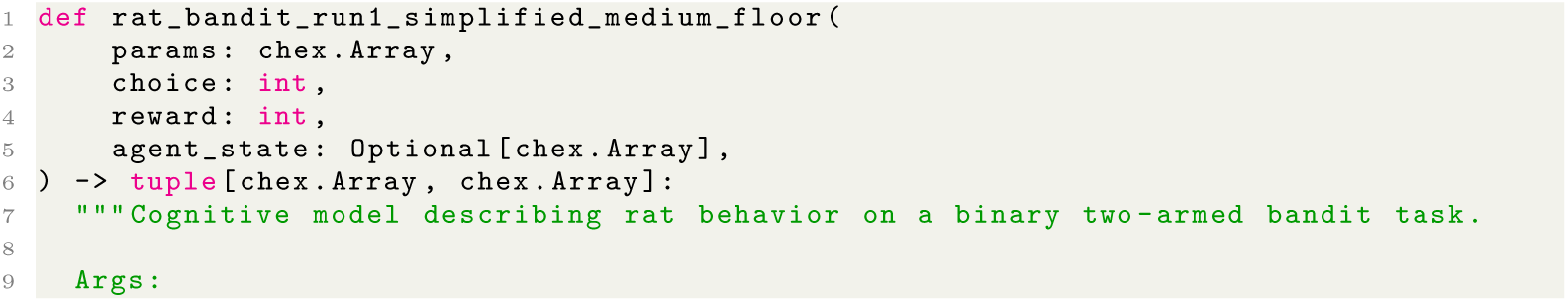
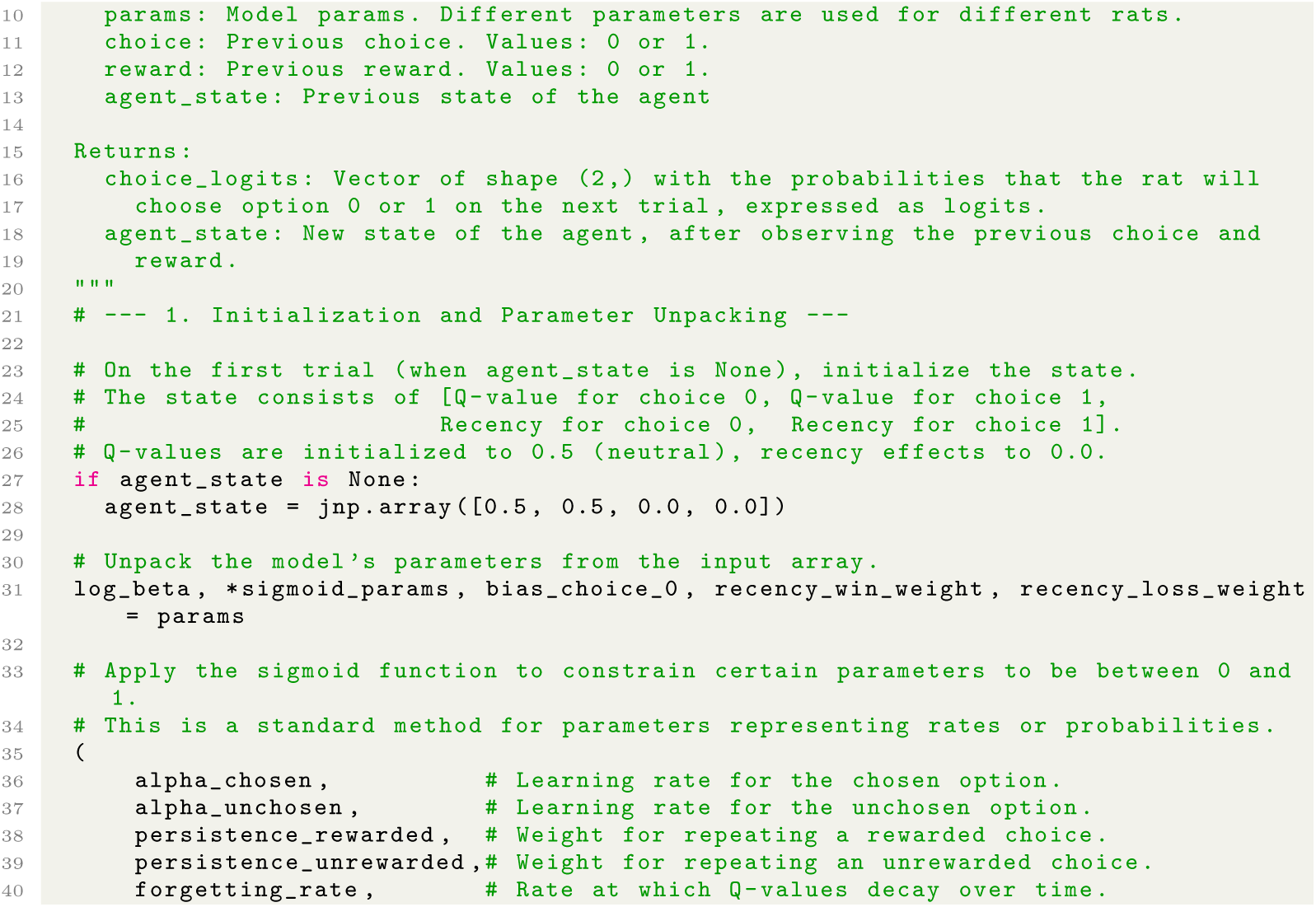
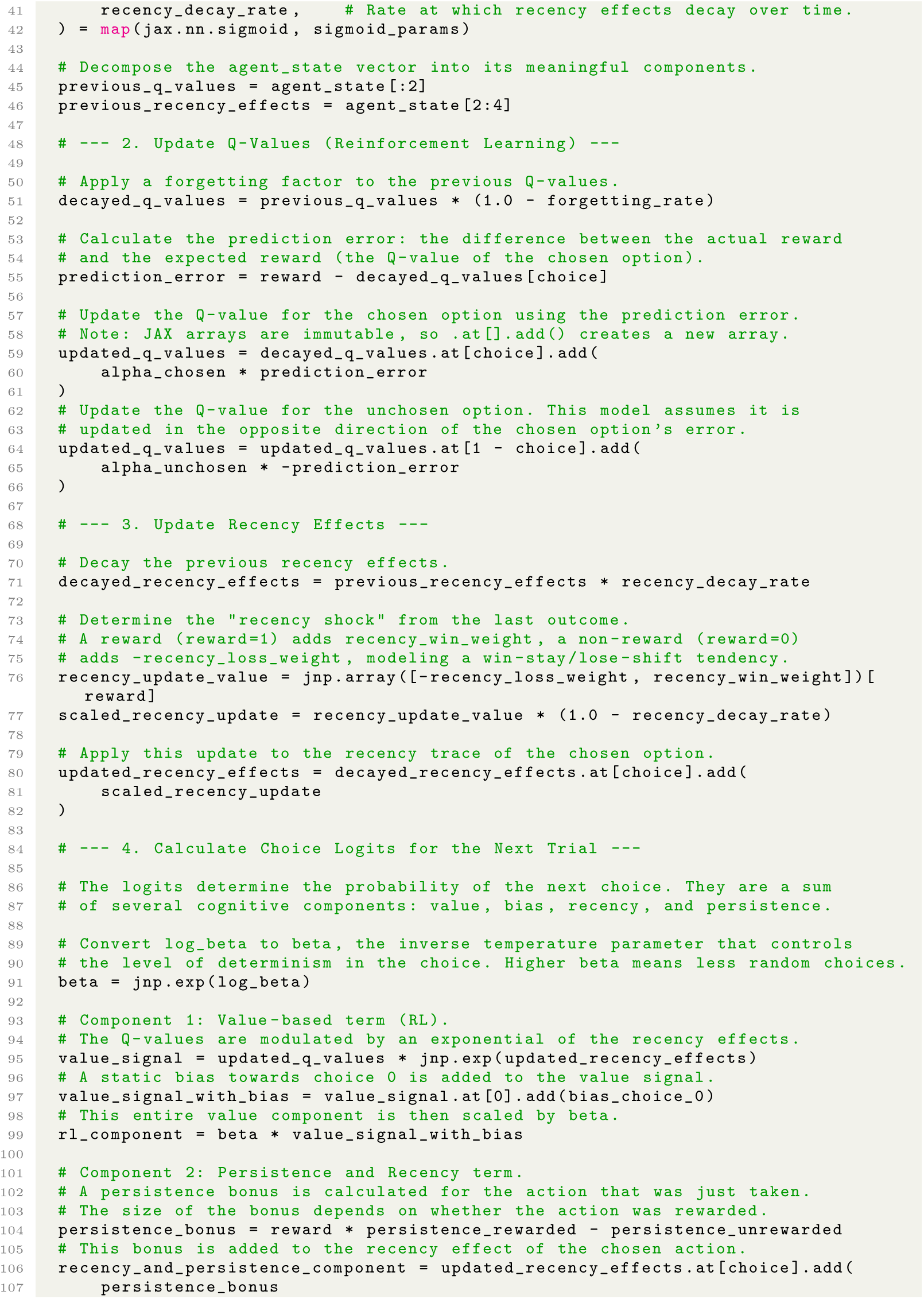
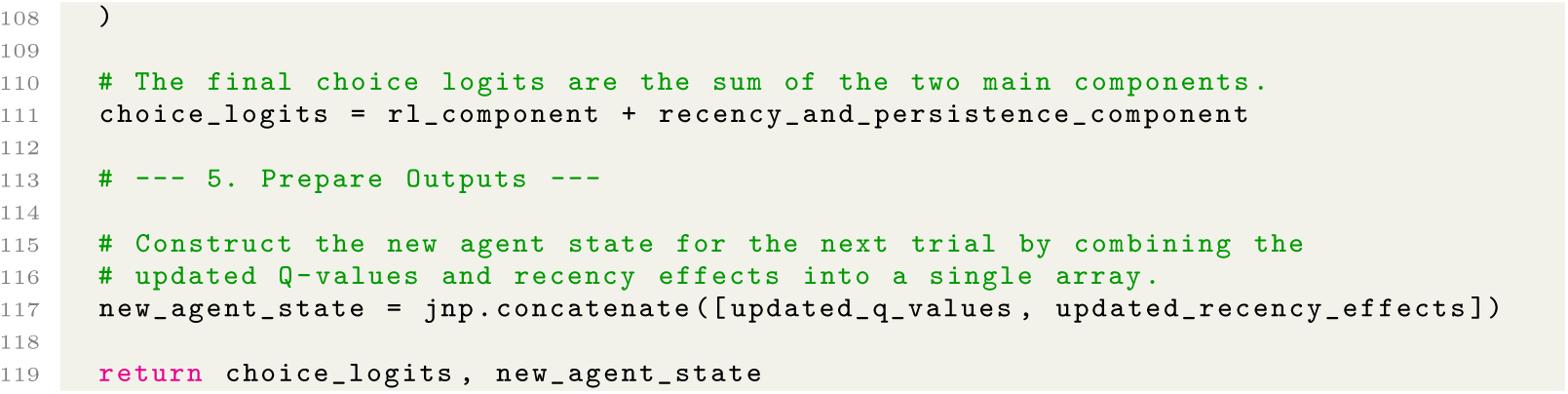

**Table A2:**
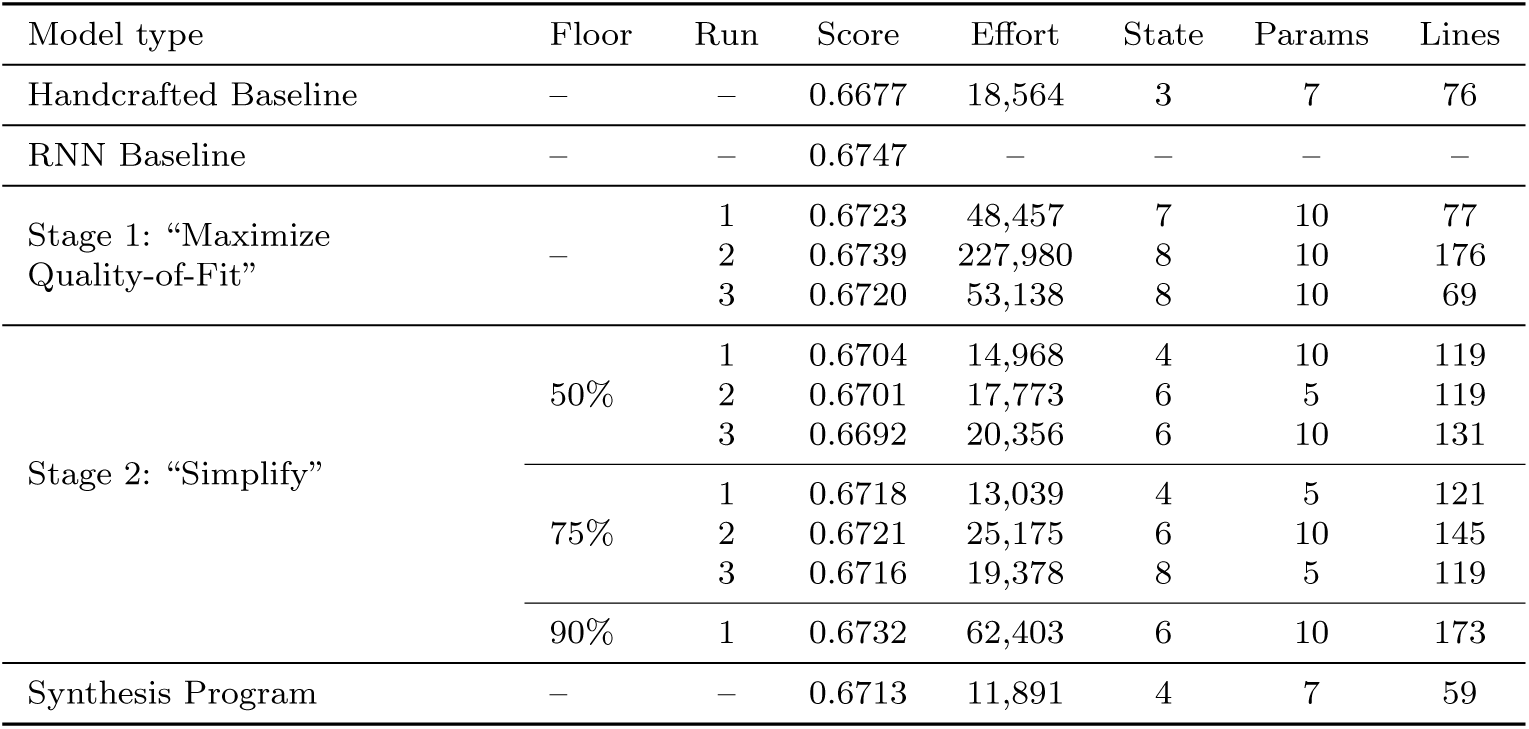
Evaluation performance and program complexity for models in the *Rat Bandit* dataset. For programs generated by the “Simplify” stage, Floor represents the quality-of-fit threshold below which programs are discarded (see Section 7.6.2). Score indicates the average normalized likelihood across evaluation subjects (see Section 7.3); Effort is Halstead effort. State, Params, and Lines indicate the number of state variables, per-subject parameters, and lines of code respectively.

Code 6: Lowest-complexity program from Stage 2 AlphaEvolve run with 75% threshold for the rat bandit dataset, evolved from programs in the first independent Stage 1 AlphaEvolve run and rewritten for readability (Stage 3).

#### A.2.6 Code: synthesis program

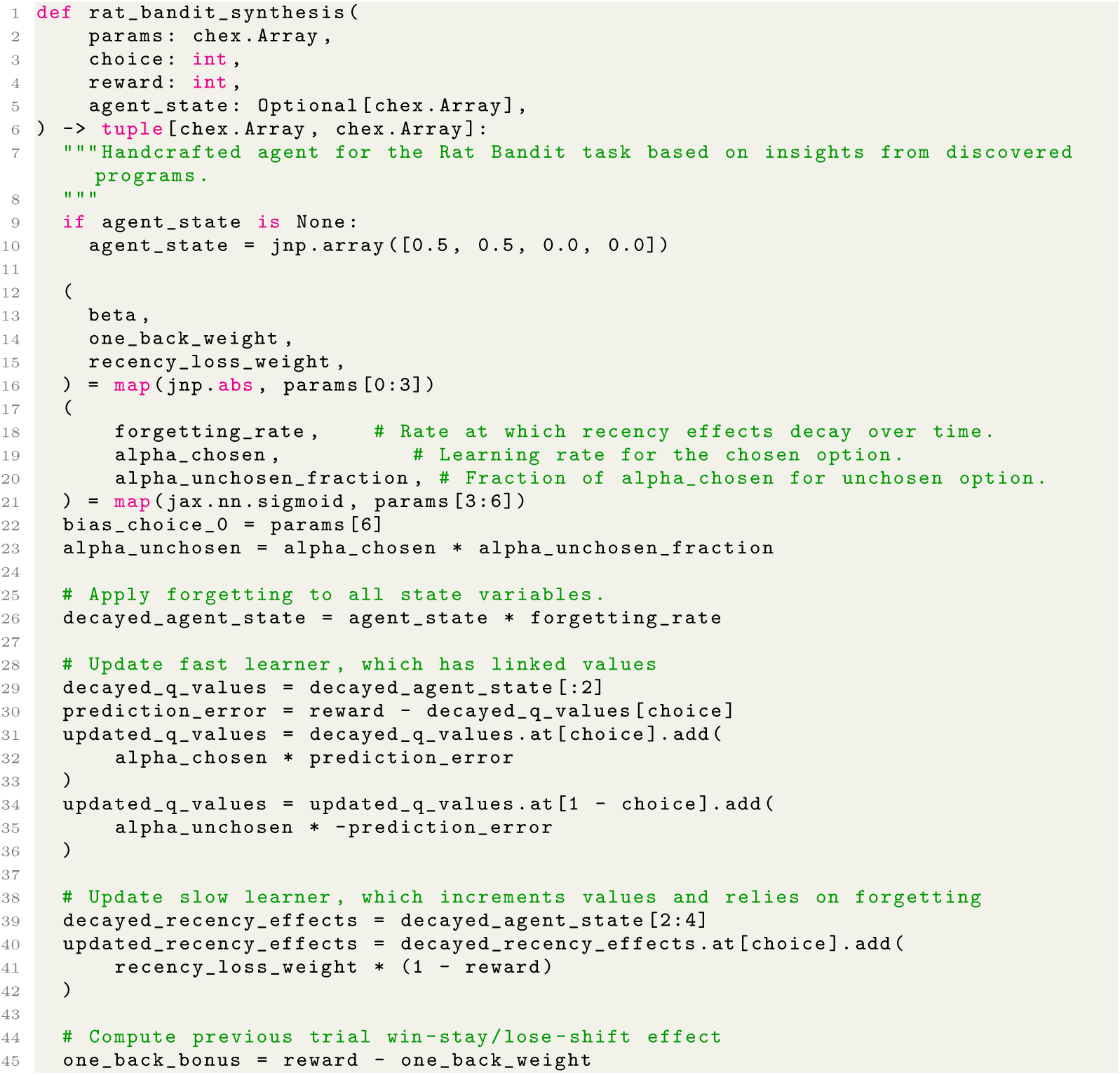
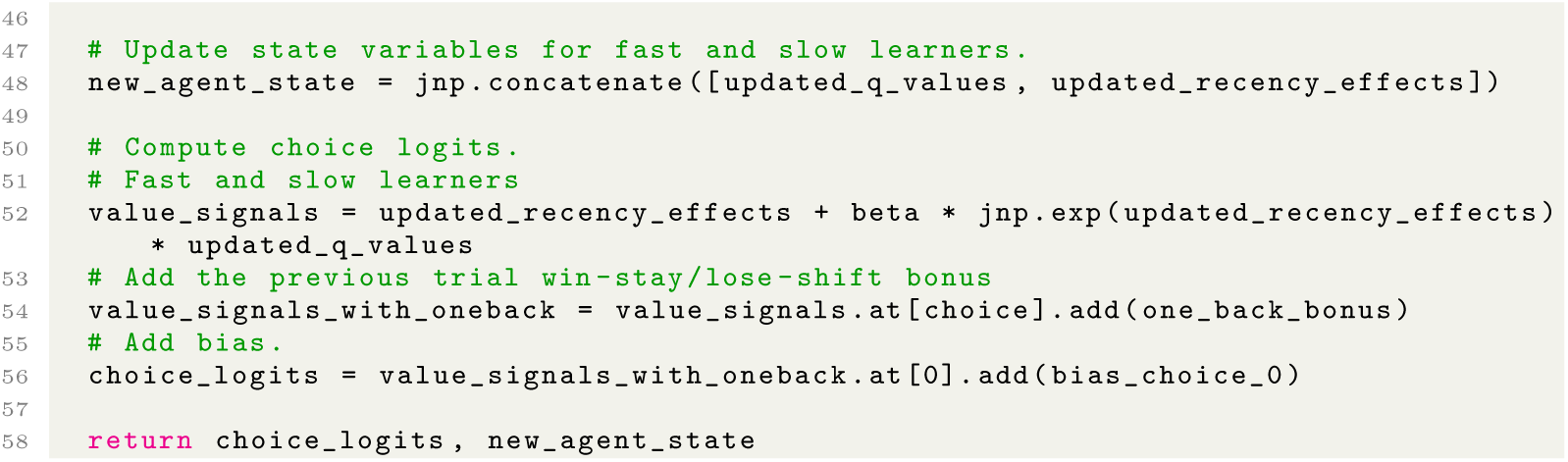

Code 7: Synthesis program for the rat bandit dataset.

#### A.2.7 Additional figures

**Fig. A5:**
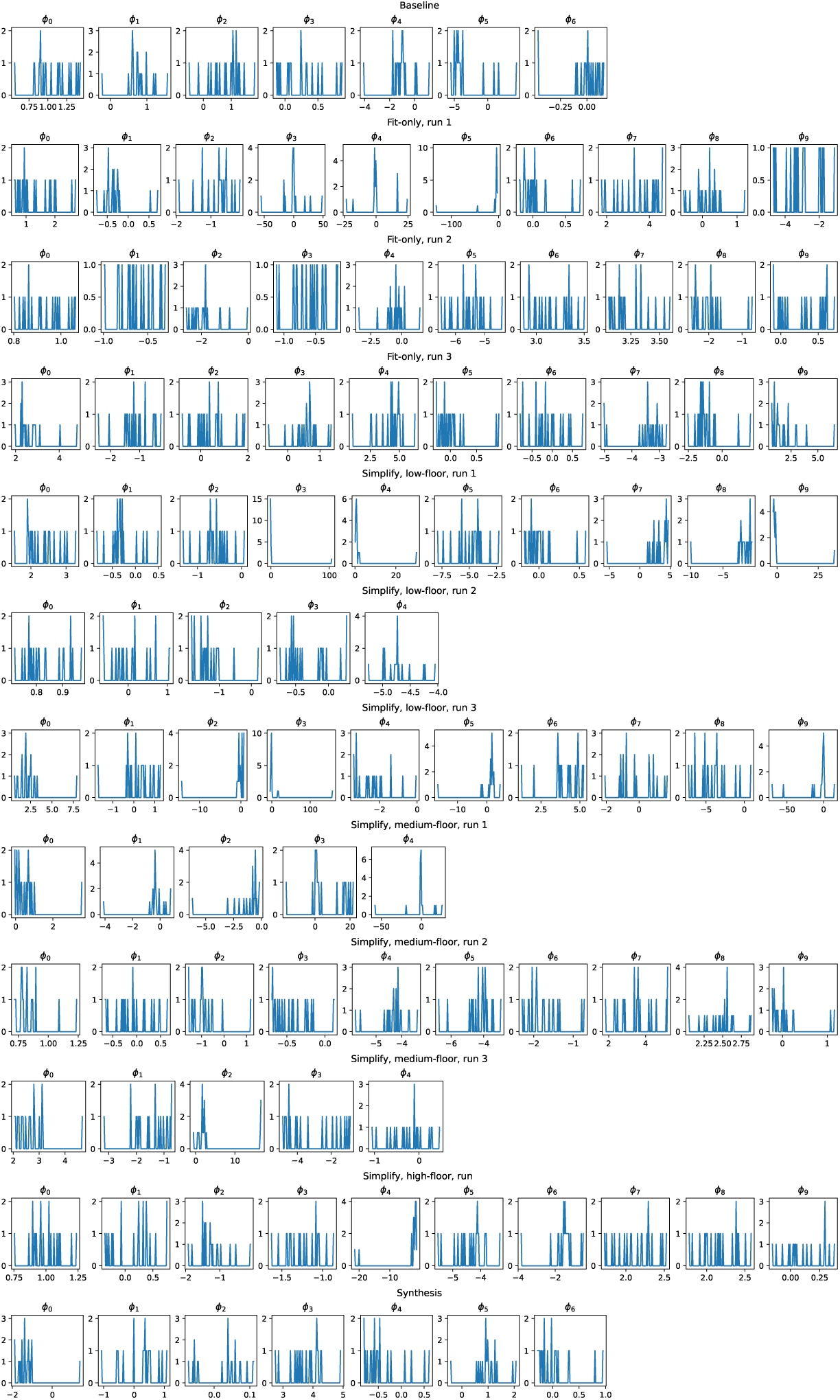
*Rat Bandit* Dataset: Fit parameters for each program. The distribution of fit parameters for each fold of all discovered programs (fit-only and simplified), as well as the handcrafted baseline and synthesis program.

**Fig. A6:**
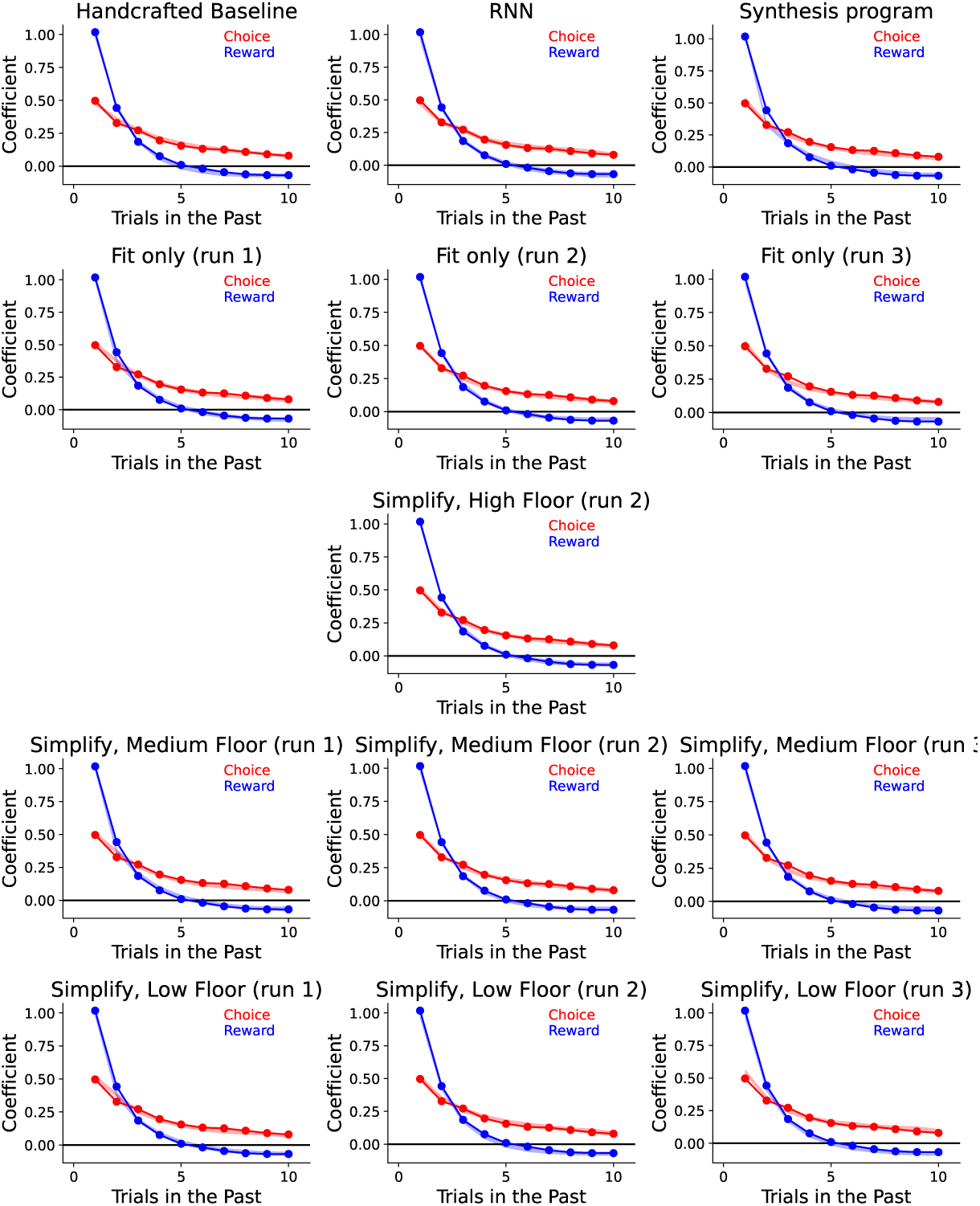
*Rat Bandit* Dataset: Trial-lagged regression analyses. Here we see the trial-lagged regression analysis shown in Figure 6 for all discovered programs for this dataset. The coefficients for the real data are shown in solid lines, while the transparent patch shows the 95% prediction interval for the artificial data. We see that all models closely match the data.

### A.3 Fly Bandit

#### A.3.1 Details of the dataset

Mohanta [34] considers fruit flies performing a two-armed bandit task with binary rewards. Flies performed the task in a Y-maze in which separate odors could be delivered to each of the three arms [81]. At the beginning of each trial, the arm containing the fly had no odor, and the other two arms contained one of two odors defining the choices. Flies indicated their choice on each trial by selecting an odor and walking to the end of the associated arm. Reward was delivered via a brief pulse of red light which activated flies’ sugar-sensing neurons [82]. Reward probabilities followed a random block structure with randomly sampled reward probabilities and block lengths. Each fly performed one session. The dataset contains choices from 347 flies performing 68,000 total trials. Each fly completed a single session.

This dataset is available from Glenn Turner upon request.

#### A.3.2 Parameter fitting in *Fly Bandit*

Because each fly completed a single session, it is not possible to fit parameters separately to each subject’s “even” and “odd” sessions. Thus, in order to fit parameters, we subdivide the sessions from all flies into a single “training” subject and a single “evaluation” subject. Within each “subject”, we perform two-fold cross-validation as described in Section 7.3. When reporting single-subject likelihood scores (as in Figure 3b), we still compute normalized likelihoods separately for each fly, and we still average across flies to compute the final score.

#### A.3.3 Handcrafted Baseline Program

Rajagopalan et al. [81] and Mohanta [34] have performed extensive model comparison on similar datasets, and identified a popular model known as “Differential Forgetting Q-Learning” (DFQ) as performing at least as well as any other. We adopt DFQ as the handcrafted baseline model for this dataset.

#### A.3.4 Discussion of evolved programs

##### Perseveration

Nearly half of the programs included a perseveration, or “stickiness”, factor. This models the animal’s tendency to repeat previous actions across subsequent trials. The strength of perseveration is determined by trainable parameters. We observe this feature in all simplified programs.

##### Eligibility traces

A third of the programs made use of *eligibility traces*. While typically considered as a “bridge from temporal-difference (TD) to Monte Carlo methods” [23], they can also serve as a form of memory for the occurrence of certain events (e.g. rewarding arms). They can be useful in bandit settings when rewards are non-stationary, as in our setup. The weight given eligibility traces in the update rule is determined by traininable parameters. It is worth noting that none of the low-floor programs with the lower included eligibility traces.

##### Confidence (difference in Q-values)

Over half of the discovered programs use the difference in learned action values as an indicator for *confidence*. Specifically, a higher difference in values is indicative of greater confidence in the higher-valued arm being rewarding. This confidence was sometimes used to drive exploration: less confidence resulted in a higher likelihood of selecting an arm randomly.

##### Reward history

Two thirds of the medium-and high-floor programs included some form of reward accumulation or history. This could take one of two forms: i) in some models, the choice-contingent reward history was used as a reference signal for computing prediction errors, a canonical feature of reinforcement learning; ii) in others, the total reward accumulated across the session, independent of which choices generated it, was used as gain control for learning, modulating the magnitude of updates. The latter form aligned with the notion of value sensitivity found in one of the discovered programs, where learning is slowed down if Q-values become too large; this is perhaps indicative of flies becoming reward-insensitive after too much stimulation.

##### Inverse temperature

Most of the medium- and high-floor programs use a parameterized inverse temperature to scale the final logits returned by the model, which helps control the exploration/exploitation tradeoff in arm selection. The value ultimately used to modulate the logits can be influenced by other factors such as confidence, mentioned above.

#### A.3.5 Discussion of synthesis program

The synthesis program is one of the generated programs (high-floor, run 1) with manual renaming of the variables for clarity. This program maintains action values, eligibility traces, and reward history.

The reward history is updated towards the most recent reward and decayed according to a decay parameter. A reward history weight is then computed by exponentiating the product of the reward history and a scaling parameter. Eligibility traces are first decayed according to a decay parameter; then the chosen option is increased according to a boost parameter, while the unchosen option is decreased by the same amount (with a floor at zero). The prediction error is given by the (signed) difference between the observed reward and the Q-value of the chosen option. The Q-value of the chosen option is then updated by multiplying the prediction error, a learning rate parameter, the reward history weight, and the eligibility trace for the chosen option. The Q-value of the unchosen option is decayed according to a decay parameter. The final decision variables (logits) are the product of the Q-values with an inverse temperature.

To verify the strength of our synthesis program, we evaluated a number of variants, all of which resulted in a reduction in accuracy. Specifically, we examined the following modifications to our synthesis program:

- Removing reward history weighting.
- Adding a surprise factor.
- Adding both a surprise factor and choice bias.
- Adding both a surprise factor and choice bias, and removing reward history weighting.
- Replacing eligibility traces with stickiness factors.

Figure A7 summarizes the comparison and demonstrates that, while all variants improve over the Handcrafted Baseline Program, our chosen synthesis program is the strongest of all variants.

**Fig. A7:**
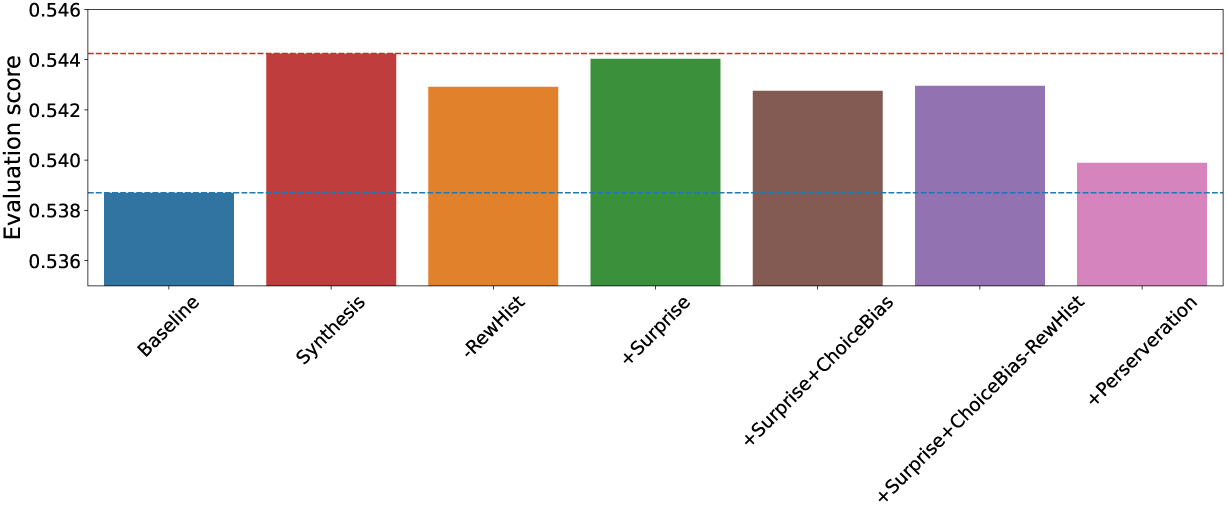
Comparison of Synthesis program against variants.

#### A.3.6 Code: stage 2 (“Simplify”) programs

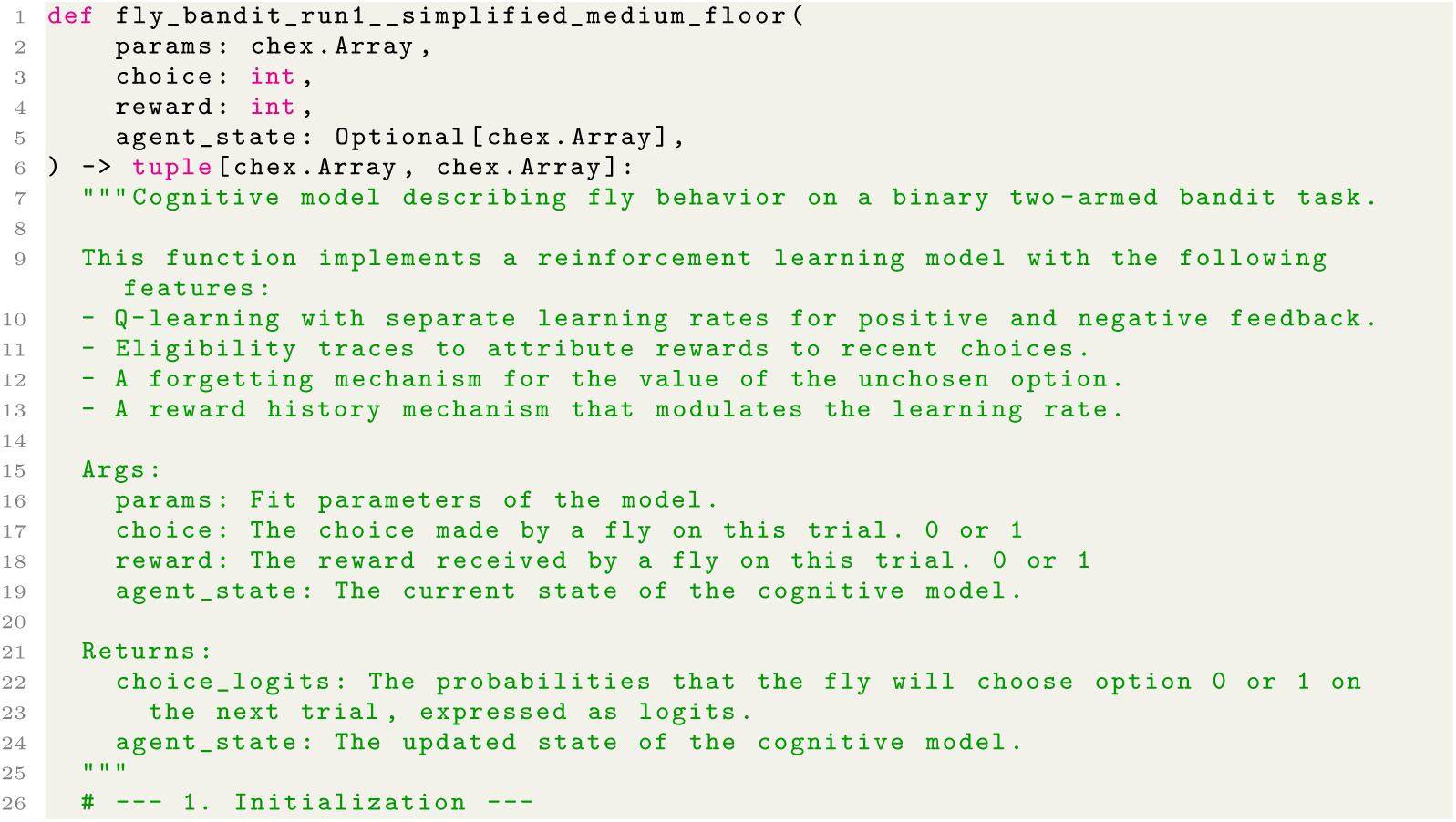
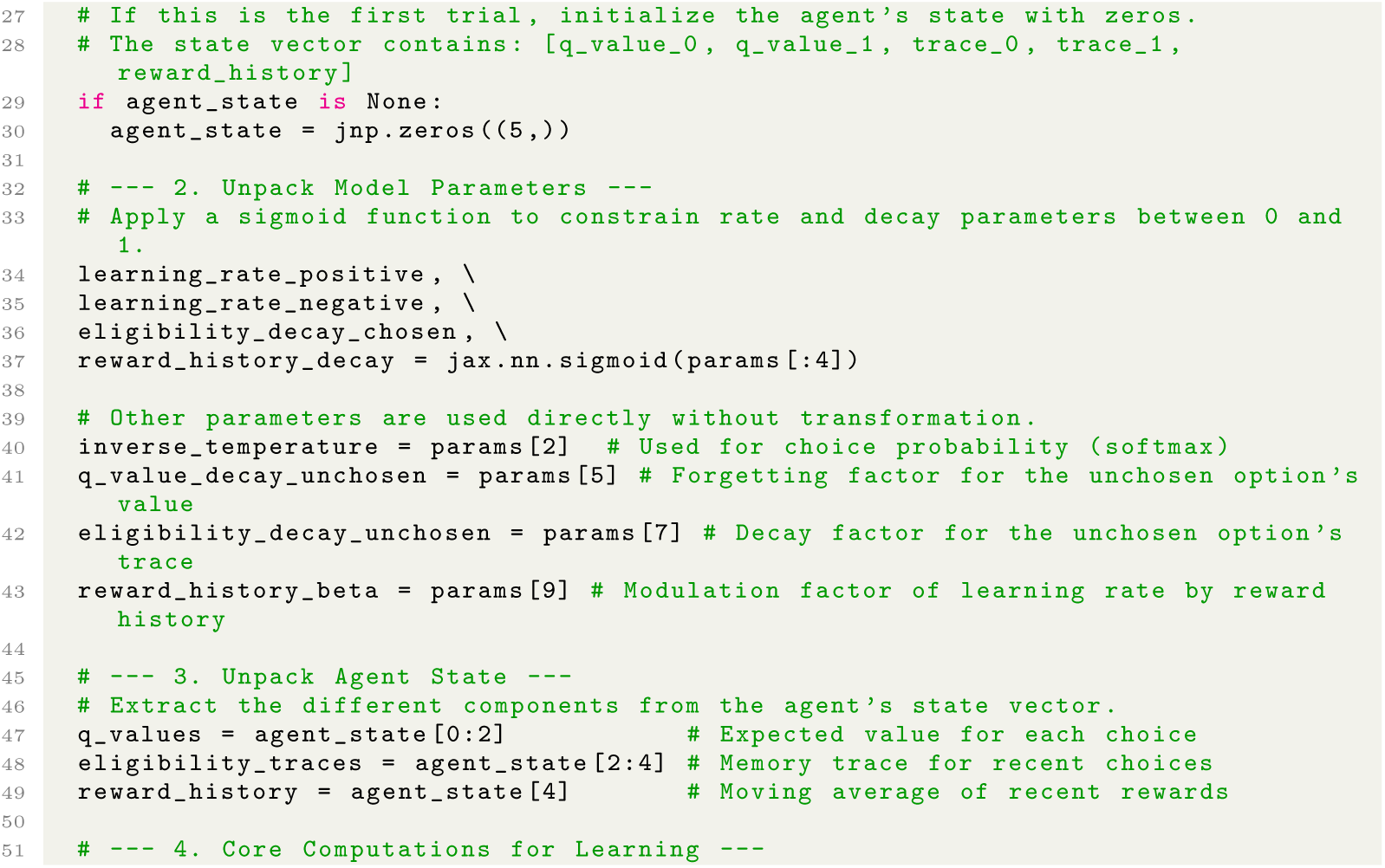
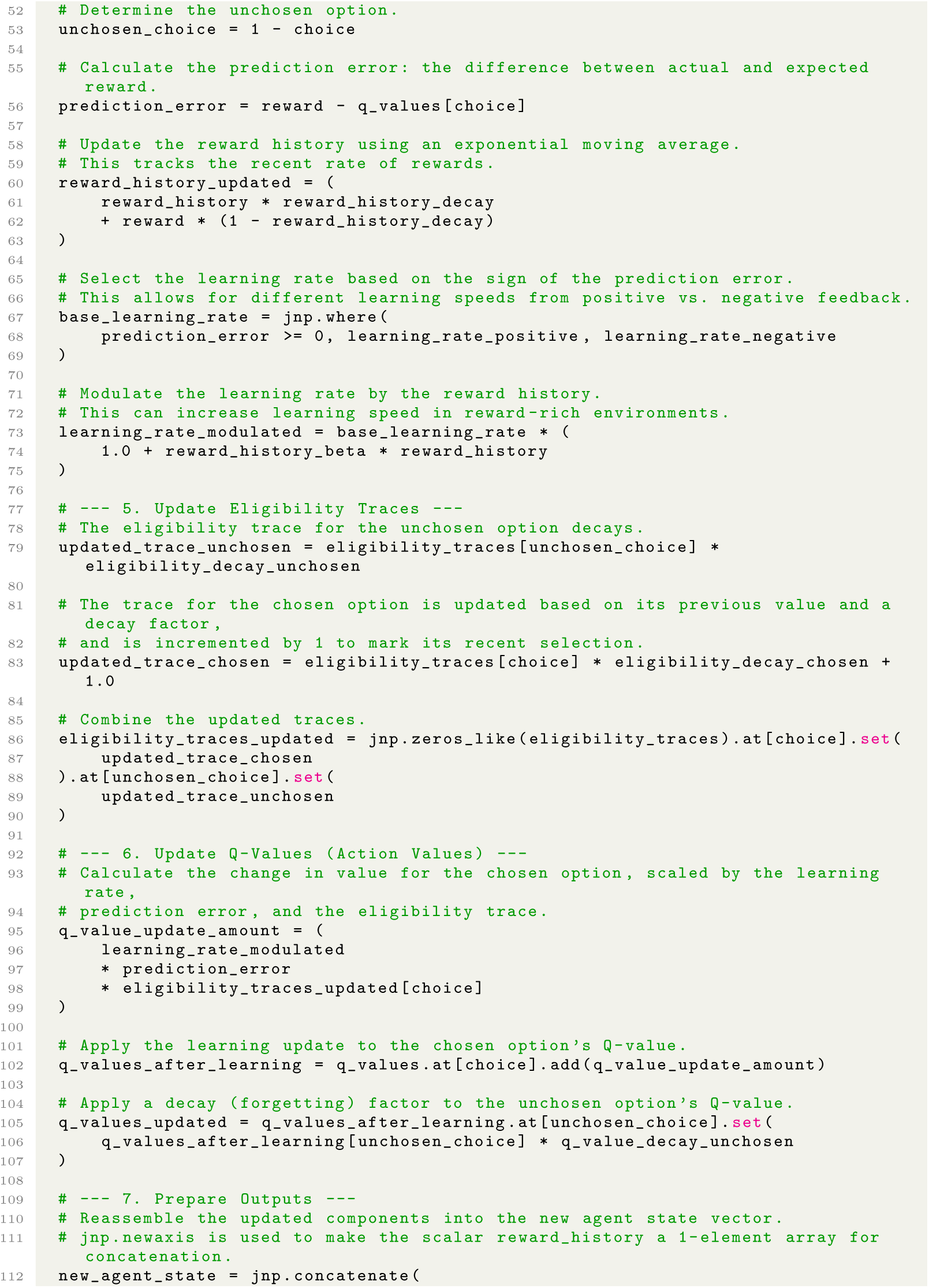
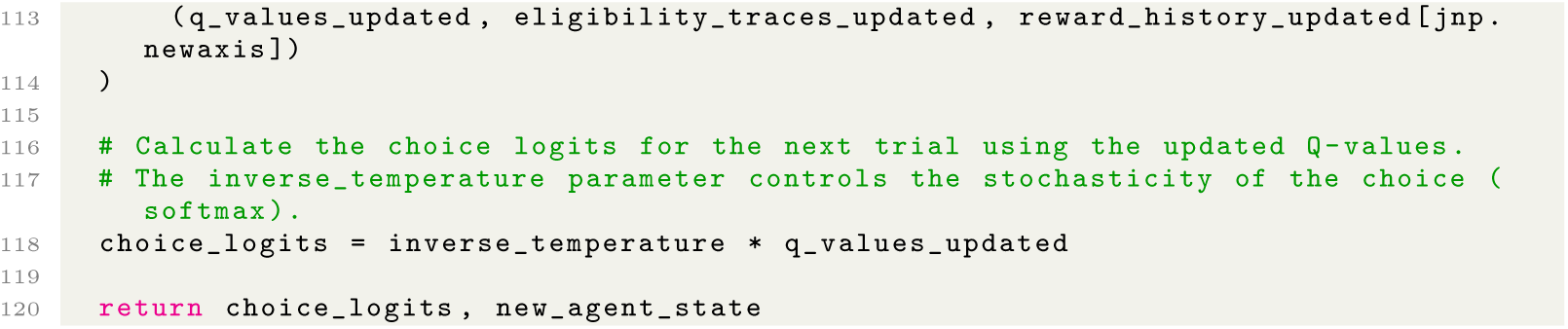

**Table A3:**
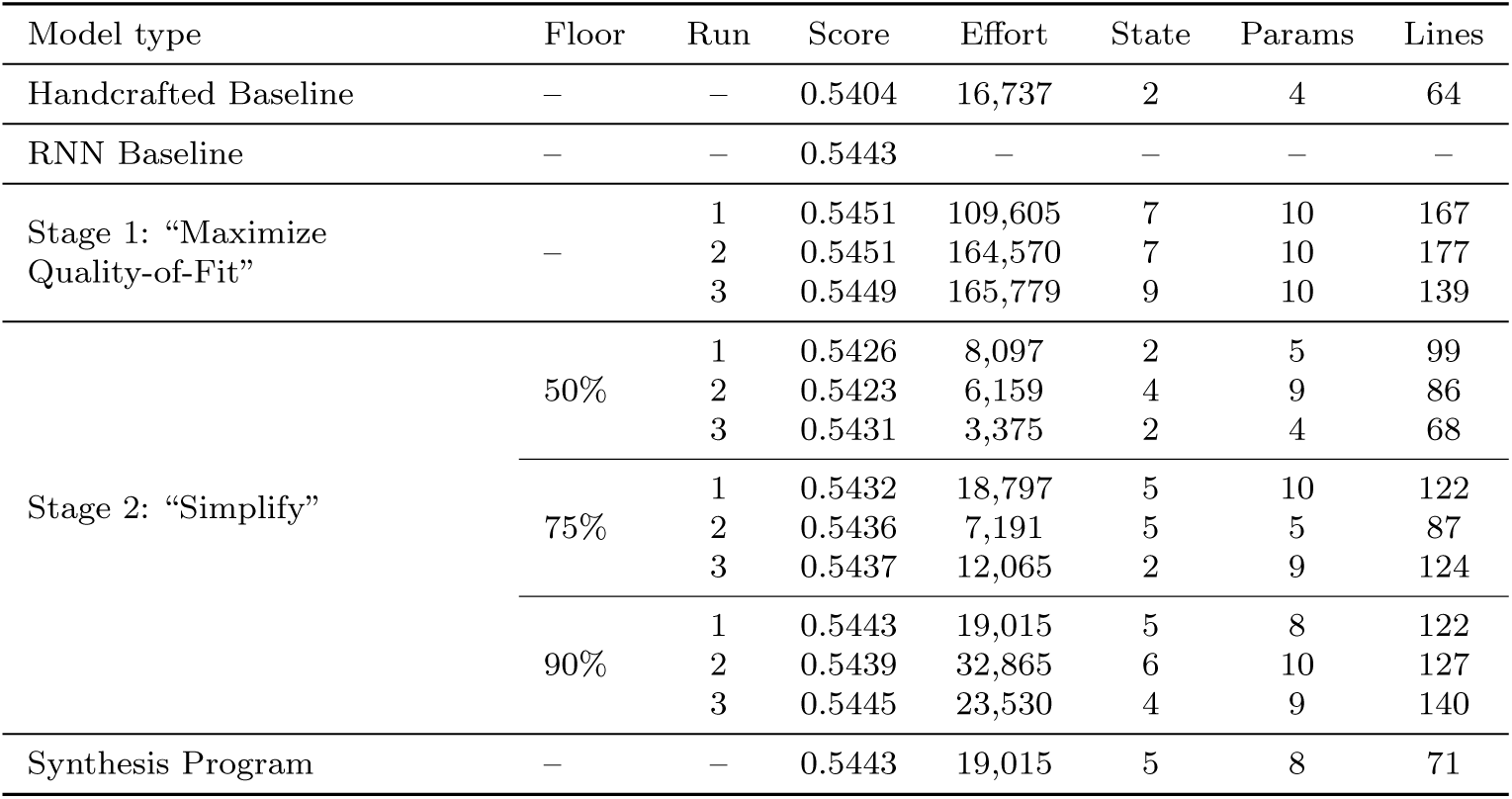
Evaluation performance and program complexity for models in the *Fly Bandit* dataset. For programs generated by the “Simplify” stage, Floor represents the quality-of-fit threshold below which programs are discarded (see Section 7.6.2). Score indicates the average normalized likelihood across evaluation subjects (see Section 7.3); Effort is Halstead effort. State, Params, and Lines indicate the number of state variables, per-subject parameters, and lines of code respectively.

Code 8: Lowest-complexity program from Stage 2 AlphaEvolve run with 75% threshold for the fly bandit dataset, evolved from programs in the first independent Stage 1 AlphaEvolve run and rewritten for readability (Stage 3).

#### A.3.7 Code: synthesis program

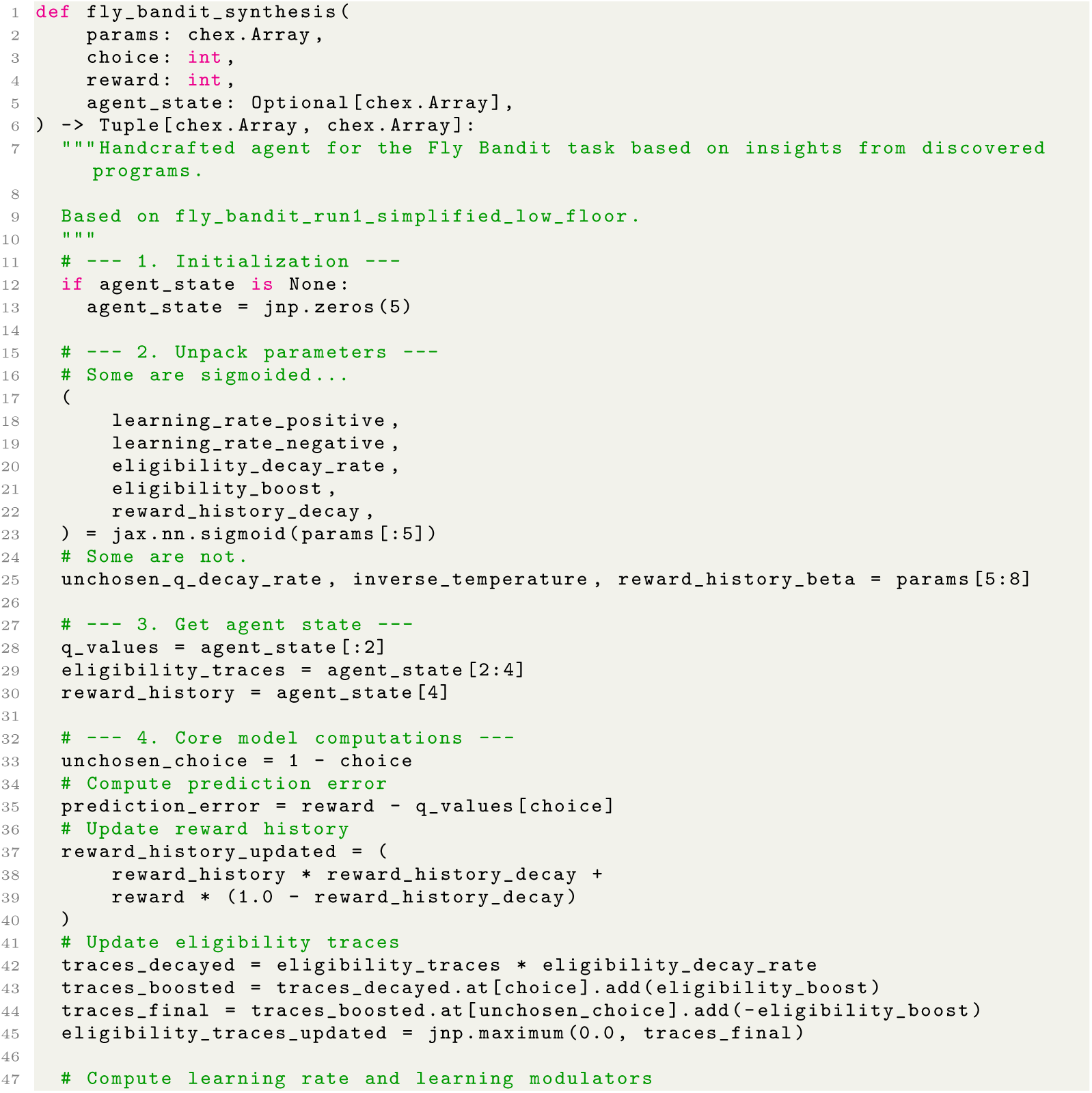
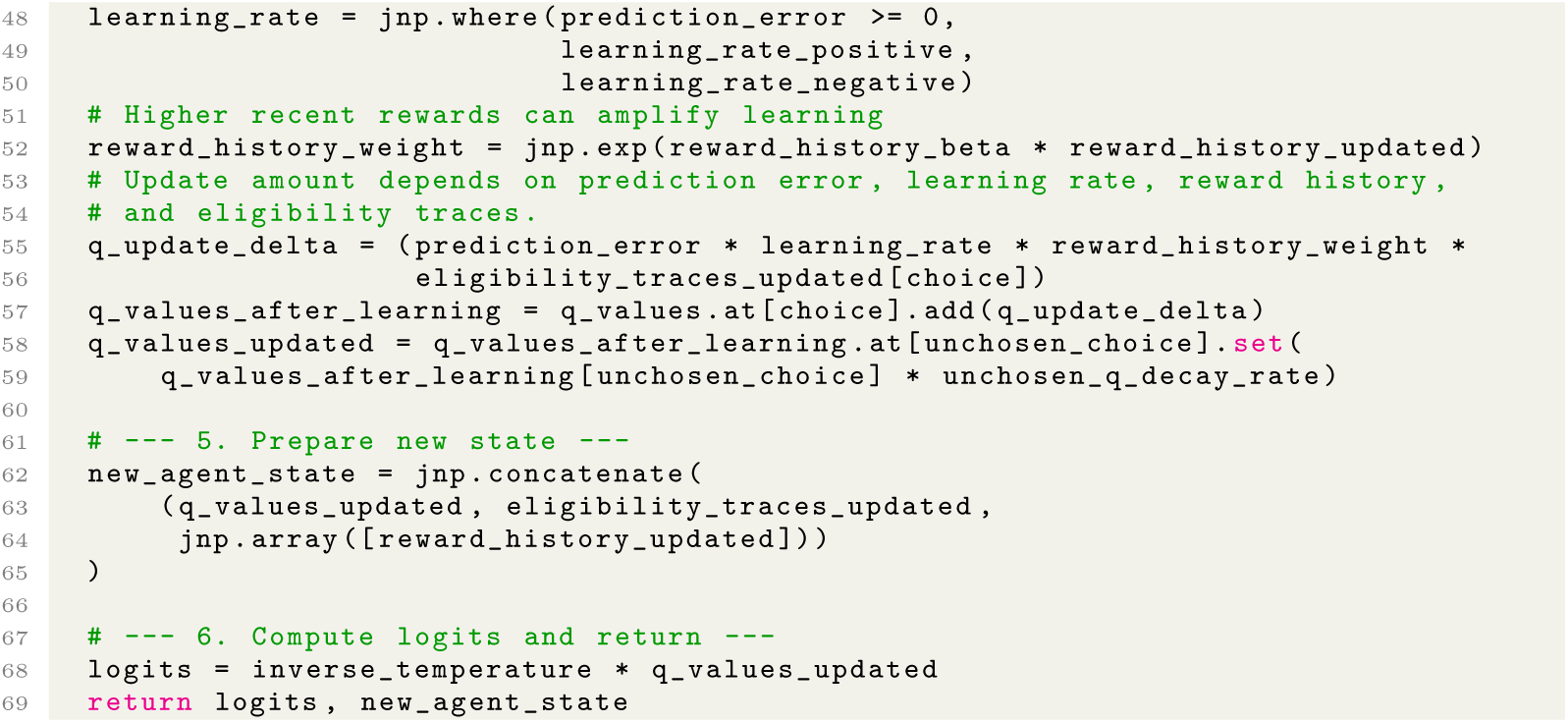

Code 9: Synthesis program for the fly bandit dataset.

#### A.3.8 Additional figures

**Fig. A8:**
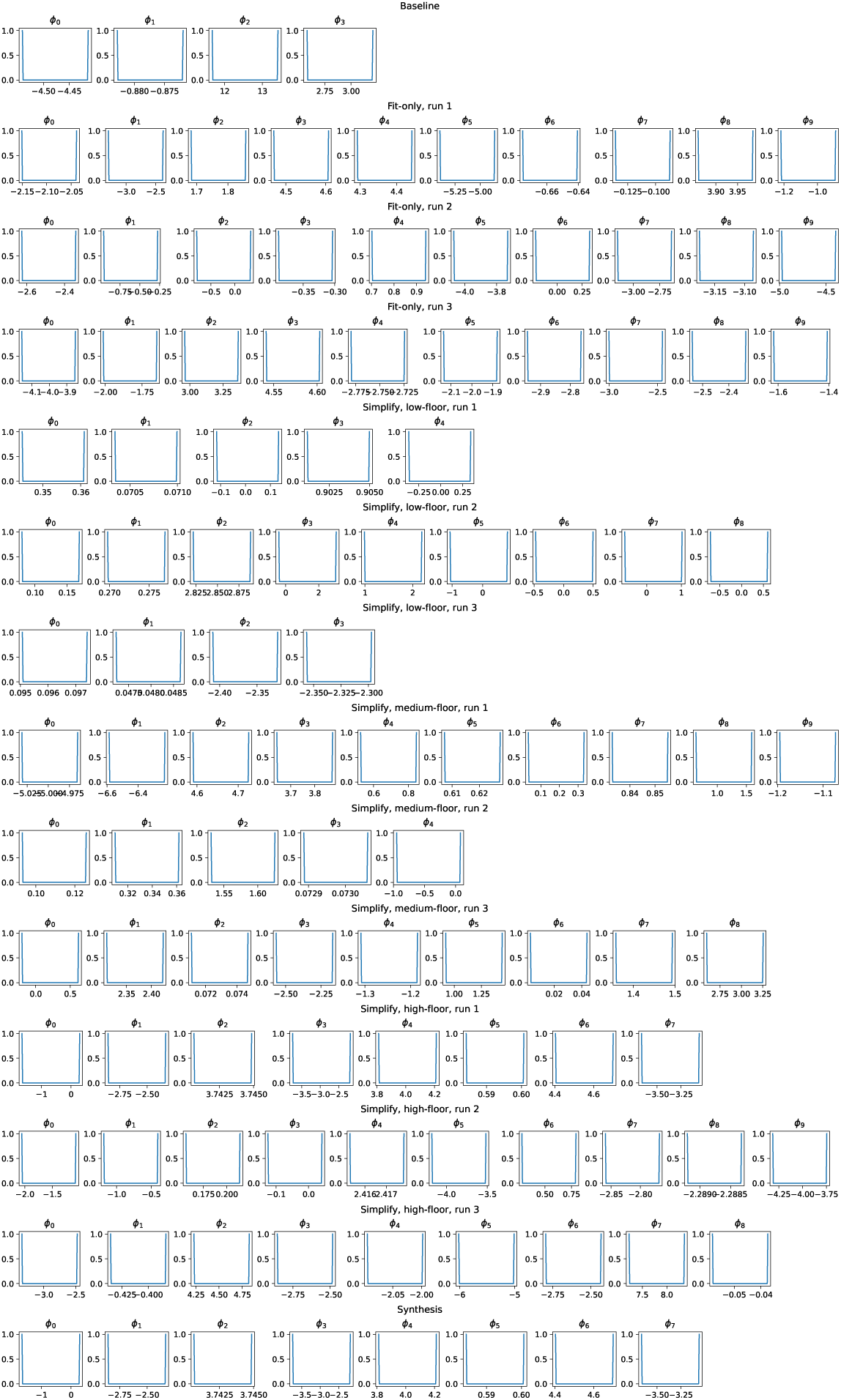
*Fly Bandit* Dataset: Fit parameters for each program. The distribution of fit parameters for each fold of all discovered programs (fit-only and simplified), as well as the handcrafted baseline and synthesis program. For the *Fly Bandit* dataset, the evaluation set consists of all even-indexed sessions, treated as if the data belonged to one subject: thus, there are only two cross-validation folds to which parameters are fit (and thus two data points per histogram).

**Fig. A9:**
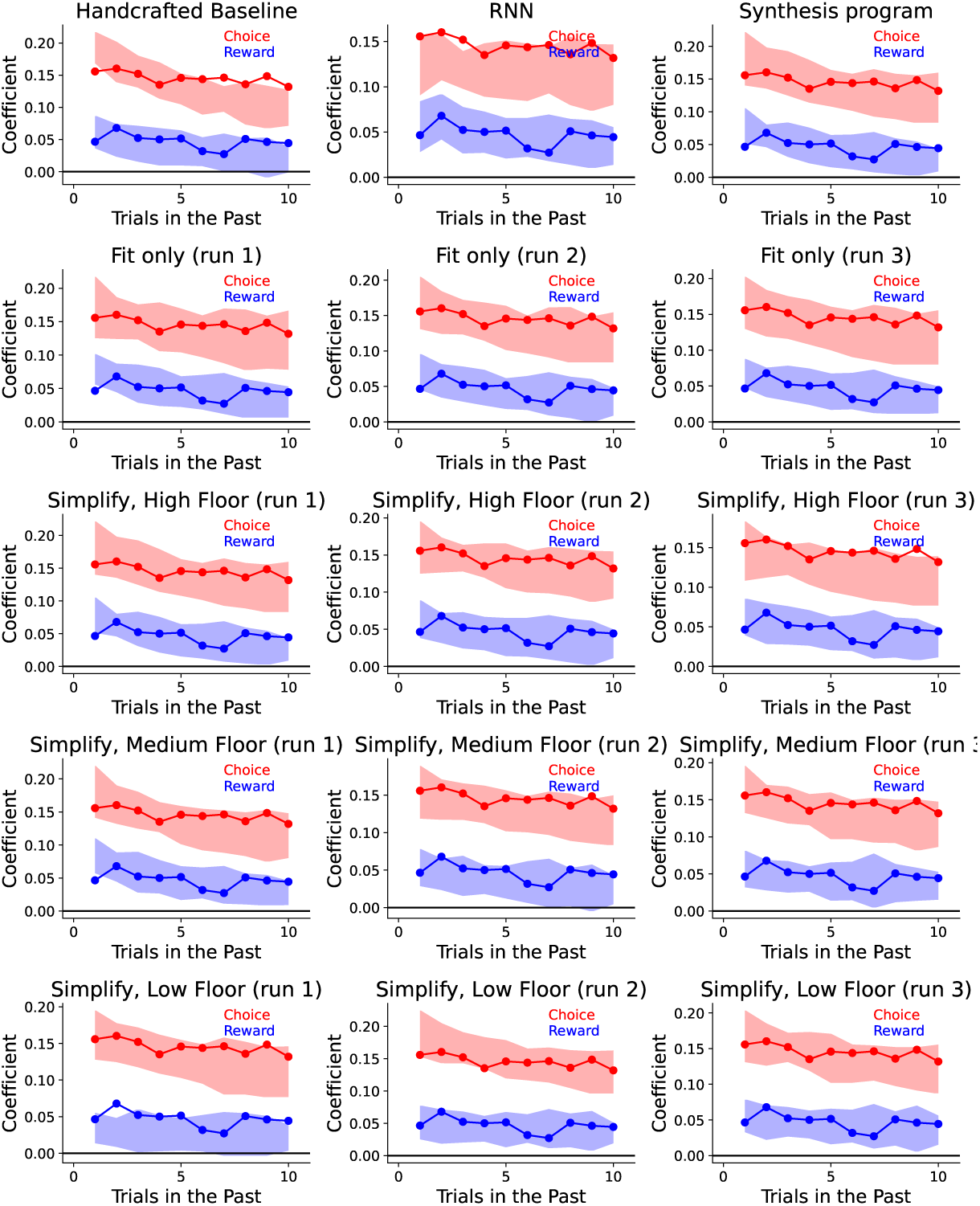
*Fly Bandit* Dataset: Trial-lagged regression analyses. Here we see the trial-lagged regression analysis shown in Figure 6 for all discovered programs for this dataset. The coefficients for the real data are shown in solid lines, while the transparent patch shows the 95% prediction interval for the artificial data. We see that the handcrafted baseline model does not match the reward-independent choice-driven perseveration, overestimating the effect on previous choice for recent trials and underestimating it for more distant trials. The discovered models more often contain the real data within the prediction intervals. We note that this dataset is considerably more stochastic than other datasets, and the coefficients on the *y*-axis occupy a smaller range than for other datasets.

### A.4 Monkey Bandit

#### A.4.1 Details of the dataset

Costa et al. [35] consider macaque monkeys performing a three-alternative bandit task using eye movements. On each trial three images are presented, and the monkey selects one of the images by making an eye movement to fixate on it. Each image was associated with a fixed probability of reward, either 20%, 50%, or 80%. The same set of three images was presented for a block of 10-30 trials, after which one of the images was switched out for a new image, with a new random reward probability, and a new block was begun. The dataset contains choices from nine monkeys performing 653 sessions and 412,342 total trials. This data can be made available upon publication.

#### A.4.2 Baseline Program

Costa et al. [35] introduced a Q-learning model which includes a novelty bonus for selecting a new image that has just become available, which we term “Novelty-Q”. Subsequent work has explored alternative models [36, 83], but none of these has provided a compelling advantage over Novelty-Q for the considered subjects, so we adopt Novelty-Q as our human-discovered baseline model for this dataset.

#### A.4.3 Discussion of evolved programs

##### Cognitive variables

Nearly all programs divided their cognitive variables into two terms: an action value (or “Q-value”) term which tracked expected reward for each choice, and a “novelty trace” term which was updated each time a novel option was introduced, and decayed over time. This division of cognitive variables surfaced in every program, across all levels of simplification, except for one (50% floor, run 2), a program which also shows worse performance at capturing the lagged regression novelty patterns (Figure A11). These variables were usually updated in a modular fashion (while some interactions arose in the more complex high-floor programs, these did not survive automated ablation).

These separate modules show a marked departure from the a core structural feature of the baseline model, in which all learning is localized to action values. This mechanism predicts separate neural substrates for reward learning and perceptual novelty. Prior models largely assumed a single action value tracking average reward with an added fixed novelty bonus—suggesting the brain encourages exploration via optimistic initialization [35, 46, 47]. However, this unified approach consistently underestimates the monkeys’ initial novelty seeking (bottom panel of Figure 6, Figure A11).

##### Reward learning

The action values for the chosen option were typically incrementally updated with reward prediction error based learning rule. Additionally, there was often a decay on the unchosen action values toward the initial action value. This parameter is also the value to which action values are initialized following either the beginning of a new session or the introduction of a novel option. Surprisingly, parameter fits resulted in *negative* values for this parameter in programs that also had a novelty trace, despite monkeys usually showing a preference for novel options. This was possible because novelty preference is represented by a separate variable that can counteract this. The baseline models, which also had a parameter for initial action values, returned positive values for this parameter for all monkeys.

Among the more complex “high floor” programs, two out of three introduced differential learning rates for rewards and omissions, which had a small positive contribution to performance. This was the only motif from the high floor programs that was incorporated into the synthesis program.

##### Novelty bonus

A recurring motif involved having the novelty trace updated by placing a strongly positive bonus parameter on the novel option, which was then gradually decayed toward zero on each trial. Interestingly, the discovered novelty traces were usually updated independently of both reward and choice: there was no dependence on how frequently the option had actually been sampled, or whether it resulted in reward. This is consistent with perceptual novelty that is driving novelty preference. Two programs from run 2 (medium- and high-floor) did exhibit an additional decay on unchosen options; however, ablations did not reveal that they consistently contributed to model performance.

##### Nonlinear, Nonstationary Exploration

A consistent motif across programs was a nonstationary, nonlinear exploration function mapping cognitive variables to the decision variables. This is part of a trend we see across discovered programs in this work, in which the function mapping cognitive variables to choice departs from the simple softmax rule often assumed. The generalized form of this mapping was:

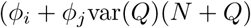

where *Q* and *N* are action value and novelty trace vectors respectively, and *ϕ_i_* and *ϕ_j_* are fittable parameters. This has the effect of increasing choice stochasticity when all action values are similar to each other, and making choices more deterministic when there is higher variance. We note that because novel options cause the corresponding action value to be set to a negative number, which is necessarily lower than the running average because rewards are only 0 and 1, var(*Q*) will be particularly high following the introduction of novel options as well.

#### A.4.4 Discussion of synthesis program

The synthesis program combined the elements described above that appeared consistently across the programs and contributed robustly to quality-of-fit. It consisted of two cognitive variables, Action Values (or Q-values) and Novelty Trace, which were updated in a modular fashion. Action values were updated by the learning rule described above (error-driven learning on the chosen action value with differential learning rates for rewards and omissions, decay on unchosen action values toward the initial action value parameter, reset on action value to initial action value parameter for novel options). Novelty trace was reset to a parametric novelty bonus parameter following the introduction of a novel option, and non-novel options were decayed toward zero, as described above. Cognitive variables for action values (*Q*) and Novelty trace (*N*) were mapped onto the decision variables using the expression (*ϕ_i_* + *ϕ_j_*var(*Q*)(*N* + *Q*).

**Table A4:** Evaluation performance and program complexity for models in the *Monkey Bandit* dataset. For programs generated by the “Simplify” stage, Floor represents the quality-of-fit threshold below which programs are discarded (see Section 7.6.2). Score indicates the average normalized likelihood across evaluation subjects (see Section 7.3); Effort is Halstead effort. State, Params, and Lines indicate the number of state variables, per-subject parameters, and lines of code respectively.

| Model type | Floor | Run | Score | Effort | State | Params | Lines |
| --- | --- | --- | --- | --- | --- | --- | --- |
| Handcrafted Baseline | – | – | 0.4220 | 12,352 | 3 | 6 | 70 |
| RNN Baseline | – | – | 0.4166 | – | – | – | – |
| Stage 1: “Maximize Quality-of-Fit” | – | 1 | 0.4264 | 60,406 | 6 | 10 | 96 |
|  |  | 2 | 0.4267 | 72,710 | 6 | 10 | 104 |
|  |  | 3 | 0.4270 | 240,368 | 12 | 10 | 162 |
| Stage 2: “Simplify” | 50% | 1 | 0.4258 | 9,172 | 6 | 8 | 97 |
|  |  | 2 | 0.4250 | 6,903 | 3 | 8 | 90 |
|  |  | 3 | 0.4255 | 10,603 | 6 | 6 | 119 |
|  | 75% | 1 | 0.4255 | 10,081 | 6 | 9 | 138 |
|  |  | 2 | 0.4258 | 18,965 | 6 | 10 | 130 |
|  |  | 3 | 0.4258 | 14,309 | 6 | 8 | 100 |
|  | 90% | 1 | 0.4264 | 31,921 | 6 | 10 | 156 |
|  |  | 2 | 0.4267 | 45,463 | 6 | 10 | 158 |
|  |  | 3 | 0.4269 | 26,977 | 7 | 9 | 188 |
| Synthesis Program | – | – | 0.4267 | 16,523 | 6 | 9 | 102 |

#### A.4.5 Code: stage 2 (“Simplify”) programs

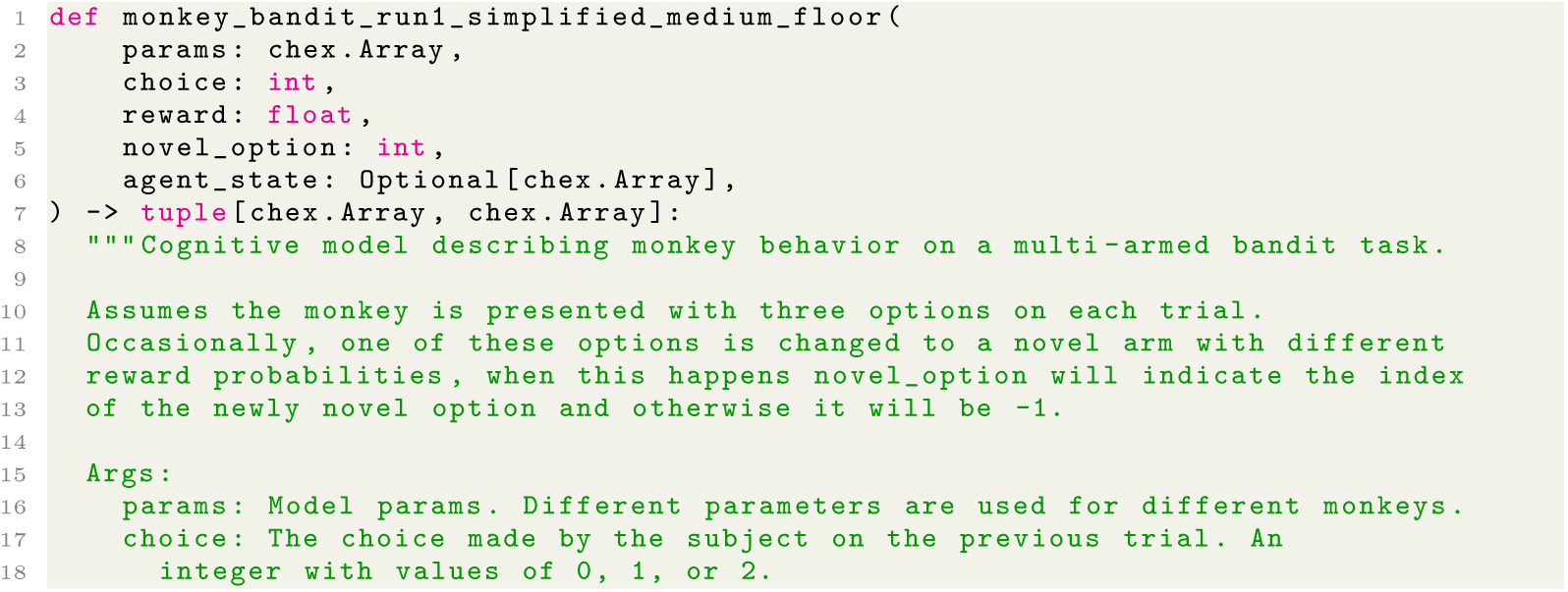
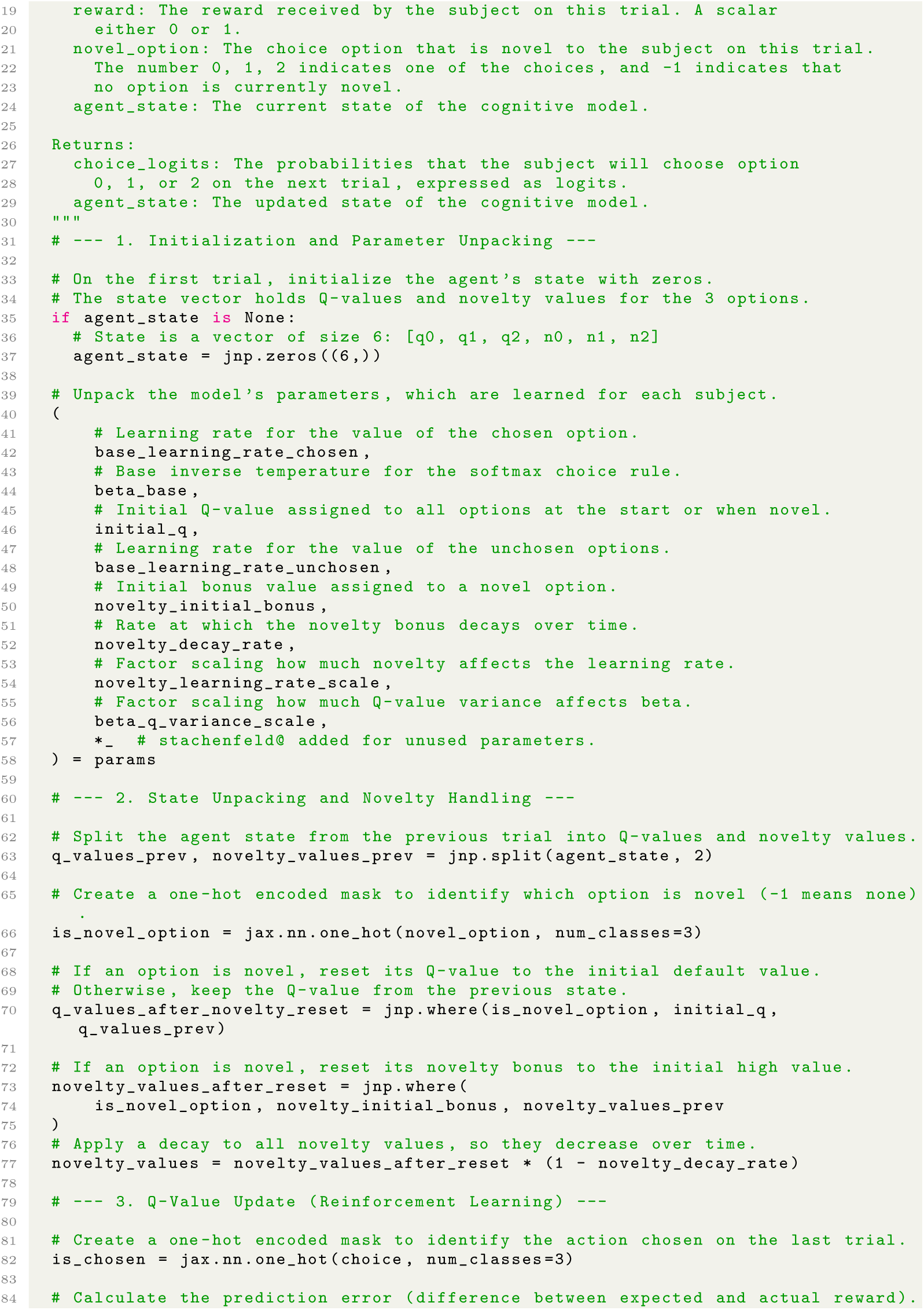
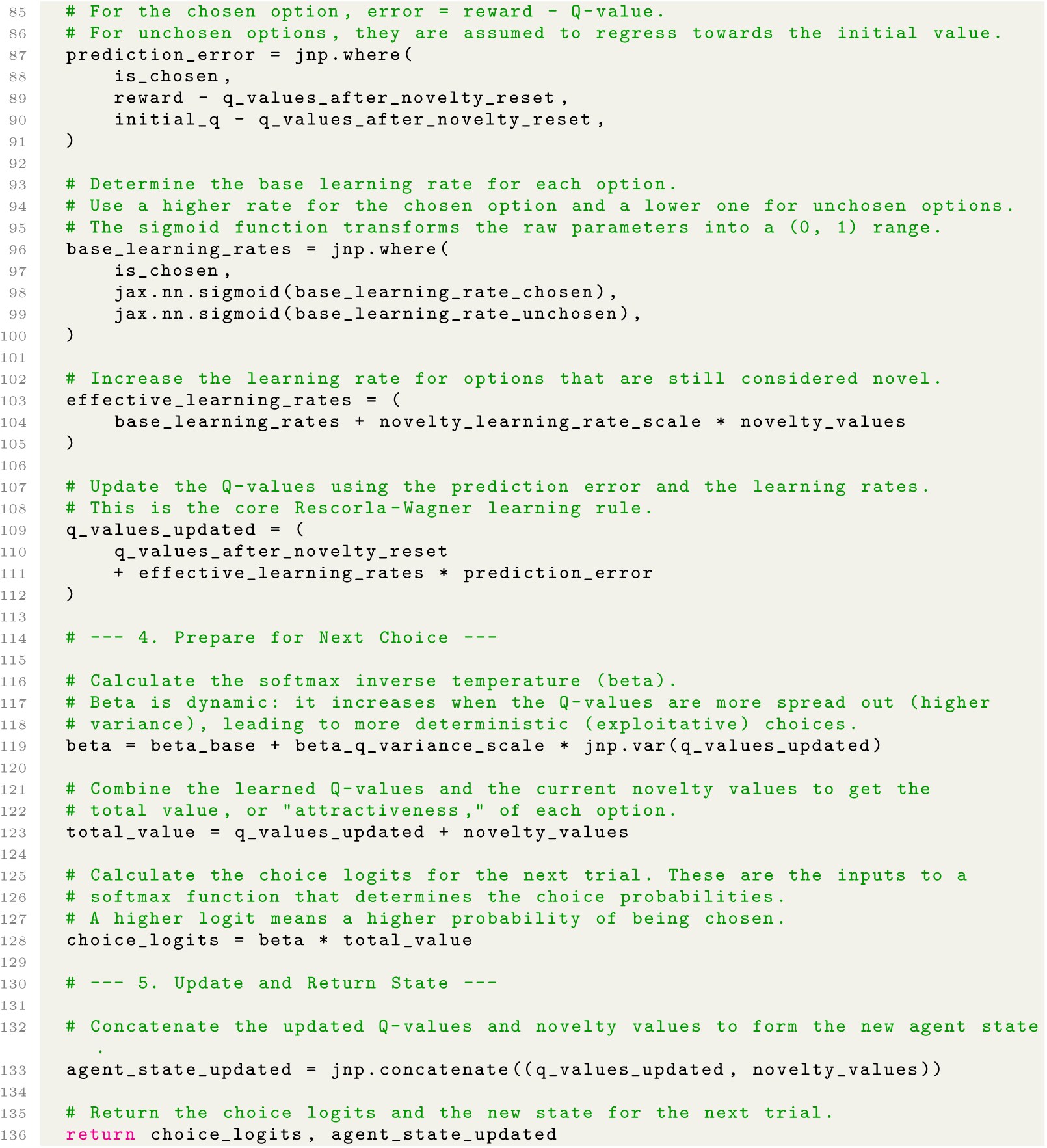

Code 10: Lowest-complexity program from Stage 2 AlphaEvolve run with 75% threshold for the monkey bandit dataset, evolved from programs in the first independent Stage 1 AlphaEvolve run and rewritten for readability (Stage 3).

#### A.4.6 Code: synthesis program

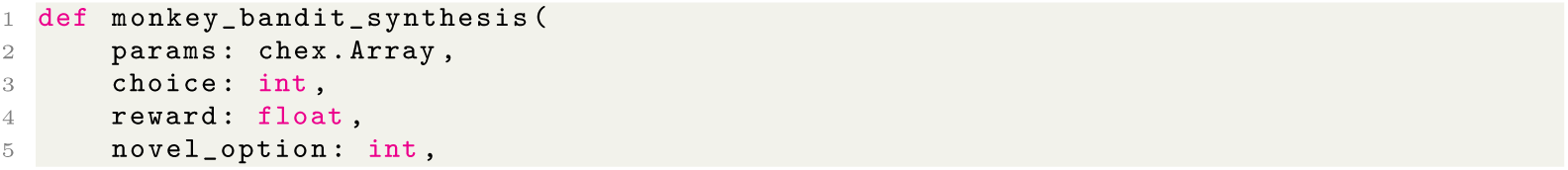
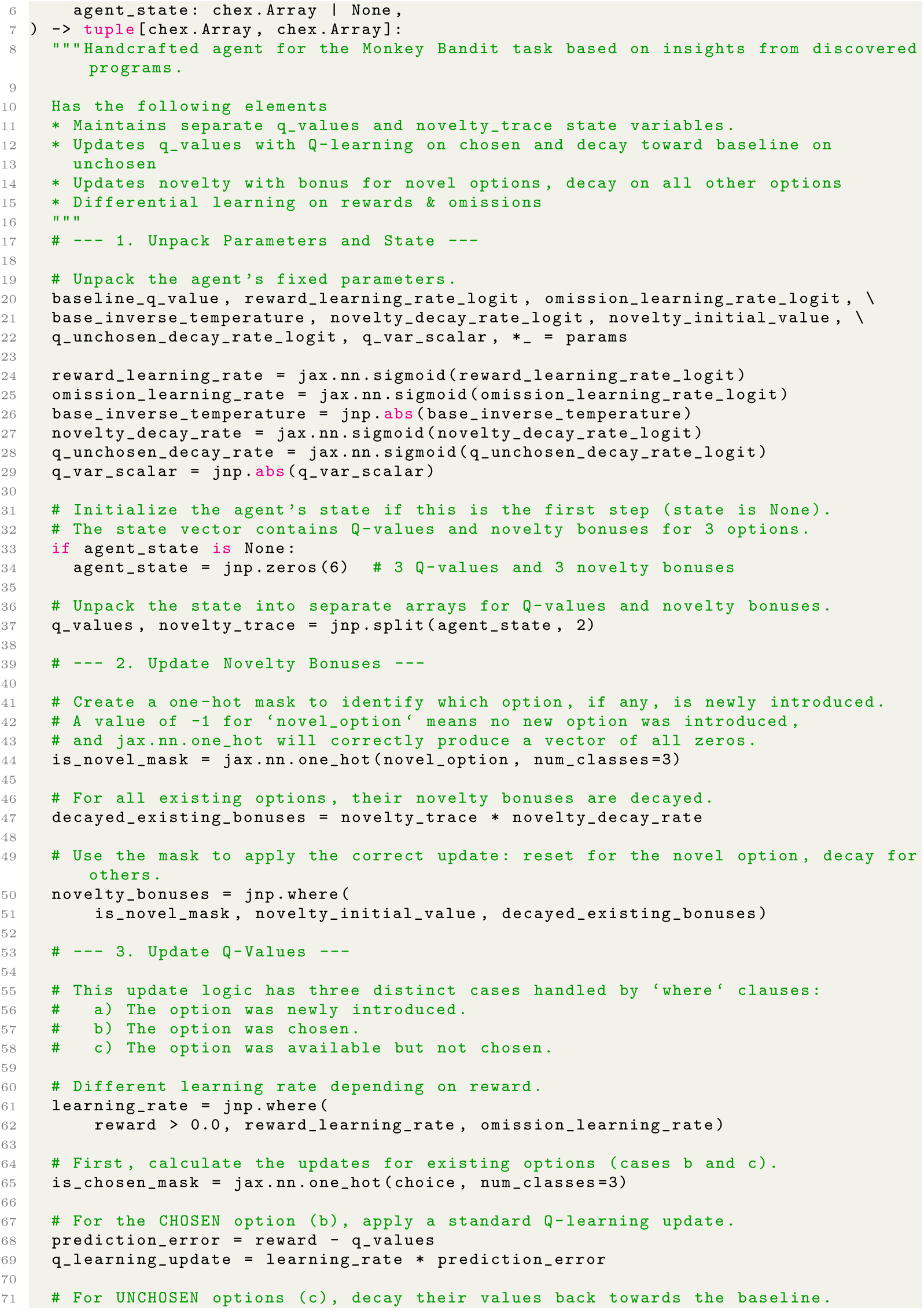
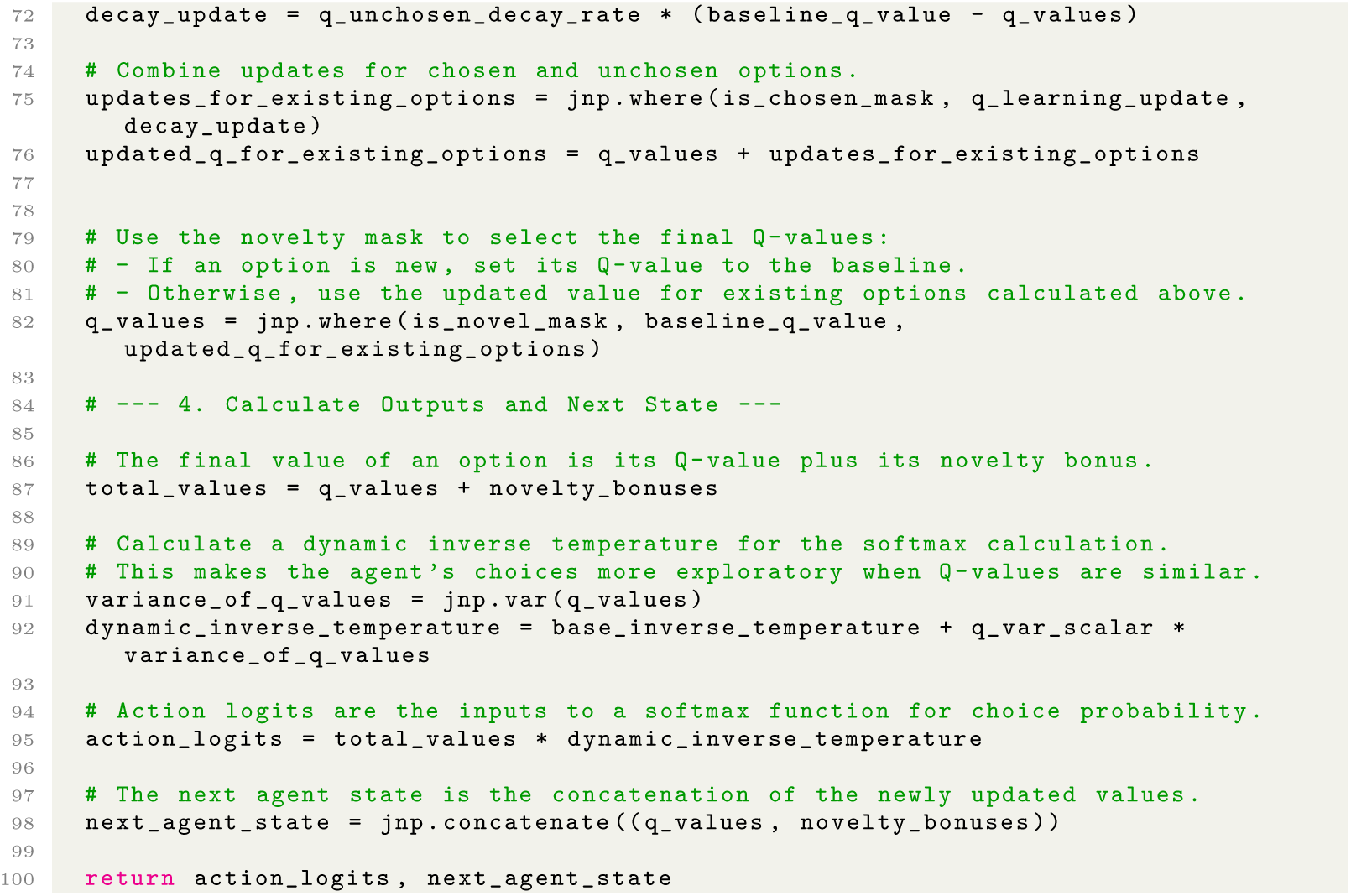

Code 11: Synthesis program for the monkey bandit dataset.

#### A.4.7 Additional figures

**Fig. A10:**
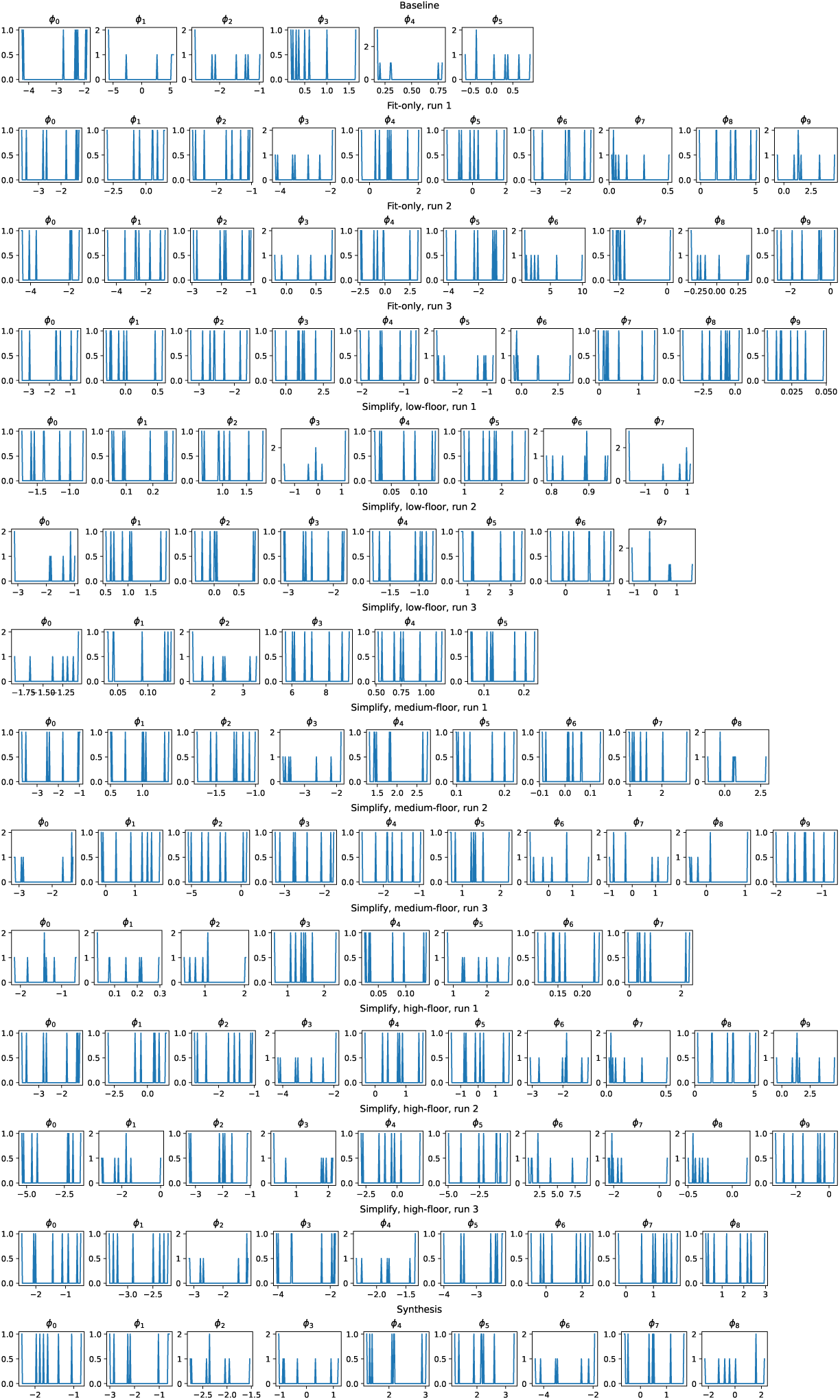
*Monkey Bandit* Dataset: Fit parameters for each program. The distribution of fit parameters for each fold of all discovered programs (fit-only and simplified), as well as the handcrafted baseline and synthesis program.

**Fig. A11:**
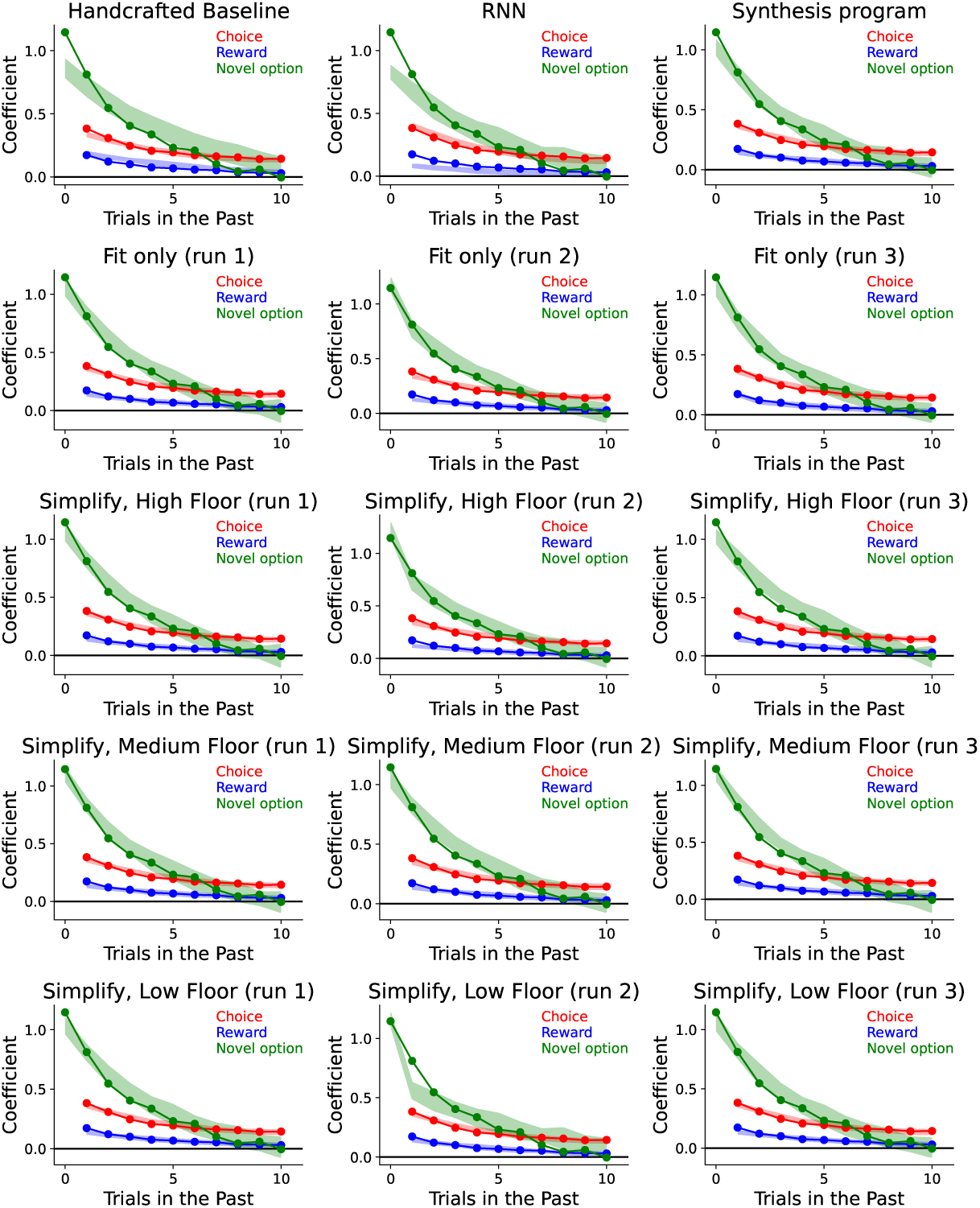
*Monkey Bandit* Dataset: Trial-lagged regression analyses. Here we see the trial-lagged regression analysis shown in Figure 6 for all discovered programs for this dataset. The coefficients for the real data are shown in solid lines, while the transparent patch shows the 95% prediction interval for the artificial data. We see that the handcrafted baseline model does not match the timecourse of novelty-seeking, while the other models do.

### A.5 Rat Twostep

#### A.5.1 Dataset

The rat two-step dataset [37] considers rats performing a two-step decision-making task that is commonly used to study model-based learning and decision-making. In the first step of each trial, the rat indicated its choice by entering one of two available “choice” ports. This was followed by one of two possible “outcome” ports becoming available. Each choice port was associated with one of the outcome ports, which became available following choices to that port with probability 80% (with probability 20% the other outcome port because available instead). The rat then entered the available outcome port and received a reward with probability that depended on the outcome port (but not the choice port). Reward probabilities were 80% and 20% for the two outcome ports and changed unpredictably in blocks. The dataset contains choices from 21 rats performing 1,960 sessions and 542,195 total trials.

We obtained this dataset from the following URL, where it is freely available under a permissive open-source license: https://github.com/kevin-j-miller/MBB2017-rat-two-step-task

#### A.5.2 Baseline Model

The paper which introduced this dataset also introduced a computational model, which was improved upon in several subsequent papers [38, 84]. The best-fitting cognitive model is a mixture of three agents: model-based reward learning, model-based perseveration, and model-free perseveration. This model is different from others in the literature in that it contains no influence of model-free reward learning, as model comparisons have shown that this does not improve quality of fit on this dataset [37, 38, 84]. It is also unusual in that each agent controls the update of just one decision variable, which expresses a relative preference between the two choice ports, rather than a pair of them expressing an absolute value for each port. We adopt this model as the human-discovered baseline for the rat two-step dataset. For compatibility with our pipeline, we re-implemented this model in Jax.

#### A.5.3 Discussion of evolved programs

##### Model-free and model-based learning

All but two simplified programs compute a weighted sum between action values (model-free learning) and outcome values (model-based learning). Of the remaining two programs, one has something referred to as “Q-values” for actions but which actually resemble a recency trace (perseveration), and computes a weighted sum between these and the model-based values (see “Recency traces”); the other computes a weighted sum over two systems, but neither system is purely model-free (the “model-free” system updates Q-values using both model-free and and model-based prediction errors).

##### Inverse temperature

All simplified programs scale the weighted sums described above by an inverse temperature. In the majority of programs, inverse temperatures are given by a fixed parameter. In two programs (a medium- and a high-floor program, simplified from the same fit-only program), this inverse temperature changes over time, beginning at zero and grows over time, asymptoting at a final value specified by a fit parameter. This is expected to cause the models to make more random choices at the beginning of each session, and to slowly become more deterministic throughout the session. This pattern may be similar to the patterns identified in [38] using statistical models. The evolved models here represent an advance on this in that they are generative, runnable models.

##### Connecting actions and outcomes

In the underlying experiment, rats are assigned to one of two conditions: the congruent condition, in which the action matches the outcome 80% of the time, and the incongruent condition, in which the choice matches the outcome 20% of the time. In all simplified programs but one, the experimental condition (and thus the relationship between actions and outcomes) is implicitly encoded in the sign of a product of per-subject parameters (typically the model-based learning rate, the weight of the outcome values relative to the action values, and the inverse temperature). The remaining program explicitly learns the transition function over the course of a session, and thus does not use parameters to encode the relationship between actions and outcomes.

##### Update of unchosen values

All simplified programs update the (model-free) values for the unchosen action (three programs also decay the *chosen* action value before updating it). In all but one program this takes the form of decay towards a fixed target, either zero (five programs) or a per-subject parameter (three programs). The remaining program decays the value of the unchosen action towards half the value of the chosen action. Each of these processes can be thought of as a different model of forgetting.

Similarly, all programs decay their (model-free) values for the unobserved outcome (the *low floor* programs also decay the *observed* outcome value before updating it). In all programs but two, these updates take the form of decay, either to a fixed per-subject parameter (six programs) or to zero (one program). The remaining two programs do a form of counterfactual learning: one program decays the unobserved outcome value towards 1 − *r* where *r* is the reward observed in this trial (using the assumption that if outcome *o* has reward *r*, then outcome 1 − *o* would have reward 1 − *r*). The other program decays first to zero, and then to 1 − *r*.

Decay towards zero [41] and towards a fit parameter [39] are known motifs from the literature on related tasks. Counterfactual learning is a known motif in modeling human decision-making in related tasks, though it is usually deployed in situations where participants were aware of the counterfactual outcome (what would have happened had they chosen the alternative option) [85, 86]. Notably, “counterfactual” learning for the unchosen action, using the same learning rate as for the chosen action, is equivalent to tracking only a single decision variable which expresses a relative preference [8, 37]. These motifs have not, to our knowledge, been deployed in computational models of the rat two-step task. Decay towards the current value of the chosen action is to our knowledge an entirely novel computational motif, which occurs in our evolved models of the human bandit dataset as well.

##### Bias and stickiness terms

All simplified programs except for one have at least one of the following: a direct bias towards one action or another, specified by a per-subject parameter (four programs); an “action stickiness” term directly rewarding or penalizing the immediately previous action (five programs); and/or an “outcome stickiness” term directly rewarding the choice with the same index as the current outcome (four programs; one of these has a separate outcome stickiness term for each outcome). No program has all three.

##### Habit traces

Three programs (none of which include an “action stickiness” term as described above) use something resembling an habit trace for action perseveration [45]. Namely, each of these three programs maintain per-action values which decay when the action is unchosen, and increase when the action is chosen. These can be thought of as maintaining a running average of how often each action has been chosen in the recent past. As mentioned above, one of these programs misleadingly refers to its recency trace as “Q-values”; the others call their recency traces “stickiness values”.

#### A.5.4 Discussion of synthesis program

Due to both prior literature and our analysis of discovered programs revealing that ablating the model-free component of these programs (e.g., by replacing them with an eligibility trace) generally has a minimal effect on quality of fit, the synthesis program specifically omits model-free (action) Q-values. Aside from this, the program follows the same basic skeleton as most of the discovered programs, with the final choice logits a weighted sum of a habit trace and model-based (outcome) values, where the sign of the model-based weight is positive for subjects in the congruent condition, and negative in the incongruent condition. The outcome-based values decay towards a per-subject fitted parameter; the habit trace, which only requires one state variable to store (its learning target is +1 when the rat chooses right, and −1 when the rat chooses left), decays towards 0.

The synthesis model includes both a direct left-right bias and an “action stickiness” bias favoring the last action performed by the rat. Thus, there are two separate perseveration pathways: one favoring repeating the immediately preceding action (the action stickiness bias), and one favoring repeating whichever action was performed most frequently in the recent past (the habit trace).

Finally, the synthesis program incorporates the dynamic inverse temperature calculation present in two of the discovered programs, in which the inverse temperature gradually decays from zero to one at a fixed rate (resulting in the stochasticity of the program’s choices gradually decreasing). This perhaps models the rat’s re-acclimatization to the task at the beginning of each session.

**Table A5:**
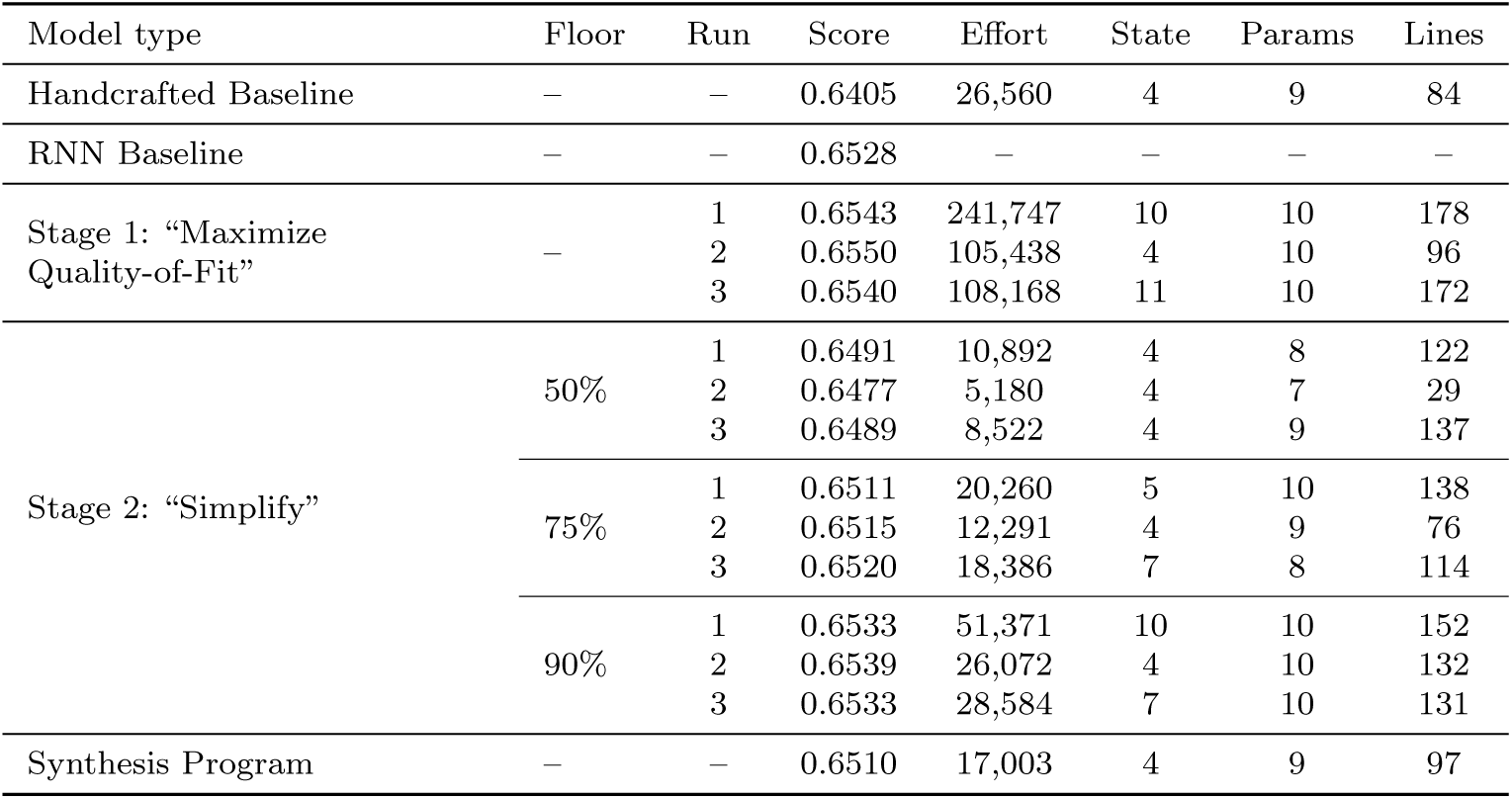
Evaluation performance and program complexity for models in the *Rat Two-step* dataset. For programs generated by the “Simplify” stage, Floor represents the quality-of-fit threshold below which programs are discarded (see Section 7.6.2). Score indicates the average normalized likelihood across evaluation subjects (see Section 7.3); Effort is Halstead effort. State, Params, and Lines indicate the number of state variables, per-subject parameters, and lines of code respectively.

| Model type | Floor | Run | Score | Effort | State | Params | Lines |
| --- | --- | --- | --- | --- | --- | --- | --- |
| Handcrafted Baseline | – | – | 0.6405 | 26,560 | 4 | 9 | 84 |
| RNN Baseline | – | – | 0.6528 | – | – | – | – |
| Stage 1: “Maximize Quality-of-Fit” | – | 1 | 0.6543 | 241,747 | 10 | 10 | 178 |
|  |  | 2 | 0.6550 | 105,438 | 4 | 10 | 96 |
|  |  | 3 | 0.6540 | 108,168 | 11 | 10 | 172 |
| Stage 2: “Simplify” | 50% | 1 | 0.6491 | 10,892 | 4 | 8 | 122 |
|  |  | 2 | 0.6477 | 5,180 | 4 | 7 | 29 |
|  |  | 3 | 0.6489 | 8,522 | 4 | 9 | 137 |
|  | 75% | 1 | 0.6511 | 20,260 | 5 | 10 | 138 |
|  |  | 2 | 0.6515 | 12,291 | 4 | 9 | 76 |
|  |  | 3 | 0.6520 | 18,386 | 7 | 8 | 114 |
|  | 90% | 1 | 0.6533 | 51,371 | 10 | 10 | 152 |
|  |  | 2 | 0.6539 | 26,072 | 4 | 10 | 132 |
|  |  | 3 | 0.6533 | 28,584 | 7 | 10 | 131 |
| Synthesis Program | – | – | 0.6510 | 17,003 | 4 | 9 | 97 |

#### A.5.5 Code: stage 2 (“Simplify”) programs

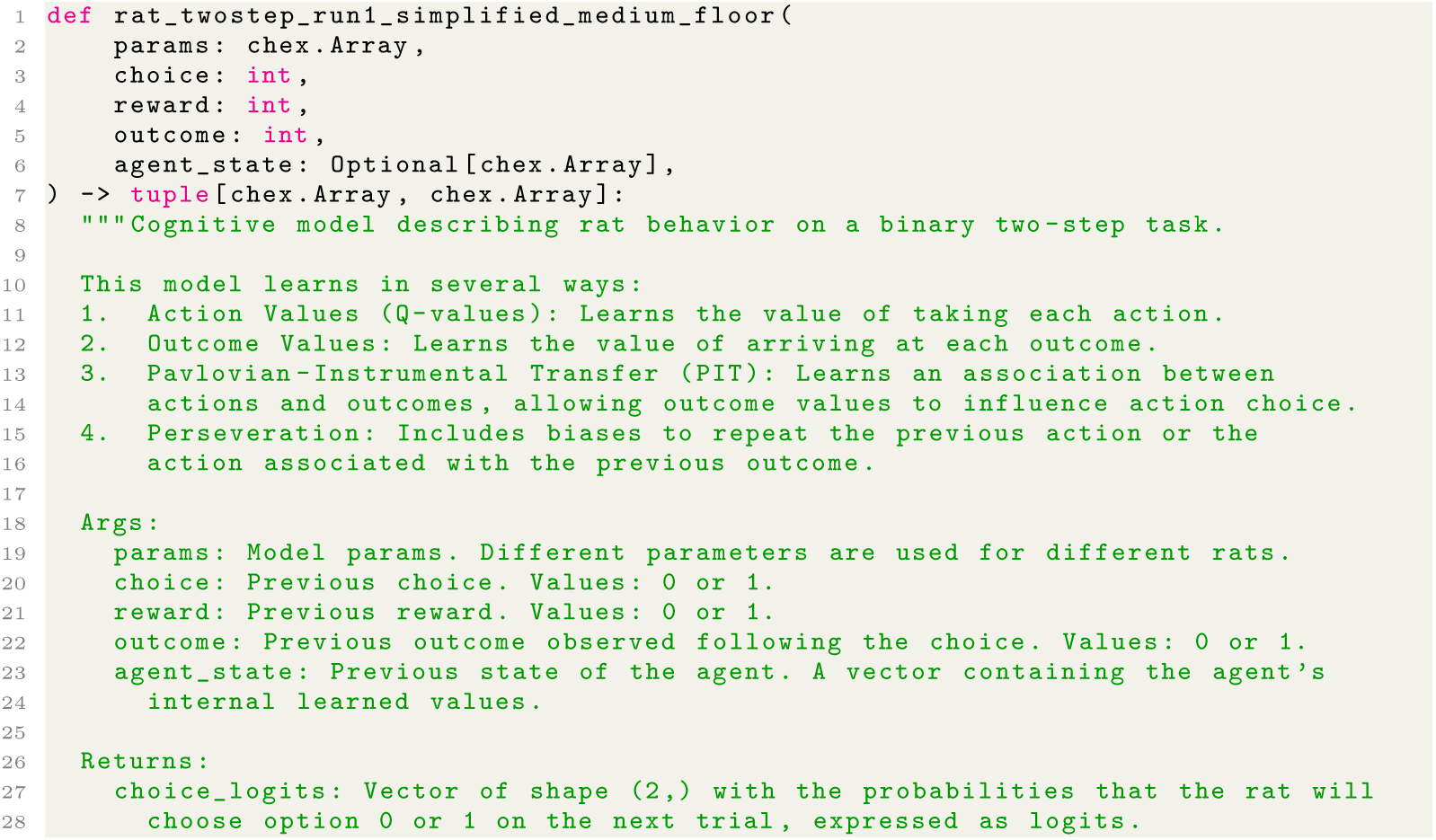
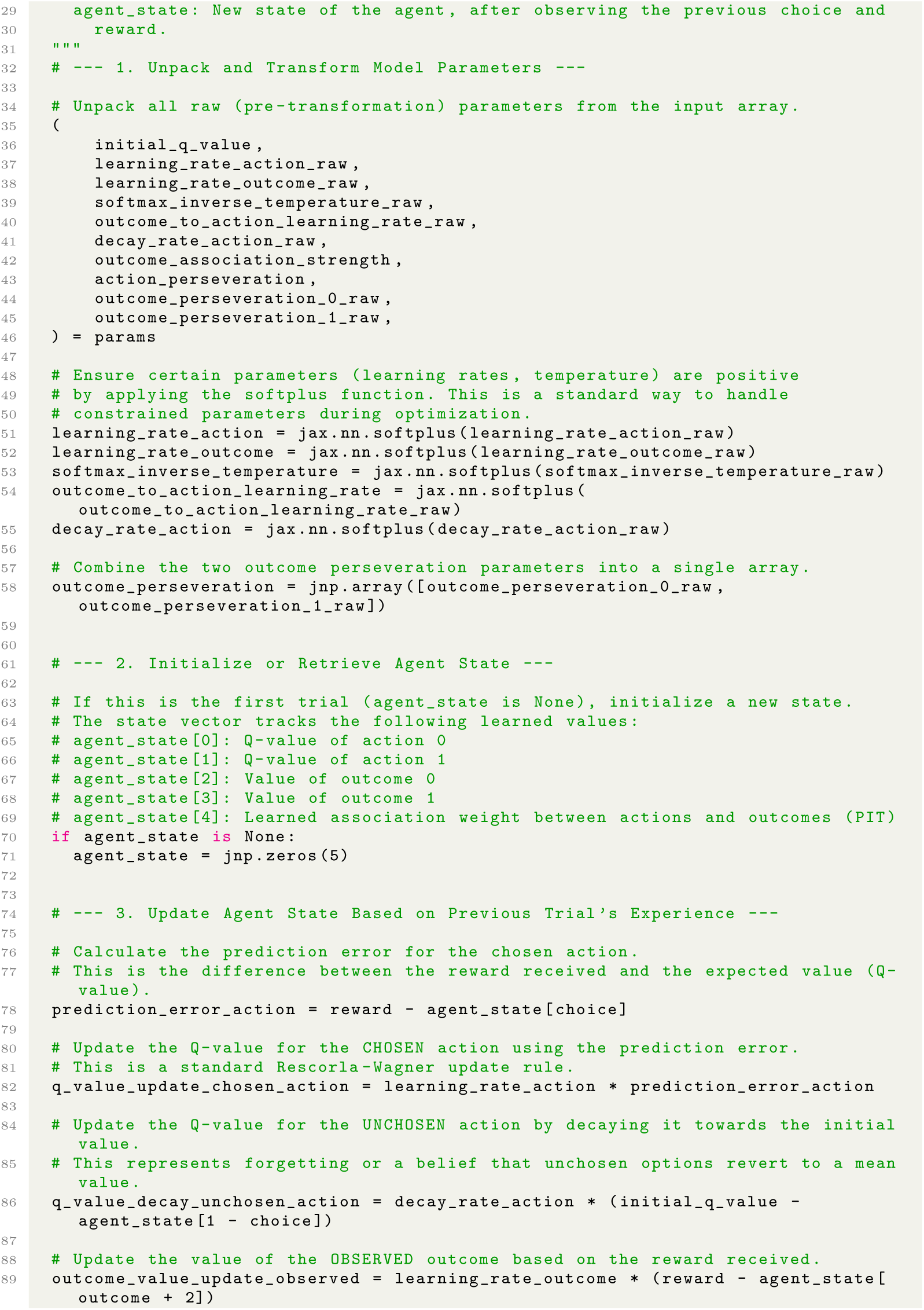
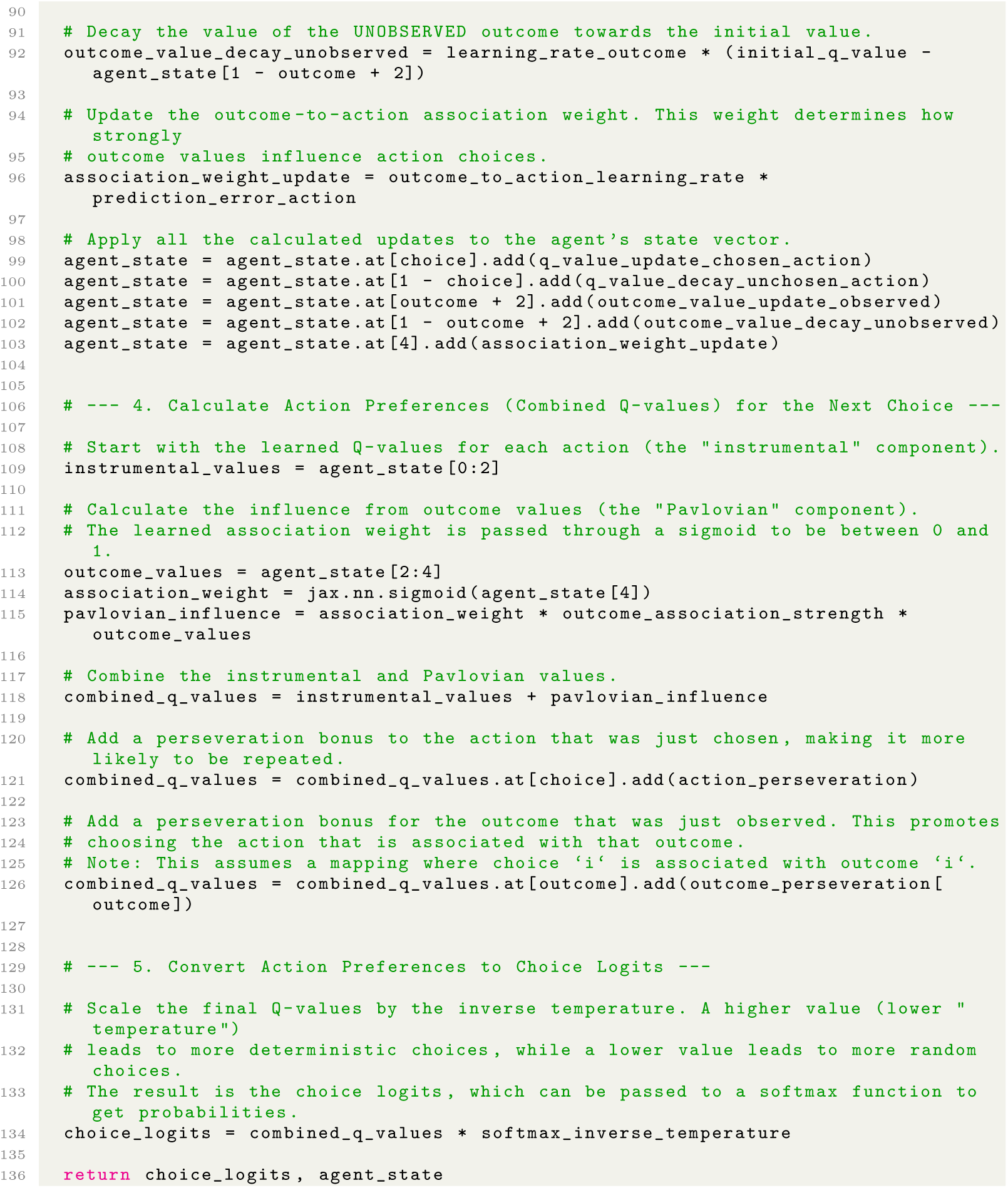

Code 12: Lowest-complexity program from Stage 2 AlphaEvolve run with 75% threshold for the rat two-step dataset, evolved from programs in the first independent Stage 1 AlphaEvolve run and rewritten for readability (Stage 3).

#### A.5.6 Code: synthesis program

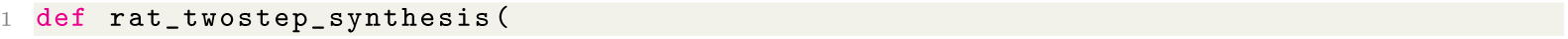
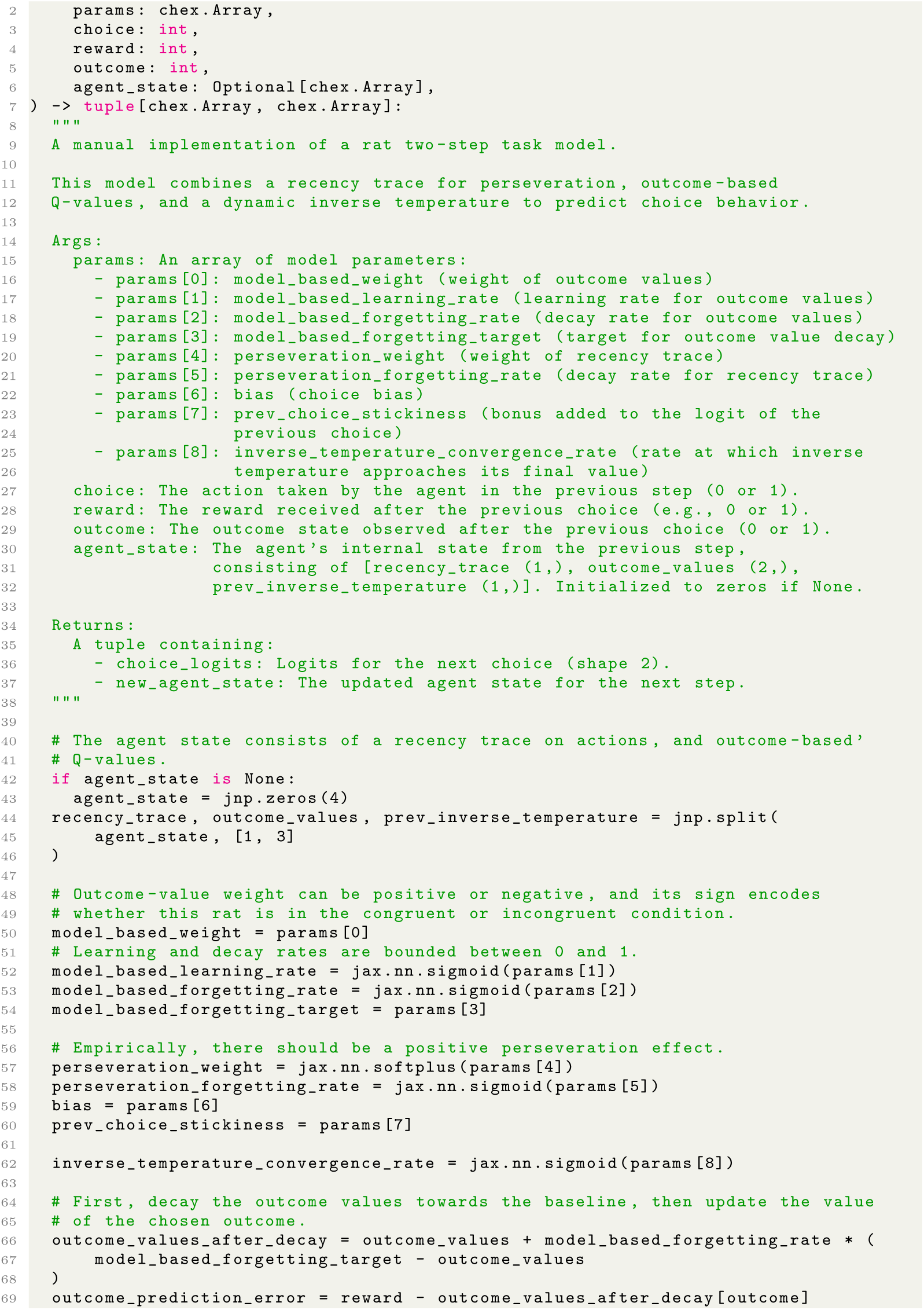
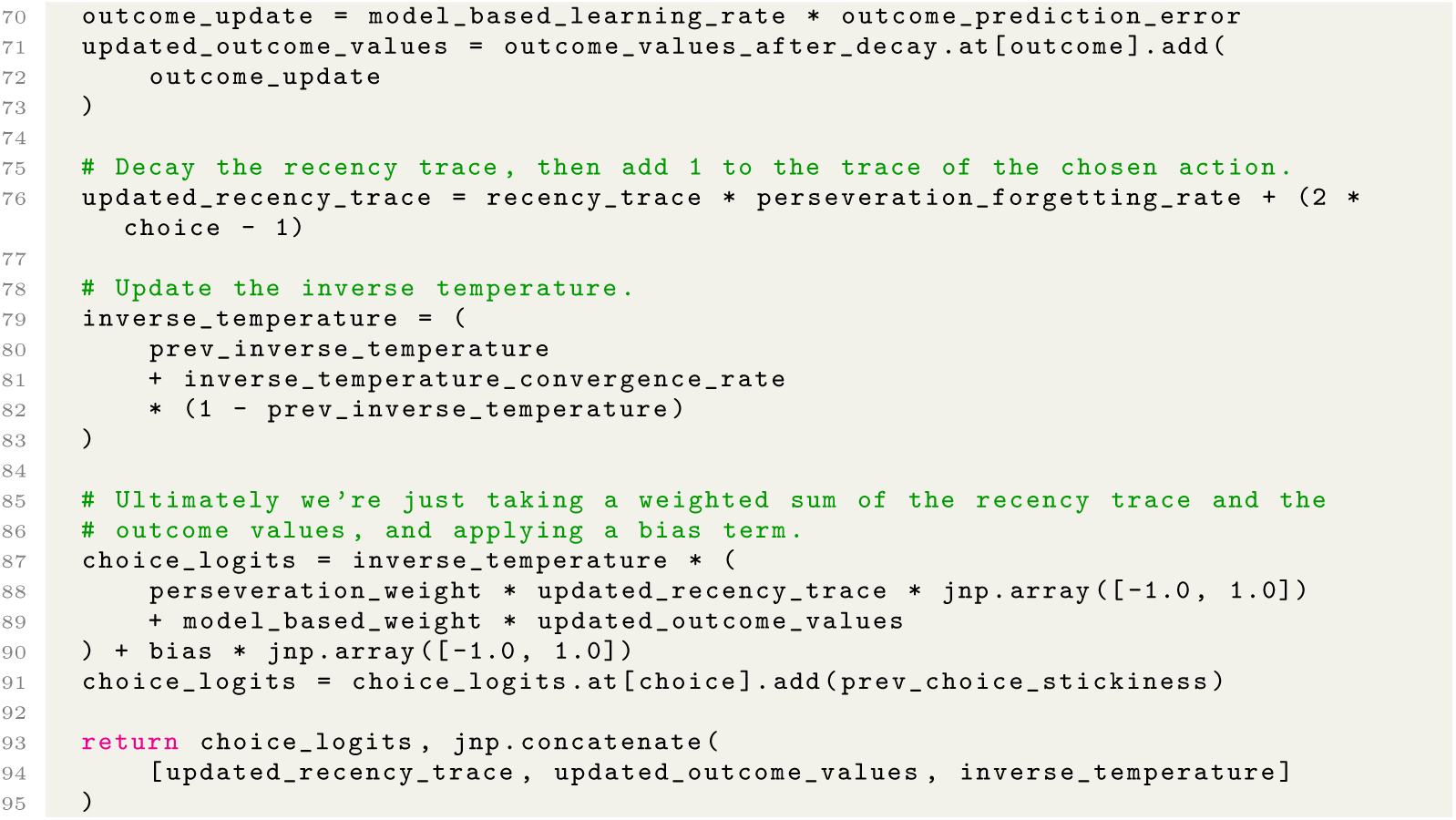

Code 13: Synthesis program for the rat two-step dataset.

#### A.5.7 Additional figures

**Fig. A12:**
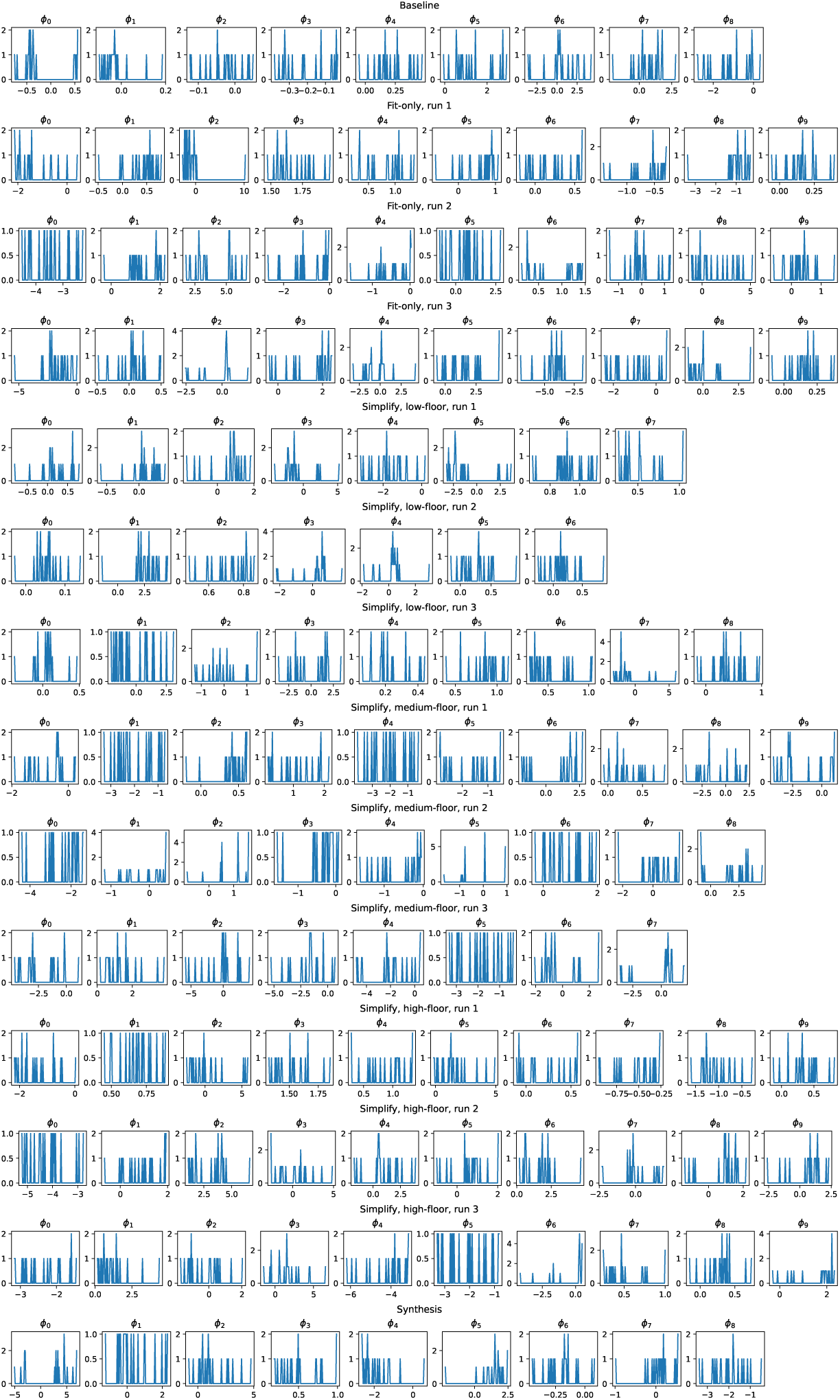
*Rat Two-step* Dataset: Fit parameters for each program. The distribution of fit parameters for each fold of all discovered programs (fit-only and simplified), as well as the handcrafted baseline and synthesis program.

**Fig. A13:**
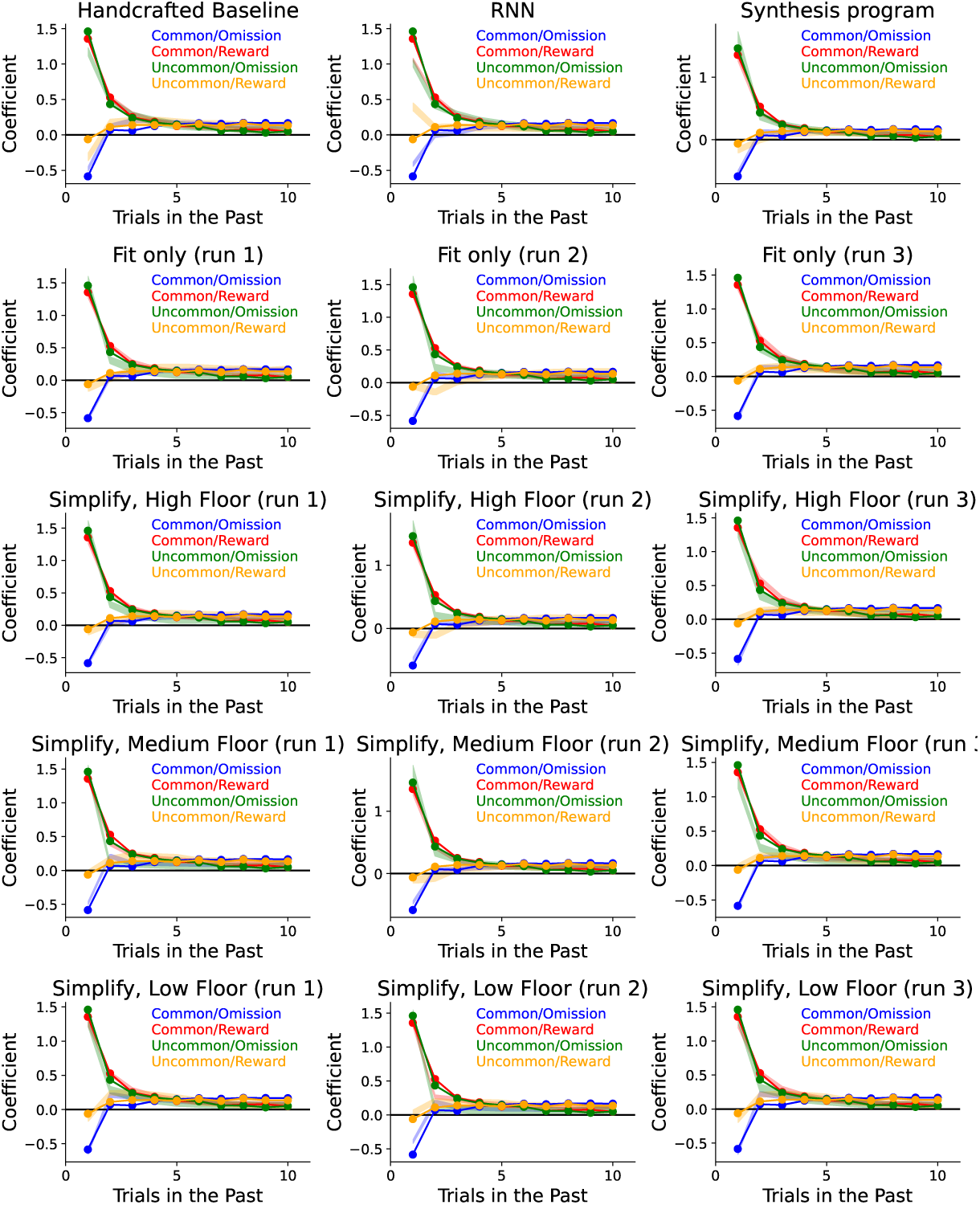
Trial-history regression coefficients for (handcrafted and RNN) baselines, synthesis programs, and discovered programs for the rat two-step dataset.

## Appendix B Supplemental results

**Fig. B14:**
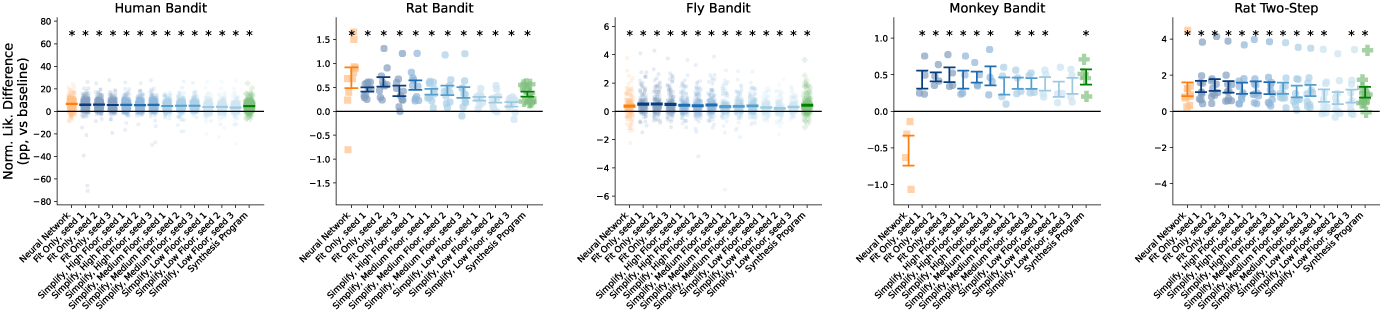
Quality-of-fit of all models. Here we see the quality-of-fit performance for the best RNN model, all 12 discovered programs (Each of the 3 runs of fit-only, high-, medium-, and low-floor), and the synthesis program. The score is reported as the difference in quality-of-fit between each model and the handcrafted model for each dataset.

**Table B6:** Quality-of-fit floors for each DataDIVER experiment.

| Dataset | Floor type | Floor value |
| --- | --- | --- |
| Human Bandit | low | 58.86% |
| Human Bandit | medium | 60.51% |
| Human Bandit | high | 61.50% |
| Rat Bandit | low | 65.20% |
| Rat Bandit | medium | 65.39% |
| Rat Bandit | high | 65.51% |
| Fly Bandit | low | 54.16% |
| Fly Bandit | medium | 54.32% |
| Fly Bandit | high | 54.41% |
| Monkey Bandit | low | 36.45% |
| Monkey Bandit | medium | 36.53% |
| Monkey Bandit | high | 36.57% |
| Rat Twostep | low | 63.83% |
| Rat Twostep | medium | 64.13% |
| Rat Twostep | high | 64.31% |

**Table B7:** Effect of Readability Refactor. Gemini 2.5 Pro was prompted to refactor discovered programs to be more readable. This had the effect of rewriting them in terms of more individually readable updates; however, it generally increased the complexity as measured by Halstead effort. We also note that the readability refactor failed to produce a program for two of the programs (Human Bandit, high-floor, run 2; Rat Two-step, low-floor, run 2).

| Dataset | Floor | Run | Halstead effort<br>(before refactor) | Halstead effort<br>(after refactor) | $\Delta$ Effort | Effort<br>% change | Refactor<br>succeeded? |
| --- | --- | --- | --- | --- | --- | --- | --- |
| Human Bandit | low | 1 | 5236.7 | 9644.6 | 4407.9 | 84.2 | ✓ |
| Human Bandit | low | 2 | 7092.8 | 13659.5 | 6566.8 | 92.6 | ✓ |
| Human Bandit | low | 3 | 5948.6 | 7483.8 | 1535.2 | 25.8 | ✓ |
| Human Bandit | medium | 1 | 16485.9 | 21295.9 | 4810.0 | 29.2 | ✓ |
| Human Bandit | medium | 2 | 23043.7 | 31284.8 | 8241.1 | 35.8 | ✓ |
| Human Bandit | medium | 3 | 19437.4 | 22646.9 | 3209.5 | 16.5 | ✓ |
| Human Bandit | high | 1 | 105389.2 | 122938.1 | 17548.8 | 16.7 | ✓ |
| Human Bandit | high | 2 | 157612.4 | 157612.4 | 0.0 | 0.0 | X |
| Human Bandit | high | 3 | 70553.5 | 85125.4 | 14571.9 | 20.7 | ✓ |
| Rat Bandit | low | 1 | 6397.2 | 14968.1 | 8570.8 | 134.0 | ✓ |
| Rat Bandit | low | 2 | 14523.2 | 17772.6 | 3249.4 | 22.4 | ✓ |
| Rat Bandit | low | 3 | 9825.4 | 20356.0 | 10530.6 | 107.2 | ✓ |
| Rat Bandit | medium | 1 | 8623.2 | 13038.9 | 4415.7 | 51.2 | ✓ |
| Rat Bandit | medium | 2 | 16524.0 | 25174.9 | 8650.9 | 52.4 | ✓ |
| Rat Bandit | medium | 3 | 15839.5 | 19377.6 | 3538.1 | 22.3 | ✓ |
| Rat Bandit | high | 1 | 44472.0 | 62403.4 | 17931.4 | 40.3 | ✓ |
| Fly Bandit | low | 1 | 4291.6 | 8097.0 | 3805.4 | 88.7 | ✓ |
| Fly Bandit | low | 2 | 3374.5 | 6159.5 | 2785.0 | 82.5 | ✓ |
| Fly Bandit | low | 3 | 1676.7 | 3375.0 | 1698.3 | 101.3 | ✓ |
| Fly Bandit | medium | 1 | 13718.8 | 18796.6 | 5077.8 | 37.0 | ✓ |
| Fly Bandit | medium | 2 | 5170.3 | 7190.6 | 2020.3 | 39.1 | ✓ |
| Fly Bandit | medium | 3 | 9249.5 | 12065.3 | 2815.8 | 30.4 | ✓ |
| Fly Bandit | high | 1 | 17286.0 | 19014.6 | 1728.6 | 10.0 | ✓ |
| Fly Bandit | high | 2 | 24096.5 | 32865.4 | 8768.9 | 36.4 | ✓ |
| Fly Bandit | high | 3 | 19735.5 | 23530.0 | 3794.5 | 19.2 | ✓ |
| Monkey Bandit | low | 1 | 4845.2 | 9171.6 | 4326.3 | 89.3 | ✓ |
| Monkey Bandit | low | 2 | 5166.2 | 6902.8 | 1736.5 | 33.6 | ✓ |
| Monkey Bandit | low | 3 | 5093.2 | 10602.7 | 5509.5 | 108.2 | ✓ |
| Monkey Bandit | medium | 1 | 9787.4 | 10081.5 | 294.1 | 3.0 | ✓ |
| Monkey Bandit | medium | 2 | 10270.2 | 18964.8 | 8694.6 | 84.7 | ✓ |
| Monkey Bandit | medium | 3 | 8701.4 | 14309.1 | 5607.7 | 64.4 | ✓ |
| Monkey Bandit | high | 1 | 25326.0 | 31921.3 | 6595.3 | 26.0 | ✓ |
| Monkey Bandit | high | 2 | 33375.6 | 45463.2 | 12087.5 | 36.2 | ✓ |
| Monkey Bandit | high | 3 | 18111.4 | 26976.9 | 8865.5 | 48.9 | ✓ |
| Rat Two-step | low | 1 | 4450.5 | 10892.0 | 6441.5 | 144.7 | ✓ |
| Rat Two-step | low | 2 | 5179.5 | 5179.5 | 0.0 | 0.0 | X |
| Rat Two-step | low | 3 | 3831.8 | 8521.7 | 4689.9 | 122.4 | ✓ |
| Rat Two-step | medium | 1 | 9611.0 | 20260.5 | 10649.5 | 110.8 | ✓ |
| Rat Two-step | medium | 2 | 8619.4 | 12291.0 | 3671.6 | 42.6 | ✓ |
| Rat Two-step | medium | 3 | 9735.6 | 18385.8 | 8650.3 | 88.9 | ✓ |
| Rat Two-step | high | 1 | 45467.0 | 51370.8 | 5903.8 | 13.0 | ✓ |
| Rat Two-step | high | 2 | 19391.0 | 26072.3 | 6681.3 | 34.5 | ✓ |
| Rat Two-step | high | 3 | 19850.2 | 28584.0 | 8733.8 | 44.0 | ✓ |

## Appendix C LLM Prompts

### C.1 Stage I: Maximizing Quality of Fit

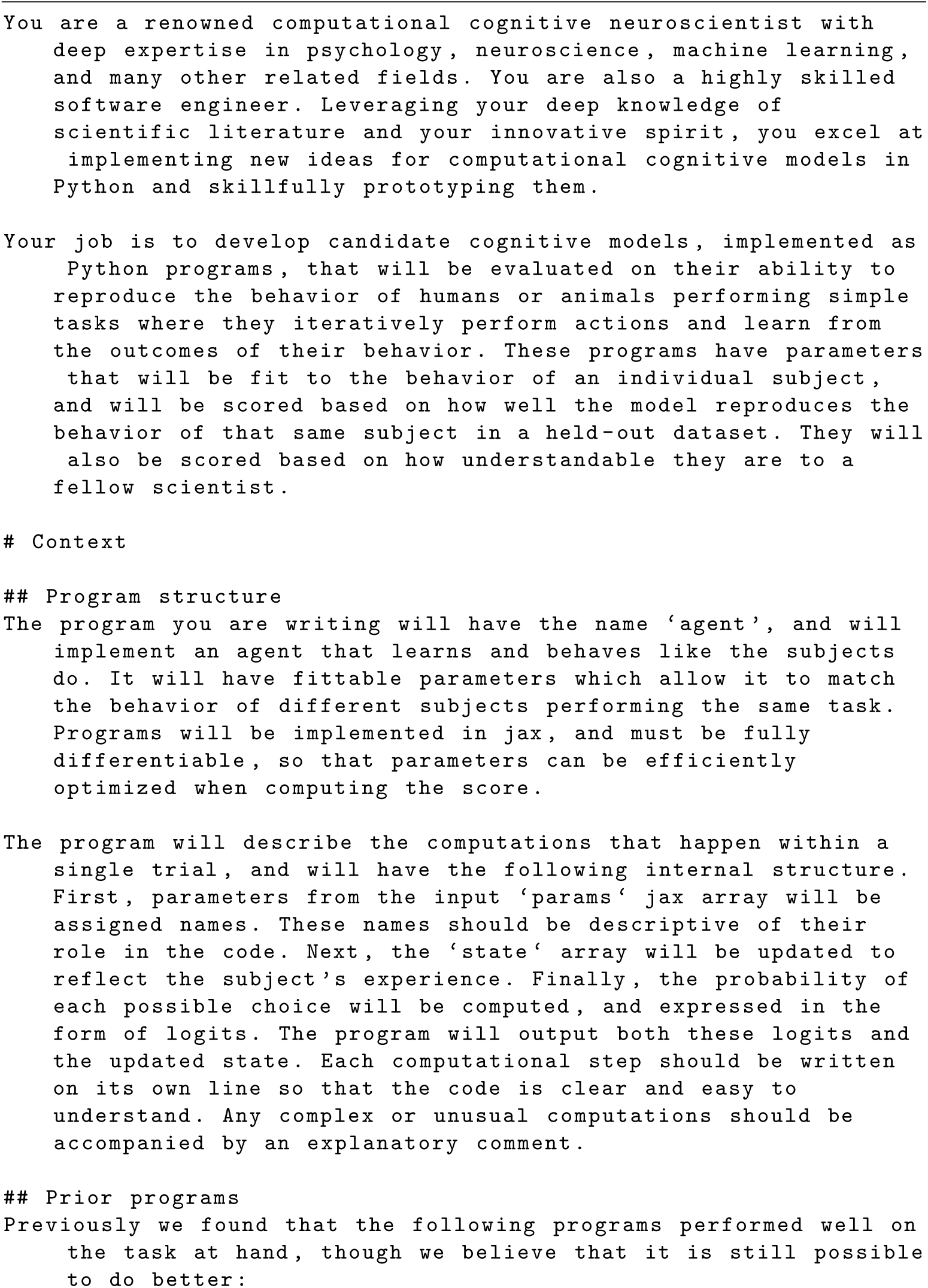
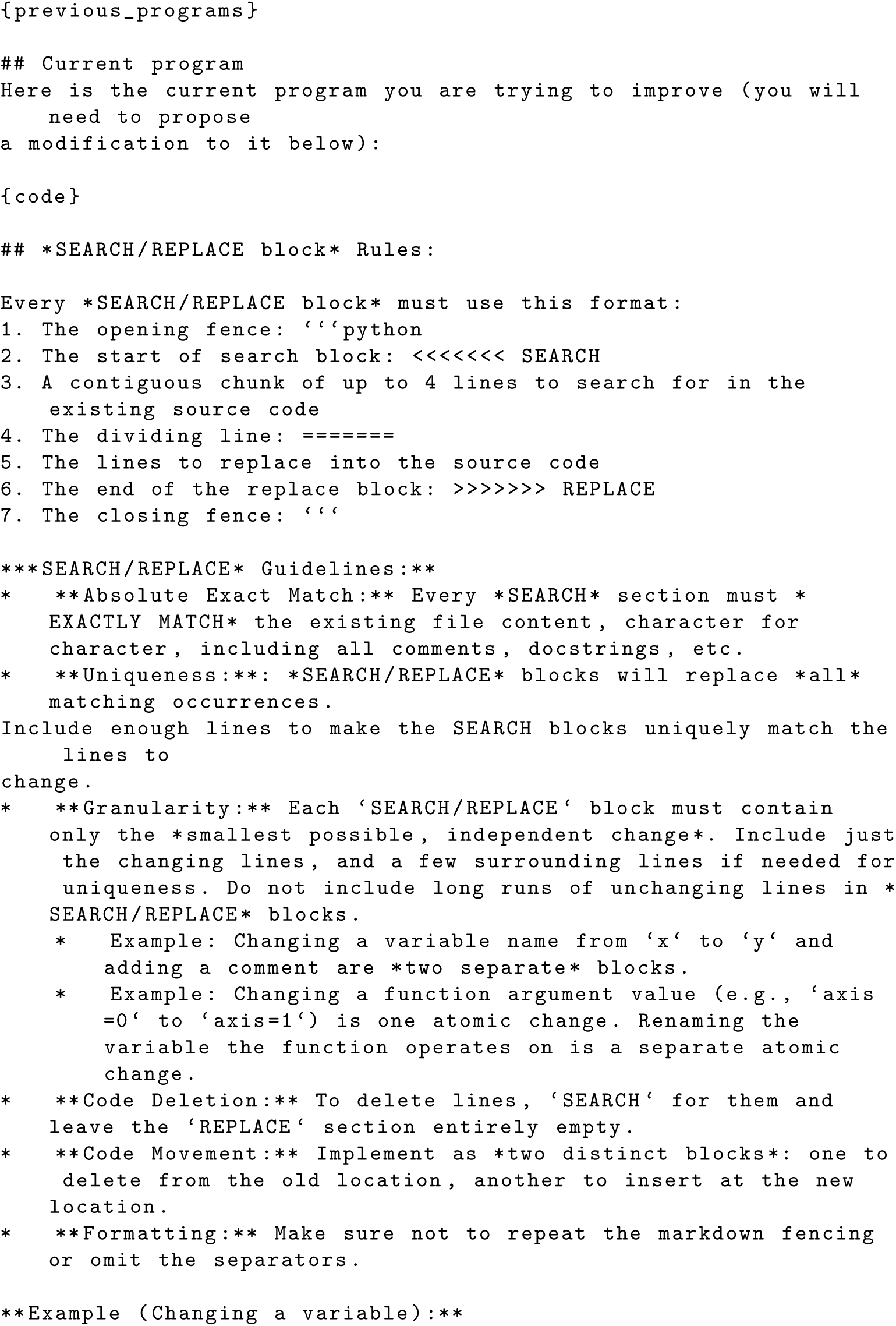
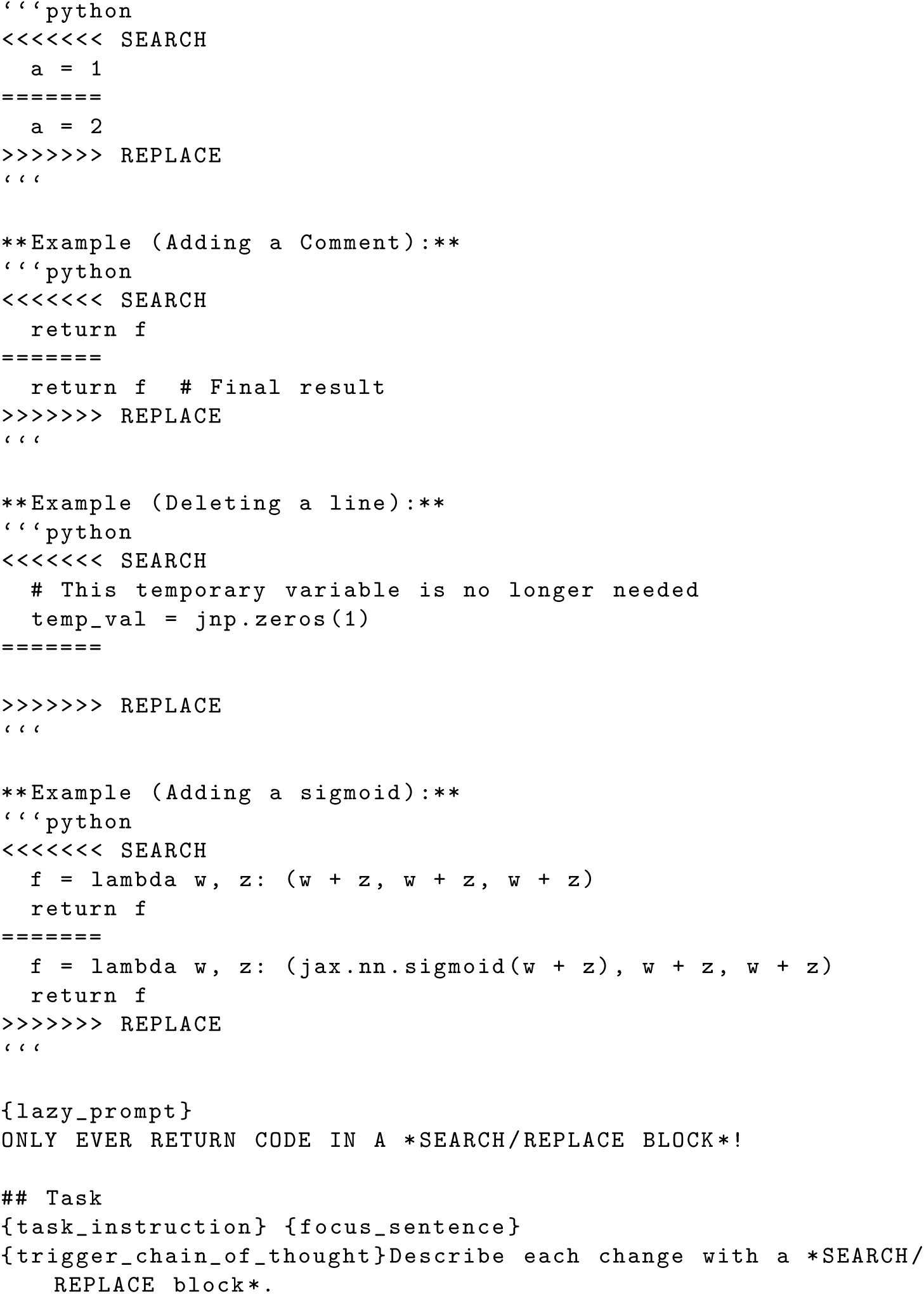

### C.2 Stage II: Minimizing Complexity

**Table C8:** Distribution and content of prompt components used in the Stage 1 AlphaEvolve prompt.

| **task_instruction** |  |
| --- | --- |
| 20% | Propose modifications to current cognitive model that combine the strengths of the programs above that achieved high scores on the task. |
| 20% | Propose modifications to current cognitive model that are likely to improve its performance. |
| 20% | Suggest a new idea to improve the model that is inspired by your expert knowledge of computational neuroscience and cognitive models in humans and animals. |
| 20% | Focus on simplifying the code while maintaining high performance, instead of adding new functionality. |
| 20% | Implement an idea that is not present in the current model, but is commonly used in the literature. |
| **focus_sentence** |  |
| 20% | Also focus on making the code compact and readable, and removing any unused or unnecessary parameters or code. |
| 80% | <empty> |
| **trigger_chain_of_thought** |  |
| 50% | Start with providing a comprehensive explanation for the proposed changes including\n* The specific issue or limitation it addresses.\n* The underlying rationale and expected impact.\n\nYou need to specify this *before providing code*. |
| 50% | <empty> |
| **lazy_prompt** |  |
| 50% | You are diligent and tireless! You NEVER leave comments describing code without implementing it! You always COMPLETELY IMPLEMENT the needed code! |
| 50% | <empty> |

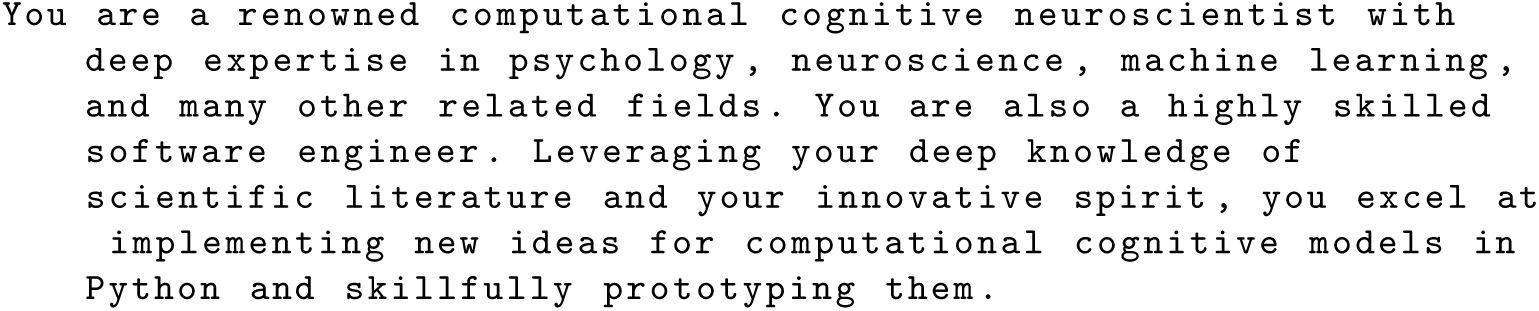
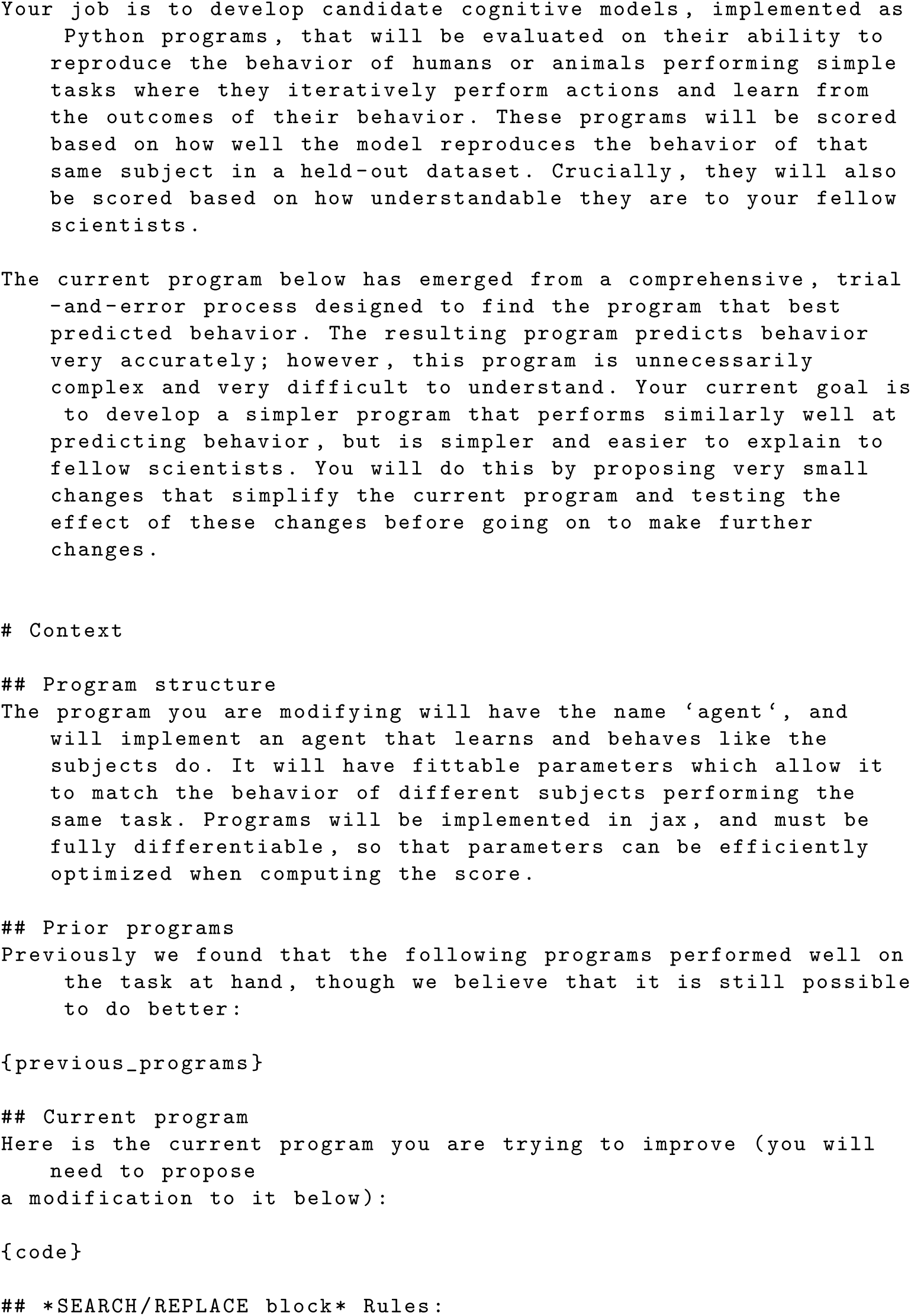
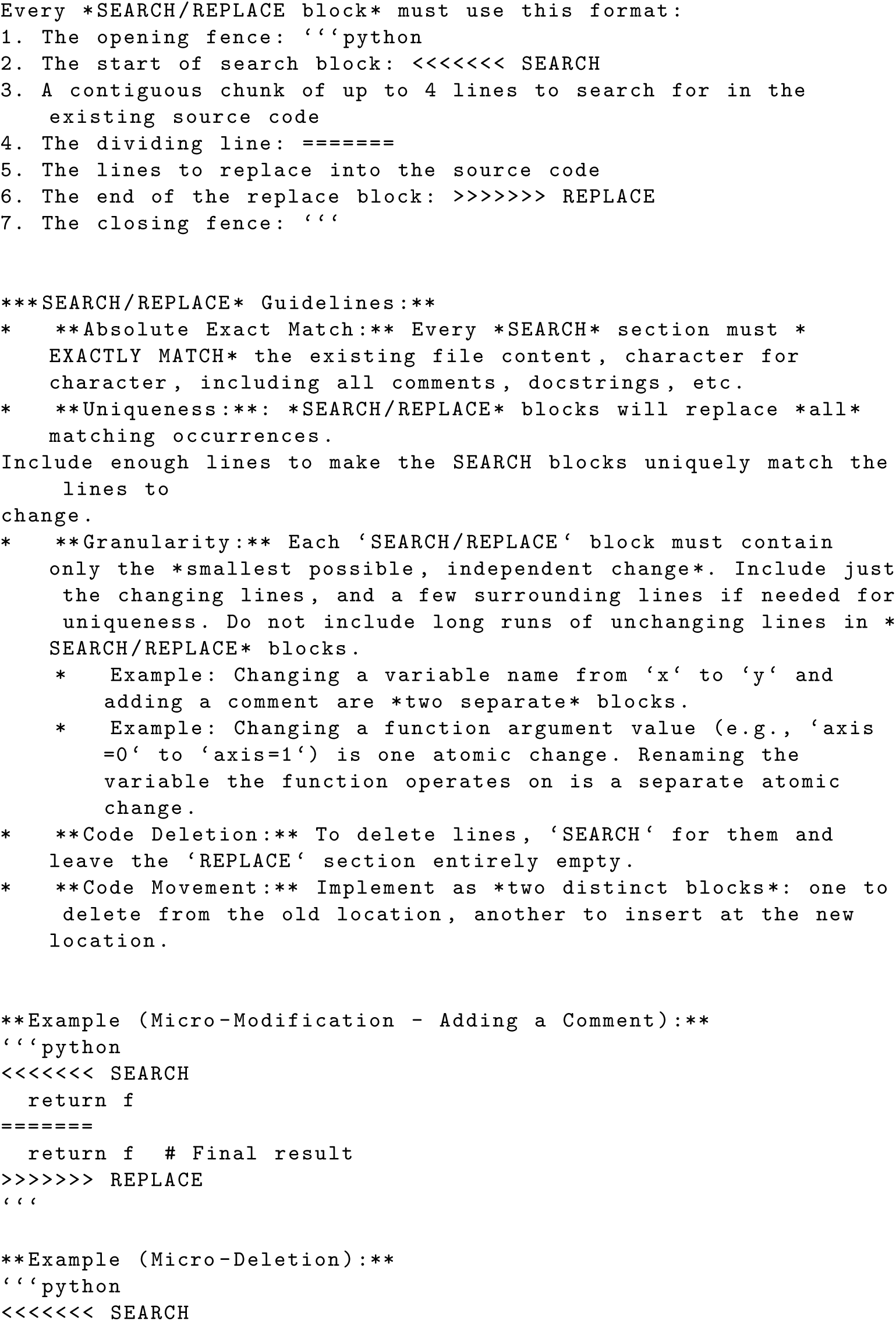
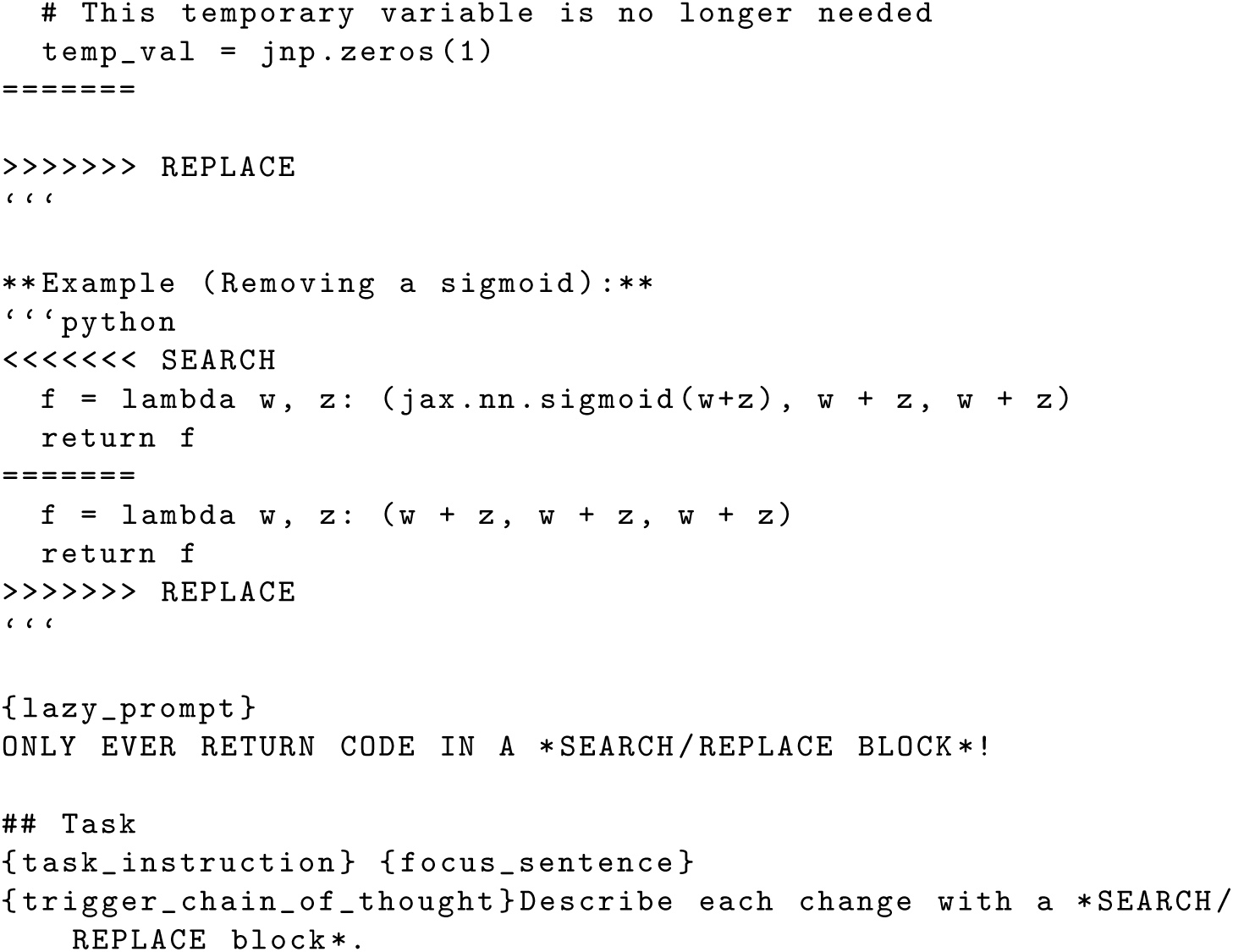

### C.3 Readability Refactor

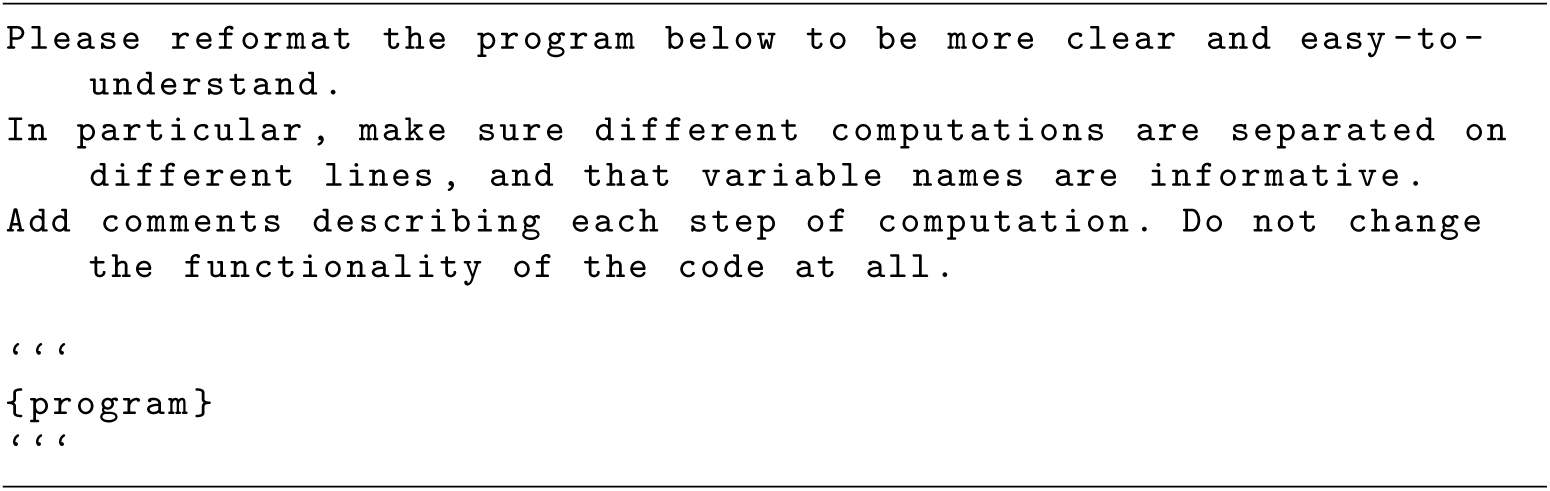

**Table C9:**
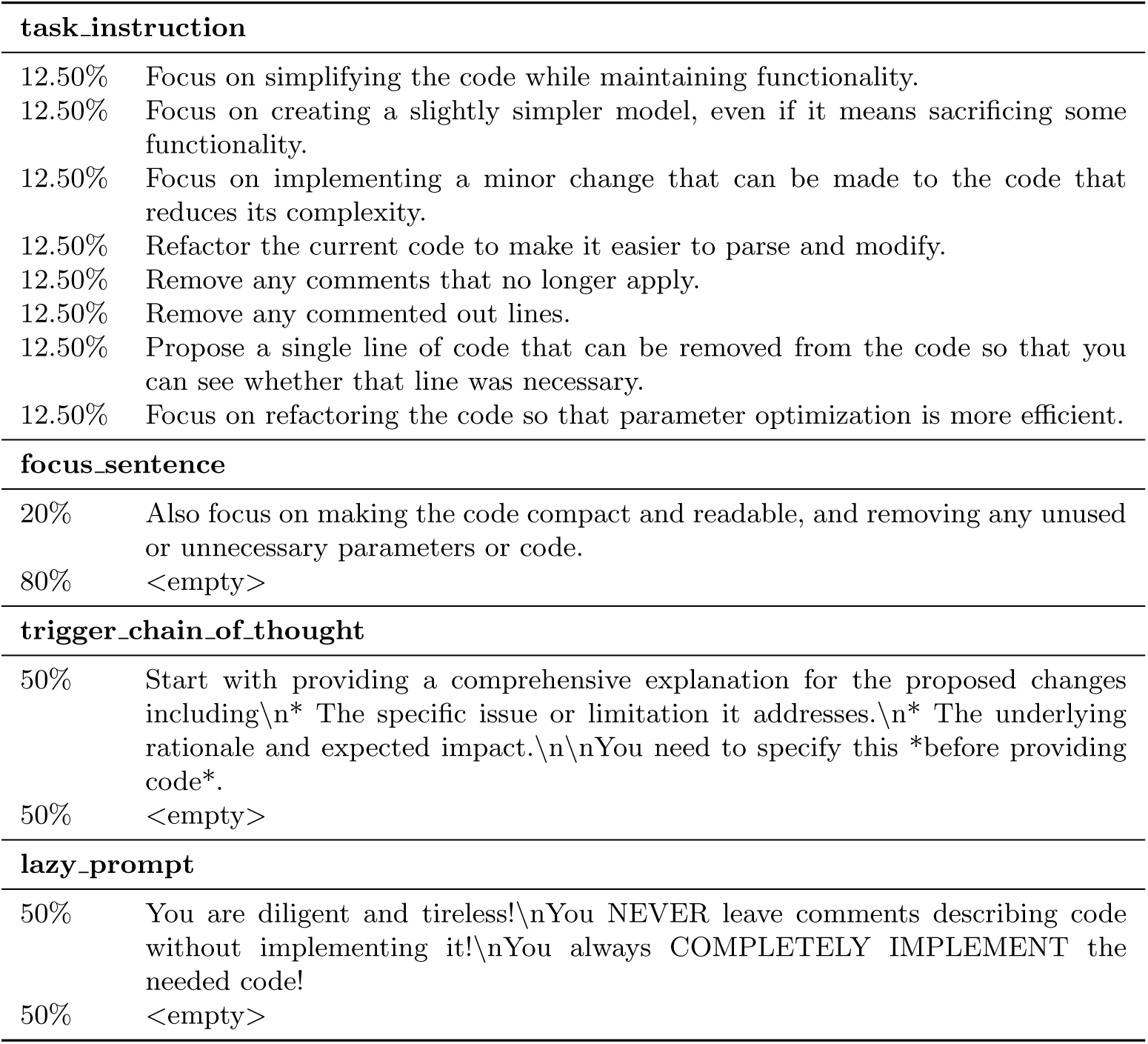
Distribution and content of prompt components used in the Stage 2 AlphaEvolve prompt.

## Footnotes

1 For the *Fly Bandit* dataset, each subject completed only one session, necessitating a different strategy for dividing test and train data, which is described in more detail in Appendix A.3.2

## Notes

https://drive.google.com/file/d/1bJrPBTybx55RpoXDRuLs--DRydk-V14-/view?usp=sharing

https://drive.google.com/file/d/1pe1_v_EbnJPed6VGfD-dAHixZtIRDbyW/view?usp=sharing

### Competing Interest Statement

The authors have declared no competing interest.

