## Supplementary material for "AI-Discovered Cognitive Models Reveal Novel Insights into Human and Animal Learning": DataDIVER Discovered Programs: fly_bandit_models.py

```
# Copyright 2026 DeepMind Technologies Limited.
#
# Licensed under the Apache License, Version 2.0 (the "License");
# you may not use this file except in compliance with the License.
# You may obtain a copy of the License at
#
#     http://www.apache.org/licenses/LICENSE-2.0
#
# Unless required by applicable law or agreed to in writing, software
# distributed under the License is distributed on an "AS IS" BASIS,
# WITHOUT WARRANTIES OR CONDITIONS OF ANY KIND, either express or implied.
# See the License for the specific language governing permissions and
# limitations under the License.

"""Saved models for the Nature 2026 submission."""
from typing import Optional, Tuple

import chex
import jax
import jax.numpy as jnp


def fly_bandit_run1_fit_only(
    params: chex.Array,
    choice: int,
    reward: int,
    agent_state: Optional[chex.Array],
) -> tuple[chex.Array, chex.Array]:
  """Cognitive model describing fly behavior on a binary two-armed bandit task.

  Args:
    params: Fit parameters of the model.
    choice: The choice made by a fly on this trial. 0 or 1
    reward: The reward received by a fly on this trial. 0 or 1
    agent_state: The current state of the cognitive model.

  Returns:
    choice_logits: The probabilities that the fly will choose option 0 or 1 on
      the next trial, expressed as logits.
    agent_state: The updated state of the cognitive model.
  """
  # Do not remove comments or TODOs from this program.

  # Initialize agent state: [Q_0, Q_1, Eligibility_Trace_0, Eligibility_Trace_1, Reward_History, Choice_Count_0, Choice_Count_1]
  if agent_state is None:
    agent_state = jnp.zeros((7,))

  # Define all parameters.
  # Learning rate for positive prediction errors.
  learning_rate_positive = jax.nn.softplus(params[0])
  # Learning rate for negative prediction errors.
  learning_rate_negative = jax.nn.softplus(params[1])
  # Inverse temperature for softmax.
  inverse_temperature = jnp.exp(params[2])
  # Eligibility trace decay rates for each action.
  eligibility_decay_rates = jax.nn.sigmoid(params[jnp.array([3, 4])])
  # Inhibition/facilitation learning rate for the unchosen action.
  inhibition_facilitation_learning_rate = jax.nn.softplus(params[5])
  # Decay rate for unchosen Q-values.
  q_decay_to_baseline_rate = jax.nn.sigmoid(params[6])
  # Baseline preference for choice 0.
  bias = params[7]
  # Decay rate for the recent reward history.
  reward_history_decay = jax.nn.sigmoid(params[8])
  # Strength of modulation of learning rates by recent reward history.
  reward_history_beta = params[9]
  # Eligibility trace boosts for each action.
  eligibility_boosts = jax.nn.softplus(params[jnp.array([10, 11])])

  # Unpack Q-values, eligibility traces, recent reward history, and choice counts from the current state.
  q_values = agent_state[0:2]
  eligibility_traces = agent_state[2:4]
  reward_history = agent_state[4]
  choice_counts = agent_state[5:7]

  # Compute prediction error (delta).
  prediction_error = reward - q_values[choice]

  # Update eligibility traces.
  # Decay and boost eligibility traces.
  # For the chosen action, decay and then boost. For the unchosen, decay and suppress.
  # Modulate eligibility trace decay rates by the absolute prediction error (surprise).
  # Larger absolute prediction error (more surprising outcome) leads to a slower decay,
  # allowing eligibility to persist longer for salient events.
  surprise_factor = 1.0 - jnp.exp(-jnp.abs(prediction_error))
  modulated_eligibility_decay_rates = (
      eligibility_decay_rates * (1.0 - surprise_factor) + 0.99 * surprise_factor
  )

  # Update eligibility traces. For the chosen action, decay and then boost. For the unchosen, decay and suppress.
  eligibility_traces = eligibility_traces * modulated_eligibility_decay_rates
  eligibility_traces = eligibility_traces.at[choice].set(
      eligibility_traces[choice] + eligibility_boosts[choice]
  )
  eligibility_traces = eligibility_traces.at[1 - choice].set(
      eligibility_traces[1 - choice] - eligibility_boosts[1 - choice]
  )

  # Update reward history: a recency-weighted average of past rewards.
  reward_history = (
      reward_history * reward_history_decay
      + reward * (1 - reward_history_decay)
  )
  # Determine the learning rate based on the sign of the prediction error.
  # Ensure eligibility traces remain non-negative.
  eligibility_traces = jnp.maximum(0.0, eligibility_traces)

  # Update choice counts for the chosen action. Add a small epsilon to avoid division by zero.
  choice_counts = choice_counts.at[choice].set(choice_counts[choice] + 1.0)
  # Calculate certainty factor: inversely proportional to the square root of choice count.
  # This makes learning rate higher when experience is low (less certain) and lower when experience is high (more certain).
  certainty_factor = 1.0 / jnp.sqrt(choice_counts[choice] + 1e-6)

  # Learning rate is modulated by a combination of recent reward history, surprise, and certainty.
  learning_rate_modulation_factor = (
      jnp.exp(reward_history_beta * reward_history) *
      jax.nn.sigmoid(jnp.abs(prediction_error) * 5.0 - 1.0) *  # Stronger attentional modulation based on surprise
      certainty_factor
  )

  chosen_learning_rate = jnp.where(
      prediction_error >= 0,
      learning_rate_positive * learning_rate_modulation_factor,
      learning_rate_negative * learning_rate_modulation_factor,
  )

  # The chosen action's Q-value is updated, scaled by its eligibility trace.
  q_values_updated = q_values.at[choice].set(
      q_values[choice] + chosen_learning_rate * prediction_error * eligibility_traces[choice]
  )


  # Index for the unchosen action.
  unchosen_choice = 1 - choice

  # Update the unchosen action's Q-value using decay towards zero and an inhibition/facilitation term.
  q_values_updated = q_values_updated.at[unchosen_choice].set(
      q_values_updated[unchosen_choice] * (1 - q_decay_to_baseline_rate)
      - inhibition_facilitation_learning_rate
      * (reward - 0.5)
      * eligibility_traces[unchosen_choice]
      * eligibility_traces[choice]
      * jnp.exp(reward_history_beta * reward_history)
      * (1 - jnp.tanh(jnp.abs(q_values[choice])))
  )

  # Combine updated Q-values, eligibility traces, reward history, and choice counts into the new agent state.
  agent_state = jnp.concatenate(
      (q_values_updated, eligibility_traces, jnp.array([reward_history]), choice_counts)
  )

  # Confidence-weighted inverse temperature.
  # Modulate inverse temperature based on the confidence in the chosen action.
  # Higher Q-value for the chosen action relative to the unchosen implies higher confidence,
  # leading to increased exploitation (higher effective inverse temperature).
  # When Q-values are close or low, uncertainty is high, leading to more exploration (lower effective inverse temperature).
  # Modulate inverse temperature based on the confidence in the chosen action
  # and the overall reward history.
  # Higher Q-value for the chosen action relative to the unchosen implies higher confidence.
  # Scale of confidence, 0 to 1.
  min_q_value = jnp.min(q_values_updated)
  confidence_in_chosen = jax.nn.sigmoid(q_values_updated[choice] - min_q_value)

  # Dynamic bounds for inverse temperature, influenced by recent reward history.
  # When reward history is high, the agent is more likely to exploit (higher base temperature).
  # When reward history is low, the agent is more likely to explore (lower base temperature).
  base_inverse_temperature = inverse_temperature * (1 + reward_history_beta * (reward_history - 0.5))
  lower_temp_bound = base_inverse_temperature * 0.5  # More exploratory baseline
  upper_temp_bound = base_inverse_temperature * 1.5  # More exploitative baseline

  # Adaptively set the effective inverse temperature using a sigmoidal scaling of confidence.
  # This allows for a more nuanced and potentially non-linear transition from exploration to exploitation.
  # A higher, sigmoidally scaled confidence pushes the temperature towards the upper bound (exploitation),
  # while lower confidence pushes it towards the lower bound (exploration).
  sigmoidal_confidence_scaling = jax.nn.sigmoid(confidence_in_chosen * 5.0 - 2.5) # Scale confidence to span a sigmoid
  
  modulated_inverse_temperature = (
      lower_temp_bound * (1.0 - sigmoidal_confidence_scaling)
      + upper_temp_bound * sigmoidal_confidence_scaling
  )
  # Ensure the modulated inverse temperature remains positive.
  modulated_inverse_temperature = jnp.maximum(1e-6, modulated_inverse_temperature)

  # Compute choice_logits using a softmax over Q-values, incorporating a bias for choice 0.
  choice_logits = jax.nn.softplus(modulated_inverse_temperature * q_values_updated + jnp.array([bias, 0.0]))

  return choice_logits, agent_state
  

def fly_bandit_run2_fit_only(
    params: chex.Array,
    choice: int,
    reward: int,
    agent_state: Optional[chex.Array],
) -> tuple[chex.Array, chex.Array]:
  """Cognitive model describing fly behavior on a binary two-armed bandit task.

  Args:
    params: Fit parameters of the model.
    choice: The choice made by a fly on this trial. 0 or 1
    reward: The reward received by a fly on this trial. 0 or 1
    agent_state: The current state of the cognitive model.

  Returns:
    choice_logits: The probabilities that the fly will choose option 0 or 1 on
      the next trial, expressed as logits.
    agent_state: The updated state of the cognitive model.
  """
  # Do not remove comments or TODOs from this program.

  # TODO: Initialize agent state. NOLINT
  if agent_state is None:
    # Initialize value estimates for choices (Q0, Q1), choice stickiness (S0, S1),
    # a reward reference point (RRP), and value uncertainty (VU).
    # Q0, Q1, S0, S1, RRP, VU0, VU1 (Value Uncertainty for each choice)
    # Initialize Q-values and Value Uncertainties to a small positive value to encourage initial exploration
    agent_state = jnp.array([0.1, 0.1, 0.0, 0.0, 0.0, 0.1, 0.1])

  # TODO: Define all parameters. NOLINT
  # Each parameter assignment should be a single line.
  # Change the names of the parameters to be descriptive.
  # Include comments describing the parameter transformation, if applicable.
  learning_rate = jax.nn.softplus(params[0])  # Learning rate (alpha) for updating value estimates.
  static_inverse_temperature = jnp.exp(params[1])  # Static component of inverse temperature (beta).
  uncertainty_weight = jnp.exp(params[2]) # Weight for scaling the dynamic inverse temperature component. # Renamed from `uncertainty_scale` for clarity.
  # Decay rate for choice perseveration, transformed with sigmoid.
  choice_perseveration_decay = jax.nn.sigmoid(params[7])
  forgetting_rate = jax.nn.sigmoid(params[3])  # Forgetting rate for decaying unchosen values.
  previous_choice_bias = jnp.exp(params[4])  # Bias to repeat the previous choice.
  choice_stickiness_lr = jax.nn.sigmoid(params[5])  # Learning rate for choice stickiness.
  # Influence of outcome valence on choice stickiness.
  outcome_valence_stickiness_weight = params[6]
  # Learning rate for adapting the reward reference point.
  reward_reference_learning_rate = jax.nn.sigmoid(params[8])
  uncertainty_learning_rate = jax.nn.sigmoid(params[9])  # Learning rate for adapting the value uncertainty.

  # Update agent_state
  # Retrieve the current reward reference point and per-choice uncertainties
  reward_reference_point = agent_state[4]
  # Update the value of the chosen option using a Rescorla-Wagner rule, now relative to the reward reference point
  raw_prediction_error = reward - reward_reference_point
  # Dynamically scale the prediction error based on the reward reference point.
  # If the reward reference point is high, scale down the RPE (less sensitive to individual outcomes).
  # If the reward reference point is low, scale up the RPE (more sensitive to individual outcomes).
  # Add a small constant to the denominator to prevent division by zero and ensure stability.
  rpe_scaling_factor = 1.0 / (1.0 + jnp.abs(reward_reference_point))
  prediction_error = raw_prediction_error * rpe_scaling_factor
  # Dynamically adjust forgetting rate for the unchosen option based on the absolute prediction error.
  # Larger absolute prediction error leads to more forgetting of the unchosen option.
  contextual_forgetting_rate = forgetting_rate * (1 + jnp.abs(raw_prediction_error)) # Use raw RPE for forgetting rate, as it's a measure of surprise.
  # Dynamically adjust the decay/update for the unchosen option based on the reward received.
  # If a reward is received from the chosen option, penalize the unchosen option's value.
  # If no reward is received, the unchosen option's value decays less, or even slightly increases,
  # as it becomes relatively more attractive compared to the unrewarded chosen option.
  unchosen_value_update_factor = jnp.where(reward == 1, 1 + learning_rate, 1 - learning_rate * 0.5) # Penalize if rewarded, less decay if unrewarded
  value_uncertainty_chosen = agent_state[5 + choice] # Get uncertainty for the chosen option
  # Adaptive forgetting: forget unchosen option more if confident in chosen option.
  # If value_uncertainty_chosen is low (high confidence), the decay factor approaches 1, leading to faster decay.
  # If value_uncertainty_chosen is high (low confidence), the decay factor approaches 0, leading to slower decay.
  adaptive_decay_factor = jax.nn.sigmoid(1 - value_uncertainty_chosen)
  # Decay the unchosen option's value towards zero (a neutral baseline).
  # The decay rate is modulated by contextual_forgetting_rate and adaptive_decay_factor,
  # and also influenced by unchosen_value_update_factor.
  agent_state = agent_state.at[1 - choice].set(
      agent_state[1 - choice] * (1 - contextual_forgetting_rate * adaptive_decay_factor)
      + (0.0 - agent_state[1 - choice]) * contextual_forgetting_rate * adaptive_decay_factor * 0.1 # Decay towards zero baseline
  )
  agent_state = agent_state.at[1 - choice].set(agent_state[1 - choice] * unchosen_value_update_factor) # Apply penalty/boost after decay
  # Adjust the learning rate for Q-value updates based on both the absolute prediction error
  # and the current uncertainty of the chosen option.
  # Larger absolute prediction error or lower uncertainty leads to a larger effective learning rate,
  # promoting faster updates when outcomes are surprising or beliefs are confident.
  # Separate prediction error into positive and negative components to modulate learning asymmetrically.
  positive_prediction_error = jnp.maximum(0.0, prediction_error)
  negative_prediction_error = jnp.minimum(0.0, prediction_error)

  # Modulate learning rate based on positive and negative surprise.
  # Positive surprises (better than expected) and negative surprises (worse than expected)
  # can have different impacts on the learning rate, reflecting distinct neural pathways.
  # Here, we allow for potentially different sensitivities to unexpected gains vs. losses.
  # A non-linear scaling (e.g., softplus) can make the learning rate more responsive to larger surprises.
  effective_learning_rate = learning_rate * (
      1.0
      + jax.nn.softplus(positive_prediction_error * 2.0)  # Stronger learning for positive surprises
      + jax.nn.softplus(-negative_prediction_error * 1.0) # Potentially different sensitivity for negative surprises
  ) / (1.0 + value_uncertainty_chosen) # Still modulated by uncertainty.
  agent_state = agent_state.at[choice].set(
      agent_state[choice] + effective_learning_rate * prediction_error
  )
  # Apply a small, constant decay to the chosen option's Q-value to prevent over-stabilization.
  agent_state = agent_state.at[choice].set(agent_state[choice] * (1 - forgetting_rate * 0.1))
  # Update the reward reference point based on the experienced reward.
  # Apply a stronger decay towards zero if no reward was received,
  # This makes the reference point more responsive to unrewarded trials.
  # allowing the reference point to reset in unrewarding contexts.
  reference_point_decay_strength = jnp.where(reward == 0, 0.5, 0.1) # Stronger decay if no reward
  agent_state = agent_state.at[4].set(
      reward_reference_point * (1 - reward_reference_learning_rate * reference_point_decay_strength)
      + reward_reference_learning_rate * prediction_error
      - reward_reference_learning_rate * 0.1 * reward_reference_point # Proportional decay towards zero.
      - reward_reference_learning_rate * 0.05 * agent_state[choice] # Decay based on chosen Q-value.
      - reward_reference_learning_rate * 0.01 * reward_reference_point # Constant passive decay towards zero.
  )
  # Update value uncertainty based on reward prediction surprise.
  # This update reflects how surprising the observed reward is, given the expected probability of reward.
  # Approximate the expected probability of reward using a sigmoid of the chosen Q-value.
  expected_reward_probability = jax.nn.sigmoid(agent_state[choice])
  # Compute the surprise (negative log-likelihood) of the observed reward.
  # This measures how unexpected the actual reward was, given the model's prediction.
  reward_surprise = - (reward * jnp.log(expected_reward_probability + 1e-6) + (1 - reward) * jnp.log(1 - expected_reward_probability + 1e-6))
  # Update uncertainty. It increases with surprise and decays over time, bounded between 0 and 1.
  # Adjust uncertainty learning rate based on reward reference point, allowing for more sensitive updates.
  dynamic_uncertainty_lr = uncertainty_learning_rate * (1 + jnp.abs(reward_reference_point))
  # Update uncertainty for the chosen option: increases with surprise, decays over time, bounded.
  new_chosen_uncertainty = value_uncertainty_chosen + dynamic_uncertainty_lr * reward_surprise * (1 - value_uncertainty_chosen)
  # Apply a general decay to uncertainty for the chosen option to allow it to decrease over time.
  new_chosen_uncertainty = new_chosen_uncertainty * (1 - uncertainty_learning_rate * 0.05)
  agent_state = agent_state.at[5 + choice].set(new_chosen_uncertainty)

  # Update uncertainty for the unchosen option: it increases if the chosen option's uncertainty decreased.
  # This reflects that knowledge about the unchosen option remains static or becomes relatively less certain
  # as information about the chosen option solidifies.
  uncertainty_decrease_chosen = value_uncertainty_chosen - new_chosen_uncertainty
  unchosen_uncertainty = agent_state[5 + (1 - choice)]
  # Increase unchosen uncertainty proportional to the decrease in chosen uncertainty, ensuring it stays bounded.
  new_unchosen_uncertainty = unchosen_uncertainty + uncertainty_decrease_chosen * uncertainty_learning_rate * 0.5 * (1 - unchosen_uncertainty)
  agent_state = agent_state.at[5 + (1 - choice)].set(new_unchosen_uncertainty)
  # Update choice stickiness values (choice history).
  # Apply general decay to all stickiness values to prevent indefinite accumulation.
  # The decay rate is higher for the unchosen option, reflecting faster forgetting of unreinforced choices.
  decayed_stickiness_chosen = agent_state[choice + 2] * (1 - choice_perseveration_decay * 0.5)
  decayed_stickiness_unchosen = agent_state[1 - choice + 2] * (1 - choice_perseveration_decay)
  # Update chosen stickiness: it increases, modulated by reward and RPE.
  # The (1 - decayed_stickiness_chosen) term keeps stickiness bounded between 0 and 1.
  new_chosen_stickiness = decayed_stickiness_chosen + choice_stickiness_lr * (
      1 + outcome_valence_stickiness_weight * reward + jnp.abs(prediction_error)
  ) * (1 - decayed_stickiness_chosen)
  agent_state = agent_state.at[choice + 2].set(new_chosen_stickiness)
  agent_state = agent_state.at[1 - choice + 2].set(decayed_stickiness_unchosen)

  # Compute choice_logits.
  # Retrieve the current value estimates for choices 0 and 1.
  q_values = agent_state[:2]
  # Add stickiness bias to q_values.
  q_values = q_values + agent_state[2:4] * previous_choice_bias
  # Retrieve per-choice uncertainties.
  learned_uncertainty_0 = agent_state[5]
  learned_uncertainty_1 = agent_state[6]
  # The inverse temperature combines a static base and a dynamic component.
  # The dynamic component promotes exploration when average uncertainty is high
  # and exploitation when Q-value or uncertainty differences are large.
  average_learned_uncertainty = (learned_uncertainty_0 + learned_uncertainty_1) / 2.0
  # Dynamic inverse temperature increases with Q-value difference and uncertainty difference (exploitation),
  # and decreases with average uncertainty (exploration).
  effective_inverse_temperature = static_inverse_temperature + (
      jnp.abs(q_values[0] - q_values[1])
      + jnp.abs(learned_uncertainty_0 - learned_uncertainty_1)
      - uncertainty_weight * jax.nn.sigmoid(average_learned_uncertainty)
  )
  # Ensure inverse temperature is always positive to prevent undefined behavior.
  effective_inverse_temperature = jnp.maximum(0.1, effective_inverse_temperature)
  choice_logits = q_values * effective_inverse_temperature

  return choice_logits, agent_state
  

def fly_bandit_run3_fit_only(
    params: chex.Array,
    choice: int,
    reward: int,
    agent_state: Optional[chex.Array],
) -> tuple[chex.Array, chex.Array]:
  """Cognitive model describing fly behavior on a binary two-armed bandit task.

  Args:
    params: Fit parameters of the model.
    choice: The choice made by a fly on this trial. 0 or 1
    reward: The reward received by a fly on this trial. 0 or 1
    agent_state: The current state of the cognitive model.

  Returns:
    choice_logits: The probabilities that the fly will choose option 0 or 1 on
      the next trial, expressed as logits.
    agent_state: The updated state of the cognitive model.
  """
  # Do not remove comments or TODOs from this program.

  if agent_state is None:
    # Initialize action values (Q-values) for choice 0 and 1.
    # Eligibility traces are initialized to 0.0, indicating no prior experience.
    initial_eligibility_trace = 0.0
    # Q_0, Q_1, ET_0, ET_1, Count_0, Count_1, Global_Surprise, Surprise_0, Surprise_1
    # Initialize choice counts to 0.0 to reflect no prior choices.
    # Initialize global and action-specific surprise to 0.0, reflecting no initial volatility.
    agent_state = jnp.array([0., 0., initial_eligibility_trace, initial_eligibility_trace, 0., 0., 0., 0., 0.])

  base_learning_rate = jax.nn.sigmoid(params[0]) # A single base learning rate.
  fixed_decay_rate = jax.nn.sigmoid(params[1]) # Fixed decay rate for unchosen actions.
  inverse_temperature = jnp.exp(params[2])
  eligibility_trace_decay = jax.nn.sigmoid(params[3]) # Decay rate for eligibility traces.
  eligibility_trace_factor = jax.nn.sigmoid(params[4]) # Factor for incrementing eligibility traces.
  surprise_reset_strength = jax.nn.sigmoid(params[5]) # Strength of surprise-driven eligibility trace reset for unchosen actions.
  chosen_action_decay_rate = jax.nn.sigmoid(params[6]) # Decay rate for chosen actions.
  eligibility_trace_baseline = jax.nn.sigmoid(params[7]) # Baseline strength of the eligibility trace.
  surprise_modulated_decay_strength = jax.nn.sigmoid(params[8]) # Strength of surprise-driven modulation of unchosen action decay.
  volatility_learning_rate = jax.nn.sigmoid(params[9]) # Rate at which global surprise is updated.

  # Calculate reward prediction error.
  q_values = agent_state[0:2]
  eligibility_traces = agent_state[2:4]
  choice_counts = agent_state[4:6] # Retrieve choice counts
  global_surprise = agent_state[6] # Retrieve global surprise
  action_surprise = agent_state[7:9] # Retrieve action-specific surprise values

  prediction_error = reward - q_values[choice]
  absolute_prediction_error = jnp.abs(prediction_error)

  # Update global surprise: A moving average of squared prediction errors.
  # This serves as a proxy for environmental volatility, with higher sensitivity to large deviations.
  global_surprise = (1 - volatility_learning_rate) * global_surprise + volatility_learning_rate * jnp.square(prediction_error)

  # Update action-specific surprise for the chosen action.
  # This tracks the magnitude of prediction error specifically for this action.
  action_surprise = action_surprise.at[choice].set(
      (1 - volatility_learning_rate) * action_surprise[choice] + volatility_learning_rate * jnp.square(prediction_error)
  )
  # When the chosen action yields a high prediction error, it also influences the perceived
  # volatility or uncertainty of the unchosen action, promoting broader re-evaluation.
  unchosen_choice = 1 - choice
  action_surprise = action_surprise.at[unchosen_choice].set(
      (1 - volatility_learning_rate) * action_surprise[unchosen_choice] + volatility_learning_rate * absolute_prediction_error * surprise_reset_strength
  )

  # Modulate the base learning rate by global surprise, absolute prediction error, eligibility trace,
  # inversely by the log of the chosen action's count, and the chosen action's specific surprise.
  # Also introduce a recency bias component, making recent experiences more impactful.
  # Modulate the base learning rate by global surprise, absolute prediction error, and eligibility trace.
  # The chosen action's count provides a form of recency and experience-dependent modulation.
  count_modulation_factor = 1.0 / (1.0 + jnp.log1p(choice_counts[choice]))
  volatility_modulation_factor = 1.0 + global_surprise + action_surprise[choice]
  learning_rate_for_update = jnp.clip(base_learning_rate * volatility_modulation_factor * absolute_prediction_error * eligibility_traces[choice] * count_modulation_factor, 0., 1.)
  unchosen_choice = 1 - choice

  eligibility_traces *= eligibility_trace_decay

  # For the unchosen action, decay its eligibility trace.
  # If there's a prediction error, reset the unchosen eligibility trace more strongly.
  eligibility_traces = eligibility_traces.at[unchosen_choice].set(
      jnp.where(
          absolute_prediction_error > 0,
          jnp.maximum(0., eligibility_traces[unchosen_choice] - eligibility_trace_baseline - jnp.clip(absolute_prediction_error * surprise_reset_strength, 0, 1)),
          eligibility_traces[unchosen_choice]
      )
  )

  # Update eligibility trace for the chosen action.
  # Increment is modulated by the absolute prediction error.
  # Decay is also modulated by prediction error: positive PE leads to slower decay, negative PE to faster decay.
  chosen_et_decay_modulation = jnp.where(prediction_error >= 0, 1. - absolute_prediction_error * surprise_reset_strength, 1. + absolute_prediction_error * surprise_reset_strength)
  chosen_et_increment_modulation = jnp.where(prediction_error >= 0, 1. + absolute_prediction_error * (1. + surprise_reset_strength), 1. + absolute_prediction_error * (1. - surprise_reset_strength))

  eligibility_traces = eligibility_traces.at[choice].set(
      eligibility_traces[choice] * jnp.clip(eligibility_trace_decay * chosen_et_decay_modulation, 0., 1.)
      + eligibility_trace_factor * chosen_et_increment_modulation
      + eligibility_trace_baseline
  )

  # Increment the choice count for the chosen action.
  choice_counts = choice_counts.at[choice].set(choice_counts[choice] + 1.0)

  # Update Q-values for both chosen and unchosen actions.
  # Chosen action's Q-value is updated using the modulated learning rate and eligibility trace.
  q_values = q_values.at[choice].set(
      q_values[choice] + learning_rate_for_update * prediction_error * eligibility_traces[choice]
  )
  # For the unchosen action, its Q-value is updated using a small fraction of the chosen action's learning rate,
  # modulated by global and action-specific surprise. This allows for spillover learning.
  # A positive prediction error for the chosen action leads to a small positive update for the unchosen,
  # while a negative prediction error leads to a small negative update, reflecting general environmental state.
  unchosen_learning_rate_modulation = 1.0 + global_surprise + action_surprise[unchosen_choice]
  unchosen_update_amount = jnp.where(
      prediction_error > 0,
      (learning_rate_for_update * unchosen_learning_rate_modulation * 0.5) * prediction_error,
      (learning_rate_for_update * unchosen_learning_rate_modulation * 0.1) * prediction_error
  )
  q_values = q_values.at[unchosen_choice].set(
      q_values[unchosen_choice] + unchosen_update_amount
  )

  q_values = q_values.at[choice].set(q_values[choice] * (1 - chosen_action_decay_rate))
  # Decay the value of the unchosen action, with its decay rate modulated by surprise and choice count.
  surprise_decay_modulation = absolute_prediction_error * surprise_modulated_decay_strength
  count_decay_modulation = 1.0 / (1.0 + jnp.log1p(choice_counts[unchosen_choice]))
  effective_decay_rate = jnp.clip(fixed_decay_rate + surprise_decay_modulation + count_decay_modulation, 0., 1.)
  q_values = q_values.at[unchosen_choice].set(q_values[unchosen_choice] * (1 - effective_decay_rate))

  # Combine updated Q-values, eligibility traces, choice counts, global surprise, and action-specific surprise into the new agent state.
  agent_state = jnp.concatenate((q_values, eligibility_traces, choice_counts, jnp.array([global_surprise]), action_surprise))

  # Compute choice logits using a softmax function over action values.
  choice_logits = inverse_temperature * q_values

  return choice_logits, agent_state


# Shortened to 100k steps then refactored, run 1, low floor.
def fly_bandit_run1_simplified_low_floor(
    params: chex.Array,
    choice: int,
    reward: int,
    agent_state: Optional[chex.Array],
) -> Tuple[chex.Array, chex.Array]:
  """
  Updates the agent's state and computes action preferences (logits) based on a
  reinforcement learning model with recency effects.

  Args:
    params: An array containing the model's parameters.
    choice: The action taken by the agent in the previous step (e.g., 0 or 1).
    reward: The reward received for the previous action.
    agent_state: The internal state of the agent from the previous step,
                 containing Q-values and a recency trace.

  Returns:
    A tuple containing:
    - choice_logits: The logits for the next choice, used to determine action probabilities.
    - new_agent_state: The updated internal state of the agent.
  """
  # --- 1. Unpack Agent State ---
  # If this is the first step (agent_state is None), initialize Q-values and
  # the recency trace to zero. Otherwise, unpack them from the provided state.
  if agent_state is None:
    q_values = 0.0
    recency_trace = 0.0
  else:
    q_values, recency_trace = agent_state

  # --- 2. Unpack Model Parameters ---
  # Assign informative names to the model parameters for better readability.
  learning_rate_positive = params[0]  # Learning rate for positive feedback (reward > Q-value)
  learning_rate_negative = params[1]  # Learning rate for negative feedback (reward < Q-value)
  inverse_temperature = params[2]     # Controls the determinism of choices (exploration vs. exploitation)
  recency_decay_rate = params[3]      # Rate at which the memory of recent choices fades
  recency_strength = params[4]        # The influence of recent choices on the next decision

  # --- 3. Update Q-values (Value Learning) ---
  # Calculate the prediction error: the difference between the actual reward received
  # and the expected reward (the current Q-value).
  prediction_error = reward - q_values

  # Select the appropriate learning rate based on whether the prediction error
  # is positive or negative. This allows the model to learn differently from
  # better-than-expected and worse-than-expected outcomes.
  learning_rate = jnp.where(prediction_error > 0,
                            learning_rate_positive,
                            learning_rate_negative)

  # Calculate the change in Q-value.
  q_value_update = learning_rate * prediction_error

  # Apply the update to get the new Q-value.
  updated_q_values = q_values + q_value_update

  # --- 4. Update Recency Trace ---
  # The recency trace captures a tendency to repeat or avoid the last action.
  # First, the existing trace decays over time.
  decayed_recency_trace = recency_decay_rate * recency_trace

  # Next, the trace is pushed towards the most recent choice.
  # The term (2 * choice - 1) maps choice 0 to -1 and choice 1 to 1.
  choice_direction = (2 * choice - 1)
  updated_recency_trace = decayed_recency_trace + choice_direction

  # --- 5. Calculate Choice Logits for the Next Action ---
  # Logits are the raw, unnormalized preferences for each action. They will
  # typically be passed to a softmax function to generate choice probabilities.

  # a) Determine the influence of recency on each potential choice.
  #    This creates a bias towards (+recency_trace) or away from (-recency_trace)
  #    the direction of the previous choice.
  recency_bias = jnp.array([-updated_recency_trace, updated_recency_trace])

  # b) Scale the recency bias by its strength parameter and use an exponential
  #    to create a multiplicative factor. This ensures the effect is always positive.
  recency_factor = jnp.exp(recency_strength * recency_bias)

  # c) Combine the value component (Q-value) with the recency factor.
  #    In this model, the learned Q-value is broadcasted and applied equally
  #    to both choices, while the recency factor differentiates them.
  value_with_recency = recency_factor * updated_q_values

  # d) Scale the result by the inverse temperature. A higher value leads to
  #    more deterministic choices, while a lower value leads to more random choices.
  choice_logits = inverse_temperature * value_with_recency

  # --- 6. Package the Updated Agent State ---
  # Combine the updated Q-values and recency trace into a new state array
  # to be used in the next step.
  new_agent_state = jnp.array([updated_q_values, updated_recency_trace])

  # --- 7. Return Results ---
  # Return the calculated logits and the new agent state.
  return choice_logits, new_agent_state


# Shortened to 100k steps then refactored, run 2, low floor.
def fly_bandit_run2_simplified_low_floor(
    params: chex.Array,
    choice: int,
    reward: int,
    agent_state: Optional[chex.Array],
) -> tuple[chex.Array, chex.Array]:
  """Cognitive model describing fly behavior on a binary two-armed bandit task.

  Args:
    params: Fit parameters of the model.
    choice: The choice made by a fly on this trial. 0 or 1
    reward: The reward received by a fly on this trial. 0 or 1
    agent_state: The current state of the cognitive model.

  Returns:
    choice_logits: The probabilities that the fly will choose option 0 or 1 on
      the next trial, expressed as logits.
    agent_state: The updated state of the cognitive model.
  """
  # --- 1. Initialization and Parameter Unpacking ---

  # On the first trial, initialize the agent's state from the learned parameters.
  # The agent's state tracks Q-values and stickiness values for each choice.
  # Structure: [q_value_choice_0, q_value_choice_1, stickiness_0, stickiness_1]
  if agent_state is None:
    initial_q_and_stickiness_values = params[5:9]
    agent_state = initial_q_and_stickiness_values

  # Unpack the model's core parameters for clarity.
  learning_rate = params[0]
  inverse_temperature_weight = params[1]
  stickiness_bias_weight = params[2]

  # Unpack the current state of the agent into its components.
  current_q_values = agent_state[0:2]
  current_stickiness_values = agent_state[2:4]


  # --- 2. Update Agent State based on Trial Outcome ---

  # a) Update Q-value for the chosen option using a delta rule (Rescorla-Wagner).
  # First, calculate the prediction error: the difference between actual and expected reward.
  prediction_error = reward - current_q_values[choice]

  # Next, calculate the update amount based on the learning rate.
  q_value_update_amount = learning_rate * prediction_error

  # Apply the update to the Q-value of the action that was just chosen.
  updated_q_values = current_q_values.at[choice].add(q_value_update_amount)


  # b) Update stickiness values to model choice perseveration.
  # First, decay both existing stickiness values by a factor of 0.9.
  decayed_stickiness_values = 0.9 * current_stickiness_values

  # Then, add a fixed amount (0.1) to the stickiness of the chosen action.
  updated_stickiness_values = decayed_stickiness_values.at[choice].add(0.1)


  # c) Reassemble the updated state vector.
  updated_agent_state = jnp.concatenate([updated_q_values, updated_stickiness_values])


  # --- 3. Compute Choice Logits for the Next Trial ---

  # a) Calculate the choice bias introduced by stickiness.
  # This encourages or discourages repeating the same choice.
  stickiness_bias = updated_stickiness_values * stickiness_bias_weight

  # b) Add the stickiness bias to the Q-values to get the final action values.
  biased_q_values = updated_q_values + stickiness_bias

  # c) Calculate a dynamic scaling factor based on the difference between the biased Q-values.
  # This term modulates the influence of the inverse temperature.
  value_difference_scaling = jnp.abs(biased_q_values[0] - biased_q_values[1])

  # d) Convert the final action values into choice logits.
  # This is done by scaling with the inverse temperature and the dynamic factor.
  # Higher inverse temperature leads to more deterministic (less random) choices.
  choice_logits = biased_q_values * inverse_temperature_weight * value_difference_scaling


  # --- 4. Return Results ---
  return choice_logits, updated_agent_state


# Shortened to 100k steps then refactored, run 3, low floor.
def fly_bandit_run3_simplified_low_floor(
    params: chex.Array,
    choice: int,
    reward: int,
    agent_state: Optional[chex.Array],
) -> tuple[chex.Array, chex.Array]:
  """
  Updates an agent's Q-values based on a choice, a reward, and a set of learning parameters.

  The update rule is a combination of a standard Q-learning update and a decay
  towards a set of initial Q-values.
  """

  # --- 1. Unpack Parameters ---
  # The `params` array contains the agent's learning hyperparameters.
  
  # `learning_rate` (alpha): Controls the step size of the learning update.
  # `decay_rate_to_initial_q`: Controls how fast Q-values drift back to their initial state.
  # `initial_q_values`: The starting Q-values, which also act as an anchor for the decay.
  learning_rate, decay_rate_to_initial_q = params[:2]
  initial_q_values = params[2:4]

  # --- 2. Update Q-Values (Agent State) ---
  # This section determines the new Q-values based on the agent's experience.

  # Check if this is the first step (i.e., the agent has no prior state).
  if agent_state is None:
    # If there is no previous state, initialize the Q-values to their starting values.
    updated_q_values = initial_q_values
  else:
    # If a previous state exists, perform a two-part update.

    # Part A: Standard Q-learning update for the chosen action.
    # Get the Q-value for the action that was just taken.
    current_q_for_choice = agent_state[choice]
    # Calculate the prediction error: the difference between the actual reward
    # and the expected reward (the current Q-value).
    prediction_error = reward - current_q_for_choice
    # Calculate the update amount based on the learning rate.
    learning_update = learning_rate * prediction_error
    # Apply the update to the Q-value of the chosen action.
    # Note: `.at[...].add(...)` is the JAX way to perform an immutable update,
    # creating a new array with the modification.
    q_values_after_learning = agent_state.at[choice].add(learning_update)

    # Part B: Decay towards the initial Q-values for all actions.
    # This term causes the agent's beliefs to slowly revert towards its initial priors.
    decay_update = decay_rate_to_initial_q * (initial_q_values - agent_state)

    # Part C: Combine the learning and decay updates to get the final Q-values.
    updated_q_values = q_values_after_learning + decay_update

  # --- 3. Calculate and Return Outputs ---
  # The function returns two arrays: a derived signal and the new agent state.

  # The first return value is a signal calculated from the updated Q-values.
  # It is computed by scaling the new Q-values by the absolute difference between them.
  # This might be used for decision-making or analyzing learning dynamics.
  q_value_difference = jnp.abs(updated_q_values[0] - updated_q_values[1])
  action_selection_signal = q_value_difference * updated_q_values

  # The second return value is the new agent state, which is simply the updated Q-values.
  # This will be passed back as the `agent_state` in the next call to this function.
  new_agent_state = updated_q_values

  return action_selection_signal, new_agent_state


# Shortened to 100k steps then refactored, run 1, medium floor.
def fly_bandit_run1_simplified_medium_floor(
    params: chex.Array,
    choice: int,
    reward: int,
    agent_state: Optional[chex.Array],
) -> tuple[chex.Array, chex.Array]:
  """Cognitive model describing fly behavior on a binary two-armed bandit task.

  This function implements a reinforcement learning model with the following features:
  - Q-learning with separate learning rates for positive and negative feedback.
  - Eligibility traces to attribute rewards to recent choices.
  - A forgetting mechanism for the value of the unchosen option.
  - A reward history mechanism that modulates the learning rate.

  Args:
    params: Fit parameters of the model.
    choice: The choice made by a fly on this trial. 0 or 1
    reward: The reward received by a fly on this trial. 0 or 1
    agent_state: The current state of the cognitive model.

  Returns:
    choice_logits: The probabilities that the fly will choose option 0 or 1 on
      the next trial, expressed as logits.
    agent_state: The updated state of the cognitive model.
  """
  # --- 1. Initialization ---
  # If this is the first trial, initialize the agent's state with zeros.
  # The state vector contains: [q_value_0, q_value_1, trace_0, trace_1, reward_history]
  if agent_state is None:
    agent_state = jnp.zeros((5,))

  # --- 2. Unpack Model Parameters ---
  # Apply a sigmoid function to constrain rate and decay parameters between 0 and 1.
  learning_rate_positive, \
  learning_rate_negative, \
  eligibility_decay_chosen, \
  reward_history_decay = jax.nn.sigmoid(params[:4])

  # Other parameters are used directly without transformation.
  inverse_temperature = params[2]  # Used for choice probability (softmax)
  q_value_decay_unchosen = params[5] # Forgetting factor for the unchosen option's value
  eligibility_decay_unchosen = params[7] # Decay factor for the unchosen option's trace
  reward_history_beta = params[9] # Modulation factor of learning rate by reward history

  # --- 3. Unpack Agent State ---
  # Extract the different components from the agent's state vector.
  q_values = agent_state[0:2]           # Expected value for each choice
  eligibility_traces = agent_state[2:4] # Memory trace for recent choices
  reward_history = agent_state[4]       # Moving average of recent rewards

  # --- 4. Core Computations for Learning ---
  # Determine the unchosen option.
  unchosen_choice = 1 - choice

  # Calculate the prediction error: the difference between actual and expected reward.
  prediction_error = reward - q_values[choice]

  # Update the reward history using an exponential moving average.
  # This tracks the recent rate of rewards.
  reward_history_updated = (
      reward_history * reward_history_decay
      + reward * (1 - reward_history_decay)
  )

  # Select the learning rate based on the sign of the prediction error.
  # This allows for different learning speeds from positive vs. negative feedback.
  base_learning_rate = jnp.where(
      prediction_error >= 0, learning_rate_positive, learning_rate_negative
  )

  # Modulate the learning rate by the reward history.
  # This can increase learning speed in reward-rich environments.
  learning_rate_modulated = base_learning_rate * (
      1.0 + reward_history_beta * reward_history
  )

  # --- 5. Update Eligibility Traces ---
  # The eligibility trace for the unchosen option decays.
  updated_trace_unchosen = eligibility_traces[unchosen_choice] * eligibility_decay_unchosen

  # The trace for the chosen option is updated based on its previous value and a decay factor,
  # and is incremented by 1 to mark its recent selection.
  updated_trace_chosen = eligibility_traces[choice] * eligibility_decay_chosen + 1.0

  # Combine the updated traces.
  eligibility_traces_updated = jnp.zeros_like(eligibility_traces).at[choice].set(
      updated_trace_chosen
  ).at[unchosen_choice].set(
      updated_trace_unchosen
  )

  # --- 6. Update Q-Values (Action Values) ---
  # Calculate the change in value for the chosen option, scaled by the learning rate,
  # prediction error, and the eligibility trace.
  q_value_update_amount = (
      learning_rate_modulated
      * prediction_error
      * eligibility_traces_updated[choice]
  )
  
  # Apply the learning update to the chosen option's Q-value.
  q_values_after_learning = q_values.at[choice].add(q_value_update_amount)
  
  # Apply a decay (forgetting) factor to the unchosen option's Q-value.
  q_values_updated = q_values_after_learning.at[unchosen_choice].set(
      q_values_after_learning[unchosen_choice] * q_value_decay_unchosen
  )

  # --- 7. Prepare Outputs ---
  # Reassemble the updated components into the new agent state vector.
  # jnp.newaxis is used to make the scalar reward_history a 1-element array for concatenation.
  new_agent_state = jnp.concatenate(
      (q_values_updated, eligibility_traces_updated, reward_history_updated[jnp.newaxis])
  )

  # Calculate the choice logits for the next trial using the updated Q-values.
  # The inverse_temperature parameter controls the stochasticity of the choice (softmax).
  choice_logits = inverse_temperature * q_values_updated

  return choice_logits, new_agent_state


# Shortened to 100k steps then refactored, run 2, medium floor.
def fly_bandit_run2_simplified_medium_floor(
    params: chex.Array,
    choice: int,
    reward: int,
    agent_state: Optional[chex.Array],
) -> tuple[chex.Array, chex.Array]:
  """Cognitive model describing fly behavior on a binary two-armed bandit task.

  Args:
    params: Fit parameters of the model.
    choice: The choice made by a fly on this trial. 0 or 1
    reward: The reward received by a fly on this trial. 0 or 1
    agent_state: The current state of the cognitive model.

  Returns:
    choice_logits: The probabilities that the fly will choose option 0 or 1 on
      the next trial, expressed as logits.
    agent_state: The updated state of the cognitive model.
  """
  # --- 1. Initialize State and Unpack Parameters ---

  # If this is the first trial (agent_state is None), initialize a new state.
  # The state vector contains: [Q-value for option 0, Q-value for option 1,
  #                         stickiness for option 0, stickiness for option 1,
  #                         accumulated reward prediction error]
  if agent_state is None:
    agent_state = jnp.zeros(5)

  # Unpack the model's fixed parameters into named variables for clarity.
  # The `*_` syntax ignores any extra parameters that might be present.
  learning_rate, forgetting_rate, previous_choice_bias, choice_stickiness_lr, *_ = params

  # Unpack the agent's current state.
  # Although not used directly here, this shows what each index represents.
  q_values_0, q_values_1, stickiness_0, stickiness_1, rpe_accumulator = agent_state


  # --- 2. Update Agent State Based on Trial Outcome ---

  # Calculate the reward prediction error (RPE). This is the difference between
  # the actual reward received and the expected reward (rpe_accumulator).
  prediction_error = reward - rpe_accumulator

  # Calculate the learning update amount, often called "delta".
  learning_update_amount = learning_rate * prediction_error

  # Apply forgetting to the Q-value of the *unchosen* option.
  # The index of the unchosen option is (1 - choice).
  state_after_forgetting = agent_state.at[1 - choice].multiply(1 - forgetting_rate)

  # Update the Q-value of the *chosen* option based on the learning amount.
  state_after_q_update = state_after_forgetting.at[choice].add(learning_update_amount)

  # Update the accumulated reward prediction error.
  state_after_rpe_update = state_after_q_update.at[4].add(learning_update_amount)

  # Decay the choice stickiness for both options.
  state_after_stickiness_decay = state_after_rpe_update.at[2:4].multiply(1 - choice_stickiness_lr)

  # Add stickiness to the *chosen* option.
  # The stickiness index for the chosen option is (2 + choice).
  updated_agent_state = state_after_stickiness_decay.at[2 + choice].add(choice_stickiness_lr)


  # --- 3. Calculate Choice Logits for the Next Trial ---

  # Extract the newly updated Q-values and stickiness values from the state.
  updated_q_values = updated_agent_state[:2]
  updated_stickiness = updated_agent_state[2:4]

  # Combine Q-values and choice stickiness to get the effective action values.
  # The `previous_choice_bias` parameter weights the influence of stickiness.
  effective_q_values = updated_q_values + updated_stickiness * previous_choice_bias

  # Calculate a scaling factor based on the absolute difference between the
  # effective Q-values. This term amplifies choice probability when values are far apart.
  q_value_difference = jnp.abs(effective_q_values[0] - effective_q_values[1])

  # The final choice logits are the effective Q-values scaled by their difference.
  choice_logits = effective_q_values * q_value_difference


  # --- 4. Return Results ---

  return choice_logits, updated_agent_state


# Shortened to 100k steps then refactored, run 3, medium floor.
def fly_bandit_run3_simplified_medium_floor(
    params: chex.Array,
    choice: int,
    reward: int,
    agent_state: Optional[chex.Array],
) -> tuple[chex.Array, chex.Array]:
  """Cognitive model describing fly behavior on a binary two-armed bandit task.

  Args:
    params: Fit parameters of the model.
    choice: The choice made by a fly on this trial. 0 or 1
    reward: The reward received by a fly on this trial. 0 or 1
    agent_state: The current state of the cognitive model.

  Returns:
    choice_logits: The probabilities that the fly will choose option 0 or 1 on
      the next trial, expressed as logits.
    agent_state: The updated state of the cognitive model.
  """
  # === 1. Unpack Parameters and Initialize State ===

  # Unpack the model's fit parameters from the input array. These parameters
  # define the agent's learning and decision-making characteristics.
  (
      learning_rate_base_raw,
      beta,
      decay_rate_unchosen_to_initial_q,
      initial_q,
      value_sensitivity,
      prediction_error_sensitivity,
      perseveration_bias,
      confidence_beta_slope,
      confidence_beta_midpoint,
  ) = params[:9]

  # If this is the first trial (agent_state is None), initialize the agent's
  # state (Q-values) to the initial_q parameter for both choices.
  if agent_state is None:
    agent_state = jnp.full(2, initial_q)

  # === 2. Update Q-Values Based on Choice and Reward ===

  # --- Update the value of the CHOSEN option ---

  # Calculate the prediction error: the difference between the actual reward
  # received and the agent's expected value (Q-value) for the chosen option.
  q_value_chosen = agent_state[choice]
  prediction_error = reward - q_value_chosen

  # The learning rate is dynamically modulated by several factors.
  # First, calculate the base learning rate by passing the raw parameter through
  # a sigmoid function, constraining it to be between 0 and 1.
  base_learning_rate = jax.nn.sigmoid(learning_rate_base_raw)

  # Second, modulate the learning rate based on the current value of the chosen option.
  # This term models how the agent's learning is sensitive to its own value estimate.
  value_sensitivity_modulation = jax.nn.sigmoid(
      value_sensitivity - jnp.abs(q_value_chosen)
  )

  # Third, modulate the learning rate based on the magnitude of the prediction error.
  # This allows learning to be faster when the outcome is more surprising.
  prediction_error_magnitude = jnp.abs(prediction_error)
  prediction_error_modulation = (
      1.0 + prediction_error_sensitivity * prediction_error_magnitude
  )

  # Combine the base learning rate with the two modulation factors.
  modulated_learning_rate = (
      base_learning_rate
      * value_sensitivity_modulation
      * prediction_error_modulation
  )

  # Update the Q-value of the chosen option using the prediction error and the
  # modulated learning rate (a Rescorla-Wagner style update).
  updated_q_value_chosen = q_value_chosen + modulated_learning_rate * prediction_error
  agent_state = agent_state.at[choice].set(updated_q_value_chosen)

  # --- Update the value of the UNCHOSEN option ---

  # The Q-value of the unchosen option decays towards the initial Q-value.
  # This models a 'forgetting' or regression-to-the-mean effect for options
  # that are not selected.
  unchosen_choice = 1 - choice
  q_value_unchosen = agent_state[unchosen_choice]
  updated_q_value_unchosen = q_value_unchosen * (
      1 - decay_rate_unchosen_to_initial_q
  ) + initial_q * decay_rate_unchosen_to_initial_q
  agent_state = agent_state.at[unchosen_choice].set(updated_q_value_unchosen)

  # === 3. Compute Choice Probabilities (Logits) for the Next Trial ===

  # Apply a perseveration bias, which adds a bonus to the value of the
  # option that was just chosen, making it more likely to be chosen again.
  # This creates a temporary, biased set of Q-values for decision-making.
  q_value_of_last_choice = agent_state[choice]
  biased_q_value_of_last_choice = q_value_of_last_choice + perseveration_bias
  biased_state = agent_state.at[choice].set(biased_q_value_of_last_choice)

  # The decision's randomness (inverse temperature, beta) is modulated by confidence.
  # Higher confidence (a larger difference in Q-values) can lead to more
  # deterministic (less random) choices.

  # First, calculate confidence as the absolute difference between the biased Q-values.
  confidence = jnp.abs(biased_state[0] - biased_state[1])

  # Use a sigmoid function to determine the scaling factor for beta based on confidence.
  # The midpoint and slope control the shape of this confidence-beta relationship.
  confidence_effect = confidence_beta_slope * (confidence - confidence_beta_midpoint)
  beta_scaling_factor = jax.nn.sigmoid(confidence_effect)

  # Calculate the final, modulated beta to be used in the choice rule.
  modulated_beta = beta * beta_scaling_factor

  # Finally, compute the choice logits by scaling the biased Q-values by the
  # modulated beta. This is the standard softmax/logistic choice rule.
  # Logits are the inputs to a softmax function for generating choice probabilities.
  choice_logits = modulated_beta * biased_state

  # Return the computed logits for the next choice and the updated agent state.
  return choice_logits, agent_state


# Shortened to 100k steps then refactored, run 1, high floor.
def fly_bandit_run1_simplified_high_floor(
    params: chex.Array,
    choice: int,
    reward: int,
    agent_state: Optional[chex.Array],
) -> tuple[chex.Array, chex.Array]:
  """Cognitive model describing fly behavior on a binary two-armed bandit task.

  Args:
    params: Fit parameters of the model.
    choice: The choice made by a fly on this trial. 0 or 1
    reward: The reward received by a fly on this trial. 0 or 1
    agent_state: The current state of the cognitive model.

  Returns:
    choice_logits: The probabilities that the fly will choose option 0 or 1 on
      the next trial, expressed as logits.
    agent_state: The updated state of the cognitive model.
  """
  # --- 1. Initialization ---
  # On the very first trial, the agent_state is None. Initialize it to a
  # vector of zeros. The state holds [q_value_0, q_value_1,
  # eligibility_trace_0, eligibility_trace_1, reward_history].
  if agent_state is None:
    agent_state = jnp.zeros(5)

  # --- 2. Unpack Parameters ---
  # The model's behavior is determined by a set of parameters.
  # Some parameters are passed through a sigmoid function to constrain them
  # between 0 and 1, which is typical for rates and probabilities.
  (
      learning_rate_positive,
      learning_rate_negative,
      eligibility_decay_rate,
      eligibility_boost,
      reward_history_decay,
  ) = jax.nn.sigmoid(params[:5])

  # Other parameters are used directly.
  unchosen_q_decay_rate, inverse_temperature, reward_history_beta = params[5:8]

  # --- 3. Unpack Agent State ---
  # Extract the internal variables of the model from the agent_state vector.
  q_values = agent_state[:2]  # Estimated value for each choice
  eligibility_traces = agent_state[2:4]  # Credit assignment traces
  reward_history = agent_state[4]  # A moving average of recent rewards

  # --- 4. Core Model Computations ---

  # --- 4a. Calculate Prediction Error ---
  # The prediction error is the difference between the actual reward received
  # and the expected reward (the Q-value of the chosen option).
  prediction_error = reward - q_values[choice]

  # --- 4b. Update Reward History ---
  # The reward history is an exponential moving average of past rewards.
  # It tracks the general richness of the environment.
  reward_history_updated = (
      reward_history * reward_history_decay
      + reward * (1.0 - reward_history_decay)
  )

  # --- 4c. Update Eligibility Traces ---
  # Eligibility traces help assign credit for outcomes to recent choices.
  # First, all traces decay.
  traces_decayed = eligibility_traces * eligibility_decay_rate
  # Then, the trace for the chosen action is boosted, and the unchosen is depressed.
  unchosen_choice = 1 - choice
  traces_boosted = traces_decayed.at[choice].add(eligibility_boost)
  traces_final = traces_boosted.at[unchosen_choice].add(-eligibility_boost)
  # Finally, ensure traces do not become negative.
  eligibility_traces_updated = jnp.maximum(0.0, traces_final)

  # --- 4d. Update Q-Values ---
  # The Q-values (expected rewards) are updated based on the outcome.

  # -- Update for the CHOSEN action --
  # Select a different learning rate depending on whether the prediction error
  # was positive (better than expected) or negative (worse than expected).
  learning_rate = jnp.where(
      prediction_error >= 0, learning_rate_positive, learning_rate_negative
  )

  # The magnitude of the update is modulated by the reward history.
  # Higher recent rewards can amplify learning.
  reward_history_weight = jnp.exp(reward_history_beta * reward_history_updated)

  # Calculate the final change for the chosen Q-value.
  # This change depends on the learning rate, prediction error, reward history,
  # and the eligibility trace for the chosen action.
  q_update_delta = (
      learning_rate * reward_history_weight * prediction_error * eligibility_traces_updated[choice]
  )

  # Apply the update to the Q-value of the chosen action.
  q_values_after_learning = q_values.at[choice].add(q_update_delta)

  # -- Update for the UNCHOSEN action --
  # The Q-value of the unchosen action decays towards zero. This represents
  # a form of forgetting.
  q_values_updated = q_values_after_learning.at[unchosen_choice].set(
      q_values_after_learning[unchosen_choice] * unchosen_q_decay_rate
  )

  # --- 5. Assemble New State ---
  # Combine the updated internal variables back into a single state vector for
  # the next trial.
  new_agent_state = jnp.concatenate(
      (q_values_updated, eligibility_traces_updated, jnp.array([reward_history_updated]))
  )

  # --- 6. Compute Next Choice Logits ---
  # The probability of choosing each option on the next trial is based on the
  # updated Q-values. The 'inverse_temperature' parameter controls the
  # randomness of the choice (higher value means more deterministic).
  # These are returned as logits, the raw inputs to a softmax function.
  choice_logits = inverse_temperature * q_values_updated

  # --- 7. Return Results ---
  return choice_logits, new_agent_state


# Shortened to 100k steps then refactored, run 2, high floor.
def fly_bandit_run2_simplified_high_floor(
    params: chex.Array,
    choice: int,
    reward: int,
    agent_state: Optional[chex.Array],
) -> tuple[chex.Array, chex.Array]:
  """
  A reinforcement learning agent model that updates its internal state based on a choice and its reward,
  and then calculates the logits for the next choice.
  """
  # --- 1. Initialization ---
  # If this is the first trial, initialize the agent's state with zeros.
  # The agent state holds [Q-value for choice 0, Q-value for choice 1, 
  #                        stickiness for 0, stickiness for 1,
  #                        reward reference point, value uncertainty].
  if agent_state is None:
    agent_state = jnp.zeros(6)

  # --- 2. Unpack and Transform Model Parameters ---
  # The `params` array contains the raw, learnable parameters of the model.
  (
      raw_learning_rate, 
      raw_static_inverse_temperature, 
      uncertainty_weight,
      raw_forgetting_rate, 
      raw_previous_choice_bias, 
      raw_choice_stickiness_lr,
      outcome_valence_stickiness_weight, 
      raw_choice_perseveration_decay,
      raw_reward_reference_learning_rate, 
      raw_uncertainty_learning_rate
  ) = params

  # Apply transformations to constrain parameters to valid ranges.
  # Sigmoid maps values to (0, 1), useful for rates and probabilities.
  learning_rate = jax.nn.sigmoid(raw_learning_rate)
  forgetting_rate = jax.nn.sigmoid(raw_forgetting_rate)
  choice_stickiness_lr = jax.nn.sigmoid(raw_choice_stickiness_lr)
  choice_perseveration_decay = jax.nn.sigmoid(raw_choice_perseveration_decay)
  reward_reference_learning_rate = jax.nn.sigmoid(raw_reward_reference_learning_rate)
  uncertainty_learning_rate = jax.nn.sigmoid(raw_uncertainty_learning_rate)

  # Exponential maps values to be positive (> 0).
  static_inverse_temperature = jnp.exp(raw_static_inverse_temperature)
  previous_choice_bias = jnp.exp(raw_previous_choice_bias)

  # --- 3. Unpack Agent's Internal State ---
  # Extract the different components from the agent's state array.
  q_values = agent_state[:2]
  stickiness_values = agent_state[2:4]
  reward_reference_point = agent_state[4]
  value_uncertainty = agent_state[5]
  
  # --- 4. Update State Based on Last Trial's Outcome ---

  # 4a. Apply forgetting to the Q-value of the *unchosen* option.
  unchosen_option_index = 1 - choice
  q_values = q_values.at[unchosen_option_index].set(
      q_values[unchosen_option_index] * (1 - forgetting_rate)
  )

  # 4b. Calculate the prediction error.
  prediction_error = reward - reward_reference_point
  prediction_error_abs = jnp.abs(prediction_error)
  
  # 4c. Update the Q-value of the *chosen* option using the prediction error.
  q_values = q_values.at[choice].set(
      q_values[choice] + learning_rate * prediction_error
  )

  # 4d. Update the reward reference point (aspirations).
  reward_reference_point_new = reward_reference_point + reward_reference_learning_rate * prediction_error

  # 4e. Update value uncertainty based on the absolute prediction error.
  uncertainty_update_term = prediction_error_abs * (prediction_error_abs - value_uncertainty)
  value_uncertainty_new = value_uncertainty + uncertainty_learning_rate * uncertainty_update_term
  
  # 4f. Update choice stickiness/perseveration values.
  # First, decay all stickiness values.
  stickiness_values = stickiness_values * (1 - choice_perseveration_decay)
  
  # Then, increase the stickiness of the chosen option.
  # This increase is modulated by the reward outcome and prediction error.
  stickiness_update_factor = (
      1 + outcome_valence_stickiness_weight * reward + prediction_error_abs
  )
  current_stickiness_of_choice = stickiness_values[choice]
  stickiness_increase = choice_stickiness_lr * stickiness_update_factor * (1 - current_stickiness_of_choice)
  
  stickiness_values = stickiness_values.at[choice].set(
      current_stickiness_of_choice + stickiness_increase
  )

  # --- 5. Assemble the New Agent State ---
  # Combine all the updated components back into a single state array for the next trial.
  agent_state = jnp.concatenate([
      q_values, 
      stickiness_values, 
      jnp.array([reward_reference_point_new, value_uncertainty_new])
  ])

  # --- 6. Calculate Choice Logits for the Next Decision ---

  # 6a. Combine Q-values with stickiness to get biased values.
  # This represents a tendency to repeat previous choices.
  perseveration_effect = stickiness_values * previous_choice_bias * (1 - choice_perseveration_decay)
  biased_q_values = q_values + perseveration_effect

  # 6b. Calculate a dynamic inverse temperature (decision noise).
  # This makes decisions more random (lower temperature) when there is high uncertainty.
  q_value_difference = jnp.abs(biased_q_values[0] - biased_q_values[1])
  uncertainty_penalty = 5 * jax.nn.sigmoid(value_uncertainty)
  
  # The exploration bonus/penalty is added to the static temperature.
  exploration_term = uncertainty_weight * (q_value_difference - uncertainty_penalty)
  dynamic_inverse_temperature = static_inverse_temperature + exploration_term
  
  # Ensure the temperature does not go below a minimum threshold.
  dynamic_inverse_temperature = jnp.maximum(0.1, dynamic_inverse_temperature)

  # 6c. Calculate the final choice logits.
  # These are the unnormalized log-probabilities for the softmax function.
  choice_logits = biased_q_values * dynamic_inverse_temperature

  return choice_logits, agent_state


# Shortened to 100k steps then refactored, run 3, high floor.
def fly_bandit_run3_simplified_high_floor(
    params: chex.Array,
    choice: int,
    reward: int,
    agent_state: Optional[chex.Array],
) -> tuple[chex.Array, chex.Array]:
  """Cognitive model describing fly behavior on a binary two-armed bandit task.

  This function has been reformatted for clarity, with computations separated
  onto different lines, informative variable names, and comments describing
  each step. The underlying functionality has not been changed.

  Args:
    params: Fit parameters of the model.
    choice: The choice made by a fly on this trial. 0 or 1
    reward: The reward received by a fly on this trial. 0 or 1
    agent_state: The current state of the cognitive model.

  Returns:
    choice_logits: The probabilities that the fly will choose option 0 or 1 on
      the next trial, expressed as logits.
    agent_state: The updated state of the cognitive model.
  """
  # --- 1. Initialization and Parameter Unpacking ---

  # If this is the first trial, initialize the agent's state to all zeros.
  # The state vector contains [Q-value for option 0, Q-value for option 1,
  #                          Eligibility trace for option 0, Eligibility trace for option 1].
  if agent_state is None:
    agent_state = jnp.zeros(4)

  # Unpack the model's behavioral parameters.
  # These are passed through a sigmoid function to constrain them between 0 and 1.
  (
      learning_rate,
      fixed_decay_rate,
      eligibility_trace_decay,
      eligibility_trace_factor,
      surprise_influence_factor,
      chosen_action_decay_rate,
      decay_offset_unchosen_et,
  ) = jax.nn.sigmoid(params[:7])

  # Unpack parameters that are not constrained between 0 and 1.
  inverse_temperature, prior_choice_bias = params[7:9]

  # --- 2. Unpack Agent State ---

  # Extract the Q-values (expected value of each choice) from the state.
  q_values = agent_state[:2]
  # Extract the eligibility traces from the state.
  eligibility_traces = agent_state[2:]

  # --- 3. Core Reinforcement Learning Calculations ---

  # Calculate the prediction error (surprise): the difference between the
  # actual reward received and the expected reward (Q-value) for the chosen option.
  prediction_error = reward - q_values[choice]
  absolute_prediction_error = jnp.abs(prediction_error)

  # Identify the index of the action that was not chosen.
  unchosen_choice = 1 - choice

  # --- 4. Update Eligibility Traces (ETs) ---

  # First, apply a passive decay to both eligibility traces.
  eligibility_traces_decayed = eligibility_traces * eligibility_trace_decay

  # For the UNCHOSEN action, the ET is further decreased based on the magnitude
  # of the prediction error (surprise). This decay only happens if there was a surprise.
  # The `jnp.maximum(0., ...)` ensures the trace does not become negative.
  is_surprise = (absolute_prediction_error > 0)
  unchosen_et_decay_amount = (
      decay_offset_unchosen_et + absolute_prediction_error * surprise_influence_factor
  )
  updated_eligibility_trace_unchosen = jnp.maximum(
      0.,
      eligibility_traces_decayed[unchosen_choice] - unchosen_et_decay_amount * is_surprise
  )

  # For the CHOSEN action, the ET is updated based on two factors:
  # 1. A decay modulated by reward and surprise.
  # 2. An increment based on a base factor and the magnitude of the surprise.
  reward_modulated_decay = eligibility_traces_decayed[choice] * (
      1. - reward * surprise_influence_factor
  )
  surprise_based_increment = eligibility_trace_factor * (
      1. + absolute_prediction_error
  )
  updated_eligibility_trace_chosen = reward_modulated_decay + surprise_based_increment

  # Reassemble the eligibility traces array with the updated values.
  eligibility_traces = eligibility_traces_decayed.at[unchosen_choice].set(
      updated_eligibility_trace_unchosen
  )
  eligibility_traces = eligibility_traces.at[choice].set(
      updated_eligibility_trace_chosen
  )

  # --- 5. Update Q-Values ---

  # First, update both Q-values based on the prediction error, scaled by the
  # eligibility traces. Note: This specific update rule applies the learning
  # to both Q-values, weighted by their respective ETs.
  learning_update = (
      learning_rate * absolute_prediction_error *
      eligibility_traces[choice] * prediction_error * eligibility_traces
  )
  q_values = q_values + learning_update

  # Second, apply a specific decay to the Q-value of the CHOSEN action.
  q_values = q_values.at[choice].set(
      q_values[choice] * (1 - chosen_action_decay_rate)
  )

  # Third, apply a separate decay to the Q-value of the UNCHOSEN action.
  # This decay is influenced by the magnitude of the surprise.
  unchosen_q_decay_factor = (
      1 - fixed_decay_rate - absolute_prediction_error * surprise_influence_factor
  )
  q_values = q_values.at[unchosen_choice].set(
      q_values[unchosen_choice] * unchosen_q_decay_factor
  )

  # --- 6. Prepare Outputs for Next Trial ---

  # Recombine the updated Q-values and eligibility traces into the new agent state.
  agent_state = jnp.concatenate((q_values, eligibility_traces))

  # Calculate the choice logits for the next trial. This is part of the
  # softmax decision rule, where Q-values are scaled by the inverse temperature.
  choice_logits = inverse_temperature * q_values

  # Apply a fixed bias to the logit for option 0, reflecting a potential
  # baseline preference for one choice over the other.
  choice_logits = choice_logits.at[0].set(choice_logits[0] + prior_choice_bias)

  return choice_logits, agent_state


def fly_bandit_synthesis(
    params: chex.Array,
    choice: int,
    reward: int,
    agent_state: Optional[chex.Array],
) -> Tuple[chex.Array, chex.Array]:
  """Synthesis program combining the main ideas from discovered programs."""
  # --- 1. Initialization ---
  if agent_state is None:
    agent_state = jnp.zeros(5)

  # --- 2. Unpack parameters ---
  # Some are sigmoided...
  (
      learning_rate_positive,
      learning_rate_negative,
      eligibility_decay_rate,
      eligibility_boost,
      reward_history_decay,
  ) = jax.nn.sigmoid(params[:5])
  # Some are not.
  unchosen_q_decay_rate, inverse_temperature, reward_history_beta = params[5:8]

  # --- 3. Get agent state ---
  q_values = agent_state[:2]
  eligibility_traces = agent_state[2:4]
  reward_history = agent_state[4]

  # --- 4. Core model computations ---
  unchosen_choice = 1 - choice
  # Compute prediction error
  prediction_error = reward - q_values[choice]
  # Update reward history
  reward_history_updated = (
      reward_history * reward_history_decay +
      reward * (1.0 - reward_history_decay)
  )
  # Update eligibility traces
  traces_decayed = eligibility_traces * eligibility_decay_rate
  traces_boosted = traces_decayed.at[choice].add(eligibility_boost)
  traces_final = traces_boosted.at[unchosen_choice].add(-eligibility_boost)
  eligibility_traces_updated = jnp.maximum(0.0, traces_final)

  # Compute learning rate and learning modulators
  learning_rate = jnp.where(prediction_error >= 0,
                            learning_rate_positive,
                            learning_rate_negative)
  # Higher recent rewards can amplify learning
  reward_history_weight = jnp.exp(reward_history_beta * reward_history_updated)
  # Update amount depends on prediction error, learning rate, reward history,
  # and eligibility traces.
  q_update_delta = (prediction_error * learning_rate * reward_history_weight *
                    eligibility_traces_updated[choice])
  q_values_after_learning = q_values.at[choice].add(q_update_delta)
  q_values_updated = q_values_after_learning.at[unchosen_choice].set(
      q_values_after_learning[unchosen_choice] * unchosen_q_decay_rate)

  # --- 5. Prepare new state ---
  new_agent_state = jnp.concatenate(
      (q_values_updated, eligibility_traces_updated,
       jnp.array([reward_history_updated]))
  )

  # --- 6. Compute logits and return ---
  logits = inverse_temperature * q_values_updated
  return logits, new_agent_state
```
