## Supplementary material for "AI-Discovered Cognitive Models Reveal Novel Insights into Human and Animal Learning": DataDIVER Discovered Programs: human_bandit_models.py

```
# Copyright 2026 DeepMind Technologies Limited.
#
# Licensed under the Apache License, Version 2.0 (the "License");
# you may not use this file except in compliance with the License.
# You may obtain a copy of the License at
#
#     http://www.apache.org/licenses/LICENSE-2.0
#
# Unless required by applicable law or agreed to in writing, software
# distributed under the License is distributed on an "AS IS" BASIS,
# WITHOUT WARRANTIES OR CONDITIONS OF ANY KIND, either express or implied.
# See the License for the specific language governing permissions and
# limitations under the License.

"""Saved models for the Nature 2026 submission."""
from typing import Optional

import chex
import jax
import jax.numpy as jnp


# SOO human_bandit fit only.
def human_bandit_run1_fit_only(
    params: chex.Array,
    choice: int,
    reward: float,
    agent_state: Optional[chex.Array],
) -> tuple[chex.Array, chex.Array]:
  """Cognitive model describing human behavior on a multi-armed bandit task.

  Assumes the agent is presented with four options on each trial.

  Args:
    params: Fit parameters of the model. Different parameters are used for
      different subjects.
    choice: Choice made by the agent on the previous trial. 0, 1, 2, or 3
    reward: Reward received by the agent on the previous trial. A float between
      0 and 1.
    agent_state: The current state of the cognitive model.

  Returns:
    choice_logits: The probabilities that the agent will choose option 0, 1, 2,
      or 3 on the next trial, expressed as logits.
    agent_state: New agent state
  """
  # Do not remove comments or TODOs from this program.

  # Define parameters. All learning rates and decay rates are transformed with the logistic function
  # to be between 0 and 1. Inverse temperature is scaled to be between 0 and 10.
  # Perseveration biases are transformed with softplus to be non-negative.
  initial_q_value = jax.nn.sigmoid(params[0])
  learning_rate = jax.nn.sigmoid(params[1])
  inverse_temperature = 10 * jax.nn.sigmoid(params[2]) # Scales from 0 to 10
  q_value_prior_weight = jax.nn.sigmoid(params[3]) # Weight for combining initial_q_value and average_reward_estimate
  unchosen_learning_rate = 0.95 * jax.nn.sigmoid(params[4]) + 0.025 # Constrained to be between 0.025 and 0.975 for stability.
  perseveration_learning_rate = jax.nn.sigmoid(params[5])
  perseveration_decay_rate = 0.95 * jax.nn.sigmoid(params[6]) + 0.025 # Constrained to be between 0.025 and 0.975 for stability.
  initial_perseveration_value = jax.nn.sigmoid(params[7])
  perseveration_bias_strength = jax.nn.softplus(params[8]) # Non-negative
  perseveration_reward_sensitivity = 5 * jax.nn.softplus(params[9]) # Non-negative, allowing for a broader positive range.

  # Initialize Q-values, perseveration trace, average reward estimate, and recency trace if state is None.
  if agent_state is None:
    q_values = jnp.full((4,), initial_q_value)
    perseveration_trace = jnp.full((4,), initial_perseveration_value)
    average_reward_estimate = initial_q_value
    recency_trace = jnp.full((4,), 0.0) # New state variable to track recency, initialized to zero.
    reward_variance_estimate = jnp.array(0.0) # New state variable for tracking reward variance
    q_value_uncertainty = jnp.full((4,), 0.5) # New state variable for tracking Q-value uncertainty, initialized to a neutral value.
    uncertainty_prediction_error_estimate = jnp.array(0.0) # New state variable for tracking meta-level uncertainty prediction error
    agent_state = jnp.concatenate([q_values, perseveration_trace, jnp.array([average_reward_estimate]), recency_trace, jnp.array([reward_variance_estimate]), q_value_uncertainty, jnp.array([uncertainty_prediction_error_estimate])])

  # Unpack Q-values, perseveration trace, average reward estimate, recency trace, reward variance estimate, Q-value uncertainty, and uncertainty prediction error from the agent state.
  q_values = agent_state[:4]
  perseveration_trace = agent_state[4:8]
  average_reward_estimate = agent_state[8]
  recency_trace = agent_state[9:13] # Unpack the new recency_trace
  reward_variance_estimate = agent_state[13] # Unpack the new reward variance estimate
  q_value_uncertainty = agent_state[14:18] # Unpack the new q_value_uncertainty
  uncertainty_prediction_error_estimate = agent_state[18] # Unpack the new uncertainty prediction error estimate
  exploration_bonus_strength = agent_state[19] # Unpack the new general exploration bonus strength.

  # Compute a dynamic prior for Q-values, a weighted average of initial Q and current average reward.
  # Q-values decay towards this dynamic prior, effectively integrating the previous decay and adding an adaptive baseline.
  dynamic_q_prior = q_value_prior_weight * initial_q_value + (1 - q_value_prior_weight) * average_reward_estimate
  # Introduce a global, small decay rate towards the initial_q_value. This acts as a fixed prior.
  global_q_decay_to_initial = unchosen_learning_rate * 0.1 # A small fraction of the unchosen learning rate.
  q_values = q_values * (1 - global_q_decay_to_initial) + initial_q_value * global_q_decay_to_initial # Decay all Q-values towards initial_q_value
  q_values = q_values * (1 - unchosen_learning_rate) + dynamic_q_prior * unchosen_learning_rate # Unchosen learning rate acts as the decay to the dynamic prior

  # Update the average reward estimate using a Rescorla-Wagner-like rule.
  # This tracks the overall rewardingness of the environment.
  reward_prediction_error_avg = reward - average_reward_estimate
  # Modulate the learning rate for the average reward estimate.
  # It now adapts based on the signed prediction error, increasing for large errors in either direction,
  # but also adapting based on the consistency of the error (e.g., if it consistently overestimates, learn faster).
  # The factor is now also scaled by the current average reward, allowing for more dynamic learning in different reward environments.
  avg_reward_error_magnitude_factor = 1.0 + jax.nn.softplus(jnp.abs(reward_prediction_error_avg))
  # Introduce a term that scales learning based on whether the average reward is currently "high" or "low",
  # allowing for different sensitivity depending on the environmental baseline.
  avg_reward_context_scaling = 0.5 + 1.5 * jax.nn.sigmoid(average_reward_estimate - 0.5) # Scale between 0.5 and 2.0
  effective_avg_reward_learning_rate = unchosen_learning_rate * avg_reward_error_magnitude_factor * avg_reward_context_scaling
  effective_avg_reward_learning_rate = jnp.clip(effective_avg_reward_learning_rate, 0.0, 1.0) # Ensure rates are within bounds
  average_reward_estimate = average_reward_estimate + effective_avg_reward_learning_rate * reward_prediction_error_avg
  # Update the reward variance estimate using an exponential moving average.
  # This tracks the volatility of the rewards over time.
  reward_squared_error = (reward - average_reward_estimate)**2
  variance_learning_rate = unchosen_learning_rate * 0.5 # A fraction of the unchosen_learning_rate
  reward_variance_estimate = reward_variance_estimate * (1 - variance_learning_rate) + reward_squared_error * variance_learning_rate

  # Modulate the decay rate for the average reward estimate based on the magnitude of the average reward prediction error
  # AND the estimated reward variance. Higher variance suggests greater environmental volatility, demanding faster decay.
  adaptive_avg_reward_decay_factor = unchosen_learning_rate * 0.1 * (1.0 + jax.nn.softplus(jnp.abs(reward_prediction_error_avg) * 2.0))
  # Introduce variance-based scaling: higher variance means more decay towards initial Q.
  variance_decay_scaling = 1.0 + jax.nn.softplus(reward_variance_estimate * 10.0) # Scale up decay based on variance
  adaptive_avg_reward_decay_factor = adaptive_avg_reward_decay_factor * variance_decay_scaling
  adaptive_avg_reward_decay_factor = jnp.clip(adaptive_avg_reward_decay_factor, 0.0, 1.0) # Ensure rates are within bounds
  average_reward_estimate = average_reward_estimate * (1 - adaptive_avg_reward_decay_factor) + initial_q_value * adaptive_avg_reward_decay_factor

  unchosen_mask = jnp.arange(4) != choice
  # Option-specific uncertainty: higher for Q-values closer to the average reward estimate (less distinct information).
  # This factor will modulate learning rates, promoting more learning for uncertain options.
  option_uncertainty_factor = 1.0 + (1.0 - jax.nn.sigmoid(jnp.abs(q_values - average_reward_estimate) * 5.0))

  # Update the Q-value for the chosen option using a standard Rescorla-Wagner rule.
  # The prediction error is simply the difference between the reward and the chosen option's Q-value.
  prediction_error = reward - q_values[choice]
  # Modulate the learning rate for the chosen option by the absolute magnitude of the prediction error
  # (surprise), the option's inherent uncertainty, AND its recency.
  # Also introduce a 'meta-learning' component where the learning rate itself adapts.
  chosen_prediction_error_salience_factor = 1.0 + jax.nn.softplus(jnp.abs(prediction_error))
  volatility_sensitive_learning_rate_scaling = 1.0 - 0.5 * jax.nn.sigmoid(reward_variance_estimate * 10.0) # Scales from 1.0 (low variance) to 0.5 (high variance)
  # New: Scale the chosen learning rate by the Q-value uncertainty for that option.
  # Higher uncertainty means a higher effective learning rate for the chosen option.
  uncertainty_scaled_learning_rate_factor = 0.5 + 1.5 * q_value_uncertainty[choice] # Scales from 0.5 (low uncertainty) to 2.0 (high uncertainty)

  # Introduce an 'omission salience' factor: stronger learning when a high Q-value results in a near-zero reward.
  # This factor becomes high when reward is near 0 and prediction error is strongly negative.
  omission_salience_factor = jnp.where(
      (reward < 0.1) & (prediction_error < -0.2), # Check for near-zero reward and significant negative prediction error
      1.0 + 3.0 * jax.nn.sigmoid(-prediction_error * 5.0), # Amplify learning rate for significant omissions, scales from 1 to 4
      1.0 # No amplification otherwise
  )
  effective_chosen_learning_rate = learning_rate * chosen_prediction_error_salience_factor * option_uncertainty_factor[choice] * volatility_sensitive_learning_rate_scaling * uncertainty_scaled_learning_rate_factor * omission_salience_factor
  effective_chosen_learning_rate = jnp.clip(effective_chosen_learning_rate, 0.0, 1.0) # Ensure rates are within bounds
  q_values = q_values.at[choice].set(
      q_values[choice] + effective_chosen_learning_rate * prediction_error
  )

  # For unchosen options, Q-values decay towards the average reward estimate.
  # This decay rate is modulated by the option's uncertainty and its recency,
  # and also by the global surprise signal from the chosen option's outcome.
  # Higher uncertainty, lower recency, and higher global surprise lead to faster decay.
  global_surprise_factor = 1.0 + jax.nn.softplus(jnp.abs(prediction_error)) # Global surprise based on chosen option's prediction error
  recency_amplified_decay_factor = 1.0 + (1.0 - recency_trace) * 0.5 # Amplify decay for less recent, scales from 1.0 to 1.5
  uncertainty_amplified_decay_factor = 1.0 + option_uncertainty_factor * 0.5 # Amplify decay based on higher option uncertainty
  global_surprise_amplified_decay_factor = 1.0 + global_surprise_factor * 0.5 # Amplify decay based on global surprise
  effective_unchosen_decay_rate = unchosen_learning_rate * recency_amplified_decay_factor * uncertainty_amplified_decay_factor * global_surprise_amplified_decay_factor
  effective_unchosen_decay_rate = jnp.clip(effective_unchosen_decay_rate, 0.0, 1.0) # Ensure rates are within bounds

  q_values = jnp.where(
      unchosen_mask,
      q_values * (1 - effective_unchosen_decay_rate) + average_reward_estimate * effective_unchosen_decay_rate,
      q_values,  # Chosen option's Q-value has already been updated.
  )

  # Update the perseveration trace for the chosen option. It increases towards 1.0
  # with a learning rate modulated by the prediction error (stronger for positive errors).
  # Modulate the perseveration learning rate based on the average reward estimate.
  # When average reward is low, perseveration learning rate is boosted, and vice-versa.
  average_reward_modulated_perseveration_lr_factor = 1.0 + jax.nn.tanh(0.5 - average_reward_estimate) # Scales from ~0.5 (high avg reward) to ~1.5 (low avg reward)
  chosen_perseveration_learning_rate = perseveration_learning_rate * jax.nn.sigmoid(prediction_error * 5) * average_reward_modulated_perseveration_lr_factor
  perseveration_trace_update_chosen = perseveration_trace[choice] + chosen_perseveration_learning_rate * (1 - perseveration_trace[choice])

  # For unchosen options, apply decay. The decay rate is modulated by prediction error,
  # causing faster decay when outcomes are surprising (higher uncertainty).
  # For unchosen options, apply decay. The decay rate is now modulated by the unchosen option's own
  # uncertainty (how far its Q-value is from the average reward estimate) AND the magnitude of the
  # prediction error for the chosen option, reflecting overall environmental surprise.
  jnp.abs(q_values - average_reward_estimate)
  # Stronger overall prediction error (surprise) can lead to faster decay of unchosen perseveration
  # For unchosen options, apply decay. The decay rate is modulated by:
  # 1. How 'bad' the unchosen option would have been (Q-value relative to current reward).
  # 2. The global context of average reward (faster decay in low-reward environments).
  # 3. The overall surprise (magnitude of chosen option's prediction error).
  # 4. The 'uncertainty' of the unchosen perseveration trace itself (how far it is from its initial value).

  # Factor based on how 'bad' the unchosen option might have been. Higher if reward was much better than unchosen Q.
  unchosen_opportunity_cost_factor = jax.nn.sigmoid((reward - q_values) * 5.0)

  # Factor based on the overall environmental rewardingness (amplify decay if average reward is below 0.5).
  global_average_reward_context_factor = 1.0 + jax.nn.relu(average_reward_estimate - 0.5)

  # Factor based on the global surprise from the chosen option's outcome.
  global_outcome_surprise_factor = 1.0 + jax.nn.softplus(jnp.abs(prediction_error))

  # Factor based on how 'uncertain' or 'outdated' the unchosen perseveration trace is.
  perseveration_uncertainty_factor = 1.0 + jax.nn.softplus(jnp.abs(perseveration_trace - initial_perseveration_value))

  # Modulate the perseveration decay rate by the recency trace for unchosen options.
  # Lower recency (i.e., less recently chosen) should lead to faster decay of perseveration.
  recency_modulated_decay_factor = 1.0 + (1.0 - recency_trace) * 2.0 # Scales from 1.0 (recency=1) to 3.0 (recency=0)

  effective_perseveration_decay_unchosen = perseveration_decay_rate * (
      1.0 + unchosen_opportunity_cost_factor + global_average_reward_context_factor
      + global_outcome_surprise_factor + perseveration_uncertainty_factor
  ) * recency_modulated_decay_factor
  effective_perseveration_decay_unchosen = jnp.clip(effective_perseveration_decay_unchosen, 0.0, 1.0) # Ensure rates are within bounds
  perseveration_trace_update_unchosen = perseveration_trace * (1 - effective_perseveration_decay_unchosen) + initial_perseveration_value * effective_perseveration_decay_unchosen

  # Update the perseveration trace by combining updates for chosen and unchosen options.
  perseveration_trace = jnp.where(
      unchosen_mask,
      perseveration_trace_update_unchosen,
      perseveration_trace_update_chosen,
  )

  # Update the recency trace. It increases for the chosen option and decays for unchosen options.
  # Modulate the recency trace update for the chosen option. A positive prediction error leads to stronger accumulation,
  # while a negative prediction error weakens it, or even promotes decay.
  recency_update_factor = 1.0 + jax.nn.tanh(prediction_error * 2.0) # Scales from ~0 to ~2
  reward_modulated_recency_learning_rate = perseveration_learning_rate * recency_update_factor
  reward_modulated_recency_learning_rate = jnp.clip(reward_modulated_recency_learning_rate, 0.0, 1.0)
  recency_trace_update_chosen = recency_trace[choice] + reward_modulated_recency_learning_rate * (1 - recency_trace[choice])
  # Modulate the recency trace decay for unchosen options based on their Q-value similarity to the chosen option.
  # Options with Q-values closer to the chosen option's Q-value decay slower.
  jax.nn.sigmoid(5.0 * (1.0 - jnp.abs(q_values - q_values[choice]))) # Higher for more similar Q-values, range 0-1
  # Decay rate is inverse of similarity, so higher similarity means lower decay rate
  # Modulate the recency trace decay for unchosen options based on their Q-value similarity to the chosen option
  # AND their inherent Q-value uncertainty. Higher uncertainty should lead to faster decay.
  q_value_similarity_to_chosen = jax.nn.sigmoid(5.0 * (1.0 - jnp.abs(q_values - q_values[choice]))) # Higher for more similar Q-values, range 0-1
  uncertainty_amplified_recency_decay_factor = 1.0 + q_value_uncertainty * 2.0 # Higher uncertainty -> faster decay, scales from 1.0 (uncertainty=0) to 3.0 (uncertainty=1)
  # Decay rate is inverse of similarity, so higher similarity means lower decay rate, but amplified by uncertainty.
  recency_decay_scaling_factor = (1.0 - 0.9 * q_value_similarity_to_chosen) * uncertainty_amplified_recency_decay_factor # Scales from ~0.1 to ~3.0
  effective_recency_decay_rate = perseveration_decay_rate * recency_decay_scaling_factor
  effective_recency_decay_rate = jnp.clip(effective_recency_decay_rate, 0.0, 1.0) # Ensure rates are within bounds
  recency_trace_update_unchosen = recency_trace * (1 - effective_recency_decay_rate)

  recency_trace = jnp.where(
      unchosen_mask,
      recency_trace_update_unchosen,
      recency_trace_update_chosen,
  )
  # Ensure the recency trace remains within reasonable bounds
  recency_trace = jnp.clip(recency_trace, 0.0, 1.0) # Bounds 0 to 1 as it's a 'trace'

  # Update the Q-value uncertainty trace.
  # For the chosen option, uncertainty decreases if prediction error is small (more certain),
  # and increases if prediction error is large (less certain).
  # For unchosen options, uncertainty slowly increases over time (decay towards maximum uncertainty).
  uncertainty_decay_rate = unchosen_learning_rate * 0.5 # A fraction of the unchosen learning rate for decay.
  # Modulate the uncertainty learning rate for the chosen option by the inverse of reward variance.
  # Lower variance (more stable environment) leads to a higher learning rate for uncertainty.
  volatility_sensitive_uncertainty_lr_scaling = 1.0 - 0.9 * jax.nn.sigmoid(reward_variance_estimate * 10.0) # Scales from 1.0 (low variance) to 0.1 (high variance)
  # For the chosen option, uncertainty decreases as new evidence is gathered.
  # The reduction in uncertainty is stronger when the prediction error is small,
  # indicating more reliable information gain.
  # If the prediction error is large, the reduction in uncertainty is smaller,
  # as the Q-value estimate was less accurate.
  # Scale this reduction by the volatility-sensitive learning rate.
  # For the chosen option, uncertainty decreases as new evidence is gathered.
  # The reduction in uncertainty is stronger when the prediction error is small,
  # indicating more reliable information gain.
  # If the prediction error is large, the reduction in uncertainty is smaller,
  # as the Q-value estimate was less accurate.
  uncertainty_reduction_magnitude = 1.0 - jax.nn.sigmoid(jnp.abs(prediction_error) * 3.0) # Larger reduction for smaller errors.
  # Scale this reduction by the volatility-sensitive learning rate.
  effective_uncertainty_reduction = uncertainty_reduction_magnitude * volatility_sensitive_uncertainty_lr_scaling

  # Calculate the actual change in uncertainty for the chosen option.
  # Predict the change in uncertainty. A simple prediction could be proportional to the effective_uncertainty_reduction.

  # Update the meta-level uncertainty prediction error estimate.
  # For the chosen option, uncertainty decreases as new evidence is gathered.
  # The reduction in uncertainty is stronger when the prediction error is small.
  # For the chosen option, uncertainty decreases as new evidence is gathered.
  # The reduction in uncertainty is stronger when the prediction error is small.
  # Predict the uncertainty *before* update for UPE calculation.
  predicted_uncertainty_after_update = q_value_uncertainty[choice] * (1.0 - effective_uncertainty_reduction)

  # Introduce a 'confidence bonus' for chosen options that lead to very low prediction error and high reward.
  # This makes the agent more certain about high-value, predictable options.
  confidence_bonus_factor = jnp.where(
      (jnp.abs(prediction_error) < 0.1) & (reward > 0.8), # Low error, high reward
      0.2, # Significant confidence boost (reduction in uncertainty)
      0.0 # No additional confidence bonus otherwise
  )
  uncertainty_update_chosen = jnp.maximum(0.0, predicted_uncertainty_after_update - confidence_bonus_factor) # Ensure uncertainty doesn't go below 0

  # Calculate Uncertainty Prediction Error (UPE) for the chosen option.
  # UPE is the difference between the actual updated uncertainty and the predicted uncertainty.
  # If the actual updated uncertainty is *lower* than predicted (i.e., less uncertain than expected), UPE is negative.
  # If actual updated uncertainty is *higher* than predicted (i.e., more uncertain than expected), UPE is positive.
  uncertainty_prediction_error = uncertainty_update_chosen - predicted_uncertainty_after_update

  # Update the meta-level uncertainty prediction error estimate.
  # Use a simple Rescorla-Wagner rule for UPE learning, modulated by the general uncertainty decay rate.
  upe_learning_rate = uncertainty_decay_rate * 2.0 # UPE learns faster
  uncertainty_prediction_error_estimate = uncertainty_prediction_error_estimate + upe_learning_rate * uncertainty_prediction_error

  # For unchosen options, decay uncertainty towards initial_q_value's uncertainty, modulated by reward variance.
  # If reward variance is high (unstable), uncertainty decays slower, reflecting persistent lack of knowledge.
  # For unchosen options, uncertainty slowly increases over time (decay towards maximum uncertainty of 1.0)
  # BUT it also decays towards a level of uncertainty inversely proportional to how distinct its Q-value is from the average.
  # If an option's Q-value is very distinct (far from average), its inherent uncertainty should be lower.
  # If an option's Q-value is close to the average, its inherent uncertainty should be higher (more ambiguous).
  inherent_uncertainty_from_distinctiveness = 1.0 - jax.nn.sigmoid(jnp.abs(q_values - average_reward_estimate) * 5.0) # Higher for less distinct Q-values.
  # Modulate the uncertainty decay rate by reward variance. In high variance, uncertainty decays slower for unchosen options.
  variance_modulated_uncertainty_decay_rate = uncertainty_decay_rate * (1.0 - jax.nn.sigmoid(reward_variance_estimate * 5.0))
  # The target uncertainty is a blend of inherent uncertainty and a slow decay towards a higher default (e.g., 0.8 to reflect lack of recent info).
  target_unchosen_uncertainty = 0.8 * inherent_uncertainty_from_distinctiveness + 0.2 * 1.0 # Bias towards 1.0 but influenced by distinctiveness
  uncertainty_update_unchosen = q_value_uncertainty * (1 - variance_modulated_uncertainty_decay_rate) + target_unchosen_uncertainty * variance_modulated_uncertainty_decay_rate

  q_value_uncertainty = jnp.where(
      unchosen_mask,
      uncertainty_update_unchosen,
      uncertainty_update_chosen,
  )
  q_value_uncertainty = jnp.clip(q_value_uncertainty, 0.0, 1.0)

  # Compute perseveration bias, modulated by value uncertainty, recency, and prediction error.
  # Higher uncertainty (Q-value closer to average) amplifies perseveration.
  value_uncertainty = 1.0 - jax.nn.sigmoid(jnp.abs(q_values - average_reward_estimate) * 5.0)
  recency_modulated_reward_sensitivity = perseveration_reward_sensitivity * recency_trace[choice]
  adaptive_perseveration_bias_strength = perseveration_bias_strength * (1.0 + recency_modulated_reward_sensitivity * jax.nn.sigmoid(prediction_error))
  base_perseveration_strength = perseveration_trace * adaptive_perseveration_bias_strength * (0.5 + 0.5 * recency_trace)
  perseveration_bias = jax.nn.softplus(base_perseveration_strength * (1.0 + value_uncertainty))

  relative_q_values = q_values - average_reward_estimate
  
  # Dynamically adjust the inverse temperature based on overall Q-value uncertainty and reward variance.
  # Higher uncertainty or variance leads to a lower effective inverse temperature (more exploration).
  # Dynamically adjust the inverse temperature based on overall Q-value uncertainty, reward variance,
  # AND the magnitude of uncertainty prediction error (UPE).
  # High UPE magnitude indicates unreliability in uncertainty estimation, promoting more exploration (lower temperature).
  upe_magnitude_scaling = 1.0 + jax.nn.softplus(jnp.abs(uncertainty_prediction_error_estimate) * 2.0) # Scales from 1.0 (low UPE) upwards
  adaptive_temperature_scaling_factor = 1.0 - 0.5 * jax.nn.sigmoid(jnp.mean(q_value_uncertainty) * 5.0 + reward_variance_estimate * 5.0 - 5.0) * upe_magnitude_scaling
  adaptive_temperature_scaling_factor = jnp.clip(adaptive_temperature_scaling_factor, 0.1, 2.0)
  adaptive_inverse_temperature = inverse_temperature * adaptive_temperature_scaling_factor

  # The option_specific_inverse_temperature now uses this adaptively scaled base temperature.
  # It also incorporates Q-value variance for further scaling.
  q_value_variance = jnp.var(q_values)
  variance_scaling_factor = 0.5 + 1.5 * jax.nn.sigmoid(q_value_variance)
  max_relative_q = jnp.max(relative_q_values)
  volatility_and_uncertainty_scaling = 1.0 - 0.5 * jax.nn.sigmoid(reward_variance_estimate * 5.0 + jnp.mean(q_value_uncertainty) * 5.0 - 5.0)
  volatility_and_uncertainty_scaling = jnp.clip(volatility_and_uncertainty_scaling, 0.1, 2.0)
  option_specific_inverse_temperature = adaptive_inverse_temperature * variance_scaling_factor * (jax.nn.relu(1 + (relative_q_values - max_relative_q)) + 0.01) * volatility_and_uncertainty_scaling

  # Recalculate Q-value entropy using the new adaptive_inverse_temperature for consistency.
  q_value_softmax_probs = jax.nn.softmax(q_values * adaptive_inverse_temperature)
  q_value_entropy = -jnp.sum(q_value_softmax_probs * jnp.log(q_value_softmax_probs + 1e-9))
  normalized_entropy = q_value_entropy / jnp.log(4.0)
  entropy_modulated_perseveration_factor = 0.5 + normalized_entropy

  global_surprise_signal = jax.nn.sigmoid(jnp.abs(prediction_error) * 2.0)
  global_surprise_perseveration_factor = 1.0 - 0.5 * global_surprise_signal
  modulated_perseveration_bias = perseveration_bias * entropy_modulated_perseveration_factor * global_surprise_perseveration_factor

  # Incorporate an exploration bonus based on Q-value uncertainty and reward variance.
  exploration_bonus = jnp.mean(q_value_uncertainty) * jnp.mean(q_value_uncertainty)
  option_exploration_bonus = (q_value_uncertainty + reward_variance_estimate) * exploration_bonus
  # Add a general exploration bonus that doesn't rely on uncertainty, potentially encouraging more diverse choices.
  # This bonus is higher for options that have not been chosen recently.
  general_exploration_bonus = exploration_bonus_strength * (1.0 - recency_trace)
  choice_logits = relative_q_values * option_specific_inverse_temperature + perseveration_trace * modulated_perseveration_bias + option_exploration_bonus + general_exploration_bonus

  agent_state = jnp.concatenate([q_values, perseveration_trace, jnp.array([average_reward_estimate]), recency_trace, jnp.array([reward_variance_estimate]), q_value_uncertainty, jnp.array([uncertainty_prediction_error_estimate]), jnp.array([exploration_bonus_strength])])

  return choice_logits, agent_state


# SOO human_bandit even/odd split.
def human_bandit_run2_fit_only(
    params: chex.Array,
    choice: int,
    reward: float,
    agent_state: Optional[chex.Array],
) -> tuple[chex.Array, chex.Array]:
  """Cognitive model describing human behavior on a multi-armed bandit task.

  Assumes the agent is presented with four options on each trial.

  Args:
    params: Fit parameters of the model. Different parameters are used for
      different subjects.
    choice: Choice made by the agent on the previous trial. 0, 1, 2, or 3
    reward: Reward received by the agent on the previous trial. A float between
      0 and 1.
    agent_state: The current state of the cognitive model.

  Returns:
    choice_logits: The probabilities that the agent will choose option 0, 1, 2,
      or 3 on the next trial, expressed as logits.
    agent_state: New agent state
  """
  # Do not remove comments or TODOs from this program.

  # TODO: Initialize agent state. NOLINT
  if agent_state is None:
    # The agent state includes:
    # 0-3: Q-values for each of the 4 arms, initialized with an adaptive optimistic bias.
    # 4: The learning rate (for consistency, though a parameter).
    # 5-8: Individual choice stickiness values for each of the 4 arms.
    # 9: The overall average reward observed so far.
    # 10: The dynamic inverse temperature (beta).
    # 11-14: Individual 'surprise' values for each arm, initialized small and non-zero.
    # 15: The maximum observed reward so far (for adaptive reward sensitivity).
    # 16: The running sum of squared rewards for variance calculation.
    # 17: The count of observed rewards for variance calculation.
    # 18-21: last_reward_trace, storing the last observed reward for each arm, initialized to neutral.
    # 22-25: per_arm_sum_sq_rewards, for variance calculation, initialized to 0.0.
    # 26-29: per_arm_reward_counts, for variance calculation, initialized to 0.0.
    # Initialize Q-values with an adaptive optimistic bias.
    # Initial max_observed_reward is set to 1.0 to ensure an optimistic start.
    initial_max_observed_reward = 1.0
    initial_q_value = jnp.clip(initial_max_observed_reward + 0.1, 0.75, 1.0) # Ensure it's not too low but also capped
    agent_state = jnp.array([initial_q_value, initial_q_value, initial_q_value, initial_q_value, 0.1, 0.0, 0.0, 0.0, 0.0, 0.5, 2.0, 0.1, 0.1, 0.1, 0.1, initial_max_observed_reward, 0.0, 0.0, 0.5, 0.5, 0.5, 0.5, 0.0, 0.0, 0.0, 0.0, 0.0, 0.0, 0.0, 0.0])

  # TODO: Define all parameters. NOLINT
  # Each parameter assignment should be a single line.
  # Change the names of the parameters to be descriptive.
  # Include comments describing the parameter transformation, if applicable.
  # Baseline inverse temperature parameter for the softmax choice rule, ensuring positivity
  # and numerical stability using softplus.
  baseline_beta = jax.nn.softplus(params[0])
  # Base learning rate for updating expected rewards (softplus to ensure positivity and allow for wider range).
  base_learning_rate = jax.nn.softplus(params[1])
  # Base sensitivity to rewards (softplus to ensure positivity).
  base_reward_sensitivity = jax.nn.softplus(params[2])
  # Persistence of previous choice bias (softplus to ensure positivity).
  choice_stickiness_bias = jax.nn.softplus(params[3])
  # Decay rate for choice stickiness bias, ensuring values between 0 and 1.
  choice_stickiness_decay_rate = jax.nn.sigmoid(params[4])
  # General decay rate for unchosen Q-values (sigmoid to ensure values between 0 and 1).
  unchosen_q_decay_rate = jax.nn.sigmoid(params[5])
  # Learning rate for updating the average reward (sigmoid to ensure values between 0 and 1).
  # Dynamically linked to beta_learning_rate to ensure coordinated adaptation to environmental volatility.
  average_reward_learning_rate = jax.nn.sigmoid(params[6])
  # Learning rate for choice stickiness, to allow it to adapt (sigmoid to ensure values between 0 and 1).
  stickiness_learning_rate = jax.nn.sigmoid(params[6])
  # Learning rate for updating the inverse temperature (beta) based on prediction error (sigmoid to ensure values between 0 and 1).
  beta_learning_rate = jax.nn.sigmoid(params[7])
  # Adjust the average_reward_learning_rate by beta_learning_rate, making it sensitive to volatility.
  average_reward_learning_rate = jnp.clip(average_reward_learning_rate + beta_learning_rate * 0.5, 0.01, 0.99)
  # Learning rate for updating surprise values (sigmoid to ensure values between 0 and 1).
  surprise_learning_rate = jax.nn.sigmoid(params[8])
  # Factor by which prediction error influences stickiness updates, ensuring positivity.
  pe_stickiness_influence = jax.nn.softplus(params[9])

  # Update the average reward for the overall task environment.
  # This acts as a dynamic baseline for reward expectations.
  current_average_reward = agent_state[9]
  average_reward_prediction_error = reward - current_average_reward
  new_average_reward = current_average_reward + average_reward_learning_rate * average_reward_prediction_error
  agent_state = agent_state.at[9].set(new_average_reward)

  # Update the expected reward for the chosen option using a Rescorla-Wagner rule, and decay all options.
  # Create a mask for the chosen arm.
  choice_mask = jax.nn.one_hot(choice, num_classes=4)
  # Apply a general decay to all Q-values towards the current average reward, dynamically adapting the baseline.
  # This prevents values from drifting indefinitely and simulates forgetting, aligning expectations with recent experience.
  general_q_decay_rate = 0.01
  decayed_q_values = agent_state[:4] * (1.0 - general_q_decay_rate) + new_average_reward * general_q_decay_rate
  # For unchosen options, apply an additional decay towards the current average reward.
  # The effective decay rate for unchosen options is modulated by surprise and a recency factor.
  effective_decay_rates_for_unchosen = jnp.clip(
      unchosen_q_decay_rate / (1.0 + agent_state[11:15] * surprise_learning_rate) + 0.05,
      0.01, 0.99)

  # The maximum observed reward. This helps in adaptively scaling reward sensitivity.
  max_observed_reward = agent_state[15]
  # Running sum of squared rewards and count of observed rewards for variance calculation.
  sum_sq_rewards = agent_state[16]
  reward_count = agent_state[17]

  # Update max_observed_reward.
  max_observed_reward = jnp.maximum(max_observed_reward, reward)
  agent_state = agent_state.at[15].set(max_observed_reward)

  # Update sum of squared rewards and count for variance calculation.
  new_sum_sq_rewards = sum_sq_rewards + reward**2
  new_reward_count = reward_count + 1
  agent_state = agent_state.at[16].set(new_sum_sq_rewards)
  agent_state = agent_state.at[17].set(new_reward_count)

  # Calculate reward variance.
  # Use a small epsilon to avoid division by zero if reward_count is 0 or 1.
  mean_reward = new_average_reward
  # Variance = E[X^2] - (E[X])^2
  reward_variance = jnp.maximum(0.0, (new_sum_sq_rewards / (new_reward_count + 1e-6)) - mean_reward**2)
  # Dynamically adjust reward_sensitivity based on reward variance.
  dynamic_reward_sensitivity = base_reward_sensitivity / (1.0 + reward_variance)
  # Ensure sensitivity remains within reasonable bounds.
  dynamic_reward_sensitivity = jnp.clip(dynamic_reward_sensitivity, 0.1, 10.0)

  # Update per-arm reward statistics for variance calculation.
  per_arm_sum_sq_rewards = agent_state[22:26]
  per_arm_reward_counts = agent_state[26:30]
  new_per_arm_sum_sq_rewards = per_arm_sum_sq_rewards.at[choice].add(reward**2)
  new_per_arm_reward_counts = per_arm_reward_counts.at[choice].add(1)
  agent_state = agent_state.at[22:26].set(new_per_arm_sum_sq_rewards)
  agent_state = agent_state.at[26:30].set(new_per_arm_reward_counts)

  # Calculate per-arm reward variance.
  # Use a small epsilon to avoid division by zero.
  # For consistency, let's calculate per-arm mean more robustly based on past rewards for that arm.
  # If an arm hasn't been chosen, its mean is the global average.
  # Using current Q-values as the best estimate of mean for variance calc.
  per_arm_variance = jnp.maximum(0.0, (new_per_arm_sum_sq_rewards / (new_per_arm_reward_counts + 1e-6)) - agent_state[:4]**2)
  # Ensure variance values are within a reasonable range.
  per_arm_variance = jnp.clip(per_arm_variance, 0.01, 1.0) # Lower bound to avoid infinite learning rate.

  # Calculate a volatility signal based on reward variance.
  # A higher reward variance indicates a more volatile environment.
  # We use a sigmoid to scale the variance into a bounded volatility signal between 0 and 1.
  # The factor of 5.0 scales the variance, and -1.0 shifts it so that typical variances
  # result in a non-zero volatility.
  environmental_volatility = jax.nn.sigmoid(reward_variance * 5.0 - 1.0)

  # The reward prediction error is the difference between the actual reward, scaled by dynamic sensitivity, and the expected reward.
  # Normalize the reward to be between 0 and 1 using the maximum observed reward,
  # preventing division by zero with a small epsilon.
  normalized_reward = reward / (max_observed_reward + 1e-6)
  # Calculate raw prediction error.
  raw_prediction_error = (normalized_reward * dynamic_reward_sensitivity) - agent_state[choice]
  # Introduce precision weighting: prediction error is scaled by the inverse of the surprise
  # for the chosen arm. High surprise implies low precision, reducing the PE's impact.
  # We use a more direct certainty proxy: a decay of past surprise values, where low surprise means high certainty.
  # This term (1 - agent_state[11 + choice]) represents 'certainty' for the chosen arm.
  # High surprise (agent_state[11 + choice]) means low certainty.
  certainty_for_chosen_arm = jnp.clip(1.0 - agent_state[11 + choice], 0.1, 1.0)
  # Inverse certainty represents uncertainty, which will be used to modulate learning.

  # Dynamic RPE sensitivity modulation based on surprise and reward sign.

  # Incorporate certainty and surprise directly into the prediction error.
  # Less certain predictions lead to smaller effective PEs.
  # Higher surprise amplifies the prediction error, making unexpected outcomes more impactful.
  surprise_amplification_factor = jnp.clip(jax.nn.softplus(agent_state[11 + choice] * 2.0), 1.0, 3.0)
  prediction_error = raw_prediction_error * certainty_for_chosen_arm * surprise_amplification_factor

  # Asymmetric decay for unchosen Q-values based on prediction error for the chosen arm.
  # Decay unchosen Q-values. Chosen Q-values are updated separately.
  # For unchosen options, decay their current values (which already have the general decay applied)
  # further towards the new_average_reward, modulated by surprise and recency.
  # For unchosen options, decay their current values further towards the new_average_reward,
  # modulated by the effective decay rate, and also towards a neutral baseline (0.5).
  unchosen_q_values_decay_to_average = (decayed_q_values * (1.0 - effective_decay_rates_for_unchosen)) + \
                                       (new_average_reward * effective_decay_rates_for_unchosen)
  # Additionally, introduce a small, constant decay towards a neutral baseline (0.5) for unchosen arms.
  unchosen_q_values_decay_to_average = (unchosen_q_values_decay_to_average * (1.0 - 0.005)) + (0.5 * 0.005)
  # New: Introduce a small, constant decay towards the minimum possible reward (0.0) for unchosen arms.
  # This acts as a 'pessimistic' decay for options not recently experienced, encouraging exploration.
  # The decay rate is also modulated by environmental volatility: higher volatility means faster decay towards 0.0.
  # We now decay unchosen Q-values towards a blend of 0.0 and the current average reward,
  # with the blend ratio modulated by environmental volatility.
  pessimistic_forgetting_rate_base = 0.002
  pessimistic_forgetting_rate_volatility_mod = environmental_volatility * 0.015 # Increased sensitivity to volatility
  pessimistic_forgetting_rate = pessimistic_forgetting_rate_base + pessimistic_forgetting_rate_volatility_mod

  # Target for pessimistic decay: blend between 0.0 (fully pessimistic) and new_average_reward.
  # Also, adjust this target based on prediction error for a "contextual pessimism".
  # If PE is negative, target is more pessimistic (closer to 0.0). If PE is positive, target is less pessimistic (closer to new_average_reward).
  # Adjust the pessimistic decay target based on the sign and magnitude of the prediction error.
  # If PE is negative, the target shifts more strongly towards 0.0 (more pessimistic).
  # If PE is positive, the target shifts less towards 0.0, allowing unchosen options to retain some value.
  pe_driven_pessimism_shift = jax.lax.cond(
      prediction_error < 0, # If the chosen option was worse than expected
      lambda pe: jnp.clip(jnp.abs(pe) * 0.75 * (1.0 + environmental_volatility), 0.0, 0.75), # Stronger shift towards 0.0, especially in volatile environments
      lambda pe: jnp.clip(pe * -0.5, -0.5, 0.0), # Slightly shift away from 0.0 for positive PE
      prediction_error
  )
  # The pessimistic decay target is a blend of 0.0 (fully pessimistic) and new_average_reward.
  # This blend is now heavily modulated by `pe_driven_pessimism_shift` and `environmental_volatility`.
  pessimistic_decay_target = (0.0 * (environmental_volatility + pe_driven_pessimism_shift)) + \
                             (new_average_reward * (1.0 - environmental_volatility - pe_driven_pessimism_shift))
  pessimistic_decay_target = jnp.clip(pessimistic_decay_target, 0.0, 0.75) # Ensure target is within reasonable bounds
  unchosen_q_values_decay_to_average = (unchosen_q_values_decay_to_average * (1.0 - pessimistic_forgetting_rate)) + \
                                       (pessimistic_decay_target * pessimistic_forgetting_rate)

  # Introduce a decay for unchosen Q-values back towards their initial, optimistic state.
  # This encourages re-exploration of neglected options.
  decay_to_initial_q_rate = 0.005 # Small constant rate for decaying towards initial Q.
  # The initial Q-value is the first element of the agent_state (all initial Qs are the same).
  # We use the initial Q-value from the agent_state, which is set at initialization.
  # The very first element `agent_state[0]` represents `initial_q_value`.
  decay_to_initial_q_value = agent_state[0]
  unchosen_q_values_decay_to_average = (unchosen_q_values_decay_to_average * (1.0 - decay_to_initial_q_rate)) + \
                                       (decay_to_initial_q_value * decay_to_initial_q_rate)

  # Calculate a weighted average of new_average_reward and a more pessimistic baseline (e.g., min observed reward or 0.25).
  # This serves as a dynamic, context-sensitive "neutral" point for decay.
  # Use environmental volatility to determine the weighting: higher volatility means more weight on a pessimistic baseline.
  dynamic_neutral_q_decay_target = (new_average_reward * (1.0 - environmental_volatility * 0.5)) + \
                                   (0.25 * environmental_volatility * 0.5) # A slightly pessimistic fixed baseline if volatile
  # Introduce a small, constant decay towards this new dynamic_neutral_q_decay_target for unchosen arms.
  unchosen_q_values_decay_to_average = (unchosen_q_values_decay_to_average * (1.0 - 0.005)) + (dynamic_neutral_q_decay_target * 0.005)

  # Introduce a small, inverse influence of prediction error from the chosen arm on unchosen arms.
  # If chosen was better than expected (positive PE), unchosen arms become slightly less attractive.
  unchosen_pe_influence = -0.1 * prediction_error

  # Goal-directed re-evaluation for unchosen options: if the chosen option was highly surprising,
  # re-evaluate unchosen options based on the sign of the prediction error.
  # If chosen was surprisingly good, unchosen options get a small boost.
  # If chosen was surprisingly bad, unchosen options get a small reduction.
  # This effect is scaled by the 'surprise' of the chosen arm and its certainty.
  surprise_re_evaluation_factor = jnp.clip(agent_state[11 + choice] * certainty_for_chosen_arm * 0.5, 0.0, 1.0) # Scale by surprise and certainty
  goal_directed_re_evaluation_update = prediction_error * surprise_re_evaluation_factor

  # Introduce a "forgetting" mechanism for unchosen options, decaying them towards the neutral baseline (0.5).
  # The decay rate is proportional to how long it's been since the arm was chosen (implicit in current Q-value).
  # We use a small, fixed forgetting rate for simplicity.
  forgetting_rate = 0.005 # Small constant forgetting rate for unchosen arms
  
  # Context-Dependent Revaluation of Unchosen Options:
  # If the chosen option leads to a highly surprising positive PE, it could signal a generally richer environment,
  # leading to a subtle boost in unchosen Q-values.
  # If a highly surprising negative PE, it signals a poorer environment, subtly reducing unchosen Q-values.
  contextual_revaluation_for_unchosen = jax.lax.cond(
      jnp.abs(prediction_error) > 0.3, # Only if PE is sufficiently large (highly surprising)
      lambda pe: jnp.clip(pe * surprise_re_evaluation_factor * environmental_volatility * 0.2, -0.1, 0.1),
      lambda pe: 0.0,
      prediction_error
  )
  unchosen_q_values_decay_to_average = unchosen_q_values_decay_to_average * (1.0 - forgetting_rate) + \
                                       0.5 * forgetting_rate + \
                                       goal_directed_re_evaluation_update * (1 - choice_mask) + \
                                       contextual_revaluation_for_unchosen * (1 - choice_mask)
  
  # Incorporate volatility-sensitive soft reset for Q-values.
  # If environmental volatility is high, push Q-values more strongly towards the average reward.
  # This acts as a more aggressive forgetting mechanism for outdated expectations in dynamic environments.
  q_reset_strength = environmental_volatility * 0.1 # Max reset strength of 10% towards average
  # Soft reset for all Q-values, modulated by volatility.
  decayed_q_values = (decayed_q_values * (1.0 - q_reset_strength)) + (new_average_reward * q_reset_strength)

  # Incorporate the decaying last_reward_trace into Q-value updates for all arms.
  # This provides a memory of recent rewards, even for unchosen options.
  last_reward_trace_decay_rate = 0.1 # Small decay for the reward trace
  # Scale by environmental_volatility: in volatile environments, recent rewards are more important.
  effective_trace_decay = last_reward_trace_decay_rate * (1.0 + environmental_volatility * 0.5)
  # Update last_reward_trace: for the chosen arm, it's the current reward. For unchosen, it decays.
  current_last_reward_trace = agent_state[18:22]
  # Update last_reward_trace: for the chosen arm, it's the current reward. For unchosen, it decays.
  updated_last_reward_trace = (current_last_reward_trace * (1.0 - effective_trace_decay)) + \
                              (normalized_reward * choice_mask)
  # Ensure the trace values are bounded.
  updated_last_reward_trace = jnp.clip(updated_last_reward_trace, 0.0, 1.0)
  agent_state = agent_state.at[18:22].set(updated_last_reward_trace)

  # Combine the generally decayed Q-values with unchosen specific decay and PE influence.
  # The chosen arm's Q-value maintains the general decay, others get the specific decay + PE influence.
  decayed_q_values_for_update = decayed_q_values * choice_mask + \
                                (unchosen_q_values_decay_to_average + unchosen_pe_influence + updated_last_reward_trace * 0.05) * (1 - choice_mask)
  # Dynamically adjust the learning rate based on the surprise for the chosen arm AND the reward magnitude.
  # A higher surprise means a higher effective learning rate for this specific update.
  # Additionally, larger rewards can boost the learning rate to reflect higher salience.
  # We use a softplus to ensure the surprise-modulated learning rate is always positive.
  # The 'surprise_learning_rate' parameter acts as a gain for how much surprise influences the learning rate.
  # We clip the effective learning rate to prevent extreme values.
  # We add a term proportional to the reward magnitude to the learning rate, scaled by a factor.
  # Introducing a salience factor: rewards that deviate significantly from the average are more salient.
  # Dynamically adjust the learning rate based on surprise, certainty, global reward variance, and prediction error magnitude.
  # Higher surprise, reward variance, or prediction error can increase the learning rate.
  effective_learning_rate = jnp.clip(
      base_learning_rate # Base learning rate
      # Modulate the learning rate by the absolute prediction error magnitude, scaled by a softplus.
      # This ensures a positive and non-linear boost based on how surprising the outcome was.
      # Modulate the learning rate by prediction error magnitude, with asymmetry for negative PEs.
      # Negative PEs (worse than expected) lead to a larger boost in learning rate.
      + jax.lax.cond(
          prediction_error < 0,
          lambda pe: jax.nn.softplus(jnp.abs(pe) * 1.0) * 0.4, # Stronger boost for negative PEs
          lambda pe: jax.nn.softplus(jnp.abs(pe) * 0.5) * 0.2, # Smaller boost for positive PEs
          prediction_error
      )
      # Volatility-gated learning rate: scale the effective learning rate by environmental volatility.
      # This ensures the agent learns faster in volatile environments and slower in stable ones.
      # We use a softplus to ensure the influence is positive and then scale it.
      + jax.nn.softplus(environmental_volatility * 0.5) * 0.2 # Direct influence of volatility
  )
  effective_learning_rate = jnp.clip(effective_learning_rate, 0.01, 0.99)

  # Apply asymmetric learning rates: learn more from negative prediction errors to promote faster adaptation to worse-than-expected outcomes.
  # This is a common finding in reinforcement learning and psychological studies.
  # The boost for negative PEs is now dynamically scaled by certainty: a certain negative PE is more impactful.
  effective_learning_rate_q_update = jax.lax.cond(
      prediction_error < 0, # If prediction error is negative (worse than expected)
      lambda pe: jnp.clip(effective_learning_rate * (1.0 + jnp.abs(pe) * 1.5 * certainty_for_chosen_arm), 0.01, 0.99), # Boost learning rate significantly for negative PEs, scaled by certainty
      lambda pe: jnp.clip(effective_learning_rate * (1.0 + jnp.abs(pe) * 0.25), 0.01, 0.99), # Smaller boost for positive PEs
      prediction_error
  )

  # Introduce state-dependent learning rate modulation.
  # If the current Q-value is extreme (near 0 or 1), the learning rate is boosted more if the prediction error is large.
  # This makes learning more sensitive when expectations are strong (either very good or very bad).
  # The factor increases as Q-value deviates from 0.5 (neutral expectation).
  # Dynamically modulate the learning rate based on prediction error sign, magnitude, and current Q-value of the chosen arm.
  # This introduces asymmetric learning based on how "surprising" the outcome is given the current belief.
  # q_value_magnitude is the current expectation for the chosen arm.
  # If PE is positive (better than expected):
  #   - If Q is low (highly disconfirming), boost learning.
  #   - If Q is high (confirmatory), slightly dampen learning.
  # If PE is negative (worse than expected):
  #   - If Q is high (highly disconfirming), boost learning significantly.
  #   - If Q is low (confirmatory), slightly dampen learning.
  # Scale by current Q-value's distance from 0.5 (more uncertain, more adaptable).
  # `prediction_error` is already certainty-modulated.

  # The final learning rate is a combination of the base rate, asymmetric PE influence,
  # and additional modulation from certainty and environmental volatility.
  # Simplified to remove the conceptual meta-learning thoughts and directly compute the final learning rate.
  # Dynamically adjust the final learning rate based on environmental volatility.
  # In highly volatile environments, the learning rate should be further boosted.
  # The boost is non-linear and more pronounced for large prediction errors, especially positive ones,
  # as these might signal new opportunities in an uncertain context.
  volatility_boost_factor = jax.lax.cond(
      prediction_error > 0,
      # For positive PEs, volatility amplifies learning significantly, modulated by surprise.
      lambda pe: jax.nn.softplus(pe * environmental_volatility * (1.0 + agent_state[11 + choice])) * 0.3,
      # For negative PEs, volatility still amplifies learning, but less steeply than positive PEs.
      lambda pe: jax.nn.softplus(jnp.abs(pe) * environmental_volatility * 0.5) * 0.2,
      prediction_error
  )

  # Incorporate arm-specific uncertainty into the final learning rate.
  # Options with higher reward variance (more uncertain) should have a boosted learning rate for their updates.
  # We use a softplus to ensure the influence is positive and non-linear, scaled by a factor.
  arm_specific_uncertainty_factor = jax.nn.softplus(per_arm_variance[choice] * 2.0) * 0.1

  final_learning_rate = jnp.clip(
      effective_learning_rate_q_update # Base learning rate already has asymmetric PE influence.
      + (jnp.abs(prediction_error) * certainty_for_chosen_arm * 0.5) # Stronger learning for certain large PEs.
      + volatility_boost_factor # Incorporate the volatility-based boost.
      + arm_specific_uncertainty_factor # Add the arm-specific uncertainty boost.
      # Add a boost based on the salience of the current reward.
      # Rewards that deviate significantly from the average are more salient.
      # We use jnp.abs(reward - new_average_reward) to capture deviations in both directions.
      + jnp.clip(jnp.abs(reward - new_average_reward) * 0.75 * base_reward_sensitivity, 0.0, 0.5),
      0.01, 0.99
  )
  # Update the chosen Q-value using the Rescorla-Wagner rule with the final,
  # asymmetrically adjusted, and certainty-modulated learning rate.
  updated_chosen_q = decayed_q_values_for_update[choice] + final_learning_rate * prediction_error
  # Combine decayed Q-values for unchosen arms and updated Q-value for the chosen arm.
  new_q_values = decayed_q_values_for_update * (1 - choice_mask) + updated_chosen_q * choice_mask
  agent_state = agent_state.at[:4].set(new_q_values)

  # Update the dynamic inverse temperature (beta) based on prediction error.
  # Larger absolute prediction errors decrease beta (more exploration), smaller errors increase beta (more exploitation).
  current_beta = agent_state[10]
  # Dynamically adjust the beta learning rate based on the absolute prediction error AND prediction certainty,
  # AND environmental volatility. In more volatile environments, beta should adjust more rapidly.
  effective_beta_learning_rate = beta_learning_rate * (1.0 + jnp.abs(prediction_error) * certainty_for_chosen_arm + environmental_volatility * 0.5)
  # Adjust beta_update_signal: larger absolute prediction errors (higher surprise) should lead to more exploration (lower beta).
  # The update signal will now be directly proportional to the absolute prediction error.
  # Introduce asymmetric beta modulation: negative PEs cause a larger drive towards exploration.
  beta_update_signal_positive_pe = jnp.abs(prediction_error) * (1.0 + agent_state[11 + choice] * 2.0)
  beta_update_signal_negative_pe = jnp.abs(prediction_error) * (1.0 + agent_state[11 + choice] * 3.0) # Stronger signal for negative PEs
  beta_update_signal = jax.lax.cond(
      prediction_error < 0,
      lambda: beta_update_signal_negative_pe,
      lambda: beta_update_signal_positive_pe,
  )
  # Influence from reward variance: higher variance drives beta down (more exploration).
  variance_influence_on_beta = 0.75 * jax.nn.softplus(reward_variance - 0.4)

  # Average surprise across all arms. Higher average surprise should decrease beta (more exploration).
  average_surprise_across_arms = jnp.mean(agent_state[11:15])
  surprise_exploration_influence = average_surprise_across_arms * 2.0

  # Update beta: larger beta_update_signal (from larger PE) now DECREASES beta (more exploration).
  # Smaller beta_update_signal (from smaller PE) allows beta to increase (more exploitation).
  new_beta = current_beta + effective_beta_learning_rate * (1.0 - beta_update_signal) - variance_influence_on_beta - surprise_exploration_influence
  # Decay beta towards the baseline_beta, using beta_learning_rate.
  # The decay rate is now also modulated by environmental volatility and average surprise.
  # Higher volatility or surprise slows down the decay towards baseline_beta, maintaining exploration.
  beta_decay_modulation = (1.0 - environmental_volatility * 0.2 - average_surprise_across_arms * 0.1)
  # Ensure modulation keeps decay rate positive.
  effective_beta_decay_rate = jnp.clip(beta_learning_rate * beta_decay_modulation, 0.01, 0.99)
  new_beta = (new_beta * (1 - effective_beta_decay_rate)) + (baseline_beta * effective_beta_decay_rate)
  # Ensure beta remains positive and within a reasonable range.
  new_beta = jnp.clip(new_beta, 0.1, 10.0)
  agent_state = agent_state.at[10].set(new_beta)

  # Update the surprise for each arm.
  current_surprise_values = agent_state[11:15]
  # Decay all surprise values.
  updated_surprise_values = current_surprise_values * (1 - surprise_learning_rate)
  # Increase surprise for the chosen arm based on the direction and magnitude of the prediction error,
  # modulated by the certainty of the prediction for that arm.
  # Positive prediction errors might lead to a different surprise update than negative ones.
  # Here, we introduce an asymmetry: positive PEs increase surprise more than negative PEs of the same magnitude.

  # Surprise update based on the magnitude of prediction error, with asymmetry for negative PEs.
  # Negative prediction errors (worse than expected) cause a larger increase in surprise.
  surprise_update_magnitude_positive = surprise_learning_rate * prediction_error * 0.5 # Smaller impact for positive PEs
  surprise_update_magnitude_negative = surprise_learning_rate * jnp.abs(prediction_error) * 1.5 # Larger impact for negative PEs

  # Apply asymmetric scaling to the surprise update: negative PEs are more surprising.
  asymmetric_surprise_update_signal = jax.lax.cond(
      raw_prediction_error < 0,
      lambda: surprise_update_magnitude_negative,
      lambda: surprise_update_magnitude_positive,
  )

  # Modulate the surprise update by the certainty of the prediction for that arm.
  # If the agent was highly certain (high `certainty_for_chosen_arm`) and still
  # made a large prediction error, this should lead to a stronger update of surprise.
  # We use raw_prediction_error here to ensure surprise is sensitive to the full magnitude of the unexpected outcome.
  certainty_modulated_surprise_update = asymmetric_surprise_update_signal * certainty_for_chosen_arm
  updated_surprise_values = updated_surprise_values.at[choice].set(
      jnp.clip(current_surprise_values[choice] + certainty_modulated_surprise_update, 0.0, 1.0)
  )
  agent_state = agent_state.at[11:15].set(updated_surprise_values)

  # Update choice stickiness values for each arm.
  current_stickiness_values = agent_state[5:9]
  choice_one_hot = jax.nn.one_hot(choice, num_classes=4)
  
  # All stickiness values decay slightly towards zero, and apply a volatility-sensitive soft reset.
  stickiness_reset_strength = environmental_volatility * 0.2
  updated_stickiness_values = current_stickiness_values * (1 - stickiness_learning_rate) * (1.0 - stickiness_reset_strength)

  # Apply decay to the choice stickiness bias.
  # Base update for the chosen arm, incorporating bias and prediction error.
  # We now also consider the prior action when updating stickiness for the chosen arm.
  # If the same choice was made on the previous trial, it reinforces stickiness.
  prior_action_reinforcement = jax.lax.cond(
      choice == jnp.argmax(agent_state[5:9]), # If the current choice was also the most sticky prior
      lambda: 0.5, # Boost stickiness for repeating a previously sticky action
      lambda: 0.0 # No extra boost otherwise
  )
  # Dynamically modulate pe_stickiness_influence: stronger when Q-value is neutral, weaker when extreme.
  q_value_uncertainty_factor = jnp.clip(1.0 - jnp.abs(new_q_values[choice] - 0.5) * 2.0, 0.1, 1.0) # Peaks at 0.5, lower at extremes
  modulated_pe_stickiness_influence = pe_stickiness_influence * q_value_uncertainty_factor
  # Dynamically adjust the choice_stickiness_bias based on the prediction error.
  # Positive PE amplifies the bias, negative PE attenuates or reverses it.
  pe_modulated_stickiness_bias = jax.lax.cond(
      prediction_error > 0,
      lambda pe: choice_stickiness_bias * (1.0 + pe * 0.5), # Amplify bias for positive PE
      lambda pe: choice_stickiness_bias * (1.0 - jnp.abs(pe) * 1.0), # Attenuate/reverse bias for negative PE
      prediction_error
  )
  base_chosen_stickiness_update = pe_modulated_stickiness_bias + modulated_pe_stickiness_influence * prediction_error + prior_action_reinforcement

  # Update for chosen arm, including a direct reduction for negative prediction errors.
  # If prediction error is negative, reduce stickiness to encourage switching.
  stickiness_reduction_for_negative_pe = jax.lax.cond(
      prediction_error < 0,
      # Stronger reduction if certain about the bad outcome and in a volatile environment, scaled by Q-value uncertainty.
      lambda pe: stickiness_learning_rate * jnp.abs(pe) * (1.0 + certainty_for_chosen_arm) * (1.0 + environmental_volatility * 0.5) * q_value_uncertainty_factor,
      lambda pe: 0.0,
      prediction_error
  )
  updated_stickiness_values = updated_stickiness_values.at[choice].set(
      current_stickiness_values[choice] + stickiness_learning_rate * base_chosen_stickiness_update - stickiness_reduction_for_negative_pe
  )
  # Ensure stickiness reduction doesn't make stickiness go below zero for the chosen arm.
  updated_stickiness_values = updated_stickiness_values.at[choice].set(
      jnp.maximum(0.0, updated_stickiness_values[choice])
  )

  # Introduce an "implicit negative learning" for unchosen options when the chosen option yields a positive PE.
  # If the chosen arm was surprisingly good (positive PE), reduce stickiness for unchosen arms.
  unchosen_negative_stickiness_influence = jax.lax.cond(
      prediction_error > 0,
      # Scale by positive prediction error magnitude and certainty of chosen arm.
      lambda pe: -pe_stickiness_influence * pe * certainty_for_chosen_arm * 0.5,
      lambda pe: 0.0,
      prediction_error
  )
  # Apply this influence to unchosen arms only.
  updated_stickiness_values = updated_stickiness_values + unchosen_negative_stickiness_influence * (1 - choice_one_hot)

  # Q-value modulated decay for stickiness: higher Q-values lead to slower decay.
  # This makes stickiness persist longer for options that are performing well.
  q_modulated_decay_factor = stickiness_learning_rate * (1.0 - jax.nn.sigmoid(new_q_values))
  updated_stickiness_values = updated_stickiness_values * (1.0 - q_modulated_decay_factor)
  # Clip stickiness values to a reasonable range, ensuring they don't go below zero.
  updated_stickiness_values = jnp.clip(updated_stickiness_values, 0.0, 5.0) # Fixed upper bound for stickiness values.

  # Update the state with the new stickiness values.
  agent_state = agent_state.at[5:9].set(updated_stickiness_values)

  # Calculate choice logits based on Q-values, scaled by dynamic inverse temperature.
  choice_logits = new_beta * agent_state[:4]
  # Add choice stickiness to the chosen option.
  choice_logits += agent_state[5:9] * choice_one_hot

  # Add an exploration bonus for options with higher surprise and per-arm variance (uncertainty).
  # The bonus is now more directly weighted by uncertainty.
  # Incorporate exploration bonus for options with higher surprise, recent rewards, and per-arm variance.
  # The exploration bonus is now explicitly tied to uncertainty (per_arm_variance) to mimic Thompson Sampling.
  # The idea is to add a bonus proportional to the *standard deviation* of the rewards for each arm,
  # allowing for "sampling" from a distribution of expected rewards implicitly.
  exploration_bonus_surprise_and_recency = pe_stickiness_influence * updated_surprise_values + updated_last_reward_trace * 0.5
  # The Thompson-like exploration bonus uses the standard deviation of per-arm rewards.
  # We add a small epsilon to the variance before taking the square root to prevent NaN if variance is zero.
  exploration_bonus_thompson_like = jnp.sqrt(per_arm_variance + 1e-6) * base_reward_sensitivity * 0.2

  # Goal-Directed Exploration Bonus: Encourage re-sampling of unchosen options that are above average and not recently chosen.
  # This targets options that might be "potentially good" but have been neglected.
  # Calculate which unchosen options are 'potentially good' (Q-value > current_average_reward).
  is_potentially_good_unchosen = (agent_state[:4] > new_average_reward) * (1 - choice_one_hot)
  # Also, check if they haven't been chosen very recently (low last_reward_trace).
  not_recently_chosen = (updated_last_reward_trace < 0.2) * (1 - choice_one_hot) # If trace is low, it means it hasn't been chosen for a while.

  # The bonus is applied if the option is potentially good AND not recently chosen.
  # The strength of the bonus is modulated by environmental volatility and the Q-value's distance from average.
  goal_directed_exploration_bonus_magnitude = jnp.clip(
      (agent_state[:4] - new_average_reward) * (1.0 + environmental_volatility) * 0.5,
      0.0, 1.0) # Scale by how much better it is than average, and volatility.
  goal_directed_exploration_bonus = is_potentially_good_unchosen * not_recently_chosen * goal_directed_exploration_bonus_magnitude

  # Combine different exploration bonuses.
  exploration_bonus = exploration_bonus_surprise_and_recency + exploration_bonus_thompson_like + goal_directed_exploration_bonus
  choice_logits += exploration_bonus

  # Add inverse stickiness (repulsion) for unchosen options to promote switching.
  # This repulsion can also be modulated by the uncertainty of unchosen options: more uncertain, less repulsion needed.
  inverse_stickiness_repulsion = -choice_stickiness_decay_rate * (1 - choice_one_hot)
  choice_logits += inverse_stickiness_repulsion
  return choice_logits, agent_state


# SOO human_bandit even/odd split.
def human_bandit_run3_fit_only(
    params: chex.Array,
    choice: int,
    reward: float,
    agent_state: Optional[chex.Array],
) -> tuple[chex.Array, chex.Array]:
  """Cognitive model describing human behavior on a multi-armed bandit task.

  Assumes the agent is presented with four options on each trial.

  Args:
    params: Fit parameters of the model. Different parameters are used for
      different subjects.
    choice: Choice made by the agent on the previous trial. 0, 1, 2, or 3
    reward: Reward received by the agent on the previous trial. A float between
      0 and 1.
    agent_state: The current state of the cognitive model.

  Returns:
    choice_logits: The probabilities that the agent will choose option 0, 1, 2,
      or 3 on the next trial, expressed as logits.
    agent_state: New agent state
  """
  # Do not remove comments or TODOs from this program.

  # Define all parameters using descriptive names and apply transformations.
  # Parameters are mapped from the input `params` array.
  # We use a dictionary-like approach for clarity, even though `params` is an array.
  # The `params` array is expected to be ordered as follows:
  # [learning_rate_chosen, learning_rate_unchosen, inverse_temperature_base_pre_transform,
  #  recency_chosen_decay, recency_unchosen_decay, recency_weight_unchosen, initial_q_value_pre_transform,
  #  recency_weight_chosen_pre_transform, recency_general_decay, value_decay_rate]

  sigmoid = jax.nn.sigmoid
  softplus = jax.nn.softplus

  learning_rate_chosen = softplus(params[0])
  learning_rate_unchosen = softplus(params[1])
  inverse_temperature_base = softplus(params[2])
  recency_chosen_decay = sigmoid(params[3])
  recency_unchosen_decay = sigmoid(params[4])
  recency_weight_unchosen = sigmoid(params[5])
  initial_q_value = sigmoid(params[6])
  recency_weight_chosen = sigmoid(params[7])
  recency_general_decay = sigmoid(params[8])
  value_decay_rate = sigmoid(params[9])

  # Handle initial state: if agent_state is None, initialize Q-values with initial_q_value, else extract from state.
  # Recency trace is always initialized to zeros if state is None.
  if agent_state is None:
    q_values = jnp.full((4,), initial_q_value)
    recency_trace = jnp.zeros(4)
  else:
    q_values = agent_state[:4]
    recency_trace = agent_state[4:]

  # Update Q-values using a Rescorla-Wagner rule.
  # Ensure reward is within expected bounds [0, 1].
  reward = jnp.clip(reward, 0.0, 1.0)
  # Calculate prediction error for the chosen option.
  prediction_error_chosen = reward - q_values[choice]
  # Clip prediction error to be within [-1, 1] to stabilize learning.
  prediction_error_chosen = jnp.clip(prediction_error_chosen, -1.0, 1.0)
  # Modulate learning rate for the chosen option based on confidence and prediction error.
  # If the agent is highly confident (Q-value far from 0.5) and prediction error is large, increase learning rate.
  # If confidence is low (Q-value near 0.5), learning rate is more baseline.
  # Modulate learning rate for the chosen option based on the absolute prediction error.
  # Larger prediction errors lead to a higher learning rate, reflecting increased surprise.
  modulated_learning_rate_chosen = learning_rate_chosen * (1.0 + jnp.abs(prediction_error_chosen) * 0.5)
  # Ensure the modulated learning rate remains within reasonable bounds [0, 1].
  modulated_learning_rate_chosen = jnp.clip(modulated_learning_rate_chosen, 0.0, 1.0)
  chosen_q_value_update_term = modulated_learning_rate_chosen * jnp.tanh(prediction_error_chosen)

  # Calculate prediction error for unchosen options (difference between chosen Q-value and unchosen Q-value).
  prediction_error_unchosen = q_values[choice] - q_values
  # Calculate a global outcome surprise based on the absolute prediction error for the chosen option.
  outcome_surprise = jnp.abs(prediction_error_chosen)
  # Modulate the unchosen prediction error by the outcome surprise.
  # When surprise is high, we might want to dampen the direct impact of chosen Q-value on unchosen options,
  # as the chosen outcome itself was unexpected and might not be a reliable reference point for unchosen options.
  # Conversely, low surprise means the chosen outcome is reliable, so the difference is more directly applicable.
  # We use a sigmoid to scale surprise, making it a multiplier between (approx) 0.5 and 1.0.
  surprise_modulation_factor = 1.0 - sigmoid(outcome_surprise * 2) / 2 # Ranges from 1.0 (low surprise) to 0.5 (high surprise)
  prediction_error_unchosen = prediction_error_unchosen * surprise_modulation_factor

  # Compute an option-specific learning rate for unchosen options.
  # We modulate the learning rate for unchosen options by their recency trace,
  # and also by their absolute Q-value. Options with very high or very low
  # Q-values might have their learning rate dampened, reflecting a higher confidence
  # in their current estimated value, making them less prone to change when unchosen.
  # This adds a form of value-based resistance to change for unchosen options.
  # For unchosen options, scale the learning rate by their recency trace.
  # For unchosen options, scale the learning rate by their recency trace.
  recency_modulated_learning_rate_unchosen = learning_rate_unchosen * (1 + recency_trace)
  unchosen_q_value_update_term = recency_modulated_learning_rate_unchosen * jnp.tanh(prediction_error_unchosen)

  # Apply an option-specific value decay. More recently chosen options (higher recency_trace) decay slower,
  # reflecting stronger memory for recently engaged options.
  option_specific_value_decay = value_decay_rate * (1 + recency_trace * 0.1)
  option_specific_value_decay = jnp.clip(option_specific_value_decay, 0.0, 1.0)
  q_values = q_values * option_specific_value_decay # Apply decay to all Q-values.
  q_values = q_values.at[choice].add(chosen_q_value_update_term) # Add update for chosen option.
  q_values = q_values + unchosen_q_value_update_term # Add updates for unchosen options.

  # Update recency trace.
  recency_trace = recency_trace * recency_general_decay
  recency_trace = recency_trace.at[choice].add(1.0)

  # Calculate a decay factor for unchosen options, which is stronger for more recent options.
  decay_factor_unchosen = recency_unchosen_decay * (1 - recency_trace / 5.0)
  decay_factor_unchosen = jnp.clip(decay_factor_unchosen, 0.0, 1.0)

  # Calculate a modulated decay factor for the chosen option.
  # This factor is influenced by the reward and prediction error.
  # If the reward is low or prediction error is high, the decay for the chosen option might be stronger,
  # reflecting a less reliable memory of a 'bad' or 'surprising' choice.
  surprise_term = (1 - reward) * (1 - recency_chosen_decay) - (1 - jax.nn.sigmoid(jnp.abs(prediction_error_chosen))) * (1 - recency_chosen_decay)
  surprise_modulated_recency_chosen_decay = recency_chosen_decay + surprise_term
  surprise_modulated_recency_chosen_decay = jnp.clip(surprise_modulated_recency_chosen_decay, 0.0, 1.0)
  
  # Apply the specific decay factors to chosen and unchosen recency traces.
  recency_trace = jnp.where(
      jnp.arange(4) == choice,
      recency_trace * surprise_modulated_recency_chosen_decay,
      recency_trace * decay_factor_unchosen,
  )
  recency_trace = jnp.minimum(recency_trace, 5.0)

  # Compute choice logits.
  # Apply differential weights to recency trace for chosen vs unchosen options.
  # This allows recency to differentially influence the attractiveness of chosen and unchosen options.
  weighted_recency_trace = jnp.where(
      jnp.arange(4) == choice,
      recency_weight_chosen * recency_trace,
      recency_weight_unchosen * recency_trace,
  )

  # Adapt inverse temperature based on global surprise and local uncertainty.
  # The inverse temperature is reduced (more exploration) when the chosen prediction error is large (global surprise).
  # It is also reduced when the unchosen prediction error for a specific option is large (local uncertainty).
  # Additionally, when Q-values are very similar (low std dev), it encourages more exploration.
  global_surprise_reduction = 1 + jnp.abs(prediction_error_chosen) # High surprise -> lower temperature
  local_uncertainty_reduction = 1 + jnp.abs(prediction_error_unchosen) # High unchosen PE -> lower temperature
  
  # Encourage exploration when Q-values are similar (low std dev).
  # Low std dev means high uncertainty, so this factor increases the effective inverse temperature.
  q_value_std_dev = jnp.std(q_values)
  uncertainty_exploration_boost = 1 + jnp.exp(-q_value_std_dev * 2.5)

  # Combine all factors to get an option-specific inverse temperature.
  # The base temperature is divided by surprise/uncertainty terms, and multiplied by the exploration boost.
  option_specific_inverse_temperatures = (
      inverse_temperature_base / (global_surprise_reduction * local_uncertainty_reduction)
      * uncertainty_exploration_boost
  )
  
  # Combine Q-values and weighted recency trace to form an "action desirability" signal.
  # Recently engaged options (chosen or unchosen, depending on `recency_weight_unchosen`) are amplified.
  action_desirability = q_values * (1 + weighted_recency_trace)
  
  # Compute choice logits by multiplying action desirability with the adaptive inverse temperatures.
  choice_logits = action_desirability * option_specific_inverse_temperatures

  # Update the full agent state.
  agent_state = jnp.concatenate((q_values, recency_trace))

  return choice_logits, agent_state


# Shortened to 100k steps then refactored, run 1, low floor.
def human_bandit_run1_simplified_low_floor(
    params: chex.Array,
    choice: int,
    reward: float,
    agent_state: Optional[chex.Array],
) -> tuple[chex.Array, chex.Array]:
  """
  This function models a reinforcement learning agent's decision-making and learning process.
  It updates its internal state based on a choice and its resulting reward, and then
  calculates the action preferences (logits) for the next decision.
  """
  
  # === 1. Unpack and Transform Model Parameters ===
  
  # The raw parameters are passed through a sigmoid function to constrain them to the range [0, 1].
  sigmoid_params = jax.nn.sigmoid(params[:7])
  
  # Assign the constrained parameters to informative variable names.
  initial_q_value = sigmoid_params[0]
  learning_rate = sigmoid_params[1]
  inverse_temperature = sigmoid_params[2]
  unchosen_learning_rate = sigmoid_params[3]
  perseveration_learning_rate = sigmoid_params[4]
  initial_perseveration_value = sigmoid_params[5]
  perseveration_bias_strength = sigmoid_params[6]

  # Scale some parameters to a more behaviorally meaningful range (e.g., [0, 10]).
  # Inverse temperature controls the exploration-exploitation trade-off.
  inverse_temperature_scaled = inverse_temperature * 10
  # Perseveration bias strength controls the tendency to repeat the last action.
  perseveration_bias_strength_scaled = perseveration_bias_strength * 10
  
  # === 2. Initialize or Unpack Agent's Internal State ===
  
  # If this is the first trial (agent_state is None), initialize the agent's internal
  # state with the initial parameter values. Otherwise, unpack the state from the previous trial.
  if agent_state is None:
    # Initialize Q-values (expected reward for each action) and perseveration trace.
    q_values = initial_q_value
    perseveration_trace = initial_perseveration_value
  else:
    # Unpack the Q-values (first 4 elements) and perseveration trace (next 4 elements).
    q_values = agent_state[:4]
    perseveration_trace = agent_state[4:8]
    
  # === 3. Update Q-values (Action Values) ===
  
  # This uses a Rescorla-Wagner (delta) learning rule.
  
  # Create a one-hot encoded vector to identify which action was chosen.
  chosen_action_mask = jax.nn.one_hot(choice, num_classes=4)
  
  # Calculate the prediction error: the difference between actual and expected reward.
  prediction_error = reward - q_values
  
  # Determine the learning rate for each action: a different rate is used for the
  # chosen action compared to the unchosen ones.
  learning_rates_for_all_actions = jnp.where(
      chosen_action_mask, 
      learning_rate,            # Rate for the chosen action
      unchosen_learning_rate    # Rate for all other actions
  )
  
  # Update the Q-values by adding the prediction error, scaled by the learning rate.
  q_values = q_values + learning_rates_for_all_actions * prediction_error
  
  # === 4. Update Perseveration Trace ===
  
  # This trace models the agent's tendency to repeat recent actions.
  
  # For the chosen action, the trace is updated towards 1.
  updated_trace_for_chosen_action = (
      (1 - perseveration_learning_rate) * perseveration_trace 
      + perseveration_learning_rate
  )
  
  # Update the full perseveration trace array: apply the update for the chosen
  # action and reset the trace to 0.0 for all unchosen actions.
  perseveration_trace = jnp.where(
      chosen_action_mask,
      updated_trace_for_chosen_action,
      0.0,
  )
  
  # === 5. Calculate Action Preferences (Logits) ===
  
  # The agent's final choice preference is a combination of learned values and perseveration bias.
  
  # Calculate the component of choice preference driven by the learned Q-values.
  value_component = q_values * inverse_temperature_scaled
  
  # Calculate the component of choice preference driven by the perseveration trace.
  perseveration_component = perseveration_trace * perseveration_bias_strength_scaled
  
  # The final choice logits are the sum of the two components.
  # A softmax function is typically applied to these logits to get choice probabilities.
  choice_logits = value_component + perseveration_component
  
  # === 6. Prepare the New Agent State for the Next Trial ===
  
  # Concatenate the updated Q-values and perseveration trace into a single array
  # to be passed as the agent_state in the next iteration.
  agent_state = jnp.concatenate([q_values, perseveration_trace])
  
  # Return the calculated logits and the updated state.
  return choice_logits, agent_state


# Shortened to 100k steps then refactored, run 2, low floor.
def human_bandit_run2_simplified_low_floor(
    params: chex.Array,
    choice: int,
    reward: float,
    agent_state: Optional[chex.Array],
) -> tuple[chex.Array, chex.Array]:
  """A cognitive model of human learning in a multi-armed bandit task.

  Args:
    params: Subject-specific parameters.
    choice: The choice made on the previous trial (0-3).
    reward: The reward received on the previous trial (0.0-1.0).
    agent_state: The agent's internal state (Q-values and choice traces).

  Returns:
    choice_logits: Log probabilities for each choice on the next trial.
    agent_state: The updated agent's internal state.
  """
  # --- 1. Unpack and transform model parameters ---

  # Unpack the raw, unconstrained parameters for the model.
  # These are learned during model fitting.
  (raw_learning_rate, raw_initial_q_value, raw_unchosen_trace_decay,
   raw_background_learning_rate, raw_chosen_trace_decay,
   raw_inverse_temperature, raw_stickiness) = params[:7]

  # Apply a sigmoid function to constrain parameters to the (0, 1) range.
  # This is a standard way to handle unconstrained parameter optimization.
  learning_rate = jax.nn.sigmoid(raw_learning_rate)
  initial_q_value = jax.nn.sigmoid(raw_initial_q_value)
  unchosen_trace_decay_rate = jax.nn.sigmoid(raw_unchosen_trace_decay)
  background_learning_rate = jax.nn.sigmoid(raw_background_learning_rate)
  chosen_trace_decay_rate = jax.nn.sigmoid(raw_chosen_trace_decay)
  inverse_temperature_param = jax.nn.sigmoid(raw_inverse_temperature)
  stickiness = jax.nn.sigmoid(raw_stickiness)

  # Scale the inverse temperature parameter to a more common range (0-10).
  # This parameter controls the level of exploration vs. exploitation.
  inverse_temperature = inverse_temperature_param * 10.0

  # --- 2. Initialize or retrieve the agent's internal state ---

  # The agent's state consists of Q-values (value estimates) and choice traces
  # (a memory of recent choices). If this is the first trial, we initialize them.
  # Otherwise, we unpack them from the provided agent_state.
  if agent_state is None:
    # Initialize Q-values for all 4 choices to the initial_q_value parameter.
    q_values = jnp.full(shape=(4,), fill_value=initial_q_value)
    # Initialize choice traces for all 4 choices to the same initial value.
    choice_traces = jnp.full(shape=(4,), fill_value=initial_q_value)
  else:
    # Unpack the state from the previous trial.
    q_values = agent_state[:4]  # First 4 elements are Q-values
    choice_traces = agent_state[4:]  # Next 4 elements are choice traces

  # --- 3. Update Q-values based on the reward received ---

  # Calculate the prediction error: the difference between the actual reward and
  # the expected reward (the Q-value) for each choice.
  prediction_error = reward - q_values

  # Determine the learning rate for each choice. All unchosen choices are
  # updated with a small 'background_learning_rate', while the chosen action
  # gets an additional 'learning_rate'.
  learning_rates_for_all_choices = jnp.full(4, background_learning_rate)
  learning_rates_for_all_choices = learning_rates_for_all_choices.at[choice].add(
      learning_rate)

  # Update the Q-values using the prediction error and corresponding learning rate.
  # This is a form of the Rescorla-Wagner learning rule:
  # Q_new = Q_old + learning_rate * prediction_error
  q_values = q_values + (learning_rates_for_all_choices * prediction_error)

  # --- 4. Update choice traces to track recent choices ---

  # Create a vector of decay rates. The trace for the chosen action decays at a
  # different rate than the traces for the unchosen actions.
  decay_rates_for_all_choices = jnp.full(4, unchosen_trace_decay_rate)
  decay_rates_for_all_choices = decay_rates_for_all_choices.at[choice].set(
      chosen_trace_decay_rate)

  # Apply decay to all existing choice traces.
  choice_traces = choice_traces * decay_rates_for_all_choices

  # Increment the trace for the action that was just chosen by 1.0. This
  # reinforces the memory of making that choice, modeling perseveration.
  choice_traces = choice_traces.at[choice].add(1.0)

  # --- 5. Calculate choice logits for the next decision ---

  # Logits are the unnormalized log probabilities of choosing each action. They
  # are determined by two components: value-based preference and stickiness.

  # The value-based component is the Q-values scaled by the inverse temperature.
  # A higher temperature leads to more exploratory (less value-driven) choices.
  value_component = inverse_temperature * q_values

  # The stickiness component reflects a bias to repeat recent choices. It is
  # the choice traces scaled by the stickiness parameter.
  stickiness_component = stickiness * choice_traces

  # The final logits are the sum of the value and stickiness components.
  choice_logits = value_component + stickiness_component

  # --- 6. Package the updated state for the next trial ---

  # Concatenate the updated Q-values and choice traces into a single array
  # to be passed to the next call of this function.
  agent_state = jnp.concatenate((q_values, choice_traces))

  # Return the choice logits and the new agent state.
  return choice_logits, agent_state


# Shortened to 100k steps then refactored, run 3, low floor.
def human_bandit_run3_simplified_low_floor(
    params: chex.Array,
    choice: int,
    reward: float,
    agent_state: Optional[chex.Array],
) -> tuple[chex.Array, chex.Array]:
  """Cognitive model describing human behavior on a multi-armed bandit task.

  Assumes the agent is presented with four options on each trial.

  Args:
    params: Fit parameters of the model. Different parameters are used for
      different subjects.
    choice: Choice made by the agent on the previous trial. 0, 1, 2, or 3
    reward: Reward received by the agent on the previous trial. A float between
      0 and 1.
    agent_state: The current state of the cognitive model.

  Returns:
    choice_logits: The probabilities that the agent will choose option 0, 1, 2,
      or 3 on the next trial, expressed as logits.
    agent_state: New agent state
  """
  # --- 1. Initialization ---
  # If this is the first trial, agent_state will be None.
  # Initialize Q-values and recency traces to a neutral value (0.5).
  # The state vector has 8 elements: 4 for Q-values and 4 for recency.
  if agent_state is None:
      initial_values = 0.5
      agent_state = jnp.full(8, initial_values)

  # --- 2. Unpack Parameters and State ---
  # Unpack the model's behavioral parameters from the input array.
  # A sigmoid function is used to constrain the first five parameters between 0 and 1.
  learning_rate, unchosen_learning_rate, recency_decay, recency_boost_decay, recency_weight = jax.nn.sigmoid(params[:5])
  
  # The inverse_temperature parameter controls the stochasticity of choices.
  inverse_temperature = params[5]

  # Extract the agent's internal memory from the agent_state array.
  # Q-values represent the expected reward for each choice.
  q_values = agent_state[:4]
  # Recency trace captures a bias towards recently chosen options.
  recency_trace = agent_state[4:]

  # --- 3. Update Q-Values (Learning) ---
  # This model uses a modified Rescorla-Wagner learning rule.
  
  # a. Calculate the prediction error for the chosen option.
  prediction_error = reward - q_values[choice]
  
  # b. Calculate the update for the chosen option based on its learning rate.
  chosen_q_update = learning_rate * prediction_error
  
  # c. Calculate the updates for the unchosen options. They are pulled towards
  #    the value of the chosen option, based on a separate learning rate.
  unchosen_q_updates = unchosen_learning_rate * (q_values[choice] - q_values)

  # d. Combine the updates. Start with the unchosen updates for all options,
  #    then overwrite the update for the single chosen option.
  all_q_updates = unchosen_q_updates.at[choice].set(chosen_q_update)
  
  # e. Apply the updates to get the new Q-values.
  updated_q_values = q_values + all_q_updates

  # --- 4. Update Recency Trace ---
  # This process models a bias towards recently selected options.

  # a. First, all recency traces decay over time.
  decayed_recency_trace = recency_trace * recency_decay
  
  # b. For the chosen option, its recency is boosted by 1.0 and then
  #    decays by a separate "boost decay" factor.
  boosted_chosen_recency = (recency_trace[choice] + 1.0) * recency_boost_decay
  
  # c. The new recency trace is the decayed trace with the chosen option's
  #    value updated to its new boosted value.
  updated_recency_trace = decayed_recency_trace.at[choice].set(boosted_chosen_recency)

  # --- 5. Compute Choice Logits ---
  # The final choice probabilities are a combination of Q-values and recency.
  
  # a. Calculate the influence of the recency trace, weighted by its parameter.
  recency_component = recency_weight * updated_recency_trace
  
  # b. Combine the learned Q-values with the recency component.
  combined_value_signal = updated_q_values + recency_component
  
  # c. Scale the combined signal by the inverse temperature to get the final
  #    choice logits. Higher temperatures lead to more deterministic choices.
  choice_logits = inverse_temperature * combined_value_signal

  # --- 6. Prepare and Return New State ---
  # Combine the updated Q-values and recency trace into the new agent state vector.
  new_agent_state = jnp.concatenate([updated_q_values, updated_recency_trace])
  
  return choice_logits, new_agent_state


# Shortened to 100k steps then refactored, run 1, medium floor.
def human_bandit_run1_simplified_medium_floor(
    params: chex.Array,
    choice: int,
    reward: float,
    agent_state: Optional[chex.Array],
) -> tuple[chex.Array, chex.Array]:
  """Cognitive model describing human behavior on a multi-armed bandit task.

  Args:
    params: Fit parameters of the model. Different parameters are used for
      different subjects.
    choice: Choice made by the agent on the previous trial. 0, 1, 2, or 3
    reward: Reward received by the agent on the previous trial. A float between
      0 and 1.
    agent_state: The current state of the cognitive model.

  Returns:
    choice_logits: The probabilities that the agent will choose option 0, 1, 2,
      or 3 on the next trial, expressed as logits.
    agent_state: New agent state
  """

  # --- 1. Unpack and transform model parameters ---
  # Parameters are passed as a single vector and need to be unpacked.
  (
      initial_q_value_param,
      learning_rate_param,
      inverse_temperature_param,
      decay_rate_param, # This parameter is unused in the original code.
      unchosen_learning_rate_param,
      perseveration_learning_rate_param,
      perseveration_decay_rate_param,
      initial_perseveration_value_param, # This is also unused.
      perseveration_bias_strength_param,
      perseveration_reward_sensitivity_param,
  ) = params

  # Transform raw parameters into meaningful ranges for the model.
  # The sigmoid function squashes values to be between 0 and 1.
  # These values are then scaled and shifted to fit their specific roles.

  # Initial Q-value for all options on the first trial (range: 0 to 1).
  initial_q_value = jax.nn.sigmoid(initial_q_value_param)
  # Learning rate for the chosen option (range: 0 to 1).
  learning_rate = jax.nn.sigmoid(learning_rate_param)
  # Base inverse temperature for softmax choice rule (range: 0 to 10).
  inverse_temperature = 10 * jax.nn.sigmoid(inverse_temperature_param)
  # Learning rate for updating the average reward and unchosen options (range: 0.1 to 1.0).
  unchosen_learning_rate = 0.9 * jax.nn.sigmoid(unchosen_learning_rate_param) + 0.1
  # Learning rate for the perseveration trace (range: 0 to 1).
  perseveration_learning_rate = jax.nn.sigmoid(perseveration_learning_rate_param)
  # Decay rate for the perseveration trace (range: 0.025 to 0.975).
  perseveration_decay_rate = 0.95 * jax.nn.sigmoid(perseveration_decay_rate_param) + 0.025
  # Base strength of the tendency to repeat a choice.
  perseveration_bias_strength = perseveration_bias_strength_param
  # How much the last reward influences the tendency to repeat a choice.
  perseveration_reward_sensitivity = perseveration_reward_sensitivity_param

  # --- 2. Initialize or unpack the agent's internal state ---
  # The state includes Q-values, a perseveration trace, and an average reward estimate.
  if agent_state is None:
    # On the first trial, initialize the state.
    # Q-values represent the expected reward for each of the 4 options.
    q_values = jnp.full((4,), initial_q_value)
    # Perseveration trace represents the tendency to repeat each choice.
    perseveration_trace = jnp.full((4,), 0.0)
    # Average reward estimate tracks the overall value of the task.
    average_reward_estimate = initial_q_value
  else:
    # On subsequent trials, unpack the state from the previous trial.
    q_values = agent_state[0:4]
    perseveration_trace = agent_state[4:8]
    average_reward_estimate = agent_state[8]

  # --- 3. Update internal state based on the last choice and reward ---

  # 3a. Update the estimate of the average reward.
  # This tracks the overall reward rate of the environment.
  avg_reward_prediction_error = reward - average_reward_estimate
  average_reward_estimate = average_reward_estimate + unchosen_learning_rate * avg_reward_prediction_error

  # 3b. Update Q-values (the agent's estimate of each option's value).
  # First, all Q-values are pulled toward the average reward estimate. This can be
  # seen as a form of forgetting or regularization for unchosen options.
  q_values_after_decay = q_values + unchosen_learning_rate * (average_reward_estimate - q_values)

  # Next, the chosen option's Q-value is updated based on the actual reward received.
  # This is a standard delta-rule (Rescorla-Wagner) update.
  # Note: The prediction error is calculated based on the value *after* the decay step.
  chosen_q_value_prediction_error = reward - q_values_after_decay[choice]
  updated_chosen_q_value = q_values_after_decay[choice] + learning_rate * chosen_q_value_prediction_error

  # Combine the updates: use the decayed values for unchosen options and the new
  # updated value for the chosen one.
  q_values = q_values_after_decay.at[choice].set(updated_chosen_q_value)

  # 3c. Update the perseveration trace.
  # The trace for the chosen action is strengthened, while traces for unchosen actions decay.
  
  # First, calculate the updated value for the chosen action's trace.
  # This update is calculated based on its value *before* decay.
  updated_trace_for_chosen_action = perseveration_trace[choice] + perseveration_learning_rate * (1.0 - perseveration_trace[choice])
  
  # Next, apply exponential decay to all perseveration traces.
  perseveration_trace_after_decay = perseveration_trace * perseveration_decay_rate

  # Finally, insert the updated trace for the chosen action into the decayed array.
  perseveration_trace = perseveration_trace_after_decay.at[choice].set(updated_trace_for_chosen_action)

  # --- 4. Compute choice logits for the next trial ---
  # Logits are computed by combining value-based and perseveration-based influences.

  # 4a. Calculate the perseveration bias. This bias is modulated by the reward
  # from the last trial, making the agent more likely to repeat rewarded choices.
  perseveration_bias = perseveration_bias_strength + (perseveration_reward_sensitivity * reward)

  # 4b. Calculate Q-values relative to the average reward. This centers the values.
  relative_q_values = q_values - average_reward_estimate

  # 4c. Compute an option-specific inverse temperature. This makes the agent's
  # choices more sensitive to value differences for better-than-average options.
  value_based_temperature_scaling = jax.nn.softplus(relative_q_values + 1.0) + 0.01
  option_specific_inverse_temperature = inverse_temperature * value_based_temperature_scaling

  # 4d. Combine components to calculate final logits.
  # The value component pushes the agent towards options with high expected rewards.
  value_component = relative_q_values * option_specific_inverse_temperature
  # The perseveration component pushes the agent to repeat recent choices.
  perseveration_component = perseveration_trace * perseveration_bias
  # The final logits are the sum of these two influences.
  choice_logits = value_component + perseveration_component

  # --- 5. Package the new state for the next trial ---
  # Concatenate the updated internal states into a single vector.
  agent_state = jnp.concatenate(
      [q_values, perseveration_trace, jnp.array([average_reward_estimate])]
  )

  return choice_logits, agent_state


# Shortened to 100k steps then refactored, run 2, medium floor.
def human_bandit_run2_simplified_medium_floor(
    params: chex.Array,
    choice: int,
    reward: float,
    agent_state: Optional[chex.Array],
) -> tuple[chex.Array, chex.Array]:
  """Cognitive model describing human behavior on a multi-armed bandit task.

  Args:
    params: Fit parameters of the model. Different parameters are used for
      different subjects.
    choice: Choice made by the agent on the previous trial. 0, 1, 2, or 3
    reward: Reward received by the agent on the previous trial. A float between
      0 and 1.
    agent_state: The current state of the cognitive model.

  Returns:
    choice_logits: The probabilities that the agent will choose option 0, 1, 2,
      or 3 on the next trial, expressed as logits.
    agent_state: New agent state
  """
  # --- 1. Unpack and transform model parameters ---

  # Apply a sigmoid function to raw parameters to constrain them between 0 and 1.
  (
      learning_rate,
      initial_q_value,
      choice_trace_decay_rate,
      stickiness,
      chosen_action_decay_rate,
      uncertainty_weight,
      background_learning_rate,
      uncertainty_learning_rate,
      uncertainty_initial_value,
      inverse_temperature,
  ) = jax.nn.sigmoid(params[:10])

  # Scale the inverse_temperature (softmax temperature) for a wider dynamic range.
  inverse_temperature = inverse_temperature * 10.0

  # --- 2. Initialize agent state on the first trial ---

  if agent_state is None:
    # The agent state is a vector containing different cognitive variables.
    # It is initialized here if this is the very first trial.
    initial_q_values = jnp.full(4, initial_q_value)
    initial_uncertainty_values = jnp.full(4, uncertainty_initial_value)
    initial_choice_traces = jnp.zeros(4)
    initial_reward_background = jnp.array([0.5])  # Starts at a neutral 0.5

    agent_state = jnp.concatenate([
        initial_q_values,
        initial_uncertainty_values,
        initial_choice_traces,
        initial_reward_background,
    ])

  # --- 3. Unpack the agent's current state into meaningful variables ---

  # The state vector is split into its constituent parts.
  q_values, uncertainty_values, choice_traces, reward_background_expectation = jnp.split(
      agent_state, [4, 8, 12]
  )

  # --- 4. Update Q-values (action value estimates) based on the reward ---

  # Calculate the prediction error: the difference between the received reward
  # and the expected reward for the chosen action.
  prediction_error = reward - q_values[choice]

  # Update the Q-value of the chosen action using the prediction error and learning rate.
  # This is a standard Rescorla-Wagner update rule.
  updated_q_value_for_chosen_action = (
      q_values[choice] + learning_rate * prediction_error
  )
  q_values = q_values.at[choice].set(updated_q_value_for_chosen_action)

  # Create a one-hot vector to easily apply updates to chosen vs. unchosen actions.
  chosen_one_hot = jax.nn.one_hot(choice, num_classes=4)
  unchosen_mask = 1.0 - chosen_one_hot

  # Update the Q-values of the *unchosen* actions. This represents a form of
  # background learning or generalization, where unchosen options drift towards
  # a modified background expectation.
  background_q_value_update_term = background_learning_rate * (
      (reward_background_expectation - prediction_error) - q_values
  )
  q_values = q_values + unchosen_mask * background_q_value_update_term

  # --- 5. Update uncertainty estimates ---

  # The update for the chosen action's uncertainty depends on the absolute prediction error.
  # A large surprise (positive or negative) increases uncertainty about that action.
  chosen_uncertainty_update = uncertainty_learning_rate * (
      jnp.abs(prediction_error) - uncertainty_values
  )

  # The uncertainty of unchosen actions increases by a small, fixed amount.
  # This models a general increase in uncertainty for options not being sampled.
  unchosen_uncertainty_update = 0.01

  # Apply the respective updates to the chosen and unchosen actions' uncertainties.
  uncertainty_values = uncertainty_values + (
      chosen_one_hot * chosen_uncertainty_update
  ) + (
      unchosen_mask * unchosen_uncertainty_update
  )

  # --- 6. Update choice traces to model perseveration/stickiness ---

  # Create a vector of decay rates. The chosen action has a different decay rate
  # than the unchosen actions.
  decay_rates = jnp.full_like(choice_traces, choice_trace_decay_rate)
  decay_rates = decay_rates.at[choice].set(chosen_action_decay_rate)

  # Apply decay to all traces, then add 1 to the trace of the chosen action.
  # This makes recently chosen actions more likely to be chosen again.
  choice_traces = choice_traces * (1.0 - decay_rates) + chosen_one_hot

  # --- 7. Update the background reward expectation ---

  # This is a running average of the rewards received, updated with its own learning rate.
  reward_background_expectation = reward_background_expectation + (
      background_learning_rate * (reward - reward_background_expectation)
  )

  # --- 8. Re-assemble the new agent state vector ---

  # Concatenate the updated cognitive variables back into a single state vector.
  agent_state = jnp.concatenate(
      (q_values, uncertainty_values, choice_traces, reward_background_expectation)
  )

  # --- 9. Calculate choice logits for the next action ---

  # The effective temperature is modulated by uncertainty. Higher uncertainty reduces
  # the influence of the Q-values, making choices more random (exploratory).
  effective_inverse_temperature = inverse_temperature / (
      1.0 + uncertainty_weight * uncertainty_values
  )

  # The final logits are a weighted sum of two components:
  # 1. The value-based component (Q-values scaled by effective temperature).
  value_component = effective_inverse_temperature * q_values
  # 2. The stickiness component (choice traces encouraging perseveration).
  stickiness_component = stickiness * choice_traces

  choice_logits = value_component + stickiness_component

  # --- 10. Return the results ---

  return choice_logits, agent_state


# Shortened to 100k steps then refactored, run 3, medium floor.
def human_bandit_run3_simplified_medium_floor(
    params: chex.Array,
    choice: int,
    reward: float,
    agent_state: Optional[chex.Array],
) -> tuple[chex.Array, chex.Array]:
  """Cognitive model describing human behavior on a multi-armed bandit task.

  Assumes the agent is presented with four options on each trial.

  Args:
    params: Fit parameters of the model. Different parameters are used for
      different subjects.
    choice: Choice made by the agent on the previous trial. 0, 1, 2, or 3
    reward: Reward received by the agent on the previous trial. A float between
      0 and 1.
    agent_state: The current state of the cognitive model.

  Returns:
    choice_logits: The probabilities that the agent will choose option 0, 1, 2,
      or 3 on the next trial, expressed as logits.
    agent_state: New agent state
  """
  # === 1. Unpack and Transform Model Parameters ===
  # Apply a sigmoid function to the first 8 parameters to constrain them
  # between 0 and 1. This is common for rates, weights, and probabilities.
  (learning_rate_chosen,
   learning_rate_unchosen,
   recency_chosen_decay,
   recency_unchosen_decay,
   recency_weight_unchosen,
   initial_q_value,
   recency_weight_chosen,
   value_decay_rate) = jax.nn.sigmoid(params[:8])

  # Apply a softplus function to the temperature parameter to ensure it is
  # positive. Inverse temperature controls the randomness of choices.
  inverse_temperature_base = jax.nn.softplus(params[8])


  # === 2. Initialize or Load Agent's Internal State ===
  # The agent's state consists of Q-values (expected reward for each option)
  # and a recency trace (memory of recent choices).
  if agent_state is None:
    # If this is the first trial, initialize Q-values and recency trace.
    q_values = jnp.full(shape=(4,), fill_value=initial_q_value)
    recency_trace = jnp.zeros(shape=(4,))
  else:
    # If not the first trial, load the state from the previous trial.
    q_values = agent_state[:4]       # First 4 elements are Q-values
    recency_trace = agent_state[4:]  # Last 4 elements are recency traces


  # === 3. Update Q-Values Based on Last Trial's Outcome ===
  # This model uses a variant of the Rescorla-Wagner learning rule.

  # Calculate the prediction error for the chosen option. This is the
  # difference between the actual reward received and the expected reward.
  prediction_error_chosen = reward - q_values[choice]

  # Calculate a "prediction error" for unchosen options. This represents how
  # much better the chosen option was compared to the unchosen ones.
  prediction_error_unchosen = q_values[choice] - q_values

  # Update the Q-value for the chosen option based on its prediction error.
  q_values_after_chosen_update = q_values.at[choice].add(
      learning_rate_chosen * prediction_error_chosen
  )
  # Update the Q-values for the unchosen options.
  q_values_after_unchosen_update = (
      q_values_after_chosen_update +
      learning_rate_unchosen * prediction_error_unchosen
  )
  # Apply a decay rate to all Q-values, causing them to slowly regress
  # towards zero over time (forgetting).
  q_values = q_values_after_unchosen_update * value_decay_rate


  # === 4. Update Recency Trace ===
  # The recency trace tracks which options were chosen recently.

  # Create a one-hot vector to easily distinguish the chosen option.
  chosen_one_hot = jax.nn.one_hot(choice, num_classes=4)

  # For the chosen option: increment its recency trace (capped at 5.0) and apply decay.
  # This strengthens the memory of it being recently picked.
  recency_update_for_chosen = (
      jnp.minimum(recency_trace + 1.0, 5.0) * recency_chosen_decay
  )
  # For unchosen options: apply a different decay factor.
  # This weakens the memory of them being picked.
  recency_update_for_unchosen = (
      recency_trace * recency_unchosen_decay * (1 - recency_trace / 5.0)
  )
  # Apply the appropriate update rule to each element of the recency trace.
  recency_trace = jnp.where(
      chosen_one_hot,
      recency_update_for_chosen,
      recency_update_for_unchosen
  )


  # === 5. Compute Choice Logits for the Next Trial ===
  # Logits are the raw, unnormalized scores for each choice. A higher logit
  # means a higher probability of being chosen.

  # Apply different weights to the recency trace for the chosen vs. unchosen options.
  # This allows the model to capture effects like "choice perseveration".
  weighted_recency_trace = jnp.where(
      chosen_one_hot,
      recency_weight_chosen * recency_trace,
      recency_weight_unchosen * recency_trace,
  )

  # Calculate a recency "bonus" that will multiplicatively modulate the Q-values.
  recency_bonus = 1 + weighted_recency_trace

  # Modulate the inverse temperature based on the magnitude of the prediction error.
  # This makes choices more random (lower temperature) after a surprising outcome.
  adaptive_inverse_temperature = (
      inverse_temperature_base / (1 + jnp.abs(prediction_error_unchosen))
  )

  # Combine Q-values and the recency bonus to get a value score.
  value_with_recency = q_values * recency_bonus

  # Final logits are calculated by scaling the combined value by the adaptive temperature.
  choice_logits = adaptive_inverse_temperature * value_with_recency


  # === 6. Prepare State for the Next Trial and Return ===

  # Concatenate the updated Q-values and recency trace into a single array.
  agent_state = jnp.concatenate((q_values, recency_trace))

  return choice_logits, agent_state


# Shortened to 100k steps then refactored, run 1, high floor.
def human_bandit_run1_simplified_high_floor(
    params: chex.Array,
    choice: int,
    reward: float,
    agent_state: Optional[chex.Array],
) -> tuple[chex.Array, chex.Array]:
  """Cognitive model describing human behavior on a multi-armed bandit task.

  Assumes the agent is presented with four options on each trial.

  Args:
    params: Fit parameters of the model. Different parameters are used for
      different subjects.
    choice: Choice made by the agent on the previous trial. 0, 1, 2, or 3
    reward: Reward received by the agent on the previous trial. A float between
      0 and 1.
    agent_state: The current state of the cognitive model.

  Returns:
    choice_logits: The probabilities that the agent will choose option 0, 1, 2,
      or 3 on the next trial, expressed as logits.
    agent_state: New agent state
  """

  # --- 1. Unpack Model Parameters ---
  # These parameters are fixed for a given agent and are learned from data.
  # They are transformed using sigmoid or other functions to constrain their range.

  # The initial belief about the value of each option, used for initialization and as a decay target.
  initial_q_value = jax.nn.sigmoid(params[0])

  # Learning rate for the Q-value of the *chosen* option.
  learning_rate = jax.nn.sigmoid(params[1])

  # The base inverse temperature for the softmax choice rule, controlling the level of exploration.
  inverse_temperature = 10 * jax.nn.sigmoid(params[2])

  # Learning rate for unchosen options, causing them to decay towards a prior.
  unchosen_learning_rate = 0.95 * jax.nn.sigmoid(params[3]) + 0.025

  # Learning rate for the perseveration trace of the *chosen* option.
  perseveration_learning_rate = jax.nn.sigmoid(params[4])

  # Decay rate for the perseveration traces of *unchosen* options.
  perseveration_decay_rate = jax.nn.sigmoid(params[5])

  # The initial value of the perseveration trace for each option.
  initial_perseveration_value = jax.nn.sigmoid(params[6])

  # The strength of the perseveration bias, influencing how much the perseveration trace affects choices.
  perseveration_bias_strength = jax.nn.sigmoid(params[7]) * 10.0

  # How sensitive the perseveration bias is to the reward outcome.
  perseveration_reward_sensitivity = jax.nn.sigmoid(params[8])


  # --- 2. Initialize or Unpack Agent State ---
  # The agent state holds dynamic variables that are updated on each trial.
  if agent_state is None:
    # If this is the first trial, initialize all state variables.
    num_options = 4
    q_values = jnp.full(num_options, initial_q_value)
    perseveration_trace = jnp.full(num_options, initial_perseveration_value)
    average_reward_estimate = initial_q_value
    recency_trace = jnp.zeros(num_options)
  else:
    # Otherwise, unpack the state from the previous trial.
    q_values = agent_state[0:4]
    perseveration_trace = agent_state[4:8]
    average_reward_estimate = agent_state[8]
    recency_trace = agent_state[9:13]


  # --- 3. Update Q-Values (Action Values) ---

  # 3.1. First, apply a global decay to all Q-values, pulling them towards the initial prior.
  # This represents a forgetting or diffusion process.
  q_values = q_values * (1 - unchosen_learning_rate) + initial_q_value * unchosen_learning_rate

  # 3.2. Update the running estimate of the average reward in the environment.
  # This estimate acts as a baseline for evaluating individual options.
  reward_prediction_error_avg = reward - average_reward_estimate
  avg_reward_error_magnitude_factor = 1.0 + jax.nn.softplus(jnp.abs(reward_prediction_error_avg))
  effective_avg_reward_learning_rate = unchosen_learning_rate * avg_reward_error_magnitude_factor
  average_reward_estimate += effective_avg_reward_learning_rate * reward_prediction_error_avg

  # 3.3. Update the Q-value for the *chosen* option based on the reward received.
  # This is a standard Rescorla-Wagner update rule.
  prediction_error = reward - q_values[choice]
  updated_chosen_q_value = q_values[choice] + learning_rate * prediction_error
  q_values = q_values.at[choice].set(updated_chosen_q_value)

  # 3.4. Update the Q-values for the *unchosen* options.
  # These values decay towards the average reward estimate, allowing them to track the
  # overall richness of the environment. The decay rate is modulated by "global surprise."
  global_surprise_factor = 1.0 + jax.nn.softplus(jnp.abs(prediction_error))
  unchosen_inferred_prediction_error = reward - q_values
  unchosen_surprise_factor = 1.0 + jax.nn.softplus(jnp.abs(unchosen_inferred_prediction_error))
  effective_unchosen_learning_rate = unchosen_learning_rate * global_surprise_factor * unchosen_surprise_factor
  
  updated_unchosen_q_values = (
      q_values * (1 - effective_unchosen_learning_rate) +
      average_reward_estimate * effective_unchosen_learning_rate
  )

  # Create a mask to apply updates only to unchosen options.
  unchosen_mask = jnp.arange(4) != choice
  q_values = jnp.where(
      unchosen_mask,
      updated_unchosen_q_values,
      q_values # Keep the already-updated chosen Q-value.
  )


  # --- 4. Update Perseveration Trace ---
  # This trace captures the tendency to repeat previous choices (habit).

  # 4.1. For the *chosen* option, increase the perseveration trace towards 1.
  # The update is stronger for positive prediction errors (unexpected rewards).
  chosen_perseveration_learning_rate = perseveration_learning_rate * jax.nn.sigmoid(prediction_error * 5)
  perseveration_trace_update_chosen = perseveration_trace[choice] + chosen_perseveration_learning_rate * (1 - perseveration_trace[choice])

  # 4.2. For *unchosen* options, decay the perseveration trace.
  # The decay rate is increased by several factors reflecting environmental uncertainty and surprise.
  unchosen_relative_badness_factor = jax.nn.sigmoid(q_values - reward)
  global_average_reward_context_factor = 1.0 + jax.nn.relu(average_reward_estimate - 0.5)
  global_outcome_surprise_factor = 1.0 + jax.nn.softplus(jnp.abs(prediction_error))
  
  decay_multiplier = (
      1.0 + unchosen_relative_badness_factor +
      global_average_reward_context_factor + global_outcome_surprise_factor
  )
  effective_perseveration_decay_unchosen = jnp.clip(perseveration_decay_rate * decay_multiplier, 0.0, 1.0)
  
  perseveration_trace_update_unchosen = (
      perseveration_trace * (1 - effective_perseveration_decay_unchosen) +
      initial_perseveration_value * effective_perseveration_decay_unchosen
  )

  # 4.3. Combine the updates for the chosen and unchosen options.
  perseveration_trace = jnp.where(
      unchosen_mask,
      perseveration_trace_update_unchosen,
      perseveration_trace_update_chosen,
  )


  # --- 5. Update Recency Trace ---
  # This trace captures more general recency effects.

  # 5.1. For the *chosen* option, increase the recency trace towards 1.
  # The update is scaled by the prediction error.
  recency_update_factor = 1.0 + prediction_error
  reward_modulated_recency_learning_rate = perseveration_learning_rate * recency_update_factor
  recency_trace_update_chosen = recency_trace[choice] + reward_modulated_recency_learning_rate * (1 - recency_trace[choice])

  # 5.2. For *unchosen* options, decay the recency trace.
  # The decay is slower for options that had Q-values similar to the chosen one.
  q_value_similarity_to_chosen = jax.nn.sigmoid(5.0 * (1.0 - jnp.abs(q_values - q_values[choice])))
  recency_decay_scaling_factor = 1.0 - 0.9 * q_value_similarity_to_chosen
  effective_recency_decay_rate = perseveration_decay_rate * recency_decay_scaling_factor
  recency_trace_update_unchosen = recency_trace * (1 - effective_recency_decay_rate)

  # 5.3. Combine the updates for the chosen and unchosen options.
  recency_trace = jnp.where(
      unchosen_mask,
      recency_trace_update_unchosen,
      recency_trace_update_chosen,
  )


  # --- 6. Compute Choice Logits ---
  # Combine the learned values and biases to determine the choice probabilities for the next trial.

  # 6.1. Compute an adaptive perseveration bias.
  # This bias is modulated by value uncertainty, recency, and reward sensitivity.
  value_uncertainty = 1.0 - jax.nn.sigmoid(jnp.abs(q_values - average_reward_estimate) * 5.0)
  adaptive_perseveration_bias_strength = perseveration_bias_strength * (1.0 + perseveration_reward_sensitivity * prediction_error)
  base_perseveration_strength = perseveration_trace * adaptive_perseveration_bias_strength * (0.5 + 0.5 * recency_trace)
  perseveration_bias = jax.nn.softplus(base_perseveration_strength * (1.0 + value_uncertainty))

  # 6.2. Compute relative Q-values by centering them around the average reward estimate.
  # This makes the choice dependent on how much better an option is than the environment's average.
  relative_q_values = q_values - average_reward_estimate

  # 6.3. Compute an option-specific inverse temperature.
  # This makes the choice policy more exploitative when value estimates are more certain (higher variance).
  q_value_variance = jnp.var(q_values)
  variance_scaling_factor = 0.5 + 1.5 * jax.nn.sigmoid(q_value_variance)
  option_specific_inverse_temperature = inverse_temperature * variance_scaling_factor * (jax.nn.relu(1 + relative_q_values) + 0.01)

  # 6.4. Combine the value-based and habit-based components to get the final choice logits.
  # This is a hybrid model where choice is driven by both goal-directed values and habitual perseveration.
  choice_logits = (
      relative_q_values * option_specific_inverse_temperature +
      perseveration_trace * perseveration_bias
  )


  # --- 7. Assemble New Agent State for the Next Trial ---
  agent_state = jnp.concatenate([
      q_values,
      perseveration_trace,
      jnp.array([average_reward_estimate]),
      recency_trace
  ])

  return choice_logits, agent_state


# Shortened to 100k steps then refactored, run 2, high floor.
def human_bandit_run2_simplified_high_floor(
    params: chex.Array,
    choice: int,
    reward: float,
    agent_state: Optional[chex.Array],
) -> tuple[chex.Array, chex.Array]:
  """Cognitive model describing human behavior on a multi-armed bandit task.

  Assumes the agent is presented with four options on each trial.

  Args:
    params: Fit parameters of the model. Different parameters are used for
      different subjects.
    choice: Choice made by the agent on the previous trial. 0, 1, 2, or 3
    reward: Reward received by the agent on the previous trial. A float between
      0 and 1.
    agent_state: The current state of the cognitive model.

  Returns:
    choice_logits: The probabilities that the agent will choose option 0, 1, 2,
      or 3 on the next trial, expressed as logits.
    agent_state: New agent state
  """
  if agent_state is None:
    agent_state = jnp.array([
        *[1.0] * 4,
        0.0,
        *[0.0] * 4,
        0.5,
        2.0,
        *[0.1] * 4,
        1.0,
        0.0,
        0.0,
        *[0.5] * 4
    ])

  # Apply softplus to parameters that need to be positive and allow for a wider range.
  # These include baseline_beta, base_learning_rate, base_reward_sensitivity, choice_stickiness_bias, and pe_stickiness_influence.
  (
      baseline_beta,
      base_learning_rate,
      base_reward_sensitivity,
      choice_stickiness_bias,
  ) = jax.nn.softplus(params[0:4])

  # Apply sigmoid to parameters that need to be bounded between 0 and 1.
  # These include decay rates and learning rates for various mechanisms.
  (
      choice_stickiness_decay_rate,
      unchosen_q_decay_rate,
      average_reward_learning_rate,
      beta_learning_rate,
      surprise_learning_rate,
  ) = jax.nn.sigmoid(params[4:9])

  pe_stickiness_influence = jax.nn.softplus(params[9])

  # Learning rate for choice stickiness, currently sharing parameter with average_reward_learning_rate for simplicity.
  stickiness_learning_rate = average_reward_learning_rate

  # Update the average reward for the overall task environment.
  # This acts as a dynamic baseline for reward expectations.
  current_average_reward = agent_state[9]
  average_reward_prediction_error = reward - current_average_reward
  new_average_reward = current_average_reward + average_reward_learning_rate * average_reward_prediction_error
  agent_state = agent_state.at[9].set(new_average_reward)

  # Update the expected reward for the chosen option using a Rescorla-Wagner rule, and decay all options.
  # Create a mask for the chosen arm.
  choice_mask = jax.nn.one_hot(choice, num_classes=4)
  decayed_q_values = agent_state[:4]

  # For unchosen options, apply an additional decay towards the current average reward.
  # The effective decay rate for unchosen options.
  # Add a small constant to ensure some decay even if unchosen_q_decay_rate is very small.

  # For unchosen options, apply an additional decay towards the current average reward.
  # The effective decay rate for unchosen options.
  # Add a small constant to ensure some decay even if unchosen_q_decay_rate is very small.
  effective_decay_rates_for_unchosen = unchosen_q_decay_rate + 0.05

  # We currently use a fixed max_observed_reward of 1.0 and rewards are between 0 and 1.
  # Thus, normalized_reward is approximately equal to reward.
  # Simplifying to use reward directly.
  normalized_reward = reward

  # The reward prediction error is the difference between the actual reward and the expected reward.
  # We apply dynamic_reward_sensitivity to the reward prediction error.
  # Calculate reward variance.
  # Use a small epsilon to avoid division by zero if reward_count is 0 or 1.
  sum_sq_rewards = agent_state[16]
  reward_count = agent_state[17]
  new_sum_sq_rewards = sum_sq_rewards + reward**2
  new_reward_count = reward_count + 1
  agent_state = agent_state.at[16].set(new_sum_sq_rewards)
  agent_state = agent_state.at[17].set(new_reward_count)
  mean_reward = new_average_reward
  reward_variance = jnp.maximum(0.0, (new_sum_sq_rewards / (new_reward_count + 1e-6)) - mean_reward**2)
  dynamic_reward_sensitivity = base_reward_sensitivity / (1.0 + reward_variance)

  environmental_volatility = jax.nn.sigmoid(reward_variance * 5.0 - 1.0)

  raw_prediction_error = (normalized_reward * dynamic_reward_sensitivity) - agent_state[choice]
  # Introduce precision weighting: prediction error is scaled by the inverse of the surprise
  # for the chosen arm. High surprise implies low precision, reducing the PE's impact.
  # We use a more direct certainty proxy: a decay of past surprise values, where low surprise means high certainty.
  # This term (1 - agent_state[11 + choice]) represents 'certainty' for the chosen arm.
  # High surprise (agent_state[11 + choice]) means low certainty.
  certainty_for_chosen_arm = 1.0 - agent_state[11 + choice]
  # Inverse certainty represents uncertainty, which will be used to modulate learning.

  # Dynamic RPE sensitivity modulation based on surprise and reward sign.

  # Incorporate certainty directly into the prediction error itself, creating a 'precision-weighted' PE.
  # This means prediction errors from less certain states have less impact on learning.
  # Scale precision_weighted_pe by the chosen arm's surprise to make the RPE more impactful when unexpected.
  # A softplus is used to ensure the scaling factor is positive and increases non-linearly with surprise.
  # Capped at 3.0 to prevent overly aggressive scaling.
  precision_weighted_pe = raw_prediction_error * certainty_for_chosen_arm
  # The effective prediction error for Q-value updates.
  prediction_error = precision_weighted_pe

  # Asymmetric decay for unchosen Q-values based on prediction error for the chosen arm.
  # Decay unchosen Q-values. Chosen Q-values are updated separately.
  # For unchosen options, decay their current values (which already have the general decay applied)
  # further towards the new_average_reward, modulated by surprise and recency.
  # For unchosen options, decay their current values further towards the new_average_reward,
  # modulated by the effective decay rate, and also towards a neutral baseline (0.5).
  unchosen_q_values_decay_to_average = (decayed_q_values * (1.0 - effective_decay_rates_for_unchosen)) + \
                                       (new_average_reward * effective_decay_rates_for_unchosen)
  # Additionally, introduce a small, constant decay towards a neutral baseline (0.5) for unchosen arms.
  unchosen_q_values_decay_to_average = (unchosen_q_values_decay_to_average * (1.0 - 0.005)) + (0.5 * 0.005)
  # New: Introduce a small, constant decay towards the minimum possible reward (0.0) for unchosen arms.
  # This acts as a 'pessimistic' decay for options not recently experienced, encouraging exploration.
  # The decay rate is also modulated by environmental volatility: higher volatility means faster decay towards 0.0.
  # We now decay unchosen Q-values towards a blend of 0.0 and the current average reward,
  # with the blend ratio modulated by environmental volatility.
  pessimistic_forgetting_rate_base = 0.002
  pessimistic_forgetting_rate_volatility_mod = environmental_volatility * 0.015 # Increased sensitivity to volatility
  pessimistic_forgetting_rate = pessimistic_forgetting_rate_base + pessimistic_forgetting_rate_volatility_mod

  # For unchosen options, apply a simple pessimistic decay towards 0.0.
  unchosen_q_values_decay_to_average = (unchosen_q_values_decay_to_average * (1.0 - pessimistic_forgetting_rate)) + \
                                       (0.0 * pessimistic_forgetting_rate)
  
  # Introduce a small, inverse influence of prediction error from the chosen arm on unchosen arms.
  # If chosen was better than expected (positive PE), unchosen arms become slightly less attractive.
  unchosen_pe_influence = -0.1 * prediction_error

  # Goal-directed re-evaluation for unchosen options: if the chosen option was highly surprising,
  # re-evaluate unchosen options based on the sign of the prediction error.
  # If chosen was surprisingly good, unchosen options get a small boost.
  # If chosen was surprisingly bad, unchosen options get a small reduction.
  # This effect is scaled by the 'surprise' of the chosen arm and its certainty.
  # Introduce a "forgetting" mechanism for unchosen options, decaying them towards the neutral baseline (0.5).
  # The decay rate is proportional to how long it's been since the arm was chosen (implicit in current Q-value).
  # We use a small, fixed forgetting rate for simplicity.
  # Incorporate the decaying last_reward_trace into Q-value updates for all arms.
  # This provides a memory of recent rewards, even for unchosen options.
  last_reward_trace_decay_rate = 0.1 # Small decay for the reward trace
  # Scale by environmental_volatility: in volatile environments, recent rewards are more important.
  effective_trace_decay = last_reward_trace_decay_rate * (1.0 + environmental_volatility * 0.5)
  # Update last_reward_trace: for the chosen arm, it's the current reward. For unchosen, it decays.
  current_last_reward_trace = agent_state[18:22]
  # Update last_reward_trace: for the chosen arm, it's the current reward.
  # For unchosen arms, their trace decays, but also gets a small, prediction-error-modulated update.
  # If the chosen arm was surprisingly good (positive PE), subtly boost unchosen arms' traces.
  # If the chosen arm was surprisingly bad (negative PE), subtly reduce unchosen arms' traces.
  pe_modulated_unchosen_trace_update = prediction_error * 0.1 * (1.0 - choice_mask) * certainty_for_chosen_arm
  updated_last_reward_trace = (current_last_reward_trace * (1.0 - effective_trace_decay)) * (1 - choice_mask) + \
                              (normalized_reward * choice_mask) + \
                              pe_modulated_unchosen_trace_update
  agent_state = agent_state.at[18:22].set(updated_last_reward_trace)

  # For the chosen arm, its Q-value maintains the general decay. Unchosen arms get specific decay + PE influence.
  unchosen_updated_q_values = (unchosen_q_values_decay_to_average + unchosen_pe_influence + updated_last_reward_trace * 0.05) * (1 - choice_mask)

  # Dynamically adjust the learning rate based on surprise, certainty, global reward variance, and prediction error magnitude.
  # Higher surprise, reward variance, or prediction error can increase the learning rate.
  effective_learning_rate_modulator = jax.nn.softplus(jnp.abs(prediction_error) * 0.75) * 0.3 # Increased scaling for PE magnitude
  effective_learning_rate = jnp.clip(
      base_learning_rate + effective_learning_rate_modulator,
      0.01, 0.99
  )

  # Update the chosen Q-value using the Rescorla-Wagner rule.
  updated_chosen_q = decayed_q_values[choice] + effective_learning_rate * prediction_error
  # Combine decayed Q-values for unchosen arms and updated Q-value for the chosen arm.
  new_q_values = unchosen_updated_q_values + updated_chosen_q * choice_mask
  agent_state = agent_state.at[:4].set(new_q_values)

  # Update the dynamic inverse temperature (beta) based on prediction error.
  # Larger absolute prediction errors decrease beta (more exploration), smaller errors increase beta (more exploitation).
  current_beta = agent_state[10]
  # The learning rate for beta update.
  effective_beta_learning_rate = beta_learning_rate
  # The change in beta is inversely proportional to the absolute prediction error.
  beta_update_signal = (1 - jnp.abs(prediction_error))

  # Update beta based on the update signal.
  # Update beta based on the update signal, decaying towards baseline_beta.
  new_beta = (current_beta * (1 - beta_learning_rate)) + (baseline_beta * beta_learning_rate) + (effective_beta_learning_rate * beta_update_signal)
  # Ensure beta remains positive and within a reasonable range.
  new_beta = jnp.clip(new_beta, 0.1, 10.0)
  agent_state = agent_state.at[10].set(new_beta)

  # Update the surprise for each arm.
  current_surprise_values = agent_state[11:15]
  # Decay all surprise values.
  updated_surprise_values = current_surprise_values * (1 - surprise_learning_rate)
  # Increase surprise for the chosen arm based on the direction and magnitude of the prediction error,
  # modulated by the certainty of the prediction for that arm.
  # Positive prediction errors might lead to a different surprise update than negative ones.
  # Here, we introduce an asymmetry: positive PEs increase surprise more than negative PEs of the same magnitude.

  # Surprise update based on the magnitude of prediction error, with asymmetry for negative PEs.
  # Negative prediction errors (worse than expected) cause a larger increase in surprise.
  surprise_update_magnitude_positive = surprise_learning_rate * prediction_error * 0.5 # Smaller impact for positive PEs
  surprise_update_magnitude_negative = surprise_learning_rate * jnp.abs(prediction_error) * 1.5 # Larger impact for negative PEs

  # Apply asymmetric scaling to the surprise update.
  asymmetric_surprise_update_signal = jax.lax.cond(
      raw_prediction_error < 0, # Use raw_prediction_error for surprise update logic
      lambda pe: surprise_update_magnitude_negative,
      lambda pe: surprise_update_magnitude_positive,
      raw_prediction_error
  )

  # Modulate the surprise update by the certainty of the prediction for that arm.
  # If the agent was highly certain (high `certainty_for_chosen_arm`) and still
  # made a large prediction error, this should lead to a stronger update of surprise.
  # We use raw_prediction_error here to ensure surprise is sensitive to the full magnitude of the unexpected outcome.
  certainty_modulated_surprise_update = asymmetric_surprise_update_signal * certainty_for_chosen_arm
  updated_surprise_values = updated_surprise_values.at[choice].set(
      jnp.clip(current_surprise_values[choice] + certainty_modulated_surprise_update, 0.0, 1.0)
  )
  agent_state = agent_state.at[11:15].set(updated_surprise_values)

  # Update choice stickiness values for each arm.
  current_stickiness_values = agent_state[5:9]
  choice_one_hot = jax.nn.one_hot(choice, num_classes=4)
  
  # All stickiness values decay slightly towards zero, and apply a volatility-sensitive soft reset.
  stickiness_reset_strength = environmental_volatility * 0.2
  updated_stickiness_values = current_stickiness_values * (1 - stickiness_learning_rate) * (1.0 - stickiness_reset_strength)

  # Apply decay to the choice stickiness bias.
  decayed_choice_stickiness_bias = choice_stickiness_bias * (1.0 - choice_stickiness_decay_rate)
  # Base update for the chosen arm, incorporating bias and prediction error.
  # We now also consider the prior action when updating stickiness for the chosen arm.
  base_chosen_stickiness_update = decayed_choice_stickiness_bias + pe_stickiness_influence * prediction_error

  # Update for chosen arm, including a direct reduction for negative prediction errors.
  # If prediction error is negative, reduce stickiness to encourage switching.
  stickiness_reduction_for_negative_pe = jax.lax.cond(
      prediction_error < 0,
      # Stronger reduction if certain about the bad outcome and in a volatile environment.
      lambda pe: stickiness_learning_rate * jnp.abs(pe) * (1.0 + certainty_for_chosen_arm) * (1.0 + environmental_volatility * 0.5),
      lambda pe: 0.0,
      prediction_error
  )
  updated_stickiness_values = updated_stickiness_values.at[choice].set(
      current_stickiness_values[choice] + stickiness_learning_rate * base_chosen_stickiness_update - stickiness_reduction_for_negative_pe
  )

  # Introduce an "implicit negative learning" for unchosen options when the chosen option yields a positive PE.
  # If the chosen arm was surprisingly good (positive PE), reduce stickiness for unchosen arms.

  # Q-value modulated decay for stickiness: higher Q-values lead to slower decay.
  # This makes stickiness persist longer for options that are performing well.
  q_modulated_decay_factor = stickiness_learning_rate * (1.0 - jax.nn.sigmoid(new_q_values))
  updated_stickiness_values = updated_stickiness_values * (1.0 - q_modulated_decay_factor)

  # Clip stickiness values to a reasonable range, ensuring they don't go below zero.
  updated_stickiness_values = jnp.clip(updated_stickiness_values, 0.0, 5.0) # Fixed upper bound for stickiness values.

  # Update the state with the new stickiness values.
  agent_state = agent_state.at[5:9].set(updated_stickiness_values)

  # Calculate choice logits based on Q-values, scaled by dynamic inverse temperature.
  choice_logits = new_beta * agent_state[:4]
  # Add choice stickiness to the chosen option.
  choice_logits += agent_state[5:9]

  # Add an exploration bonus for options with higher surprise.
  exploration_bonus = pe_stickiness_influence * updated_surprise_values
  choice_logits += exploration_bonus

  # Add inverse stickiness (repulsion) for unchosen options to promote switching.
  inverse_stickiness_repulsion = -choice_stickiness_decay_rate * (1 - choice_one_hot)
  choice_logits += inverse_stickiness_repulsion

  # Introduce a recency-weighted reward for unchosen options into logits.
  # This encourages exploration of recently "good" but unchosen arms.
  # Create a decaying trace of recent rewards for each arm.
  # For the chosen arm, it's the current reward. For unchosen, it decays.
  # This is a new state variable, we assume it's initialized to zeros for all arms.
  # Given the current state structure, we need to add a new set of indices for this.
  # Let's assume for now this is handled by existing indices or a conceptual placeholder.
  # For implementation: this would require expanding agent_state.
  # To avoid changing state initialization for now, let's conceptualize it as
  # an influence from the *previous* trial's reward and chosen arm.
  return choice_logits, agent_state
  # EVOLVE-BLOCK-END


# Shortened to 100k steps then refactored, run 3, high floor.
def human_bandit_run3_simplified_high_floor(
    params: chex.Array,
    choice: int,
    reward: float,
    agent_state: Optional[chex.Array],
) -> tuple[chex.Array, chex.Array]:
  """Cognitive model describing human behavior on a multi-armed bandit task.

  Assumes the agent is presented with four options on each trial.

  Args:
    params: Fit parameters of the model.
    choice: Choice made by the agent on the previous trial. 0, 1, 2, or 3.
    reward: Reward received by the agent on the previous trial. A float between
      0 and 1.
    agent_state: The current state of the cognitive model.

  Returns:
    choice_logits: The probabilities that the agent will choose option 0, 1, 2,
      or 3 on the next trial, expressed as logits.
    agent_state: New agent state.
  """
  # Helper function to apply the sigmoid transformation.
  sigmoid = jax.nn.sigmoid

  # --- 1. Unpack and Transform Model Parameters ---
  # Parameters are transformed to ensure they fall within a sensible range.
  # Sigmoid maps them to (0, 1), suitable for rates and weights.
  # Softplus maps them to be positive, suitable for temperature.

  # Learning rate for the value of the chosen option.
  learning_rate_chosen = sigmoid(params[0])
  # Learning rate for the values of unchosen options.
  learning_rate_unchosen = sigmoid(params[1])
  # Decay rate for the recency trace of the chosen option.
  recency_chosen_decay = sigmoid(params[2])
  # Decay rate for the recency trace of unchosen options.
  recency_unchosen_decay = sigmoid(params[3])
  # Weight of the recency trace on the chosen option's choice probability.
  recency_weight_chosen = sigmoid(params[4])
  # Weight of the recency trace on the unchosen options' choice probabilities.
  recency_weight_unchosen = sigmoid(params[5])
  # Initial value estimate (Q-value) for all options.
  initial_q_value = sigmoid(params[6])
  # General rate at which Q-values decay towards zero (forgetting).
  value_decay_rate = sigmoid(params[7])
  # Base inverse temperature for the softmax choice rule. Higher values lead to
  # more deterministic (exploitative) choices. Clipped for stability.
  inverse_temperature_base = jnp.clip(jax.nn.softplus(params[8]), 0.1, 10.0)

  # --- 2. Initialize or Unpack Agent State ---
  # The agent's state consists of the Q-values and a recency trace for each option.
  if agent_state is None:
    # On the first trial, initialize Q-values and recency traces.
    q_values = jnp.full((4,), initial_q_value)
    recency_trace = jnp.zeros(4)
  else:
    # On subsequent trials, unpack the state from the previous trial.
    q_values = agent_state[:4]
    recency_trace = agent_state[4:8]

  # --- 3. Update Q-Values (Value Estimates) ---

  # a. Calculate the update for the CHOSEN option.
  # The prediction error is the difference between the actual reward and the expected reward.
  prediction_error_chosen = reward - q_values[choice]

  # A "novelty bonus" increases learning for options that haven't been chosen recently.
  novelty_bonus = 1.0 + jnp.exp(-recency_trace[choice] * 0.5)

  # The learning rate is modulated by this novelty bonus.
  modulated_learning_rate_chosen = jnp.clip(
      learning_rate_chosen * novelty_bonus, 0.0, 1.0)
  
  # The final update term for the chosen option's Q-value.
  chosen_q_value_update_term = modulated_learning_rate_chosen * prediction_error_chosen

  # b. Calculate the update for the UNCHOSEN options (fictive learning).
  # This is a form of counterfactual thinking: "what if I had chosen another option?".
  # The "error" is the difference between the chosen Q-value and the other Q-values.
  value_difference_from_chosen = q_values[choice] - q_values

  # This update is modulated by the "surprise" from the chosen option.
  # If the chosen outcome was very surprising, fictive learning is reduced.
  surprise_modulation = 1.0 - sigmoid(jnp.abs(prediction_error_chosen) * 2) / 2
  prediction_error_unchosen = value_difference_from_chosen * surprise_modulation

  # The update term for unchosen options is also boosted by their recency.
  recency_boost_unchosen = 1.0 + recency_trace
  unchosen_q_value_update_term = (
      learning_rate_unchosen * recency_boost_unchosen * prediction_error_unchosen
  )

  # c. Apply all updates to the Q-values.
  # First, all Q-values decay towards zero, modeling forgetting.
  # The decay is stronger for options that haven't been chosen recently.
  recency_modulated_decay = jnp.clip(value_decay_rate * (1 + recency_trace * 0.1), 0.0, 1.0)
  q_values = q_values * recency_modulated_decay

  # Add the update term to the chosen option's Q-value.
  q_values = q_values.at[choice].add(chosen_q_value_update_term)

  # Add the update terms to all unchosen options' Q-values.
  # Note: The chosen option's term in `unchosen_q_value_update_term` will be zero.
  q_values = q_values + unchosen_q_value_update_term

  # --- 4. Update Recency Traces ---
  # The recency trace tracks how recently each option has been chosen.

  # a. First, perform a simple decay and increment.
  # All traces decay.
  recency_trace = recency_trace * recency_chosen_decay
  # The chosen option's trace is incremented by 1.
  recency_trace = recency_trace.at[choice].add(1.0)

  # b. Then, perform a second, more complex, state-dependent decay.
  # The decay for the chosen option is modulated by "surprise" (reward prediction error).
  # Positive surprise (reward > expectation) slows decay; negative surprise speeds it up.
  surprise_modulated_recency_chosen_decay = (
      recency_chosen_decay + (0.5 - reward) * (1 - recency_chosen_decay)
  )

  # The decay for unchosen options depends on their recency and value difference from the chosen option.
  unchosen_recency_factor = 1.0 - recency_trace / 5.0
  unchosen_value_diff_factor = 2.0 - jnp.abs(q_values[choice] - q_values)
  decay_factor_unchosen = jnp.clip(
      recency_unchosen_decay * unchosen_recency_factor * unchosen_value_diff_factor,
      0.0, 1.0
  )

  # This term adds a novelty-based component to the unchosen decay.
  unchosen_novelty_decay_component = jnp.exp(-recency_trace * 0.5)
  final_unchosen_decay_factor = (
      decay_factor_unchosen +
      (1.0 - decay_factor_unchosen) * unchosen_novelty_decay_component
  )

  # Apply the separate decay factors to the chosen and unchosen options.
  recency_trace = jnp.where(
      jnp.arange(4) == choice,
      recency_trace * surprise_modulated_recency_chosen_decay, # Decay for chosen
      recency_trace * final_unchosen_decay_factor,             # Decay for unchosen
  )

  # --- 5. Calculate Choice Logits for the Next Trial ---
  # Logits are calculated based on Q-values, recency, and exploration parameters.

  # a. Calculate a recency bonus.
  # The recency trace is weighted differently for the chosen vs. unchosen options.
  weighted_recency_trace = jnp.where(
      jnp.arange(4) == choice,
      recency_weight_chosen * recency_trace,
      recency_weight_unchosen * recency_trace,
  )
  # This creates a "stickiness" or recency effect on choice.
  recency_bonus = 1.0 + weighted_recency_trace

  # b. Calculate an adaptive, option-specific inverse temperature.
  # The base temperature is adapted based on surprise, promoting exploration after unexpected outcomes.
  surprise_based_temperature_scaling = 1.0 + jnp.abs(prediction_error_chosen) * 0.5
  adaptive_inverse_temperature = inverse_temperature_base / surprise_based_temperature_scaling

  # Exploration is also boosted when there is high uncertainty about Q-values (high standard deviation).
  uncertainty_exploration_boost = 1.0 + jnp.exp(-jnp.std(q_values) * 2.5)

  # The final inverse temperature is option-specific, modulated by fictive errors and uncertainty.
  fictive_error_scaling = 1.0 + jnp.abs(prediction_error_unchosen)
  option_specific_inverse_temperatures = (
      adaptive_inverse_temperature / fictive_error_scaling * uncertainty_exploration_boost
  )

  # c. Combine all components to get the final choice logits.
  # The final value is a product of the Q-value, the recency bonus, and the temperature scaling.
  choice_logits = (
      q_values * recency_bonus * option_specific_inverse_temperatures
  )

  # --- 6. Package the New Agent State ---
  # The new state is a concatenation of the updated Q-values and recency traces.
  agent_state = jnp.concatenate((q_values, recency_trace))

  return choice_logits, agent_state


def human_bandit_synthesis(
    params: chex.Array,
    choice: int,
    reward: float,
    agent_state: chex.Array | None,
) -> tuple[chex.Array, chex.Array]:
  """Handcrafted agent for the Human Bandit task based on insights from discovered programs.

  Has the following elements
  * Maintains separate q_values, choice_trace, and average_reward state variables.
  * Updates q_values on chosen by replacing with recent reward.
  * Updates q_values with decay toward average reward on unchosen
  * Increment choice_trace for choice, clip at 5. Quadratic reset on unchosen.
  * Nonlinearity on q-values: rel_q * softplus(rel_q + 1)
  """

  # === 1. Unpack and Transform Model Parameters ===
  # Apply a sigmoid function to the first 8 parameters to constrain them
  # between 0 and 1. This is common for rates, weights, and probabilities.

  (raw_initial_q_value,
   raw_inverse_temperature_base,
   learning_rate_unchosen_logit,
   recency_chosen_decay_logit,
   recency_unchosen_decay_logit,
   recency_weight_chosen,
   *_) = params

  initial_q_value = jax.nn.sigmoid(raw_initial_q_value)
  inverse_temperature_base = 10 * jax.nn.sigmoid(raw_inverse_temperature_base)
  learning_rate_unchosen = 0.9 * jax.nn.sigmoid(learning_rate_unchosen_logit) + 0.1
  recency_chosen_decay = jax.nn.sigmoid(recency_chosen_decay_logit)
  recency_unchosen_decay = 0.95 * jax.nn.sigmoid(recency_unchosen_decay_logit) + 0.025

  # === 2. Initialize or Load Agent's Internal State ===
  # The agent's state consists of Q-values (expected reward for each option),
  # a recency trace (memory of recent choices), and average reward
  if agent_state is None:
    # If this is the first trial, initialize Q-values and recency trace.
    q_values = jnp.full(shape=(4,), fill_value=initial_q_value)
    choice_trace = jnp.zeros(shape=(4,))
    average_reward = initial_q_value
  else:
    # If not the first trial, load the state from the previous trial.
    q_values = agent_state[:4]       # First 4 elements are Q-values
    choice_trace = agent_state[4:8]  # Last 4 elements are recency traces
    average_reward = agent_state[8]

  # === 3. Update Q-Values Based on Last Trial's Outcome ===

  # 3a. Update the estimate of the average reward.
  # This tracks the overall reward rate of the environment.

  avg_reward_prediction_error = reward - average_reward
  average_reward += learning_rate_unchosen * avg_reward_prediction_error

  q_values += (
      learning_rate_unchosen * (average_reward - q_values)
  )

  # Update the Q-value for the chosen option by replacing with recent reward
  q_values = q_values.at[choice].set(reward)

  # === 4. Update Recency Trace ===
  # The recency trace tracks which options were chosen recently.

  # Create a one-hot vector to easily distinguish the chosen option.
  chosen_one_hot = jax.nn.one_hot(choice, num_classes=4)

  # Update recency trace.
  recency_update_for_chosen = (
      jnp.minimum(choice_trace + 1.0, 5.0) * recency_chosen_decay
  )
  recency_update_for_unchosen = (
      choice_trace * recency_unchosen_decay * (1 - choice_trace / 5.0)
  )

  # Apply the appropriate update rule to each element of the recency trace.
  choice_trace = jnp.where(
      chosen_one_hot,
      recency_update_for_chosen,
      recency_update_for_unchosen,
  )

  # === 5. Compute Choice Logits for the Next Trial ===
  # Logits are the raw, unnormalized scores for each choice. A higher logit
  # means a higher probability of being chosen.

  choice_trace_component = recency_weight_chosen * choice_trace  # one weight on all.

  # Nonlinearity on q-values.
  relative_q_values = q_values - average_reward
  value_based_temperature_scaling = jax.nn.softplus(relative_q_values + 1.0) + 0.01
  adaptive_inverse_temperature = inverse_temperature_base * value_based_temperature_scaling
  value_component = adaptive_inverse_temperature * relative_q_values

  choice_logits = value_component + choice_trace_component

  # === 6. Prepare State for the Next Trial and Return ===

  # Concatenate the updated Q-values and recency trace into a single array.
  agent_state = jnp.concatenate((q_values, choice_trace, average_reward[jnp.newaxis]))

  return choice_logits, agent_state
```
