## Supplementary material for "AI-Discovered Cognitive Models Reveal Novel Insights into Human and Animal Learning": DataDIVER Discovered Programs: monkey_bandit_models.py

```
# Copyright 2026 DeepMind Technologies Limited.
#
# Licensed under the Apache License, Version 2.0 (the "License");
# you may not use this file except in compliance with the License.
# You may obtain a copy of the License at
#
#     http://www.apache.org/licenses/LICENSE-2.0
#
# Unless required by applicable law or agreed to in writing, software
# distributed under the License is distributed on an "AS IS" BASIS,
# WITHOUT WARRANTIES OR CONDITIONS OF ANY KIND, either express or implied.
# See the License for the specific language governing permissions and
# limitations under the License.

"""Saved models for the Nature 2026 submission."""
from typing import Optional, Tuple

import chex
import jax
import jax.numpy as jnp


def monkey_bandit_run1_fit_only(
    params: chex.Array,
    choice: int,
    reward: float,
    novel_option: int,
    agent_state: Optional[chex.Array],
) -> tuple[chex.Array, chex.Array]:
  """Cognitive model describing monkey behavior on a multi-armed bandit task.

  Assumes the monkey is presented with three options on each trial.
  Occasionally, one of these options is changed to a novel arm with different
  reward probabilities, when this happens novel_option will indicate the index
  of the newly novel option and otherwise it will be -1.

  Args:
    params: Model params. Different parameters are used for different monkeys.
    choice: The choice made by the subject on the previous trial. An
      integer with values of 0, 1, or 2.
    reward: The reward received by the subject on this trial. A scalar
      either 0 or 1.
    novel_option: The choice option that is novel to the subject on this trial.
      The number 0, 1, 2 indicates one of the choices, and -1 indicates that
      no option is currently novel.
    agent_state: The current state of the cognitive model.

  Returns:
    choice_logits: The probabilities that the subject will choose option
      0, 1, or 2 on the next trial, expressed as logits.
    agent_state: The updated state of the cognitive model.
  """
  # Do not remove comments or TODOs from this program.

  if agent_state is None:
    agent_state = jnp.zeros((6,))  # Initialize Q-values and novelty values for three options

  # Define all parameters.
  base_learning_rate_chosen = jax.nn.sigmoid(params[0])
  beta_base = jax.nn.softplus(params[1])
  fictive_reward_context_scale = jax.nn.sigmoid(params[4])
  novelty_initial_bonus = jax.nn.softplus(params[5])
  novelty_decay_rate = jax.nn.sigmoid(params[6])
  novelty_learning_rate_scale = params[7]
  beta_avg_q_scale = jax.nn.sigmoid(params[8])
  beta_spread_scale = jax.nn.sigmoid(params[9])
  base_learning_rate_unchosen = jax.nn.sigmoid(params[3])

  # agent_state[0:3] = Q-values, agent_state[3:6] = novelty_values
  q_values, novelty_values = agent_state[0:3], agent_state[3:6]

  # Initialize Q-values and novelty bonus for the novel option.
  other_q_values_mask = (novel_option != jnp.arange(3))
  # Set the Q-value for the novel option, using the context-adjusted initial Q.
  # Calculate the average Q-value of the non-novel options.
  average_other_q = jnp.sum(q_values * other_q_values_mask) / 2.0
  # The initial Q for a novel option is a weighted average of a fixed value (params[2])
  # and the average Q of other options.
  initial_q_context_adjusted = (1 - beta_avg_q_scale) * params[2] + beta_avg_q_scale * average_other_q
  q_values = jnp.where(novel_option == jnp.arange(3), initial_q_context_adjusted, q_values)
  novelty_values = jnp.where(
      novel_option == jnp.arange(3), novelty_initial_bonus, novelty_values
  )
  novelty_values = novelty_values * (1 - novelty_decay_rate)

  # Update Q-values for unchosen options using fictive learning.
  # Fictive reward context is a weighted average of the actual reward and a baseline (params[2]).
  fictive_reward_context = (
      reward * (1 - fictive_reward_context_scale) + params[2] * fictive_reward_context_scale
  )
  fictive_prediction_error = fictive_reward_context - q_values
  unchosen_mask = jnp.arange(3) != choice
  novelty_scaled_learning_rate = novelty_learning_rate_scale * novelty_values
  learning_rates_for_unchosen = (
      base_learning_rate_unchosen + novelty_scaled_learning_rate * jnp.abs(fictive_prediction_error)
  )
  q_values = jnp.where(unchosen_mask, q_values + learning_rates_for_unchosen * fictive_prediction_error, q_values)

  # Update the Q-value of the chosen option.
  prediction_error = reward - q_values[choice]
  learning_rate_for_chosen = (
      base_learning_rate_chosen + novelty_scaled_learning_rate[choice] * jnp.abs(prediction_error)
  )
  q_values = q_values.at[choice].set(q_values[choice] + learning_rate_for_chosen * prediction_error)
  q_values_with_novelty = q_values + novelty_values
  avg_abs_q = jnp.mean(jnp.abs(q_values))
  q_spread = jnp.max(q_values) - jnp.min(q_values)
  beta = beta_base + beta_avg_q_scale * avg_abs_q + beta_spread_scale * q_spread

  # Compute choice_logits using a softmax over the Q-values.
  choice_logits = beta * q_values_with_novelty

  # Update the full agent_state for the next trial.
  agent_state = jnp.concatenate((q_values, novelty_values))

  return choice_logits, agent_state
  

def monkey_bandit_run2_fit_only(
    params: chex.Array,
    choice: int,
    reward: float,
    novel_option: int,
    agent_state: Optional[chex.Array],
) -> tuple[chex.Array, chex.Array]:
  """Cognitive model describing monkey behavior on a multi-armed bandit task.

  Assumes the monkey is presented with three options on each trial.
  Occasionally, one of these options is changed to a novel arm with different
  reward probabilities, when this happens novel_option will indicate the index
  of the newly novel option and otherwise it will be -1.

  Args:
    params: Model params. Different parameters are used for different monkeys.
    choice: The choice made by the subject on the previous trial. An
      integer with values of 0, 1, or 2.
    reward: The reward received by the subject on this trial. A scalar
      either 0 or 1.
    novel_option: The choice option that is novel to the subject on this trial.
      The number 0, 1, 2 indicates one of the choices, and -1 indicates that
      no option is currently novel.
    agent_state: The current state of the cognitive model.

  Returns:
    choice_logits: The probabilities that the subject will choose option
      0, 1, or 2 on the next trial, expressed as logits.
    agent_state: The updated state of the cognitive model.
  """
  # Do not remove comments or TODOs from this program.

  # Initialize agent state: first 3 elements are Q-values, next 3 are novelty bonuses.
  # Q-values are initialized to prior_q_initial_value, novelty bonuses to novelty_baseline_value.
  if agent_state is None:
    # Define parameters needed for initialization early.
    prior_q_initial_value = params[4]
    novelty_baseline_value = params[8]
    agent_state = jnp.concatenate([
        jnp.full((3,), prior_q_initial_value),
        jnp.full((3,), novelty_baseline_value)
    ])

  # Define all parameters.
  positive_learning_rate = jax.nn.sigmoid(params[0])
  negative_learning_rate = jax.nn.sigmoid(params[1])
  unchosen_initial_learning_rate = jax.nn.sigmoid(params[2])
  inverse_temperature = jnp.exp(params[3])
  unchosen_q_baseline = params[5]
  q_variance_scaler = params[6]
  novelty_bonus_decay_rate = jax.nn.softplus(params[7])
  novelty_baseline_value = params[8]
  novel_q_initial_value = params[9]
  prediction_error = reward - agent_state[choice]
  chosen_novelty_bonus = agent_state[choice + 3]

  # Modulate prediction error by novelty bonus, enhancing learning for novel options.
  modulated_prediction_error = prediction_error * (1.0 + jax.nn.softplus(chosen_novelty_bonus))
  chosen_learning_rate = jnp.where(prediction_error > 0, positive_learning_rate, negative_learning_rate) * jnp.sqrt(jnp.abs(prediction_error))
  q_chosen_update = chosen_learning_rate * modulated_prediction_error
  q_chosen = agent_state[choice] + q_chosen_update
  agent_state = agent_state.at[choice].set(q_chosen)

  unchosen_mask = jnp.ones_like(agent_state, dtype=bool).at[choice].set(False)
  q_unchosen_decay_rate = unchosen_initial_learning_rate / 2.0
  # Unchosen Q-values decay towards a baseline and are updated vicariously.
  agent_state = jnp.where(
      unchosen_mask,
      agent_state * (1 - q_unchosen_decay_rate) + unchosen_q_baseline * q_unchosen_decay_rate + unchosen_initial_learning_rate * 0.2 * reward,
      agent_state
  )

  is_novel_option = novel_option != -1

  # Update novelty bonuses: decay for unchosen options
  novelty_bonuses = agent_state[3:6]
  novelty_bonuses = jnp.where(
      jnp.arange(3) == choice,
      novelty_bonuses[choice],
      novelty_bonuses * (1 - novelty_bonus_decay_rate) + novelty_baseline_value * novelty_bonus_decay_rate
  )
  agent_state = agent_state.at[3:6].set(novelty_bonuses)

  # If there's a novel option, re-initialize its Q-value and novelty bonus
  agent_state = jnp.where(
      is_novel_option,
      agent_state.at[novel_option].set(novel_q_initial_value).at[novel_option + 3].set(jax.nn.sigmoid(novel_q_initial_value)),
      agent_state
  )

  effective_q_values = agent_state[0:3] + agent_state[3:6]
  q_value_variance = jnp.var(agent_state[0:3]) + 1e-6
  base_inverse_temperature = inverse_temperature * (1 + q_variance_scaler * q_value_variance)
  option_inverse_temperatures = jnp.where(
      jnp.arange(3) == novel_option,
      base_inverse_temperature * (1.0 - novelty_bonus_decay_rate),
      base_inverse_temperature
  )

  choice_logits = effective_q_values * option_inverse_temperatures

  return choice_logits, agent_state
  

def monkey_bandit_run3_fit_only(
    params: chex.Array,
    choice: int,
    reward: float,
    novel_option: int,
    agent_state: Optional[chex.Array],
) -> tuple[chex.Array, chex.Array]:
  """Cognitive model describing monkey behavior on a multi-armed bandit task.

  Assumes the monkey is presented with three options on each trial.
  Occasionally, one of these options is changed to a novel arm with different
  reward probabilities, when this happens novel_option will indicate the index
  of the newly novel option and otherwise it will be -1.

  Args:
    params: Model params. Different parameters are used for different monkeys.
    choice: The choice made by the subject on the previous trial. An
      integer with values of 0, 1, or 2.
    reward: The reward received by the subject on this trial. A scalar
      either 0 or 1.
    novel_option: The choice option that is novel to the subject on this trial.
      The number 0, 1, 2 indicates one of the choices, and -1 indicates that
      no option is currently novel.
    agent_state: The current state of the cognitive model.

  Returns:
    choice_logits: The probabilities that the subject will choose option
      0, 1, or 2 on the next trial, expressed as logits.
    agent_state: The updated state of the cognitive model.
  """
  # Do not remove comments or TODOs from this program.

  # Initialize agent state if not provided.
  if agent_state is None:
    # Stores Q-values, choice counts, predictability estimates, and prediction errors.
    # The long-term prediction error dimension is removed for simplification.
    agent_state = jnp.zeros((3, 4))
    # Initialize prediction errors to a neutral value.
    agent_state = agent_state.at[:, 3].set(0.5)

  # Define all parameters.
  # Each parameter assignment is a single line.
  # Change the names of the parameters to be descriptive.
  # Include comments describing the parameter transformation, if applicable.
  learning_rate = jax.nn.sigmoid(params[0])  # Base learning rate for updating expected values, constrained to (0,1).
  beta = jnp.exp(params[1])  # Inverse temperature for softmax choice, constrained to be positive.
  forgetting_rate_base = jax.nn.sigmoid(params[2]) # Base forgetting rate for unchosen options, constrained to (0,1).
  prior_initial_q = params[3] # Prior or default initial value for novel options.
  baseline_q_value = params[4] # Baseline value for unchosen options to decay towards.
  choice_count_decay_modifier = jnp.exp(params[5]) # Parameter controlling how choice counts modify forgetting, constrained to be positive.
  value_weighting_beta = jnp.exp(params[6]) # Inverse temperature for softmax weighting of Q-values for novel option initialization, constrained to be positive.
  beta_scale = jnp.exp(params[7]) # Scaling factor for dynamic beta based on Q-value variance, constrained to be positive.
  exploration_bonus_strength = jnp.exp(params[8]) # Strength of the uncertainty-driven exploration bonus, constrained to be positive.
  recency_weight = params[9] # Weight for the recency bias, boosting or depressing the last chosen option.

  # Update agent_state (Q-values, choice counts, predictability estimates, and prediction errors) based on previous choice and reward.
  chosen_mask = jnp.arange(3) == choice
  q_values = agent_state[:, 0]
  choice_counts = agent_state[:, 1]
  predictability_estimates = agent_state[:, 2] # Predictability estimates for each option.
  prediction_errors = agent_state[:, 3] # Stores recent short-term prediction errors for each option.

  # Determine the learning rate for Q-value updates.
  # This learning rate is modulated by Q-value variance, inverse choice count, and average prediction error.
  q_learning_rate = learning_rate * (1 + beta_scale * jnp.var(q_values)) / (1 + jnp.sqrt(choice_counts + 1)) * (1 + jnp.mean(jnp.abs(prediction_errors)))

  # Update Q-values for the chosen option, with an eligibility trace for unchosen options.
  eligibility_trace_strength = 0.1 # Base strength of the eligibility trace for unchosen options.
  q_delta = q_learning_rate * (reward - q_values)
  # Modulate eligibility trace for unchosen options by the absolute prediction error of the chosen option.
  eligibility_trace_modifier = jnp.abs(reward - q_values[choice])
  q_values = q_values + q_delta * chosen_mask + q_delta * (1 - chosen_mask) * eligibility_trace_strength * eligibility_trace_modifier

  # Determine the learning rate for predictability updates.
  # This learning rate is modulated by current predictability and short-term prediction errors.
  predictability_learning_rate = learning_rate * 0.5 * (1 - predictability_estimates) * (1 + jnp.abs(prediction_errors))

  # Update predictability estimates for the chosen option based on prediction error magnitude.
  predictability_delta = predictability_learning_rate * (1 - jnp.abs(reward - q_values))
  # Apply the same modulated eligibility trace for predictability estimates.
  predictability_estimates = predictability_estimates + predictability_delta * chosen_mask + predictability_delta * (1 - chosen_mask) * eligibility_trace_strength * eligibility_trace_modifier
  predictability_estimates = jnp.clip(predictability_estimates, 0.0, 1.0)

  # Apply adaptive forgetting to the chosen option if there's a negative prediction error, scaled by error magnitude and Q-value variance.
  chosen_forgetting_rate = jnp.where(chosen_mask * (reward - q_values < 0),
                                     forgetting_rate_base * jnp.abs(reward - q_values) * jnp.var(q_values), 0.0)
  q_values = q_values * (1 - chosen_forgetting_rate) + chosen_forgetting_rate * baseline_q_value

  # Update choice counts for the chosen option.
  choice_counts = choice_counts + chosen_mask

  # Calculate forgetting rate for unchosen options, inversely proportional to choice count.
  forgetting_modifier = forgetting_rate_base / (1 + choice_count_decay_modifier * choice_counts)
  forgetting_rate_for_unchosen = forgetting_modifier * (1 - chosen_mask)

  # Calculate the mean Q-value and a discrepancy factor for adaptive forgetting.
  mean_q_value = jnp.mean(q_values)
  q_discrepancy_factor = 1 + jnp.abs(q_values - mean_q_value) * 0.1

  # Estimate local Q-value volatility.
  q_value_volatility = jnp.abs(q_values - mean_q_value) * 0.5

  # Apply adaptive forgetting to unchosen options, scaled by Q-value discrepancy and volatility.
  adaptive_forgetting_rate = forgetting_rate_for_unchosen * q_discrepancy_factor * (1 + q_value_volatility)
  q_values = q_values * (1 - adaptive_forgetting_rate * jnp.abs(q_values - baseline_q_value)) + \
             adaptive_forgetting_rate * baseline_q_value * jnp.abs(q_values - baseline_q_value)

  # Decay choice counts for unchosen options and apply a global decay.
  choice_count_decay_rate_unchosen = forgetting_rate_base / (1 + choice_count_decay_modifier * choice_counts) * q_discrepancy_factor * (1 - chosen_mask)
  global_choice_count_decay = forgetting_rate_base * 0.1
  choice_counts = jnp.maximum(0, choice_counts - choice_count_decay_rate_unchosen - global_choice_count_decay)

  # Apply a simpler decay to predictability estimates for all options.
  predictability_estimates = predictability_estimates * (1 - forgetting_rate_base * 0.05)
  predictability_estimates = jnp.clip(predictability_estimates, 0.1, 1.0)

  # Apply a simpler decay to prediction errors for all options, introducing a dual-decay mechanism.
  # Apply a simpler decay to prediction errors for all options.
  # The fast and slow decay distinction is removed to simplify the model.
  fast_decay_rate = forgetting_rate_base * 0.1
  prediction_errors = jnp.maximum(0.0, prediction_errors * (1 - fast_decay_rate))

  # Handle novel options by resetting Q-value, choice count, predictability, and prediction errors.
  novel_mask = jnp.arange(3) == novel_option
  non_novel_mask = ~novel_mask
  average_q_non_novel = jnp.sum(q_values * non_novel_mask) / (jnp.sum(non_novel_mask) + 1e-6)

  # Dynamically adjust prior_initial_q based on recent reward history and Q-value variance.
  recent_reward_history_influence = jnp.mean(q_values)
  q_value_variance_influence = jnp.var(q_values)

  # Compute an adaptive prior for novel options.
  adaptive_prior_for_novel = prior_initial_q + (recent_reward_history_influence - 0.5) * 0.5 - q_value_variance_influence * 0.1

  # Initialize the novel option's Q-value by blending the adaptive prior with existing Q-values.
  q_value_weights = jax.nn.softmax(q_values * value_weighting_beta)
  value_weighted_average_q = jnp.sum(q_values * q_value_weights)
  novel_q_initialization_value = (adaptive_prior_for_novel + average_q_non_novel + value_weighted_average_q) / 3.0
  q_values = jnp.where(novel_mask, novel_q_initialization_value, q_values)
  choice_counts = jnp.where(novel_mask, 0, choice_counts)
  predictability_estimates = jnp.where(novel_mask, 0.0, predictability_estimates) # Novel options start with maximal unpredictability (0.0).
  prediction_errors = jnp.where(novel_mask, 1.0, prediction_errors) # Strong initial surprise for short-term PE.

  # Recombine components into the agent_state. Note the removal of prediction_errors_long_term.
  agent_state = jnp.stack([q_values, choice_counts, predictability_estimates, prediction_errors], axis=-1)

  # Dynamically adjust beta: higher standard deviation of Q-values leads to higher beta, and
  # higher average predictability leads to higher beta (more exploitation).
  average_predictability = jnp.mean(predictability_estimates)
  dynamic_beta = beta * (1 + beta_scale * jnp.std(q_values) + average_predictability)

  # Add an uncertainty-driven exploration bonus, scaled by the variance of predictability estimates.
  predictability_uncertainty_factor = (1 + jnp.var(predictability_estimates))
  uncertainty_bonus = exploration_bonus_strength * (1 + (1 - predictability_estimates)) / (1 + jnp.sqrt(choice_counts + 1)) * predictability_uncertainty_factor

  # Compute choice_logits using softmax over Q-values with dynamic beta, uncertainty bonus, and recency bias.
  recency_bonus_chosen = recency_weight * chosen_mask * (reward - 0.5) * 4
  choice_logits = dynamic_beta * q_values + uncertainty_bonus + recency_bonus_chosen

  return choice_logits, agent_state


# Shortened to 100k steps then refactored, run 1, low floor.
def monkey_bandit_run1_simplified_low_floor(
    params: chex.Array,
    choice: int,
    reward: float,
    novel_option: int,
    agent_state: Optional[chex.Array],
) -> tuple[chex.Array, chex.Array]:
  """
  Calculates agent's action logits and updates its internal state.

  The agent uses a combination of Q-learning and a novelty bonus system.
  It updates Q-values based on reward, decays unchosen options towards a
  baseline, and manages novelty bonuses that decay over time.
  """
  # --- 1. Unpack Parameters and State ---

  # Unpack the agent's fixed parameters.
  # Note: `exploration_sensitivity` is unpacked but not used in the original logic.
  baseline_q_value, learning_rate, base_inverse_temperature, \
      exploration_sensitivity, decay_to_baseline_rate, \
      novelty_bonus_magnitude, novelty_bonus_retention, *_ = params  # stachenfeld@ added *_ to ignore unused params

  # Initialize the agent's state if this is the first step (state is None).
  # The state vector contains Q-values and novelty bonuses for 3 options.
  if agent_state is None:
    current_state = jnp.zeros(6)
  else:
    current_state = agent_state

  # Unpack the state into separate arrays for Q-values and novelty bonuses.
  q_values, novelty_bonuses = jnp.split(current_state, 2)

  # --- 2. Update Novelty Bonuses ---

  # Create a one-hot mask to identify which option, if any, is newly introduced.
  # A value of -1 for `novel_option` means no new option was introduced,
  # and jax.nn.one_hot will correctly produce a vector of all zeros.
  is_novel_mask = jax.nn.one_hot(novel_option, num_classes=3)

  # For any newly introduced option, its novelty bonus is set to a fixed magnitude.
  new_option_bonus = novelty_bonus_magnitude

  # For all existing options, their novelty bonuses are decayed.
  decayed_existing_bonuses = novelty_bonuses * novelty_bonus_retention

  # Use the mask to apply the correct update: reset for the novel option, decay for others.
  novelty_bonuses = jnp.where(is_novel_mask, new_option_bonus, decayed_existing_bonuses)

  # --- 3. Update Q-Values ---

  # This update logic has three distinct cases handled by nested `where` clauses:
  #   a) The option was newly introduced.
  #   b) The option was chosen.
  #   c) The option was available but not chosen.

  # First, calculate the updates for existing options (cases b and c).
  is_chosen_mask = jax.nn.one_hot(choice, num_classes=3)
  
  # For the CHOSEN option (b), apply a standard Q-learning update.
  prediction_error = reward - q_values
  q_learning_update = learning_rate * prediction_error
  
  # For UNCHOSEN options (c), decay their values back towards the baseline.
  decay_update = decay_to_baseline_rate * (baseline_q_value - q_values)
  
  # Combine updates for chosen and unchosen options.
  updates_for_existing_options = jnp.where(is_chosen_mask, q_learning_update, decay_update)
  updated_q_for_existing_options = q_values + updates_for_existing_options
  
  # Now, handle the NOVEL option case (a).
  # A newly introduced option's Q-value is initialized to the baseline.
  new_option_q_value = baseline_q_value

  # Use the novelty mask to select the final Q-values:
  # - If an option is new, set its Q-value to the baseline.
  # - Otherwise, use the updated value for existing options calculated above.
  q_values = jnp.where(is_novel_mask, new_option_q_value, updated_q_for_existing_options)

  # --- 4. Calculate Outputs and Next State ---

  # The final value of an option is its Q-value plus its novelty bonus.
  total_values = q_values + novelty_bonuses

  # Calculate a dynamic inverse temperature for the softmax calculation.
  # This makes the agent's choices more exploratory when Q-values are similar.
  variance_of_q_values = jnp.var(q_values)
  dynamic_inverse_temperature = base_inverse_temperature + variance_of_q_values

  # Action logits are the inputs to a softmax function for choice probability.
  action_logits = total_values * dynamic_inverse_temperature

  # The next agent state is the concatenation of the newly updated values.
  next_agent_state = jnp.concatenate((q_values, novelty_bonuses))

  return action_logits, next_agent_state


# Shortened to 100k steps then refactored, run 2, low floor.
def monkey_bandit_run2_simplified_low_floor(
    params: chex.Array,
    choice: int,
    reward: float,
    novel_option: int,
    agent_state: Optional[chex.Array],
) -> tuple[chex.Array, chex.Array]:
  """
  Updates agent state based on action and reward, and computes next action logits.

  This agent uses a Q-learning-style update rule that includes:
  1. A decay mechanism that pulls all Q-values towards a baseline.
  2. A standard learning update for the chosen action based on prediction error.
  3. A novelty mechanism that resets the value of new options.
  4. An exploration strategy that considers Q-value variance and novelty.
  """

  # --- 1. Initialization and Parameter Unpacking ---

  # If this is the first step (agent_state is None), initialize Q-values to zeros.
  # The agent's state represents the estimated Q-value for each of the 3 options.
  q_values = jnp.zeros((3,)) if agent_state is None else agent_state

  # Unpack the agent's learnable parameters into named variables for clarity.
  (learning_rate_logit,
   inverse_temperature,
   novelty_bonus,
   decay_rate_logit,
   baseline_q_value,
   q_variance_scaler,
   novelty_exploration_bonus, *_) = params  # stachenfeld@ added *_ to ignore unused params

  # Convert logits to rates (values between 0 and 1) using the sigmoid function.
  learning_rate = jax.nn.sigmoid(learning_rate_logit)
  decay_rate = jax.nn.sigmoid(decay_rate_logit)

  # --- 2. Q-Value Updates ---

  # Step 2a: Calculate the standard Q-learning update for the chosen action.
  # The update is based on the prediction error (the difference between the
  # actual reward and the expected reward/Q-value).
  prediction_error = reward - q_values[choice]
  updated_q_for_choice = q_values[choice] + learning_rate * prediction_error

  # Step 2b: Apply a decay to all Q-values, pulling them towards a baseline.
  # This models forgetting or a natural reversion to a mean value over time.
  q_values_after_decay = q_values + decay_rate * (baseline_q_value - q_values)

  # Step 2c: Combine the updates. The decayed values are used for all unchosen
  # options, while the newly learned value is used for the chosen option.
  q_values_after_learning = q_values_after_decay.at[choice].set(updated_q_for_choice)

  # Step 2d: Apply a novelty bonus. The Q-value of a newly introduced "novel"
  # option is reset to a specific `novelty_bonus` value.
  is_novel_option_mask = jax.nn.one_hot(novel_option, num_classes=3)

  # This calculation effectively sets the Q-value of the novel option to `novelty_bonus`.
  # For the novel option (where mask=1): new_q = old_q + 1 * (bonus - old_q) = bonus.
  # For other options (where mask=0): new_q = old_q + 0 * (...) = old_q.
  q_values_after_novelty_bonus = q_values_after_learning + is_novel_option_mask * (
      novelty_bonus - q_values_after_learning
  )

  # The final, updated Q-values will become the new state for the next step.
  new_agent_state = q_values_after_novelty_bonus

  # --- 3. Compute Action Selection Logits for the Next Step ---

  # Step 3a: Calculate base logits by scaling the new Q-values by an inverse temperature.
  # A higher inverse_temperature leads to more deterministic choices.
  base_logits = new_agent_state * inverse_temperature

  # Step 3b: Modulate the logits by the variance of the Q-values. This is a form of
  # uncertainty-based exploration: higher variance in Q-values can be used to
  # make the agent's choices more deterministic (less random).
  q_value_variance = jnp.var(new_agent_state)
  variance_scaling_factor = 1 + q_variance_scaler * q_value_variance
  scaled_logits = base_logits * variance_scaling_factor

  # Step 3c: Add a direct exploration bonus to the novel option.
  # This encourages the agent to select the novel option, regardless of its Q-value.
  final_logits = scaled_logits + novelty_exploration_bonus * is_novel_option_mask

  # --- 4. Return Results ---

  # Return the computed logits (to be used for the next action selection)
  # and the updated agent state for the next iteration.
  return final_logits, new_agent_state


# Shortened to 100k steps then refactored, run 3, low floor.
def monkey_bandit_run3_simplified_low_floor(
    params: chex.Array,
    choice: int,
    reward: float,
    novel_option: int,
    agent_state: Optional[chex.Array],
) -> Tuple[chex.Array, chex.Array]:
  """
  Updates agent state based on a choice and reward, and computes action logits.

  This agent uses a Q-learning-like update rule, incorporates a novelty bonus
  for exploration, and adjusts its decision-making temperature based on the
  variance of its Q-values.

  Args:
    params: An array containing the agent's parameters:
      - initial_q_value: The starting Q-value for new options.
      - learning_rate: The rate at which Q-values are updated.
      - base_inverse_temperature: The base temperature for the softmax policy.
      - inverse_temperature_sensitivity_to_variance: How much Q-value variance
        affects the policy temperature.
      - novelty_bonus_magnitude: The initial bonus for exploring a new option.
      - novelty_bonus_decay: The rate at which the novelty bonus decreases.
    choice: The index of the action taken by the agent.
    reward: The reward received for the chosen action.
    novel_option: The index of an option that is newly available.
    agent_state: The internal state of the agent, containing Q-values and
      novelty bonus states. None if this is the first step.

  Returns:
    A tuple containing:
      - choice_logits: The logits for the next action selection.
      - agent_state: The updated internal state of the agent.
  """
  # --- 1. Unpack Parameters and State ---

  # Unpack the agent's parameters for easier access.
  initial_q_value = params[0]
  learning_rate = params[1]
  base_inverse_temperature = params[2]
  inverse_temperature_sensitivity_to_variance = params[3]
  novelty_bonus_magnitude = params[4]
  novelty_bonus_decay = params[5]

  # Initialize state if it's the first step; otherwise, unpack the existing state.
  # The state consists of two arrays: one for Q-values and one for the novelty bonus.
  if agent_state is None:
    # `jnp.zeros` is a placeholder for initialization. The logic below will
    # correctly set the values for the first interaction.
    q_values = jnp.zeros(3)
    novelty_bonus_state = jnp.zeros(3)
  else:
    q_values, novelty_bonus_state = agent_state

  # --- 2. Update Q-Values ---

  # Create a one-hot mask to identify the newly available option.
  novel_option_mask = jax.nn.one_hot(novel_option, num_classes=3)

  # --- Q-value update for existing (non-novel) options ---
  
  # a. Calculate the standard temporal difference (TD) update for the chosen action.
  choice_mask = jax.nn.one_hot(choice, num_classes=3)
  td_error = reward - q_values
  td_update = q_values + learning_rate * td_error * choice_mask

  # b. Apply a decay to all existing Q-values, pulling them towards the initial value.
  # This acts as a form of forgetting or regularization.
  q_values_for_existing_options = (
      td_update * (1.0 - learning_rate) + initial_q_value * learning_rate
  )
  
  # --- Combine updates for novel and existing options ---
  # If an option is novel, reset its Q-value to the initial value.
  # Otherwise, apply the TD and decay update calculated above.
  updated_q_values = jnp.where(
      novel_option_mask,
      initial_q_value,
      q_values_for_existing_options
  )

  # --- 3. Update Novelty Bonus ---

  # Decay the novelty bonus for all existing options.
  decayed_novelty_bonus = novelty_bonus_state * (1.0 - novelty_bonus_decay)
  
  # Reset the bonus for the novel option to its maximum magnitude.
  updated_novelty_bonus_state = jnp.where(
      novel_option_mask,
      novelty_bonus_magnitude,
      decayed_novelty_bonus
  )

  # --- 4. Prepare for Next Action Selection ---

  # a. Combine Q-values and the novelty bonus to get the final values for decision-making.
  # This encourages exploration of options with a high novelty bonus.
  effective_values = updated_q_values + updated_novelty_bonus_state

  # b. Calculate a dynamic inverse temperature (softmax beta).
  # The temperature is increased when Q-value variance is high, promoting more
  # exploration when the agent is uncertain.
  q_value_variance = jnp.var(updated_q_values)
  dynamic_inverse_temperature = (
      base_inverse_temperature +
      inverse_temperature_sensitivity_to_variance * q_value_variance
  )

  # c. Calculate the final choice logits for the softmax policy.
  choice_logits = dynamic_inverse_temperature * effective_values

  # --- 5. Package and Return State ---

  # Stack the updated arrays to form the new agent state.
  updated_agent_state = jnp.stack([updated_q_values, updated_novelty_bonus_state])

  return choice_logits, updated_agent_state


# Shortened to 100k steps then refactored, run 1, medium floor.
def monkey_bandit_run1_simplified_medium_floor(
    params: chex.Array,
    choice: int,
    reward: float,
    novel_option: int,
    agent_state: Optional[chex.Array],
) -> tuple[chex.Array, chex.Array]:
  """Cognitive model describing monkey behavior on a multi-armed bandit task.

  Assumes the monkey is presented with three options on each trial.
  Occasionally, one of these options is changed to a novel arm with different
  reward probabilities, when this happens novel_option will indicate the index
  of the newly novel option and otherwise it will be -1.

  Args:
    params: Model params. Different parameters are used for different monkeys.
    choice: The choice made by the subject on the previous trial. An
      integer with values of 0, 1, or 2.
    reward: The reward received by the subject on this trial. A scalar
      either 0 or 1.
    novel_option: The choice option that is novel to the subject on this trial.
      The number 0, 1, 2 indicates one of the choices, and -1 indicates that
      no option is currently novel.
    agent_state: The current state of the cognitive model.

  Returns:
    choice_logits: The probabilities that the subject will choose option
      0, 1, or 2 on the next trial, expressed as logits.
    agent_state: The updated state of the cognitive model.
  """
  # --- 1. Initialization and Parameter Unpacking ---

  # On the first trial, initialize the agent's state with zeros.
  # The state vector holds Q-values and novelty values for the 3 options.
  if agent_state is None:
    # State is a vector of size 6: [q0, q1, q2, n0, n1, n2]
    agent_state = jnp.zeros((6,))

  # Unpack the model's parameters, which are learned for each subject.
  (
      # Learning rate for the value of the chosen option.
      base_learning_rate_chosen,
      # Base inverse temperature for the softmax choice rule.
      beta_base,
      # Initial Q-value assigned to all options at the start or when novel.
      initial_q,
      # Learning rate for the value of the unchosen options.
      base_learning_rate_unchosen,
      # Initial bonus value assigned to a novel option.
      novelty_initial_bonus,
      # Rate at which the novelty bonus decays over time.
      novelty_decay_rate,
      # Factor scaling how much novelty affects the learning rate.
      novelty_learning_rate_scale,
      # Factor scaling how much Q-value variance affects beta.
      beta_q_variance_scale,
      *_  # stachenfeld@ added for unused parameters.
  ) = params

  # --- 2. State Unpacking and Novelty Handling ---

  # Split the agent state from the previous trial into Q-values and novelty values.
  q_values_prev, novelty_values_prev = jnp.split(agent_state, 2)

  # Create a one-hot encoded mask to identify which option is novel (-1 means none).
  is_novel_option = jax.nn.one_hot(novel_option, num_classes=3)

  # If an option is novel, reset its Q-value to the initial default value.
  # Otherwise, keep the Q-value from the previous state.
  q_values_after_novelty_reset = jnp.where(is_novel_option, initial_q, q_values_prev)

  # If an option is novel, reset its novelty bonus to the initial high value.
  novelty_values_after_reset = jnp.where(
      is_novel_option, novelty_initial_bonus, novelty_values_prev
  )
  # Apply a decay to all novelty values, so they decrease over time.
  novelty_values = novelty_values_after_reset * (1 - novelty_decay_rate)

  # --- 3. Q-Value Update (Reinforcement Learning) ---

  # Create a one-hot encoded mask to identify the action chosen on the last trial.
  is_chosen = jax.nn.one_hot(choice, num_classes=3)

  # Calculate the prediction error (difference between expected and actual reward).
  # For the chosen option, error = reward - Q-value.
  # For unchosen options, they are assumed to regress towards the initial value.
  prediction_error = jnp.where(
      is_chosen,
      reward - q_values_after_novelty_reset,
      initial_q - q_values_after_novelty_reset,
  )

  # Determine the base learning rate for each option.
  # Use a higher rate for the chosen option and a lower one for unchosen options.
  # The sigmoid function transforms the raw parameters into a (0, 1) range.
  base_learning_rates = jnp.where(
      is_chosen,
      jax.nn.sigmoid(base_learning_rate_chosen),
      jax.nn.sigmoid(base_learning_rate_unchosen),
  )

  # Increase the learning rate for options that are still considered novel.
  effective_learning_rates = (
      base_learning_rates + novelty_learning_rate_scale * novelty_values
  )

  # Update the Q-values using the prediction error and the learning rates.
  # This is the core Rescorla-Wagner learning rule.
  q_values_updated = (
      q_values_after_novelty_reset
      + effective_learning_rates * prediction_error
  )

  # --- 4. Prepare for Next Choice ---

  # Calculate the softmax inverse temperature (beta).
  # Beta is dynamic: it increases when the Q-values are more spread out (higher
  # variance), leading to more deterministic (exploitative) choices.
  beta = beta_base + beta_q_variance_scale * jnp.var(q_values_updated)

  # Combine the learned Q-values and the current novelty values to get the
  # total value, or "attractiveness," of each option.
  total_value = q_values_updated + novelty_values

  # Calculate the choice logits for the next trial. These are the inputs to a
  # softmax function that determines the choice probabilities.
  # A higher logit means a higher probability of being chosen.
  choice_logits = beta * total_value

  # --- 5. Update and Return State ---

  # Concatenate the updated Q-values and novelty values to form the new agent state.
  agent_state_updated = jnp.concatenate((q_values_updated, novelty_values))

  # Return the choice logits and the new state for the next trial.
  return choice_logits, agent_state_updated


# Shortened to 100k steps then refactored, run 2, medium floor.
def monkey_bandit_run2_simplified_medium_floor(
    params: chex.Array,
    choice: int,
    reward: float,
    novel_option: int,
    agent_state: Optional[chex.Array],
) -> tuple[chex.Array, chex.Array]:
  """Cognitive model describing monkey behavior on a multi-armed bandit task.

  Occasionally, one of these options is changed to a novel arm with different
  reward probabilities, when this happens novel_option will indicate the index
  of the newly novel option and otherwise it will be -1.

  Args:
    params: Model params. Different parameters are used for different monkeys.
    choice: The choice made by the subject on the previous trial. An
      integer with values of 0, 1, or 2.
    reward: The reward received by the subject on this trial. A scalar
      either 0 or 1.
    novel_option: The choice option that is novel to the subject on this trial.
      The number 0, 1, 2 indicates one of the choices, and -1 indicates that
      no option is currently novel.
    agent_state: The current state of the cognitive model.

  Returns:
    choice_logits: The probabilities that the subject will choose option
      0, 1, or 2 on the next trial, expressed as logits.
    agent_state: The updated state of the cognitive model.
  """
  # --- 1. Initialization and Parameter Unpacking ---

  # On the first trial, initialize the agent's state to a vector of six zeros.
  # The state vector is structured as:
  # [Q-value_option_0, Q-value_option_1, Q-value_option_2,
  #  Novelty_bonus_0,  Novelty_bonus_1,  Novelty_bonus_2]
  if agent_state is None:
    agent_state = jnp.zeros((6,))

  # Unpack the raw model parameters. Using softplus ensures some parameters are positive.
  learning_rate_raw = params[0]
  inverse_temperature_raw = params[1]
  novelty_bonus_initial_raw = params[2]
  decay_rate_raw = params[3]
  variance_influence_raw = params[5]

  # Apply the softplus transformation to ensure positivity for rates and temperatures.
  learning_rate = jax.nn.softplus(learning_rate_raw)
  inverse_temperature = jax.nn.softplus(inverse_temperature_raw)
  novelty_bonus_initial = jax.nn.softplus(novelty_bonus_initial_raw)
  decay_rate = jax.nn.softplus(decay_rate_raw)
  
  # These parameters are used directly without transformation.
  baseline_q_value = params[4]
  variance_influence = params[5] # Note: used raw, not softplus-transformed
  novel_q_initial_bias = params[9]

  # --- 2. Q-Value Update (Learning and Forgetting) ---

  # First, apply a decay rule to all Q-values, pulling them toward a baseline.
  # This models a "forgetting" or reversion-to-the-mean process for all options.
  # Note: This decay is applied to the full agent_state, but only affects the
  # Q-values (first 3 elements) because the novelty bonuses are updated later.
  state_after_decay = agent_state * (1 - decay_rate) + baseline_q_value * decay_rate
  
  # Calculate the prediction error: the difference between actual reward and expected reward (Q-value).
  # This update is based on the Q-value *before* the decay was applied.
  prediction_error = reward - agent_state[choice]
  
  # Update the Q-value for the chosen option using a standard Rescorla-Wagner learning rule.
  learned_q_value_for_choice = agent_state[choice] + learning_rate * prediction_error
  
  # Combine the two steps: the state now contains decayed values for unchosen options
  # and the newly learned value for the chosen option.
  state_after_q_update = state_after_decay.at[choice].set(learned_q_value_for_choice)

  # --- 3. Handle Introduction of a Novel Option ---

  # If a novel option was introduced on this trial (`novel_option` is 0, 1, or 2)...
  # ...reset its Q-value to an initial bias and set its novelty bonus to a high initial value.
  # This is done using jnp.where for conditional, parallelizable execution.
  state_if_novel = state_after_q_update.at[novel_option].set(novel_q_initial_bias)
  state_if_novel = state_if_novel.at[novel_option + 3].set(novelty_bonus_initial)

  # If no option was novel (`novel_option` is -1), the state remains unchanged.
  state_after_novelty_reset = jnp.where(
      novel_option >= 0,
      state_if_novel,
      state_after_q_update
  )

  # --- 4. Novelty Bonus Update ---

  # Implement a "use-it-or-lose-it" decay for novelty bonuses.
  # First, get the current novelty bonuses from the state.
  current_novelty_bonuses = state_after_novelty_reset[3:6]
  
  # Decay all novelty bonuses by a fixed factor (0.9).
  decayed_novelty_bonuses = current_novelty_bonuses * 0.9

  # Overwrite the decayed bonus for the *chosen* option with its original,
  # undecayed value. This means the chosen option's novelty is "consumed"
  # without decay, while unchosen options' novelty fades.
  bonuses_after_selective_decay = decayed_novelty_bonuses.at[choice].set(current_novelty_bonuses[choice])

  # Update the main state vector with the new novelty bonuses.
  final_agent_state = state_after_novelty_reset.at[3:6].set(bonuses_after_selective_decay)

  # --- 5. Calculate Choice Logits for the Next Trial ---

  # Extract the final Q-values and novelty bonuses for the decision calculation.
  final_q_values = final_agent_state[0:3]
  final_novelty_bonuses = final_agent_state[3:6]
  
  # The total value of each option is its learned Q-value plus its current novelty bonus.
  total_option_values = final_q_values + final_novelty_bonuses
  
  # Calculate the variance across the Q-values. This is used to modulate exploration.
  # Higher variance (more certainty about which option is best) leads to more exploitation.
  q_value_variance = jnp.var(final_q_values)
  
  # Create a scaling factor based on Q-value variance.
  variance_scaler = 1 + variance_influence * q_value_variance
  
  # Calculate the choice logits. These are the inputs to a softmax function
  # that determines the probability of choosing each option on the next trial.
  choice_logits = total_option_values * inverse_temperature * variance_scaler

  return choice_logits, final_agent_state


# Shortened to 100k steps then refactored, run 3, medium floor.
def monkey_bandit_run3_simplified_medium_floor(
    params: chex.Array,
    choice: int,
    reward: float,
    novel_option: int,
    agent_state: Optional[chex.Array],
) -> tuple[chex.Array, chex.Array]:
  """Cognitive model describing monkey behavior on a multi-armed bandit task."""

  # --- 1. Unpack model parameters ---
  # These parameters define the agent's learning and decision-making rules.
  initial_q_value = params[0]
  learning_rate = params[1]
  base_inverse_temperature = params[2]
  inverse_temperature_sensitivity_to_q_variance = params[3]
  unchosen_decay_rate = params[4]
  novelty_bonus_magnitude = params[5]
  novelty_bonus_decay = params[6]
  novelty_learning_rate_impact_on_learning_rate = params[7]

  # --- 2. Initialize or retrieve agent's internal state ---
  # The agent's state consists of the Q-values (expected reward) and the
  # current novelty bonus for each of the 3 options.
  if agent_state is None:
    # If this is the first trial, initialize Q-values and novelty bonuses.
    q_values = jnp.full((3,), initial_q_value)
    novelty_bonuses = jnp.full((3,), 0.0)
  else:
    # Otherwise, unpack the state from the previous trial.
    q_values, novelty_bonuses = agent_state

  # --- 3. Update Q-values based on the chosen action and reward ---

  # Calculate the prediction error: the difference between the received reward
  # and the expected value (Q-value) of the chosen option.
  prediction_error = reward - q_values[choice]

  # The learning rate is modulated by the novelty of the chosen option.
  # A higher novelty bonus for the chosen option can increase the learning rate for it.
  novelty_modulated_learning_rate = learning_rate * (
      1 + novelty_learning_rate_impact_on_learning_rate * novelty_bonuses[choice]
  )

  # Calculate the new Q-value for the chosen option using the prediction error
  # and the modulated learning rate.
  updated_q_value_for_chosen_option = q_values[choice] + novelty_modulated_learning_rate * prediction_error

  # The Q-values of unchosen options decay towards the initial Q-value.
  # This represents forgetting or a belief that the environment is changing.
  decayed_q_values = q_values * (1 - unchosen_decay_rate) + initial_q_value * unchosen_decay_rate

  # Combine the updates: use the decayed values for all options, then
  # overwrite the value for the option that was just chosen.
  q_values_after_learning = decayed_q_values.at[choice].set(updated_q_value_for_chosen_option)

  # Create a mask to identify which option, if any, was replaced with a novel one.
  # `novel_option` is an integer (0, 1, or 2) indicating the novel slot, or -1 if none.
  novel_mask = jax.nn.one_hot(novel_option, num_classes=3)

  # If an option was replaced, its Q-value is reset to the initial value.
  # Otherwise, we keep the value that was just updated through learning.
  final_q_values = jnp.where(
      novel_mask,
      initial_q_value,              # Value if the option is novel
      q_values_after_learning,      # Value if the option is not novel
  )

  # --- 4. Update the novelty bonus for each option ---

  # The bonus for any newly introduced option is reset to its maximum magnitude.
  # The bonuses for all other, non-novel options decay exponentially.
  decayed_novelty_bonuses = novelty_bonuses * (1. - novelty_bonus_decay)
  updated_novelty_bonuses = jnp.where(
      novel_mask,
      novelty_bonus_magnitude,  # Value if the option is novel
      decayed_novelty_bonuses,  # Value if the option is not novel
  )

  # --- 5. Calculate action probabilities for the next decision ---

  # The "inverse temperature" controls the exploration/exploitation trade-off.
  # A higher value leads to more deterministic (exploitative) choices.
  # It is modulated by the variance of the Q-values, increasing exploration
  # when the agent is more uncertain about the option values.
  q_value_variance = jnp.var(final_q_values)
  inverse_temperature = base_inverse_temperature + inverse_temperature_sensitivity_to_q_variance * q_value_variance

  # The agent's preference for each option (logits) is the sum of its
  # learned value (Q-value) and its transient novelty bonus. These logits
  # are the inputs to a softmax function to determine choice probabilities.
  effective_values = final_q_values + updated_novelty_bonuses
  choice_logits = inverse_temperature * effective_values

  # --- 6. Package the updated state for the next trial ---
  new_agent_state = jnp.stack((final_q_values, updated_novelty_bonuses))

  # Return the choice logits and the new agent state.
  return choice_logits, new_agent_state


# Shortened to 100k steps then refactored, run 1, high floor.
def monkey_bandit_run1_simplified_high_floor(
    params: chex.Array,
    choice: int,
    reward: float,
    novel_option: int,
    agent_state: Optional[chex.Array],
) -> tuple[chex.Array, chex.Array]:
  """Cognitive model describing monkey behavior on a multi-armed bandit task.

  Assumes the monkey is presented with three options on each trial.
  Occasionally, one of these options is changed to a novel arm with different
  reward probabilities, when this happens novel_option will indicate the index
  of the newly novel option and otherwise it will be -1.

  Args:
    params: Model params. Different parameters are used for different monkeys.
    choice: The choice made by the subject on the previous trial. An
      integer with values of 0, 1, or 2.
    reward: The reward received by the subject on this trial. A scalar
      either 0 or 1.
    novel_option: The choice option that is novel to the subject on this trial.
      The number 0, 1, 2 indicates one of the choices, and -1 indicates that
      no option is currently novel.
    agent_state: The current state of the cognitive model.

  Returns:
    choice_logits: The probabilities that the subject will choose option
      0, 1, or 2 on the next trial, expressed as logits.
    agent_state: The updated state of the cognitive model.
  """
  # --- 1. Unpack or Initialize Agent State ---
  # The agent's state consists of Q-values (expected reward for each option)
  # and novelty values (a bonus for choosing new options).
  # If this is the first trial (agent_state is None), they are initialized to zeros.
  if agent_state is not None:
    # On subsequent trials, unpack the state from the previous step.
    # The state is a single array of size 6.
    q_values, novelty_values = jnp.split(agent_state, 2)
  else:
    # On the first trial, initialize state to all zeros.
    initial_state = jnp.zeros(6)
    q_values, novelty_values = jnp.split(initial_state, 2)

  # --- 2. Unpack and Transform Model Parameters ---
  # Parameters are passed in a raw format and need to be transformed by
  # activation functions (e.g., sigmoid, softplus) to ensure they are in a
  # valid range (e.g., learning rates between 0 and 1).
  (p_base_learning_rate_chosen, p_beta_base, initial_q_value,
   p_base_learning_rate_unchosen, p_fictive_reward_context_scale,
   p_novelty_initial_bonus, p_novelty_decay_rate, novelty_q_learning_rate_scale,
   p_beta_avg_q_scale, p_beta_spread_scale) = params

  # Apply sigmoid to constrain learning rates and scales between 0 and 1.
  base_learning_rate_chosen = jax.nn.sigmoid(p_base_learning_rate_chosen)
  base_learning_rate_unchosen = jax.nn.sigmoid(p_base_learning_rate_unchosen)
  fictive_reward_context_scale = jax.nn.sigmoid(p_fictive_reward_context_scale)
  novelty_decay_rate = jax.nn.sigmoid(p_novelty_decay_rate)
  beta_avg_q_scale = jax.nn.sigmoid(p_beta_avg_q_scale)
  beta_spread_scale = jax.nn.sigmoid(p_beta_spread_scale)

  # Apply softplus to ensure beta and novelty bonus are non-negative.
  beta_base = jax.nn.softplus(p_beta_base)
  novelty_initial_bonus = jax.nn.softplus(p_novelty_initial_bonus)

  # --- 3. Handle the Introduction of a Novel Option ---
  # If an option is novel on this trial, its Q-value and novelty bonus are reset.
  is_novel_mask = (novel_option == jnp.arange(3))

  # Calculate the Q-value to assign to the new option. This is a weighted
  # average of a default initial value and the mean of the Q-values of the
  # other, non-novel options.
  mean_of_other_q_values = jnp.mean(q_values * (1 - is_novel_mask)) * 1.5
  new_q_for_novel_option = ((1.0 - beta_avg_q_scale) * initial_q_value +
                            beta_avg_q_scale * mean_of_other_q_values)

  # Update the Q-values array: replace the value for the novel option.
  q_values = jnp.where(is_novel_mask, new_q_for_novel_option, q_values)

  # Update the novelty values array: reset the bonus for the novel option.
  novelty_values = jnp.where(is_novel_mask, novelty_initial_bonus, novelty_values)

  # --- 4. Decay All Novelty Bonuses ---
  # The novelty bonus for all options decays over time, making them less attractive.
  novelty_decay_rate_multiplier = 1.0 - novelty_decay_rate
  novelty_values = novelty_values * novelty_decay_rate_multiplier

  # --- 5. Calculate Prediction Errors for Learning ---
  # The prediction error is the difference between the actual outcome and the
  # expected outcome (Q-value). This error signal drives learning.
  chosen_mask = (jnp.arange(3) == choice)

  # For unchosen options, a "fictive reward" is calculated. This allows the
  # agent to learn from the outcomes of options it didn't pick. The target
  # for unchosen options is a mix of the actual reward and a default value.
  fictive_reward_term_for_unchosen = (
      (initial_q_value - reward) * fictive_reward_context_scale)

  # The prediction error for the chosen option is simply (reward - Q_value).
  # For unchosen options, the fictive reward term is added to this difference.
  reward_minus_q = reward - q_values
  prediction_errors = reward_minus_q + jnp.where(
      chosen_mask, 0.0, fictive_reward_term_for_unchosen)

  # --- 6. Calculate Modulated Learning Rates ---
  # The learning rate determines how much the Q-values are updated.
  # It differs for chosen vs. unchosen options and is modulated by novelty.
  base_learning_rates = jnp.where(chosen_mask,
                                  base_learning_rate_chosen,
                                  base_learning_rate_unchosen)

  # The learning rate is increased for options with a high novelty bonus,
  # scaled by the magnitude of the prediction error. This is a form of
  # uncertainty-driven learning.
  novelty_learning_rate_boost = (novelty_q_learning_rate_scale *
                                 novelty_values *
                                 jnp.abs(prediction_errors))

  # The final learning rate is the sum of the base rate and the novelty boost.
  effective_learning_rates = base_learning_rates + novelty_learning_rate_boost

  # --- 7. Update Q-Values ---
  # Update the Q-values using the calculated learning rates and prediction errors.
  # This is the core reinforcement learning step.
  q_values_update = effective_learning_rates * prediction_errors
  q_values = q_values + q_values_update

  # --- 8. Calculate Action-Selection Policy (Logits) ---
  # The agent's choice on the next trial is determined by the Q-values and
  # novelty bonuses, passed through a softmax function. We calculate the
  # inputs to the softmax, which are called logits.

  # The "decision value" for each option is its Q-value plus its current novelty bonus.
  decision_values = q_values + novelty_values

  # The 'beta' parameter (inverse temperature) controls the exploration-exploitation
  # trade-off. Here, it is dynamically adjusted based on the current Q-values.
  # A higher beta means more exploitation (less random choices).
  average_absolute_q = jnp.mean(jnp.abs(q_values))
  q_value_range = jnp.ptp(q_values)  # Peak-to-peak (max - min)

  # Beta is a sum of a base value and terms scaled by the average Q and the range of Qs.
  dynamic_beta = (beta_base +
                  beta_avg_q_scale * average_absolute_q +
                  beta_spread_scale * q_value_range)

  # The final choice logits are the decision values scaled by the dynamic beta.
  choice_logits = dynamic_beta * decision_values

  # --- 9. Package State for Next Trial ---
  # Combine the updated Q-values and novelty values into a single array
  # to be passed as the agent_state in the next iteration.
  agent_state = jnp.concatenate((q_values, novelty_values))

  return choice_logits, agent_state


# Shortened to 100k steps then refactored, run 2, high floor.
def monkey_bandit_run2_simplified_high_floor(
    params: chex.Array,
    choice: int,
    reward: float,
    novel_option: int,
    agent_state: Optional[chex.Array],
) -> tuple[chex.Array, chex.Array]:
  """Cognitive model describing monkey behavior on a multi-armed bandit task.

  Assumes the monkey is presented with three options on each trial.
  Occasionally, one of these options is changed to a novel arm with different
  reward probabilities, when this happens novel_option will indicate the index
  of the newly novel option and otherwise it will be -1.

  Args:
    params: Model params. Different parameters are used for different monkeys.
    choice: The choice made by the subject on the previous trial. An
      integer with values of 0, 1, or 2.
    reward: The reward received by the subject on this trial. A scalar
      either 0 or 1.
    novel_option: The choice option that is novel to the subject on this trial.
      The number 0, 1, 2 indicates one of the choices, and -1 indicates that
      no option is currently novel.
    agent_state: The current state of the cognitive model.

  Returns:
    choice_logits: The probabilities that the subject will choose option
      0, 1, or 2 on the next trial, expressed as logits.
    agent_state: The updated state of the cognitive model.
  """
  # --- 1. Unpack and Transform Model Parameters ---

  # Unpack the 1D parameter array into named variables for clarity.
  (positive_learning_rate_param, negative_learning_rate_param,
   unchosen_initial_learning_rate_param, log_inverse_temperature,
   prior_q_initial_value, unchosen_q_baseline, q_variance_scaler,
   novelty_bonus_decay_rate_param, novelty_baseline_value,
   novel_q_initial_value) = params

  # Transform raw parameters into their usable form.
  # Learning rates are passed through a sigmoid to constrain them between 0 and 1.
  learning_rate_params = jnp.array([
      positive_learning_rate_param,
      negative_learning_rate_param,
      unchosen_initial_learning_rate_param
  ])
  positive_learning_rate, negative_learning_rate, unchosen_initial_learning_rate = jax.nn.sigmoid(
      learning_rate_params
  )

  # Inverse temperature and decay rate are passed through softplus to ensure they are positive.
  softplus_params = jnp.array([log_inverse_temperature, novelty_bonus_decay_rate_param])
  inverse_temperature, novelty_bonus_decay_rate = jax.nn.softplus(softplus_params)

  # --- 2. Initialize Agent State ---

  # If this is the first trial, initialize the agent's state.
  # The state vector contains Q-values (beliefs about reward) for the 3 options,
  # followed by novelty bonuses for the 3 options.
  if agent_state is None:
    initial_q_values = jnp.array([
        prior_q_initial_value, prior_q_initial_value, prior_q_initial_value
    ])
    initial_novelty_bonuses = jnp.array([0.0, 0.0, 0.0])
    agent_state = jnp.concatenate([initial_q_values, initial_novelty_bonuses])

  # --- 3. Update Q-values based on choice and reward ---

  # Extract current Q-values and novelty bonuses from the agent state.
  current_q_values = agent_state[0:3]
  current_novelty_bonuses = agent_state[3:6]

  # Calculate the prediction error: the difference between the received reward and the expected reward (Q-value).
  prediction_error = reward - current_q_values[choice]

  # The learning rate for the chosen option is dynamic.
  # It depends on whether the outcome was better or worse than expected (prediction_error sign).
  base_learning_rate = jnp.where(prediction_error > 0,
                                 positive_learning_rate,
                                 negative_learning_rate)
  # The learning rate is also scaled by the magnitude of the prediction error and a novelty-based term.
  chosen_option_novelty_bonus = current_novelty_bonuses[choice]
  novelty_modulation_factor = 1.0 + jax.nn.softplus(chosen_option_novelty_bonus)
  chosen_option_learning_rate = base_learning_rate * jnp.abs(prediction_error) * novelty_modulation_factor

  # Update the Q-value for the chosen option using the Rescorla-Wagner rule.
  q_value_update_for_chosen = chosen_option_learning_rate * prediction_error
  updated_q_chosen = current_q_values[choice] + q_value_update_for_chosen

  # Update Q-values for unchosen options. This model assumes they decay towards
  # a baseline and are updated vicariously based on the reward received.
  # NOTE: This update is calculated for and applied to the *entire* state vector,
  # affecting both Q-values and novelty bonuses, before the chosen Q-value is corrected.
  decay_towards_baseline = 0.5 * (unchosen_q_baseline - agent_state)
  vicarious_reward_update = 0.2 * reward
  general_state_update = unchosen_initial_learning_rate * (decay_towards_baseline + vicarious_reward_update)
  
  # Apply the general update and then set the specific, corrected value for the chosen option's Q-value.
  state_after_general_update = agent_state + general_state_update
  state_after_q_update = state_after_general_update.at[choice].set(updated_q_chosen)

  # --- 4. Update Novelty Bonuses ---

  # Novelty bonuses for unchosen options decay towards a baseline.
  # The chosen option's novelty bonus is not updated in this step.
  current_novelty_bonuses_after_q = state_after_q_update[3:6]
  decayed_bonuses = (current_novelty_bonuses_after_q * (1 - novelty_bonus_decay_rate)
                     + novelty_baseline_value * novelty_bonus_decay_rate)

  # Restore the chosen option's novelty bonus to its value before the decay step.
  chosen_option_bonus_before_decay = current_novelty_bonuses_after_q[choice]
  updated_novelty_bonuses = decayed_bonuses.at[choice].set(chosen_option_bonus_before_decay)

  # Place the updated novelty bonuses back into the agent state.
  state_after_novelty_update = state_after_q_update.at[3:6].set(updated_novelty_bonuses)

  # --- 5. Handle Introduction of a Novel Option ---

  # If a new option was introduced on this trial (`novel_option` >= 0),
  # reset its Q-value and novelty bonus to their initial values.
  is_novel_option_present = (novel_option >= 0)
  state_if_novel = state_after_novelty_update.at[novel_option].set(novel_q_initial_value)
  state_if_novel = state_if_novel.at[novel_option + 3].set(0.0)
  
  # Use jnp.where to conditionally apply the reset. If no novel option, state remains unchanged.
  updated_agent_state = jnp.where(is_novel_option_present, state_if_novel, state_after_novelty_update)

  # --- 6. Calculate Choice Logits for the Next Trial ---

  # Final Q-values and novelty bonuses from the updated state.
  final_q_values = updated_agent_state[0:3]
  final_novelty_bonuses = updated_agent_state[3:6]

  # The effective value of each option is its Q-value plus its novelty bonus.
  effective_q_values = final_q_values + final_novelty_bonuses

  # The inverse temperature (choice stochasticity) is modulated by the variance of Q-values.
  # Higher variance leads to more deterministic choices (higher inverse temperature).
  q_variance = jnp.var(final_q_values)
  variance_scaling_factor = 1.0 + q_variance_scaler * (q_variance + 1e-6) # 1e-6 for stability
  base_inverse_temperature = inverse_temperature * variance_scaling_factor

  # Further modulate the inverse temperature for the novel option, making the
  # agent initially less sensitive to its Q-value.
  is_option_novel_array = (jnp.arange(3) == novel_option)
  novelty_temperature_reduction = 1.0 - novelty_bonus_decay_rate
  modulated_inverse_temperature = jnp.where(
      is_option_novel_array,
      base_inverse_temperature * novelty_temperature_reduction,
      base_inverse_temperature
  )

  # Calculate choice logits using a softmax rule: logits are effective values scaled by temperature.
  choice_logits = effective_q_values * modulated_inverse_temperature

  return choice_logits, updated_agent_state


# Shortened to 100k steps then refactored, run 3, high floor.
def monkey_bandit_run3_simplified_high_floor(
    params: chex.Array,
    choice: int,
    reward: float,
    novel_option: int,
    agent_state: Optional[chex.Array],
) -> tuple[chex.Array, chex.Array]:
  """Cognitive model describing monkey behavior on a multi-armed bandit task.

  Assumes the monkey is presented with three options on each trial.
  Occasionally, one of these options is changed to a novel arm with different
  reward probabilities, when this happens novel_option will indicate the index
  of the newly novel option and otherwise it will be -1.

  Args:
    params: Model params. Different parameters are used for different monkeys.
    choice: The choice made by the subject on the previous trial. An
      integer with values of 0, 1, or 2.
    reward: The reward received by the subject on this trial. A scalar
      either 0 or 1.
    novel_option: The choice option that is novel to the subject on this trial.
      The number 0, 1, 2 indicates one of the choices, and -1 indicates that
      no option is currently novel.
    agent_state: The current state of the cognitive model.

  Returns:
    choice_logits: The probabilities that the subject will choose option
      0, 1, or 2 on the next trial, expressed as logits.
    agent_state: The updated state of the cognitive model.
  """
  # === 1. Unpack and transform model parameters ===
  # Model parameters are passed in a single array. Here we assign them to
  # named variables for clarity.

  # The initial value for all Q-values. This is the assumed value before any
  # learning has occurred. It is also used as a baseline to decay unchosen
  # option values towards.
  initial_q_value = params[0]

  # Other parameters are transformed using the softplus function (a smooth
  # version of relu) to ensure they are always positive.
  (
      learning_rate_for_reward_0,
      learning_rate_for_reward_1,
      learning_rate_for_unchosen,
      novelty_bonus_decay,
      prediction_error_history_decay,
      inverse_temperature_sensitivity_to_q_variance,
      base_inverse_temperature_untransformed,
      novelty_bonus_magnitude,
  ) = jax.nn.softplus(params[1:9])


  # === 2. Unpack or initialize the agent's internal state ===
  # The agent's state carries information from one trial to the next.
  # If this is the first trial (`agent_state` is None), we initialize the state.
  if agent_state is None:
    # Q-values: The agent's estimate of the value of each of the 3 options.
    q_values = jnp.full(3, initial_q_value)
    # Novelty bonuses: An extra value added to novel options to encourage
    # exploration. Initialized to zero.
    novelty_bonuses = jnp.zeros(3)
    # Recency-weighted PE: A running average of the absolute prediction error,
    # which influences choice randomness (inverse temperature).
    recency_weighted_prediction_error = 0.0
  else:
    # If not the first trial, unpack the state from the previous trial.
    # The agent_state is a flat array containing all state variables.
    q_values = agent_state[0:3]
    novelty_bonuses = agent_state[3:6]
    recency_weighted_prediction_error = agent_state[6]


  # === 3. Update Q-values based on the last choice and reward ===

  # Calculate the prediction error: the difference between the reward received
  # and the value that was expected for the chosen option.
  prediction_error = reward - q_values[choice]

  # Determine the learning rate for the chosen option. The model uses
  # different learning rates depending on whether a reward was received.
  learning_rate_for_chosen = jnp.where(
      reward == 1, learning_rate_for_reward_1, learning_rate_for_reward_0
  )

  # Calculate the value update for the chosen option based on its learning rate
  # and the prediction error.
  update_for_chosen_option = learning_rate_for_chosen * prediction_error

  # For the options that were *not* chosen, their Q-values decay towards the
  # initial Q-value. This represents a form of forgetting.
  updates_for_unchosen_options = learning_rate_for_unchosen * (
      initial_q_value - q_values
  )

  # Combine the updates: start with the decay for all options, then
  # overwrite the entry for the chosen option with its specific update.
  q_value_updates = updates_for_unchosen_options.at[choice].set(
      update_for_chosen_option
  )

  # The magnitude of the update is scaled by the current novelty bonus.
  # A higher novelty bonus leads to a larger change in the Q-value.
  scaled_q_value_updates = q_value_updates * (1 + novelty_bonuses)

  # Apply the final scaled updates to the current Q-values.
  q_values = q_values + scaled_q_value_updates


  # === 4. Handle novel options, if any ===
  # If one of the options was replaced with a new one, we reset its state.

  # Create a boolean mask to identify which option is novel.
  # `novel_option` is -1 if no option is novel this trial.
  is_novel_mask = jnp.arange(3) == novel_option

  # Reset the Q-value of the novel option to the initial default value.
  q_values = jnp.where(is_novel_mask, initial_q_value, q_values)

  # Update the novelty bonuses for all options.
  # If an option is novel, its bonus is reset to the full magnitude.
  # Otherwise, its existing bonus decays over time.
  decayed_bonuses = novelty_bonuses * (1 - novelty_bonus_decay)
  novelty_bonuses = jnp.where(
      is_novel_mask, novelty_bonus_magnitude, decayed_bonuses
  )


  # === 5. Update the recency-weighted prediction error history ===
  # This is an exponential moving average of the absolute prediction error,
  # tracking recent surprise or environmental volatility.
  previous_error_contribution = recency_weighted_prediction_error * (
      1 - prediction_error_history_decay
  )
  current_error_contribution = (
      jnp.abs(prediction_error) * prediction_error_history_decay
  )
  recency_weighted_prediction_error = (
      previous_error_contribution + current_error_contribution
  )


  # === 6. Pack the updated state variables into a single array for output ===
  # The agent state must be a single array to be handled by JAX's machinery.
  # Structure: [q0, q1, q2, bonus0, bonus1, bonus2, prediction_error_history]
  agent_state = jnp.array([
      *q_values, *novelty_bonuses, recency_weighted_prediction_error
  ])


  # === 7. Calculate choice logits for the next trial ===
  # Logits are the unnormalized log-probabilities for the softmax function.
  # A higher logit for an option means it's more likely to be chosen.

  # The 'inverse temperature' (beta) controls the exploration vs. exploitation
  # tradeoff. It is dynamically adjusted based on Q-value variance and recent
  # prediction errors.
  q_value_variance = jnp.var(q_values)
  variance_term = (
      inverse_temperature_sensitivity_to_q_variance * q_value_variance
  )
  
  # The raw value for the inverse temperature before transformation.
  # Higher variance increases it (more exploitation).
  # Higher recent error decreases it (more exploration).
  untransformed_inverse_temperature = (
      base_inverse_temperature_untransformed
      + variance_term
      - recency_weighted_prediction_error
  )

  # Ensure the inverse temperature is positive using softplus and add a small
  # constant to prevent it from becoming zero.
  inverse_temperature = (
      jax.nn.softplus(untransformed_inverse_temperature) + 0.01
  )

  # The final decision values are a combination of the learned Q-values
  # and the exploration-driving novelty bonuses.
  decision_values = q_values + novelty_bonuses

  # Calculate choice logits by scaling the decision values by the inverse
  # temperature.
  choice_logits = inverse_temperature * decision_values

  return choice_logits, agent_state


# Program constructed by manually combining insights from discovered programs.
def monkey_bandit_synthesis(
    params: chex.Array,
    choice: int,
    reward: float,
    novel_option: int,
    agent_state: chex.Array | None,
) -> tuple[chex.Array, chex.Array]:
  """Handcrafted agent for the Monkey Bandit task based on insights from discovered programs.

  Has the following elements
  * Maintains separate q_values and novelty_trace state variables.
  * Updates q_values with Q-learning on chosen and decay toward baseline on
    unchosen
  * Updates novelty with bonus for novel options, decay on all other options
  * Differential learning on rewards & omissions
  """
  # --- 1. Unpack Parameters and State ---

  # Unpack the agent's fixed parameters.
  baseline_q_value, reward_learning_rate_logit, omission_learning_rate_logit, \
  base_inverse_temperature, novelty_decay_rate_logit, novelty_initial_value, \
  q_unchosen_decay_rate_logit, q_var_scalar, *_ = params

  reward_learning_rate = jax.nn.sigmoid(reward_learning_rate_logit)
  omission_learning_rate = jax.nn.sigmoid(omission_learning_rate_logit)
  base_inverse_temperature = jnp.abs(base_inverse_temperature)
  novelty_decay_rate = jax.nn.sigmoid(novelty_decay_rate_logit)
  q_unchosen_decay_rate = jax.nn.sigmoid(q_unchosen_decay_rate_logit)
  q_var_scalar = jnp.abs(q_var_scalar)

  # Initialize the agent's state if this is the first step (state is None).
  # The state vector contains Q-values and novelty bonuses for 3 options.
  if agent_state is None:
    agent_state = jnp.zeros(6)  # 3 Q-values and 3 novelty bonuses

  # Unpack the state into separate arrays for Q-values and novelty bonuses.
  q_values, novelty_trace = jnp.split(agent_state, 2)

  # --- 2. Update Novelty Bonuses ---

  # Create a one-hot mask to identify which option, if any, is newly introduced.
  # A value of -1 for `novel_option` means no new option was introduced,
  # and jax.nn.one_hot will correctly produce a vector of all zeros.
  is_novel_mask = jax.nn.one_hot(novel_option, num_classes=3)

  # For all existing options, their novelty bonuses are decayed.
  decayed_existing_bonuses = novelty_trace * novelty_decay_rate

  # Use the mask to apply the correct update: reset for the novel option, decay for others.
  novelty_bonuses = jnp.where(
      is_novel_mask, novelty_initial_value, decayed_existing_bonuses)

  # --- 3. Update Q-Values ---

  # This update logic has three distinct cases handled by `where` clauses:
  #   a) The option was newly introduced.
  #   b) The option was chosen.
  #   c) The option was available but not chosen.

  # Different learning rate depending on reward.
  learning_rate = jnp.where(
      reward > 0.0, reward_learning_rate, omission_learning_rate)

  # First, calculate the updates for existing options (cases b and c).
  is_chosen_mask = jax.nn.one_hot(choice, num_classes=3)
  
  # For the CHOSEN option (b), apply a standard Q-learning update.
  prediction_error = reward - q_values
  q_learning_update = learning_rate * prediction_error
  
  # For UNCHOSEN options (c), decay their values back towards the baseline.
  decay_update = q_unchosen_decay_rate * (baseline_q_value - q_values)
  
  # Combine updates for chosen and unchosen options.
  updates_for_existing_options = jnp.where(is_chosen_mask, q_learning_update, decay_update)
  updated_q_for_existing_options = q_values + updates_for_existing_options


  # Use the novelty mask to select the final Q-values:
  # - If an option is new, set its Q-value to the baseline.
  # - Otherwise, use the updated value for existing options calculated above.
  q_values = jnp.where(is_novel_mask, baseline_q_value, updated_q_for_existing_options)

  # --- 4. Calculate Outputs and Next State ---

  # The final value of an option is its Q-value plus its novelty bonus.
  total_values = q_values + novelty_bonuses

  # Calculate a dynamic inverse temperature for the softmax calculation.
  # This makes the agent's choices more exploratory when Q-values are similar.
  variance_of_q_values = jnp.var(q_values)
  dynamic_inverse_temperature = base_inverse_temperature + q_var_scalar * variance_of_q_values

  # Action logits are the inputs to a softmax function for choice probability.
  action_logits = total_values * dynamic_inverse_temperature

  # The next agent state is the concatenation of the newly updated values.
  next_agent_state = jnp.concatenate((q_values, novelty_bonuses))

  return action_logits, next_agent_state
```
