## Supplementary material for "AI-Discovered Cognitive Models Reveal Novel Insights into Human and Animal Learning": DataDIVER Discovered Programs: rat_bandit_models.py

```
# Copyright 2026 DeepMind Technologies Limited.
#
# Licensed under the Apache License, Version 2.0 (the "License");
# you may not use this file except in compliance with the License.
# You may obtain a copy of the License at
#
#     http://www.apache.org/licenses/LICENSE-2.0
#
# Unless required by applicable law or agreed to in writing, software
# distributed under the License is distributed on an "AS IS" BASIS,
# WITHOUT WARRANTIES OR CONDITIONS OF ANY KIND, either express or implied.
# See the License for the specific language governing permissions and
# limitations under the License.

"""Saved models for the Nature 2026 submission."""
from typing import Optional, Tuple

import chex
import jax
import jax.numpy as jnp


def rat_bandit_run1_fit_only(
    params: chex.Array,
    choice: int,
    reward: int,
    agent_state: Optional[chex.Array],
) -> tuple[chex.Array, chex.Array]:
  """Cognitive model describing rat behavior on a binary two-armed bandit task.

  Args:
    params: Model params. Different parameters are used for different rats.
    choice: Previous choice. Values: 0 or 1.
    reward: Previous reward. Values: 0 or 1.
    agent_state: Previous state of the agent

  Returns:
    choice_logits: Vector of shape (2,) with the probabilities that the rat will
      choose option 0 or 1 on the next trial, expressed as logits.
    agent_state: New state of the agent, after observing the previous choice and
      reward.
  """
  # Do not remove comments or TODOs from this program.

  # TODO: Initialize agent state. NOLINT
  if agent_state is None:
    # Initialize Q-values for choices 0 and 1.
    # These are initialized to 0.5, then subject to one forgetting step, effectively setting the baseline.
    # The recency trackers are initialized to account for one forgetting step, providing a baseline.
    # The last element stores the previous choice for persistence.
    forgetting_rate = jax.nn.sigmoid(params[5])
    q_initial = 0.5 * (1.0 - forgetting_rate) + 0.1 * forgetting_rate
    recency_decay_rate = jax.nn.sigmoid(params[7]) # Use the actual decay rate from params[7]
    recency_initial = 0.1 * (1.0 - recency_decay_rate)
    agent_state = jnp.array([q_initial, q_initial, 0.0, recency_initial, recency_initial, recency_initial, recency_initial])

  beta = jax.nn.softplus(params[0])
  alpha_chosen = jax.nn.sigmoid(params[1])
  alpha_unchosen = jax.nn.sigmoid(params[2])
  persistence_rewarded = jax.nn.sigmoid(params[3])
  persistence_unrewarded = jax.nn.sigmoid(params[4])
  forgetting_rate = jax.nn.sigmoid(params[5])
  bias_choice_0 = params[6]
  recency_decay_rate = jax.nn.sigmoid(params[7])
  recency_win_weight = params[8]
  recency_loss_weight = params[9]

  # Update Q-values with forgetting (decaying past values)
  agent_state = agent_state.at[:2].set(agent_state[:2] * (1.0 - forgetting_rate))
  prediction_error = reward - agent_state[choice]
  agent_state = agent_state.at[choice].add(alpha_chosen * prediction_error)
  unchosen_choice = 1 - choice
  agent_state = agent_state.at[unchosen_choice].add(alpha_unchosen * -prediction_error)
  # Update recency trackers
  agent_state = agent_state.at[3:7].set(agent_state[3:7] * recency_decay_rate)
  agent_state = agent_state.at[3 + choice].add(reward * (1.0 - recency_decay_rate))
  agent_state = agent_state.at[5 + choice].add((1 - reward) * (1.0 - recency_decay_rate))
  # Store the current choice for use in the next trial's persistence calculation.
  agent_state = agent_state.at[2].set(jnp.asarray(choice, float))

  q_values = agent_state[:2]
  # Calculate recency terms for each choice
  recency_terms = recency_win_weight * agent_state[3:5] - recency_loss_weight * agent_state[5:7]

  # Adjust Q-values with recency terms and bias
  adjusted_q_values = q_values * (1.0 + recency_terms)
  adjusted_q_values = adjusted_q_values.at[0].add(bias_choice_0)

  choice_logits = beta * adjusted_q_values
  previous_choice_int = jnp.asarray(agent_state[2], int)
  
  choice_logits = choice_logits.at[previous_choice_int].add(
      jnp.where(reward == 1, persistence_rewarded, -persistence_unrewarded)
  )
  choice_logits = choice_logits + recency_terms

  return choice_logits, agent_state
  

def rat_bandit_run2_fit_only(
    params: chex.Array,
    choice: int,
    reward: int,
    agent_state: Optional[chex.Array],
) -> tuple[chex.Array, chex.Array]:
  """Cognitive model describing rat behavior on a binary two-armed bandit task.

  Args:
    params: Model params. Different parameters are used for different rats.
    choice: Previous choice. Values: 0 or 1.
    reward: Previous reward. Values: 0 or 1.
    agent_state: Previous state of the agent

  Returns:
    choice_logits: Vector of shape (2,) with the probabilities that the rat will
      choose option 0 or 1 on the next trial, expressed as logits.
    agent_state: New state of the agent, after observing the previous choice and
      reward.
  """
  # Do not remove comments or TODOs from this program.

  # Initialize agent state: 2 Q-values (one for each choice) and a choice trace.
  # agent_state structure: [Q0, Q1, GhostQ0, GhostQ1, ChoiceTrace0, ChoiceTrace1, PredictionErrorTrace0, PredictionErrorTrace1]
  if agent_state is None:
    # Initialize Q-values and GhostQ-values to a fittable parameter, choice traces and prediction error traces to zero.
    agent_state = jnp.array([params[0], params[0], params[0], params[0], 0.0, 0.0, 0.0, 0.0])

  # Define all parameters.
  # Each parameter assignment should be a single line.
  # Change the names of the parameters to be descriptive.
  positive_base_learning_rate = jax.nn.sigmoid(params[1])
  negative_base_learning_rate = jax.nn.sigmoid(params[2])
  unchosen_q_update_learning_rate = jax.nn.sigmoid(params[3])
  chosen_q_ghost_influence_rate = jax.nn.sigmoid(params[4])
  ghost_learning_rate = jax.nn.sigmoid(params[5])
  inverse_temperature_0 = jnp.exp(params[6]) # Inverse temperature for choice 0
  inverse_temperature_1 = jnp.exp(params[7]) # Inverse temperature for choice 1
  trace_learning_rate = jax.nn.sigmoid(params[8])
  choice_bias = params[9]
  stickiness_weight = jax.nn.sigmoid(params[9])
  # TODO: Consider adding a parameter for the base value of unchosen_recency_decay_rate. NOLINT (stachenfeld@ added).
  # TODO: Consider adding a parameter for the prediction error trace modulation factor. NOLINT (stachenfeld@ added).

  # Update agent_state.
  # Each component of the agent state update should be a single line.
  q_values = agent_state[0:2]
  ghost_q_values = agent_state[2:4]
  choice_traces = agent_state[4:6]
  prediction_error_traces = agent_state[6:8]

  prediction_error = reward - q_values[choice]
  # Q-value update for chosen option
  # Asymmetric learning rates are applied based on the sign of the prediction error.
  # The update is also modulated by the prediction error trace for the chosen option, amplifying updates when past PE was high.
  pe_trace_modulation = 1 + jax.nn.softplus(jnp.abs(prediction_error_traces[choice]))
  chosen_lr = jnp.where(
      prediction_error >= 0,
      positive_base_learning_rate,
      negative_base_learning_rate,
  )
  q_values_chosen_update = chosen_lr * prediction_error * pe_trace_modulation
  q_values = q_values.at[choice].set(q_values[choice] + q_values_chosen_update)

  # Ghost Q-value influence on chosen Q-value
  # When prediction error is negative, the ghost Q-value pulls the chosen Q-value towards it.
  # When prediction error is positive, the pull is also scaled by the PE, making it stronger for larger positive surprises.
  scaled_ghost_influence_base = chosen_q_ghost_influence_rate * jax.nn.sigmoid(jnp.abs(prediction_error))
  q_values_chosen_ghost_pull = jnp.where(
      prediction_error < 0,
      scaled_ghost_influence_base * (ghost_q_values[choice] - q_values[choice]),
      scaled_ghost_influence_base * prediction_error * (ghost_q_values[choice] - q_values[choice]),
  )
  q_values = q_values.at[choice].set(q_values[choice] + q_values_chosen_ghost_pull)

  # Q-value update for unchosen option
  # The unchosen Q-value updates towards its ghost value.
  # This update is modulated by reward and confidence.
  # Higher reward increases the learning rate for the unchosen option.
  reward_modulated_unchosen_lr = unchosen_q_update_learning_rate * (1 + reward)

  # Confidence modulation: Higher Q-value difference (more confidence in chosen option)
  # or larger PE for chosen option reduces the learning rate for the unchosen option.
  epsilon = 1e-6 # For numerical stability
  q_value_contrast = jnp.abs(q_values[choice] - q_values[1 - choice]) / (jnp.abs(q_values[choice]) + jnp.abs(q_values[1 - choice]) + epsilon)
  confidence_reduction_factor = jax.nn.sigmoid(q_value_contrast) * jax.nn.sigmoid(jnp.abs(prediction_error))
  confidence_modulated_unchosen_lr = reward_modulated_unchosen_lr * (1 - confidence_reduction_factor)

  # Additionally, if PE is negative, the unchosen Q-value is pulled further away from the chosen Q-value.
  unchosen_negative_pe_boost = jnp.where(
      prediction_error < 0,
      confidence_modulated_unchosen_lr * (q_values[choice] - q_values[1 - choice]),
      0.0,
  )
  q_values_unchosen_update = confidence_modulated_unchosen_lr * (ghost_q_values[1 - choice] - q_values[1 - choice]) + unchosen_negative_pe_boost
  q_values = q_values.at[1 - choice].set(q_values[1 - choice] + q_values_unchosen_update)

  # Ghost Q-value update for unchosen option
  # The learning rate for ghost Q-values is modulated by prediction error and the discrepancy
  # between the chosen Q-value and its ghost, reflecting uncertainty.
  ghost_prediction_error_unchosen = reward - ghost_q_values[1 - choice]
  chosen_q_ghost_discrepancy = jax.nn.softplus(jnp.abs(q_values[choice] - ghost_q_values[choice]))
  ghost_lr_modulation_factor = (1 + jax.nn.sigmoid(jnp.abs(prediction_error))) * (1 + chosen_q_ghost_discrepancy)
  ghost_learning_rate_for_unchosen = ghost_learning_rate * ghost_lr_modulation_factor
  ghost_q_values = ghost_q_values.at[1 - choice].set(
      ghost_q_values[1 - choice] + ghost_learning_rate_for_unchosen * ghost_prediction_error_unchosen
  )

  # Ghost Q-value update for chosen option
  # The target for the chosen ghost Q-value is the reward if positive, otherwise the initial Q-value (params[0]).
  dynamic_prior_target_chosen = jnp.where(reward == 1, reward, params[0])
  ghost_learning_rate_for_chosen = ghost_learning_rate * ghost_lr_modulation_factor
  ghost_q_values_chosen_update = ghost_learning_rate_for_chosen * (dynamic_prior_target_chosen - ghost_q_values[choice])
  ghost_q_values = ghost_q_values.at[choice].set(ghost_q_values[choice] + ghost_q_values_chosen_update)

  # Choice trace update
  # All traces decay, then the chosen trace gets a boost.
  # Decay rates can be modulated.
  # Chosen trace learning rate is modulated by absolute prediction error (surprise).
  chosen_trace_learning_rate = trace_learning_rate * (1 + jax.nn.sigmoid(jnp.abs(prediction_error)))
  choice_traces = choice_traces * (1 - trace_learning_rate)
  choice_traces = choice_traces.at[choice].set(choice_traces[choice] + chosen_trace_learning_rate)

  # Unchosen trace decay is modulated by positive PE and reward for the chosen option.
  positive_pe_boost_for_unchosen_decay = jnp.where(prediction_error > 0, jax.nn.sigmoid(prediction_error), 0.0)
  decay_rate_unchosen = trace_learning_rate * (1 + positive_pe_boost_for_unchosen_decay + reward)
  choice_traces = choice_traces.at[1 - choice].set(choice_traces[1 - choice] * (1 - decay_rate_unchosen))

  # Add a small, decaying influence to the unchosen option's trace,
  # reflecting its continued presence as an alternative, but fading if not chosen.
  unchosen_recency_decay_rate = trace_learning_rate * 1.5
  choice_traces = choice_traces.at[1 - choice].set(
      choice_traces[1 - choice] * (1 - unchosen_recency_decay_rate) + (1 - reward) * 0.1
  )

  # Prediction error trace update
  # Asymmetric decay for PE trace: faster for positive PE, slower for negative PE.
  pe_trace_decay_rate = jnp.where(
      prediction_error >= 0,
      trace_learning_rate * 2.0,
      trace_learning_rate * 0.5,
  )
  prediction_error_traces = prediction_error_traces.at[choice].set(
      prediction_error_traces[choice] * (1 - pe_trace_decay_rate) + prediction_error * trace_learning_rate
  )

  agent_state = jnp.concatenate([q_values, ghost_q_values, choice_traces, prediction_error_traces])

  # Dynamic Inverse Temperature based on Q-value difference and volatility.
  # Small Q-value difference -> high uncertainty -> more exploration (lower inverse temperature).
  # High Q-value volatility (from PE trace) -> high uncertainty -> more exploration.
  jnp.abs(q_values[0] - q_values[1])
  q_value_volatility = jnp.mean(jax.nn.softplus(prediction_error_traces))
  jax.nn.sigmoid(q_value_volatility)

  # Modulate inverse temperature by the average absolute prediction error trace.
  # Higher average PE trace (more recent prediction errors) implies higher uncertainty,
  # leading to a lower effective inverse temperature (more exploration).
  # The modulation factor is designed to be between 0 and 1, where 1 means no modulation (max temp)
  # and lower values mean more exploration (lower temp).
  exploration_modulator = 1 - jax.nn.sigmoid(q_value_volatility) # Sigmoid scales volatility to [0,1], then invert for exploration.

  # Apply dynamic modulation to inverse temperatures.
  dynamic_inverse_temperature_0 = inverse_temperature_0 * exploration_modulator
  dynamic_inverse_temperature_1 = inverse_temperature_1 * exploration_modulator
  choice_logits = jnp.array([dynamic_inverse_temperature_0 * q_values[0], dynamic_inverse_temperature_1 * q_values[1]])

  # Add choice bias and stickiness.
  choice_logits = choice_logits + jnp.array([choice_bias, -choice_bias])
  # Stickiness is stronger with higher rewards, promoting perseveration.
  reward_modulated_stickiness_weight = stickiness_weight * (1 + jax.nn.softplus(reward))
  choice_logits = choice_logits + reward_modulated_stickiness_weight * choice_traces

  return choice_logits, agent_state
  

def rat_bandit_run3_fit_only(
    params: chex.Array,
    choice: int,
    reward: int,
    agent_state: Optional[chex.Array],
) -> tuple[chex.Array, chex.Array]:
  """Cognitive model describing rat behavior on a binary two-armed bandit task.

  Args:
    params: Model params. Different parameters are used for different rats.
    choice: Previous choice. Values: 0 or 1.
    reward: Previous reward. Values: 0 or 1.
    agent_state: Previous state of the agent

  Returns:
    choice_logits: Vector of shape (2,) with the probabilities that the rat will
      choose option 0 or 1 on the next trial, expressed as logits.
    agent_state: New state of the agent, after observing the previous choice and
      reward.
  """
  # Do not remove comments or TODOs from this program.

  # TODO: Initialize agent state. NOLINT
  if agent_state is None:
    # State: [Q_value_0, Q_value_1, eligibility_trace_0, eligibility_trace_1, last_reward_0, last_reward_1, action_bias_0, action_bias_1]
    # Initialize Q-values and last_rewards to the sigmoid of params[9], eligibility traces to 0, and action biases to 0.
    agent_state = jnp.array([jax.nn.sigmoid(params[9]), jax.nn.sigmoid(params[9]), 0.0, 0.0, jax.nn.sigmoid(params[9]), jax.nn.sigmoid(params[9]), 0.0, 0.0])

  # Define all parameters.
  # Each parameter assignment should be a single line.
  # Include comments describing the parameter transformation, if applicable.
  inverse_temperature = jnp.exp(jax.nn.softplus(params[0]))
  learning_rate = jax.nn.sigmoid(params[1])
  negative_pe_scaling_factor = jax.nn.sigmoid(params[2])
  perseveration_strength = jax.nn.softplus(params[3])
  eligibility_decay_rate = jax.nn.sigmoid(params[4])
  global_bias_0 = params[5]
  q_decay_rate = jax.nn.sigmoid(params[6])
  reward_history_learning_rate = jax.nn.sigmoid(params[7])
  action_bias_learning_rate = jax.nn.sigmoid(params[8])
  initial_q_value_param = jax.nn.softplus(params[9])

  q_values = agent_state[:2]
  eligibility_traces = agent_state[2:4]
  last_rewards = agent_state[4:6]
  action_biases = agent_state[6:8]

  prediction_error = reward - q_values[choice]
  # Smoothly blend learning rates based on prediction error, inspired by asymmetric learning.
  # Modulate learning rate by the absolute magnitude of prediction error to capture 'surprise'.
  surprise_factor = jax.nn.sigmoid(jnp.abs(prediction_error) * 5.0) # Scale absolute prediction error for sensitivity
  # Blend between baseline learning rate and a scaled version based on surprise and PE direction
  current_learning_rate = learning_rate * (surprise_factor * (jax.nn.sigmoid(prediction_error * 10.0) + (1 - jax.nn.sigmoid(prediction_error * 10.0)) * negative_pe_scaling_factor) + (1 - surprise_factor))
  q_values = q_values * (1 - q_decay_rate) + last_rewards * q_decay_rate
  q_values = q_values.at[choice].set(q_values[choice] + current_learning_rate * prediction_error)
  # Incorporate an exploration bonus: unchosen options get a small, persistent eligibility boost.
  # This encourages exploration of options that haven't been recently chosen, mimicking stochastic exploration.
  exploration_bonus = jnp.array([1.0, 1.0]) - jax.nn.one_hot(choice, num_classes=2)
  eligibility_traces = eligibility_traces * eligibility_decay_rate + (1 - eligibility_decay_rate) * (jax.nn.one_hot(choice, num_classes=2) * (1 + initial_q_value_param * jnp.abs(prediction_error)) + exploration_bonus * 0.1)
  action_biases = action_biases * (1 - action_bias_learning_rate) + jax.nn.one_hot(choice, num_classes=2) * action_bias_learning_rate * (0.5 + reward)
  last_rewards = last_rewards.at[choice].set(last_rewards[choice] + reward_history_learning_rate * prediction_error)
  agent_state = jnp.concatenate([q_values, eligibility_traces, last_rewards, action_biases])

  choice_logits = inverse_temperature * (q_values - last_rewards) + perseveration_strength * (eligibility_traces + action_biases)
  choice_logits = choice_logits.at[0].add(global_bias_0)

  return choice_logits, agent_state
#################################
## HUMAN-CONSTRUCTED PROGRAMS  ##
## Made by kevinjmiller, with  ##
## inspiration from DataDIVER  ##
## Programs                    ##
#################################


def synthesis_base_program(
    params: chex.Array,
    choice: int,
    reward: int,
    agent_state: Optional[chex.Array],
) -> tuple[chex.Array, chex.Array]:
  """Modified rat_bandit_204369309_refactored, added parameter constraints.

  rat_bandit_204369309_refactored is the simplest of the medium-floor shorten
  programs and is quite readable. We'll use it as our base program, and add
  further simplifications. First we'll add parameter constraints to make it
  easier to fit. These exclude bad regions of the parameter space.
  """
  if agent_state is None:
    agent_state = jnp.array([0.5, 0.5, 0.0, 0.0])

  (
    beta,
    persistence_rewarded,
    persistence_unrewarded,
    recency_win_weight,
    recency_loss_weight,
  ) = map(jnp.abs, params[0:5])
  (
      forgetting_rate,       # Rate at which Q-values decay over time.
      recency_decay_rate,    # Rate at which recency effects decay over time.
      alpha_chosen,          # Learning rate for the chosen option.
      alpha_unchosen_fraction, # Fraction of alpha_chosen for unchosen option.
  ) = map(jax.nn.sigmoid, params[5:9])
  bias_choice_0 = params[9]
  alpha_unchosen = alpha_chosen * alpha_unchosen_fraction

  previous_q_values = agent_state[:2]
  previous_recency_effects = agent_state[2:4]

  decayed_q_values = previous_q_values * (1.0 - forgetting_rate)

  prediction_error = reward - decayed_q_values[choice]

  updated_q_values = decayed_q_values.at[choice].add(
      alpha_chosen * prediction_error
  )
  updated_q_values = updated_q_values.at[1 - choice].add(
      alpha_unchosen * -prediction_error
  )

  decayed_recency_effects = previous_recency_effects * recency_decay_rate

  # The recency loss weight param was unconstrained and fitting to negative
  # values. So we constrain it to be positive above, and flip the sign here.
  # recency_update_value = jnp.array([-recency_loss_weight, recency_win_weight])[reward]
  recency_update_value = jnp.array([recency_loss_weight, recency_win_weight])[reward]
  scaled_recency_update = recency_update_value * (1.0 - recency_decay_rate)

  updated_recency_effects = decayed_recency_effects.at[choice].add(
      scaled_recency_update
  )

  # No longer log beta, just constrain it to be positive above.
  # beta = jnp.exp(log_beta)

  value_signal = updated_q_values * jnp.exp(updated_recency_effects)
  value_signal_with_bias = value_signal.at[0].add(bias_choice_0)
  rl_component = beta * value_signal_with_bias

  persistence_bonus = reward * persistence_rewarded - persistence_unrewarded
  recency_and_persistence_component = updated_recency_effects.at[choice].add(
      persistence_bonus
  )

  choice_logits = rl_component + recency_and_persistence_component

  new_agent_state = jnp.concatenate([updated_q_values, updated_recency_effects])

  return choice_logits, new_agent_state


def ablate_persistence(
    params: chex.Array,
    choice: int,
    reward: int,
    agent_state: Optional[chex.Array],
) -> tuple[chex.Array, chex.Array]:
  """Remove persistence_rewarded from the model, fixing it to 1.

  Automatic ablations found that this had little/no effect on quality of fit.
  This program modifies synthesis_base_program. Changes are indicated by
  comments, also the parameter persistence_rewarded is removed.
  """

  if agent_state is None:
    agent_state = jnp.array([0.5, 0.5, 0.0, 0.0])

  (
    beta,
    persistence_unrewarded,
    recency_win_weight,
    recency_loss_weight,
  ) = map(jnp.abs, params[0:4])
  (
      forgetting_rate,       # Rate at which Q-values decay over time.
      recency_decay_rate,    # Rate at which recency effects decay over time.
      alpha_chosen,          # Learning rate for the chosen option.
      alpha_unchosen_fraction, # Fraction of alpha_chosen for unchosen option.
  ) = map(jax.nn.sigmoid, params[4:8])
  bias_choice_0 = params[8]
  alpha_unchosen = alpha_chosen * alpha_unchosen_fraction

  previous_q_values = agent_state[:2]
  previous_recency_effects = agent_state[2:4]

  decayed_q_values = previous_q_values * (1.0 - forgetting_rate)

  prediction_error = reward - decayed_q_values[choice]

  updated_q_values = decayed_q_values.at[choice].add(
      alpha_chosen * prediction_error
  )
  updated_q_values = updated_q_values.at[1 - choice].add(
      alpha_unchosen * -prediction_error
  )

  decayed_recency_effects = previous_recency_effects * recency_decay_rate

  recency_update_value = jnp.array([recency_loss_weight, recency_win_weight])[reward]
  scaled_recency_update = recency_update_value * (1.0 - recency_decay_rate)

  updated_recency_effects = decayed_recency_effects.at[choice].add(
      scaled_recency_update
  )


  value_signal = updated_q_values * jnp.exp(updated_recency_effects)
  value_signal_with_bias = value_signal.at[0].add(bias_choice_0)
  rl_component = beta * value_signal_with_bias

  # Remove persistence_rewarded, fixing it to 1
  # persistence_bonus = reward * persistence_rewarded - persistence_unrewarded
  persistence_bonus = reward - persistence_unrewarded
  recency_and_persistence_component = updated_recency_effects.at[choice].add(
      persistence_bonus
  )


  choice_logits = rl_component + recency_and_persistence_component

  new_agent_state = jnp.concatenate([updated_q_values, updated_recency_effects])

  return choice_logits, new_agent_state


def ablate_forgetting(
    params: chex.Array,
    choice: int,
    reward: int,
    agent_state: Optional[chex.Array],
) -> tuple[chex.Array, chex.Array]:
  """Remove forgetting from the model, fixing it to 0.

  Automatic ablations found that this had little/no effect on quality of fit.
  This program modifies synthesis_base_program. Changes are indicated by comments, also
  the parameter forgetting_rate is removed.
  """
  if agent_state is None:
    agent_state = jnp.array([0.5, 0.5, 0.0, 0.0])

  (
    beta,
    persistence_rewarded,
    persistence_unrewarded,
    recency_win_weight,
    recency_loss_weight,
  ) = map(jnp.abs, params[0:5])
  (
      recency_decay_rate,    # Rate at which recency effects decay over time.
      alpha_chosen,          # Learning rate for the chosen option.
      alpha_unchosen_fraction, # Fraction of alpha_chosen for unchosen option.
  ) = map(jax.nn.sigmoid, params[5:8])
  bias_choice_0 = params[8]
  alpha_unchosen = alpha_chosen * alpha_unchosen_fraction

  previous_q_values = agent_state[:2]
  previous_recency_effects = agent_state[2:4]

  # No forgetting for q-values
  decayed_q_values = previous_q_values # * (1.0 - forgetting_rate)

  prediction_error = reward - decayed_q_values[choice]

  updated_q_values = decayed_q_values.at[choice].add(
      alpha_chosen * prediction_error
  )
  updated_q_values = updated_q_values.at[1 - choice].add(
      alpha_unchosen * -prediction_error
  )

  decayed_recency_effects = previous_recency_effects * recency_decay_rate

  recency_update_value = jnp.array([recency_loss_weight, recency_win_weight])[reward]
  scaled_recency_update = recency_update_value * (1.0 - recency_decay_rate)

  updated_recency_effects = decayed_recency_effects.at[choice].add(
      scaled_recency_update
  )

  value_signal = updated_q_values * jnp.exp(updated_recency_effects)
  value_signal_with_bias = value_signal.at[0].add(bias_choice_0)
  rl_component = beta * value_signal_with_bias

  persistence_bonus = reward * persistence_rewarded - persistence_unrewarded
  recency_and_persistence_component = updated_recency_effects.at[choice].add(
      persistence_bonus
  )


  choice_logits = rl_component + recency_and_persistence_component

  new_agent_state = jnp.concatenate([updated_q_values, updated_recency_effects])

  return choice_logits, new_agent_state


def ablate_beta(
    params: chex.Array,
    choice: int,
    reward: int,
    agent_state: Optional[chex.Array],
) -> tuple[chex.Array, chex.Array]:
  """Remove beta from the model, fixing it to 1.
  Automatic ablations found that this had little/no effect on quality of fit.
  This program modifies base_program
  This program modifies synthesis_base_program. Changes are indicated by
  comments, also the parameter beta is removed.
  """
  if agent_state is None:
    agent_state = jnp.array([0.5, 0.5, 0.0, 0.0])

  (
    persistence_rewarded,
    persistence_unrewarded,
    recency_win_weight,
    recency_loss_weight,
  ) = map(jnp.abs, params[0:4])
  (
      forgetting_rate,       # Rate at which Q-values decay over time.
      recency_decay_rate,    # Rate at which recency effects decay over time.
      alpha_chosen,          # Learning rate for the chosen option.
      alpha_unchosen_fraction, # Fraction of alpha_chosen for unchosen option.
  ) = map(jax.nn.sigmoid, params[4:8])
  bias_choice_0 = params[8]
  alpha_unchosen = alpha_chosen * alpha_unchosen_fraction

  previous_q_values = agent_state[:2]
  previous_recency_effects = agent_state[2:4]

  decayed_q_values = previous_q_values * (1.0 - forgetting_rate)

  prediction_error = reward - decayed_q_values[choice]

  updated_q_values = decayed_q_values.at[choice].add(
      alpha_chosen * prediction_error
  )
  updated_q_values = updated_q_values.at[1 - choice].add(
      alpha_unchosen * -prediction_error
  )

  decayed_recency_effects = previous_recency_effects * recency_decay_rate

  recency_update_value = jnp.array([recency_loss_weight, recency_win_weight])[reward]
  scaled_recency_update = recency_update_value * (1.0 - recency_decay_rate)

  updated_recency_effects = decayed_recency_effects.at[choice].add(
      scaled_recency_update
  )

  value_signal = updated_q_values * jnp.exp(updated_recency_effects)
  value_signal_with_bias = value_signal.at[0].add(bias_choice_0)
  # Remove beta here
  # rl_component = beta * value_signal_with_bias
  rl_component = value_signal_with_bias

  persistence_bonus = reward * persistence_rewarded - persistence_unrewarded
  recency_and_persistence_component = updated_recency_effects.at[choice].add(
      persistence_bonus
  )


  choice_logits = rl_component + recency_and_persistence_component

  new_agent_state = jnp.concatenate([updated_q_values, updated_recency_effects])

  return choice_logits, new_agent_state


def link_forgetting(
    params: chex.Array,
    choice: int,
    reward: int,
    agent_state: Optional[chex.Array],
) -> tuple[chex.Array, chex.Array]:
  """Modifies base_program. Links for forgetting rates across Q-values and recency effects.

  Model comparison found that ablate_forgetting did affect quality of fit.
  Parameter estimation found that forgetting was often similar between the two.
  
  This program modifies synthesis_base_program. Changes are indicated by
  comments, also the parameter forgetting_rate is replaced by a shared
  forgetting_rate for Q-values and recency effects.
  """
  if agent_state is None:
    agent_state = jnp.array([0.5, 0.5, 0.0, 0.0])

  (
    beta,
    persistence_rewarded,
    persistence_unrewarded,
    recency_win_weight,
    recency_loss_weight,
  ) = map(jnp.abs, params[0:5])
  (
      forgetting_rate,    # Rate at which recency effects decay over time.
      alpha_chosen,          # Learning rate for the chosen option.
      alpha_unchosen_fraction, # Fraction of alpha_chosen for unchosen option.
  ) = map(jax.nn.sigmoid, params[5:8])
  bias_choice_0 = params[8]
  alpha_unchosen = alpha_chosen * alpha_unchosen_fraction

  previous_q_values = agent_state[:2]
  previous_recency_effects = agent_state[2:4]

  # Link forgetting rates: use forgetting_rate directly for Q-values
  decayed_q_values = previous_q_values * forgetting_rate

  prediction_error = reward - decayed_q_values[choice]

  updated_q_values = decayed_q_values.at[choice].add(
      alpha_chosen * prediction_error
  )
  updated_q_values = updated_q_values.at[1 - choice].add(
      alpha_unchosen * -prediction_error
  )

  # Link forgetting rates: use shared forgetting_rate for recency effects
  decayed_recency_effects = previous_recency_effects * forgetting_rate

  recency_update_value = jnp.array([recency_loss_weight, recency_win_weight])[reward]
  scaled_recency_update = recency_update_value * (1.0 - forgetting_rate)

  updated_recency_effects = decayed_recency_effects.at[choice].add(
      scaled_recency_update
  )

  value_signal = updated_q_values * jnp.exp(updated_recency_effects)
  value_signal_with_bias = value_signal.at[0].add(bias_choice_0)
  rl_component = beta * value_signal_with_bias

  persistence_bonus = reward * persistence_rewarded - persistence_unrewarded
  recency_and_persistence_component = updated_recency_effects.at[choice].add(
      persistence_bonus
  )


  choice_logits = rl_component + recency_and_persistence_component

  new_agent_state = jnp.concatenate([updated_q_values, updated_recency_effects])

  return choice_logits, new_agent_state


def link_forgetting_ablate_persistence(
    params: chex.Array,
    choice: int,
    reward: int,
    agent_state: Optional[chex.Array],
) -> tuple[chex.Array, chex.Array]:
  """Modifies base_program. Links forgetting rates and ablates persistence."""

  if agent_state is None:
    agent_state = jnp.array([0.5, 0.5, 0.0, 0.0])

  (
    beta,
    persistence_unrewarded,
    recency_win_weight,
    recency_loss_weight,
  ) = map(jnp.abs, params[0:4])
  (
      forgetting_rate,       # Rate at which Q-values decay over time.
      alpha_chosen,          # Learning rate for the chosen option.
      alpha_unchosen_fraction, # Fraction of alpha_chosen for unchosen option.
  ) = map(jax.nn.sigmoid, params[4:7])
  bias_choice_0 = params[7]
  alpha_unchosen = alpha_chosen * alpha_unchosen_fraction

  previous_q_values = agent_state[:2]
  previous_recency_effects = agent_state[2:4]

  # Link forgetting rates: use shared forgetting_rate for Q-values
  decayed_q_values = previous_q_values * forgetting_rate

  prediction_error = reward - decayed_q_values[choice]

  updated_q_values = decayed_q_values.at[choice].add(
      alpha_chosen * prediction_error
  )
  updated_q_values = updated_q_values.at[1 - choice].add(
      alpha_unchosen * -prediction_error
  )

  # Link forgetting rates: use shared forgetting_rate for recency effects
  decayed_recency_effects = previous_recency_effects * forgetting_rate

  recency_update_value = jnp.array([recency_loss_weight, recency_win_weight])[reward]
  scaled_recency_update = recency_update_value * (1.0 - forgetting_rate)

  updated_recency_effects = decayed_recency_effects.at[choice].add(
      scaled_recency_update
  )

  value_signal = updated_q_values * jnp.exp(updated_recency_effects)
  value_signal_with_bias = value_signal.at[0].add(bias_choice_0)
  rl_component = beta * value_signal_with_bias

  # Remove persistence_rewarded
  persistence_bonus = reward - persistence_unrewarded
  recency_and_persistence_component = updated_recency_effects.at[choice].add(
      persistence_bonus
  )


  choice_logits = rl_component + recency_and_persistence_component

  new_agent_state = jnp.concatenate([updated_q_values, updated_recency_effects])

  return choice_logits, new_agent_state


def link_forgetting_ablate_persistence_simplified(
    params: chex.Array,
    choice: int,
    reward: int,
    agent_state: Optional[chex.Array],
) -> tuple[chex.Array, chex.Array]:
  """Simplification of link_forgetting_ablate_persistence to improve Halstead.
  """
  if agent_state is None:
    agent_state = jnp.array([0.5, 0.5, 0.0, 0.0])

  (
    beta,
    one_back_weight,
    recency_win_weight,
    recency_loss_weight,
  ) = map(jnp.abs, params[0:4])
  (
      forgetting_rate,    # Rate at which recency effects decay over time.
      alpha_chosen,          # Learning rate for the chosen option.
      alpha_unchosen_fraction, # Fraction of alpha_chosen for unchosen option.
  ) = map(jax.nn.sigmoid, params[4:7])
  bias_choice_0 = params[7]
  alpha_unchosen = alpha_chosen * alpha_unchosen_fraction

  # Apply forgetting to all state variables.
  decayed_agent_state = agent_state * forgetting_rate

  # Update fast learner, which has linked values
  decayed_q_values = decayed_agent_state[:2]
  prediction_error = reward - decayed_q_values[choice]
  updated_q_values = decayed_q_values.at[choice].add(
      alpha_chosen * prediction_error
  )
  updated_q_values = updated_q_values.at[1 - choice].add(
      alpha_unchosen * -prediction_error
  )

  # Update slow learner, which increments values and relies on forgetting
  decayed_recency_effects = decayed_agent_state[2:4]
  recency_update_value = jnp.array([recency_loss_weight, recency_win_weight])[reward]
  updated_recency_effects = decayed_recency_effects.at[choice].add(
      recency_update_value
  )

  # Compute previous trial win-stay/lose-shift effect
  one_back_bonus = reward - one_back_weight

  # Update state variables for fast and slow learners.
  new_agent_state = jnp.concatenate([updated_q_values, updated_recency_effects])

  # Compute choice logits.
  # Fast and slow learners
  value_signals = updated_recency_effects + beta * jnp.exp(updated_recency_effects) * updated_q_values
  # Add the previous trial win-stay/lose-shift bonus
  value_signals_with_oneback = value_signals.at[choice].add(one_back_bonus)
  # Add bias.
  choice_logits = value_signals_with_oneback.at[0].add(bias_choice_0)

  return choice_logits, new_agent_state


def link_forgetting_ablate_persistence_nowinweight_simplified(
    params: chex.Array,
    choice: int,
    reward: int,
    agent_state: Optional[chex.Array],
) -> tuple[chex.Array, chex.Array]:
  """Remove win-weight param from link_forgetting_ablate_persistence_simplified.
  """
  if agent_state is None:
    agent_state = jnp.array([0.5, 0.5, 0.0, 0.0])

  (
    beta,
    one_back_weight,
    recency_loss_weight,
  ) = map(jnp.abs, params[0:3])
  (
      forgetting_rate,    # Rate at which recency effects decay over time.
      alpha_chosen,          # Learning rate for the chosen option.
      alpha_unchosen_fraction, # Fraction of alpha_chosen for unchosen option.
  ) = map(jax.nn.sigmoid, params[3:6])
  bias_choice_0 = params[6]
  alpha_unchosen = alpha_chosen * alpha_unchosen_fraction

  # Apply forgetting to all state variables.
  decayed_agent_state = agent_state * forgetting_rate

  # Update fast learner, which has linked values
  decayed_q_values = decayed_agent_state[:2]
  prediction_error = reward - decayed_q_values[choice]
  updated_q_values = decayed_q_values.at[choice].add(
      alpha_chosen * prediction_error
  )
  updated_q_values = updated_q_values.at[1 - choice].add(
      alpha_unchosen * -prediction_error
  )

  # Update slow learner, which increments values and relies on forgetting
  decayed_recency_effects = decayed_agent_state[2:4]
  updated_recency_effects = decayed_recency_effects.at[choice].add(
      recency_loss_weight * (1 - reward)
  )

  # Compute previous trial win-stay/lose-shift effect
  one_back_bonus = reward - one_back_weight

  # Update state variables for fast and slow learners.
  new_agent_state = jnp.concatenate([updated_q_values, updated_recency_effects])

  # Compute choice logits.
  # Fast and slow learners
  value_signals = updated_recency_effects + beta * jnp.exp(updated_recency_effects) * updated_q_values
  # Add the previous trial win-stay/lose-shift bonus
  value_signals_with_oneback = value_signals.at[choice].add(one_back_bonus)
  # Add bias.
  choice_logits = value_signals_with_oneback.at[0].add(bias_choice_0)

  return choice_logits, new_agent_state


def yoked(
    params: chex.Array,
    choice: int,
    reward: int,
    agent_state: Optional[chex.Array],
) -> tuple[chex.Array, chex.Array]:
  """Yoked q-values to use just one variable for both Q-left and Q-right."""
  # --- 1. Initialization and Parameter Unpacking ---

  # On the first trial (when agent_state is None), initialize the state.
  # The state consists of [reward_seeking, recency_0, recency_1].
  if agent_state is None:
    agent_state = jnp.array([0.5, 0.5, 0.0, 0.0])

  (
      alpha_chosen,
      recency_decay_rate,
  ) = map(jax.nn.sigmoid, params[0:2])
  (
    beta,
    one_back_weight,
    recency_win_weight,
    recency_loss_weight,
  ) = map(jnp.abs, params[2:6])
  bias_choice_0 = params[6]

  q_values = agent_state[0:2]
  previous_recency_effects = agent_state[2:4]

  # --- 2. Update Reward-Seeking (Reinforcement Learning) ---
  prediction_error = reward - q_values[choice]
  partially_updated_q_values = q_values.at[choice].add(alpha_chosen * prediction_error)
  updated_q_values = partially_updated_q_values.at[1 - choice].add(-alpha_chosen * prediction_error)

  # --- 3. Update Recency Effects ---
  decayed_recency_effects = previous_recency_effects * recency_decay_rate
  recency_update_value = jnp.array([recency_loss_weight, recency_win_weight])[reward]
  updated_recency_effects = decayed_recency_effects.at[choice].add(
      recency_update_value
  )

  # --- 4. Calculate Choice Logits ---
  value_and_recency = beta * updated_q_values * jnp.exp(updated_recency_effects) + updated_recency_effects
  # Add bias.
  value_recency_and_bias = value_and_recency.at[0].add(bias_choice_0)
  # Add one-back weight. This mediates win-stay/lose-shift.
  choice_logits = value_recency_and_bias.at[choice].add(reward - one_back_weight)

  new_agent_state = jnp.concatenate([updated_q_values, updated_recency_effects])

  return choice_logits, new_agent_state

#######################################
## AUTOMATICALLY GENERATED PROGRAMS  ##
## Made by DataDIVER                 ##
#######################################


# Shortened to 100k steps then refactored, run 1, low floor.
def rat_bandit_run1_simplified_low_floor(
    params: chex.Array,
    choice: int,
    reward: int,
    agent_state: Optional[chex.Array],
) -> tuple[chex.Array, chex.Array]:
  """Cognitive model describing rat behavior on a binary two-armed bandit task.

  Args:
    params: Model params. Different parameters are used for different rats.
    choice: Previous choice. Values: 0 or 1.
    reward: Previous reward. Values: 0 or 1.
    agent_state: Previous state of the agent.

  Returns:
    choice_logits: Vector of shape (2,) with the probabilities that the rat will
      choose option 0 or 1 on the next trial, expressed as logits.
    agent_state: New state of the agent, after observing the previous choice and
      reward.
  """

  # --- 1. Initialize or Unpack Agent State ---
  # The agent's state tracks two key components:
  # - Q-values: The learned values for choosing option 0 or 1.
  # - Net Recency: A tracker for recent wins vs. losses for each option.
  
  # On the first trial, agent_state is None, so we initialize it.
  # Q-values start at 0.5 (neutral), and recency trackers start at 0.0.
  if agent_state is None:
    initial_q_values = jnp.array([0.5, 0.5])
    initial_net_recency = jnp.array([0.0, 0.0])
    agent_state = jnp.concatenate([initial_q_values, initial_net_recency])

  # Unpack the state from the previous trial for clarity.
  old_q_values = agent_state[:2]
  old_net_recency = agent_state[2:4]

  # --- 2. Unpack and Transform Model Parameters ---
  # Unpack the raw model parameters.
  (beta, alpha_chosen_raw, alpha_unchosen_raw,
   persistence, persistence_reward_effect, forgetting_rate_raw,
   bias_choice_0, recency_decay_rate_raw, recency_win_weight,
   recency_loss_weight) = params

  # Transform raw parameters into a usable range (0 to 1) using a sigmoid.
  # This is common for rates (learning, forgetting, decay).
  alpha_chosen = jax.nn.sigmoid(alpha_chosen_raw)
  alpha_unchosen = jax.nn.sigmoid(alpha_unchosen_raw)
  forgetting_rate = jax.nn.sigmoid(forgetting_rate_raw)
  recency_decay_rate = jax.nn.sigmoid(recency_decay_rate_raw)

  # --- 3. Update Q-Values (Learning) ---
  # This update follows a modified Q-learning rule with forgetting.

  # First, apply a forgetting factor to both Q-values, causing them to decay
  # towards their initial value (or zero, depending on the model interpretation).
  q_values_after_forgetting = old_q_values * (1.0 - forgetting_rate)
  
  # Calculate the prediction error: the difference between the actual reward
  # and the expected reward (the Q-value of the chosen option).
  prediction_error = reward - old_q_values[choice]
  
  # Update the Q-value of the CHOSEN option based on the prediction error.
  # This is the standard Rescorla-Wagner update.
  updated_q_for_chosen = q_values_after_forgetting[choice] + alpha_chosen * prediction_error
  
  # Update the Q-value of the UNCHOSEN option. This model assumes a form of
  # counterfactual learning, where the unchosen option's value moves in the
  # opposite direction of the chosen one.
  updated_q_for_unchosen = q_values_after_forgetting[1 - choice] - alpha_unchosen * prediction_error

  # Reconstruct the full Q-value vector.
  new_q_values = q_values_after_forgetting.at[choice].set(updated_q_for_chosen)
  new_q_values = new_q_values.at[1 - choice].set(updated_q_for_unchosen)
  
  # --- 4. Update Net Recency Trackers ---
  # These trackers capture the recent history of wins and losses for each choice.
  
  # First, decay the previous recency values.
  recency_after_decay = old_net_recency * recency_decay_rate
  
  # Calculate the new information to add for the chosen option.
  # This value is positive for a win and negative for a loss.
  recency_update_value = (reward * recency_win_weight - recency_loss_weight)
  
  # Integrate the new information into the recency tracker for the chosen option.
  # The (1 - decay_rate) term ensures the update is properly scaled.
  recency_increment = recency_update_value * (1.0 - recency_decay_rate)
  
  # Add the increment to the chosen option's tracker.
  new_net_recency = recency_after_decay.at[choice].add(recency_increment)
  
  # --- 5. Assemble New Agent State ---
  # Combine the updated components into the new state vector for the next trial.
  new_agent_state = jnp.concatenate([new_q_values, new_net_recency])

  # --- 6. Calculate Choice Logits for the Next Trial ---
  # Logits determine the probability of choosing each option on the next trial.
  
  # Start with the learned Q-values and add a static bias for option 0.
  value_with_bias = new_q_values.at[0].add(bias_choice_0)
  
  # Add the net recency trackers to the values.
  total_learned_value = value_with_bias + new_net_recency
  
  # Scale the total values by beta (inverse temperature), which controls
  # the level of choice stochasticity (exploration vs. exploitation).
  base_logits = beta * total_learned_value

  # Calculate the persistence bonus, which encourages repeating the last choice.
  # The bonus is larger if the last choice was rewarded.
  persistence_bonus = persistence + persistence_reward_effect * reward
  
  # Add the persistence bonus to the logit of the option that was just chosen.
  choice_logits = base_logits.at[choice].add(persistence_bonus)
  
  return choice_logits, new_agent_state


# Shortened to 100k steps then refactored, run 2, low floor.
def rat_bandit_run2_simplified_low_floor(
    params: chex.Array,
    choice: int,
    reward: int,
    agent_state: Optional[chex.Array],
) -> tuple[chex.Array, chex.Array]:
  """Cognitive model describing rat behavior on a binary two-armed bandit task.

  This model incorporates several cognitive mechanisms:
  1.  **Reinforcement Learning:** Learns the value of choices (Q-values) from
      rewards, using different learning rates for positive and negative outcomes.
  2.  **Ghost Values:** A latent value system that interacts with the primary
      Q-values and allows for counterfactual updates.
  3.  **Perseveration/Stickiness:** A tendency to repeat recent choices,
      modeled via a decaying choice trace.

  Args:
    params: Model params. Different parameters are used for different rats.
    choice: Previous choice. Values: 0 or 1.
    reward: Previous reward. Values: 0 or 1.
    agent_state: Previous state of the agent.

  Returns:
    choice_logits: Vector of shape (2,) with the probabilities that the rat will
      choose option 0 or 1 on the next trial, expressed as logits.
    agent_state: New state of the agent, after observing the previous choice and
      reward.
  """
  # --- 1. Unpack and transform model parameters ---

  # Unpack the flat parameter array into meaningful groups.
  (initial_q_value, *learning_rate_params, inverse_temperature_param,
   _unused_choice_bias, stickiness_weight) = params

  # Apply transformations to constrain parameter ranges.
  # Sigmoid squashes learning rate-related parameters to be between 0 and 1.
  (positive_learning_rate, negative_learning_rate, q_ghost_pull_rate,
   unchosen_ghost_reward_influence, ghost_q_decay_rate, trace_learning_rate) = \
      jax.nn.sigmoid(jnp.array(learning_rate_params))

  # Exponential ensures the temperature parameter is positive.
  inverse_temperature = jnp.exp(inverse_temperature_param)

  # --- 2. Initialize or unpack agent's internal state ---

  # If this is the first trial (no previous state), initialize the agent's state.
  if agent_state is None:
    # The state vector consists of [q_values, ghost_q_values, choice_traces].
    # All are initialized with the same starting value parameter.
    # The total size is 6 (2 for each component).
    agent_state = jnp.array([initial_q_value] * 6)

  # Unpack the agent state vector into its three core components.
  # q_values: Standard reinforcement learning action values for each choice. (2 elements)
  # ghost_q_values: Latent values that interact with q_values. (2 elements)
  # choice_traces: A memory of recent choices to model stickiness. (2 elements)
  q_values, ghost_q_values, choice_traces = jnp.split(agent_state, [2, 4])

  # --- 3. Update state based on previous choice and reward ---

  # a) Standard Q-learning update for the chosen action.
  # Calculate the prediction error: the difference between actual reward and expected value.
  prediction_error = reward - q_values[choice]

  # Use different learning rates for positive (better than expected) and
  # negative (worse than expected) prediction errors.
  learning_rate_for_update = jnp.where(
      prediction_error >= 0,
      positive_learning_rate,
      negative_learning_rate
  )

  # Calculate the update amount and apply it to the Q-value for the chosen action.
  q_value_update = learning_rate_for_update * prediction_error
  q_values = q_values.at[choice].add(q_value_update)

  # b) Interaction between standard Q-values and "ghost" Q-values.
  # The main Q-values are "pulled" towards the ghost Q-values.
  q_values_pulled_by_ghost = q_ghost_pull_rate * (ghost_q_values - q_values)
  q_values = q_values + q_values_pulled_by_ghost

  # The ghost Q-values decay towards the main Q-values.
  ghost_q_decay = ghost_q_decay_rate * (q_values - ghost_q_values)
  ghost_q_values = ghost_q_values + ghost_q_decay

  # The ghost Q-value for the *unchosen* action is updated by the reward.
  # This is a form of counterfactual learning.
  unchosen_choice = 1 - choice
  counterfactual_error = reward - ghost_q_values[unchosen_choice]
  ghost_q_counterfactual_update = unchosen_ghost_reward_influence * counterfactual_error
  ghost_q_values = ghost_q_values.at[unchosen_choice].add(ghost_q_counterfactual_update)

  # c) Update choice traces to model perseveration (stickiness).
  # All traces decay over time.
  choice_traces = choice_traces * (1 - trace_learning_rate)

  # The trace for the chosen action is strengthened (set closer to 1).
  choice_traces = choice_traces.at[choice].add(trace_learning_rate)

  # --- 4. Reconstruct agent state and compute choice logits for the next trial ---

  # Combine the updated components back into a single state vector for the next trial.
  agent_state = jnp.concatenate([q_values, ghost_q_values, choice_traces])

  # Calculate the choice logits. This involves two components: value and stickiness.
  # The value component is based on the learned Q-values.
  # The inverse_temperature parameter controls the greediness of the choice.
  value_based_logits = inverse_temperature * q_values

  # The stickiness component adds a bias to repeat the previous choice.
  stickiness_bias = stickiness_weight * choice_traces

  # The final logits are a sum of the value and stickiness components.
  choice_logits = value_based_logits + stickiness_bias

  # --- 5. Return the results ---
  return choice_logits, agent_state


# Shortened to 100k steps then refactored, run 3, low floor.
def rat_bandit_run3_simplified_low_floor(
    params: chex.Array,
    choice: int,
    reward: int,
    agent_state: Optional[chex.Array],
) -> tuple[chex.Array, chex.Array]:
  """
  A reinforcement learning agent that updates its state based on a choice and reward.
  """
  # --- 1. Initialize Agent State ---
  # If this is the first trial, create a new state vector. Otherwise, use the
  # provided state from the previous trial.
  # The state vector contains:
  # [q_value_choice_0, q_value_choice_1,
  #  reward_eligibility_trace_0, reward_eligibility_trace_1,
  #  no_reward_eligibility_trace_0, no_reward_eligibility_trace_1]
  if agent_state is None:
    agent_state = jnp.array([0.5, 0.5, 0.0, 0.0, 0.0, 0.0])

  # --- 2. Unpack and Transform Model Parameters ---
  # Parameters are extracted from the input array and transformed into valid
  # ranges (e.g., positive for temperature, or between 0 and 1 for rates).

  # Inverse temperature controls policy stochasticity. Softplus ensures it is positive.
  inverse_temperature = jax.nn.softplus(params[0])

  # Other parameters are mapped to the (0, 1) range using the sigmoid function.
  learning_rate = jax.nn.sigmoid(params[1])
  unchosen_learning_rate = jax.nn.sigmoid(params[2])
  perseveration_strength_reward = jax.nn.sigmoid(params[3])
  perseveration_strength_no_reward = jax.nn.sigmoid(params[4])
  reward_eligibility_decay_rate = jax.nn.sigmoid(params[5])
  no_reward_eligibility_decay_rate = jax.nn.sigmoid(params[6])
  _unused_global_bias_0 = jax.nn.sigmoid(params[7])
  q_decay_rate = jax.nn.sigmoid(params[8])
  q_decay_baseline = jax.nn.sigmoid(params[9])

  # --- 3. Unpack Agent State Components ---
  # The agent's state vector is split into its meaningful parts.
  q_values, eligibility_traces_reward, eligibility_traces_no_reward = jnp.split(
      agent_state, [2, 4]
  )

  # --- 4. Apply Q-value Decay ---
  # Q-values gradually decay towards a baseline value, modeling a forgetting process.
  q_values_decayed = q_values + q_decay_rate * (q_decay_baseline - q_values)

  # --- 5. Update Q-values based on Reward ---
  # This section implements a modified Rescorla-Wagner learning rule.

  # Create a one-hot vector to isolate the chosen action.
  # e.g., if choice is 1, this becomes [0, 1].
  choice_vector = jax.nn.one_hot(choice, num_classes=2)

  # Calculate the prediction error: the difference between the actual reward
  # and the expected reward (the Q-value of the chosen action).
  q_value_of_chosen_action = q_values_decayed[choice]
  prediction_error = reward - q_value_of_chosen_action

  # Calculate the updates for the chosen and unchosen actions. The chosen
  # action is updated by the learning rate, while the unchosen action is
  # updated by a separate (negative) rate.
  chosen_action_update = learning_rate * prediction_error
  unchosen_action_update = -unchosen_learning_rate * prediction_error

  # Create a full update vector using the one-hot choice vector.
  q_value_updates = (
      chosen_action_update * choice_vector
      + unchosen_action_update * (1 - choice_vector)
  )

  # Apply the updates to the decayed Q-values to get the new Q-values.
  q_values_updated = q_values_decayed + q_value_updates

  # --- 6. Update Eligibility Traces ---
  # Eligibility traces track recent actions to model perseveration (the tendency
  # to repeat actions). They are updated separately for rewarded and unrewarded trials.

  # First, decay the existing reward eligibility traces.
  # Then, add 1 to the trace for the chosen action if a reward was received.
  eligibility_traces_reward_updated = (
      reward_eligibility_decay_rate * eligibility_traces_reward
      + reward * choice_vector
  )

  # First, decay the existing no-reward eligibility traces.
  # Then, add 1 to the trace for the chosen action if no reward was received.
  eligibility_traces_no_reward_updated = (
      no_reward_eligibility_decay_rate * eligibility_traces_no_reward
      + (1 - reward) * choice_vector
  )

  # --- 7. Reassemble the Updated Agent State ---
  # The updated components are combined back into a single state vector.
  updated_agent_state = jnp.concatenate(
      [
          q_values_updated,
          eligibility_traces_reward_updated,
          eligibility_traces_no_reward_updated,
      ]
  )

  # --- 8. Calculate Action Logits for the Next Choice ---
  # Logits determine the probability of choosing each action. They are a sum
  # of influences from Q-values and perseveration tendencies.

  # 1. Influence from the learned value (Q-values) of each action.
  q_value_influence = inverse_temperature * q_values_updated

  # 2. Influence from the tendency to repeat a recently rewarded action.
  reward_perseveration_influence = (
      perseveration_strength_reward * eligibility_traces_reward_updated
  )

  # 3. Influence from the tendency to repeat a recently unrewarded action.
  no_reward_perseveration_influence = (
      perseveration_strength_no_reward * eligibility_traces_no_reward_updated
  )

  # The final logits are the sum of these three influences.
  choice_logits = (
      q_value_influence
      + reward_perseveration_influence
      + no_reward_perseveration_influence
  )

  # --- 9. Return Logits and New State ---
  # Return the logits for the next action selection and the updated agent state.
  return choice_logits, updated_agent_state


# Shortened to 100k steps then refactored, run 1, medium floor.
def rat_bandit_run1_simplified_medium_floor(
    params: chex.Array,
    choice: int,
    reward: int,
    agent_state: Optional[chex.Array],
) -> tuple[chex.Array, chex.Array]:
  """Cognitive model describing rat behavior on a binary two-armed bandit task.

  Args:
    params: Model params. Different parameters are used for different rats.
    choice: Previous choice. Values: 0 or 1.
    reward: Previous reward. Values: 0 or 1.
    agent_state: Previous state of the agent

  Returns:
    choice_logits: Vector of shape (2,) with the probabilities that the rat will
      choose option 0 or 1 on the next trial, expressed as logits.
    agent_state: New state of the agent, after observing the previous choice and
      reward.
  """
  # --- 1. Initialization and Parameter Unpacking ---

  # On the first trial (when agent_state is None), initialize the state.
  # The state consists of [Q-value for choice 0, Q-value for choice 1,
  #                       Recency for choice 0,  Recency for choice 1].
  # Q-values are initialized to 0.5 (neutral), recency effects to 0.0.
  if agent_state is None:
    agent_state = jnp.array([0.5, 0.5, 0.0, 0.0])

  # Unpack the model's parameters from the input array.
  log_beta, *sigmoid_params, bias_choice_0, recency_win_weight, recency_loss_weight = params

  # Apply the sigmoid function to constrain certain parameters to be between 0 and 1.
  # This is a standard method for parameters representing rates or probabilities.
  (
      alpha_chosen,          # Learning rate for the chosen option.
      alpha_unchosen,        # Learning rate for the unchosen option.
      persistence_rewarded,  # Weight for repeating a rewarded choice.
      persistence_unrewarded,# Weight for repeating an unrewarded choice.
      forgetting_rate,       # Rate at which Q-values decay over time.
      recency_decay_rate,    # Rate at which recency effects decay over time.
  ) = map(jax.nn.sigmoid, sigmoid_params)

  # Decompose the agent_state vector into its meaningful components.
  previous_q_values = agent_state[:2]
  previous_recency_effects = agent_state[2:4]

  # --- 2. Update Q-Values (Reinforcement Learning) ---

  # Apply a forgetting factor to the previous Q-values.
  decayed_q_values = previous_q_values * (1.0 - forgetting_rate)

  # Calculate the prediction error: the difference between the actual reward
  # and the expected reward (the Q-value of the chosen option).
  prediction_error = reward - decayed_q_values[choice]

  # Update the Q-value for the chosen option using the prediction error.
  # Note: JAX arrays are immutable, so .at[].add() creates a new array.
  updated_q_values = decayed_q_values.at[choice].add(
      alpha_chosen * prediction_error
  )
  # Update the Q-value for the unchosen option. This model assumes it is
  # updated in the opposite direction of the chosen option's error.
  updated_q_values = updated_q_values.at[1 - choice].add(
      alpha_unchosen * -prediction_error
  )

  # --- 3. Update Recency Effects ---

  # Decay the previous recency effects.
  decayed_recency_effects = previous_recency_effects * recency_decay_rate

  # Determine the "recency shock" from the last outcome.
  # A reward (reward=1) adds recency_win_weight, a non-reward (reward=0)
  # adds -recency_loss_weight, modeling a win-stay/lose-shift tendency.
  recency_update_value = jnp.array([-recency_loss_weight, recency_win_weight])[reward]
  scaled_recency_update = recency_update_value * (1.0 - recency_decay_rate)

  # Apply this update to the recency trace of the chosen option.
  updated_recency_effects = decayed_recency_effects.at[choice].add(
      scaled_recency_update
  )

  # --- 4. Calculate Choice Logits for the Next Trial ---

  # The logits determine the probability of the next choice. They are a sum
  # of several cognitive components: value, bias, recency, and persistence.

  # Convert log_beta to beta, the inverse temperature parameter that controls
  # the level of determinism in the choice. Higher beta means less random choices.
  beta = jnp.exp(log_beta)

  # Component 1: Value-based term (RL).
  # The Q-values are modulated by an exponential of the recency effects.
  value_signal = updated_q_values * jnp.exp(updated_recency_effects)
  # A static bias towards choice 0 is added to the value signal.
  value_signal_with_bias = value_signal.at[0].add(bias_choice_0)
  # This entire value component is then scaled by beta.
  rl_component = beta * value_signal_with_bias

  # Component 2: Persistence and Recency term.
  # A persistence bonus is calculated for the action that was just taken.
  # The size of the bonus depends on whether the action was rewarded.
  persistence_bonus = reward * persistence_rewarded - persistence_unrewarded
  # This bonus is added to the recency effect of the chosen action.
  recency_and_persistence_component = updated_recency_effects.at[choice].add(
      persistence_bonus
  )

  # The final choice logits are the sum of the two main components.
  choice_logits = rl_component + recency_and_persistence_component

  # --- 5. Prepare Outputs ---

  # Construct the new agent state for the next trial by combining the
  # updated Q-values and recency effects into a single array.
  new_agent_state = jnp.concatenate([updated_q_values, updated_recency_effects])

  return choice_logits, new_agent_state


# Shortened to 100k steps then refactored, run 2, medium floor.
def rat_bandit_run2_simplified_medium_floor(
    params: chex.Array,
    choice: int,
    reward: int,
    agent_state: Optional[chex.Array],
) -> tuple[chex.Array, chex.Array]:
  """Cognitive model describing rat behavior on a binary two-armed bandit task.

  Args:
    params: Model params. Different parameters are used for different rats.
    choice: Previous choice. Values: 0 or 1.
    reward: Previous reward. Values: 0 or 1.
    agent_state: Previous state of the agent.

  Returns:
    choice_logits: Vector of shape (2,) with the probabilities that the rat will
      choose option 0 or 1 on the next trial, expressed as logits.
    agent_state: New state of the agent, after observing the previous choice and
      reward.
  """
  # --- 1. Unpack and Transform Model Parameters ---

  # Unpack the raw, fitted parameters from the input array.
  initial_q_value, positive_learning_rate_raw, negative_learning_rate_raw, \
  q_ghost_pull_rate_raw, unchosen_ghost_learning_rate_raw, \
  chosen_ghost_pull_rate_raw, trace_learning_rate_raw, \
  stickiness_weight, inverse_temperature_raw, choice_bias = params

  # Transform raw parameters into a usable range.
  # Learning rates are bounded between 0 and 1 using a sigmoid function.
  positive_learning_rate, negative_learning_rate, q_ghost_pull_rate, \
  unchosen_ghost_learning_rate, chosen_ghost_pull_rate, trace_learning_rate = jax.nn.sigmoid(
      jnp.array([
          positive_learning_rate_raw,
          negative_learning_rate_raw,
          q_ghost_pull_rate_raw,
          unchosen_ghost_learning_rate_raw,
          chosen_ghost_pull_rate_raw,
          trace_learning_rate_raw,
      ])
  )
  # Inverse temperature (decision noise) is kept positive using an exponential function.
  inverse_temperature = jnp.exp(inverse_temperature_raw)


  # --- 2. Unpack or Initialize Agent's Internal State ---

  if agent_state is None:
    # If this is the first trial, initialize the agent's state.
    # Q-values represent the expected reward for each choice.
    q_values = jnp.array([initial_q_value, initial_q_value])
    # Ghost Q-values are a secondary trace of expected rewards.
    ghost_q_values = jnp.array([initial_q_value, initial_q_value])
    # Choice traces track the recent history of choices.
    choice_traces = jnp.array([0.0, 0.0])
  else:
    # If not the first trial, unpack the state from the previous step.
    q_values, ghost_q_values, choice_traces = agent_state


  # --- 3. Core Reinforcement Learning Computations ---

  # Calculate the prediction error: the difference between the actual reward
  # and the expected reward (Q-value) for the chosen action.
  prediction_error = reward - q_values[choice]

  # Create a factor that scales some learning rates based on the magnitude
  # of the prediction error. Larger errors lead to larger updates.
  prediction_error_sensitivity_factor = 1 + jnp.abs(prediction_error)

  # Determine which learning rate to use based on whether the outcome was
  # better (positive error) or worse (negative error) than expected.
  learning_rate_for_chosen_action = jnp.where(
      prediction_error >= 0,
      positive_learning_rate,
      negative_learning_rate
  )


  # --- 4. Update Internal State Variables ---

  # --- 4a. Update Q-values ---
  # First, calculate the standard Q-learning update for the chosen action.
  q_values_after_rl_update = q_values.at[choice].add(
      learning_rate_for_chosen_action * prediction_error
  )
  # Next, "pull" the updated Q-values towards the ghost Q-values. This blends
  # the standard Q-values with the secondary ghost trace.
  updated_q_values = (1 - q_ghost_pull_rate) * q_values_after_rl_update \
                   + q_ghost_pull_rate * ghost_q_values

  # --- 4b. Update Ghost Q-values ---
  # Identify the action that was *not* chosen.
  unchosen_action = 1 - choice

  # Update the ghost Q-value for the *unchosen* action. It learns from a
  # hypothetical reward target, which is the average of the actual reward and
  # the updated Q-value of the unchosen action.
  ghost_q_target_for_unchosen = 0.5 * (reward + updated_q_values[unchosen_action])
  ghost_q_error_for_unchosen = ghost_q_target_for_unchosen - ghost_q_values[unchosen_action]
  ghost_q_values_intermediate = ghost_q_values.at[unchosen_action].add(
      unchosen_ghost_learning_rate * prediction_error_sensitivity_factor * ghost_q_error_for_unchosen
  )

  # Update the ghost Q-value for the *chosen* action. It is pulled towards the
  # newly updated main Q-value for that action.
  ghost_q_target_for_chosen = updated_q_values[choice]
  ghost_q_error_for_chosen = ghost_q_target_for_chosen - ghost_q_values_intermediate[choice]
  updated_ghost_q_values = ghost_q_values_intermediate.at[choice].add(
      chosen_ghost_pull_rate * prediction_error_sensitivity_factor * ghost_q_error_for_chosen
  )

  # --- 4c. Update Choice Traces ---
  # This trace tracks choice history, creating a "stickiness" or perseveration effect.
  # First, all traces decay over time.
  decayed_choice_traces = (1 - trace_learning_rate) * choice_traces
  # Then, the trace for the chosen action is increased, modulated by the
  # prediction error sensitivity.
  trace_update_increment = trace_learning_rate * prediction_error_sensitivity_factor * (1 - choice_traces[choice])
  updated_choice_traces = decayed_choice_traces.at[choice].add(
      (1 - trace_learning_rate) * trace_update_increment
  )

  # --- 5. Combine State and Calculate Next Choice Probabilities ---

  # Bundle the updated internal variables into the new agent state.
  updated_agent_state = jnp.array([
      updated_q_values,
      updated_ghost_q_values,
      updated_choice_traces
  ])

  # Calculate the choice logits for the next trial. This combines three factors:
  # 1. The learned Q-values (value-based preference).
  # 2. A static bias for one choice over the other.
  # 3. The choice trace (stickiness/perseveration).
  value_component = inverse_temperature * updated_q_values
  bias_component = jnp.array([choice_bias, 0.0])
  stickiness_component = stickiness_weight * updated_choice_traces

  choice_logits = value_component + bias_component + stickiness_component

  return choice_logits, updated_agent_state


# Shortened to 100k steps then refactored, run 3, medium floor.
def rat_bandit_run3_simplified_medium_floor(
    params: chex.Array,
    choice: int,
    reward: int,
    agent_state: Optional[chex.Array],
) -> tuple[chex.Array, chex.Array]:
  """Cognitive model describing rat behavior on a binary two-armed bandit task.

  Args:
    params: Model params. Different parameters are used for different rats.
    choice: Previous choice. Values: 0 or 1.
    reward: Previous reward. Values: 0 or 1.
    agent_state: Previous state of the agent

  Returns:
    choice_logits: Vector of shape (2,) with the probabilities that the rat will
      choose option 0 or 1 on the next trial, expressed as logits.
    agent_state: New state of the agent, after observing the previous choice and
      reward.
  """

  # --- 1. Unpack and Transform Model Parameters ---
  # Raw parameters are unpacked from the input array.
  (inverse_temperature_raw, *sigmoid_params_raw, global_bias_for_choice_0,
   prediction_error_eligibility_weight, prediction_error_asymmetry_weight) = params

  # Transform raw parameters to ensure they are in a valid range.
  # Use exp to ensure inverse_temperature is positive.
  inverse_temperature = jnp.exp(inverse_temperature_raw)
  # Use a sigmoid function to constrain rates and strengths between 0 and 1.
  (learning_rate, perseveration_strength, eligibility_decay_rate, q_decay_rate,
   reward_history_learning_rate, action_bias_learning_rate) = jax.nn.sigmoid(
       jnp.array(sigmoid_params_raw))


  # --- 2. Initialize or Unpack Agent State ---
  # The agent's state tracks internal variables across trials.
  if agent_state is None:
    # If this is the first trial, initialize the agent's state.
    # The state vector contains Q-values, eligibility traces, reward history,
    # and action biases. Here, all are initialized to a default value.
    initial_value = jax.nn.sigmoid(prediction_error_eligibility_weight)
    agent_state = jnp.full(8, initial_value)

  # For subsequent trials, unpack the existing state into its components.
  # Each component is a vector of size 2 (one for each choice option).
  q_values, eligibility_traces, reward_history, action_biases = jnp.split(agent_state, 4)


  # --- 3. Update State Based on Last Trial's Outcome ---
  # Create a one-hot encoded vector for the previous choice.
  # e.g., if choice was 0, one_hot_choice = [1, 0].
  one_hot_choice = jax.nn.one_hot(choice, num_classes=2)

  # a) Update the tracked average reward for each action (reward history).
  # This is a simple delta-rule update for the chosen action.
  reward_history_delta = reward - reward_history
  reward_history_updated = reward_history + reward_history_learning_rate * one_hot_choice * reward_history_delta

  # b) Compute the prediction error: the difference between actual reward
  # and the expected reward (Q-value) for the chosen action.
  prediction_error = reward - q_values[choice]

  # c) Update Q-values (action values).
  # First, Q-values decay towards the long-term average reward history.
  q_decay_term = q_decay_rate * (reward_history_updated - q_values)
  # Second, the Q-value for the chosen action is updated by the prediction error.
  # The update can be asymmetric for positive vs. negative prediction errors.
  asymmetric_prediction_error = prediction_error + prediction_error_asymmetry_weight * jnp.abs(prediction_error)
  q_learning_term = one_hot_choice * learning_rate * asymmetric_prediction_error
  # Combine decay and learning to get the final updated Q-values.
  q_values_updated = q_values + q_decay_term + q_learning_term

  # d) Update eligibility traces. These track recent actions to influence
  # perseveration (tendency to repeat actions).
  # The trace for the chosen action is boosted, modulated by prediction error "salience".
  prediction_error_salience = 1 + prediction_error_eligibility_weight * jnp.abs(prediction_error)
  eligibility_trace_target = one_hot_choice * prediction_error_salience
  # Traces decay towards their new target values.
  eligibility_traces_updated = eligibility_traces + eligibility_decay_rate * (eligibility_trace_target - eligibility_traces)

  # e) Update action biases. This is another mechanism for perseveration,
  # capturing a tendency to repeat rewarded actions.
  # The bias for the chosen action moves towards 2 if rewarded (reward=1) and 0 if not.
  # The bias for the unchosen action decays towards 0.
  action_bias_target = one_hot_choice * (2 * reward)
  action_biases_updated = action_biases + action_bias_learning_rate * (action_bias_target - action_biases)


  # --- 4. Prepare for Next Trial ---
  # Reconstruct the agent's state vector for the next trial.
  new_agent_state = jnp.concatenate([
      q_values_updated,
      eligibility_traces_updated,
      reward_history_updated,
      action_biases_updated
  ])

  # Compute the choice logits (action preferences) for the upcoming trial.
  # Logits are computed by summing three distinct components.

  # Component a) Value-based preference (exploitation):
  # The difference between Q-values and the average reward, scaled by temperature.
  value_component = inverse_temperature * (q_values_updated - reward_history_updated)

  # Component b) Global bias:
  # A constant preference for one action over the other (here, for choice 0).
  global_bias_component = jnp.array([global_bias_for_choice_0, 0.0])

  # Component c) Perseveration/recency effects:
  # A tendency to repeat recent actions, driven by eligibility traces and action biases.
  perseveration_component = perseveration_strength * (eligibility_traces_updated + action_biases_updated)

  # Sum the components to get the final choice logits.
  choice_logits = value_component + global_bias_component + perseveration_component

  return choice_logits, new_agent_state


# Shortened to 100k steps then refactored, run 2, high floor.
def rat_bandit_run2_simplified_high_floor(
    params: chex.Array,
    choice: int,
    reward: int,
    agent_state: Optional[chex.Array],
) -> tuple[chex.Array, chex.Array]:
  """Cognitive model describing rat behavior on a binary two-armed bandit task.

  Args:
    params: Model params. Different parameters are used for different rats.
    choice: Previous choice. Values: 0 or 1.
    reward: Previous reward. Values: 0 or 1.
    agent_state: Previous state of the agent

  Returns:
    choice_logits: Vector of shape (2,) with the probabilities that the rat will
      choose option 0 or 1 on the next trial, expressed as logits.
    agent_state: New state of the agent, after observing the previous choice and
      reward.
  """
  # --- 1. Unpack and Transform Model Parameters ---

  # Unpack the raw, untransformed parameters from the input array.
  # These parameters are transformed before use to ensure they are in a valid range.
  (initial_q_value, positive_learning_rate_param,
   negative_learning_rate_param, unchosen_q_update_learning_rate_param,
   chosen_q_ghost_influence_rate_param, ghost_learning_rate_param,
   trace_learning_rate_param, inverse_temperature_0_param,
   inverse_temperature_1_param, choice_bias) = params

  # Transform learning rate parameters using a sigmoid function to constrain them
  # to the range (0, 1).
  positive_base_learning_rate = jax.nn.sigmoid(positive_learning_rate_param)
  negative_base_learning_rate = jax.nn.sigmoid(negative_learning_rate_param)
  unchosen_q_update_learning_rate = jax.nn.sigmoid(
      unchosen_q_update_learning_rate_param)
  chosen_q_ghost_influence_rate = jax.nn.sigmoid(
      chosen_q_ghost_influence_rate_param)
  ghost_learning_rate = jax.nn.sigmoid(ghost_learning_rate_param)
  trace_learning_rate = jax.nn.sigmoid(trace_learning_rate_param)

  # Transform inverse temperature parameters using an exponential function to
  # ensure they are positive.
  inverse_temperature_0 = jnp.exp(inverse_temperature_0_param)
  inverse_temperature_1 = jnp.exp(inverse_temperature_1_param)
  inverse_temperatures = jnp.array([inverse_temperature_0, inverse_temperature_1])

  # --- 2. Initialize or Unpack Agent State ---

  # If this is the first trial (agent_state is None), initialize the state.
  # The state vector contains:
  #   - 2 Q-values (one for each choice)
  #   - 2 "ghost" Q-values (representing counterfactual outcomes)
  #   - 2 choice traces (a memory of recent choices)
  if agent_state is None:
    initial_q_values = jnp.array([initial_q_value, initial_q_value])
    initial_ghost_q_values = jnp.array([initial_q_value, initial_q_value])
    initial_choice_traces = jnp.zeros(2)
    agent_state = jnp.concatenate(
        [initial_q_values, initial_ghost_q_values, initial_choice_traces])

  # Unpack the agent state vector into its components for easier access.
  q_values, ghost_q_values, choice_traces = jnp.split(agent_state, [2, 4])

  # --- 3. Update Q-value for the CHOSEN arm ---

  # Calculate the prediction error: the difference between the actual reward
  # and the expected reward (Q-value).
  prediction_error = reward - q_values[choice]

  # Select the learning rate based on the sign of the prediction error.
  # A different learning rate is used for positive (better than expected)
  # and negative (worse than expected) outcomes.
  learning_rate_for_chosen_arm = jnp.where(
      prediction_error >= 0,
      positive_base_learning_rate,
      negative_base_learning_rate,
  )

  # Calculate the standard reinforcement learning update term.
  standard_rl_update = learning_rate_for_chosen_arm * prediction_error

  # Calculate the influence of the "ghost" Q-value, which pulls the chosen
  # Q-value towards the ghost value.
  ghost_influence_on_chosen_q = chosen_q_ghost_influence_rate * (
      ghost_q_values[choice] - q_values[choice])

  # Update the Q-value for the chosen arm by combining the standard update
  # and the ghost influence.
  updated_q_value_chosen = (
      q_values[choice] + standard_rl_update + ghost_influence_on_chosen_q)
  q_values = q_values.at[choice].set(updated_q_value_chosen)

  # --- 4. Update Q-value for the UNCHOSEN arm ---

  # Identify the unchosen arm (if choice is 0, unchosen is 1, and vice-versa).
  unchosen_arm = 1 - choice

  # Calculate the influence of the ghost Q-value on the unchosen arm. This
  # term is modulated by the received reward.
  ghost_influence_on_unchosen_q = (
      unchosen_q_update_learning_rate * (1 + reward) *
      (ghost_q_values[unchosen_arm] - q_values[unchosen_arm]))

  # Calculate a "counterfactual perseveration" term. If the outcome was worse
  # than expected (negative prediction error), this pulls the unchosen Q-value
  # towards the chosen Q-value, potentially promoting perseveration.
  perseveration_term = unchosen_q_update_learning_rate * (
      q_values[choice] - q_values[unchosen_arm])
  counterfactual_perseveration = jnp.where(
      prediction_error < 0, perseveration_term, 0.0)

  # Update the Q-value for the unchosen arm.
  updated_q_value_unchosen = (q_values[unchosen_arm] +
                              ghost_influence_on_unchosen_q +
                              counterfactual_perseveration)
  q_values = q_values.at[unchosen_arm].set(updated_q_value_unchosen)

  # --- 5. Update Ghost Q-values ---

  # Define the target for the ghost Q-value update.
  # The ghost value for the CHOSEN arm targets the initial default Q-value.
  # The ghost value for the UNCHOSEN arm targets the reward that was actually
  # received from making the chosen action.
  ghost_q_values_target = jnp.array([initial_q_value, initial_q_value])
  ghost_q_values_target = ghost_q_values_target.at[unchosen_arm].set(reward)

  # Update the ghost Q-values using a simple exponential moving average toward
  # the target.
  ghost_q_update = ghost_learning_rate * (
      ghost_q_values_target - ghost_q_values)
  ghost_q_values = ghost_q_values + ghost_q_update

  # --- 6. Update Choice Traces ---

  # Apply a general decay to both choice traces.
  choice_traces = choice_traces * (1 - trace_learning_rate)

  # Apply an additional, reward-modulated decay to the UNCHOSEN arm's trace.
  # Note: This update is applied to the already-decayed trace from the previous step.
  reward_modulated_decay_factor = 1 - trace_learning_rate * (1 + reward)
  updated_unchosen_trace = choice_traces[unchosen_arm] * reward_modulated_decay_factor
  choice_traces = choice_traces.at[unchosen_arm].set(updated_unchosen_trace)

  # Strengthen the trace for the CHOSEN arm, reinforcing the memory of this choice.
  updated_chosen_trace = choice_traces[choice] + trace_learning_rate * 2
  choice_traces = choice_traces.at[choice].set(updated_chosen_trace)

  # --- 7. Re-assemble Agent State and Calculate Choice Logits ---

  # Combine the updated components back into a single state vector for the next trial.
  agent_state = jnp.concatenate([q_values, ghost_q_values, choice_traces])

  # Calculate the contribution of Q-values to the choice logits, weighted by
  # inverse temperatures. A higher inverse temperature leads to more
  # deterministic choices based on Q-values.
  q_value_component = inverse_temperatures * q_values

  # Add a constant side bias to the logits.
  bias_component = jnp.array([choice_bias, -choice_bias])

  # Calculate the contribution of the choice traces to the logits. The
  # influence of the trace is modulated by the choice bias and previous reward.
  trace_weight = jax.nn.sigmoid(choice_bias) * (1 + jax.nn.softplus(reward))
  trace_component = trace_weight * choice_traces

  # Combine all components to get the final choice logits.
  choice_logits = q_value_component + bias_component + trace_component

  # Return the calculated logits and the new agent state.
  return choice_logits, agent_state
```
