## Supplementary material for "AI-Discovered Cognitive Models Reveal Novel Insights into Human and Animal Learning": DataDIVER Discovered Programs: rat_twostep_models.py

```
# Copyright 2026 DeepMind Technologies Limited.
#
# Licensed under the Apache License, Version 2.0 (the "License");
# you may not use this file except in compliance with the License.
# You may obtain a copy of the License at
#
#     http://www.apache.org/licenses/LICENSE-2.0
#
# Unless required by applicable law or agreed to in writing, software
# distributed under the License is distributed on an "AS IS" BASIS,
# WITHOUT WARRANTIES OR CONDITIONS OF ANY KIND, either express or implied.
# See the License for the specific language governing permissions and
# limitations under the License.

"""Saved models for the Nature 2026 submission."""
from typing import Optional, Tuple

import chex
import jax
import jax.numpy as jnp


def rat_twostep_run1_fit_only(
    params: chex.Array,
    choice: int,
    reward: int,
    outcome: int,
    agent_state: Optional[chex.Array],
) -> tuple[chex.Array, chex.Array]:
  """Cognitive model describing rat behavior on a binary two-step task.

  Args:
    params: Model params. Different parameters are used for different rats.
    choice: Previous choice. Values: 0 or 1.
    reward: Previous reward. Values: 0 or 1.
    outcome: Previous outcome observed following the choice. Values: 0 or 1.
    agent_state: Previous state of the agent

  Returns:
    choice_logits: Vector of shape (2,) with the probabilities that the rat will
      choose option 0 or 1 on the next trial, expressed as logits.
    agent_state: New state of the agent, after observing the previous choice and
      reward.
  """
  # Do not remove comments or TODOs from this program.

  # TODO: Initialize agent state. NOLINT
  if agent_state is None:
    # Initialize Q-values for stage 1 actions (choices)
    q_s1 = jnp.zeros((2,))
    # Initialize Q-values for stage 2 actions (outcomes)
    # Initialized to an optimistic value to encourage early exploration.
    q_s2 = jnp.full((2,), 1.0)
    # Initialize transition strengths from stage 1 choices to stage 2 outcomes.
    # These strengths will be softmaxed to obtain probabilities.
    # Initialized to a small value (e.g., 0.01) to start with near-equal probabilities.
    transition_strengths = jnp.full((2, 2), 0.01)
    # Initialize reward consistency tracker for each stage 1 choice.
    # Higher values indicate more consistent rewards for that choice.
    # Initialize reward consistency tracker for each stage 1 choice.
    # Higher values indicate more consistent rewards for that choice.
    reward_consistency_tracker = jnp.full((2,), 0.5)
    agent_state = jnp.concatenate([q_s1, q_s2.flatten(), transition_strengths.flatten(), reward_consistency_tracker])

  # TODO: Define all parameters. NOLINT
  # Each parameter assignment should be a single line.
  # Change the names of the parameters to be descriptive.
  # Include comments describing the parameter transformation, if applicable.
  learning_rate_s1_base = jax.nn.sigmoid(params[0])
  learning_rate_s2 = jax.nn.softplus(params[1])
  learning_rate_transition_strength = jax.nn.softplus(params[2])
  inverse_temperature = jnp.exp(params[3])
  model_based_weight = jax.nn.sigmoid(params[4])
  learning_rate_s2_decay = jax.nn.softplus(params[5])
  stickiness_bias = jnp.tanh(params[6])
  outcome_perseveration_weight = jnp.tanh(params[7])
  reward_consistency_sensitivity = jnp.tanh(params[8])
  prior_choice_bias = jnp.tanh(params[9])

  # TODO: Update agent_state. NOLINT
  # Do not update more than one element at a time.
  # Each component of the agent state update should be a single line.
  # Include comments describing the computation in each line.
  #  agent_state[i] = ...

  # Unpack agent state
  q_s1 = agent_state[0:2]
  q_s2 = agent_state[2:4]
  transition_strengths = agent_state[4:8].reshape((2, 2))
  reward_consistency_tracker = agent_state[8:10]

  # Update reward consistency tracker for the chosen stage 1 action.
  prediction_error_s1_mf = reward - q_s1[choice]
  salience_modulation_consistency = jax.nn.sigmoid(jnp.abs(prediction_error_s1_mf))
  consistency_learning_rate_chosen = learning_rate_s1_base * salience_modulation_consistency
  reward_consistency_tracker = reward_consistency_tracker.at[choice].set(
      reward_consistency_tracker[choice] * (1 - consistency_learning_rate_chosen) + reward * consistency_learning_rate_chosen
  )

  # Update Q-value for the unchosen stage 2 outcome.
  # For the unchosen stage 2 outcome, its value decays towards zero.
  # The decay rate is modulated by how uncertain the agent is about this outcome.
  # Smaller Q-value (closer to initial uncertainty) leads to stronger decay.
  uncertainty_modulation_unchosen = jax.nn.sigmoid(1 - q_s2[1 - outcome]) # Higher when Q-value is low
  effective_learning_rate_s2_unchosen = learning_rate_s2_decay * uncertainty_modulation_unchosen
  q_s2 = q_s2.at[1 - outcome].set(q_s2[1 - outcome] * (1 - effective_learning_rate_s2_unchosen))
  # The unchosen stage 1 action's Q-value also decays, but is modulated by a different base rate.
  q_s1 = q_s1.at[1 - choice].set(q_s1[1 - choice] * (1 - learning_rate_s1_base))

  effective_reward_s2 = reward + outcome_perseveration_weight
  prediction_error_s2 = effective_reward_s2 - q_s2[outcome]
  # Modulate learning rate by the absolute prediction error, reflecting surprise.
  salience_surprise_s2 = jax.nn.sigmoid(jnp.abs(prediction_error_s2))
  effective_learning_rate_s2 = learning_rate_s2 * salience_surprise_s2
  q_s2 = q_s2.at[outcome].add(effective_learning_rate_s2 * prediction_error_s2)

  # Update transition strengths.
  p_s1_to_s2 = jax.nn.softmax(transition_strengths, axis=1)

  prediction_error_s1_mf = reward - q_s1[choice]
  expected_value_s2_for_chosen_s1 = jnp.sum(p_s1_to_s2[choice, :] * q_s2)
  prediction_error_s1_mb = reward - expected_value_s2_for_chosen_s1

  # Modulate transition learning rate by the absolute model-based prediction error.
  salience_transition = jax.nn.sigmoid(jnp.abs(prediction_error_s1_mb))
  effective_learning_rate_transition = learning_rate_transition_strength * salience_transition
  transition_strengths_update = jnp.zeros_like(transition_strengths).at[choice, outcome].set(
      effective_learning_rate_transition
  ).at[choice, 1 - outcome].set(
      -effective_learning_rate_transition
  ).at[1 - choice, outcome].set(
      -effective_learning_rate_transition
  ).at[1 - choice, 1 - outcome].set(
      effective_learning_rate_transition
  )
  transition_strengths = transition_strengths + transition_strengths_update

  consistency_modulation = jax.nn.sigmoid(reward_consistency_sensitivity * (reward_consistency_tracker[choice] - 0.5) * 4)
  consistency_modulation_scaled = (consistency_modulation - 0.5) * 2
  positive_learning_rate_s1 = learning_rate_s1_base * (1 + consistency_modulation_scaled)
  negative_learning_rate_s1 = learning_rate_s1_base * (1 - consistency_modulation_scaled)

  consistency_decay_rate = 0.02
  reward_consistency_tracker = reward_consistency_tracker.at[1 - choice].set(
      reward_consistency_tracker[1 - choice] * (1 - consistency_decay_rate) + 0.5 * consistency_decay_rate
  )
  unchosen_consistency_learning_rate = learning_rate_s1_base * (1 + reward_consistency_sensitivity * (0.5 - reward_consistency_tracker[1 - choice]))
  expected_value_s2_for_unchosen = jnp.sum(p_s1_to_s2[1 - choice, :] * q_s2)
  counterfactual_prediction_error_consistency = expected_value_s2_for_unchosen - q_s1[1 - choice]
  counterfactual_update_strength = unchosen_consistency_learning_rate * model_based_weight * jax.nn.sigmoid(jnp.abs(prediction_error_s1_mf))
  reward_consistency_tracker = reward_consistency_tracker.at[1 - choice].add(
      counterfactual_update_strength * counterfactual_prediction_error_consistency
  )
  reward_consistency_tracker = jnp.clip(reward_consistency_tracker, 0.0, 1.0)

  # Modulate learning rates based on consistency, and then apply RPE-driven salience.
  consistency_scaling_factor = (1 + reward_consistency_sensitivity * (reward_consistency_tracker[choice] - 0.5) * 2)

  # For MF RPE:
  # Modulate the prediction error by consistency: consistent rewards make errors more salient.
  mf_modulated_prediction_error = prediction_error_s1_mf * consistency_scaling_factor
  mf_effective_learning_rate = jnp.where(
      mf_modulated_prediction_error >= 0,
      positive_learning_rate_s1 * jax.nn.sigmoid(mf_modulated_prediction_error), # Stronger learning for positive surprises
      negative_learning_rate_s1 * jax.nn.sigmoid(-mf_modulated_prediction_error)  # Stronger learning for negative surprises
  )

  # For MB RPE:
  # Modulate the prediction error by consistency: consistent rewards make errors more salient.
  mb_modulated_prediction_error = prediction_error_s1_mb * consistency_scaling_factor
  mb_effective_learning_rate = jnp.where(
      mb_modulated_prediction_error >= 0,
      positive_learning_rate_s1 * jax.nn.sigmoid(mb_modulated_prediction_error), # Stronger learning for positive surprises
      negative_learning_rate_s1 * jax.nn.sigmoid(-mb_modulated_prediction_error)  # Stronger learning for negative surprises
  )

  # Combine MF and MB updates with their respective effective learning rates and modulated prediction errors.
  q_s1_update_amount = (1 - model_based_weight) * (mf_effective_learning_rate * mf_modulated_prediction_error) + \
                       model_based_weight * (mb_effective_learning_rate * mb_modulated_prediction_error)
  q_s1 = q_s1.at[choice].add(q_s1_update_amount)

  # Repack agent state.
  agent_state = jnp.concatenate([q_s1, q_s2, transition_strengths.flatten(), reward_consistency_tracker])

  # Compute choice_logits.


  value_mf = q_s1
  value_mb = jnp.sum(p_s1_to_s2 * q_s2, axis=1)

  combined_value = (1 - model_based_weight) * value_mf + model_based_weight * value_mb
  choice_logits = inverse_temperature * combined_value
  choice_logits = choice_logits.at[choice].add(stickiness_bias)
  choice_logits = choice_logits.at[0].add(prior_choice_bias)
  choice_logits = choice_logits.at[1].add(-prior_choice_bias)

  return choice_logits, agent_state


def rat_twostep_run2_fit_only(
    params: chex.Array,
    choice: int,
    reward: int,
    outcome: int,
    agent_state: Optional[chex.Array],
) -> tuple[chex.Array, chex.Array]:
  """Cognitive model describing rat behavior on a binary two-step task.

  Args:
    params: Model params. Different parameters are used for different rats.
    choice: Previous choice. Values: 0 or 1.
    reward: Previous reward. Values: 0 or 1.
    outcome: Previous outcome observed following the choice. Values: 0 or 1.
    agent_state: Previous state of the agent

  Returns:
    choice_logits: Vector of shape (2,) with the probabilities that the rat will
      choose option 0 or 1 on the next trial, expressed as logits.
    agent_state: New state of the agent, after observing the previous choice and
      reward.
  """
  # Do not remove comments or TODOs from this program.

  # Initialize agent state.
  if agent_state is None:
    # Initialize Q-values for two choices and outcome values for two outcomes.
    agent_state = jnp.zeros((2, 2)) # agent_state[0, :] are Q-values, agent_state[1, :] are outcome_values

  # Define all parameters.
  # Define all parameters.
  learning_rate = jnp.exp(params[0])  # Learning rate for Q-value updates.
  inverse_temperature = jnp.exp(params[1])  # Inverse temperature for softmax choice.
  retention_rate = jax.nn.sigmoid(params[2])  # Retention rate for Q-values and outcome values (decay).
  outcome_learning_rates = jnp.exp(params[3:5])
  outcome_weight = params[5]
  stickiness_bias = params[6]
  outcome_stickiness_bias = params[7]
  outcome_sensitivity = jax.nn.sigmoid(params[8:10])

  q_values, outcome_values = agent_state  # Separate Q-values and outcome_values from the agent state.

  q_chosen = q_values[choice]  # Get Q-value of the chosen option.
  q_td_error = reward - q_chosen  # Calculate TD error for Q-value update.

  # Introduce a state-dependent learning rate for Q-values.
  state_dependent_learning_rate = learning_rate * jax.nn.sigmoid(jnp.square(q_td_error))

  outcome_values = outcome_values * retention_rate

  dynamic_q_retention = retention_rate + (1 - retention_rate) * jnp.abs(q_td_error)
  q_values = q_values * dynamic_q_retention
  q_values_chosen_update = q_chosen + state_dependent_learning_rate * (
      jnp.abs(q_td_error) * q_td_error
      + 0.05 # Add a small recency bonus to the chosen Q-value update.
      # Surprise-modulated influence from the unchosen outcome value, reflecting a cognitive shift when expectations are violated.
      + outcome_sensitivity[1 - outcome] * (outcome_values[1 - outcome] - q_chosen) * (1 - jax.nn.sigmoid(jnp.abs(q_td_error)))
      + (1 - jax.nn.sigmoid(jnp.abs(q_td_error))) * (outcome_values[outcome] - q_chosen)
      + outcome_sensitivity[1] * outcome_sensitivity[outcome] * (outcome_values[1 - outcome] - q_chosen)
  )
  q_values = q_values.at[choice].set(q_values_chosen_update)
  # Dynamically adjust the recency decay rate for the unchosen option, increasing it with higher surprise.
  dynamic_recency_decay_rate = 0.01 * (1 + jnp.abs(q_td_error))
  # Update the unchosen Q-value with a surprise-modulated influence from the chosen Q-value and a penalty for no reward.
  # This makes the unchosen option's update more sensitive when the previous trial's outcome was surprising.
  # Introduce an outcome certainty factor: higher surprise (abs(q_td_error)) means lower certainty, leading to a stronger update for the unchosen option.
  outcome_certainty_factor = 1.0 + jnp.exp(jnp.abs(q_td_error) - 1.0) # Sigmoidal-like effect: higher TD error, higher factor
  q_values_unchosen_update = q_values[1 - choice] * (1 - dynamic_recency_decay_rate) + learning_rate * (
      outcome_sensitivity[outcome] * (outcome_values[outcome] - q_values[1 - choice])
      + 0.5 * (q_values[choice] - q_values[1 - choice]) * outcome_certainty_factor # Surprise-modulated influence from chosen Q
      - (1 - reward) * 0.1 * q_values[1 - choice] * outcome_certainty_factor # Surprise-modulated penalty for no reward
  )
  q_values = q_values.at[1 - choice].set(q_values_unchosen_update)
  relative_q_surprise = (q_values[choice] - q_values[1 - choice]) * outcome_sensitivity[outcome]
  outcome_td_error = reward + q_values[choice] - outcome_values[outcome] + relative_q_surprise
  outcome_values_observed_update = outcome_values[outcome] + outcome_learning_rates[outcome] * outcome_td_error
  outcome_values_observed_update += learning_rate * (q_values[choice] - outcome_values[outcome])
  outcome_values = outcome_values.at[outcome].set(outcome_values_observed_update)
  # For the unobserved outcome, update its value based on the *absence* of reward, modulated by its own learning rate.
  # This reflects counterfactual learning, where the lack of an outcome provides an informative prediction error.
  outcome_td_error_unobserved = (1 - reward) - outcome_values[1 - outcome]
  outcome_values_unobserved_update = outcome_values[1 - outcome] * retention_rate + outcome_learning_rates[1 - outcome] * outcome_td_error_unobserved
  outcome_values = outcome_values.at[1 - outcome].set(outcome_values_unobserved_update)

  agent_state = jnp.stack([q_values, outcome_values])

  contextual_outcome_weight = outcome_weight * (1 + jnp.abs(q_values))
  choice_logits = inverse_temperature * (q_values + contextual_outcome_weight * outcome_values * outcome_sensitivity)
  choice_logits = choice_logits.at[choice].add(stickiness_bias * jnp.abs(q_values[choice]) * (1 + jnp.abs(q_td_error)))
  choice_logits = choice_logits.at[outcome].add(outcome_stickiness_bias * jnp.abs(outcome_values[outcome]) * (1 + jnp.abs(q_td_error)))
  exploration_bonus = params[9]
  choice_logits = choice_logits.at[1 - choice].add(exploration_bonus * jnp.abs(q_td_error) * (1 - reward))

  return choice_logits, agent_state


def rat_twostep_run3_fit_only(
    params: chex.Array,
    choice: int,
    reward: int,
    outcome: int,
    agent_state: Optional[chex.Array],
) -> tuple[chex.Array, chex.Array]:
  """Cognitive model describing rat behavior on a binary two-step task.

  Args:
    params: Model params. Different parameters are used for different rats.
    choice: Previous choice. Values: 0 or 1.
    reward: Previous reward. Values: 0 or 1.
    outcome: Previous outcome observed following the choice. Values: 0 or 1.
    agent_state: Previous state of the agent

  Returns:
    choice_logits: Vector of shape (2,) with the probabilities that the rat will
      choose option 0 or 1 on the next trial, expressed as logits.
    agent_state: New state of the agent, after observing the previous choice and
      reward.
  """
  # Do not remove comments or TODOs from this program.

  # TODO: Initialize agent state. NOLINT
  if agent_state is None:
    # Initialize Q-values for choices (2), Q-values for outcomes (2), stickiness bonus for each choice (2), current inverse temperature (1), eligibility traces for choices (2), and eligibility traces for outcomes (2).
    agent_state = jnp.zeros(11)

  # TODO: Define all parameters. NOLINT
  # Each parameter assignment should be a single line.
  # Learning rate for choice-reward associations.
  learning_rate_choice = jax.nn.softplus(params[0])
  # Learning rate for outcome-reward associations.
  learning_rate_outcome = jax.nn.softplus(params[1])
  # Weight for outcome Q-values in the combined Q-value calculation.
  outcome_weight = params[2]
  # Baseline inverse temperature for softmax choice rule.
  # This serves as the initial value and long-term target for the dynamic inverse temperature.
  inverse_temperature_baseline = jnp.exp(params[3])
  # Decay rate for choice Q-values.
  choice_decay_rate = jax.nn.sigmoid(params[4])
  # Decay rate for outcome Q-values.
  outcome_decay_rate = jax.nn.sigmoid(params[5])
  # Decay rate for stickiness.
  stickiness_decay_rate = jax.nn.sigmoid(params[6])
  # Learning rate for the dynamic inverse temperature.
  # This determines how quickly the inverse temperature adapts to reward fluctuations.
  inverse_temperature_learning_rate = jax.nn.sigmoid(params[7])
  # Baseline Q-value for the outcome associated with a choice.
  outcome_q_baseline = jax.nn.sigmoid(params[8])
  # Decay rate for eligibility trace.
  lambda_eligibility_trace = jax.nn.sigmoid(params[9])
  # Magnitude of bias for choice 0. Positive value favors choice 0.
  # This parameter is now at index 9.
  choice_bias_magnitude = params[9]
  # TODO: Update agent_state. NOLINT
  # Do not update more than one element at a time.
  # Each component of the agent state update should be a single line.
  # Include comments describing the computation in each line.
  # Extract Q-values for choices, outcomes, stickiness, current inverse temperature, and eligibility traces from the agent state.
  q_values_choices = agent_state[:2]
  q_values_outcomes = agent_state[2:4]
  stickiness_values = agent_state[4:6]
  current_inverse_temperature = agent_state[6]
  eligibility_traces_choices = agent_state[7:9] # Eligibility traces for choices
  eligibility_traces_outcomes = agent_state[9:11] # Eligibility traces for outcomes

  # Update eligibility traces for choices: the chosen action's trace is set to 1, unchosen decay.
  other_choice = 1 - choice
  eligibility_traces_choices = eligibility_traces_choices.at[choice].set(1.0)
  eligibility_traces_choices = eligibility_traces_choices.at[other_choice].set(eligibility_traces_choices[other_choice] * lambda_eligibility_trace)
  
  # Compute prediction error for choices.
  prediction_error_choice = reward - q_values_choices[choice]

  # Update eligibility traces for outcomes: the chosen outcome's trace is set to 1, unchosen decay.
  other_outcome = 1 - outcome
  eligibility_traces_outcomes = eligibility_traces_outcomes.at[outcome].set(1.0)
  # Modulate the eligibility trace decay for outcomes by the absolute magnitude of the choice prediction error.
  # A larger choice prediction error reduces the effective decay rate, making the outcome eligibility trace persist longer.
  # This allows for more sustained learning from outcomes when initial choices yield surprising results.
  # A small constant (1.0) is added to the denominator to prevent division by zero and ensure the scaling factor is well-behaved.
  modulated_lambda_eligibility_trace_outcome = lambda_eligibility_trace / (1.0 + jnp.abs(prediction_error_choice))
  eligibility_traces_outcomes = eligibility_traces_outcomes.at[other_outcome].set(eligibility_traces_outcomes[other_outcome] * modulated_lambda_eligibility_trace_outcome)

  # Update Q-values for choices based on prediction error, with an adaptive learning rate and eligibility traces.
  # The learning rate is adaptively modulated by the absolute prediction error and inversely scaled by the absolute Q-value.
  # An uncertainty bonus further boosts learning for larger prediction errors.
  # The learning rate is adaptively modulated by the square of the absolute prediction error and inversely scaled by the absolute Q-value.
  # A small constant (1.0) is added to the denominator to prevent division by zero.
  state_dependent_learning_rate = learning_rate_choice * jnp.square(jnp.abs(prediction_error_choice)) / (1.0 + jnp.abs(q_values_choices[choice]))
  q_values_choices = q_values_choices + state_dependent_learning_rate * prediction_error_choice * eligibility_traces_choices
  
  # Unchosen Q-value for choices decays towards a blend of its initial value (0) and the updated chosen Q-value.
  # This provides a mild vicarious learning effect for the unchosen option.
  decay_target_choice = 0.5 * q_values_choices[choice] # Changed to remove the redundant `+ 0.5 * 0.0`
  q_values_choices = q_values_choices.at[other_choice].set(
      q_values_choices[other_choice] * (1 - choice_decay_rate) + decay_target_choice * choice_decay_rate
  )

  # Compute prediction error for outcomes.
  prediction_error_outcome = reward - q_values_outcomes[outcome]
  # Update Q-values for outcomes based on prediction error, scaled by the eligibility traces for outcomes.
  # This allows credit to propagate back to previously experienced outcomes.
  q_values_outcomes = q_values_outcomes + learning_rate_outcome * prediction_error_outcome * eligibility_traces_outcomes
  # Unchosen Q-value for outcomes decays towards a learned baseline.
  # This makes the outcome Q-value update more robust by anchoring it to a learnable baseline.
  q_values_outcomes = q_values_outcomes.at[other_outcome].set(
      q_values_outcomes[other_outcome] * (1 - outcome_decay_rate) + outcome_q_baseline * outcome_decay_rate
  )

  # Update stickiness for the chosen action, increasing its value. Decay is modulated by reward and prediction error.
  # A higher reward reduces the effective decay rate, making stickiness persist longer for rewarded actions.
  # A larger absolute prediction error also increases the decay rate, making the agent less sticky when outcomes are surprising.
  prediction_error_outcome_magnitude = jnp.abs(prediction_error_outcome)
  # A higher reward reduces the effective decay rate, making stickiness persist longer for rewarded actions.
  # A larger absolute prediction error also increases the decay rate, making the agent less sticky when outcomes are surprising.
  reward_and_prediction_error_modulated_decay_rate = stickiness_decay_rate * (1 + prediction_error_outcome_magnitude) * (1 - reward)
  stickiness_values = stickiness_values.at[choice].set(
      stickiness_values[choice] * (1 - reward_and_prediction_error_modulated_decay_rate) + 1.0 * reward_and_prediction_error_modulated_decay_rate
  )
  # For the unchosen action, stickiness decays towards a negative value, promoting switching.
  # This makes the unchosen option more attractive following an undesirable outcome of the chosen action.
  stickiness_values = stickiness_values.at[other_choice].set(
      stickiness_values[other_choice] * (1 - stickiness_decay_rate) - (1 - reward) * stickiness_decay_rate
  )

  # Update the current inverse temperature.
  # It adjusts towards the baseline, and is also influenced by prediction errors (positive errors increase exploitation, negative decrease).
  # The learning rate for inverse temperature is dynamically modulated by the magnitude of prediction errors and the stability of Q-values.
  # Larger prediction errors and less stable Q-values (indicating higher uncertainty) will lead to more exploration (lower inverse temperature).
  total_prediction_error_magnitude = jnp.abs(prediction_error_choice) + jnp.abs(prediction_error_outcome)
  
  # Calculate Q-value stability (inverse of recent update magnitude) for the chosen action.
  # A larger update implies lower stability, hence lower confidence.
  q_update_magnitude = jnp.abs(state_dependent_learning_rate * prediction_error_choice * eligibility_traces_choices[choice])
  # Ensure stability factor is well-behaved, preventing division by zero and extreme values.
  q_stability_factor = 1.0 / (1.0 + q_update_magnitude) # Higher stability when updates are small.

  # Dynamically modulate the inverse temperature learning rate: larger prediction errors and lower Q-stability lead to slower adaptation to baseline.
  # This makes the inverse temperature more stable (and promotes exploration) when the environment is uncertain or Q-values are unstable.
  dynamic_inverse_temperature_learning_rate = inverse_temperature_learning_rate * jnp.exp(-total_prediction_error_magnitude) * q_stability_factor
  
  # Inverse temperature change is influenced by prediction errors. Positive errors increase exploitation, negative decrease.
  inverse_temperature_change = (prediction_error_choice + jnp.abs(prediction_error_outcome)) * dynamic_inverse_temperature_learning_rate
  
  current_inverse_temperature = current_inverse_temperature * (1 - dynamic_inverse_temperature_learning_rate) + \
                                inverse_temperature_baseline * dynamic_inverse_temperature_learning_rate + \
                                inverse_temperature_change
  # Ensure inverse temperature remains positive by clipping.
  current_inverse_temperature = jnp.maximum(current_inverse_temperature, 0.1)

  # Reconstruct the full agent state with updated Q-values, stickiness, current inverse temperature, and eligibility traces for choices and outcomes.
  agent_state = jnp.concatenate((q_values_choices, q_values_outcomes, stickiness_values, jnp.array([current_inverse_temperature]), eligibility_traces_choices, eligibility_traces_outcomes))

  # TODO: Compute choice_logits. NOLINT
  # Each component of the choice logit computation should be a single line.
  # Include comments describing the computation in each line.
  # Combine Q-values from choices and outcomes, weighted by outcome_weight, to compute overall action values.
  combined_q_values = (1 - outcome_weight) * q_values_choices + outcome_weight * q_values_outcomes

  # Compute logits for the softmax choice rule using the dynamic inverse temperature.
  choice_logits = combined_q_values * current_inverse_temperature
  # Add stickiness values as a bonus to the logits.
  choice_logits = choice_logits + stickiness_values
  # Apply choice bias: positive choice_bias_magnitude favors choice 0, negative favors choice 1.
  choice_logits = choice_logits + jnp.array([choice_bias_magnitude, -choice_bias_magnitude])

  return choice_logits, agent_state


# Shortened to 100k steps then refactored, run 1, low floor.
def rat_twostep_run1_simplified_low_floor(
    params: chex.Array,
    choice: int,
    reward: int,
    outcome: int,
    agent_state: Optional[chex.Array],
) -> tuple[chex.Array, chex.Array]:
  """
  Updates agent's beliefs (Q-values and V-values) based on an action and its outcome.

  This agent uses a reinforcement learning model where:
  - Q-values represent the expected reward for taking a specific action (choice).
  - V-values represent the expected reward associated with a specific outcome state.
  
  The agent's decision-making is based on the difference between these Q and V values.

  Args:
    params: An array containing the agent's fixed parameters (learning rates, etc.).
    choice: The action taken by the agent (e.g., 0 or 1).
    reward: The scalar reward received.
    outcome: The resulting state or outcome observed (e.g., 0 or 1).
    agent_state: The agent's internal state (Q and V values) from the previous step.
                 It is None on the very first step of a sequence.

  Returns:
    A tuple containing:
      - action_logits: The computed values (logits) used for making the next decision.
      - new_agent_state: The updated internal state to be used in the next step.
  """

  # --------------------------------------------------------------------------
  # 1. Initialize or retrieve the agent's internal state
  # --------------------------------------------------------------------------

  # If this is the first step (agent_state is None), initialize Q-values and V-values to zeros.
  # Otherwise, unpack the existing Q-values and V-values from the provided agent_state.
  # Both q_values and v_values are arrays of shape (2,).
  if agent_state is None:
    q_values, v_values = jnp.zeros((2, 2))
  else:
    q_values, v_values = agent_state

  # --------------------------------------------------------------------------
  # 2. Unpack the agent's fixed parameters for clarity
  # --------------------------------------------------------------------------

  # Learning rates for the two possible choices (actions)
  q_learning_rate_for_choice_0 = params[0]
  q_learning_rate_for_choice_1 = params[1]
  
  # Softmax inverse temperature (beta), controls exploration vs. exploitation
  inverse_temperature = params[2]
  
  # Learning rate for the outcome values (V-values)
  v_learning_rate = params[3]
  
  # Rate at which Q-values decay towards a baseline
  q_decay_rate = params[4]
  baseline_q_value = params[5]
  
  # Rate at which V-values decay towards a baseline
  v_decay_rate = params[6]
  baseline_v_value = params[7]

  # --------------------------------------------------------------------------
  # 3. Update Q-values (Action Values)
  # --------------------------------------------------------------------------

  # Step 3a: Apply a decay to all Q-values, pulling them toward a baseline value.
  # The sigmoid function scales the decay rate to be between 0 and 1.
  q_decay_factor = jax.nn.sigmoid(q_decay_rate)
  q_values_after_decay = q_values + q_decay_factor * (baseline_q_value - q_values)

  # Step 3b: Update the Q-value for the action that was actually chosen.
  # This is a standard reinforcement learning update based on prediction error.
  q_prediction_error = reward - q_values_after_decay[choice]
  
  # Select the learning rate specific to the choice that was made.
  q_learning_rates = jnp.array([q_learning_rate_for_choice_0, q_learning_rate_for_choice_1])
  active_q_learning_rate = q_learning_rates[choice]
  
  # Apply the learning update to the Q-value of the chosen action.
  q_value_update = active_q_learning_rate * q_prediction_error
  updated_q_values = q_values_after_decay.at[choice].add(q_value_update)

  # --------------------------------------------------------------------------
  # 4. Update V-values (Outcome Values)
  # --------------------------------------------------------------------------

  # Step 4a: Apply a decay to all V-values, pulling them toward a baseline value.
  v_values_after_decay = v_values + v_decay_rate * (baseline_v_value - v_values)

  # Step 4b: Update the V-value for the outcome that was actually observed.
  # This learns the intrinsic value of arriving in a particular outcome state.
  v_prediction_error = reward - v_values_after_decay[outcome]
  
  # Apply the learning update to the V-value of the observed outcome.
  v_value_update = v_learning_rate * v_prediction_error
  updated_v_values = v_values_after_decay.at[outcome].add(v_value_update)

  # --------------------------------------------------------------------------
  # 5. Compute action logits for the next decision
  # --------------------------------------------------------------------------
  
  # The agent's preference for an action is based on the difference between its
  # action-value (Q) and its outcome-value (V). This is an "advantage" term.
  advantage = updated_q_values - updated_v_values

  # Scale the advantage by the inverse temperature to produce the final logits.
  # Higher inverse temperature leads to more deterministic (exploitative) choices.
  action_logits = inverse_temperature * advantage

  # --------------------------------------------------------------------------
  # 6. Package the new state for the next step
  # --------------------------------------------------------------------------

  # Combine the updated Q-values and V-values into a single state array.
  new_agent_state = jnp.stack((updated_q_values, updated_v_values))

  return action_logits, new_agent_state


# Shortened to 100k steps then refactored, run 2, low floor.
def rat_twostep_run2_simplified_low_floor(
    params: chex.Array,
    choice: int,
    reward: int,
    outcome: int,
    agent_state: Optional[chex.Array],
) -> tuple[chex.Array, chex.Array]:
  learning_rate, inverse_temperature, retention_rate, *_unused_params, outcome_weight, stickiness, outcome_stickiness = params
  q_values, outcome_values = (jnp.zeros((2, 2)) if agent_state is None else agent_state)

  q_values = q_values.at[1 - choice].multiply(retention_rate).at[choice].add(
      learning_rate * (reward - q_values[choice])
  )

  outcome_values = outcome_values * params[5:7]
  outcome_values = outcome_values.at[outcome].add(
      params[3 + outcome] * (reward - outcome_values[outcome])
  )

  choice_logits = (
      inverse_temperature * (q_values + outcome_weight * outcome_values)
      .at[choice].add(stickiness)
      .at[outcome].add(outcome_stickiness)
  )

  return choice_logits, jnp.stack([q_values, outcome_values])
  # EVOLVE-BLOCK-END


# Shortened to 100k steps then refactored, run 3, low floor.
def rat_twostep_run3_simplified_low_floor(
    params: chex.Array,
    choice: int,
    reward: int,
    outcome: int,
    agent_state: Optional[chex.Array],
) -> tuple[chex.Array, chex.Array]:
  """
  Computes the agent's next action probabilities and updates its internal state.

  This agent uses a Q-learning variant to update two separate sets of values:
  1. Q-values associated with choices (e.g., left vs. right).
  2. Q-values associated with outcomes (e.g., reward vs. no reward).
  The final action selection is based on a combination of these two value sets.

  Args:
      params: An array of the agent's fixed parameters, such as learning rates,
              decay rates, and temperature.
      choice: The action taken by the agent in the previous step (e.g., 0 or 1).
      reward: The reward received for the previous action.
      outcome: The outcome observed after the previous action (e.g., 0 or 1).
      agent_state: The internal state (Q-values) of the agent from the previous
                   step. It is None on the first step.

  Returns:
      A tuple containing:
      - choice_logits: The logits (unnormalized log-probabilities) for the next
                       action selection.
      - new_agent_state: The updated internal state of the agent for the next step.
  """

  # --- 1. Initialize Agent State ---
  
  # If this is the first trial (agent_state is None), initialize a new state.
  # The state is a vector of four Q-values:
  # [Q(choice=0), Q(choice=1), Q(outcome=0), Q(outcome=1)]
  if agent_state is None:
    agent_state = jnp.zeros(4)

  # --- 2. Unpack Parameters ---

  # Assign descriptive names to the elements of the `params` array for clarity.
  learning_rate_choice  = params[0] # Rate for updating choice Q-values.
  learning_rate_outcome = params[1] # Rate for updating outcome Q-values.
  outcome_weight        = params[2] # NOTE: This parameter is unpacked but not used in the logic.
  inverse_temperature   = params[3] # Controls the greediness of the policy.
  choice_decay_rate     = params[4] # Rate at which choice Q-values decay to a baseline.
  outcome_decay_rate    = params[5] # Rate at which outcome Q-values decay to a baseline.
  outcome_q_baseline    = params[6] # Baseline value for outcome Q-values decay.
  choice_q_baseline     = params[7] # Baseline value for choice Q-values decay.
  choice_bias           = params[8] # A constant bias added to the logit of the first choice.

  # --- 3. Define the Q-Value Update Rule ---

  # This helper function encapsulates the learning and decay logic.
  def _update_q_values(
      q_values: chex.Array,
      index_to_update: int,
      learning_rate: float,
      decay_rate: float,
      baseline: float,
  ) -> chex.Array:
    """Updates a set of Q-values based on a decay and a reward prediction error."""
    
    # First, apply a decay to all Q-values in the set.
    # They are pulled towards a baseline value, which can model forgetting.
    decayed_q_values = q_values - decay_rate * (q_values - baseline)

    # Next, calculate the prediction error for the specific choice or outcome.
    # This is the difference between the actual reward and the expected reward
    # (the current, decayed Q-value).
    prediction_error = reward - decayed_q_values[index_to_update]

    # Calculate the amount to update the Q-value by.
    update_amount = learning_rate * prediction_error
    
    # Add the update amount to the Q-value for the specific index that was chosen or observed.
    # This is a pure function; it returns a new array with the updated value.
    updated_q_values = decayed_q_values.at[index_to_update].add(update_amount)
    
    return updated_q_values

  # --- 4. Update Q-Values for the Previous Choice ---
  
  # Separate the Q-values for choices from the full agent state.
  q_values_for_choices = agent_state[:2]
  
  # Apply the update rule to the Q-values associated with the agent's `choice`.
  q_values_choices_updated = _update_q_values(
      q_values=q_values_for_choices,
      index_to_update=choice,
      learning_rate=learning_rate_choice,
      decay_rate=choice_decay_rate,
      baseline=choice_q_baseline
  )

  # --- 5. Update Q-Values for the Observed Outcome ---
  
  # Separate the Q-values for outcomes from the full agent state.
  q_values_for_outcomes = agent_state[2:]

  # Apply the update rule to the Q-values associated with the observed `outcome`.
  q_values_outcomes_updated = _update_q_values(
      q_values=q_values_for_outcomes,
      index_to_update=outcome,
      learning_rate=learning_rate_outcome,
      decay_rate=outcome_decay_rate,
      baseline=outcome_q_baseline
  )

  # --- 6. Calculate Action Logits for the Next Decision ---
  
  # The value of an action is the sum of its choice-value and its outcome-value.
  # This assumes a 1-to-1 mapping (e.g., choice 0 is associated with outcome 0).
  combined_q_values = q_values_choices_updated + q_values_outcomes_updated

  # Scale the combined values by the inverse temperature. A higher value leads
  # to a more deterministic (less random) choice.
  scaled_q_values = combined_q_values * inverse_temperature

  # Add a static bias to the first choice's logit. This models a default
  # preference for one action over the other, regardless of learned values.
  choice_logits = scaled_q_values.at[0].add(choice_bias)

  # --- 7. Create the New Agent State ---

  # Re-concatenate the updated Q-value arrays to form the new agent state.
  new_agent_state = jnp.concatenate(
      [q_values_choices_updated, q_values_outcomes_updated]
  )
  
  # --- 8. Return Results ---

  # Return the calculated logits and the new state for the next step.
  return choice_logits, new_agent_state


# Shortened to 100k steps then refactored, run 1, medium floor.
def rat_twostep_run1_simplified_medium_floor(
    params: chex.Array,
    choice: int,
    reward: int,
    outcome: int,
    agent_state: Optional[chex.Array],
) -> tuple[chex.Array, chex.Array]:
  """Cognitive model describing rat behavior on a binary two-step task.

  This model learns in several ways:
  1.  Action Values (Q-values): Learns the value of taking each action.
  2.  Outcome Values: Learns the value of arriving at each outcome.
  3.  Pavlovian-Instrumental Transfer (PIT): Learns an association between
      actions and outcomes, allowing outcome values to influence action choice.
  4.  Perseveration: Includes biases to repeat the previous action or the
      action associated with the previous outcome.

  Args:
    params: Model params. Different parameters are used for different rats.
    choice: Previous choice. Values: 0 or 1.
    reward: Previous reward. Values: 0 or 1.
    outcome: Previous outcome observed following the choice. Values: 0 or 1.
    agent_state: Previous state of the agent. A vector containing the agent's
      internal learned values.

  Returns:
    choice_logits: Vector of shape (2,) with the probabilities that the rat will
      choose option 0 or 1 on the next trial, expressed as logits.
    agent_state: New state of the agent, after observing the previous choice and
      reward.
  """
  # --- 1. Unpack and Transform Model Parameters ---

  # Unpack all raw (pre-transformation) parameters from the input array.
  (
      initial_q_value,
      learning_rate_action_raw,
      learning_rate_outcome_raw,
      softmax_inverse_temperature_raw,
      outcome_to_action_learning_rate_raw,
      decay_rate_action_raw,
      outcome_association_strength,
      action_perseveration,
      outcome_perseveration_0_raw,
      outcome_perseveration_1_raw,
  ) = params

  # Ensure certain parameters (learning rates, temperature) are positive
  # by applying the softplus function. This is a standard way to handle
  # constrained parameters during optimization.
  learning_rate_action = jax.nn.softplus(learning_rate_action_raw)
  learning_rate_outcome = jax.nn.softplus(learning_rate_outcome_raw)
  softmax_inverse_temperature = jax.nn.softplus(softmax_inverse_temperature_raw)
  outcome_to_action_learning_rate = jax.nn.softplus(outcome_to_action_learning_rate_raw)
  decay_rate_action = jax.nn.softplus(decay_rate_action_raw)

  # Combine the two outcome perseveration parameters into a single array.
  outcome_perseveration = jnp.array([outcome_perseveration_0_raw, outcome_perseveration_1_raw])


  # --- 2. Initialize or Retrieve Agent State ---

  # If this is the first trial (agent_state is None), initialize a new state.
  # The state vector tracks the following learned values:
  # agent_state[0]: Q-value of action 0
  # agent_state[1]: Q-value of action 1
  # agent_state[2]: Value of outcome 0
  # agent_state[3]: Value of outcome 1
  # agent_state[4]: Learned association weight between actions and outcomes (PIT)
  if agent_state is None:
    agent_state = jnp.zeros(5)
  

  # --- 3. Update Agent State Based on Previous Trial's Experience ---

  # Calculate the prediction error for the chosen action.
  # This is the difference between the reward received and the expected value (Q-value).
  prediction_error_action = reward - agent_state[choice]

  # Update the Q-value for the CHOSEN action using the prediction error.
  # This is a standard Rescorla-Wagner update rule.
  q_value_update_chosen_action = learning_rate_action * prediction_error_action
  
  # Update the Q-value for the UNCHOSEN action by decaying it towards the initial value.
  # This represents forgetting or a belief that unchosen options revert to a mean value.
  q_value_decay_unchosen_action = decay_rate_action * (initial_q_value - agent_state[1 - choice])
  
  # Update the value of the OBSERVED outcome based on the reward received.
  outcome_value_update_observed = learning_rate_outcome * (reward - agent_state[outcome + 2])
  
  # Decay the value of the UNOBSERVED outcome towards the initial value.
  outcome_value_decay_unobserved = learning_rate_outcome * (initial_q_value - agent_state[1 - outcome + 2])
  
  # Update the outcome-to-action association weight. This weight determines how strongly
  # outcome values influence action choices.
  association_weight_update = outcome_to_action_learning_rate * prediction_error_action
  
  # Apply all the calculated updates to the agent's state vector.
  agent_state = agent_state.at[choice].add(q_value_update_chosen_action)
  agent_state = agent_state.at[1 - choice].add(q_value_decay_unchosen_action)
  agent_state = agent_state.at[outcome + 2].add(outcome_value_update_observed)
  agent_state = agent_state.at[1 - outcome + 2].add(outcome_value_decay_unobserved)
  agent_state = agent_state.at[4].add(association_weight_update)


  # --- 4. Calculate Action Preferences (Combined Q-values) for the Next Choice ---

  # Start with the learned Q-values for each action (the "instrumental" component).
  instrumental_values = agent_state[0:2]

  # Calculate the influence from outcome values (the "Pavlovian" component).
  # The learned association weight is passed through a sigmoid to be between 0 and 1.
  outcome_values = agent_state[2:4]
  association_weight = jax.nn.sigmoid(agent_state[4])
  pavlovian_influence = association_weight * outcome_association_strength * outcome_values

  # Combine the instrumental and Pavlovian values.
  combined_q_values = instrumental_values + pavlovian_influence
  
  # Add a perseveration bonus to the action that was just chosen, making it more likely to be repeated.
  combined_q_values = combined_q_values.at[choice].add(action_perseveration)
  
  # Add a perseveration bonus for the outcome that was just observed. This promotes
  # choosing the action that is associated with that outcome.
  # Note: This assumes a mapping where choice `i` is associated with outcome `i`.
  combined_q_values = combined_q_values.at[outcome].add(outcome_perseveration[outcome])


  # --- 5. Convert Action Preferences to Choice Logits ---

  # Scale the final Q-values by the inverse temperature. A higher value (lower "temperature")
  # leads to more deterministic choices, while a lower value leads to more random choices.
  # The result is the choice logits, which can be passed to a softmax function to get probabilities.
  choice_logits = combined_q_values * softmax_inverse_temperature

  return choice_logits, agent_state


# Shortened to 100k steps then refactored, run 2, medium floor.
def rat_twostep_run2_simplified_medium_floor(
    params: chex.Array,
    choice: int,
    reward: int,
    outcome: int,
    agent_state: Optional[chex.Array],
) -> tuple[chex.Array, chex.Array]:
  """
  Models an agent's learning and decision-making process for a single trial.
  """
  # --- 1. Unpack and transform model parameters ---
  # Parameters are often stored in log-space for optimization, so we exponentiate them.
  learning_rate = jnp.exp(params[0])
  inverse_temperature = jnp.exp(params[1])
  
  # Separate learning rates for the two possible outcomes.
  outcome_learning_rates = jnp.exp(params[3:5])
  
  # Stickiness parameters encourage repeating previous actions or choices that led to a specific outcome.
  stickiness = params[6]
  outcome_stickiness = params[7]
  
  # Sensitivity to the value of outcomes when making a choice.
  outcome_sensitivity = params[8:]

  # --- 2. Initialize or retrieve agent's internal state ---
  # The agent's state consists of its learned Q-values and outcome values.
  # If this is the first trial (agent_state is None), initialize values to zero.
  if agent_state is None:
    q_values = jnp.zeros(2)
    outcome_values = jnp.zeros(2)
  else:
    q_values, outcome_values = agent_state

  # --- 3. Update Q-values (action values) ---
  # This is a simple reinforcement learning update rule (Rescorla-Wagner).
  # The value of the chosen action is updated to be closer to a target value of 1.
  # Q_new = Q_old * (1 - alpha) + alpha * R, where R=1.
  q_values = (q_values * (1 - learning_rate)).at[choice].add(learning_rate)

  # --- 4. Update outcome values ---
  # First, define the target values for this trial's outcomes.
  # The observed outcome's target is the received reward.
  # The unobserved outcome's target is (1 - reward).
  target_outcomes = jnp.zeros(2).at[outcome].set(reward).at[1 - outcome].set(1 - reward)
  
  # Update the outcome values based on the prediction error (target - current value).
  prediction_error = target_outcomes - outcome_values
  outcome_values = outcome_values + outcome_learning_rates * prediction_error

  # --- 5. Store the updated values in the agent state ---
  # Stack the updated Q-values and outcome values to pass to the next trial.
  agent_state = jnp.stack([q_values, outcome_values])

  # --- 6. Calculate choice probabilities (logits) for the next decision ---
  # The logits determine the probability of choosing each action. A higher logit means a higher probability.
  
  # Start with a base value combining Q-values and outcome values.
  # The inverse_temperature parameter controls how deterministic the choices are.
  base_logits = inverse_temperature * (q_values + outcome_values * outcome_sensitivity)
  
  # Add a "stickiness" bonus to the logit of the action that was just chosen.
  # This encourages the agent to repeat its last action.
  q_stickiness_bonus = stickiness * q_values[choice]
  logits_with_q_stickiness = base_logits.at[choice].add(q_stickiness_bonus)
  
  # Add an "outcome stickiness" bonus to the logit of the action associated with the last outcome.
  # This encourages the agent to choose actions that lead to the same outcome again.
  outcome_stickiness_bonus = outcome_stickiness * outcome_values[outcome]
  choice_logits = logits_with_q_stickiness.at[outcome].add(outcome_stickiness_bonus)

  # --- 7. Return results ---
  # Return the choice logits for the next action and the updated agent state.
  return choice_logits, agent_state


# Shortened to 100k steps then refactored, run 3, medium floor.
def rat_twostep_run3_simplified_medium_floor(
    params: chex.Array,
    choice: int,
    reward: int,
    outcome: int,
    agent_state: Optional[chex.Array],
) -> Tuple[chex.Array, chex.Array]:
  """
  Updates the agent's state and calculates action logits for the next step.

  This agent model includes several components:
  1. Q-learning for choices: Learns the value of taking a specific action.
  2. Q-learning for outcomes: Learns the value associated with a specific outcome.
  3. Stickiness/Perseveration: A tendency to repeat the last chosen action.
  4. Dynamic Inverse Temperature: The exploration/exploitation rate is learned over time.
  """

  # 1. Unpack and transform model parameters.
  # Parameters that are rates or weights (0-1 range) are passed through a sigmoid function.
  sigmoid_params = jax.nn.sigmoid(params[:7])
  learning_rate_choice = sigmoid_params[0]
  learning_rate_outcome = sigmoid_params[1]
  choice_retention_rate = sigmoid_params[2]  # How much of the old Q-choice value is kept (1 - decay)
  outcome_retention_rate = sigmoid_params[3] # How much of the old Q-outcome value is kept (1 - decay)
  stickiness_decay_rate = sigmoid_params[4]
  inverse_temperature_learning_rate = sigmoid_params[5]
  outcome_q_baseline = sigmoid_params[6]

  # Parameters that are not constrained to the 0-1 range.
  raw_params = params[7:]
  outcome_weight = raw_params[0]             # How much to weigh outcome Q-values vs. choice Q-values.
  inverse_temperature_baseline = raw_params[1] # The target value for the inverse temperature.
  choice_bias = raw_params[2]                # A bias added to the first choice's logit.

  # 2. Unpack the agent's internal state (or initialize it if this is the first step).
  # The agent's state is a single vector containing all its internal variables.
  if agent_state is None:
    # If no state is provided, initialize all values to zero.
    # The state vector has 7 elements: 2 for choice Q-values, 2 for outcome Q-values,
    # 2 for stickiness, and 1 for the inverse temperature.
    current_state_vector = jnp.zeros(7)
  else:
    current_state_vector = agent_state

  # Split the state vector into its meaningful components.
  # [0:2]: Q-values for the two choices.
  # [2:4]: Q-values for the two outcomes.
  # [4:6]: Stickiness values for the two choices.
  # [6:7]: The current inverse temperature (a scalar value).
  split_indices = [2, 4, 6]
  q_values_choices, q_values_outcomes, stickiness_values, current_inverse_temperature = \
      jnp.split(current_state_vector, split_indices)

  # 3. Update the agent's internal state based on the last action, reward, and outcome.

  # 3a. Update the Q-values associated with the *choice*.
  # This is a two-part update:
  #   - First, all choice Q-values decay based on the `choice_retention_rate`.
  #   - Second, the Q-value for the action that was actually chosen is updated
  #     using a standard delta rule: Q_new = Q_old + learning_rate * (reward - Q_old).
  q_values_choices_updated = (q_values_choices * choice_retention_rate).at[choice].set(
      q_values_choices[choice] + learning_rate_choice * (reward - q_values_choices[choice])
  )

  # 3b. Update the Q-values associated with the *outcome*.
  # This is similar to the choice Q-value update but uses outcome-specific parameters.
  # A baseline value is also added.
  q_values_outcomes_updated = (q_values_outcomes * outcome_retention_rate + outcome_q_baseline).at[outcome].set(
      q_values_outcomes[outcome] + learning_rate_outcome * (reward - q_values_outcomes[outcome])
  )

  # 3c. Update the stickiness values.
  #   - First, all stickiness values decay towards 0.
  #   - Second, the stickiness for the chosen action is increased.
  # This creates a tendency to repeat the same action (perseveration).
  stickiness_values_updated = (stickiness_values * (1 - stickiness_decay_rate)).at[choice].add(
      stickiness_decay_rate
  )

  # 3d. Update the inverse temperature (softmax temperature).
  # This value controls the agent's exploration vs. exploitation.
  # It is updated to move from its current value towards a baseline value.
  current_inverse_temperature_updated = current_inverse_temperature + \
      inverse_temperature_learning_rate * (inverse_temperature_baseline - current_inverse_temperature)

  # 4. Re-assemble the updated state variables into a single array for the next time step.
  agent_state = jnp.concatenate(
      (q_values_choices_updated,
       q_values_outcomes_updated,
       stickiness_values_updated,
       current_inverse_temperature_updated)
  )

  # 5. Calculate the logits for the next choice.
  # Logits are the raw values that will be fed into a softmax function to determine choice probabilities.

  # 5a. Combine the choice-based and outcome-based Q-values using the outcome_weight.
  combined_q_values = (1 - outcome_weight) * q_values_choices_updated + \
                      outcome_weight * q_values_outcomes_updated

  # 5b. Scale the combined Q-values by the learned inverse temperature.
  # A higher inverse temperature leads to more deterministic (exploitative) choices.
  scaled_values = combined_q_values * current_inverse_temperature_updated

  # 5c. Add the choice stickiness term to promote/discourage repeating actions.
  values_with_stickiness = scaled_values + stickiness_values_updated

  # 5d. Add a fixed bias to the first choice (e.g., a left/right bias).
  choice_logits = values_with_stickiness.at[0].add(choice_bias)

  # 6. Return the calculated logits and the new agent state.
  return choice_logits, agent_state


# Shortened to 100k steps then refactored, run 1, high floor.
def rat_twostep_run1_simplified_high_floor(
    params: chex.Array,
    choice: int,
    reward: int,
    outcome: int,
    agent_state: Optional[chex.Array],
) -> tuple[chex.Array, chex.Array]:
  """Cognitive model describing rat behavior on a binary two-step task.

  Args:
    params: Model params. Different parameters are used for different rats.
    choice: Previous choice. Values: 0 or 1.
    reward: Previous reward. Values: 0 or 1.
    outcome: Previous outcome observed following the choice. Values: 0 or 1.
    agent_state: Previous state of the agent

  Returns:
    choice_logits: Vector of shape (2,) with the probabilities that the rat will
      choose option 0 or 1 on the next trial, expressed as logits.
    agent_state: New state of the agent, after observing the previous choice and
      reward.
  """
  # --- 1. Initialize Agent State ---
  # If this is the first trial, initialize the agent's internal state with
  # neutral default values.
  if agent_state is None:
    agent_state = jnp.array([
        # q_s1: Q-values for the two choices in the first stage (S1)
        0., 0.,
        # q_s2: Q-values for the two outcomes in the second stage (S2)
        0.5, 0.5,
        # transition_strengths: Unnormalized transition counts from S1 to S2
        0.01, 0.01, 0.01, 0.01,
        # reward_consistency_tracker: Tracks how consistently a choice is rewarded
        0.5, 0.5
    ])

  # --- 2. Unpack and Transform Model Parameters ---
  # Unpack the learned parameters for this specific agent.
  (
      learning_rate_s1_base_logit,
      learning_rate_s2,
      learning_rate_transition_strength_logit,
      inverse_temperature_log,
      model_based_weight_logit,
      learning_rate_s2_decay_logit,
      stickiness_bias,
      outcome_perseveration_weight,
      reward_consistency_sensitivity,
      prior_choice_bias,
  ) = params

  # Transform parameters from logit or log space into their usable ranges.
  # Sigmoid maps logits to (0, 1), and exp maps log-space values to (> 0).
  learning_rate_s1_base = jax.nn.sigmoid(learning_rate_s1_base_logit)
  learning_rate_transition_strength = jax.nn.sigmoid(learning_rate_transition_strength_logit)
  inverse_temperature = jnp.exp(inverse_temperature_log)
  model_based_weight = jax.nn.sigmoid(model_based_weight_logit)
  learning_rate_s2_decay = jax.nn.sigmoid(learning_rate_s2_decay_logit)

  # --- 3. Unpack Agent State ---
  # Unpack the agent's internal state vector into meaningful components.
  q_s1, q_s2, transition_strengths_flat, reward_consistency_tracker = jnp.split(
      agent_state, [2, 4, 8]
  )
  # Reshape the flat transition strengths into a 2x2 matrix (choice -> outcome).
  transition_strengths = transition_strengths_flat.reshape((2, 2))

  # --- 4. Model-Free (MF) Value Updates ---
  # Calculate the simple model-free prediction error at the first stage (S1).
  # This is the difference between the received reward and the expected value.
  prediction_error_s1_mf = reward - q_s1[choice]

  # Update a tracker for how consistently the chosen action has been rewarded.
  # The learning rate for this tracker is modulated by the magnitude of the prediction error.
  consistency_learning_rate = learning_rate_s1_base * jax.nn.sigmoid(jnp.abs(prediction_error_s1_mf))
  reward_consistency_update = consistency_learning_rate * (reward - reward_consistency_tracker[choice])
  reward_consistency_tracker = reward_consistency_tracker.at[choice].add(reward_consistency_update)

  # Calculate the prediction error at the second stage (S2).
  # This includes a term for perseverating on the same outcome.
  prediction_error_s2 = reward + outcome_perseveration_weight - q_s2[outcome]
  
  # Update the Q-value for the observed outcome based on the S2 prediction error.
  q_s2_update_for_observed_outcome = learning_rate_s2 * prediction_error_s2
  q_s2 = q_s2.at[outcome].add(q_s2_update_for_observed_outcome)
  
  # Apply a decay to the Q-value of the unobserved outcome.
  q_s2 = q_s2.at[1 - outcome].multiply(1 - learning_rate_s2_decay)

  # Apply a decay to the Q-value of the unchosen action in S1.
  q_s1 = q_s1.at[1 - choice].multiply(1 - learning_rate_s1_base)

  # --- 5. Model-Based (MB) Value Updates ---
  # Convert the raw transition strengths into probabilities using softmax.
  p_s1_to_s2 = jax.nn.softmax(transition_strengths, axis=1)

  # Calculate the model-based value of each S1 choice by considering the
  # values of the possible S2 outcomes, weighted by transition probabilities.
  value_mb = jnp.sum(p_s1_to_s2 * q_s2, axis=1)

  # Calculate the model-based prediction error for the first stage (S1).
  prediction_error_s1_mb = reward - value_mb[choice]

  # Update the transition strengths: increase the strength for the observed
  # transition and decrease it for the transition from the unchosen action
  # to the same outcome.
  transition_strengths = transition_strengths.at[choice, outcome].add(learning_rate_transition_strength)
  transition_strengths = transition_strengths.at[1 - choice, outcome].add(-learning_rate_transition_strength)

  # --- 6. Hybrid Q-Value Update for S1 ---
  # Modulate the base learning rate by the reward consistency.
  # If a choice is consistently rewarded, its learning rate increases.
  reward_consistency_term = reward_consistency_sensitivity * (2.0 * reward_consistency_tracker[choice] - 1.0)
  effective_learning_rate_s1 = learning_rate_s1_base * (1.0 + reward_consistency_term)

  # Combine the model-free and model-based prediction errors using the MB weight.
  combined_prediction_error = (
      (1 - model_based_weight) * prediction_error_s1_mf +
      model_based_weight * prediction_error_s1_mb
  )

  # Update the Q-value for the chosen action in S1 using the combined error.
  q_s1_update = effective_learning_rate_s1 * combined_prediction_error
  q_s1 = q_s1.at[choice].add(q_s1_update)

  # --- 7. Prepare for Next Trial ---
  # Reassemble the updated components into the new agent state vector.
  agent_state = jnp.concatenate([
      q_s1,
      q_s2,
      transition_strengths.flatten(),
      reward_consistency_tracker
  ])

  # Combine the model-free (q_s1) and model-based (value_mb) values.
  combined_action_values = (1 - model_based_weight) * q_s1 + model_based_weight * value_mb

  # Scale the action values by the inverse temperature to determine choice stochasticity.
  scaled_action_values = inverse_temperature * combined_action_values

  # Add a stickiness bias to increase the likelihood of repeating the previous choice.
  choice_logits_with_stickiness = scaled_action_values.at[choice].add(stickiness_bias)

  # Add a prior bias towards choosing action 0.
  final_choice_logits = choice_logits_with_stickiness.at[0].add(prior_choice_bias)

  # --- 8. Return Outputs ---
  # Return the final choice logits and the updated agent state.
  return final_choice_logits, agent_state


# Shortened to 100k steps then refactored, run 2, high floor.
def rat_twostep_run2_simplified_high_floor(
    params: chex.Array,
    choice: int,
    reward: int,
    outcome: int,
    agent_state: Optional[chex.Array],
) -> tuple[chex.Array, chex.Array]:
  """Cognitive model describing rat behavior on a binary two-step task.

  Args:
    params: Model params. Different parameters are used for different rats.
    choice: Previous choice. Values: 0 or 1.
    reward: Previous reward. Values: 0 or 1.
    outcome: Previous outcome observed following the choice. Values: 0 or 1.
    agent_state: Previous state of the agent

  Returns:
    choice_logits: Vector of shape (2,) with the probabilities that the rat will
      choose option 0 or 1 on the next trial, expressed as logits.
    agent_state: New state of the agent, after observing the previous choice and
      reward.
  """
  # --- 1. Initialize or Unpack Agent State ---

  # On the first trial, initialize the agent's state (Q-values and outcome
  # values) to zeros. The state is a 2x2 array where:
  # - row 0: Q-values for choice 0 and 1
  # - row 1: Outcome values for outcome 0 and 1
  if agent_state is None:
    agent_state = jnp.zeros((2, 2))

  # Unpack the agent's internal state into Q-values and outcome-values.
  q_values, outcome_values = agent_state

  # --- 2. Unpack and Transform Model Parameters ---

  # Parameters are passed as a flat vector. We assign them to meaningful names
  # and apply transformations (e.g., exp, sigmoid) to constrain their ranges.
  learning_rate = jnp.exp(params[0])              # Rate for Q-value updates (alpha)
  inverse_temperature = jnp.exp(params[1])        # Choice stochasticity (beta)
  retention_rate = jax.nn.sigmoid(params[2])      # Forgetting/decay factor
  
  # Learning rates for the two outcome values, constrained to be in [0, 1].
  outcome_learning_rates = jax.nn.sigmoid(params[3:5])
  
  outcome_weight = params[5]                      # Weight of outcome values in choice
  stickiness_bias = params[6]                     # Bias to repeat the same choice
  outcome_stickiness_bias = params[7]             # Bias related to the observed outcome
  
  # Sensitivity to each outcome value in the choice rule.
  outcome_sensitivity = params[8:10]

  # --- 3. Update Agent's Internal Values Based on Previous Trial ---

  # Store the Q-value of the action that was actually chosen *before* any decay.
  q_chosen_pre_decay = q_values[choice]

  # Identify the outcome that was *not* observed (the counterfactual outcome).
  opposite_outcome = 1 - outcome

  # Apply a decay/forgetting factor to all existing Q-values.
  q_values = q_values * retention_rate

  # Calculate the prediction error for the chosen action. This is based on the
  # observed reward and the value of the *counterfactual* outcome.
  # Note: The pre-decay Q-value is used in this calculation.
  prediction_error_chosen_q = reward - q_chosen_pre_decay + (
      outcome_values[opposite_outcome] - q_chosen_pre_decay
  )
  
  # Update the Q-value of the chosen action. This overwrites the decayed value.
  updated_q_chosen = q_chosen_pre_decay + learning_rate * prediction_error_chosen_q
  q_values = q_values.at[choice].set(updated_q_chosen)

  # Update the Q-value for the *unchosen* action, as if it had led to the
  # observed outcome. This is a form of counterfactual learning.
  prediction_error_unchosen_q = outcome_values[outcome] - q_values[opposite_outcome]
  q_values = q_values.at[opposite_outcome].add(
      learning_rate * prediction_error_unchosen_q
  )

  # Update the value of the *observed* outcome. The target for this update is
  # the sum of the reward and the value of the chosen action (before decay).
  learning_rate_for_outcome = outcome_learning_rates[outcome]
  prediction_error_observed_outcome = (
      reward + q_chosen_pre_decay - outcome_values[outcome]
  )
  outcome_values = outcome_values.at[outcome].add(
      learning_rate_for_outcome * prediction_error_observed_outcome
  )

  # Update the value of the *unobserved* outcome. It both decays and learns
  # towards a target of '1 - reward'.
  learning_rate_for_unobserved_outcome = outcome_learning_rates[opposite_outcome]
  prediction_error_unobserved_outcome = (1 - reward) - outcome_values[opposite_outcome]
  
  updated_unobserved_outcome_value = (
      outcome_values[opposite_outcome] * retention_rate +
      learning_rate_for_unobserved_outcome * prediction_error_unobserved_outcome
  )
  outcome_values = outcome_values.at[opposite_outcome].set(updated_unobserved_outcome_value)
  
  # --- 4. Prepare Outputs for the Next Trial ---

  # Re-assemble the updated values into the new agent state.
  agent_state = jnp.stack([q_values, outcome_values])

  # Calculate the choice logits for the next decision.
  # Start with a base value combining Q-values and outcome-values.
  base_logits = q_values + outcome_weight * outcome_values * outcome_sensitivity

  # Scale the logits by the inverse temperature. A higher value leads to more
  # deterministic choices.
  scaled_logits = inverse_temperature * base_logits

  # Add a "stickiness" bias to the previously chosen action, making it more
  # likely to be repeated. The bias is proportional to the action's Q-value.
  logits_with_choice_stickiness = scaled_logits.at[choice].add(
      stickiness_bias * q_values[choice]
  )

  # Add an outcome-specific "stickiness" bias. This increases the tendency
  # to choose actions associated with the previously observed outcome.
  final_logits = logits_with_choice_stickiness.at[outcome].add(
      outcome_stickiness_bias * outcome_values[outcome]
  )
  
  choice_logits = final_logits

  return choice_logits, agent_state


# Shortened to 100k steps then refactored, run 3, high floor.
def rat_twostep_run3_simplified_high_floor(
    params: chex.Array,
    choice: int,
    reward: int,
    outcome: int,
    agent_state: Optional[chex.Array],
) -> tuple[chex.Array, chex.Array]:
  """Cognitive model describing rat behavior on a binary two-step task.

  Args:
    params: Model params. Different parameters are used for different rats.
    choice: Previous choice. Values: 0 or 1.
    reward: Previous reward. Values: 0 or 1.
    outcome: Previous outcome observed following the choice. Values: 0 or 1.
    agent_state: Previous state of the agent

  Returns:
    choice_logits: Vector of shape (2,) with the probabilities that the rat will
      choose option 0 or 1 on the next trial, expressed as logits.
    agent_state: New state of the agent, after observing the previous choice and
      reward.
  """
  # --- 1. Initialization ---
  # If this is the first trial, initialize the agent's state to a zero vector.
  # The agent's state tracks Q-values, stickiness, and inverse temperature.
  if agent_state is None:
    agent_state = jnp.zeros(7)

  # --- 2. Unpack and Transform Model Parameters ---
  # Parameters are transformed into appropriate ranges.
  # Sigmoid is used to constrain rates and weights between 0 and 1.
  learning_rate_choice, \
  learning_rate_outcome, \
  q_choice_decay_rate, \
  q_outcome_decay_rate, \
  stickiness_decay_rate, \
  inverse_temperature_learning_rate = jax.nn.sigmoid(params[:6])

  # These parameters are used directly without transformation.
  outcome_weight, \
  outcome_q_baseline, \
  choice_bias, \
  log_baseline_inverse_temperature = params[6:10]

  # --- 3. Unpack Previous Agent State ---
  # The agent state is a vector containing different cognitive variables.
  previous_q_values_for_choices, \
  previous_q_values_for_outcomes, \
  previous_stickiness_values, \
  previous_inverse_temperature = jnp.split(agent_state, [2, 4, 6])

  # --- 4. Update State Components based on Previous Trial ---

  # --- 4a. Update Q-values for Choices (Model-Free Component) ---
  # This is a standard reinforcement learning update for the chosen option.
  other_choice = 1 - choice
  prediction_error_choice = reward - previous_q_values_for_choices[choice]
  updated_q_value_for_chosen = previous_q_values_for_choices[choice] + \
                               learning_rate_choice * prediction_error_choice

  # The unchosen option's Q-value decays towards a fraction of the updated chosen value.
  decayed_q_value_for_unchosen = previous_q_values_for_choices[other_choice] * \
                                 (1 - q_choice_decay_rate)
  decay_target_for_unchosen = (0.5 * updated_q_value_for_chosen) * q_choice_decay_rate
  updated_q_value_for_unchosen = decayed_q_value_for_unchosen + decay_target_for_unchosen

  # Assemble the new choice Q-values.
  new_q_values_for_choices = previous_q_values_for_choices.at[choice].set(updated_q_value_for_chosen)
  new_q_values_for_choices = new_q_values_for_choices.at[other_choice].set(updated_q_value_for_unchosen)

  # --- 4b. Update Q-values for Outcomes (Model-Based Component) ---
  # This update is for the value of the outcome that was observed.
  other_outcome = 1 - outcome
  prediction_error_outcome = reward - previous_q_values_for_outcomes[outcome]
  updated_q_value_for_observed = previous_q_values_for_outcomes[outcome] + \
                                 learning_rate_outcome * prediction_error_outcome

  # The unobserved outcome's Q-value decays towards a baseline.
  decayed_q_value_for_unobserved = previous_q_values_for_outcomes[other_outcome] * \
                                   (1 - q_outcome_decay_rate)
  decay_target_for_unobserved = outcome_q_baseline * q_outcome_decay_rate
  updated_q_value_for_unobserved = decayed_q_value_for_unobserved + decay_target_for_unobserved

  # Assemble the new outcome Q-values.
  new_q_values_for_outcomes = previous_q_values_for_outcomes.at[outcome].set(updated_q_value_for_observed)
  new_q_values_for_outcomes = new_q_values_for_outcomes.at[other_outcome].set(updated_q_value_for_unobserved)

  # --- 4c. Update Choice Stickiness (Perseveration/Switching) ---
  # Stickiness increases for the chosen option if unrewarded, promoting switching.
  stickiness_update_amount = stickiness_decay_rate * (1 - reward)
  updated_stickiness_for_chosen = previous_stickiness_values[choice] + stickiness_update_amount

  # The unchosen option's stickiness decays and is reduced by the same amount.
  decayed_stickiness_for_unchosen = previous_stickiness_values[other_choice] * (1 - stickiness_decay_rate)
  updated_stickiness_for_unchosen = decayed_stickiness_for_unchosen - stickiness_update_amount

  # Assemble the new stickiness values.
  new_stickiness_values = previous_stickiness_values.at[choice].set(updated_stickiness_for_chosen)
  new_stickiness_values = new_stickiness_values.at[other_choice].set(updated_stickiness_for_unchosen)

  # --- 4d. Update Inverse Temperature (Exploration/Exploitation) ---
  # The inverse temperature (decision randomness) slowly adapts towards a baseline.
  target_inverse_temperature = jnp.exp(log_baseline_inverse_temperature)
  temperature_difference = target_inverse_temperature - previous_inverse_temperature
  new_inverse_temperature = previous_inverse_temperature + \
                            inverse_temperature_learning_rate * temperature_difference

  # --- 5. Assemble the New Agent State for the Next Trial ---
  new_agent_state = jnp.concatenate((
      new_q_values_for_choices,
      new_q_values_for_outcomes,
      new_stickiness_values,
      new_inverse_temperature
  ))

  # --- 6. Compute Choice Logits for the Next Trial ---
  # Combine the model-free and model-based Q-values using a learned weight.
  weighted_average_q_values = (1 - outcome_weight) * new_q_values_for_choices + \
                              outcome_weight * new_q_values_for_outcomes

  # Scale the combined values by the inverse temperature to determine their influence on choice.
  value_component = weighted_average_q_values * new_inverse_temperature

  # Add stickiness and a constant bias to the final logits.
  choice_logits = value_component + new_stickiness_values
  choice_logits = choice_logits.at[0].add(choice_bias)

  # --- 7. Return Results ---
  return choice_logits, new_agent_state


def rat_twostep_synthesis(
    params: chex.Array,
    choice: int,
    reward: int,
    outcome: int,
    agent_state: Optional[chex.Array],
) -> tuple[chex.Array, chex.Array]:
  """
  A manual implementation of a rat two-step task model.

  This model combines a recency trace for perseveration, outcome-based
  Q-values, and a dynamic inverse temperature to predict choice behavior.

  Args:
    params: An array of model parameters:
      - params[0]: model_based_weight (weight of outcome values)
      - params[1]: model_based_learning_rate (learning rate for outcome values)
      - params[2]: model_based_forgetting_rate (decay rate for outcome values)
      - params[3]: model_based_forgetting_target (target for outcome value decay)
      - params[4]: perseveration_weight (weight of recency trace)
      - params[5]: perseveration_forgetting_rate (decay rate for recency trace)
      - params[6]: bias (choice bias)
      - params[7]: prev_choice_stickiness (bonus added to the logit of the
                   previous choice)
      - params[8]: inverse_temperature_convergence_rate (rate at which inverse
                   temperature approaches its final value)
    choice: The action taken by the agent in the previous step (0 or 1).
    reward: The reward received after the previous choice (e.g., 0 or 1).
    outcome: The outcome state observed after the previous choice (0 or 1).
    agent_state: The agent's internal state from the previous step,
                 consisting of [recency_trace (1,), outcome_values (2,),
                 prev_inverse_temperature (1,)]. Initialized to zeros if None.

  Returns:
    A tuple containing:
      - choice_logits: Logits for the next choice (shape 2).
      - new_agent_state: The updated agent state for the next step.
  """

  # The agent state consists of a recency trace on actions, and outcome-based'
  # Q-values.
  if agent_state is None:
    agent_state = jnp.zeros(4)
  recency_trace, outcome_values, prev_inverse_temperature = jnp.split(
      agent_state, [1, 3]
  )

  # Outcome-value weight can be positive or negative, and its sign encodes
  # whether this rat is in the congruent or incongruent condition.
  model_based_weight = params[0]
  # Learning and decay rates are bounded between 0 and 1.
  model_based_learning_rate = jax.nn.sigmoid(params[1])
  model_based_forgetting_rate = jax.nn.sigmoid(params[2])
  model_based_forgetting_target = params[3]

  # Empirically, there should be a positive perseveration effect.
  perseveration_weight = jax.nn.softplus(params[4])
  perseveration_forgetting_rate = jax.nn.sigmoid(params[5])
  bias = params[6]
  prev_choice_stickiness = params[7]

  inverse_temperature_convergence_rate = jax.nn.sigmoid(params[8])

  # First, decay the outcome values towards the baseline, then update the value
  # of the chosen outcome.
  outcome_values_after_decay = outcome_values + model_based_forgetting_rate * (
      model_based_forgetting_target - outcome_values
  )
  outcome_prediction_error = reward - outcome_values_after_decay[outcome]
  outcome_update = model_based_learning_rate * outcome_prediction_error
  updated_outcome_values = outcome_values_after_decay.at[outcome].add(
      outcome_update
  )

  # Decay the recency trace, then add 1 to the trace of the chosen action.
  updated_recency_trace = recency_trace * perseveration_forgetting_rate + (2 * choice - 1)

  # Update the inverse temperature.
  inverse_temperature = (
      prev_inverse_temperature
      + inverse_temperature_convergence_rate
      * (1 - prev_inverse_temperature)
  )

  # Ultimately we're just taking a weighted sum of the recency trace and the
  # outcome values, and applying a bias term.
  choice_logits = inverse_temperature * (
      perseveration_weight * updated_recency_trace * jnp.array([-1.0, 1.0])
      + model_based_weight * updated_outcome_values
  ) + bias * jnp.array([-1.0, 1.0])
  choice_logits = choice_logits.at[choice].add(prev_choice_stickiness)

  return choice_logits, jnp.concatenate(
      [updated_recency_trace, updated_outcome_values, inverse_temperature]
  )
```
