## Supplementary figures and images for "AI-Discovered Cognitive Models Reveal Novel Insights into Human and Animal Learning"

### ablation_performance_fly_bandit_run1_high_floor_20260420.pdf

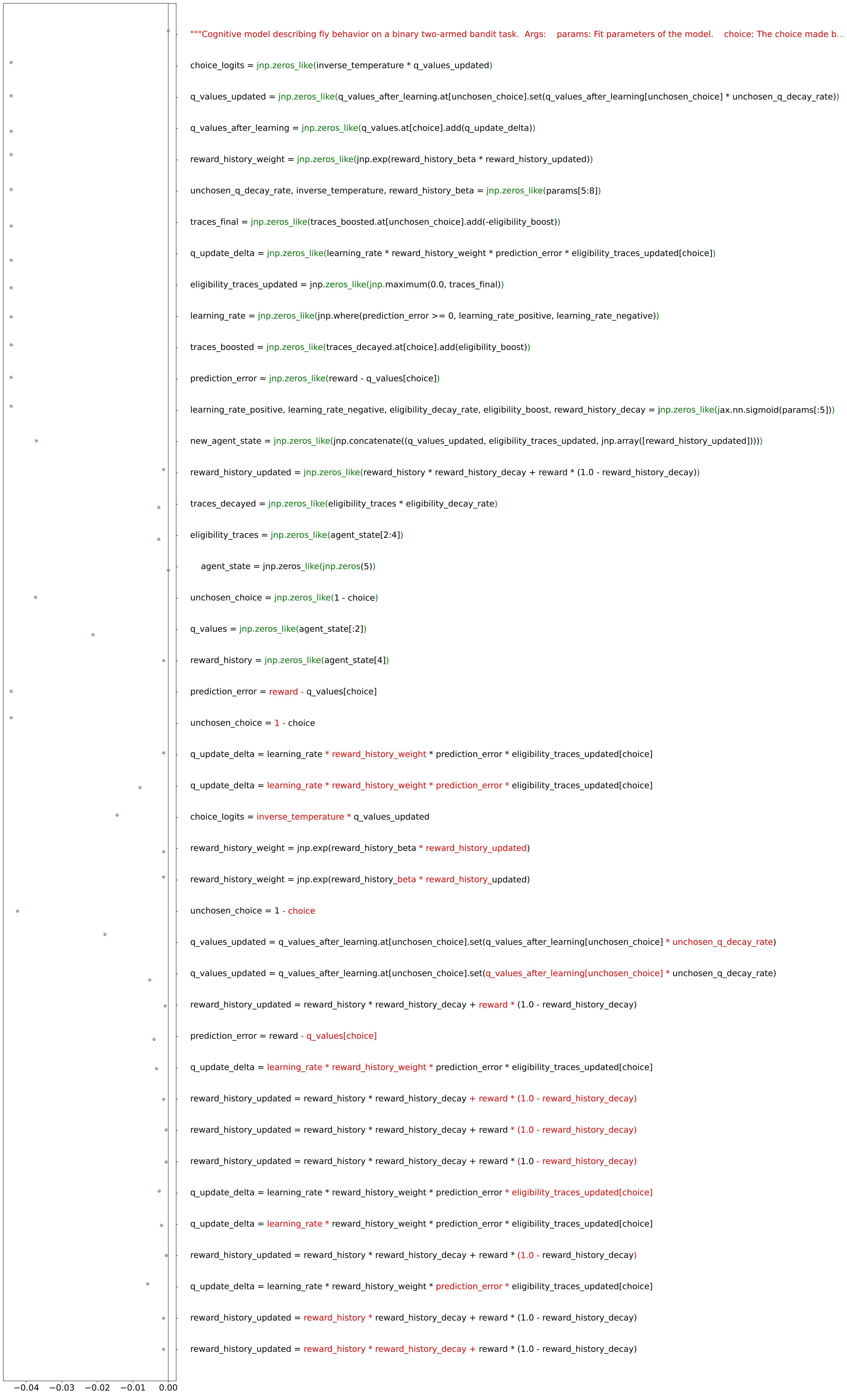

### ablation_performance_fly_bandit_run1_low_floor_20260420.pdf

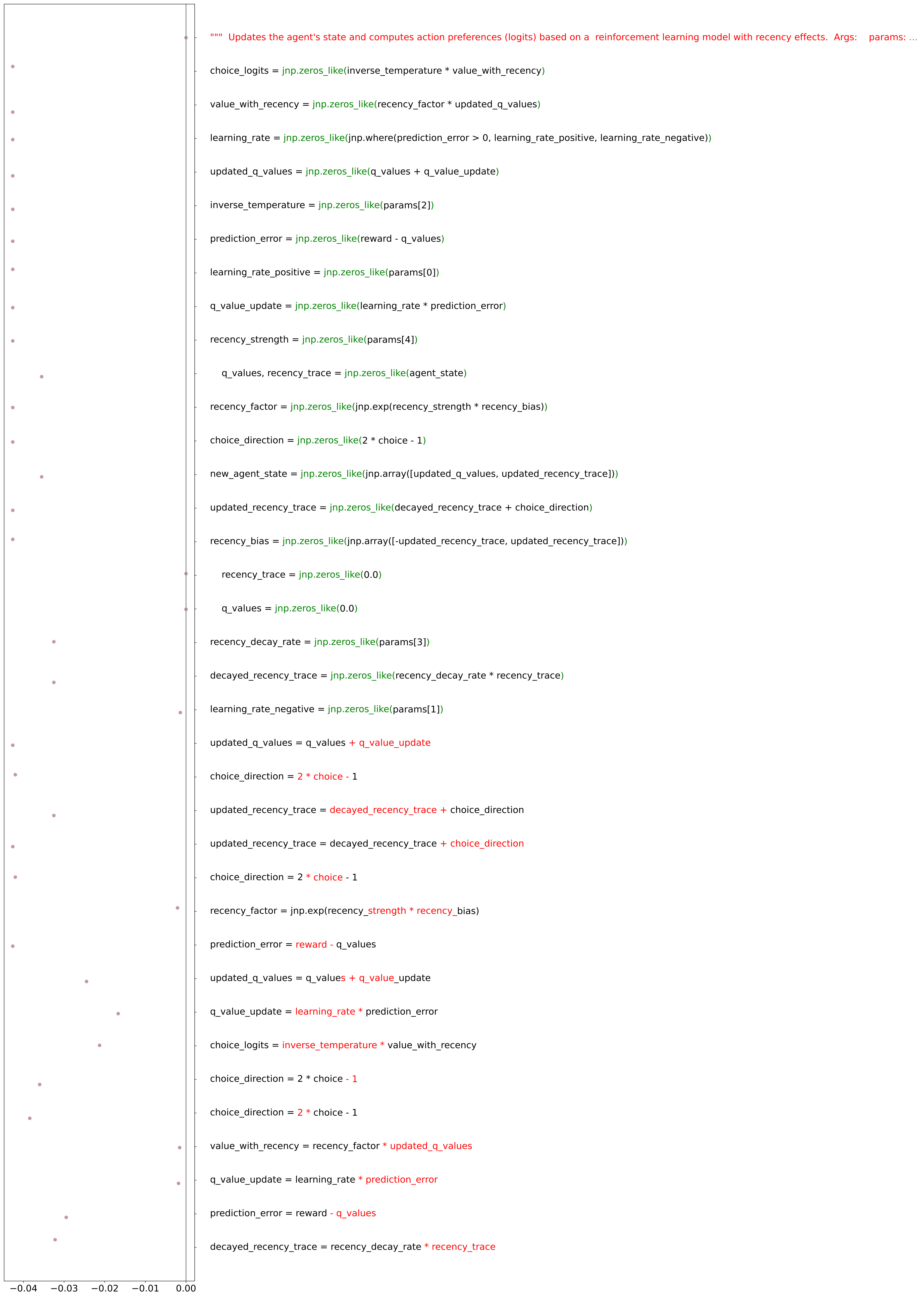

### ablation_performance_fly_bandit_run1_medium_floor_20260420.pdf

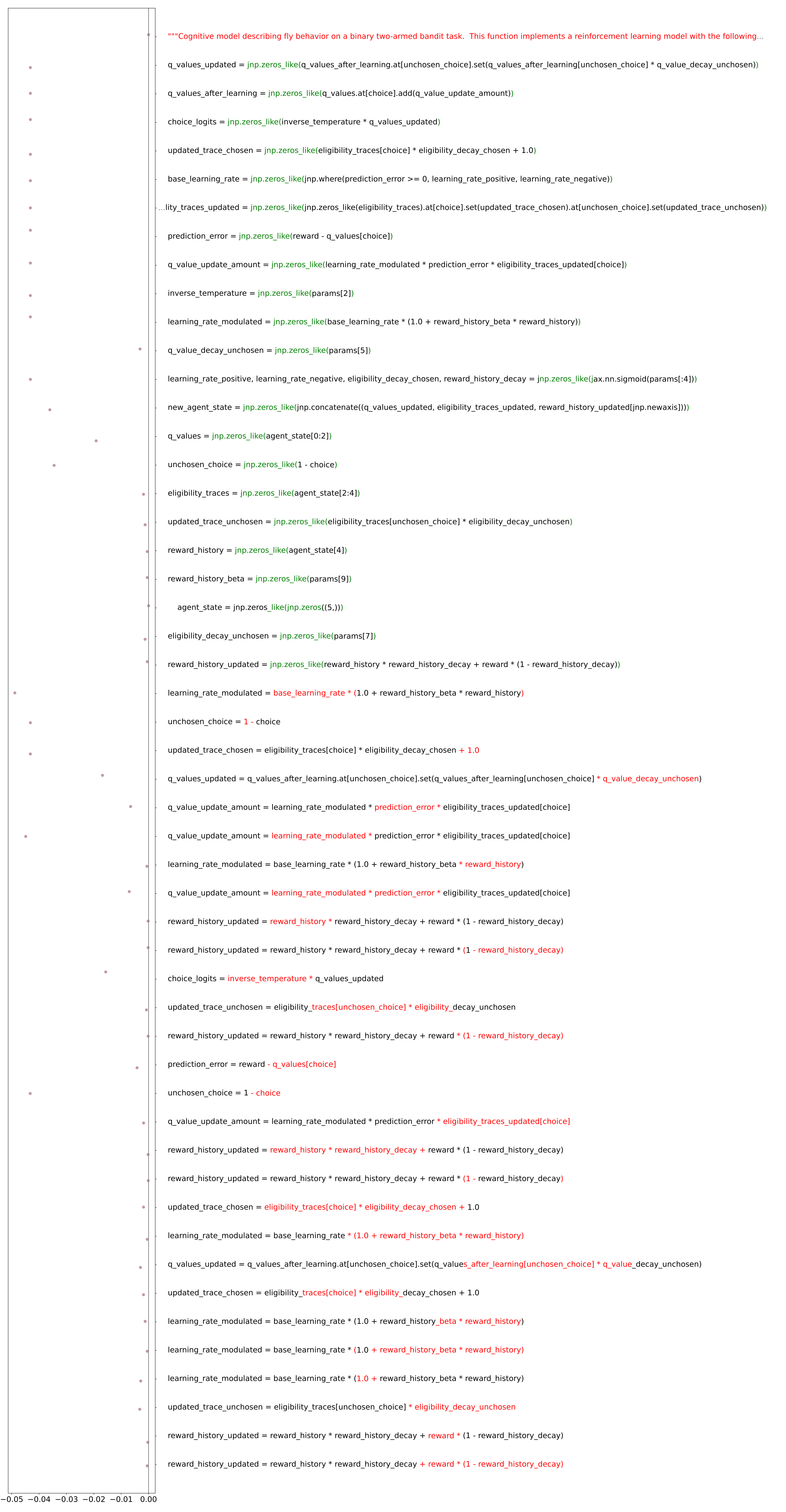

### ablation_performance_fly_bandit_run2_high_floor_20260420.pdf

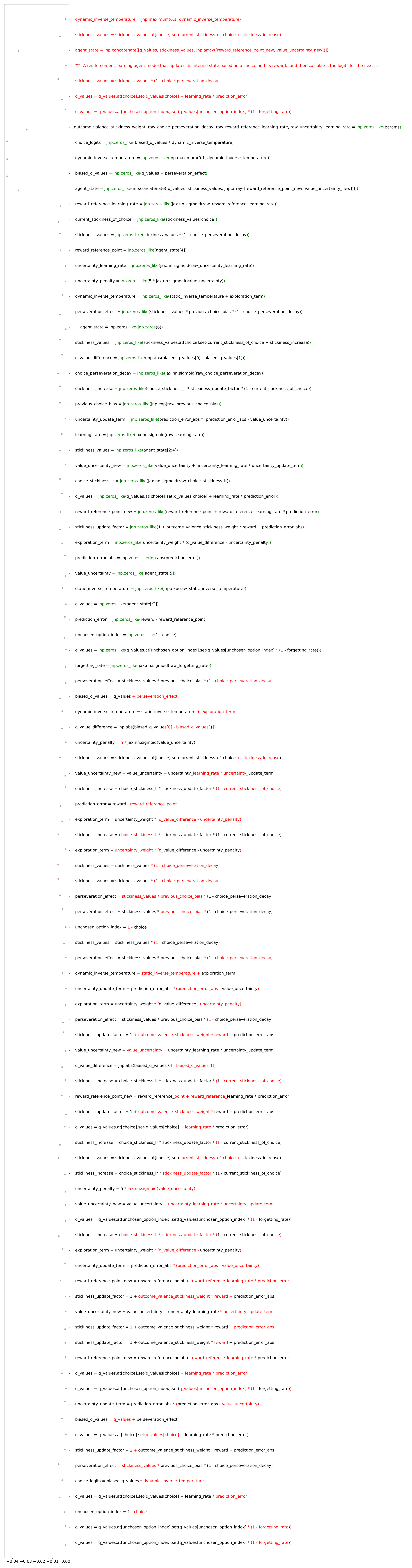

### ablation_performance_fly_bandit_run2_low_floor_20260420.pdf

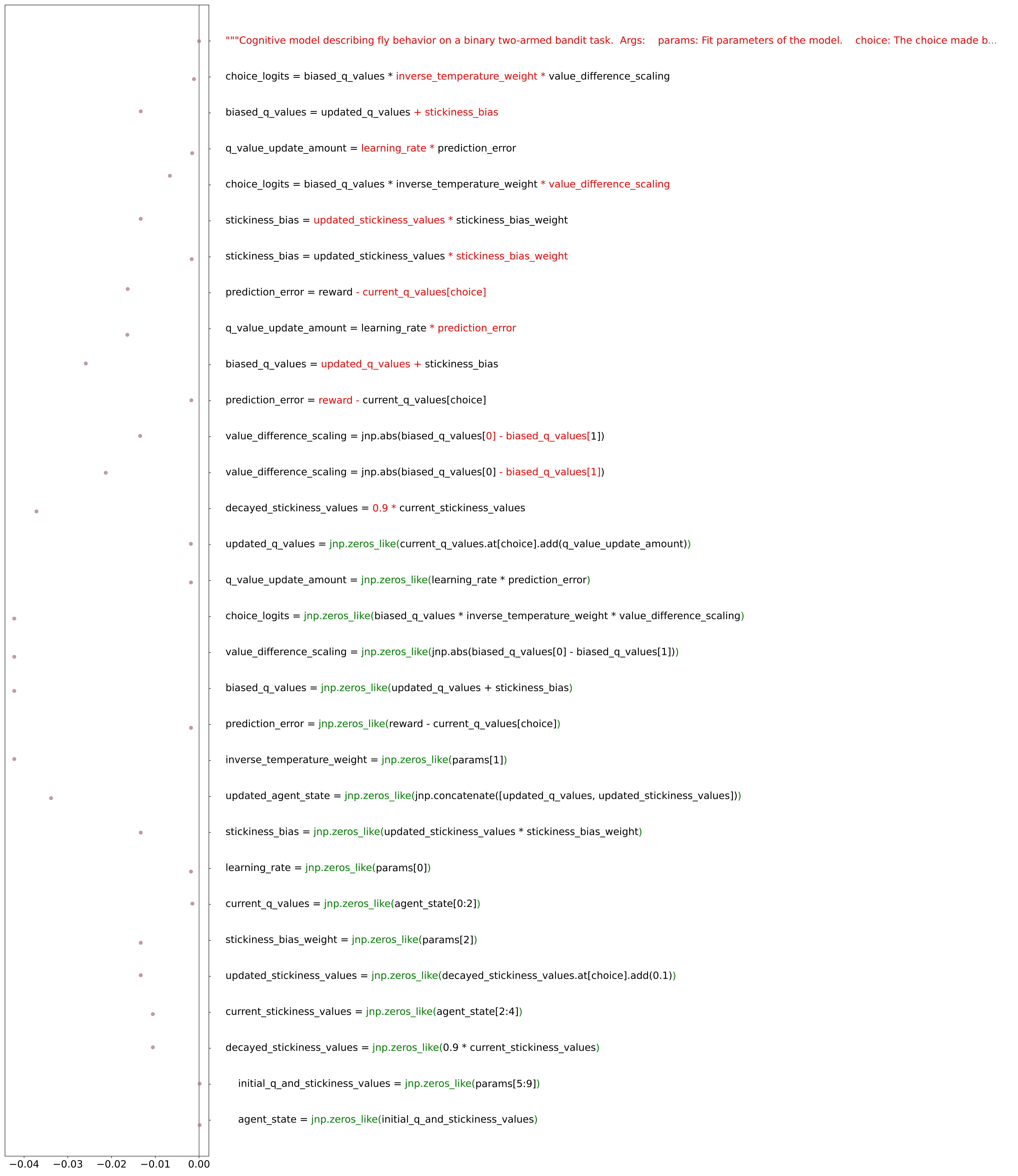

### ablation_performance_fly_bandit_run2_medium_floor_20260420.pdf

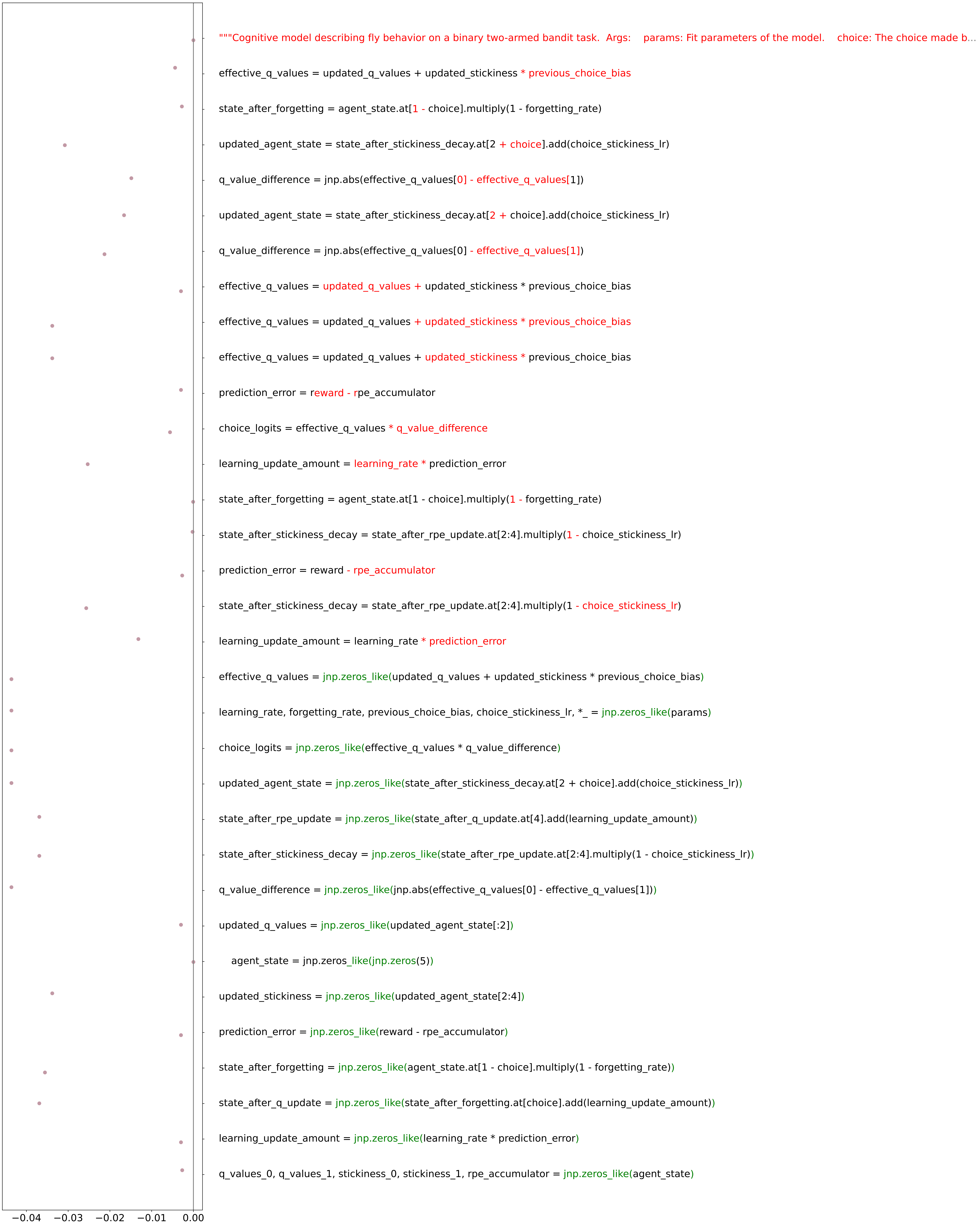

### ablation_performance_fly_bandit_run3_high_floor_20260420.pdf

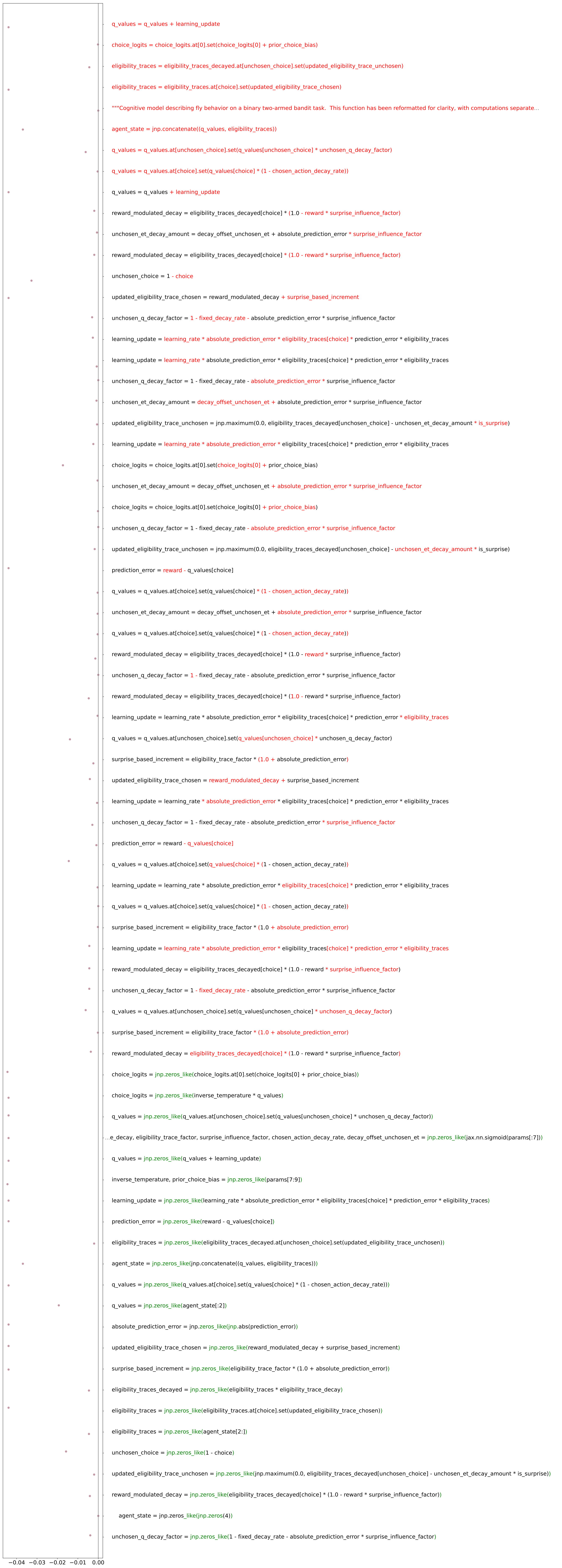

### ablation_performance_fly_bandit_run3_low_floor_20260420.pdf

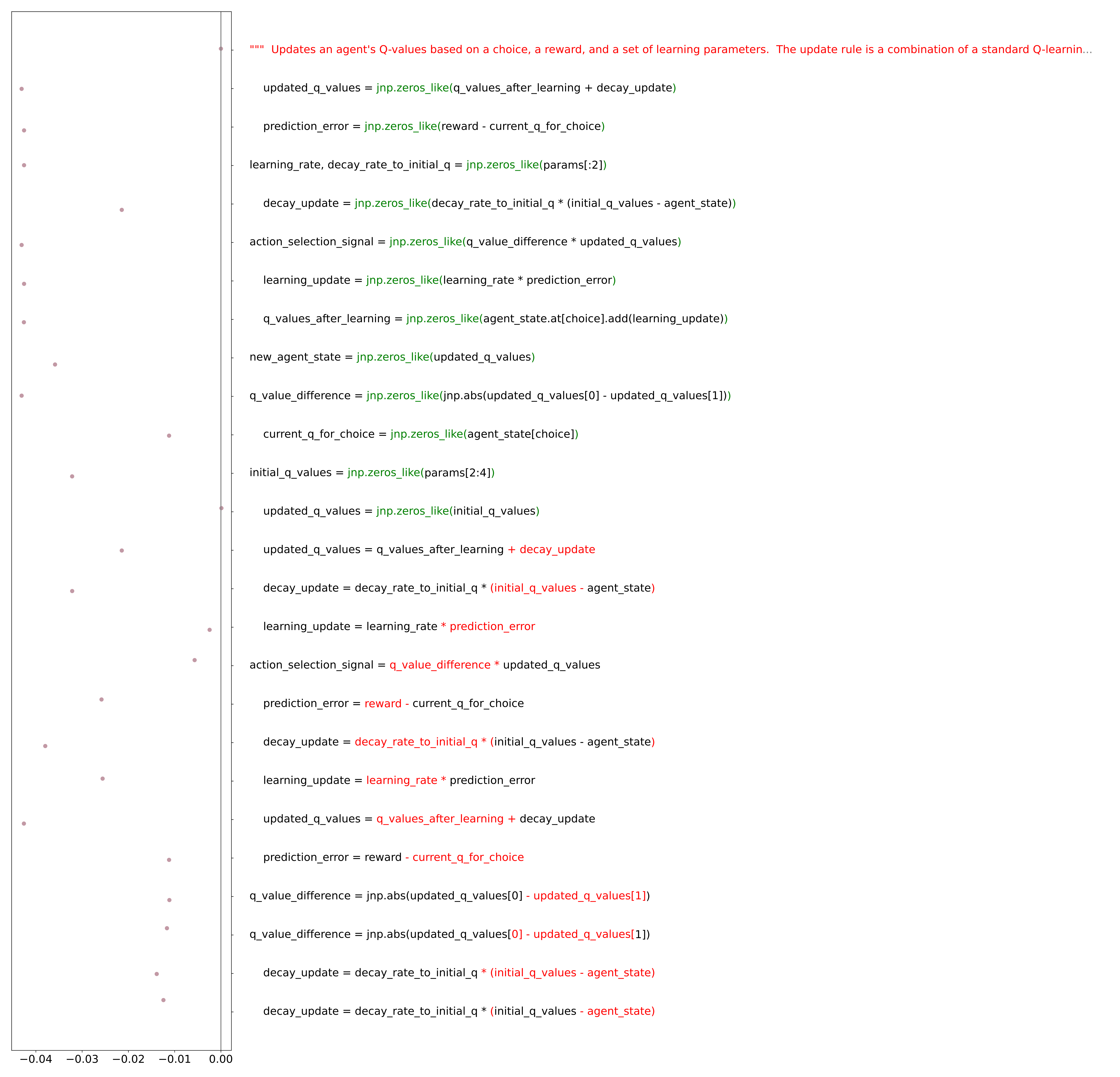

### ablation_performance_fly_bandit_run3_medium_floor_20260420.pdf

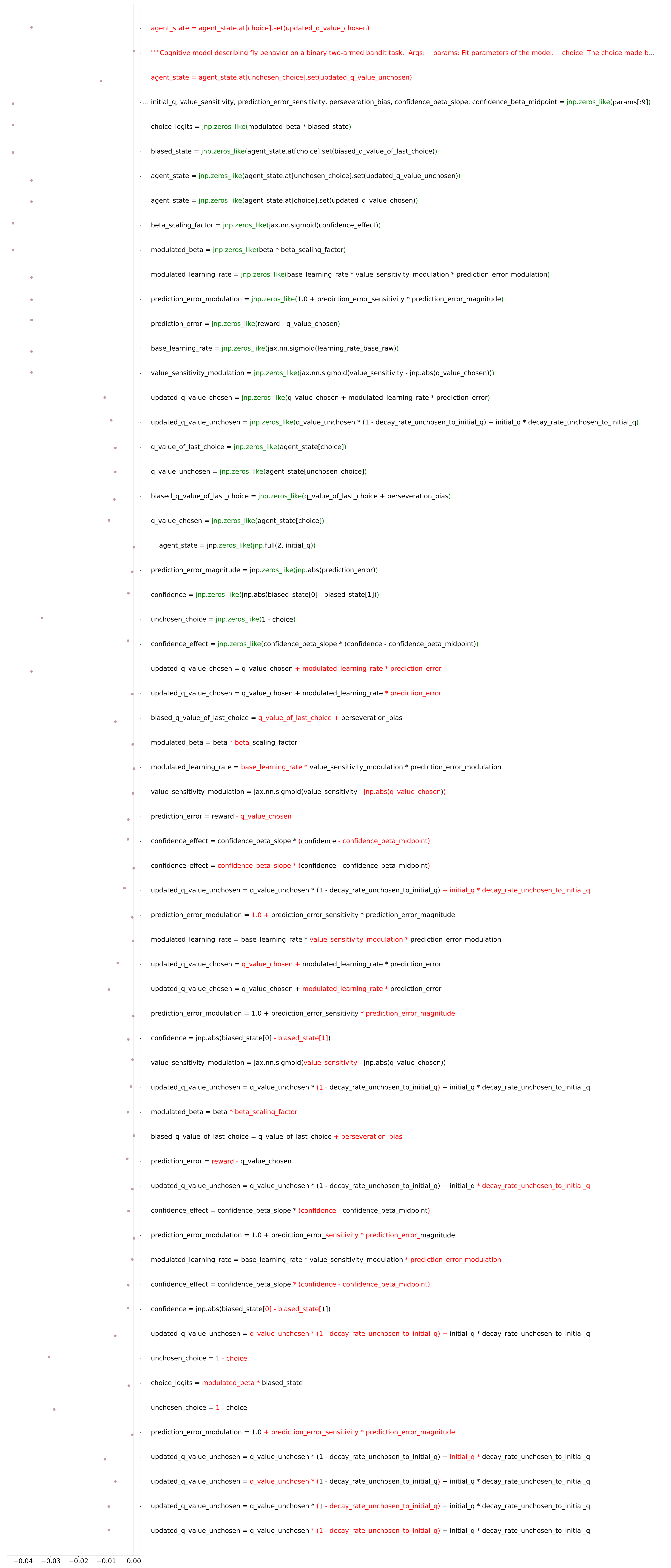

### ablation_performance_human_bandit_run1_high_floor_refactored_20260420.pdf

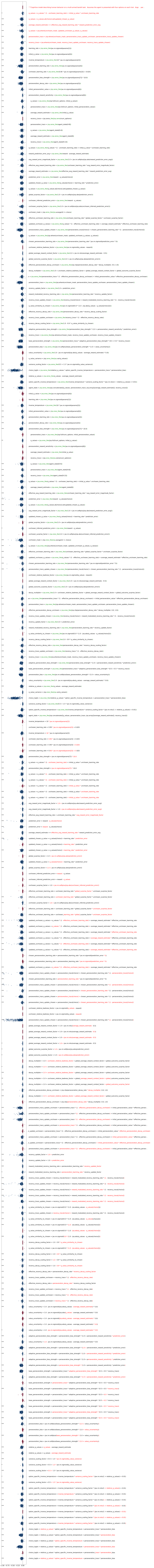

### ablation_performance_human_bandit_run1_low_floor_refactored_20260420.pdf

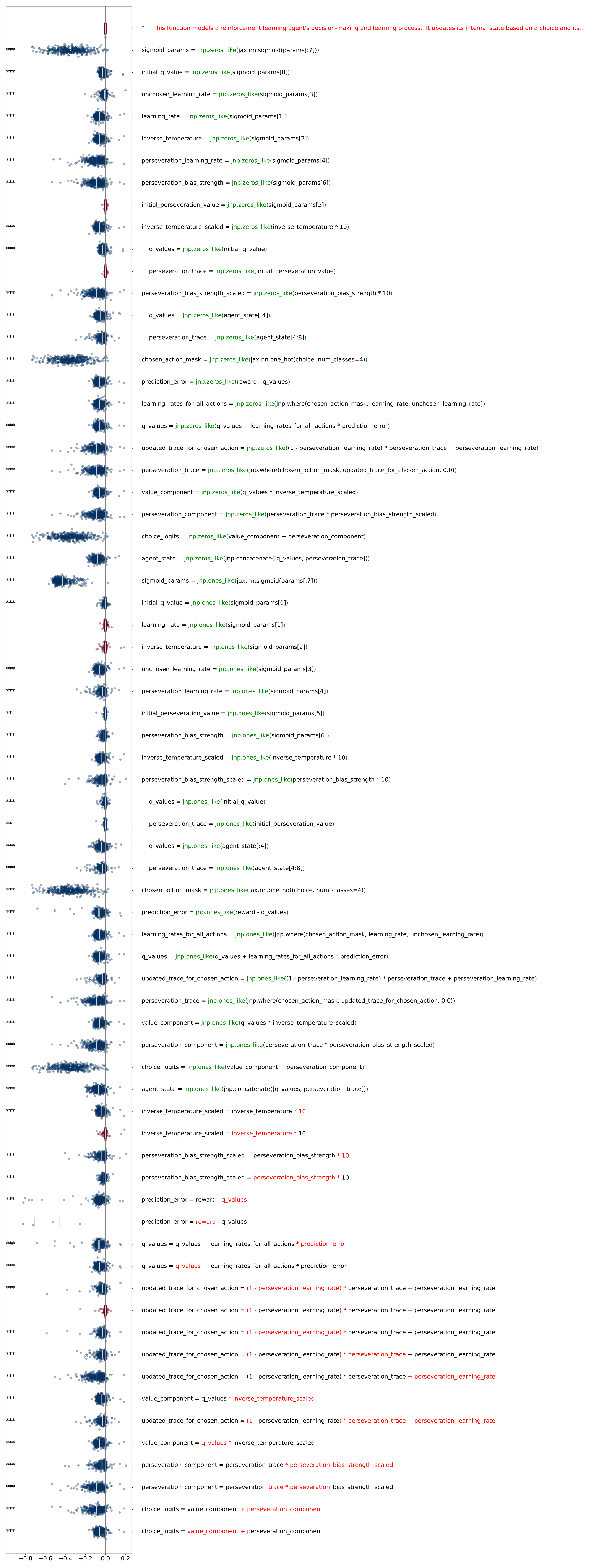

### ablation_performance_human_bandit_run1_medium_floor_refactored_20260420.pdf

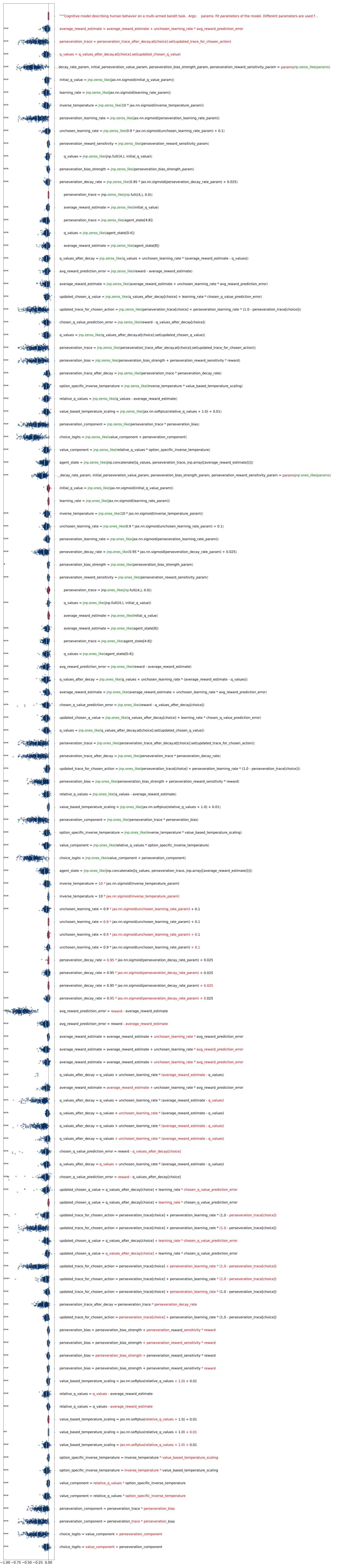

### ablation_performance_human_bandit_run2_low_floor_refactored_20260420.pdf

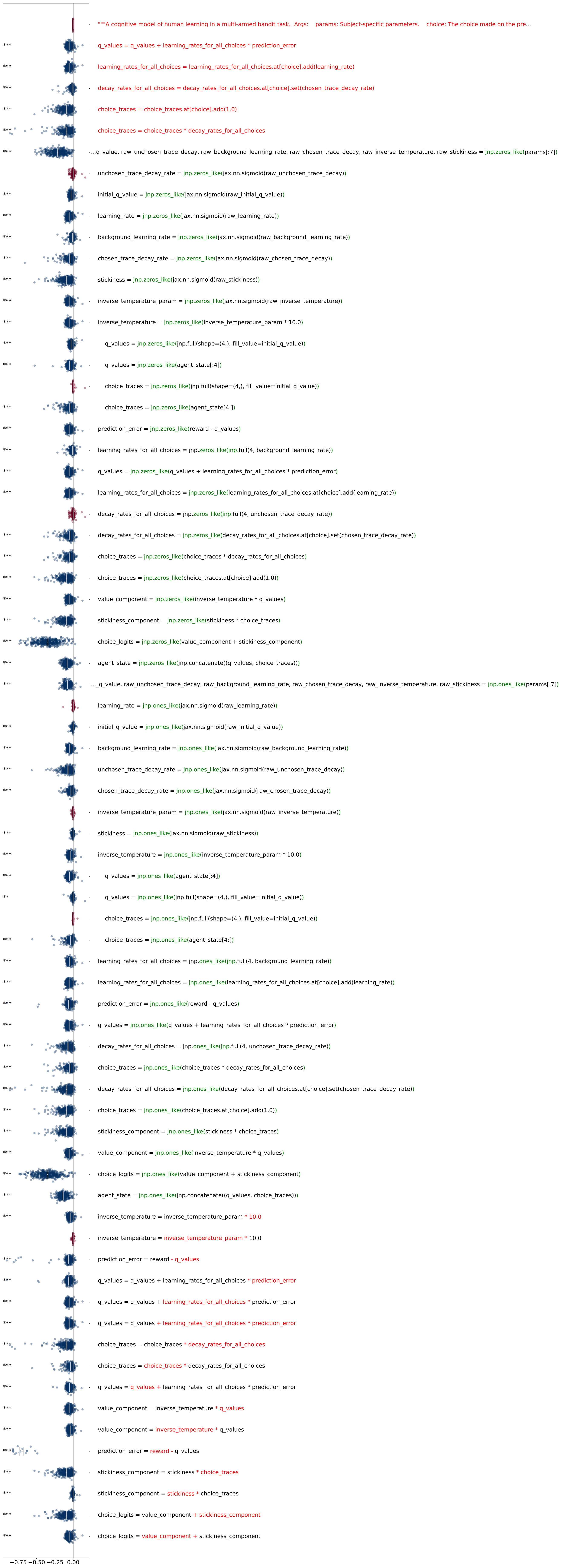

### ablation_performance_human_bandit_run2_medium_floor_refactored_20260420.pdf

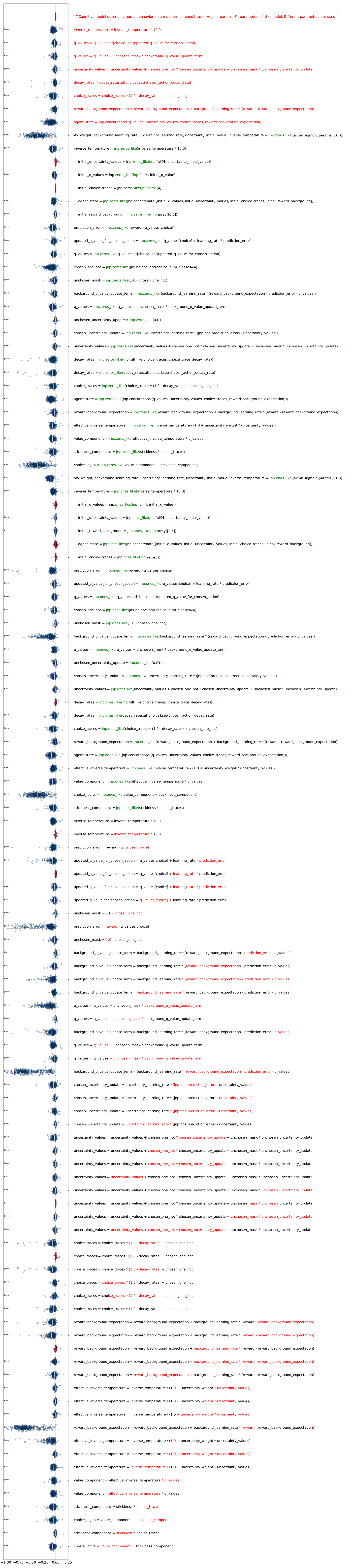

### ablation_performance_human_bandit_run3_high_floor_refactored_20260420.pdf

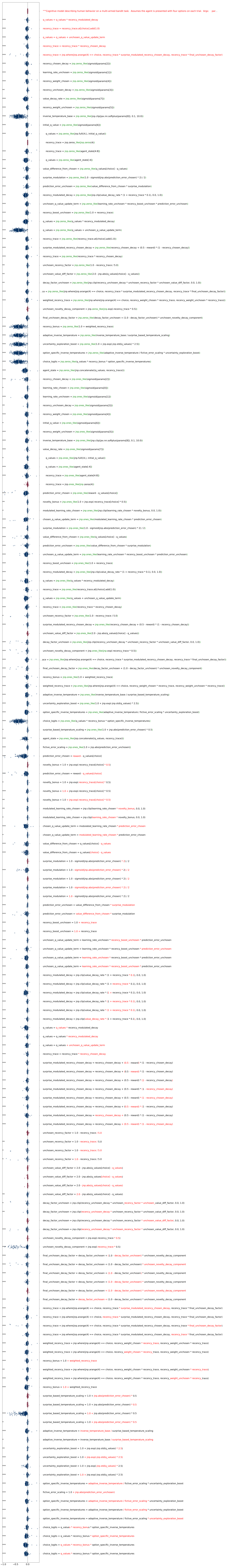

### ablation_performance_human_bandit_run3_low_floor_refactored_20260420.pdf

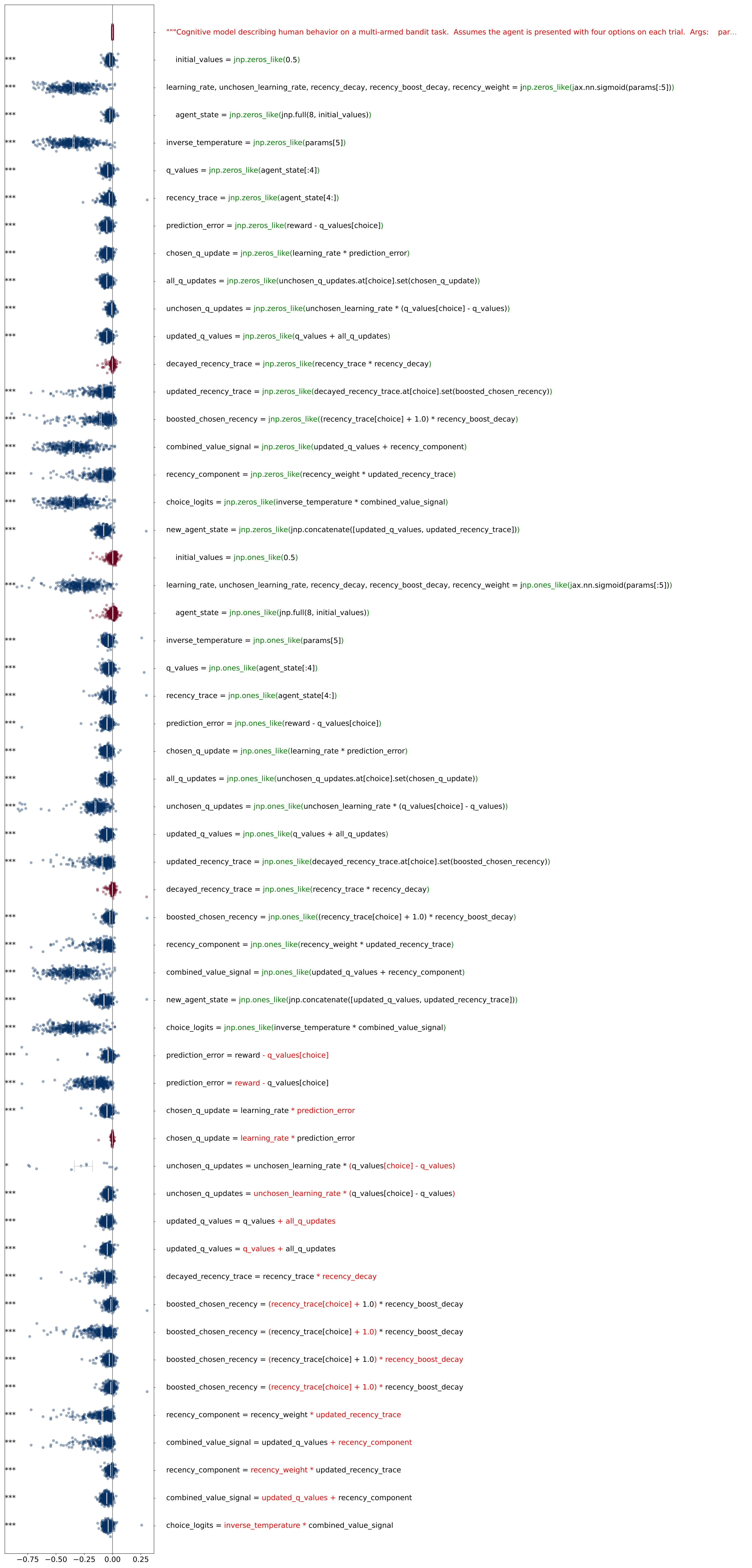

### ablation_performance_human_bandit_run3_medium_floor_refactored_20260420.pdf

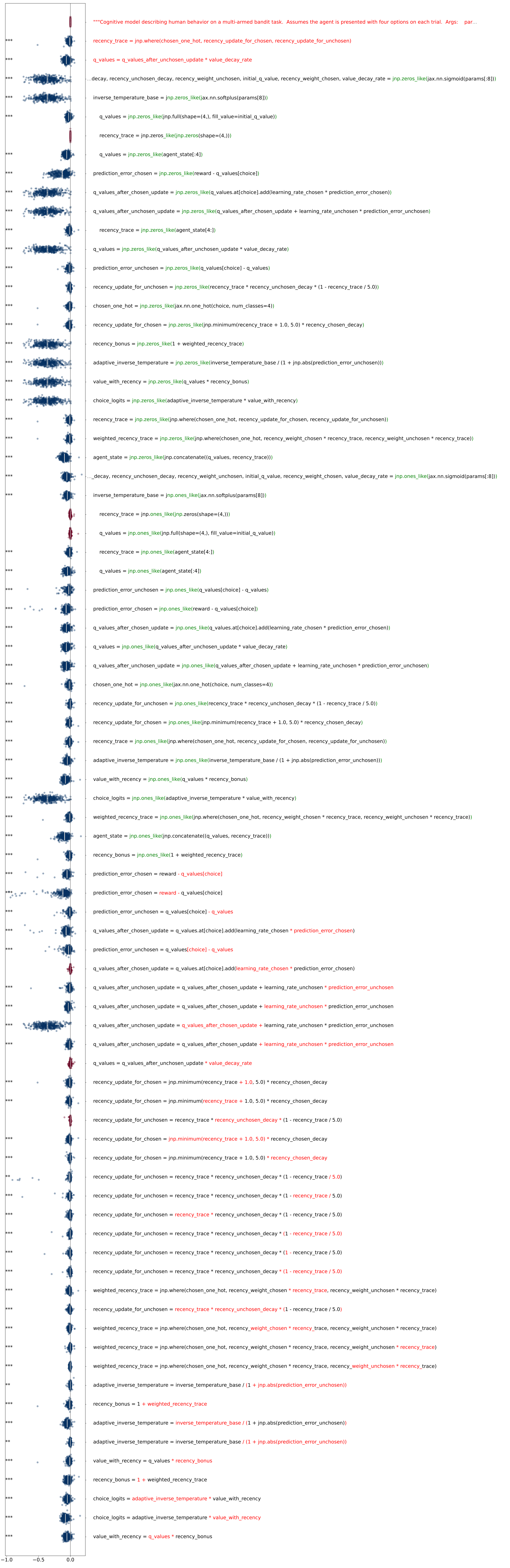

### ablation_performance_monkey_bandit_run1_high_floor_refactored_20260420.pdf

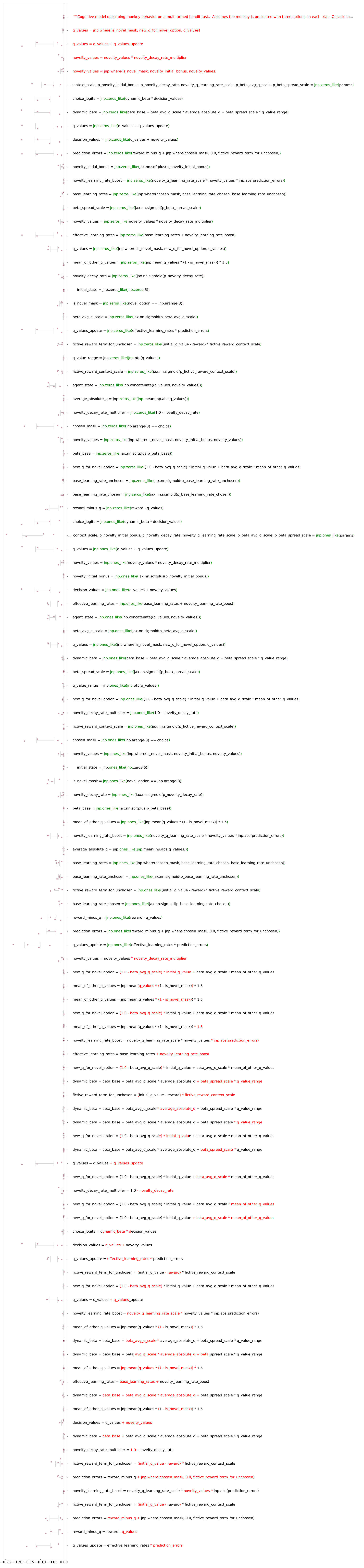

### ablation_performance_monkey_bandit_run1_low_floor_refactored_20260428.pdf

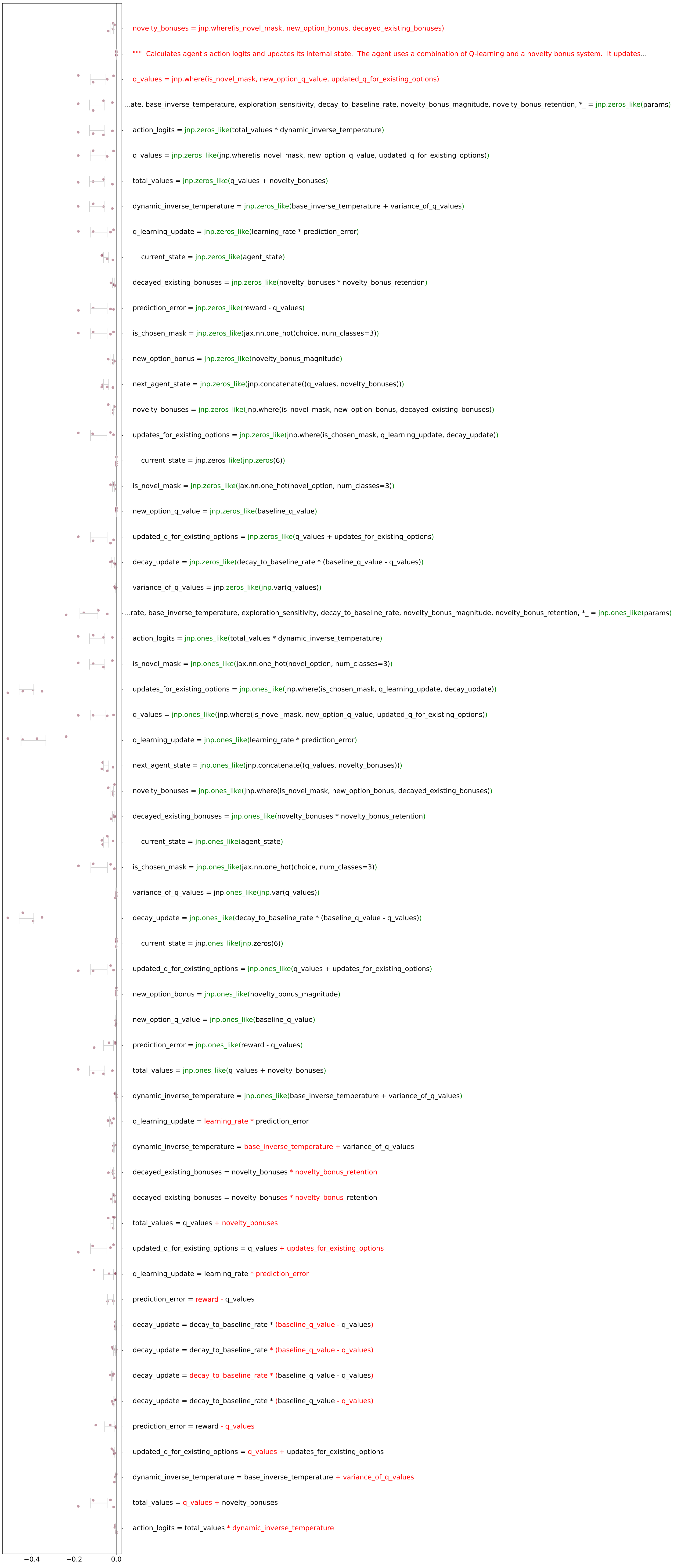

### ablation_performance_monkey_bandit_run1_medium_floor_refactored_20260428.pdf

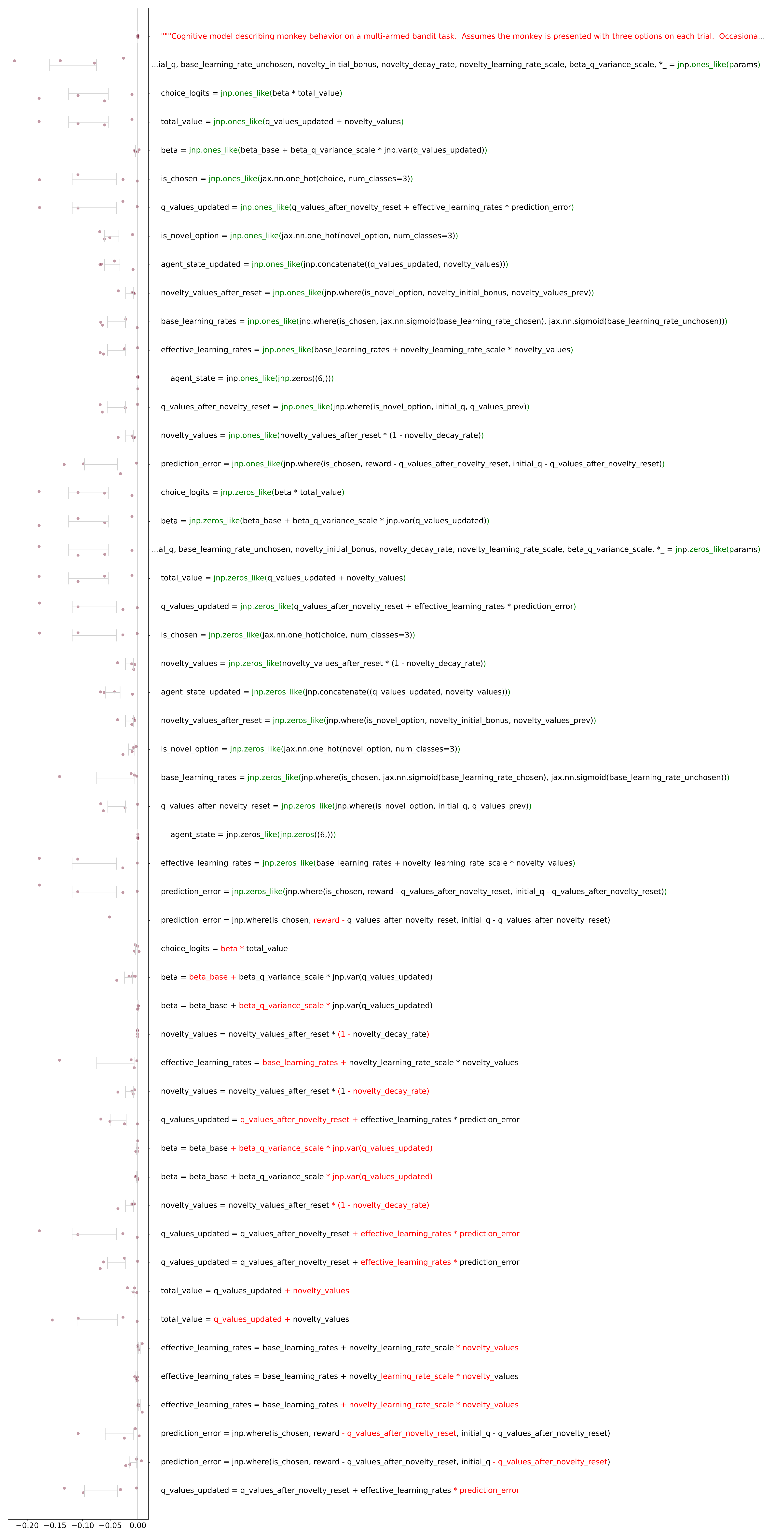

### ablation_performance_monkey_bandit_run2_high_floor_refactored_20260420.pdf

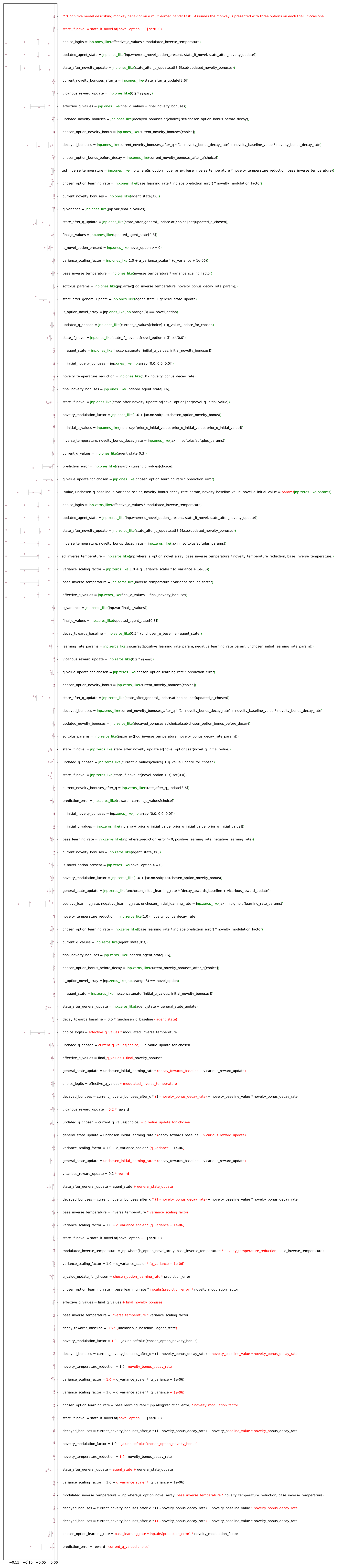

### ablation_performance_monkey_bandit_run2_low_floor_refactored_20260428.pdf

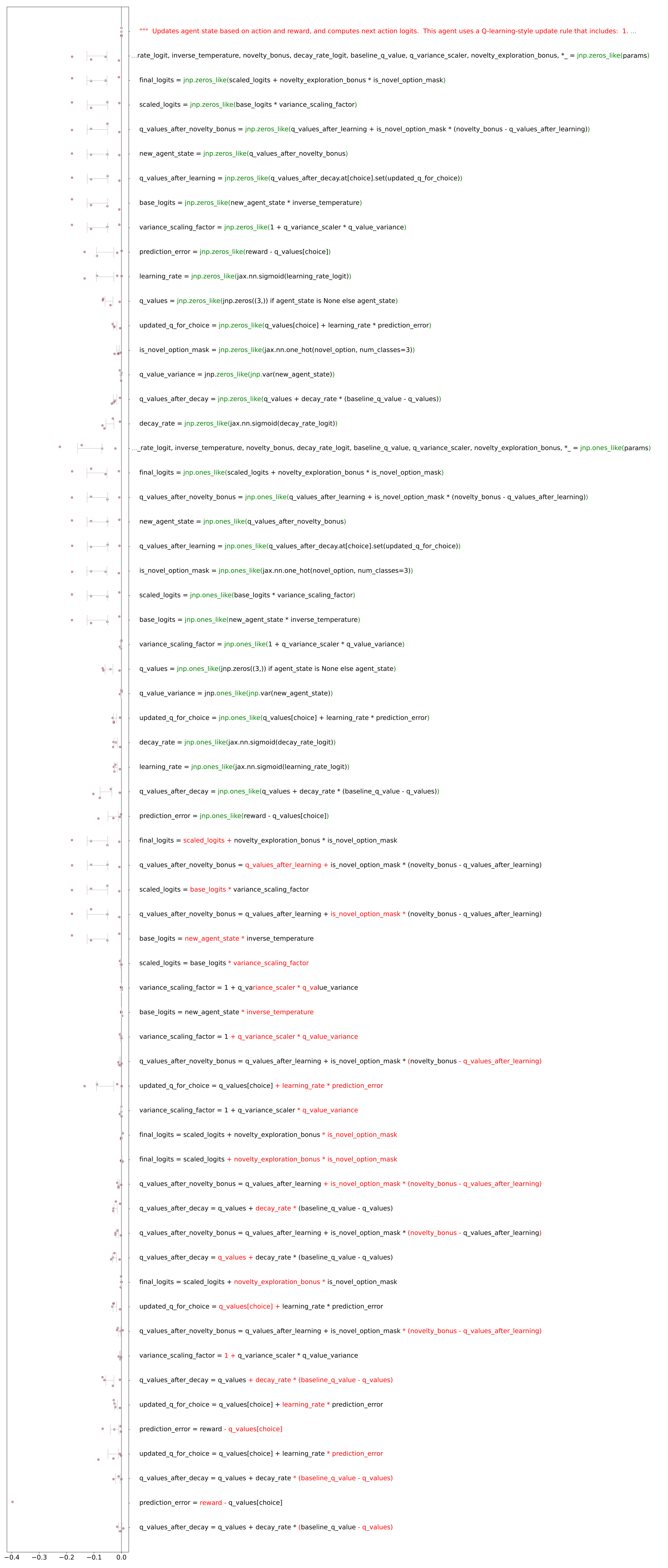

### ablation_performance_monkey_bandit_run2_medium_floor_refactored_20260420.pdf

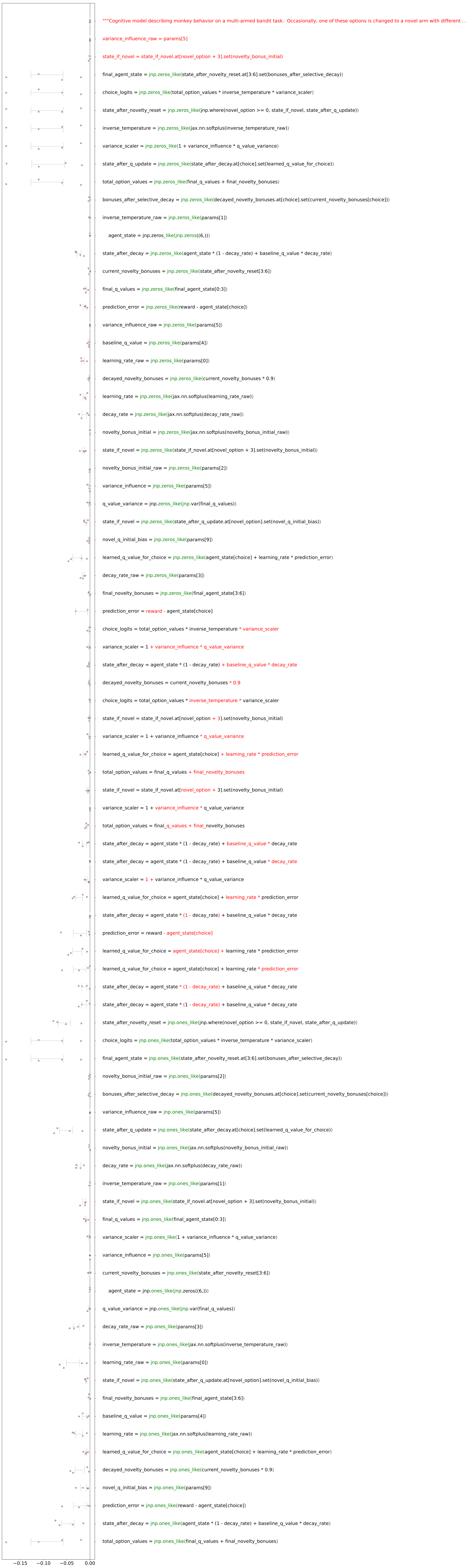

### ablation_performance_monkey_bandit_run3_high_floor_refactored_20260420.pdf

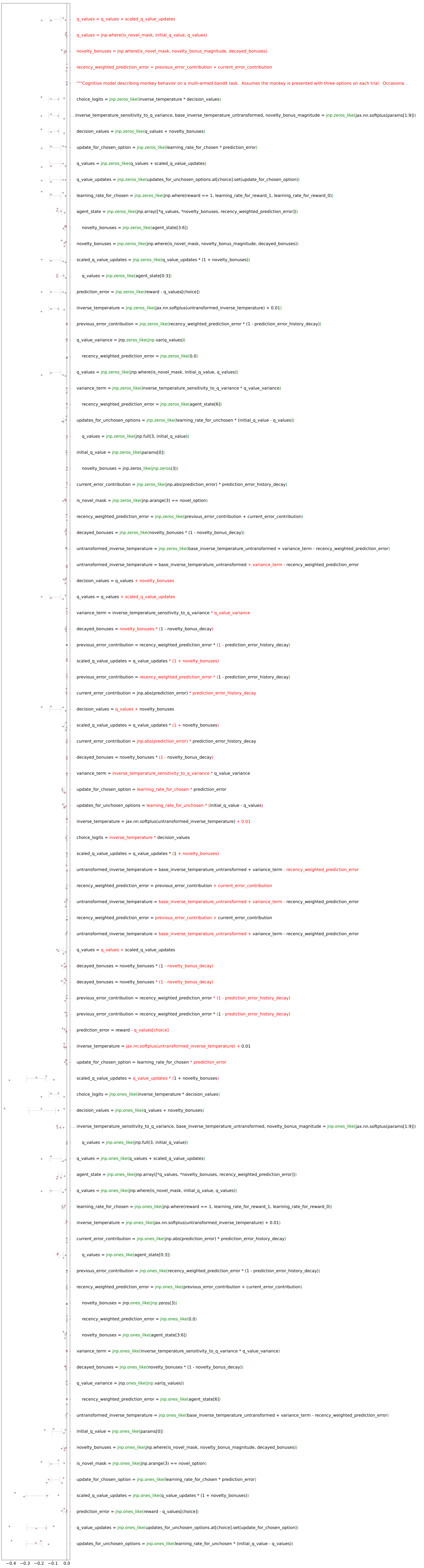

### ablation_performance_monkey_bandit_run3_low_floor_20260420.pdf

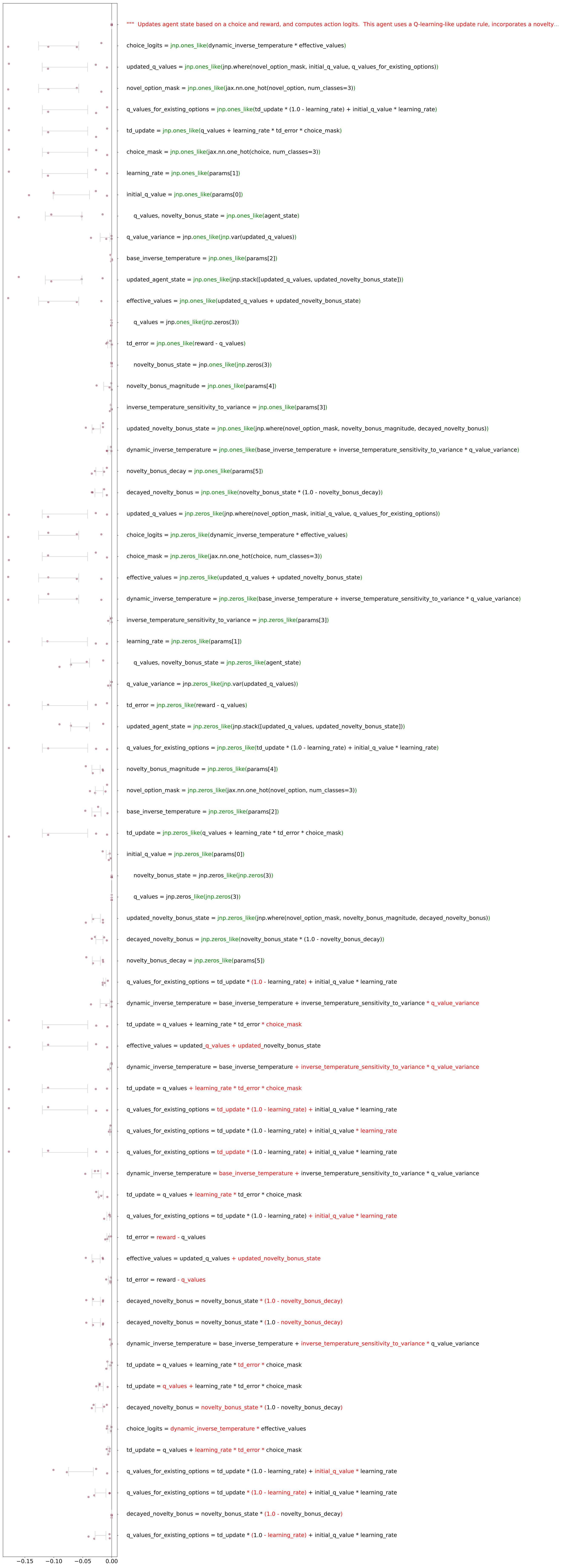

### ablation_performance_monkey_bandit_run3_low_floor_refactored_20260420.pdf

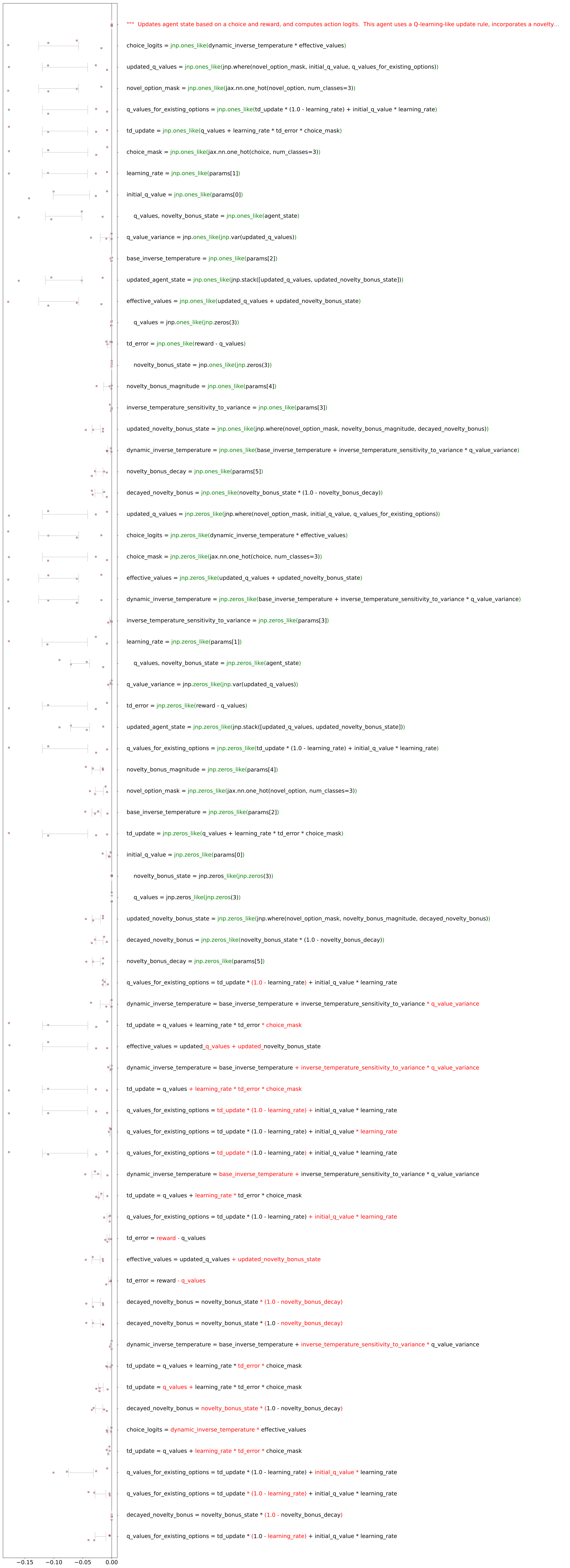

### ablation_performance_monkey_bandit_run3_medium_floor_refactored_20260420.pdf

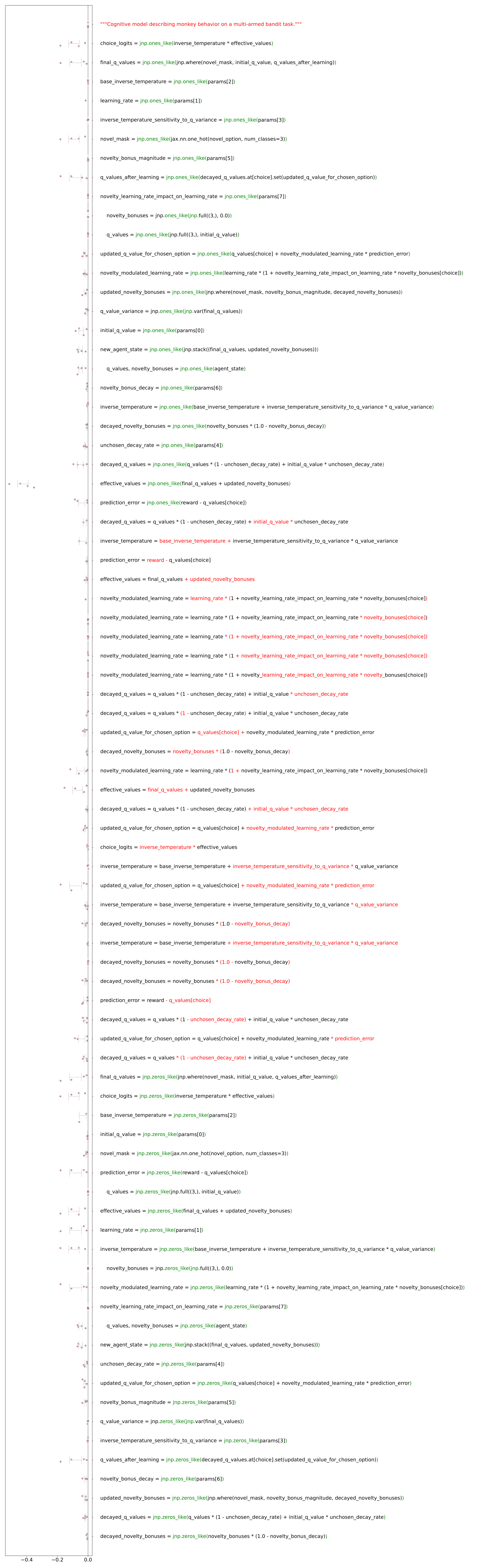

### ablation_performance_rat_bandit_run1_low_floor_20260420.pdf

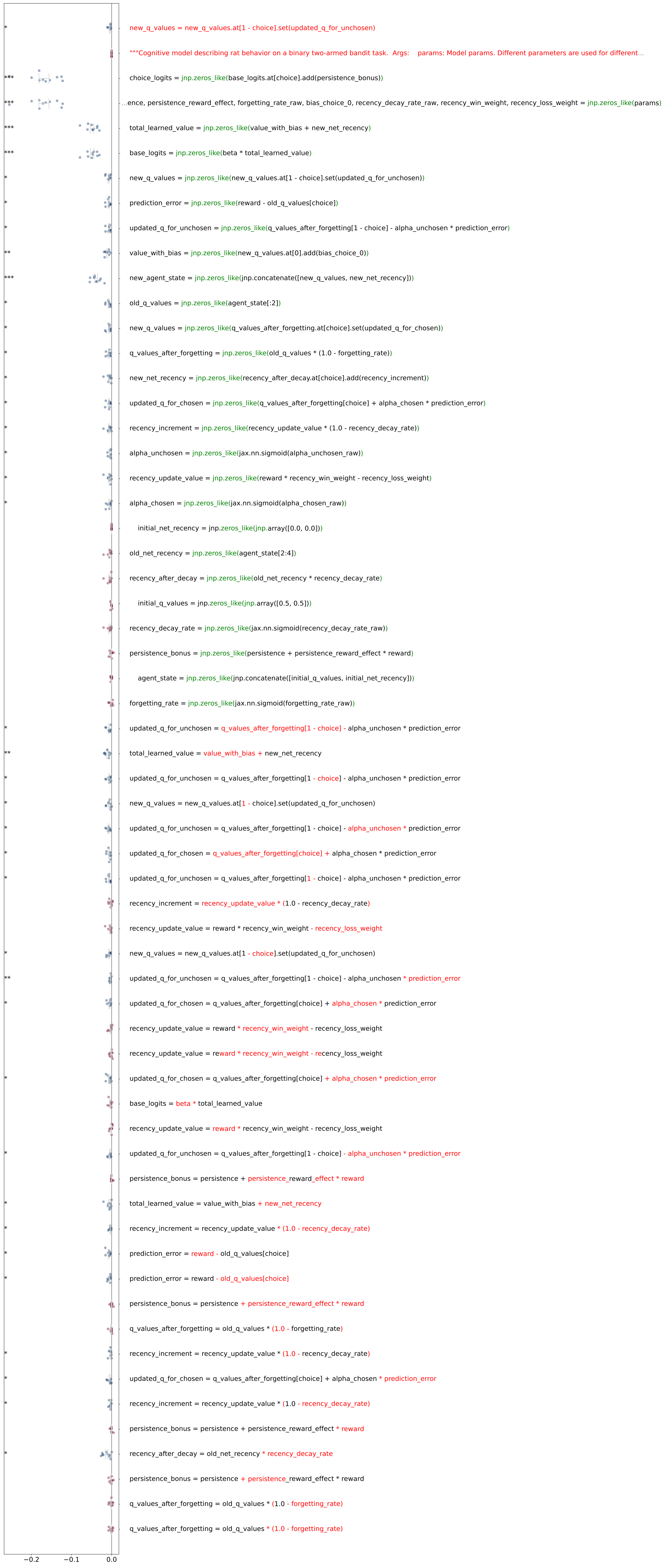

### ablation_performance_rat_bandit_run1_medium_floor_20260420.pdf

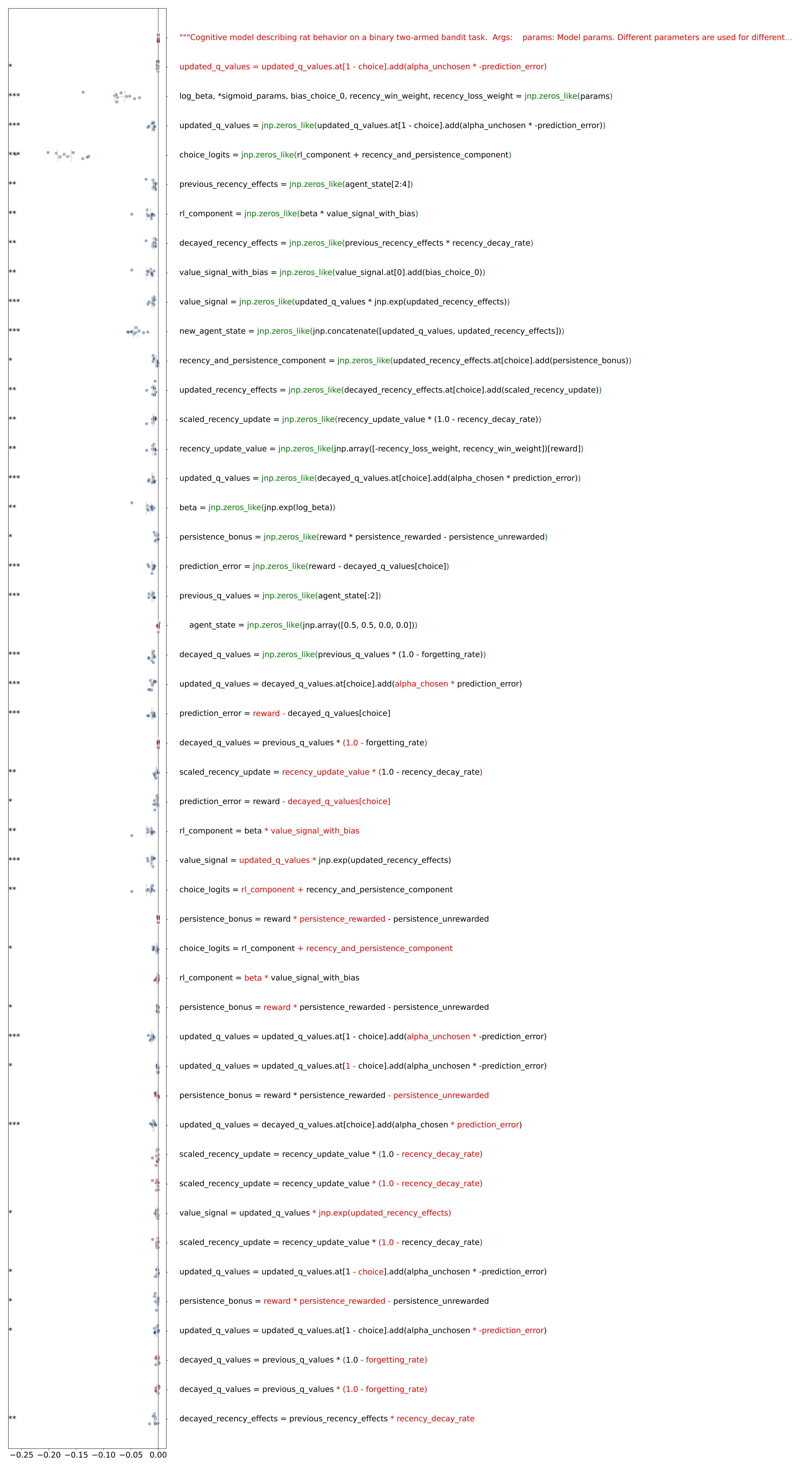

### ablation_performance_rat_bandit_run2_high_floor_20260420.pdf

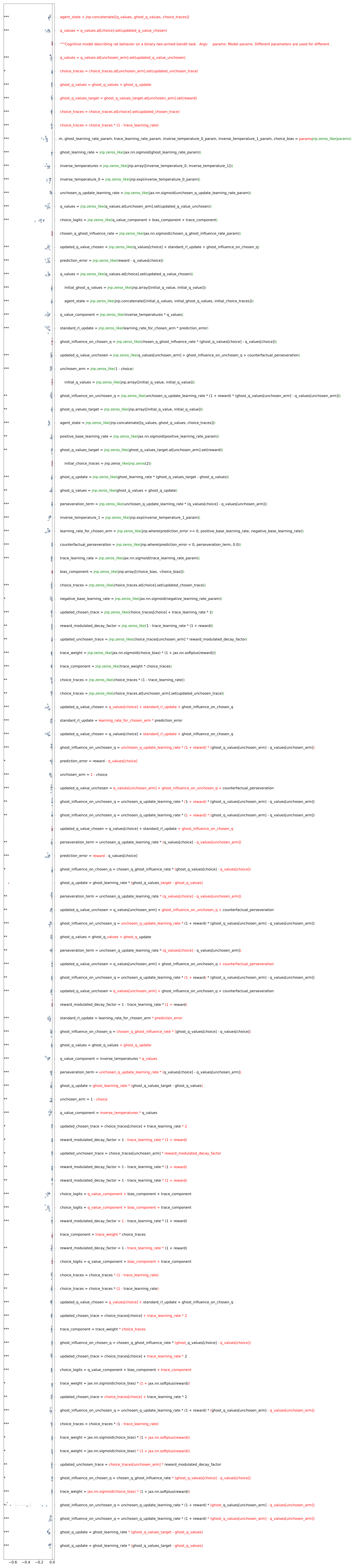

### ablation_performance_rat_bandit_run2_low_floor_20260420.pdf

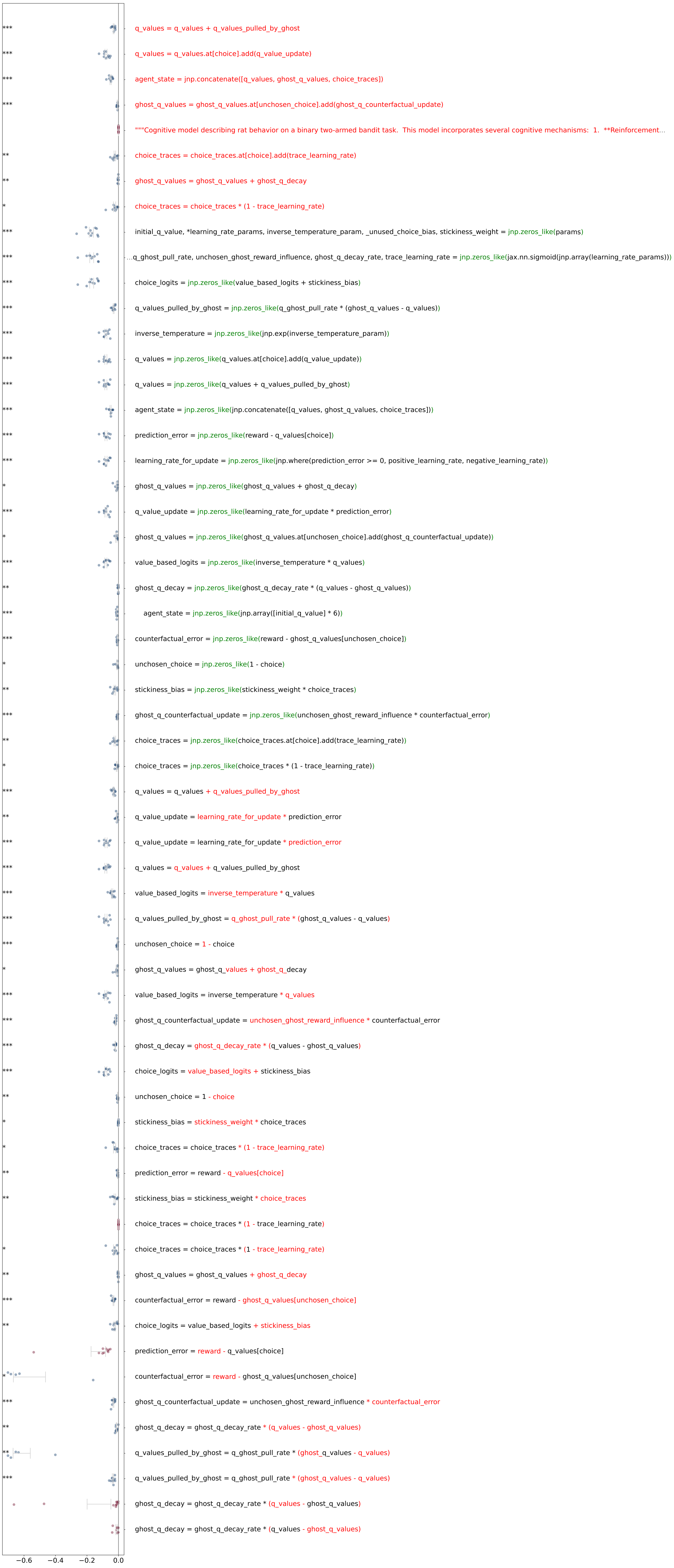

### ablation_performance_rat_bandit_run2_medium_floor_20260420.pdf

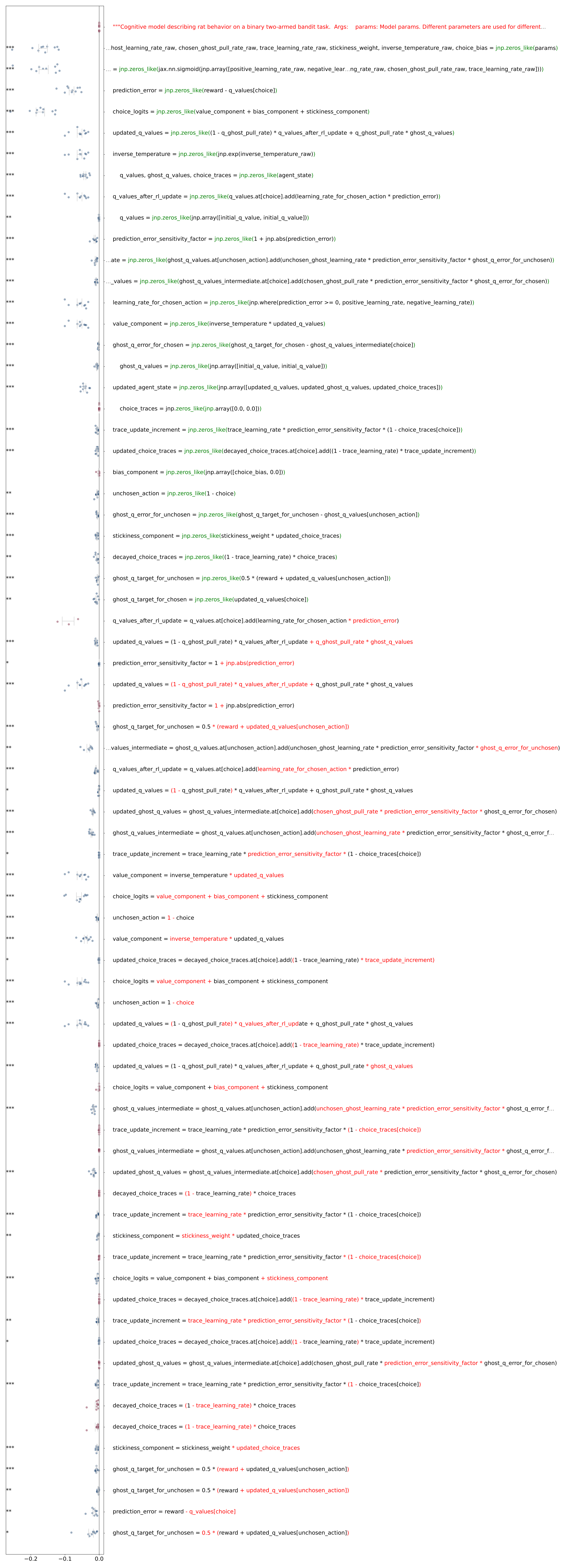

### ablation_performance_rat_bandit_run3_low_floor_20260420.pdf

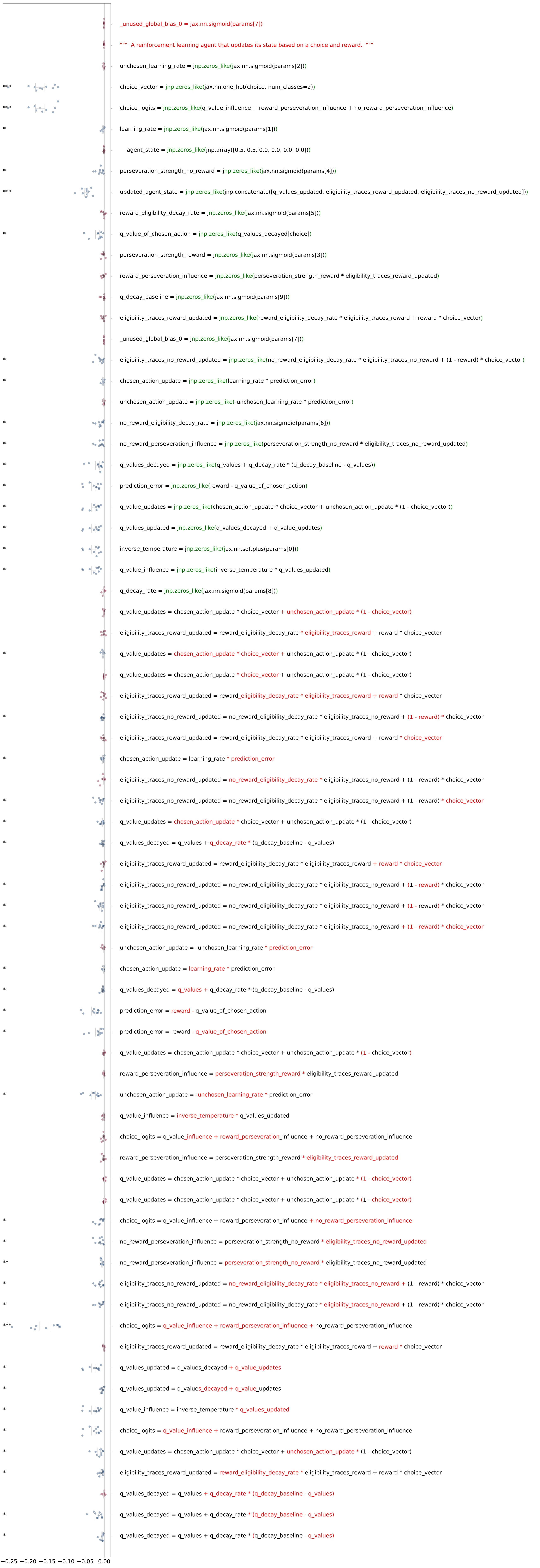

### ablation_performance_rat_bandit_run3_medium_floor_20260420.pdf

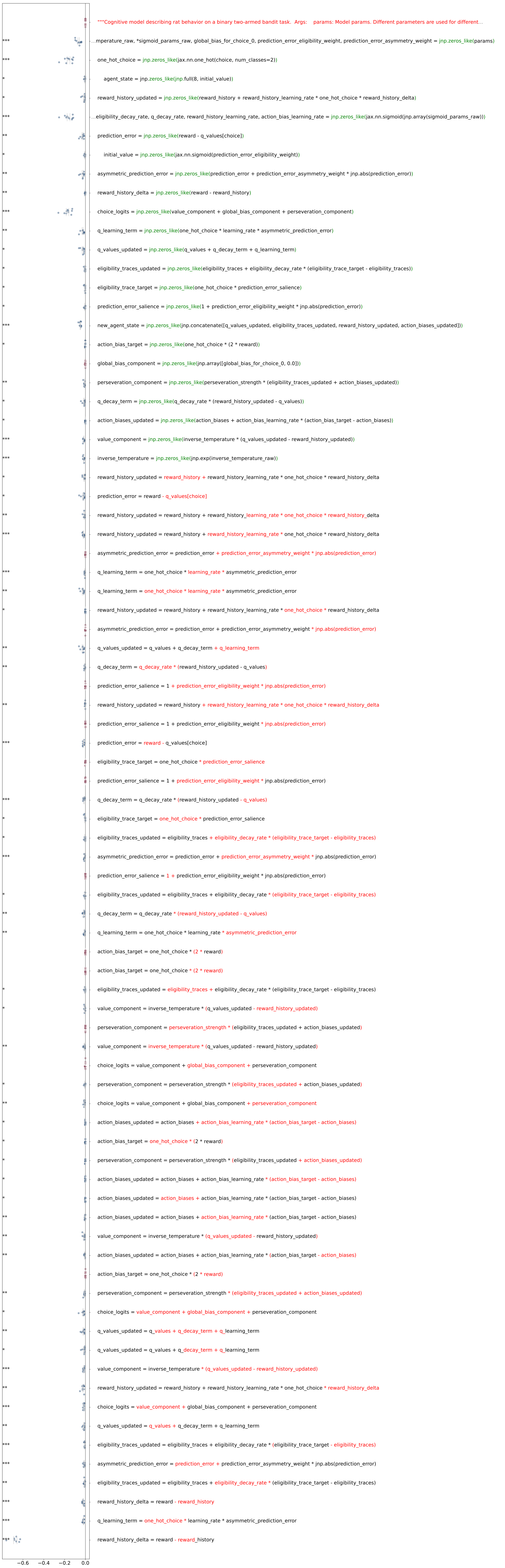

### ablation_performance_rat_twostep_run1_high_floor_20260420.pdf

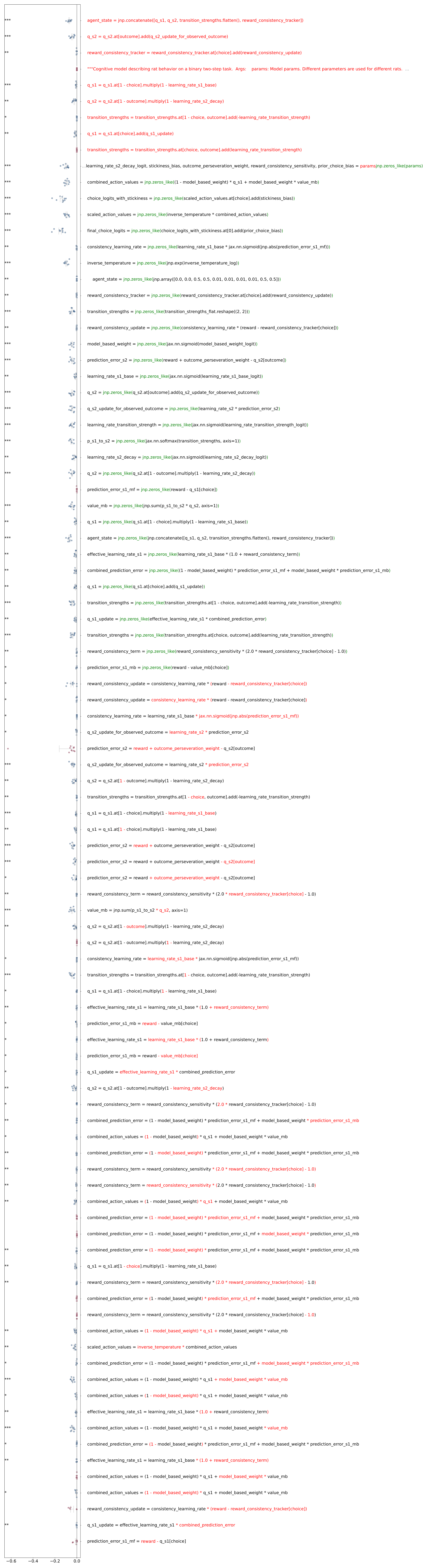

### ablation_performance_rat_twostep_run1_low_floor_20260420.pdf

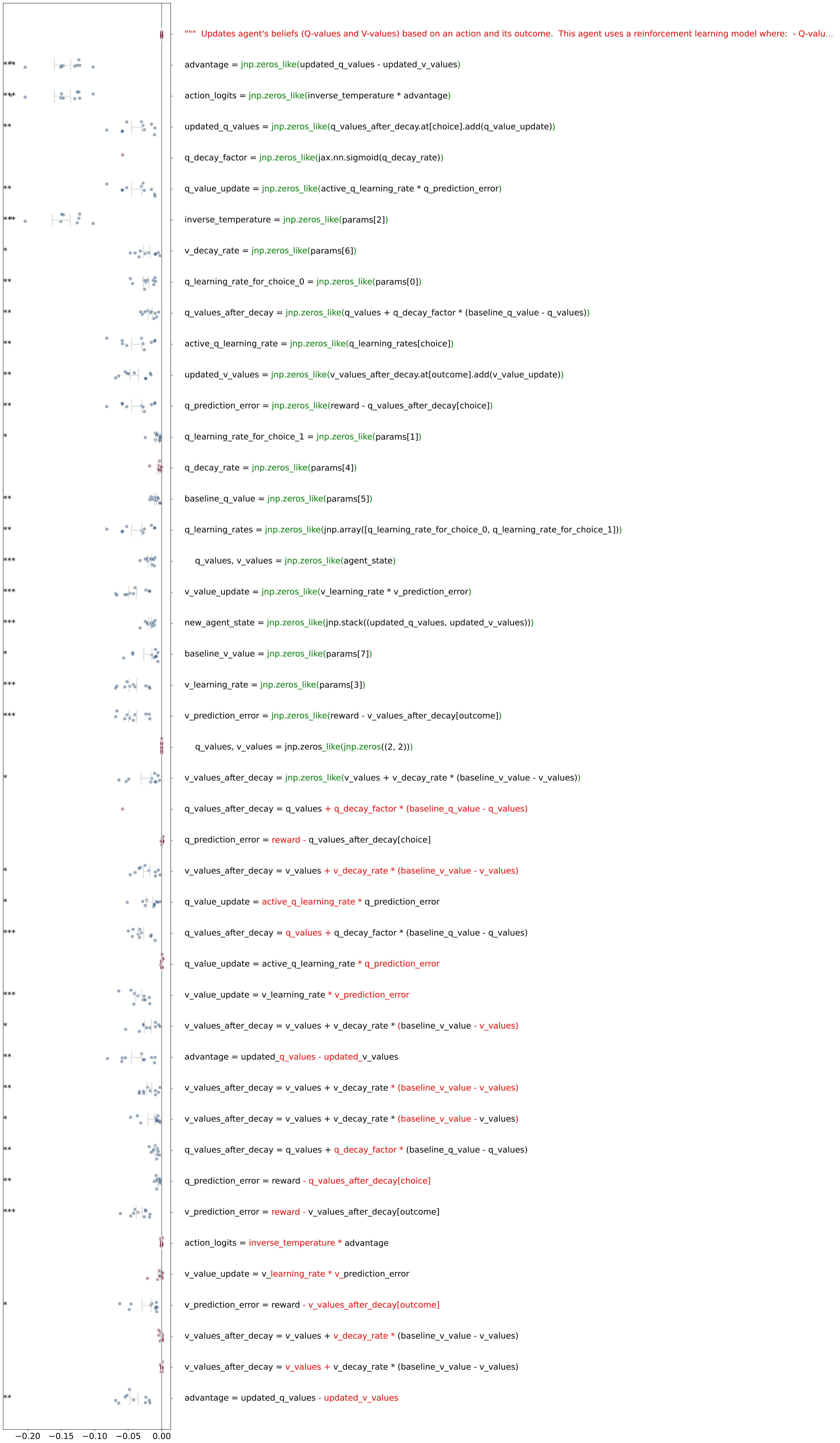

### ablation_performance_rat_twostep_run1_medium_floor_20260420.pdf

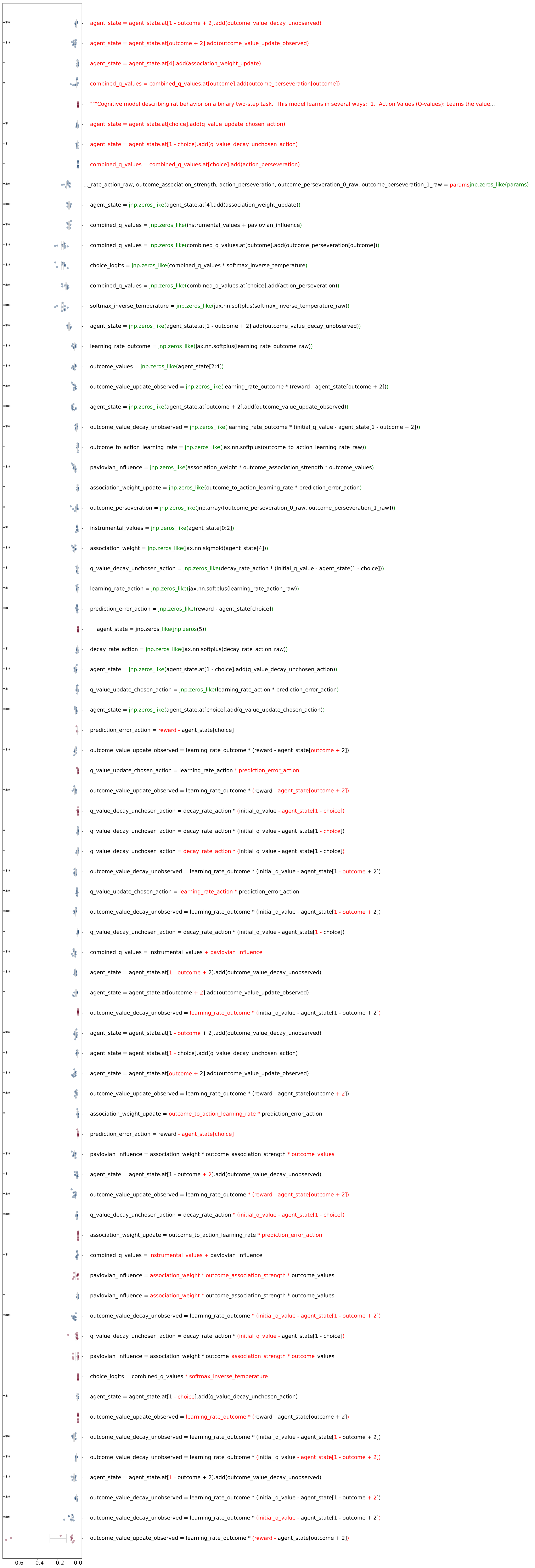

### ablation_performance_rat_twostep_run2_high_floor_20260420.pdf

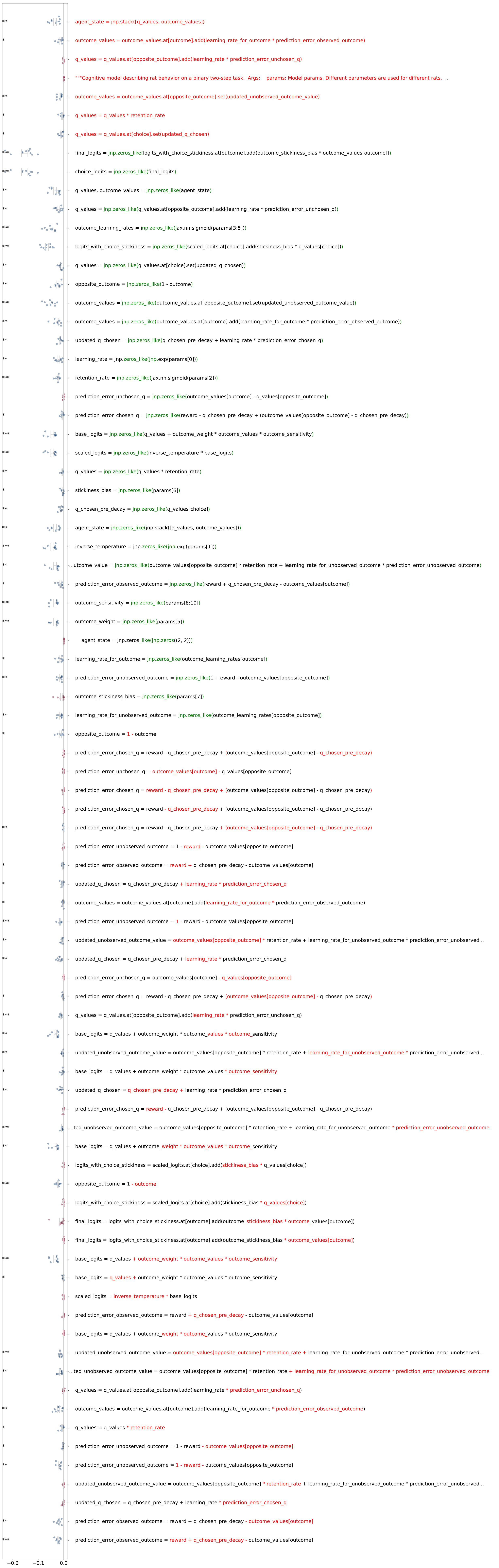

### ablation_performance_rat_twostep_run2_low_floor_shortenonly_20260420.pdf

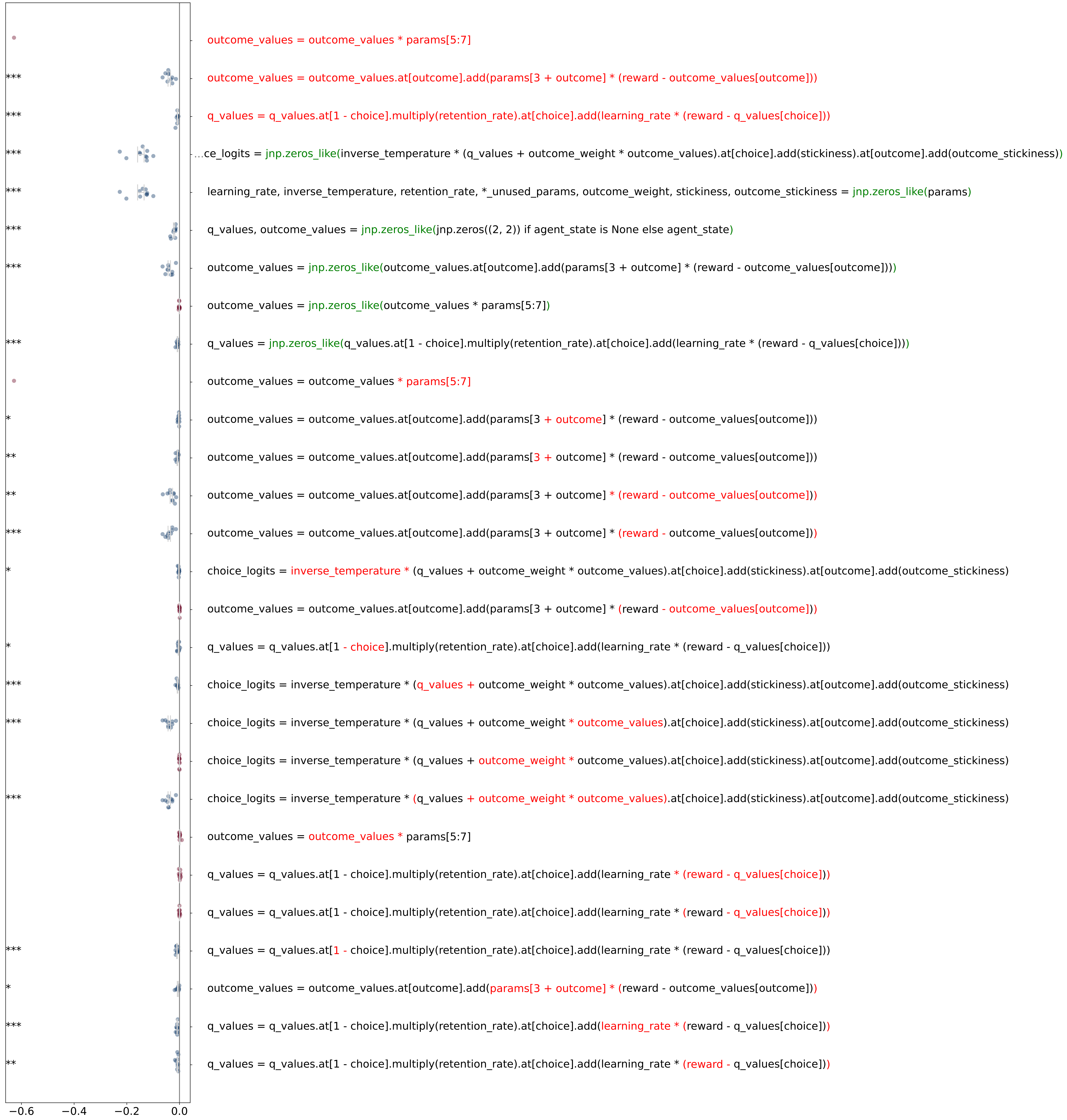

### ablation_performance_rat_twostep_run2_medium_floor_20260420.pdf

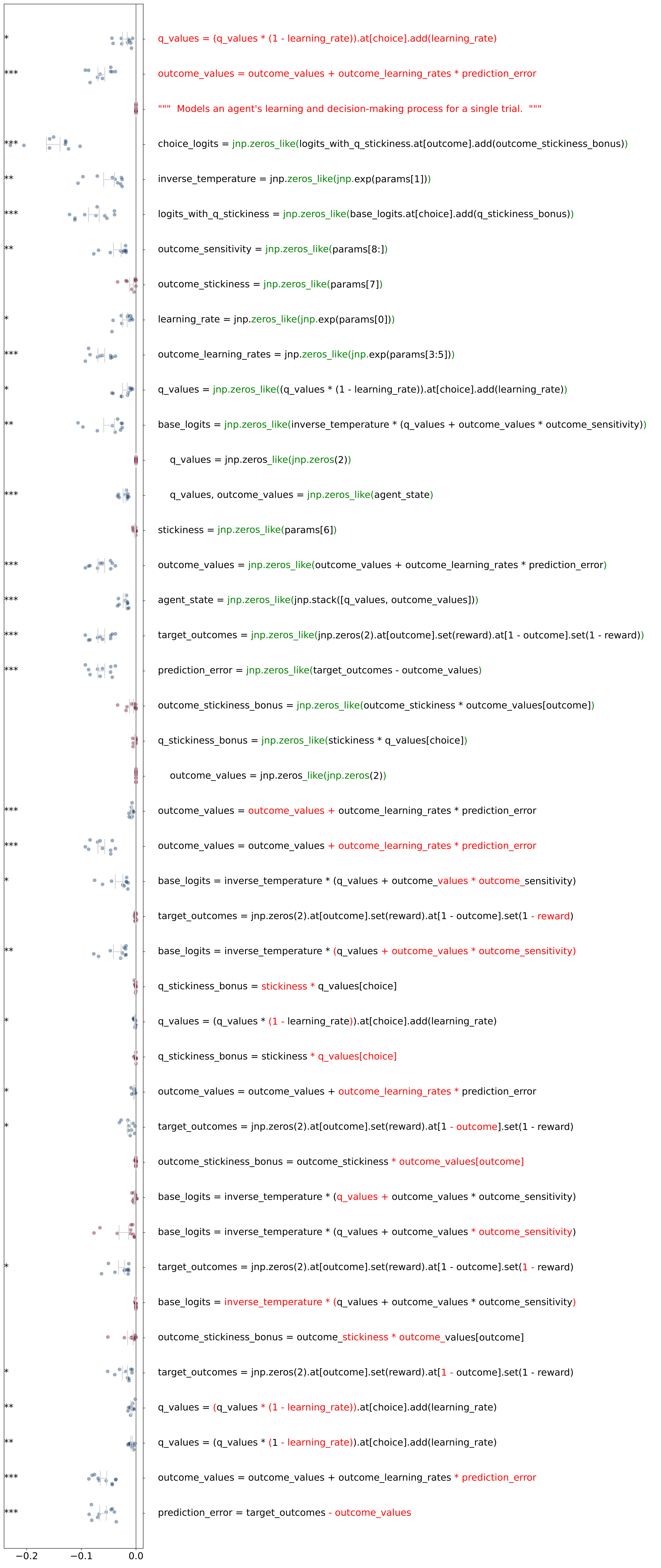

### ablation_performance_rat_twostep_run3_high_floor_20260420.pdf

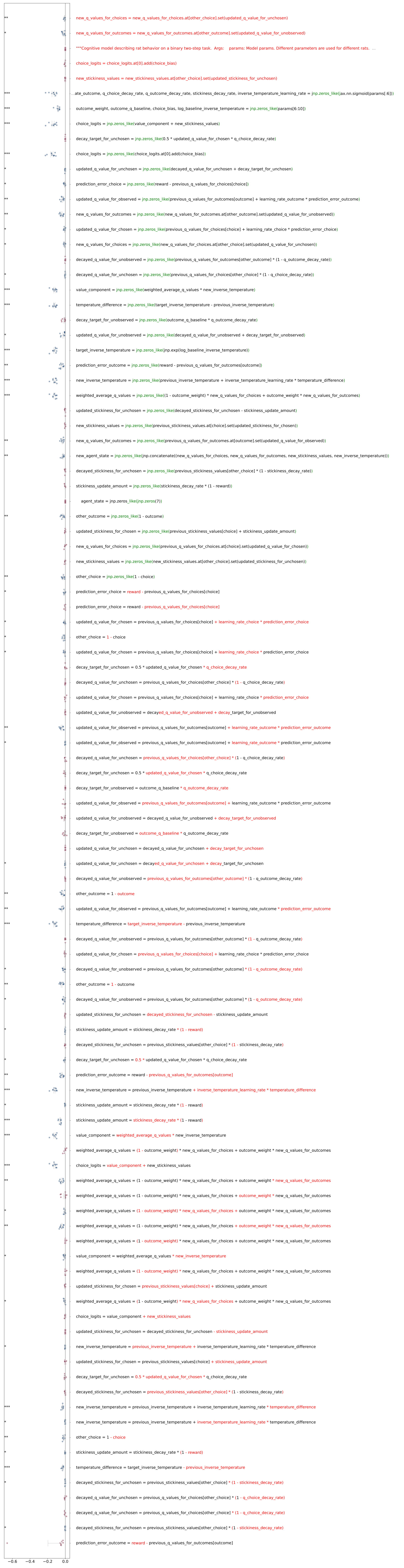

### ablation_performance_rat_twostep_run3_low_floor_20260420.pdf

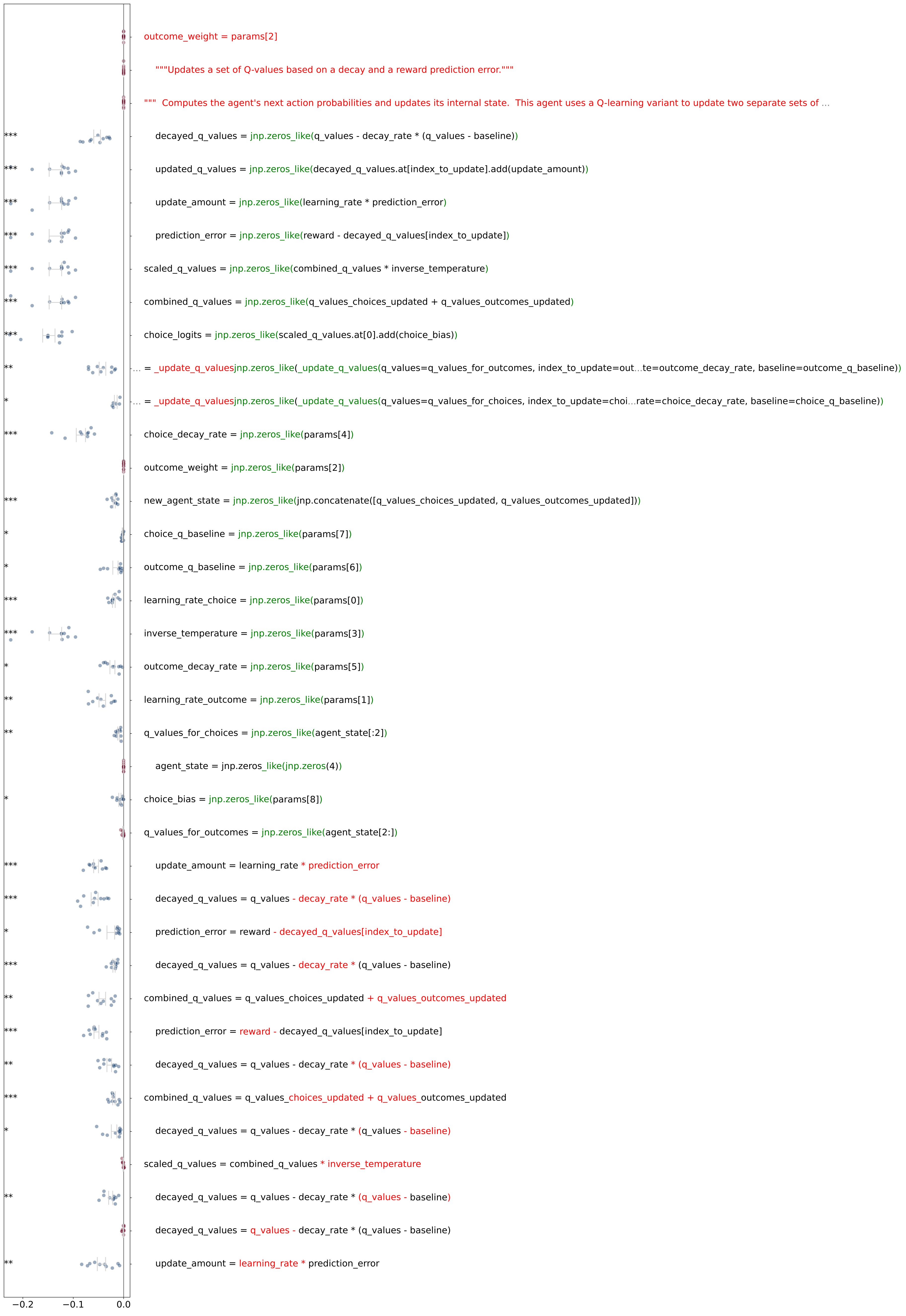

### ablation_performance_rat_twostep_run3_medium_floor_20260420.pdf

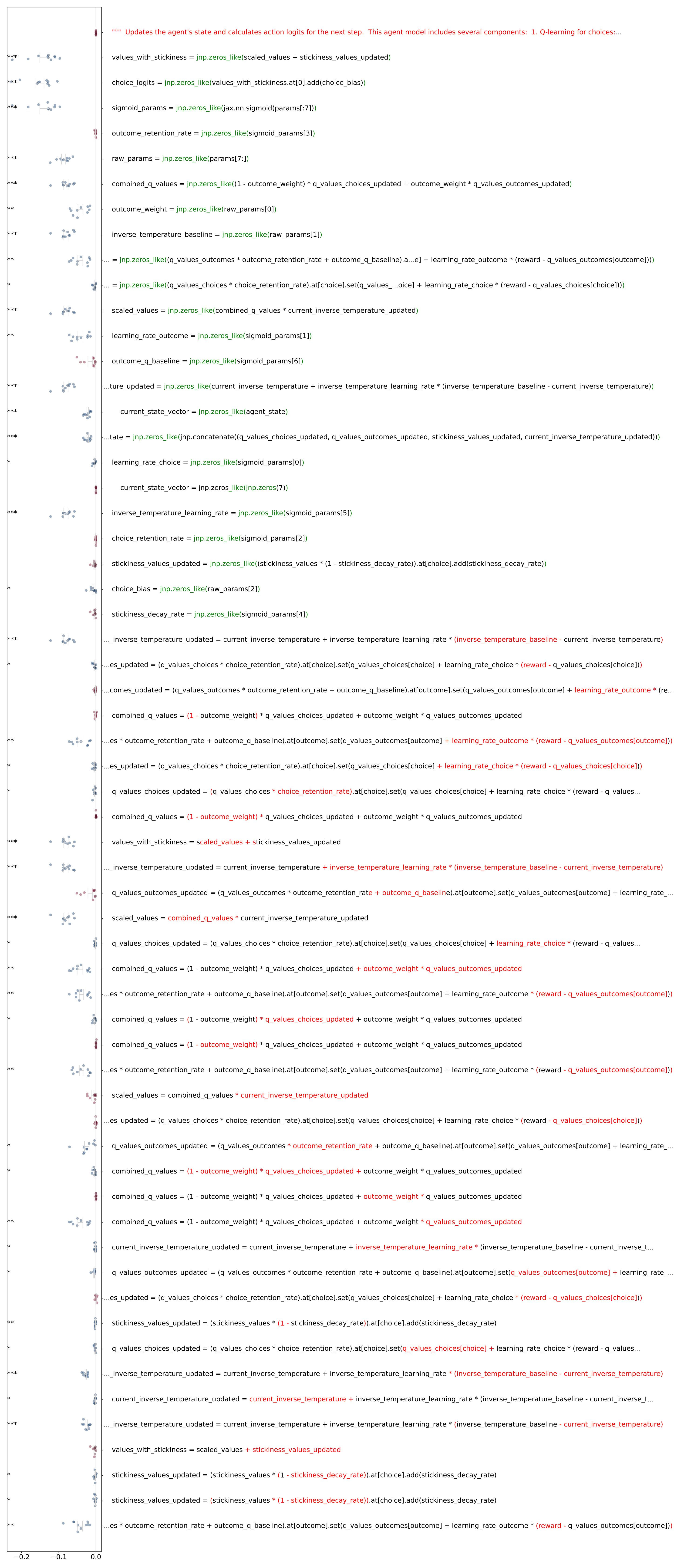
